# Transcriptional evaluation of the ductus arteriosus at the single-cell level uncovers a requirement for vimentin for complete closure

**DOI:** 10.1101/2021.10.30.466605

**Authors:** Jocelynda Salvador, Gloria E. Hernandez, Feiyang Ma, Cyrus W. Abrahamson, Matteo Pellegrini, Robert Goldman, Karen M. Ridge, M. Luisa Iruela-Arispe

## Abstract

**OBJECTIVE:** Failure to close the ductus arteriosus immediately post-birth, patent ductus arteriosus (PDA), accounts for up to 10% of all congenital heart defects. Despite significant advances in PDA management options, including pharmacological treatment targeting the prostaglandin pathway, a proportion of patients fail to respond and must undergo surgical intervention. Thus, further refinement of the cellular and molecular mechanisms that govern vascular remodeling of this vessel is required.

**APPROACH AND RESULTS:** As anticipated, single-cell RNA sequencing on the ductus arteriosus in mouse embryos at E18.5, P0.5, and P5, revealed broad transcriptional alterations in the endothelial, smooth muscle, and fibroblast cell compartments. Making use of these data sets, vimentin emerged as an interesting candidate for further investigation. Subsequent studies demonstrated that, in fact, mice with genetic deletion of vimentin fail to complete vascular remodeling of the ductus arteriosus, as per presence of a functional lumen.

**CONCLUSIONS:** Through single-cell RNA-sequencing and by tracking closure of the ductus arteriosus postnatally in mice, we uncovered the unexpected contribution of vimentin in driving complete closure of the ductus arteriosus potentially through regulation of the Notch signaling pathway.

**HIGHLIGHTS:**

- Single-cell RNA-sequencing on the ductus arteriosus at E18.5, P0.5, and P5 reveals how the ductus arteriosus undergoes drastic transcriptional changes at the single-cell level.
- Endothelial cells increase levels of Vimentin, Notch1 and Jag1 transcripts soon after birth (P0.5), concurrent with ductus arteriosus closure.
- Loss of vimentin, the major intermediate filament protein of endothelial cells, prevents proper permanent closure of the ductus arteriosus.

## INTRODUCTION

During fetal life, the ductus arteriosus, or ductus Botalli, is the vessel that connects the left pulmonary artery to the descending aorta. This structure is critical during fetal development as it diverts the blood emanating from the right ventricle away from the high-resistance pulmonary circulation and into the systemic circulation. At birth a combination of physical and biochemical changes including: the inflation of the lungs, the resulting changes in hemodynamics and alterations in prostaglandin levels trigger the closure of the ductus arteriosus. This essential physiological event prevents the mixture of non-oxygenated blood (from the pulmonary artery) with oxygenated blood (from the aorta).

Abnormal persistence of this vessel, a condition termed patent ductus arteriosus (PDA), frequently occurs in pre-term infants and it is associated with subsequent mortality (1–3). Clinical management of PDA ranges from surgical ligation to pharmacotherapy with cyclooxygenase inhibitors (1–3). Unfortunately, a percentage of patients with PDA fail to respond to these inhibitors. Consequently, a more refined understanding of the processes involved in closure of the ductus arteriosus could offer novel targets for intervention and improved therapies.

At the cellular level, the postnatal closure of the ductus arteriosus is thought to involve the contribution of smooth muscle cells, endothelial cells, and fibroblasts leading to the eventual transformation of this embryonic vessel into a ligament (1–3). Initially, a rapid and robust constriction of smooth muscle cells restricts blood flow. Subsequently, changes in endothelial and smooth muscle cells promote complete closure of the lumen, a process that in humans takes one to three weeks. While much it is known about the initial stage, less is understood about the molecular drivers associated with the later steps. Physiologically, it is understood that the initial constriction of the ductus arteriosus relies on increased arterial pO2, decreased blood pressure in the DA lumen, reduction in circulating prostaglandin E2 (PGE2) and PGE2 receptor levels. These events trigger rapid smooth muscle cell contraction through changes in intracellular calcium levels and a drastic reduction in PGE2 signaling upon dissociation of the placenta which is the main source of PGE2. Nonetheless, complete closure of the lumen requires the participation of the endothelium through relatively unknown mechanisms.

Here we sought to characterize the transcriptional changes involved in the closure of the ductus arteriosus by performing single cell RNAseq prior to, during and shortly after closure using the mouse as a model system. Our second goal was to identify novel regulators of developmentally programmed vascular remodeling.

## MATERIALS AND METHODS

### Mice

For scRNA-seq experiments, C57BL/6 mice at the following stages were used: E18.5, P0.5 and P5. Libraries were generated from ducti of pooled littermates (8-9 mice per library, both sexes). The 129/Sv6 vimentin-deficient (*Vim−/−*) mice were a gift from Albee Messing (Madison, WI)

### Single-cell isolation

For cell isolation, aorta and ducti were dissected in versine; vessels were minced into small pieces and placed in 1mL of digestion buffer containing DNase1, 1M HEPES, Libase, and HBSS at 37C for 20mins. Once tissue was digested into a single cell suspension, cells were washed, pelleted, and passed through a 40μm filter. The final cell suspension was in 0.04%BSA.

scRNA-seq libraries were generated using 10X Genomics Chromium Single Cell 3’ Library & Gel Bead Kit v3. Cells were loaded accordingly following the 10X Genomics protocol with an estimated targeted cell recovery of 6000 cells. Sequencing was performed on NovaSeq6000 (Pair-end, 100 base pairs per read). The digital expression matrix was generated by demultiplexing, barcode processing, and gene unique molecular index counting using the Cellranger count pipeline (version 4.0.0, 10X Genomics). Multiple samples were merged using the Cellranger aggr pipeline. To identify different cell types and find signature genes for each cell type, the R package Seurat (version 3.1.2) was used to analyze the digital expression matrix. Specifically, cells that express <100 genes or <500 transcripts were filtered out. The data were normalized using the NormalizeData function with a scale factor of 10,000. The genes were then scaled and centered using the ScaleData function. Principal component analysis (PCA), Uniform Manifold Approximation and Projection (UMAP) were used to reduce the dimensionality of the data. Cell clusters were identified using the FindClusters function. The cluster marker genes were found using the FindAllMarkers function. Cell types were annotated based on the cluster marker genes. Heatmaps, violin plots and gene expression plots were generated by DoHeatmap, VlnPlot, FeaturePlot functions, respectively. Data will be deposited on GEO and an accession number will be included upon acceptance of the manuscript.

### Immunofluorescence

For whole-mount aorta staining, mice were euthanized and perfused with 2% paraformaldehyde (PFA). The aortae were dissected from the backbone, cleaned, filleted open, and pinned down on silicone-coated plates with the endothelium facing up and left in 2% PFA overnight. The following day, aortae were washed three times in 1x PBS, blocked for 1hr, and primary antibodies were added in fresh blocking buffer overnight. On the second day the aortae were washed three times in 1x PBS and secondaries were added in fresh blocking buffer and allowed to incubate for 2 hrs followed by three washes. After the washes they were mounted onto slides using ProLong Gold Antifade Mounting Medium (Thermo Fischer Scientific P36930) covered with a coverslip and sealed with nail polish. The following primary antibodies and concentrations were used: Anti-Vimentin (Encor Biotech, # CPCA-Vim, 1:1000), anti-VE-cadherin goat polyclonal (discontinued- Santa Cruz Biotechnology #sc-6458, 1:200), anti-ERG (ABCAM #ab92513, 1:200). PECAM antibody was graciously provided by Dr. Bill Muller (Northwestern). The following secondary antibodies were used: Donkey anti-Rabbit Alexa Fluor™ 568 (Invitrogen A10042) used at 1:400; FITC-Donkey anti-Chicken (Thermo Fisher SA1-72000) used at 1:400; Donkey Anti-Goat Alexa Fluor® 647 (Abcam ab150135) used at 1:400; Alexa Fluor® 488 AffiniPure Goat Anti-Armenian Hamster (Jackson ImmunoResearch Laboratories127-545-160) used at 1:400; and all preparations were stained with DAPI (Thermo Fisher D1306) 1:500. Images were taken on a A1R HD25 confocal microscope (Nikon) using 20x, 40x and 60x objectives.

For histological sections: ducti arteriosus were dissected, fixed in 2% PFA overnight, washed 3 times in PBS and embedded first in Histogel under a dissecting microscope and later in paraffin. Blocks were sectioned at 5μM and stained with H&E by Northwestern Mouse Histology and Phenotyping Laboratory. Other sections were processed with the following primary antibodies: Anti-vimentin (Encor Biotech, # CPCA-Vim), Anti-Smooth Muscle alpha actin– Cy3 (C6198), Anti-smooth muscle Myosin heavy chain 11 (Abcam ab224804), Anti-Calponin 1 (Abcam ab46794), Anti-SM22 alpha (Abcam ab14106). The following secondary antibodies were used: Alexa-Fluor Donkey anti-Rabbit 568, FITC-Donkey anti-Chicken, and all preparations were also stained with DAPI (Thermo Fisher D1306).

Measurements of the ducti at several ages were done on an Echo Revolve microscope equipped with a micrometer for X and Y measurements. The ducti still attached to the dorsal aorta were carefully dissected and pinned onto silicone-coated dishes. Measurements were obtained always at the same distance from the aorta for accurate comparisons using 1.25X and 4X objectives.

### Statistics

To quantify closure of the DA, a ratio of the outer width of the ductus arteriosus (DA) to the width of the descending aorta (AO) was calculated for each aorta at each time point. Measurements were done on the Echo Revolve using the annotation tool, and statistics were performed on Graphpad Prism. An unpaired Student’s t-test was used to compare between Wild-type and Vimentin KO measurements at each developmental time point. A p-value of less than

0.05 was considered statistically significant.

## RESULTS

### Changes in Vimentin expression during the closure of the ductus arteriosus

Single-cell RNAseq (scRNA-seq) transcriptomics on the ductus arteriosus (DA) and aorta were conducted at developmental stages prior to constriction, at peak constriction, and at post-vessel occlusion (E18.5, P0.5, and P5; **Figure 1A; Supplementary Figure1A, B**). Each DA were carefully dissected and enzymatically digested to generate 3 independent scRNA-seq libraries, without FACS-sorting (**Figure 1A**). Using dimensionality reduction by uniform manifold approximation and projection (UMAP), we identified 5 distinct cell types and assigned cellular identities based on canonical lineage markers (endothelial cells, vascular smooth muscle cells, fibroblasts, sympathetic nerve cells, and leukocytes) (**Figure 1B-D, Supplementary Figure 1C**).

**Figure 1.**
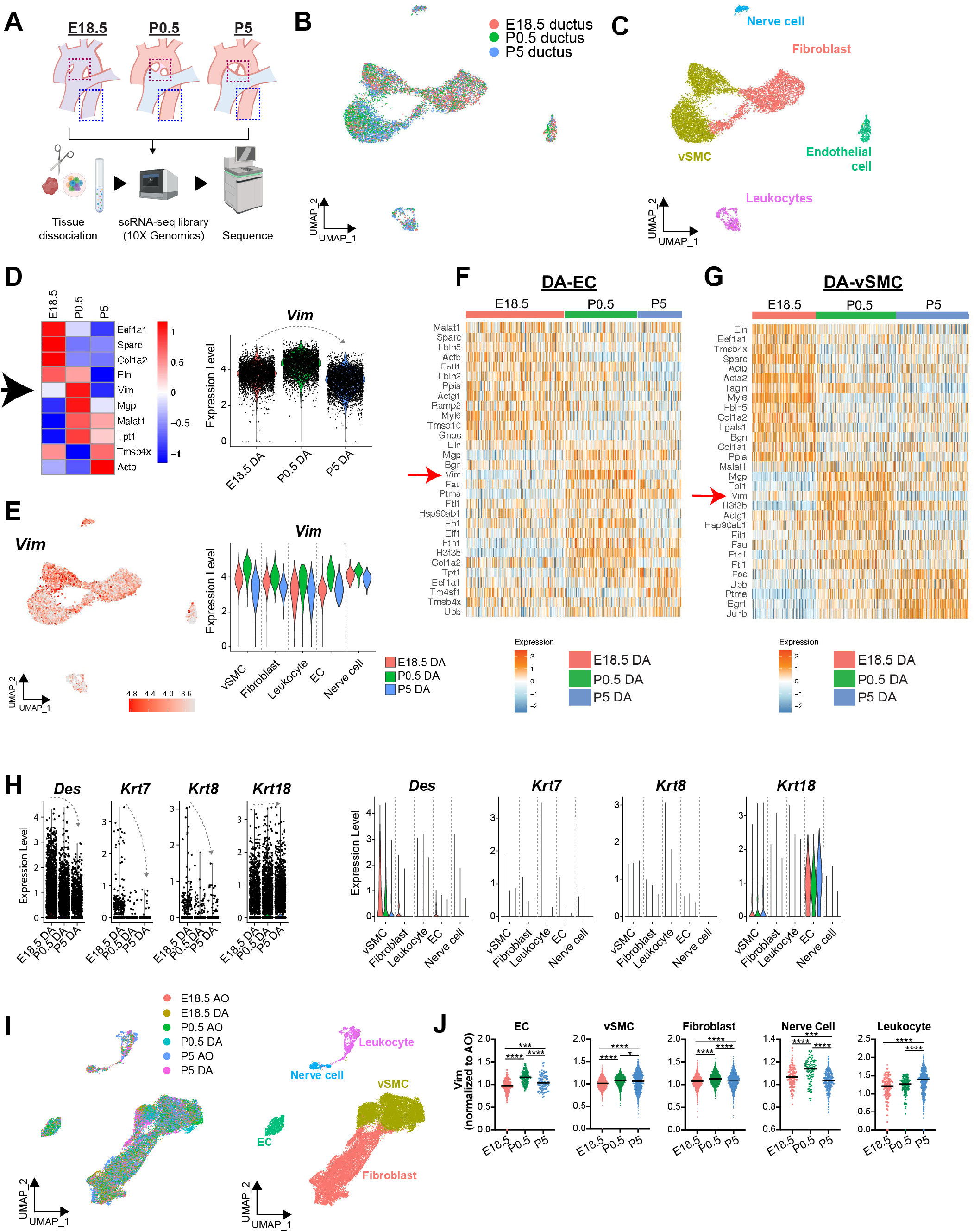
Vimentin expression increases in all vascular cell types during the closure of the ductus arteriosus. **A.** Schematic illustrating the isolation of ductus arteriosus (magenta dotted line) and aortic (blue dotted line) cells for single-cell RNA sequencing. **B**. Uniform manifold approximation and projection (UMAP) plot of cells from three independent libraries: E18.5, P0.5, and P5 ductus arteriosus. 8-9 ductus arteriosus (ductus, DA) were used per library. **C**. UMAP analysis and heat map-style representation of canonical lineage markers (color key represents expression level). **D**. Left: heat map of the top 10 genes in each ductus arteriosus library at the respective time points (E18.5, P0.5 and P5). Arrow identifies vimentin (*Vim*) as the transcript that shows the highest expression at P0.5 with drastic dropped at P5. Right: graph illustrating bell-shape expression pattern of vimentin. **E**. Left: UMAP representation of Vimentin (*Vim*) expression. Right: Vim expression levels in each cell type at the indicated time-points. **F**. Heat map of the top 30 genes in the ductus arteriosus – endothelial compartment (DA-EC). Note clear and sharp upregulation of vimentin at P0.5 (red arrow). **G**. Heat map of the top 30 genes in the ductus arteriosus – vascular smooth muscle cell compartment (DA-vSMC). Note clear and sharp upregulation of vimentin at P0.5 (red arrow). **H**. Violin plots showing overall expression of selected intermediate filaments in the ductus arteriosus at each given time point shown as normalized gene expression per cell. Dotted grey arrows note expression trends through time. Also shown on the right, selected intermediate filament expression in the ductus arteriosus at each time point per cell type. **I**. UMAP plot of cells from 6 libraries: the DA and aorta (AO) at E18.5, P0.5, and P5, isolated from the same mice and cellular identities assigned to each cluster (UMAP on the right). **J**. Vim expression in each cell type at the given time points, normalized to aortic cell *vim* expression.

Evaluation of the top 10 genes expressed at all time points, revealed two long non-coding RNAs (*Malat1* and *Tpt1*); four extracellular matrix transcripts (*Sparc, Col1a2, Eln, and Mgp*) and three cytoskeletal proteins (*Vim, Tmsb4x and Actb*) (**Figure 1D**). The 10^th^ gene, not surprisingly, was eukaryotic translation elongation factor 1 alpha (*Eef1a1*) involved in modulation of cytoskeleton, and that also exhibits chaperone-like activity, controls proliferation and cell death. Interestingly, these three transcripts exhibit a time-specific pattern of upregulation being prior to, during, or after constriction. Our particular interest when analyzing the data was to identify transcripts that sharply increased at P0.5, the time of DA closure and that returned to steady-state or lower by P5, a profile clearly followed by the intermediate filament Vimentin (*Vim*). Importantly, vimentin was highly expressed by vascular smooth muscle cells (vSMCs), fibroblasts, and endothelial cells (EC) (**Figure 1E**). Furthermore, its expression pattern fulfilled our criteria by exhibiting a bell-shaped curve with a peak at P0.5, particularly in vSMC and ECs a time of active DA remodeling. Closer evaluation of the top 30 genes in DA from EC (**Figure 1F**) and vSMC (**Figure 1G**) reveals in closer detail the peak of expression in relation to other highly abundant transcripts. The scRNAseq dataset presented was further mined to answer questions related to specific processes and transcripts during the closure of the ductus arteriosus.

The potential contribution of vimentin in the closure of the ductus arteriosus has not been previously explored. It is pertinent as intermediate filaments are well known to provide mechanical strain and resilience (4). Interestingly, vimentin’s expression was unique amongst other intermediate filaments in the ductus arteriosus. Other members of the family, such as desmin and keratins were also expressed, but at lower levels and with patterns restricted to specific cell types; desmin in smooch muscle and keratins in endothelial cells (**Figure 1H**). Furthermore, neither desmin nor keratins showed an expression pattern that peaked at P0.5 and decrease thereafter.

To clarify whether the increased expression of *Vim* at P0.5 was specific to the DA instead of a general developmental pattern that extended to other vessels, we took advantage of scRNA-seq generated from aorta of the same mice (minus ducti) (**Figure 1I**). Expression levels of *Vim* in the aortae were then used to normalize *Vim* transcripts of the DA to further assess levels per cell type and independent of global developmental patterns. These findings further confirmed sharp increases at P0.5 specifically in ducti endothelial cells, smooth muscle cells, and fibroblasts indicating that across these cell types, *Vim* expression increases during the constriction of the DA at P0.5 (**Figure 1J**).

To directly test the potential contribution of vimentin in the closure of the DA, we took advantage of an established vimentin knock-out mouse model (*Vim−/−*). Genomic deletion of vimentin was initially characterized by Colucci-Guyon in 1994 (5). *Vim−/−* mice are viable and fertile. Evaluation of vimentin in the aortic endothelium confirmed the lack of this type III intermediate filament in the knock-out mice, but a contrasting robust expression was noted in wild-type controls (**Figure 2A**). Transverse sections of adult dorsal aorta also showed high levels of vimentin in wild-type smooth muscle cells of the tunica media and its absence in null mice (**Figure 2B**). Interestingly, *en face* evaluation of the aorta of *Vim−/−* mice revealed the retention of an ostium in the area associated with the connection to the ductus arteriosus (**Figure 2C**).

**Figure 2:**
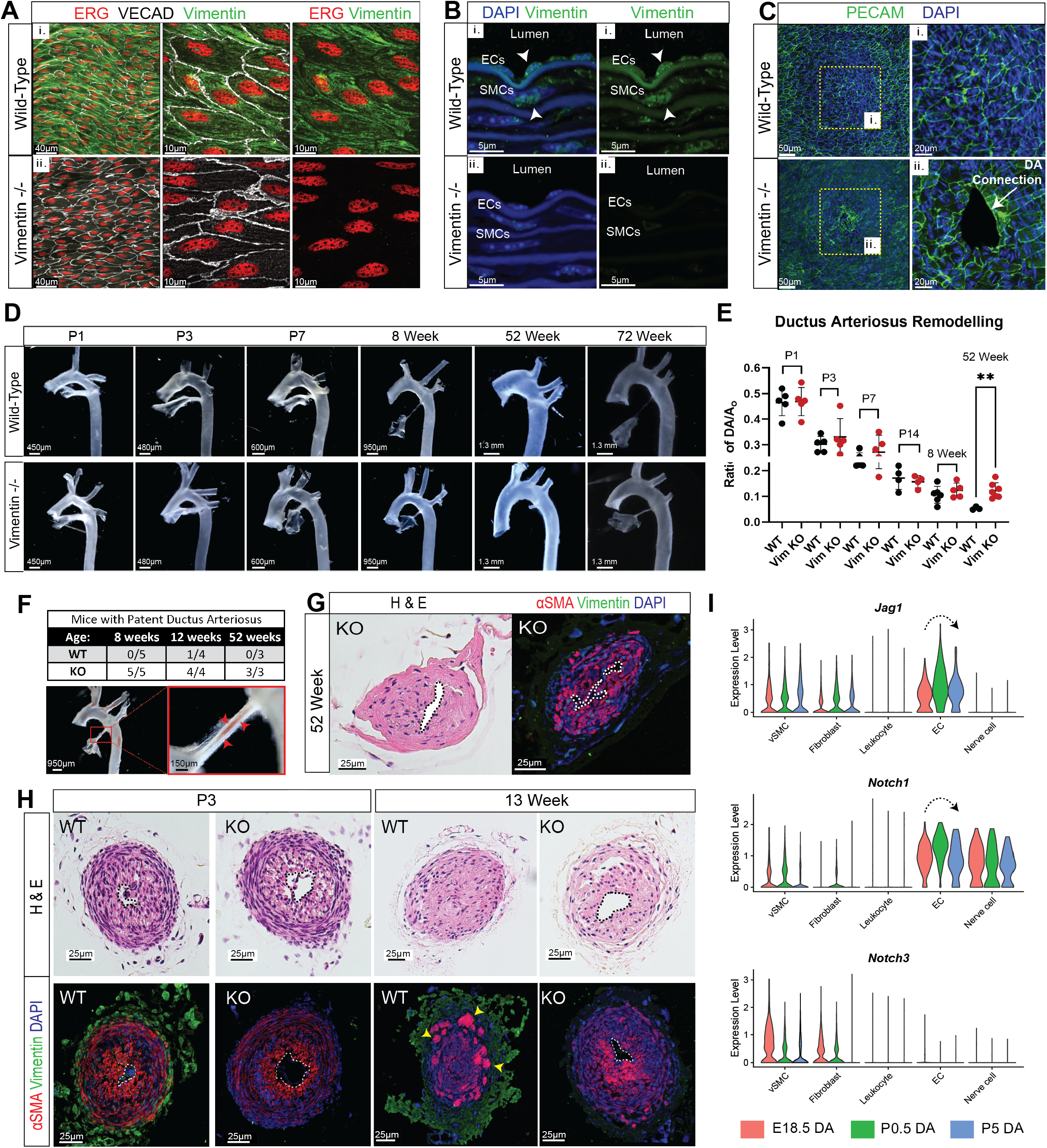
Loss of Vimentin leads to patent ductus arteriosus. **A.** *En face* immunofluorescence preparation of aortae confirm the presence and absence of vimentin in the endothelium of the wild-type (WT) (i) and knock-out (*Vim−/−*) mice (ii), respectively. **B.** Transverse sections of the descending aorta of 8-week old WT and Vim−/− mice demonstrate the abundance of vimentin in endothelial cells (ECs) and smooth muscle cells (SMCs) in the intimal and medial layers of the aorta, respectively. **C.** *En face* staining of the aortic endothelium in the region of the ductus arteriosus opening along the aorta in 52 week old WT and *Vim−/−* mice. Note, *Vim−/−* reveal patency of the ductus arteriosus (ii). **D**. Light imaging of aortae at different developmental timepoints highlights the closure of the ductus arteriosus and the enlargement of the aorta in Wild-type and *Vim−/−* mice from P1 to 52 weeks of age. **E.** Quantification of the width of the ductus arteriosus (DA) normalized to the width of the descending aorta (AO) from bright field images shown in Figure 2D. (n=6 per group, **p<0.001). **F.** Quantification of proportion of adult mice (8weeks-52weeks) with patent ductus arteriosus. n=3-5 mice per time point. **G.** Transversal sections of the DA from 52 week *Vim−/−* mice stained with H&E and stained for alpha smooth muscle alpha actin (red), vimentin (green) and DAPI (blue). Despite lack of observable differences in outer widths of the vessel in **2D & 2E** the lumen is retained and SMCs still retain smooth muscle cell markers. **H.** Transverse sections of the ductus arteriosus in WT and *Vim−/−* mice at P3 and 13 weeks of age stained for vimentin (green) and the smooth muscle marker, alpha-SMA (red). Note clear retention of lumen at 13 weeks in *Vim−/−* in contrast to controls. Also note alterations in smooth muscle cells in WT mice (yellow arrows) not detected in *Vim−/−* mice. **I.** Violin plots showing overall expression of Jag1, Notch1, and Notch3 in the ductus arteriosus at each given timepoint shown as normalized gene expression per cell. Dotted grey arrows note expression trends through time.

The dynamics of DA closure were then tracked in wild-type and *Vim*−/− mice at time points ranging from P1 to 72 weeks of age (**Figure 2D**). Both groups showed striking and similar changes in the width of the DA with marked constrictions of the tunica media. To rigorously quantify the progress of closure, we evaluated 28 control mice and 31 Vim−/− mice from P1 to 1 year of age (Figure 2E). The data showed differences only in adult mice, indicating a lack of complete remodeling in mice deleted for Vim. Macroscopic evaluation rendered the wild-type and *Vim* −/− mice nearly indistinguishable, except for the clear presence of blood cells in the DA of *Vim*−/− mice at both early and late stages. Using the presence of blood in the DA as a read-out for patency, we found that all adult *Vim*−/− mice evaluated showed incomplete closure (**Figure 2F).** This was further confirmed by cross-sections of *Vim*−/− DA from mice at 52w of age (**Figure 2G**) and was consistent with the openings noted in the lateral aspects of the aorta of *Vim*−/− where these ducti connect (**Figure 2C**). Additional histological evaluations of ducti from control and *Vim*−/− mice revealed clear retention of the patency and absence of full cellular remodeling of both the tunica intima and media (**Figure 2H**). Combined these findings support the conclusion that vimentin is required for complete remodeling of the ductus arteriosus and obliteration of the vascular lumen. We were naturally intrigued about potential underlying mechanisms and evaluation of the literature revealed links between vimentin and Notch (6,7).

It is well-accepted that signaling between endothelial cells and vascular smooth muscle cells through the Notch pathway contributes to arterial remodeling. Additionally, it has been demonstrated that the absence of the Notch receptor Jagged1 in endothelial cells, results in patent ductus arteriosus (8). Interestingly, vimentin plays an important role in Notch transactivation by endothelial cells (6). Specifically, depletion of vimentin reduces Jagged1-Notch signaling due to alterations in cellular stiffness which likely impair the pulling force required in Jagged1-Notch interactions that precede ADAM cleavage. Therefore, we explored expression patterns of Notch signaling in the ductus arteriosus. Interestingly, we found that Jag1 and Notch1 expression in endothelial and vascular smooth muscle cells increased at P0.5, coincident with Vimentin upregulation during DA remodeling **(Figure 2I**). Therefore, the concurrent expression of vimentin and Notch1/Jag1 suggests PDA may emerge in *Vim*−/− mice because of ineffective signaling between Jag1 (in endothelial cells) and Notch1/3 (in smooth muscle cells).

## DISCUSSION

Resolution of individual transcriptional changes at the cellular level offers an unprecedented opportunity to seek answers to questions that require complex developmental events. In the case of the remodeling of the ductus arteriosus, scRNAseq revealed a large number of transcriptional alterations in the endothelial, smooth muscle and fibroblast compartments which will likely offer opportunities for exploration by multiple laboratories.

Seeking transcripts that sharply increase in the aorta shortly after birth, we identified vimentin, an intermediate filament protein highly expressed in both endothelial and smooth muscle cells. Interestingly, while viable and fertile (5), we found that mice with genomic deletion of vimentin exhibit incomplete closure of the ductus arteriosus due to failure in the last stages of vascular remodeling. In fact, our data show that *Vim*−/− mice at 1 year of age, still exhibit an open lumen with visible circulating blood. We also found alterations in the remodeling of the tunica media, retention of smooth muscle cell markers in the layers proximal to the endothelium and a lack of complete transition of smooth muscle cells towards a fibroblastic phenotype.

Intermediate filaments cooperate with other elements of the cytoskeleton to provide structural support and mechanical integration between the cell surface, organelles and the nucleus (4,9,10). However, intermediate filaments distinguish themselves from other members of the cytoskeleton by their higher mechanical integrity and resistance to rupture (9,11). This is particularly important for cells that either withstand and/or impose physical forces, such as those occurring during vascular remodeling. While other types of intermediate filaments are expressed by smooth muscle (e.g., desmin) and by endothelial cells (e.g., keratins 8 and 18), the levels of vimentin supersede those in both cell types. Furthermore, in addition to their roles in maintaining mechanical integrity, vimentin undergoes impressive spatial rearrangement in smooth muscle cells during contraction and upon stimulation by several agonists (12,13). Importantly, vimentin has been implicated in the distribution of Ca^2+/^calmodulin-dependent protein kinase II (CamKII), a kinase critical in regulating smooth muscle cell contraction (14). In endothelial cells, vimentin has been implicated in barrier function and cell adhesion (15), but no studies have explored the potential role of vimentin during vascular remodeling.

The process of complete closure of the DA is known to require smooth muscle cell migration and proliferation, disruption of the internal elastic lamina, alterations in extracellular matrix production, endothelial cell proliferation and monocyte adhesion(2,3,8,16–22). Many of these events require Notch signaling, particularly those associated with smooth muscle cell contraction and endothelial cell proliferation (6,8). Recently, vimentin was shown to contribute to the mechanochemical transduction pathway that regulates multilayer cross-talk and structural homeostasis through the Notch signaling pathway (6). While additional mechanistic experiments in the context of DA remodeling are required to definitively establish a causal link, the low stiffness due to lack of vimentin is consistent with deficiencies in force generation needed for Notch signaling.

Our findings indicate that the initial stages of DA remodeling are not compromised by the absence of vimentin. Instead, the later stages that promote complete closure of the lumen are the ones impaired. Little is known about the cellular and molecular processes associated with the closure of a vascular lumen. Our findings highlight the exquisite requirement of vimentin for the completion of this process in the ductus arteriosus.

## NONSTANDARD ABBREVIATIONS AND ACROYNMS

DA: Ductus Arteriosus
PDA: Patent Ductus Arteriosus
AO: Aorta
scRNA-seq: single-cell RNA-sequencing
UMAP: uniform manifold approximation and projection
Vim: Vimentin

## ACKNOWLEDGEMENTS

We would like to thank the Jonsson Comprehensive Center at UCLA for sequencing of scRNAseq libraries and the Mouse Histology and Phenotyping core at Northwestern University. A special thanks to Michelle Steel and Snezana Mirkov for support with animal colonies, assistance with husbandry and mouse experimentation.

## CONTRIBUTIONS

**JC and GEH** designed and performed experiments, wrote and edited the manuscript.
**FM** performed bioinformatic analysis of scRNAseq data.
**CWA** performed experiments.
**KMR** provided the vimentin KO mouse and intellectual input.
**RG** and **MP** provided intellectual discussion.
**MLIA** conceived the study, designed the experiments, wrote and edited the manuscript.
All authors discussed the results and had the opportunity to comment on the manuscript.

## REFERENCES

1. Schneider DJ, Moore JW. Patent ductus arteriosus. Circulation. 2006 Oct 24;114(17):1873–1882.

2. Hsu H-W, Lin T-Y, Liu Y-C, Yeh J-L, Hsu J-H. Molecular Mechanisms Underlying Remodeling of Ductus Arteriosus: Looking beyond the Prostaglandin Pathway. Int J Mol Sci. 2021 Mar 22;22(6).

3. Hung Y-C, Yeh J-L, Hsu J-H. Molecular mechanisms for regulating postnatal ductus arteriosus closure. Int J Mol Sci. 2018 Jun 25;19(7).

4. Patteson AE, Vahabikashi A, Pogoda K, Adam SA, Mandal K, Kittisopikul M, et al. Vimentin protects cells against nuclear rupture and DNA damage during migration. J Cell Biol. 2019 Dec 2;218(12):4079–4092.

5. Colucci-Guyon E, Portier MM, Dunia I, Paulin D, Pournin S, Babinet C. Mice lacking vimentin develop and reproduce without an obvious phenotype. Cell. 1994 Nov 18;79(4):679–694.

6. van Engeland NCA, Suarez Rodriguez F, Rivero-Müller A, Ristori T, Duran CL, Stassen OMJA, et al. Vimentin regulates Notch signaling strength and arterial remodeling in response to hemodynamic stress. Sci Rep. 2019 Aug 27;9(1):12415.

7. Antfolk D, Sjöqvist M, Cheng F, Isoniemi K, Duran CL, Rivero-Muller A, et al. Selective regulation of Notch ligands during angiogenesis is mediated by vimentin. Proc Natl Acad Sci USA. 2017 Jun 6;114(23):E4574–E4581.

8. Feng X, Krebs LT, Gridley T. Patent ductus arteriosus in mice with smooth muscle-specific Jag1 deletion. Development. 2010 Dec;137(24):4191–4199.

9. Chang L, Goldman RD. Intermediate filaments mediate cytoskeletal crosstalk. Nat Rev Mol Cell Biol. 2004 Aug;5(8):601–613.

10. Lowery J, Kuczmarski ER, Herrmann H, Goldman RD. Intermediate filaments play a pivotal role in regulating cell architecture and function. J Biol Chem. 2015 Jul 10;290(28):17145–17153.

11. Köster S, Weitz DA, Goldman RD, Aebi U, Herrmann H. Intermediate filament mechanics in vitro and in the cell: from coiled coils to filaments, fibers and networks. Curr Opin Cell Biol. 2015 Feb;32:82–91.

12. Li Q-F, Spinelli AM, Wang R, Anfinogenova Y, Singer HA, Tang DD. Critical role of vimentin phosphorylation at Ser-56 by p21-activated kinase in vimentin cytoskeleton signaling. J Biol Chem. 2006 Nov 10;281(45):34716–34724.

13. Tang DD, Bai Y, Gunst SJ. Silencing of p21-activated kinase attenuates vimentin phosphorylation on Ser-56 and reorientation of the vimentin network during stimulation of smooth muscle cells by 5-hydroxytryptamine. Biochem J. 2005 Jun 15;388(Pt 3):773–783.

14. Marganski WA, Gangopadhyay SS, Je H-D, Gallant C, Morgan KG. Targeting of a novel Ca+2/calmodulin-dependent protein kinase II is essential for extracellular signal-regulated kinase-mediated signaling in differentiated smooth muscle cells. Circ Res. 2005 Sep 16;97(6):541–549.

15. Dave JM, Bayless KJ. Vimentin as an integral regulator of cell adhesion and endothelial sprouting. Microcirculation. 2014 May;21(4):333–344.

16. Mueller PP, Drynda A, Goltz D, Hoehn R, Hauser H, Peuster M. Common signatures for gene expression in postnatal patients with patent arterial ducts and stented arteries. Cardiol Young. 2009 Aug;19(4):352–359.

17. Hsieh Y-T, Liu NM, Ohmori E, Yokota T, Kajimura I, Akaike T, et al. Transcription profiles of the ductus arteriosus in Brown-Norway rats with irregular elastic fiber formation. Circ J. 2014 Mar 19;78(5):1224–1233.

18. Shelton EL, Ector G, Galindo CL, Hooper CW, Brown N, Wilkerson I, et al. Transcriptional profiling reveals ductus arteriosus-specific genes that regulate vascular tone. Physiol Genomics. 2014 Jul 1;46(13):457–466.

19. Goyal R, Goyal D, Longo LD, Clyman RI. Microarray gene expression analysis in ovine ductus arteriosus during fetal development and birth transition. Pediatr Res. 2016 Jun 3;80(4):610–618.

20. Liu NM, Yokota T, Maekawa S, Lü P, Zheng Y-W, Taniguchi H, et al. Transcription profiles of endothelial cells in the rat ductus arteriosus during a perinatal period. PLoS One. 2013 Sep 27;8(9):e73685.

21. Saito J, Kojima T, Tanifuji S, Kato Y, Oka S, Ichikawa Y, et al. Transcriptome Analysis Reveals Differential Gene Expression between the Closing Ductus Arteriosus and the Patent Ductus Arteriosus in Humans. J Cardiovasc Dev Dis. 2021 Apr 16;8(4).

22. Bökenkamp R, Raz V, Venema A, DeRuiter MC, van Munsteren C, Olive M, et al. Differential temporal and spatial progerin expression during closure of the ductus arteriosus in neonates. PLoS One. 2011 Sep 6;6(9):e23975.

