## Supplementary Figure 1 and Table 1 for "Transcriptional evaluation of the ductus arteriosus at the single-cell level uncovers a requirement for vimentin for complete closure"

Salvador, Hernandez et al.

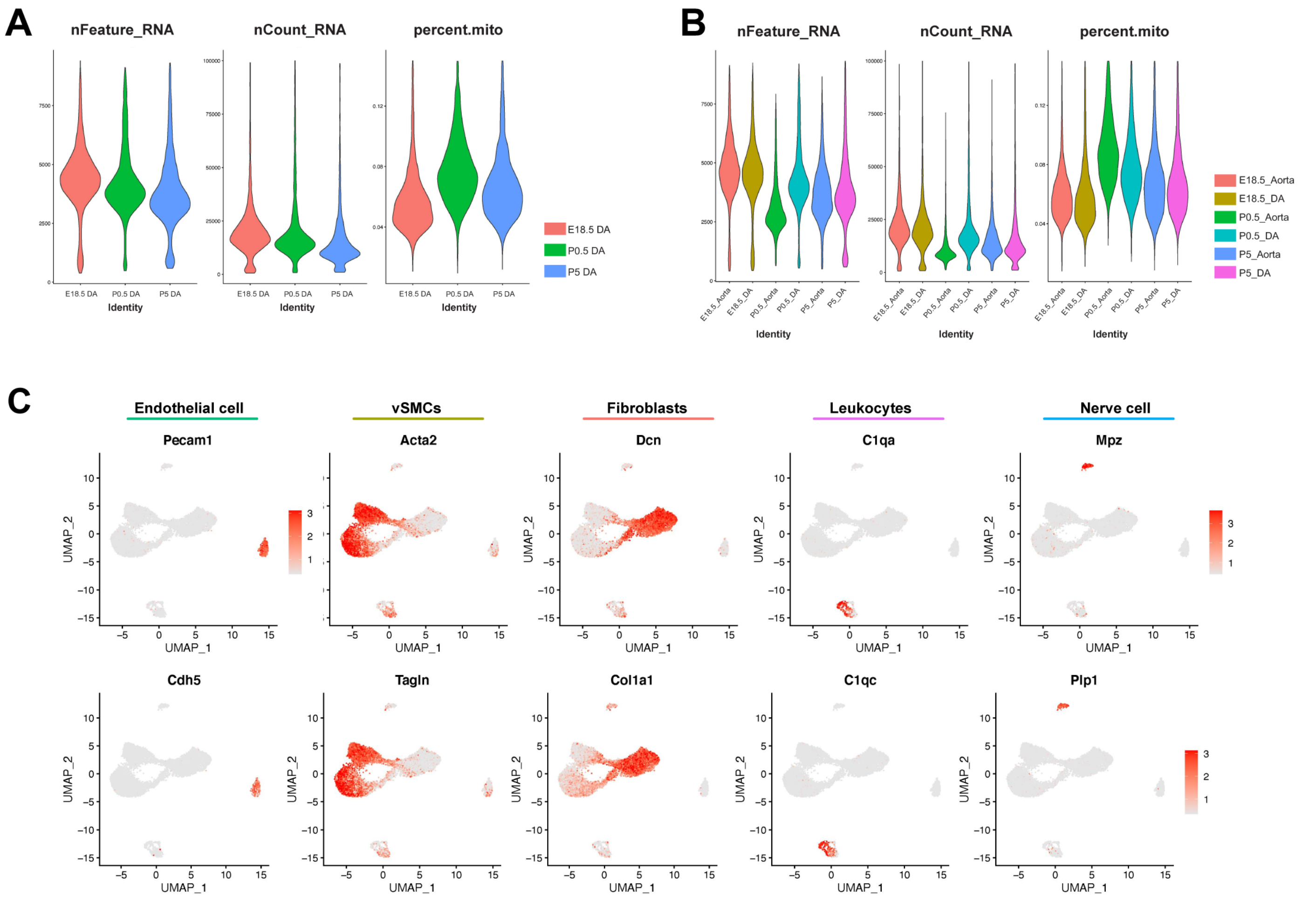


**Supplementary Figure 1**:

- **A.** Evaluation of data quality of the three libraries generated (E18.5, P0.5, and P5). The average number of genes detected per cell (nFeature), the total number of molecules detected per cell (nCount), and the perfect of mitochondrial genes per cell (percent.mito) for each library. Cells having more than 3,000 genes, at least 1,000 unique molecular identifiers, and at most 10% mitochondrial gene expression were selected.
- **B.** Comparison between Aorta and Ductus Arteriosus (DA) libraries at each time point using the same parameters as in panel **a**.
- **C.** UMAP analysis and heat-map style representation of canonical lineage markers (color key represents expression level).

**Supplementary Table 1:** Average expression per gene in each cell cluster derived from data generated from scRNAseq libraries: E18.5, P0.5,P5.

| Gene Symbol | Endothelial cell | Smooth muscle | Fibroblast |
|---|---|---|---|
| Eln | 5.592 | 4.835 | 3.890 |
| Malat1 | 4.998 | 5.722 | 5.412 |
| Tpt1 | 4.324 | 3.998 | 4.171 |
| Sparc | 4.213 | 3.901 | 4.537 |
| Eef1a1 | 3.987 | 4.017 | 4.129 |
| Mgp | 3.895 | 4.117 | 3.086 |
| Fbln5 | 3.877 | 3.229 | 2.135 |
| Tm4sf1 | 3.807 | 1.934 | 1.081 |
| Bgn | 3.699 | 3.079 | 3.022 |
| Tmsb4x | 3.646 | 3.897 | 3.470 |
| Vim | 3.578 | 3.891 | 3.682 |
| Actb | 3.409 | 3.913 | 3.435 |
| Fstl1 | 3.356 | 2.722 | 2.887 |
| Fau | 3.265 | 3.036 | 3.097 |
| Ftl1 | 3.232 | 2.900 | 3.233 |
| Fbln2 | 3.231 | 1.243 | 2.026 |
| Ubb | 3.220 | 3.396 | 3.471 |
| Ptma | 3.207 | 3.196 | 2.747 |
| Hsp90ab1 | 3.185 | 3.141 | 2.953 |
| Fn1 | 3.175 | 2.440 | 2.152 |
| Actg1 | 3.063 | 3.143 | 2.928 |
| Ppia | 3.037 | 2.875 | 2.605 |
| Ramp2 | 3.032 | 0.120 | 0.964 |
| Myl6 | 2.973 | 3.428 | 2.644 |
| Eif1 | 2.934 | 3.057 | 3.018 |
| Fth1 | 2.882 | 3.002 | 2.995 |
| Col1a2 | 2.833 | 3.170 | 4.587 |
| Tmsb10 | 2.817 | 1.836 | 2.409 |
| H3f3b | 2.808 | 3.229 | 2.785 |
| Gnas | 2.737 | 2.573 | 3.034 |
| Itm2b | 2.723 | 2.203 | 2.323 |
| H3f3a | 2.700 | 2.849 | 2.544 |
| Egfl7 | 2.672 | 0.078 | 0.092 |
| Pecam1 | 2.671 | 0.074 | 0.059 |
| Gm42418 | 2.663 | 2.366 | 2.428 |
| Serf2 | 2.625 | 2.289 | 2.543 |
| Hspa8 | 2.611 | 2.662 | 2.505 |
| Cst3 | 2.575 | 2.386 | 2.816 |
| Ifitm2 | 2.571 | 1.915 | 2.360 |
| Rack1 | 2.564 | 2.616 | 2.658 |
| Tagln2 | 2.555 | 1.692 | 1.478 |
| Col5a2 | 2.534 | 1.102 | 2.146 |
| Bsg | 2.515 | 1.679 | 1.930 |
| Htra1 | 2.496 | 1.359 | 0.854 |

| | | | |
|---|---|---|---|
| Rbp1 | 2.489 | 1.290 | 1.996 |
| Meg3 | 2.484 | 2.506 | 3.821 |
| Cpe | 2.483 | 2.240 | 1.984 |
| Eef2 | 2.477 | 2.489 | 2.531 |
| Apoe | 2.406 | 1.030 | 1.346 |
| Ctsh | 2.387 | 0.357 | 0.208 |
| Oaz1 | 2.382 | 2.292 | 2.189 |
| Calm1 | 2.337 | 2.486 | 1.800 |
| Id3 | 2.329 | 2.600 | 2.296 |
| Cd34 | 2.312 | 0.428 | 1.335 |
| Naca | 2.311 | 2.431 | 2.302 |
| Chchd2 | 2.290 | 2.102 | 1.932 |
| Cd81 | 2.282 | 2.242 | 2.360 |
| Crip2 | 2.246 | 0.939 | 0.617 |
| Lgals1 | 2.238 | 3.142 | 3.220 |
| Gapdh | 2.238 | 2.068 | 1.800 |
| Gxylt2 | 2.234 | 1.211 | 1.264 |
| Eef1b2 | 2.234 | 2.131 | 2.294 |
| Gnai2 | 2.231 | 1.730 | 1.633 |
| Ubc | 2.217 | 2.618 | 2.447 |
| Serpinh1 | 2.214 | 2.213 | 2.721 |
| Cav1 | 2.206 | 1.139 | 0.445 |
| Fkbp1a | 2.178 | 1.382 | 1.360 |
| Itgb1 | 2.161 | 1.932 | 1.516 |
| Cald1 | 2.158 | 2.698 | 1.749 |
| Gabarap | 2.157 | 1.960 | 2.049 |
| Cox4i1 | 2.151 | 2.206 | 2.047 |
| Cldn5 | 2.145 | 0.048 | 0.043 |
| Hspg2 | 2.139 | 1.636 | 1.028 |
| Ybx1 | 2.120 | 2.022 | 1.881 |
| Timp2 | 2.117 | 1.694 | 1.984 |
| Ddx5 | 2.112 | 2.279 | 1.962 |
| Fabp5 | 2.104 | 0.607 | 1.073 |
| Ptprb | 2.095 | 0.043 | 0.070 |
| Igfbp4 | 2.094 | 0.796 | 2.339 |
| Cfl1 | 2.088 | 2.073 | 1.836 |
| Col3a1 | 2.085 | 1.874 | 4.433 |
| Ctsl | 2.084 | 1.748 | 2.211 |
| Vwf | 2.082 | 0.030 | 0.035 |
| Clic1 | 2.080 | 1.471 | 1.501 |
| Npm1 | 2.070 | 2.113 | 2.150 |
| Ifitm3 | 2.057 | 1.415 | 1.660 |
| Ltbp4 | 2.042 | 1.598 | 1.476 |
| Arpc1b | 2.037 | 1.770 | 1.769 |
| Tspan6 | 2.025 | 0.388 | 0.397 |
| Ppib | 2.010 | 1.847 | 2.250 |
| Jund | 1.983 | 2.695 | 2.711 |
| Crip1 | 1.982 | 1.421 | 1.601 |

| | | | |
|---|---|---|---|
| Cd63 | 1.981 | 2.334 | 2.418 |
| Dynll1 | 1.978 | 1.981 | 1.819 |
| Csrp2 | 1.975 | 2.602 | 1.900 |
| Gng5 | 1.969 | 1.536 | 1.610 |
| Cd93 | 1.965 | 0.085 | 0.060 |
| Klf2 | 1.962 | 2.474 | 2.173 |
| Cavin2 | 1.957 | 0.357 | 0.414 |
| Tmem158 | 1.954 | 0.351 | 0.889 |
| Ptms | 1.939 | 2.122 | 1.814 |
| Rbm3 | 1.938 | 1.961 | 1.788 |
| Hsp90b1 | 1.926 | 1.516 | 1.690 |
| Btf3 | 1.915 | 1.972 | 1.836 |
| Selenow | 1.908 | 1.668 | 1.860 |
| Myl12a | 1.900 | 1.626 | 1.537 |
| Nedd4 | 1.898 | 2.057 | 2.106 |
| Nfib | 1.896 | 1.686 | 2.170 |
| Bmp4 | 1.892 | 0.156 | 0.215 |
| Cox8a | 1.891 | 1.743 | 1.679 |
| Prdx1 | 1.888 | 1.734 | 1.645 |
| Slc25a4 | 1.880 | 2.213 | 2.007 |
| Slc25a3 | 1.876 | 1.719 | 1.468 |
| Calr | 1.867 | 1.601 | 1.798 |
| Cytl1 | 1.862 | 0.057 | 0.033 |
| Hmgb1 | 1.858 | 1.871 | 1.682 |
| Sem1 | 1.857 | 1.505 | 1.451 |
| Lamp1 | 1.847 | 1.352 | 1.425 |
| Tsc22d1 | 1.844 | 2.559 | 1.769 |
| Dstn | 1.838 | 2.466 | 1.706 |
| Ndufa4 | 1.836 | 1.553 | 1.358 |
| Pam | 1.828 | 0.987 | 0.955 |
| Sfrp2 | 1.824 | 2.119 | 1.583 |
| Uqcrh | 1.823 | 1.698 | 1.692 |
| Ctla2a | 1.810 | 0.030 | 0.034 |
| Anxa2 | 1.808 | 1.397 | 1.927 |
| Rrbp1 | 1.793 | 1.763 | 2.111 |
| Rhoa | 1.789 | 1.661 | 1.578 |
| S100a11 | 1.787 | 1.701 | 1.957 |
| Mfap5 | 1.786 | 1.087 | 2.449 |
| Acta2 | 1.779 | 3.768 | 1.757 |
| Emcn | 1.777 | 0.016 | 0.021 |
| Gm10076 | 1.777 | 1.506 | 1.793 |
| Rcn3 | 1.770 | 1.518 | 2.089 |
| Ctnnb1 | 1.763 | 1.436 | 1.123 |
| Pfn1 | 1.758 | 1.854 | 1.457 |
| Atp5e | 1.756 | 1.595 | 1.694 |
| Tubb5 | 1.743 | 1.854 | 1.832 |
| Icam2 | 1.742 | 0.020 | 0.033 |
| Tmem59 | 1.739 | 1.464 | 1.517 |

| | | | |
|---|---|---|---|
| Aplp2 | 1.739 | 1.528 | 1.032 |
| Arpc2 | 1.734 | 1.928 | 1.599 |
| Eif3f | 1.732 | 1.572 | 1.685 |
| Atp5g2 | 1.731 | 1.614 | 1.379 |
| Pfdn5 | 1.730 | 1.708 | 1.662 |
| Vcan | 1.728 | 1.395 | 1.364 |
| Atp5h | 1.728 | 1.568 | 1.510 |
| Laptm4a | 1.727 | 1.662 | 1.889 |
| Dad1 | 1.725 | 1.478 | 1.442 |
| Sec61g | 1.713 | 1.599 | 1.859 |
| Sptbn1 | 1.710 | 1.305 | 1.672 |
| Akr1a1 | 1.707 | 1.345 | 1.406 |
| Cox7c | 1.705 | 1.663 | 1.505 |
| Eif4a1 | 1.700 | 1.776 | 1.641 |
| Foxp1 | 1.696 | 1.489 | 1.152 |
| Anxa5 | 1.689 | 1.698 | 1.786 |
| Rbm39 | 1.689 | 1.708 | 1.542 |
| Atp5j | 1.689 | 1.435 | 1.435 |
| Atp5l | 1.684 | 1.627 | 1.516 |
| S100a10 | 1.683 | 1.487 | 2.243 |
| Cox6c | 1.681 | 1.633 | 1.633 |
| Tpm3 | 1.679 | 1.200 | 1.280 |
| Ywhae | 1.677 | 1.321 | 1.417 |
| Ecscr | 1.676 | 0.035 | 0.063 |
| Gja4 | 1.674 | 0.061 | 0.031 |
| Tagln | 1.673 | 3.650 | 1.472 |
| Esam | 1.669 | 0.135 | 0.026 |
| Gnb2 | 1.665 | 1.510 | 1.578 |
| Eif4g2 | 1.663 | 1.437 | 1.305 |
| Prdx2 | 1.662 | 1.459 | 1.478 |
| Elob | 1.661 | 1.635 | 1.689 |
| Eif5a | 1.661 | 1.620 | 1.595 |
| Ppp3ca | 1.661 | 0.575 | 0.649 |
| Mmp2 | 1.659 | 1.106 | 1.467 |
| Morf4l1 | 1.658 | 1.648 | 1.627 |
| Eif3h | 1.657 | 1.570 | 1.435 |
| Mtch1 | 1.654 | 1.207 | 1.776 |
| Sat1 | 1.650 | 0.747 | 0.790 |
| P4hb | 1.649 | 1.168 | 1.393 |
| Fos | 1.648 | 3.511 | 3.217 |
| Dap | 1.644 | 0.777 | 1.137 |
| Ppic | 1.644 | 1.742 | 1.625 |
| Ehd2 | 1.643 | 0.857 | 0.663 |
| Atp5b | 1.640 | 1.592 | 1.456 |
| Fabp4 | 1.640 | 0.986 | 1.047 |
| Actn1 | 1.639 | 1.806 | 0.757 |
| Pdia3 | 1.637 | 1.308 | 1.499 |
| H2afz | 1.632 | 1.359 | 1.344 |

| | | | |
|---|---|---|---|
| Cox7a2l | 1.632 | 1.241 | 1.269 |
| Eef1g | 1.630 | 1.576 | 1.653 |
| Cdc42 | 1.627 | 1.448 | 1.498 |
| Gpx4 | 1.621 | 1.679 | 1.669 |
| Sumo2 | 1.621 | 1.619 | 1.518 |
| Serbp1 | 1.619 | 1.567 | 1.419 |
| Ldha | 1.619 | 1.423 | 1.286 |
| Ssr4 | 1.616 | 1.337 | 1.591 |
| Psap | 1.615 | 1.536 | 1.573 |
| Edn1 | 1.614 | 0.283 | 0.098 |
| Tmed9 | 1.612 | 1.386 | 1.395 |
| Map1lc3b | 1.609 | 1.502 | 1.359 |
| Ace | 1.608 | 0.832 | 0.326 |
| Hint1 | 1.607 | 1.629 | 1.682 |
| Hnrnpk | 1.602 | 1.656 | 1.467 |
| Hspa5 | 1.601 | 1.647 | 1.830 |
| Zyx | 1.598 | 1.173 | 1.078 |
| Sh3bgrl3 | 1.597 | 0.817 | 0.855 |
| Rtn4 | 1.590 | 1.350 | 1.218 |
| Nfix | 1.587 | 1.709 | 1.861 |
| Arf1 | 1.587 | 1.403 | 1.428 |
| Hmgn1 | 1.586 | 1.925 | 1.776 |
| Aes | 1.576 | 1.220 | 1.383 |
| Atp5d | 1.574 | 1.467 | 1.410 |
| Emp3 | 1.571 | 1.500 | 1.499 |
| Jun | 1.571 | 2.261 | 2.426 |
| Capns1 | 1.568 | 1.324 | 1.302 |
| Atp5a1 | 1.568 | 1.465 | 1.389 |
| Ece1 | 1.565 | 0.350 | 0.320 |
| Mrfap1 | 1.564 | 1.653 | 1.558 |
| Arf5 | 1.563 | 1.285 | 1.616 |
| Pdia6 | 1.562 | 1.160 | 1.537 |
| H2-D1 | 1.558 | 0.236 | 0.221 |
| Rac1 | 1.555 | 1.159 | 1.200 |
| Pmepa1 | 1.550 | 1.729 | 1.385 |
| Rab11a | 1.546 | 0.549 | 0.568 |
| Pdlim3 | 1.542 | 1.347 | 0.675 |
| Heg1 | 1.536 | 0.284 | 0.691 |
| Tmem176b | 1.533 | 0.885 | 0.648 |
| Prdx4 | 1.532 | 1.221 | 1.285 |
| Aldoa | 1.531 | 1.284 | 1.126 |
| Hnrnpa1 | 1.526 | 1.481 | 1.254 |
| Lox | 1.525 | 1.178 | 1.590 |
| Sec61b | 1.525 | 1.356 | 1.475 |
| Tie1 | 1.518 | 0.012 | 0.014 |
| Cox7a2 | 1.517 | 1.365 | 1.262 |
| S100a16 | 1.517 | 0.425 | 0.851 |
| Sub1 | 1.515 | 1.460 | 1.451 |

| | | | |
|---|---|---|---|
| App | 1.515 | 1.611 | 1.479 |
| Ppp1ca | 1.512 | 1.221 | 1.293 |
| Selenof | 1.512 | 1.480 | 1.429 |
| Rhoc | 1.509 | 1.483 | 1.570 |
| Psmb1 | 1.506 | 1.293 | 1.244 |
| Atp5j2 | 1.504 | 1.357 | 1.384 |
| Gnb1 | 1.501 | 1.102 | 1.054 |
| Tmed10 | 1.499 | 1.202 | 1.296 |
| Uqcrb | 1.498 | 1.369 | 1.205 |
| Csrp1 | 1.491 | 1.998 | 0.925 |
| Clta | 1.491 | 1.448 | 1.420 |
| Ctsd | 1.486 | 1.134 | 1.258 |
| Ptgis | 1.485 | 1.668 | 1.331 |
| Bpgm | 1.478 | 0.138 | 0.087 |
| Cox6b1 | 1.476 | 1.283 | 1.288 |
| Cox5b | 1.475 | 1.338 | 1.292 |
| Calu | 1.475 | 1.203 | 1.273 |
| Cdh5 | 1.472 | 0.012 | 0.016 |
| Sh3glb1 | 1.471 | 1.134 | 1.143 |
| Sfrp1 | 1.469 | 1.213 | 1.768 |
| Psma7 | 1.468 | 1.141 | 1.106 |
| Anxa3 | 1.468 | 0.409 | 0.514 |
| Txndc5 | 1.468 | 0.695 | 0.891 |
| Atpif1 | 1.463 | 1.496 | 1.295 |
| Rabac1 | 1.462 | 1.361 | 1.356 |
| Eif3k | 1.461 | 1.365 | 1.380 |
| Cd9 | 1.453 | 1.549 | 0.783 |
| Ssr2 | 1.451 | 1.202 | 1.567 |
| H19 | 1.450 | 0.997 | 2.081 |
| Procr | 1.448 | 0.050 | 0.091 |
| Npc2 | 1.447 | 1.002 | 1.361 |
| Arf4 | 1.446 | 1.386 | 1.697 |
| Ncam1 | 1.444 | 1.802 | 0.675 |
| Pebp1 | 1.442 | 1.419 | 1.452 |
| Ephx2 | 1.439 | 0.139 | 0.101 |
| Clu | 1.436 | 0.819 | 0.296 |
| Ncl | 1.435 | 1.619 | 1.434 |
| Arpc3 | 1.434 | 1.178 | 1.090 |
| Nrp1 | 1.433 | 0.437 | 1.294 |
| Ywhab | 1.429 | 1.019 | 1.057 |
| Csnk1a1 | 1.428 | 1.538 | 1.219 |
| Nupr1 | 1.427 | 1.095 | 1.251 |
| Rras | 1.425 | 1.220 | 1.031 |
| Srsf3 | 1.425 | 1.408 | 1.213 |
| Mfap2 | 1.420 | 1.632 | 2.305 |
| Ube2d3 | 1.419 | 1.533 | 1.456 |
| Atp5c1 | 1.418 | 1.338 | 1.294 |
| Gata6 | 1.414 | 1.201 | 1.081 |

| | | | |
|---|---|---|---|
| Tpm4 | 1.412 | 1.313 | 1.165 |
| Gpx1 | 1.407 | 1.306 | 1.034 |
| Dst | 1.405 | 1.204 | 0.855 |
| Arhgdia | 1.403 | 1.034 | 1.101 |
| Ccdc80 | 1.403 | 1.206 | 1.715 |
| Pkm | 1.396 | 1.229 | 1.014 |
| Tmed2 | 1.396 | 1.275 | 1.390 |
| Col1a1 | 1.395 | 2.917 | 4.542 |
| Rbx1 | 1.395 | 1.188 | 1.183 |
| Gkn3 | 1.394 | 0.033 | 0.046 |
| Sfr1 | 1.392 | 1.429 | 1.652 |
| Pabpc1 | 1.392 | 1.407 | 1.519 |
| Krtcap2 | 1.392 | 1.061 | 1.373 |
| Atp5f1 | 1.390 | 1.320 | 1.297 |
| Hnrnpa0 | 1.390 | 1.406 | 1.363 |
| Tuba1a | 1.388 | 1.463 | 1.999 |
| Gng11 | 1.387 | 0.546 | 0.525 |
| Fblim1 | 1.387 | 0.965 | 0.383 |
| Ubl5 | 1.383 | 0.962 | 0.950 |
| Kctd12 | 1.381 | 0.219 | 0.592 |
| Rdx | 1.381 | 1.012 | 1.138 |
| Tma7 | 1.375 | 1.149 | 1.113 |
| Ndufa13 | 1.372 | 1.426 | 1.409 |
| Iqgap1 | 1.370 | 1.235 | 0.817 |
| Ywhaq | 1.370 | 1.203 | 1.088 |
| Erp29 | 1.368 | 1.120 | 1.401 |
| Nedd8 | 1.367 | 1.219 | 1.184 |
| Clec14a | 1.367 | 0.089 | 0.099 |
| Bnip2 | 1.366 | 0.881 | 0.597 |
| Rhob | 1.366 | 1.696 | 1.398 |
| Mmrn2 | 1.366 | 0.078 | 0.050 |
| Phlda3 | 1.364 | 0.363 | 0.630 |
| Tmbim6 | 1.362 | 1.007 | 0.977 |
| Son | 1.360 | 1.500 | 1.252 |
| Ccnd2 | 1.359 | 1.162 | 1.241 |
| Hsp90aa1 | 1.358 | 1.858 | 1.699 |
| Cdh13 | 1.354 | 0.455 | 0.066 |
| Atp6v0e | 1.354 | 1.106 | 1.033 |
| Selenom | 1.352 | 1.517 | 1.721 |
| Tceal9 | 1.351 | 1.436 | 1.538 |
| Gdi2 | 1.347 | 1.216 | 1.271 |
| Fam3c | 1.344 | 0.531 | 0.418 |
| Kif5b | 1.343 | 1.238 | 1.020 |
| Cfh | 1.337 | 0.045 | 0.286 |
| Txn1 | 1.337 | 1.413 | 1.398 |
| Vcp | 1.335 | 1.266 | 1.316 |
| Txnip | 1.335 | 1.264 | 1.042 |
| Msn | 1.334 | 1.141 | 0.757 |

| | | | |
|---|---|---|---|
| Tek | 1.334 | 0.015 | 0.155 |
| Ltbp3 | 1.333 | 1.199 | 0.987 |
| Atox1 | 1.332 | 0.893 | 0.996 |
| Ier2 | 1.331 | 2.185 | 2.084 |
| Selenok | 1.330 | 1.325 | 1.346 |
| Mapk3 | 1.330 | 0.496 | 0.506 |
| Fhl1 | 1.327 | 1.401 | 1.170 |
| Hmcn1 | 1.326 | 0.260 | 0.288 |
| Luc7l2 | 1.326 | 1.272 | 1.227 |
| Mbnl1 | 1.326 | 0.937 | 0.654 |
| Lmo2 | 1.325 | 0.016 | 0.083 |
| Tpm1 | 1.325 | 2.252 | 1.500 |
| Ndufb11 | 1.322 | 1.205 | 1.175 |
| Cnbp | 1.315 | 1.154 | 1.099 |
| Atp5g1 | 1.312 | 1.212 | 1.209 |
| Eif3e | 1.311 | 1.377 | 1.372 |
| Tbca | 1.311 | 1.186 | 1.114 |
| Cox5a | 1.311 | 0.972 | 0.795 |
| Bri3 | 1.306 | 1.745 | 1.404 |
| Smdt1 | 1.304 | 1.030 | 1.096 |
| Fkbp10 | 1.303 | 0.980 | 1.081 |
| Eif3a | 1.303 | 1.223 | 1.096 |
| Mfge8 | 1.302 | 1.127 | 0.921 |
| Eng | 1.299 | 0.249 | 0.388 |
| Maged1 | 1.299 | 1.276 | 1.387 |
| Myl12b | 1.299 | 0.445 | 0.926 |
| Ddost | 1.298 | 0.965 | 1.267 |
| Swi5 | 1.296 | 1.070 | 1.257 |
| Btg1 | 1.293 | 1.588 | 1.063 |
| Hnrnpa2b1 | 1.293 | 1.219 | 1.146 |
| Nfic | 1.291 | 1.037 | 1.114 |
| Uba52 | 1.285 | 0.991 | 1.176 |
| Cavin3 | 1.283 | 1.196 | 1.221 |
| Rab2a | 1.281 | 1.162 | 1.232 |
| Fkbp9 | 1.281 | 0.879 | 1.312 |
| Ptp4a2 | 1.281 | 1.207 | 1.332 |
| Nsa2 | 1.281 | 1.260 | 1.368 |
| Slc38a2 | 1.280 | 1.756 | 1.329 |
| Ctsb | 1.275 | 1.124 | 1.193 |
| Trim47 | 1.274 | 0.726 | 0.550 |
| Dnaja1 | 1.273 | 1.465 | 1.517 |
| Eif5 | 1.272 | 1.062 | 1.007 |
| Tspo | 1.271 | 1.150 | 0.852 |
| Pcbp2 | 1.271 | 1.351 | 1.173 |
| Cdc42ep3 | 1.269 | 1.018 | 0.579 |
| Anp32a | 1.268 | 0.893 | 0.884 |
| Snrpe | 1.268 | 1.120 | 1.092 |
| Uqcr11 | 1.266 | 1.008 | 0.938 |

| | | | |
|---|---|---|---|
| Sox4 | 1.265 | 1.231 | 1.025 |
| Hdlbp | 1.264 | 1.085 | 1.287 |
| Tomm20 | 1.261 | 1.263 | 0.948 |
| Ptov1 | 1.261 | 1.346 | 1.435 |
| Rpn2 | 1.259 | 0.954 | 1.181 |
| Ehd4 | 1.259 | 0.570 | 0.349 |
| Ndufb9 | 1.258 | 1.260 | 1.189 |
| Rtraf | 1.257 | 1.241 | 1.293 |
| Psma3 | 1.256 | 1.237 | 1.143 |
| Ckap4 | 1.254 | 1.280 | 1.646 |
| Park7 | 1.253 | 1.090 | 1.104 |
| Eef1d | 1.252 | 1.255 | 1.341 |
| Ndufa2 | 1.250 | 1.216 | 1.236 |
| Atp2b1 | 1.249 | 0.797 | 1.097 |
| Reep5 | 1.249 | 1.242 | 1.149 |
| Sulf1 | 1.247 | 0.873 | 0.775 |
| Cyba | 1.246 | 0.888 | 0.896 |
| Plpp1 | 1.244 | 0.716 | 0.670 |
| Bmp6 | 1.243 | 0.005 | 0.009 |
| Ndufc2 | 1.242 | 1.065 | 0.971 |
| Cavin1 | 1.242 | 1.199 | 0.890 |
| Atp5o.1 | 1.242 | 1.120 | 1.219 |
| Atxn7l3b | 1.239 | 1.097 | 1.197 |
| Srrm2 | 1.238 | 1.270 | 1.232 |
| Psmb6 | 1.237 | 1.189 | 1.062 |
| Spcs1 | 1.236 | 1.253 | 1.229 |
| Lmcd1 | 1.236 | 0.529 | 0.197 |
| Hnrnpf | 1.233 | 1.201 | 1.114 |
| Edf1 | 1.228 | 0.976 | 0.968 |
| Ube2i | 1.226 | 0.946 | 0.915 |
| Lmna | 1.225 | 2.072 | 1.773 |
| Marcks | 1.225 | 0.718 | 1.585 |
| Ahnak | 1.224 | 1.559 | 1.724 |
| Set | 1.224 | 1.218 | 1.118 |
| Ctsz | 1.223 | 0.879 | 1.137 |
| Snrpb | 1.223 | 1.198 | 1.094 |
| Fus | 1.222 | 1.430 | 1.440 |
| Sec62 | 1.222 | 1.135 | 1.229 |
| Rbpms | 1.219 | 1.598 | 0.750 |
| Calm2 | 1.218 | 1.688 | 1.555 |
| Kdelr2 | 1.218 | 0.922 | 1.210 |
| Grn | 1.216 | 0.826 | 1.025 |
| Nfia | 1.213 | 1.193 | 1.431 |
| Junb | 1.212 | 2.925 | 2.574 |
| Nap1l1 | 1.211 | 1.079 | 1.211 |
| Dennd5b | 1.210 | 0.277 | 0.249 |
| Mical2 | 1.209 | 1.046 | 0.560 |
| Ddah2 | 1.209 | 0.710 | 1.226 |

| | | | |
|---|---|---|---|
| Ndufa6 | 1.209 | 1.165 | 0.977 |
| Fis1 | 1.207 | 0.774 | 0.838 |
| Eif3i | 1.203 | 1.060 | 1.086 |
| Anp32b | 1.201 | 1.169 | 1.204 |
| Mcl1 | 1.200 | 1.355 | 1.278 |
| Prelid1 | 1.198 | 0.871 | 0.968 |
| Uqcr10 | 1.197 | 1.073 | 1.036 |
| Efnb2 | 1.195 | 0.396 | 0.186 |
| Ube2s | 1.195 | 1.189 | 1.093 |
| Snx3 | 1.193 | 0.936 | 0.927 |
| Id2 | 1.192 | 1.431 | 1.186 |
| Atp6v1g1 | 1.192 | 1.135 | 0.991 |
| Pde3a | 1.189 | 1.049 | 0.512 |
| Arl1 | 1.189 | 1.068 | 1.240 |
| Psmb4 | 1.188 | 1.076 | 1.103 |
| Maged2 | 1.188 | 1.303 | 1.881 |
| Sf3b1 | 1.187 | 1.198 | 1.100 |
| Serinc3 | 1.186 | 0.783 | 0.604 |
| Hnrnpa3 | 1.182 | 1.271 | 1.131 |
| Tmed3 | 1.180 | 0.948 | 1.468 |
| D8Ertd738e | 1.179 | 1.088 | 1.051 |
| Rab14 | 1.176 | 1.081 | 1.025 |
| Kdelr1 | 1.175 | 0.967 | 1.182 |
| H1f0 | 1.174 | 0.823 | 0.916 |
| F11r | 1.174 | 0.168 | 0.121 |
| Tmem234 | 1.173 | 0.910 | 0.896 |
| Fndc3b | 1.171 | 1.081 | 1.003 |
| Atp1a1 | 1.171 | 0.819 | 0.552 |
| Eif4h | 1.169 | 0.969 | 0.904 |
| Cnpy2 | 1.169 | 0.959 | 1.158 |
| Uqcrq | 1.169 | 0.950 | 0.832 |
| Cgnl1 | 1.167 | 0.333 | 0.352 |
| Rcn1 | 1.167 | 0.870 | 1.346 |
| Srgn | 1.166 | 0.027 | 0.029 |
| F2r | 1.166 | 0.504 | 0.250 |
| Cyr61 | 1.163 | 0.942 | 1.004 |
| Wwtr1 | 1.161 | 0.672 | 0.500 |
| Srsf2 | 1.157 | 1.204 | 1.049 |
| Egr1 | 1.155 | 2.958 | 2.818 |
| Vdac2 | 1.154 | 1.166 | 0.937 |
| 1810037I17R | 1.153 | 0.870 | 0.718 |
| Ndufb5 | 1.147 | 1.024 | 1.020 |
| Nisch | 1.147 | 1.163 | 1.263 |
| Ywhaz | 1.145 | 1.124 | 0.955 |
| Klf4 | 1.145 | 1.291 | 1.543 |
| Ndufa7 | 1.144 | 1.170 | 1.168 |
| Cope | 1.143 | 0.879 | 1.023 |
| Mcfd2 | 1.140 | 0.356 | 0.494 |

| | | | |
|---|---|---|---|
| Ablim1 | 1.136 | 0.568 | 0.822 |
| Cox6a1 | 1.134 | 0.990 | 0.942 |
| Tcf25 | 1.134 | 1.004 | 1.101 |
| Srp9 | 1.133 | 1.005 | 0.991 |
| Psmd8 | 1.133 | 1.100 | 0.904 |
| Capzb | 1.133 | 0.989 | 0.879 |
| Psmb5 | 1.131 | 1.027 | 0.929 |
| Tmem50a | 1.131 | 1.079 | 1.032 |
| Zbtb20 | 1.127 | 1.127 | 1.303 |
| Slc9a3r2 | 1.127 | 0.205 | 0.301 |
| Atp5k | 1.126 | 0.982 | 0.972 |
| Mbnl2 | 1.125 | 1.030 | 1.017 |
| Scp2 | 1.125 | 0.930 | 0.640 |
| Bmx | 1.124 | 0.048 | 0.034 |
| H2-K1 | 1.121 | 0.178 | 0.177 |
| Sfpq | 1.121 | 1.048 | 0.999 |
| Tmem167 | 1.120 | 0.924 | 1.127 |
| Sar1a | 1.120 | 0.813 | 1.045 |
| Efna5 | 1.118 | 0.468 | 0.380 |
| Tns1 | 1.118 | 1.264 | 0.792 |
| Fxyd5 | 1.118 | 0.238 | 0.205 |
| 7-Sep | 1.117 | 1.212 | 1.268 |
| Srsf5 | 1.117 | 1.423 | 1.277 |
| Txndc17 | 1.115 | 0.898 | 1.003 |
| H2afv | 1.115 | 1.063 | 1.072 |
| Nenf | 1.114 | 1.255 | 1.619 |
| Arpc5 | 1.114 | 0.824 | 0.835 |
| Pcbp1 | 1.112 | 0.996 | 1.027 |
| Vdac3 | 1.108 | 0.925 | 0.900 |
| Minos1 | 1.107 | 1.074 | 0.966 |
| Emilin1 | 1.105 | 1.043 | 1.000 |
| Eif2s2 | 1.104 | 1.339 | 1.278 |
| Psma4 | 1.104 | 1.015 | 0.998 |
| Psma2 | 1.103 | 1.068 | 0.933 |
| Snrpd2 | 1.102 | 1.053 | 1.066 |
| Sri | 1.098 | 0.734 | 0.944 |
| Prkar1a | 1.098 | 1.048 | 0.852 |
| Golim4 | 1.097 | 0.970 | 0.859 |
| Anapc5 | 1.096 | 1.055 | 1.267 |
| Brk1 | 1.093 | 0.978 | 0.918 |
| Lrpap1 | 1.093 | 0.747 | 0.814 |
| Myct1 | 1.092 | 0.019 | 0.007 |
| Myh9 | 1.090 | 0.917 | 0.554 |
| Pomp | 1.089 | 0.933 | 0.907 |
| Car8 | 1.088 | 0.009 | 0.017 |
| Creb3l2 | 1.088 | 0.882 | 0.601 |
| Abhd17a | 1.088 | 0.496 | 0.564 |
| Sdf4 | 1.087 | 0.688 | 0.822 |

| | | | |
|---|---|---|---|
| Paip2 | 1.087 | 0.952 | 0.975 |
| Pdlim1 | 1.085 | 0.550 | 0.286 |
| Dynlrb1 | 1.084 | 1.057 | 0.984 |
| Emc10 | 1.084 | 0.968 | 0.988 |
| Arglu1 | 1.083 | 0.926 | 0.813 |
| Flna | 1.081 | 1.897 | 1.088 |
| Ndufb7 | 1.081 | 0.825 | 0.754 |
| Ube2m | 1.081 | 0.903 | 0.907 |
| Pon2 | 1.076 | 0.258 | 0.213 |
| Ostc | 1.075 | 0.899 | 1.181 |
| Tcf4 | 1.075 | 0.705 | 0.822 |
| Nfkbia | 1.073 | 1.347 | 1.228 |
| Itm2c | 1.072 | 0.689 | 0.814 |
| Trir | 1.071 | 0.892 | 0.824 |
| Atp5g3 | 1.070 | 1.015 | 0.933 |
| Cyb5r3 | 1.069 | 0.978 | 0.692 |
| Psmb2 | 1.069 | 0.903 | 0.904 |
| Tomm7 | 1.069 | 0.915 | 0.855 |
| Cd151 | 1.067 | 0.315 | 0.250 |
| Mxd4 | 1.066 | 0.900 | 0.897 |
| Atrx | 1.065 | 1.017 | 0.928 |
| Canx | 1.065 | 0.883 | 0.863 |
| Ttc28 | 1.064 | 0.879 | 0.750 |
| Top1 | 1.064 | 1.030 | 0.993 |
| Morf4l2 | 1.061 | 1.018 | 1.128 |
| Bnip3l | 1.060 | 0.864 | 0.974 |
| Neat1 | 1.059 | 1.473 | 0.964 |
| Tpr | 1.058 | 0.975 | 0.914 |
| Lman1 | 1.057 | 0.709 | 0.914 |
| Prdx5 | 1.057 | 1.000 | 1.131 |
| Psmb3 | 1.057 | 0.901 | 0.786 |
| Dbi | 1.056 | 1.264 | 1.141 |
| Cox7b | 1.056 | 1.035 | 0.977 |
| Mgst3 | 1.056 | 0.445 | 0.130 |
| Tprgl | 1.055 | 0.886 | 0.893 |
| Glg1 | 1.055 | 1.006 | 0.972 |
| Ddx3x | 1.055 | 1.097 | 0.977 |
| Chmp2a | 1.052 | 0.837 | 0.829 |
| AU021092 | 1.051 | 0.007 | 0.008 |
| Ybx3 | 1.049 | 0.982 | 1.230 |
| Psmd7 | 1.049 | 1.068 | 0.914 |
| Rapgef5 | 1.047 | 0.233 | 0.044 |
| Rbm25 | 1.046 | 1.028 | 0.895 |
| Tmod3 | 1.046 | 0.353 | 0.429 |
| Atp6v1f | 1.045 | 0.956 | 0.823 |
| Rnf7 | 1.044 | 0.872 | 0.780 |
| Ninj1 | 1.044 | 0.604 | 0.283 |
| Csde1 | 1.044 | 1.021 | 0.940 |

| | | | |
|---|---|---|---|
| Cct2 | 1.042 | 1.073 | 0.931 |
| Snrpg | 1.041 | 0.884 | 1.008 |
| Ehd1 | 1.041 | 0.403 | 0.588 |
| Ran | 1.040 | 0.962 | 0.945 |
| Eif3c | 1.039 | 0.951 | 0.891 |
| Purb | 1.039 | 1.163 | 0.910 |
| Slc25a5 | 1.037 | 1.080 | 0.735 |
| Actn4 | 1.035 | 0.896 | 0.575 |
| B2m | 1.034 | 0.793 | 0.906 |
| Prrc2c | 1.034 | 1.143 | 1.187 |
| Ube2b | 1.034 | 1.180 | 1.036 |
| Tnfaip1 | 1.033 | 0.392 | 0.473 |
| Notch1 | 1.032 | 0.249 | 0.127 |
| Atp6v0b | 1.031 | 0.883 | 0.776 |
| Mdh2 | 1.030 | 0.997 | 0.847 |
| Pgam1 | 1.030 | 0.837 | 0.890 |
| Pafah1b1 | 1.029 | 0.977 | 0.967 |
| Tbrg1 | 1.028 | 0.911 | 1.126 |
| H2afj | 1.028 | 0.762 | 0.695 |
| Mxra8 | 1.027 | 1.144 | 1.274 |
| Nrip1 | 1.027 | 0.785 | 0.826 |
| Sparcl1 | 1.026 | 0.585 | 0.476 |
| Smad7 | 1.025 | 0.691 | 0.365 |
| Mmp14 | 1.025 | 1.442 | 1.509 |
| Plagl1 | 1.025 | 0.674 | 1.553 |
| Hnrnpab | 1.025 | 0.900 | 0.880 |
| Bzw1 | 1.024 | 0.880 | 0.975 |
| Ankrd11 | 1.024 | 1.009 | 0.989 |
| Ddah1 | 1.022 | 0.104 | 0.189 |
| Ppp1r14b | 1.022 | 0.863 | 1.019 |
| Tuba1b | 1.020 | 1.063 | 1.216 |
| Sod3 | 1.020 | 0.433 | 0.588 |
| Cct5 | 1.018 | 1.002 | 0.985 |
| Sptan1 | 1.017 | 0.660 | 0.714 |
| Tinagl1 | 1.016 | 0.234 | 0.051 |
| Myzap | 1.016 | 0.017 | 0.011 |
| Ndufb8 | 1.016 | 0.883 | 0.862 |
| 2010107E04F | 1.015 | 0.849 | 0.945 |
| Ppp2ca | 1.014 | 1.071 | 0.998 |
| Dusp1 | 1.014 | 1.879 | 1.394 |
| 2410015M20 | 1.013 | 0.862 | 0.914 |
| Nucb1 | 1.012 | 0.768 | 0.800 |
| Kmt2e | 1.011 | 0.928 | 0.885 |
| Anxa6 | 1.011 | 1.108 | 1.038 |
| Hspe1 | 1.011 | 1.163 | 1.219 |
| Selenop | 1.010 | 0.644 | 1.117 |
| Calcrl | 1.010 | 0.090 | 0.073 |
| Stub1 | 1.009 | 0.860 | 0.995 |

| | | | |
|---|---|---|---|
| Ndufa11 | 1.009 | 1.112 | 1.018 |
| Colgalt1 | 1.009 | 0.762 | 1.006 |
| Paics | 1.008 | 0.560 | 0.568 |
| Mcrip1 | 1.008 | 0.766 | 1.024 |
| Acer2 | 1.007 | 0.028 | 0.026 |
| Hnrnpu | 1.006 | 1.025 | 0.992 |
| Ypel3 | 1.006 | 0.814 | 1.020 |
| Tpi1 | 1.004 | 0.931 | 0.846 |
| Ssb | 1.003 | 1.001 | 0.933 |
| Tax1bp1 | 1.001 | 0.899 | 0.887 |
| Nrep | 1.001 | 0.770 | 0.936 |
| Pnrc1 | 1.000 | 1.331 | 1.228 |
| Pcsk5 | 0.999 | 0.373 | 0.457 |
| Sod1 | 0.998 | 0.519 | 0.497 |
| Tubb4b | 0.997 | 0.862 | 0.782 |
| Fbn1 | 0.997 | 1.504 | 2.095 |
| Eif4a2 | 0.997 | 1.157 | 1.032 |
| Serpinb6b | 0.996 | 0.373 | 0.101 |
| Hsbp1 | 0.996 | 0.918 | 0.941 |
| Eno1 | 0.994 | 0.721 | 0.582 |
| Sec11a | 0.992 | 0.996 | 1.036 |
| H13 | 0.990 | 0.660 | 0.679 |
| Adamts1 | 0.990 | 1.675 | 1.183 |
| Nxn | 0.989 | 0.716 | 0.581 |
| Thra | 0.988 | 1.413 | 1.297 |
| Ttc3 | 0.988 | 1.035 | 1.181 |
| Npdc1 | 0.987 | 0.730 | 0.759 |
| Cd2ap | 0.987 | 0.515 | 0.418 |
| Tm9sf3 | 0.986 | 0.897 | 0.844 |
| Icam1 | 0.986 | 0.351 | 0.525 |
| Srsf11 | 0.983 | 0.943 | 0.913 |
| Ifngr1 | 0.983 | 0.338 | 0.340 |
| Tmem14c | 0.982 | 0.762 | 0.744 |
| Usmg5 | 0.982 | 0.874 | 0.858 |
| Ndufs5 | 0.982 | 0.939 | 0.877 |
| Mtdh | 0.981 | 1.158 | 1.065 |
| Cdk4 | 0.980 | 0.811 | 0.949 |
| Fermt2 | 0.979 | 1.435 | 0.868 |
| Tcp1 | 0.978 | 1.022 | 0.977 |
| Gpi1 | 0.978 | 0.931 | 0.758 |
| Pdap1 | 0.976 | 0.883 | 0.928 |
| S100a13 | 0.976 | 0.615 | 0.763 |
| Capza2 | 0.975 | 0.954 | 0.825 |
| Nid1 | 0.975 | 0.688 | 1.558 |
| Mif | 0.974 | 0.877 | 0.836 |
| Arpp19 | 0.973 | 0.824 | 0.752 |
| Pbrm1 | 0.973 | 1.066 | 0.922 |
| Smpdl3a | 0.972 | 0.537 | 0.390 |

| | | | |
|---|---|---|---|
| Gsk3b | 0.969 | 0.886 | 0.844 |
| Map4k4 | 0.968 | 0.953 | 0.883 |
| Col5a1 | 0.967 | 1.268 | 2.094 |
| Scand1 | 0.967 | 0.935 | 0.957 |
| Sin3b | 0.967 | 0.705 | 0.764 |
| Cirbp | 0.967 | 1.094 | 0.819 |
| Id1 | 0.964 | 1.110 | 0.689 |
| St3gal5 | 0.964 | 0.468 | 0.318 |
| Tmem2 | 0.963 | 0.154 | 0.086 |
| Eny2 | 0.963 | 0.867 | 0.904 |
| Tulp4 | 0.963 | 0.404 | 0.405 |
| Spag9 | 0.961 | 0.737 | 0.775 |
| Pdlim7 | 0.961 | 1.055 | 0.762 |
| Mlf2 | 0.961 | 0.905 | 0.773 |
| 1110008F13F | 0.960 | 0.806 | 0.651 |
| Dctn2 | 0.959 | 0.843 | 0.883 |
| Tspan7 | 0.958 | 0.121 | 0.179 |
| Ski | 0.957 | 0.953 | 0.934 |
| Ccnd3 | 0.957 | 0.933 | 1.173 |
| Serinc1 | 0.956 | 0.957 | 0.777 |
| Pgls | 0.956 | 0.838 | 0.950 |
| Eif4g1 | 0.955 | 0.819 | 0.843 |
| 1110004F10F | 0.954 | 0.871 | 0.897 |
| Eif3m | 0.954 | 0.994 | 0.896 |
| Reep3 | 0.953 | 0.822 | 0.636 |
| Eif5b | 0.953 | 0.964 | 0.881 |
| Zfp706 | 0.952 | 0.872 | 0.852 |
| Ddx39b | 0.952 | 0.939 | 0.888 |
| Vamp8 | 0.952 | 0.795 | 0.719 |
| Cct7 | 0.949 | 0.848 | 0.762 |
| Sec61a1 | 0.948 | 0.662 | 0.730 |
| Filip1l | 0.948 | 0.774 | 0.683 |
| Imp3 | 0.947 | 0.764 | 0.716 |
| Srpr | 0.947 | 0.649 | 0.675 |
| Snu13 | 0.946 | 0.999 | 0.943 |
| Rheb | 0.946 | 1.042 | 0.812 |
| Rsrp1 | 0.945 | 1.052 | 1.121 |
| Ncor1 | 0.945 | 0.985 | 0.942 |
| Synpo | 0.945 | 0.683 | 0.371 |
| Hnrnpdl | 0.944 | 1.099 | 0.909 |
| Bst2 | 0.944 | 0.175 | 0.251 |
| 1810058I24R | 0.943 | 1.232 | 1.160 |
| Nfe2l1 | 0.940 | 0.576 | 0.517 |
| Ndufa8 | 0.940 | 0.778 | 0.740 |
| Rab7 | 0.939 | 0.720 | 0.736 |
| Ndufb4 | 0.939 | 0.736 | 0.724 |
| Tsn | 0.938 | 0.820 | 0.884 |
| Timm13 | 0.938 | 0.770 | 0.826 |

| | | | |
|---|---|---|---|
| Chd4 | 0.935 | 0.978 | 0.817 |
| Klf13 | 0.935 | 0.994 | 0.810 |
| Manf | 0.934 | 0.783 | 0.911 |
| Sumo1 | 0.934 | 0.907 | 0.896 |
| Fam198b | 0.934 | 0.588 | 0.363 |
| Eid1 | 0.933 | 1.259 | 1.379 |
| Sptssa | 0.931 | 0.835 | 0.716 |
| Mapre1 | 0.931 | 0.881 | 0.874 |
| Cyb5a | 0.930 | 0.773 | 0.923 |
| Macf1 | 0.929 | 0.945 | 0.679 |
| Rab11b | 0.929 | 0.823 | 0.789 |
| Cbx3 | 0.929 | 0.942 | 1.013 |
| S1pr1 | 0.927 | 0.094 | 0.051 |
| Metap2 | 0.926 | 0.910 | 0.799 |
| Ndufb10 | 0.926 | 0.881 | 0.862 |
| Ctnna1 | 0.926 | 0.806 | 0.582 |
| Sf3b2 | 0.926 | 0.981 | 0.965 |
| Apbb2 | 0.925 | 0.232 | 0.269 |
| Mecom | 0.925 | 0.634 | 0.524 |
| Bag1 | 0.925 | 0.803 | 0.895 |
| Srp14 | 0.924 | 0.898 | 0.870 |
| Cltc | 0.923 | 0.885 | 0.742 |
| Grb10 | 0.922 | 1.080 | 1.346 |
| Ergic3 | 0.922 | 0.790 | 0.958 |
| Cdc37 | 0.922 | 0.803 | 0.778 |
| Akt1 | 0.919 | 0.719 | 0.848 |
| Litaf | 0.919 | 0.343 | 0.464 |
| Kdelr3 | 0.918 | 0.704 | 1.028 |
| Tspan3 | 0.917 | 1.022 | 1.090 |
| Psmc3 | 0.917 | 0.811 | 0.793 |
| Mrps24 | 0.916 | 0.768 | 0.773 |
| Rnf187 | 0.915 | 0.855 | 0.828 |
| Tln1 | 0.915 | 1.034 | 0.751 |
| Ube2l3 | 0.913 | 0.788 | 0.761 |
| Rap1b | 0.912 | 0.938 | 1.022 |
| Ap2m1 | 0.912 | 0.815 | 0.825 |
| Fbxl7 | 0.912 | 0.580 | 0.501 |
| Wnk1 | 0.912 | 0.772 | 0.812 |
| Itpr2 | 0.911 | 0.098 | 0.243 |
| Glrx3 | 0.911 | 0.827 | 1.020 |
| Pnp | 0.910 | 0.583 | 0.830 |
| Tmem256 | 0.908 | 0.796 | 0.944 |
| Klf3 | 0.908 | 0.434 | 0.609 |
| Bclaf1 | 0.907 | 0.989 | 0.945 |
| Ywhah | 0.906 | 1.088 | 0.900 |
| Dazap2 | 0.906 | 0.579 | 0.470 |
| G3bp2 | 0.905 | 0.693 | 0.509 |
| Ost4 | 0.905 | 0.684 | 0.768 |

| | | | |
|---|---|---|---|
| Ssfa2 | 0.905 | 0.166 | 0.091 |
| Gabarapl2 | 0.905 | 0.693 | 0.606 |
| Rer1 | 0.905 | 0.838 | 0.868 |
| Polr2m | 0.904 | 0.751 | 0.824 |
| Plpp3 | 0.904 | 0.415 | 0.916 |
| Lims1 | 0.903 | 0.573 | 0.435 |
| Cnn3 | 0.902 | 1.154 | 1.001 |
| Ubxn1 | 0.900 | 0.845 | 0.816 |
| Ufc1 | 0.900 | 0.739 | 0.837 |
| Rnh1 | 0.899 | 0.821 | 0.821 |
| Ndufab1 | 0.899 | 0.787 | 0.666 |
| Spcs2 | 0.898 | 0.839 | 0.798 |
| Esd | 0.898 | 0.994 | 1.001 |
| Hdgf | 0.896 | 0.894 | 0.752 |
| Copz2 | 0.896 | 0.817 | 1.121 |
| Krt18 | 0.895 | 0.314 | 0.053 |
| Dnajb6 | 0.895 | 0.804 | 0.707 |
| Rsrc2 | 0.895 | 0.899 | 0.826 |
| Zcrb1 | 0.895 | 0.781 | 0.675 |
| Ssr3 | 0.894 | 0.806 | 0.902 |
| 0610012G03I | 0.894 | 0.596 | 0.722 |
| Rab1a | 0.894 | 0.733 | 0.789 |
| Polr1d | 0.893 | 1.034 | 0.913 |
| Raly | 0.893 | 0.938 | 0.815 |
| Cenpb | 0.892 | 0.708 | 0.732 |
| Pbx1 | 0.892 | 1.423 | 1.082 |
| Wasf2 | 0.891 | 0.769 | 0.722 |
| Ahsg | 0.887 | 0.018 | 0.021 |
| Dync1i2 | 0.887 | 0.771 | 0.754 |
| P3h3 | 0.886 | 0.795 | 0.942 |
| Uqcrc1 | 0.885 | 0.733 | 0.641 |
| Arpc1a | 0.885 | 0.636 | 0.766 |
| Smoc1 | 0.884 | 0.713 | 0.184 |
| Emc7 | 0.883 | 0.761 | 0.634 |
| Cav2 | 0.882 | 0.332 | 0.077 |
| Sulf2 | 0.882 | 1.037 | 0.969 |
| Ap2s1 | 0.881 | 0.721 | 0.768 |
| Tmem176a | 0.880 | 0.360 | 0.270 |
| Arhgap31 | 0.880 | 0.338 | 0.246 |
| Nktr | 0.879 | 1.030 | 1.134 |
| Nol7 | 0.878 | 0.727 | 0.621 |
| Pdcd5 | 0.878 | 1.049 | 0.906 |
| Ndufv3 | 0.877 | 0.938 | 0.852 |
| Cct8 | 0.877 | 0.946 | 0.923 |
| Ranbp1 | 0.877 | 0.794 | 0.801 |
| Palmd | 0.877 | 0.015 | 0.015 |
| Ube2d2a | 0.876 | 0.881 | 0.814 |
| Ddb1 | 0.875 | 0.723 | 0.835 |

| | | | |
|---|---|---|---|
| Epas1 | 0.875 | 0.214 | 0.057 |
| Serpinb6a | 0.875 | 0.926 | 1.248 |
| Psma6 | 0.874 | 0.878 | 0.750 |
| Nuak1 | 0.873 | 0.308 | 0.214 |
| Txnl1 | 0.873 | 0.769 | 0.827 |
| Slc3a2 | 0.873 | 0.851 | 0.705 |
| Ubxn4 | 0.872 | 0.853 | 0.767 |
| Ndufa12 | 0.872 | 0.839 | 0.730 |
| Lamp2 | 0.871 | 0.872 | 0.987 |
| Aurkaip1 | 0.871 | 0.591 | 0.556 |
| Cthrc1 | 0.871 | 0.526 | 1.167 |
| Ift20 | 0.870 | 0.788 | 0.946 |
| Tra2a | 0.869 | 0.890 | 0.890 |
| Snrnp70 | 0.869 | 1.038 | 0.940 |
| Cracr2b | 0.868 | 0.106 | 0.067 |
| Cd47 | 0.868 | 0.507 | 0.850 |
| Tmx1 | 0.868 | 0.503 | 0.416 |
| Bax | 0.867 | 0.767 | 0.669 |
| Eva1b | 0.867 | 1.352 | 1.091 |
| Pttg1ip | 0.867 | 0.739 | 0.818 |
| Gnaq | 0.866 | 0.504 | 0.427 |
| Map1lc3a | 0.866 | 1.516 | 1.353 |
| Phb2 | 0.865 | 0.773 | 0.809 |
| Surf4 | 0.865 | 0.609 | 0.783 |
| Csnk1e | 0.863 | 0.533 | 0.521 |
| Bex3 | 0.863 | 0.966 | 0.924 |
| Hexa | 0.862 | 0.464 | 0.686 |
| Saraf | 0.862 | 0.804 | 0.613 |
| Ndufc1 | 0.861 | 0.776 | 0.795 |
| Mrpl20 | 0.861 | 0.812 | 0.651 |
| Fuca1 | 0.861 | 0.675 | 0.805 |
| Meis2 | 0.860 | 0.614 | 0.822 |
| Pnisr | 0.860 | 0.975 | 0.894 |
| Sf3b5 | 0.860 | 0.643 | 0.767 |
| Ube2r2 | 0.859 | 0.959 | 0.983 |
| Bptf | 0.859 | 0.915 | 0.869 |
| Scd2 | 0.858 | 0.646 | 0.799 |
| Cd200 | 0.858 | 1.009 | 0.636 |
| Fundc2 | 0.856 | 0.741 | 0.853 |
| Ctbp1 | 0.855 | 0.723 | 0.780 |
| Emp1 | 0.854 | 0.446 | 0.836 |
| Puf60 | 0.854 | 0.707 | 0.751 |
| Mydgf | 0.854 | 0.591 | 0.748 |
| Btg2 | 0.853 | 2.801 | 2.241 |
| Tmem258 | 0.853 | 0.794 | 0.932 |
| Mrpl52 | 0.853 | 0.923 | 0.930 |
| Klf6 | 0.851 | 1.187 | 1.271 |
| Tmem165 | 0.851 | 0.833 | 0.801 |

| | | | |
|---|---|---|---|
| Tm2d2 | 0.851 | 0.763 | 0.722 |
| Myl9 | 0.850 | 2.563 | 1.267 |
| Plk2 | 0.850 | 0.147 | 0.324 |
| Gsn | 0.849 | 1.488 | 2.448 |
| Ndufs2 | 0.848 | 0.851 | 0.734 |
| Matr3 | 0.846 | 0.917 | 0.818 |
| Gata2 | 0.846 | 0.008 | 0.005 |
| Ssbp4 | 0.845 | 0.566 | 0.543 |
| Qk | 0.845 | 0.657 | 0.693 |
| Ndufa3 | 0.844 | 0.676 | 0.811 |
| Romo1 | 0.844 | 0.783 | 0.813 |
| Sra1 | 0.843 | 0.513 | 0.721 |
| Dpysl3 | 0.843 | 0.948 | 1.170 |
| Tgfb1 | 0.842 | 0.455 | 0.221 |
| Creb5 | 0.842 | 0.851 | 0.817 |
| Fzd4 | 0.842 | 0.157 | 0.433 |
| Ptbp1 | 0.841 | 0.685 | 0.664 |
| Tmem109 | 0.840 | 0.516 | 0.428 |
| Ier3ip1 | 0.840 | 0.633 | 0.791 |
| Arl6ip5 | 0.840 | 0.808 | 0.527 |
| Dctn3 | 0.840 | 0.528 | 0.606 |
| Ppig | 0.839 | 0.789 | 0.737 |
| Lamtor1 | 0.839 | 0.704 | 0.698 |
| Aldh2 | 0.839 | 0.881 | 0.901 |
| Gtf2h5 | 0.838 | 0.778 | 0.871 |
| Bcap31 | 0.837 | 0.805 | 0.767 |
| Higd1a | 0.836 | 0.774 | 0.810 |
| Clk1 | 0.836 | 1.025 | 0.933 |
| Snhg18 | 0.836 | 1.013 | 1.199 |
| Gna11 | 0.835 | 0.720 | 0.695 |
| Prdx6 | 0.835 | 0.926 | 0.637 |
| Thoc7 | 0.835 | 0.843 | 0.764 |
| Cops9 | 0.835 | 0.740 | 0.735 |
| U2af1 | 0.834 | 0.850 | 0.708 |
| Dhh | 0.834 | 0.007 | 0.007 |
| Fkbp8 | 0.833 | 0.754 | 0.786 |
| Adipor1 | 0.833 | 0.477 | 0.415 |
| Tceal8 | 0.833 | 0.782 | 0.903 |
| Tanc2 | 0.833 | 0.613 | 0.422 |
| Dek | 0.833 | 1.108 | 0.797 |
| Cct4 | 0.831 | 0.923 | 0.856 |
| Thbs1 | 0.831 | 1.039 | 0.597 |
| Rhoj | 0.831 | 0.627 | 1.131 |
| Csnk2b | 0.830 | 0.650 | 0.581 |
| Trip11 | 0.830 | 0.536 | 0.531 |
| H2afy | 0.829 | 0.679 | 0.830 |
| Ktn1 | 0.828 | 0.873 | 0.493 |
| Ndufs6 | 0.828 | 0.738 | 0.738 |

| | | | |
|---|---|---|---|
| Hnrnpd | 0.827 | 0.860 | 0.742 |
| Slc50a1 | 0.827 | 0.613 | 0.826 |
| Uchl1 | 0.827 | 0.489 | 0.203 |
| Map2k2 | 0.827 | 0.697 | 0.746 |
| Tomm22 | 0.826 | 0.726 | 0.621 |
| Rock2 | 0.825 | 1.041 | 0.774 |
| Arpc4 | 0.825 | 0.638 | 0.610 |
| Nono | 0.825 | 0.717 | 0.701 |
| Hnrnpm | 0.824 | 0.731 | 0.711 |
| Senp6 | 0.823 | 0.783 | 0.723 |
| Calm3 | 0.822 | 0.717 | 0.792 |
| Hotairm1 | 0.822 | 0.478 | 0.371 |
| Sox17 | 0.822 | 0.006 | 0.005 |
| Rasip1 | 0.821 | 0.012 | 0.014 |
| Wwc2 | 0.821 | 0.501 | 0.545 |
| Rtn3 | 0.820 | 0.623 | 0.644 |
| Psma1 | 0.818 | 0.735 | 0.627 |
| Nop10 | 0.818 | 0.629 | 0.651 |
| Sh3bgrl | 0.817 | 1.094 | 0.834 |
| Smc1a | 0.815 | 0.778 | 0.734 |
| Hmgn2 | 0.815 | 0.761 | 0.860 |
| Hnrnpc | 0.815 | 0.714 | 0.714 |
| Clic4 | 0.815 | 1.326 | 0.774 |
| Arhgdib | 0.815 | 0.203 | 0.226 |
| Cetn3 | 0.813 | 0.817 | 0.772 |
| Dynlt3 | 0.813 | 0.626 | 0.416 |
| Atraid | 0.813 | 0.955 | 1.035 |
| Pdcd6 | 0.813 | 0.757 | 0.756 |
| Zfp36l1 | 0.812 | 1.317 | 1.540 |
| Srrm1 | 0.812 | 0.889 | 0.838 |
| Dync1h1 | 0.812 | 0.673 | 0.671 |
| Atf4 | 0.812 | 1.114 | 1.191 |
| Gpx7 | 0.812 | 0.615 | 1.089 |
| Rwdd1 | 0.811 | 0.730 | 0.757 |
| Rbm8a | 0.811 | 0.752 | 0.655 |
| Ppp1cc | 0.811 | 0.694 | 0.699 |
| Cmas | 0.810 | 0.274 | 0.311 |
| Tra2b | 0.810 | 1.050 | 0.894 |
| Copb2 | 0.810 | 0.793 | 0.942 |
| Sf3b6 | 0.809 | 0.747 | 0.752 |
| Plec | 0.808 | 0.683 | 0.519 |
| Tmem30a | 0.808 | 0.536 | 0.458 |
| Cct3 | 0.808 | 0.740 | 0.700 |
| P4ha3 | 0.807 | 0.299 | 0.384 |
| Nucb2 | 0.807 | 0.289 | 0.535 |
| Dnajc3 | 0.807 | 0.781 | 0.801 |
| Prmt1 | 0.807 | 0.863 | 0.743 |
| Ube2h | 0.807 | 0.553 | 0.456 |

| | | | |
|---|---|---|---|
| Idh2 | 0.805 | 1.296 | 1.190 |
| Rgs3 | 0.805 | 0.239 | 0.303 |
| Psmd4 | 0.805 | 0.778 | 0.687 |
| Rexo2 | 0.804 | 1.021 | 0.717 |
| Yif1b | 0.803 | 0.616 | 0.647 |
| Snrpd1 | 0.802 | 0.722 | 0.713 |
| Lamtor5 | 0.802 | 0.714 | 0.635 |
| Psma5 | 0.802 | 0.779 | 0.711 |
| Ndufa5 | 0.801 | 0.821 | 0.806 |
| Vapa | 0.800 | 1.018 | 0.829 |
| Mdk | 0.800 | 1.078 | 1.554 |
| Cdv3 | 0.799 | 0.695 | 0.654 |
| St13 | 0.799 | 0.774 | 0.799 |
| Gm11808 | 0.799 | 0.605 | 0.706 |
| Gltp | 0.799 | 0.262 | 0.305 |
| Flnb | 0.799 | 0.180 | 0.220 |
| Copa | 0.798 | 0.773 | 0.890 |
| Hectd1 | 0.798 | 0.763 | 0.718 |
| Taldo1 | 0.797 | 0.678 | 0.593 |
| Rbms1 | 0.796 | 0.689 | 0.846 |
| Smim7 | 0.796 | 0.612 | 0.732 |
| B230219D22I | 0.795 | 0.767 | 0.635 |
| Ssr1 | 0.795 | 0.667 | 0.728 |
| Trappc6b | 0.794 | 0.671 | 0.617 |
| Leprot | 0.794 | 0.532 | 0.620 |
| Kazald1 | 0.794 | 0.613 | 0.571 |
| Tnfrsf1a | 0.793 | 0.592 | 0.699 |
| She | 0.792 | 0.006 | 0.010 |
| Plvap | 0.792 | 0.083 | 0.072 |
| Lrrc58 | 0.791 | 0.573 | 0.564 |
| Taf10 | 0.791 | 0.397 | 0.491 |
| Igfbp5 | 0.791 | 0.709 | 1.015 |
| Cnih1 | 0.791 | 0.639 | 0.680 |
| Rrp1 | 0.790 | 0.769 | 0.677 |
| Trappc4 | 0.790 | 0.504 | 0.503 |
| Anapc11 | 0.790 | 0.658 | 0.820 |
| Mpc2 | 0.790 | 0.990 | 0.951 |
| Uqcrfs1 | 0.790 | 0.708 | 0.607 |
| Pura | 0.789 | 0.738 | 0.719 |
| Rock1 | 0.789 | 0.938 | 0.621 |
| Sypl | 0.789 | 0.581 | 0.410 |
| Ndufv2 | 0.789 | 0.840 | 0.673 |
| Vamp3 | 0.788 | 0.458 | 0.444 |
| Ddx50 | 0.787 | 0.960 | 1.029 |
| Cdkn1c | 0.787 | 0.749 | 2.201 |
| Tecr | 0.787 | 0.718 | 0.871 |
| Ptbp3 | 0.787 | 0.543 | 0.441 |
| Orai1 | 0.786 | 0.164 | 0.170 |

| | | | |
|---|---|---|---|
| Nipbl | 0.785 | 0.746 | 0.689 |
| Plod1 | 0.785 | 0.247 | 0.421 |
| Fgfr1 | 0.783 | 0.968 | 0.981 |
| Drap1 | 0.783 | 0.975 | 0.883 |
| Nrbp1 | 0.782 | 0.553 | 0.563 |
| Vkorc1 | 0.782 | 0.605 | 1.104 |
| Uqcrc2 | 0.782 | 0.796 | 0.659 |
| Card19 | 0.781 | 0.471 | 0.423 |
| Nxf1 | 0.780 | 0.665 | 0.617 |
| Eif4b | 0.780 | 0.768 | 0.676 |
| Auts2 | 0.779 | 1.209 | 0.909 |
| Selenos | 0.779 | 1.007 | 1.081 |
| Psenen | 0.779 | 0.624 | 0.599 |
| Chmp4b | 0.778 | 0.914 | 0.877 |
| Cbfa2t3 | 0.777 | 0.118 | 0.103 |
| Ube2n | 0.777 | 0.793 | 0.624 |
| Thbd | 0.777 | 0.246 | 0.342 |
| Bcam | 0.777 | 0.299 | 0.086 |
| Kcnq1ot1 | 0.776 | 0.947 | 1.511 |
| B4galt1 | 0.776 | 0.356 | 0.552 |
| Lima1 | 0.775 | 0.343 | 0.763 |
| Capn2 | 0.775 | 0.806 | 0.737 |
| Cenpt | 0.774 | 0.052 | 0.073 |
| Nos3 | 0.774 | 0.008 | 0.009 |
| S100a6 | 0.774 | 1.034 | 1.870 |
| Picalm | 0.774 | 0.864 | 0.639 |
| Slco2a1 | 0.773 | 0.056 | 0.047 |
| Ghr | 0.773 | 0.645 | 0.370 |
| Fam162a | 0.773 | 0.629 | 0.528 |
| Klhdc2 | 0.773 | 0.598 | 0.621 |
| Timp3 | 0.773 | 0.691 | 0.358 |
| Vdac1 | 0.772 | 0.877 | 0.811 |
| Sugt1 | 0.772 | 0.610 | 0.542 |
| Ppp1r2 | 0.772 | 0.607 | 0.638 |
| Elk3 | 0.771 | 0.315 | 0.292 |
| Abracl | 0.771 | 0.718 | 0.756 |
| Rbbp6 | 0.771 | 0.829 | 0.744 |
| Psme1 | 0.770 | 0.567 | 0.535 |
| Mat2a | 0.770 | 0.942 | 1.006 |
| Cpq | 0.768 | 0.589 | 0.502 |
| Sdcbp | 0.767 | 0.977 | 0.976 |
| Guk1 | 0.766 | 0.565 | 0.485 |
| Fubp1 | 0.766 | 0.643 | 0.626 |
| Tmpo | 0.765 | 0.526 | 0.500 |
| Hdac7 | 0.765 | 0.363 | 0.435 |
| Ppp1r12a | 0.764 | 1.669 | 0.752 |
| Asap2 | 0.764 | 0.313 | 0.134 |
| Ostf1 | 0.764 | 0.461 | 0.501 |

| | | | |
|---|---|---|---|
| Hmgn3 | 0.762 | 0.635 | 0.577 |
| Mrps14 | 0.762 | 0.677 | 0.619 |
| Pgk1 | 0.762 | 0.574 | 0.500 |
| Napa | 0.761 | 0.572 | 0.607 |
| Pkd1 | 0.761 | 0.898 | 0.659 |
| Pigt | 0.761 | 0.397 | 0.576 |
| Marveld1 | 0.760 | 0.757 | 0.519 |
| Syf2 | 0.760 | 0.708 | 0.599 |
| Mbd2 | 0.760 | 0.626 | 0.600 |
| Socs2 | 0.759 | 0.950 | 0.595 |
| Rab12 | 0.758 | 0.671 | 0.619 |
| Eif1ax | 0.758 | 0.858 | 0.650 |
| Hp1bp3 | 0.758 | 0.711 | 0.628 |
| Higd2a | 0.758 | 0.623 | 0.503 |
| Mrpl33 | 0.758 | 0.828 | 0.816 |
| Ecm1 | 0.758 | 0.915 | 0.917 |
| Brd2 | 0.758 | 0.839 | 0.725 |
| Pofut2 | 0.757 | 0.775 | 0.891 |
| Itga5 | 0.757 | 0.355 | 0.243 |
| Ppdpf | 0.757 | 0.471 | 0.523 |
| Atp6v1e1 | 0.756 | 0.569 | 0.487 |
| Mrpl54 | 0.756 | 0.654 | 0.668 |
| Ppp4r2 | 0.756 | 0.569 | 0.573 |
| Ube2e3 | 0.755 | 0.700 | 0.577 |
| Bace2 | 0.755 | 0.102 | 0.453 |
| Mrps33 | 0.755 | 0.514 | 0.604 |
| Rab6a | 0.754 | 0.748 | 0.687 |
| Ralbp1 | 0.754 | 0.741 | 0.670 |
| Cops6 | 0.754 | 0.686 | 0.655 |
| Sdhb | 0.754 | 0.632 | 0.673 |
| Zfp91 | 0.753 | 0.705 | 0.588 |
| Polr2f | 0.753 | 0.818 | 0.817 |
| Luzp1 | 0.753 | 0.531 | 0.487 |
| Ndufs7 | 0.752 | 0.662 | 0.668 |
| Eif6 | 0.751 | 0.605 | 0.615 |
| Eif1b | 0.751 | 0.744 | 0.717 |
| Azin1 | 0.750 | 0.446 | 0.366 |
| Srsf9 | 0.750 | 0.748 | 0.642 |
| Stmn2 | 0.750 | 0.013 | 0.047 |
| Aimp1 | 0.750 | 0.790 | 0.768 |
| Sdc1 | 0.750 | 0.280 | 0.330 |
| Twist1 | 0.749 | 0.174 | 0.365 |
| Eif4e2 | 0.748 | 0.669 | 0.820 |
| Sys1 | 0.748 | 0.538 | 0.641 |
| Gclm | 0.748 | 0.358 | 0.317 |
| BC031181 | 0.748 | 0.603 | 0.504 |
| Adipor2 | 0.748 | 0.212 | 0.212 |
| Cebpd | 0.748 | 1.834 | 1.934 |

| | | | |
|---|---|---|---|
| Rnf144a | 0.748 | 0.163 | 0.366 |
| Jag1 | 0.748 | 0.445 | 0.290 |
| Cux1 | 0.747 | 0.727 | 0.425 |
| Iscu | 0.747 | 0.707 | 0.470 |
| Ndufb6 | 0.747 | 0.675 | 0.605 |
| Aamp | 0.747 | 0.702 | 0.688 |
| Dph3 | 0.747 | 0.685 | 0.712 |
| Ier3 | 0.746 | 1.656 | 1.305 |
| Odc1 | 0.744 | 0.622 | 0.635 |
| Actr3 | 0.744 | 0.919 | 0.697 |
| Peg3 | 0.744 | 1.087 | 1.810 |
| Palld | 0.744 | 0.593 | 0.536 |
| C1qtnf6 | 0.743 | 0.663 | 0.908 |
| Dnajc8 | 0.742 | 0.705 | 0.618 |
| Oat | 0.742 | 1.159 | 1.146 |
| Adam10 | 0.740 | 0.233 | 0.193 |
| Map1b | 0.740 | 1.567 | 0.789 |
| Rala | 0.740 | 0.612 | 0.579 |
| Ash1l | 0.740 | 0.800 | 0.725 |
| Nfat5 | 0.739 | 0.752 | 0.657 |
| Mrpl30 | 0.739 | 0.727 | 0.752 |
| Smarca2 | 0.737 | 0.666 | 0.884 |
| Yipf3 | 0.737 | 0.599 | 0.714 |
| Selenoh | 0.737 | 0.446 | 0.527 |
| Ppp1r11 | 0.737 | 0.430 | 0.407 |
| Mtpn | 0.736 | 0.692 | 0.623 |
| Ifi27 | 0.735 | 0.556 | 0.815 |
| Wdr26 | 0.735 | 0.715 | 0.647 |
| Pcmtd1 | 0.733 | 0.459 | 0.429 |
| Actr10 | 0.733 | 0.708 | 0.608 |
| Cbx6 | 0.733 | 0.761 | 0.770 |
| Mef2a | 0.733 | 0.877 | 0.573 |
| Sgk1 | 0.733 | 0.352 | 0.241 |
| Dpm3 | 0.732 | 0.590 | 0.762 |
| Psmd13 | 0.732 | 0.609 | 0.572 |
| Ctsa | 0.732 | 0.616 | 0.645 |
| Lpp | 0.730 | 1.519 | 0.886 |
| Lman2 | 0.730 | 0.523 | 0.574 |
| C1d | 0.730 | 0.606 | 0.595 |
| Cbx1 | 0.729 | 0.628 | 0.617 |
| Cfl2 | 0.729 | 0.846 | 0.649 |
| Zfr | 0.729 | 0.670 | 0.714 |
| Tnfaip8 | 0.729 | 0.444 | 0.205 |
| Lmo4 | 0.728 | 0.736 | 1.136 |
| Marcksl1 | 0.728 | 0.578 | 0.738 |
| Psmd2 | 0.728 | 0.645 | 0.593 |
| Eloc | 0.728 | 0.700 | 0.598 |
| Clec1a | 0.728 | 0.004 | 0.002 |

| | | | |
|---|---|---|---|
| Rab18 | 0.728 | 0.770 | 0.703 |
| Bmp1 | 0.727 | 0.612 | 1.062 |
| Kctd10 | 0.727 | 0.533 | 0.432 |
| Pls3 | 0.727 | 1.204 | 0.887 |
| 9530068E07F | 0.726 | 0.806 | 0.704 |
| Smap1 | 0.725 | 0.754 | 0.757 |
| Ppp2r1a | 0.725 | 0.571 | 0.536 |
| Rex1bd | 0.724 | 0.671 | 0.625 |
| Psmc6 | 0.723 | 0.818 | 0.662 |
| Akap9 | 0.723 | 0.600 | 0.524 |
| Ass1 | 0.723 | 0.033 | 0.027 |
| Mff | 0.723 | 0.606 | 0.579 |
| Lamtor4 | 0.722 | 0.651 | 0.666 |
| Psmc4 | 0.722 | 0.669 | 0.559 |
| Hras | 0.721 | 0.599 | 0.519 |
| Flt1 | 0.721 | 0.153 | 0.148 |
| Ubn1 | 0.721 | 0.596 | 0.706 |
| Ndufb2 | 0.720 | 0.734 | 0.659 |
| Thrap3 | 0.720 | 0.703 | 0.710 |
| Pitpna | 0.719 | 0.621 | 0.516 |
| Igf2r | 0.719 | 0.606 | 0.851 |
| Lrrfip1 | 0.719 | 0.823 | 0.349 |
| Tkt | 0.718 | 0.916 | 0.854 |
| Ddx6 | 0.718 | 0.713 | 0.655 |
| Il11ra1 | 0.718 | 1.366 | 1.469 |
| Aaed1 | 0.718 | 0.543 | 0.461 |
| Sec31a | 0.717 | 0.655 | 0.787 |
| Ilk | 0.717 | 1.166 | 1.007 |
| Ndufs3 | 0.716 | 0.662 | 0.640 |
| Timm10b | 0.716 | 0.504 | 0.572 |
| Tapbp | 0.716 | 0.484 | 0.439 |
| Ube3a | 0.715 | 0.751 | 0.755 |
| Ndrg1 | 0.715 | 0.401 | 0.217 |
| Xbp1 | 0.715 | 0.613 | 0.732 |
| Strn3 | 0.714 | 0.631 | 0.607 |
| Smarce1 | 0.713 | 0.654 | 0.559 |
| 4931406P16F | 0.713 | 0.296 | 0.358 |
| Atxn10 | 0.713 | 0.670 | 0.862 |
| Ctdsp2 | 0.712 | 0.904 | 0.915 |
| Cwc15 | 0.711 | 0.651 | 0.538 |
| Hook3 | 0.711 | 0.581 | 0.706 |
| Timm8b | 0.710 | 0.564 | 0.520 |
| Plod3 | 0.710 | 0.351 | 0.441 |
| Ccnl1 | 0.710 | 1.134 | 1.146 |
| Supt4a | 0.710 | 0.626 | 0.641 |
| Vcam1 | 0.710 | 0.196 | 0.434 |
| Lrrc59 | 0.710 | 0.461 | 0.605 |
| Crim1 | 0.709 | 0.267 | 0.137 |

| | | | |
|---|---|---|---|
| Banf1 | 0.709 | 0.607 | 0.586 |
| Mcam | 0.708 | 0.567 | 0.068 |
| Prpf40a | 0.708 | 0.608 | 0.631 |
| Golgb1 | 0.708 | 0.615 | 0.655 |
| Nckap1 | 0.708 | 0.490 | 0.455 |
| Stab1 | 0.707 | 0.026 | 0.038 |
| Ssu72 | 0.707 | 0.543 | 0.517 |
| Wdr83os | 0.707 | 0.630 | 0.566 |
| Atp6ap1 | 0.706 | 0.521 | 0.493 |
| Slc25a11 | 0.706 | 0.432 | 0.464 |
| Akap13 | 0.706 | 0.718 | 0.795 |
| Ubap2l | 0.706 | 0.712 | 0.772 |
| Tacc1 | 0.706 | 0.753 | 0.469 |
| Nop53 | 0.705 | 0.659 | 0.589 |
| Commd3 | 0.704 | 0.724 | 0.631 |
| Rsu1 | 0.704 | 1.003 | 0.750 |
| Pdcd10 | 0.704 | 0.675 | 0.510 |
| Cpd | 0.703 | 0.344 | 0.345 |
| Rab10 | 0.703 | 0.669 | 0.638 |
| Ryk | 0.703 | 0.684 | 0.787 |
| Itm2a | 0.703 | 0.348 | 2.155 |
| Gimap1 | 0.702 | 0.004 | 0.003 |
| Srp19 | 0.702 | 0.581 | 0.629 |
| Ergic2 | 0.702 | 0.618 | 0.615 |
| Shisa5 | 0.702 | 0.444 | 0.664 |
| Ogt | 0.701 | 0.660 | 0.712 |
| Adh5 | 0.700 | 0.574 | 0.535 |
| Skil | 0.700 | 0.699 | 0.518 |
| Phactr2 | 0.700 | 0.414 | 0.607 |
| Ppp1r15a | 0.700 | 1.284 | 1.105 |
| Klf7 | 0.699 | 0.626 | 0.664 |
| Fosb | 0.699 | 1.641 | 1.668 |
| Sap18 | 0.699 | 0.679 | 0.646 |
| Aprt | 0.698 | 0.704 | 0.806 |
| Sec13 | 0.697 | 0.538 | 0.595 |
| Khdrbs1 | 0.697 | 0.644 | 0.664 |
| Trim35 | 0.697 | 0.544 | 0.646 |
| Zfp704 | 0.696 | 0.516 | 0.549 |
| Coro1c | 0.695 | 0.483 | 0.290 |
| Cdk2ap1 | 0.695 | 0.548 | 0.515 |
| Smim10l1 | 0.694 | 0.484 | 0.537 |
| Psmd11 | 0.694 | 0.667 | 0.597 |
| Psmd6 | 0.693 | 0.708 | 0.609 |
| Slc43a3 | 0.693 | 0.388 | 0.621 |
| Nudc | 0.692 | 0.683 | 0.726 |
| Mast4 | 0.692 | 0.832 | 0.752 |
| Fnbp1l | 0.692 | 0.283 | 0.260 |
| Ech1 | 0.692 | 0.807 | 0.922 |

| | | | |
|---|---|---|---|
| Tpm2 | 0.692 | 2.728 | 1.290 |
| Creg1 | 0.692 | 0.284 | 0.386 |
| Slk | 0.692 | 0.472 | 0.464 |
| Pgrmc1 | 0.692 | 0.816 | 0.630 |
| Lpcat3 | 0.691 | 0.391 | 0.313 |
| Ptges3 | 0.691 | 0.734 | 0.704 |
| Cyc1 | 0.690 | 0.586 | 0.565 |
| Rtf1 | 0.690 | 0.675 | 0.683 |
| Acadl | 0.689 | 0.659 | 0.555 |
| Ext2 | 0.689 | 0.470 | 0.464 |
| Add1 | 0.688 | 0.474 | 0.641 |
| Rnf10 | 0.688 | 0.714 | 0.695 |
| Stmn1 | 0.688 | 0.467 | 0.641 |
| Cd24a | 0.688 | 0.401 | 0.239 |
| Mycbp2 | 0.687 | 0.541 | 0.529 |
| Dnaja2 | 0.686 | 0.630 | 0.592 |
| Vps28 | 0.686 | 0.512 | 0.462 |
| Phc2 | 0.684 | 0.345 | 0.286 |
| Zmynd11 | 0.684 | 0.675 | 0.624 |
| Vps29 | 0.684 | 0.657 | 0.618 |
| Glmp | 0.684 | 0.348 | 0.405 |
| Tgoln1 | 0.684 | 0.350 | 0.322 |
| Foxn3 | 0.684 | 1.086 | 0.843 |
| Pfdn1 | 0.683 | 0.631 | 0.573 |
| Hbb-bs | 0.683 | 0.663 | 0.743 |
| Psmc1 | 0.683 | 0.549 | 0.540 |
| Tcn2 | 0.682 | 0.159 | 0.247 |
| Irf6 | 0.681 | 0.005 | 0.011 |
| Mbd3 | 0.681 | 0.655 | 0.696 |
| Srsf6 | 0.681 | 0.737 | 0.746 |
| Med28 | 0.680 | 0.492 | 0.495 |
| Skap2 | 0.680 | 0.392 | 0.330 |
| Jcad | 0.680 | 0.157 | 0.202 |
| Mt1 | 0.679 | 2.236 | 1.595 |
| Tpp1 | 0.679 | 0.407 | 0.326 |
| Ndufb3 | 0.679 | 0.656 | 0.571 |
| Srp72 | 0.679 | 0.633 | 0.729 |
| Tomm5 | 0.679 | 0.615 | 0.712 |
| Hyal2 | 0.679 | 0.220 | 0.226 |
| Trmt112 | 0.678 | 0.632 | 0.673 |
| Cycs | 0.678 | 0.562 | 0.493 |
| Chd3 | 0.678 | 0.742 | 0.709 |
| Ccnl2 | 0.678 | 0.668 | 0.607 |
| Ndel1 | 0.677 | 0.580 | 0.655 |
| Ankrd17 | 0.677 | 0.554 | 0.613 |
| 1700020I14R | 0.677 | 0.465 | 0.422 |
| Aff4 | 0.676 | 0.795 | 0.653 |
| Ccdc12 | 0.676 | 0.373 | 0.441 |

| | | | |
|---|---|---|---|
| Cfdp1 | 0.676 | 0.765 | 0.828 |
| 1500015O10I | 0.675 | 0.163 | 0.304 |
| Rnf11 | 0.675 | 0.575 | 0.510 |
| Ddx24 | 0.675 | 0.589 | 0.547 |
| Smim14 | 0.675 | 0.860 | 0.937 |
| Zwint | 0.674 | 0.335 | 0.411 |
| Nucks1 | 0.673 | 0.634 | 0.621 |
| Zc3h15 | 0.673 | 0.652 | 0.679 |
| Degs1 | 0.672 | 0.756 | 0.630 |
| Frrs1 | 0.672 | 0.054 | 0.041 |
| Cyp1b1 | 0.672 | 0.573 | 0.524 |
| Polr2j | 0.672 | 0.632 | 0.590 |
| Eif3g | 0.671 | 0.665 | 0.612 |
| Dag1 | 0.671 | 0.804 | 0.616 |
| Gls | 0.671 | 0.485 | 0.517 |
| Eprs | 0.671 | 0.885 | 0.968 |
| Tmco1 | 0.671 | 0.606 | 0.575 |
| Ndufs4 | 0.670 | 0.567 | 0.679 |
| Pigp | 0.670 | 0.513 | 0.631 |
| Snrpc | 0.669 | 0.649 | 0.613 |
| Nme1 | 0.669 | 0.608 | 0.600 |
| Tmem147 | 0.669 | 0.590 | 0.588 |
| Arid5b | 0.669 | 1.095 | 0.859 |
| Gtf2a2 | 0.668 | 0.695 | 0.591 |
| Arhgef12 | 0.668 | 0.607 | 0.522 |
| Arl2bp | 0.667 | 0.475 | 0.396 |
| Pgm5 | 0.667 | 0.185 | 0.104 |
| Tsen34 | 0.667 | 0.671 | 0.678 |
| Hnrnpr | 0.667 | 0.711 | 0.665 |
| Psmc2 | 0.666 | 0.618 | 0.530 |
| Erh | 0.666 | 0.602 | 0.563 |
| Txndc9 | 0.666 | 0.642 | 0.632 |
| Suclg1 | 0.666 | 0.661 | 0.646 |
| Gorasp2 | 0.666 | 0.492 | 0.552 |
| Tpd52l2 | 0.666 | 0.530 | 0.530 |
| Dnajc10 | 0.665 | 0.501 | 0.522 |
| Hnrnph1 | 0.665 | 0.777 | 0.670 |
| Lamtor2 | 0.665 | 0.647 | 0.710 |
| Map7d1 | 0.665 | 0.517 | 0.450 |
| Pnn | 0.664 | 0.675 | 0.617 |
| Arhgap5 | 0.664 | 0.721 | 0.708 |
| Dcun1d5 | 0.664 | 0.624 | 0.619 |
| Cers4 | 0.663 | 0.501 | 0.330 |
| Acin1 | 0.663 | 0.674 | 0.588 |
| Bod1l | 0.663 | 0.591 | 0.578 |
| Papola | 0.662 | 0.553 | 0.575 |
| Rbm5 | 0.661 | 0.603 | 0.572 |
| Prcp | 0.660 | 0.108 | 0.127 |

| | | | |
|---|---|---|---|
| Psmd1 | 0.660 | 0.575 | 0.539 |
| Ccar1 | 0.659 | 0.644 | 0.537 |
| Bmpr2 | 0.659 | 0.491 | 0.421 |
| Nbdy | 0.659 | 0.565 | 0.477 |
| Tgfb1i1 | 0.658 | 1.129 | 0.779 |
| Ccser2 | 0.658 | 0.606 | 0.526 |
| Mettl9 | 0.658 | 0.774 | 0.813 |
| Phf5a | 0.658 | 0.537 | 0.498 |
| Fkbp7 | 0.658 | 0.555 | 0.852 |
| Foxc1 | 0.657 | 0.520 | 0.235 |
| Timm17a | 0.657 | 0.603 | 0.482 |
| Plxnd1 | 0.657 | 0.085 | 0.079 |
| Ppp1r12c | 0.656 | 0.562 | 0.418 |
| Wbp2 | 0.656 | 0.450 | 0.311 |
| Eif3d | 0.656 | 0.605 | 0.602 |
| Polr2a | 0.656 | 0.569 | 0.605 |
| Eif3l | 0.655 | 0.628 | 0.602 |
| Emc3 | 0.655 | 0.471 | 0.448 |
| Arl6ip1 | 0.655 | 0.437 | 0.498 |
| Rcn2 | 0.655 | 0.724 | 0.634 |
| Zfand5 | 0.655 | 0.821 | 0.725 |
| Zfp292 | 0.655 | 0.567 | 0.543 |
| Grasp | 0.654 | 0.143 | 0.283 |
| Csad | 0.654 | 0.628 | 0.555 |
| Scx | 0.654 | 0.245 | 0.038 |
| Pdcd4 | 0.654 | 0.412 | 0.621 |
| Camk2d | 0.654 | 0.934 | 0.587 |
| Pqlc1 | 0.654 | 0.214 | 0.185 |
| Ralb | 0.654 | 0.407 | 0.325 |
| Prkcsh | 0.653 | 0.482 | 0.571 |
| Usp34 | 0.653 | 0.645 | 0.563 |
| Mrpl17 | 0.653 | 0.525 | 0.544 |
| Acp1 | 0.653 | 0.710 | 0.750 |
| 2900097C17F | 0.652 | 0.653 | 0.736 |
| Chmp5 | 0.651 | 0.492 | 0.471 |
| Sumo3 | 0.651 | 0.584 | 0.669 |
| Bcl7c | 0.651 | 0.723 | 0.934 |
| Ndufa10 | 0.651 | 0.592 | 0.599 |
| Mrpl58 | 0.651 | 0.447 | 0.408 |
| Glud1 | 0.651 | 0.552 | 0.502 |
| Elovl1 | 0.651 | 0.447 | 0.434 |
| Pea15a | 0.650 | 0.327 | 0.488 |
| Tbcb | 0.649 | 0.585 | 0.548 |
| Denr | 0.649 | 0.527 | 0.639 |
| Etfb | 0.648 | 0.631 | 0.743 |
| Snw1 | 0.648 | 0.519 | 0.539 |
| Snrpd3 | 0.647 | 0.639 | 0.634 |
| Rfk | 0.647 | 0.146 | 0.143 |

| | | | |
|---|---|---|---|
| Ewsr1 | 0.647 | 0.659 | 0.528 |
| Eif4a3 | 0.647 | 0.534 | 0.502 |
| Gja5 | 0.647 | 0.003 | 0.003 |
| Tcf7l1 | 0.647 | 0.458 | 0.475 |
| Mrpl34 | 0.646 | 0.398 | 0.361 |
| Nras | 0.646 | 0.427 | 0.495 |
| Myh10 | 0.646 | 1.188 | 1.011 |
| Cltb | 0.646 | 0.477 | 0.481 |
| Ufm1 | 0.645 | 0.497 | 0.556 |
| Psmc5 | 0.645 | 0.629 | 0.596 |
| Tnrc6a | 0.645 | 0.782 | 0.643 |
| Scaf11 | 0.645 | 0.749 | 0.717 |
| Cox14 | 0.645 | 0.560 | 0.540 |
| Yipf5 | 0.645 | 0.552 | 0.733 |
| Parva | 0.644 | 0.619 | 0.520 |
| Mrps21 | 0.644 | 0.547 | 0.559 |
| Mageh1 | 0.644 | 0.450 | 0.458 |
| Txn2 | 0.643 | 0.489 | 0.602 |
| Slc44a2 | 0.642 | 0.449 | 0.298 |
| Jpt1 | 0.642 | 0.491 | 0.693 |
| Cmip | 0.642 | 0.211 | 0.283 |
| Snx18 | 0.641 | 0.478 | 0.542 |
| Golga7 | 0.641 | 0.379 | 0.473 |
| Skp1a | 0.641 | 0.565 | 0.514 |
| Cd164 | 0.640 | 0.518 | 0.499 |
| Mgat2 | 0.640 | 0.547 | 0.643 |
| Stat3 | 0.640 | 0.913 | 0.777 |
| Tex261 | 0.640 | 0.390 | 0.560 |
| Pten | 0.640 | 0.528 | 0.539 |
| Fkbp3 | 0.640 | 0.675 | 0.713 |
| Mier1 | 0.639 | 0.578 | 0.572 |
| Atp1b3 | 0.638 | 0.638 | 0.352 |
| Slc29a1 | 0.638 | 0.683 | 0.476 |
| Snrnp27 | 0.638 | 0.574 | 0.535 |
| Ndufa1 | 0.638 | 0.540 | 0.506 |
| Snx17 | 0.638 | 0.521 | 0.516 |
| Tcf12 | 0.636 | 0.550 | 0.542 |
| Ptger4 | 0.635 | 0.354 | 0.249 |
| Lasp1 | 0.635 | 0.301 | 0.441 |
| Uqcc2 | 0.635 | 0.563 | 0.627 |
| Lsr | 0.635 | 0.006 | 0.004 |
| Nsd1 | 0.634 | 0.533 | 0.542 |
| Use1 | 0.634 | 0.650 | 0.611 |
| Rab5c | 0.633 | 0.523 | 0.481 |
| Nfe2l2 | 0.633 | 0.511 | 0.457 |
| Rflnb | 0.633 | 0.117 | 0.234 |
| Arid4b | 0.633 | 0.748 | 0.595 |
| Adgrf5 | 0.633 | 0.023 | 0.012 |

| | | | |
|---|---|---|---|
| Ccdc28b | 0.633 | 0.253 | 0.312 |
| Naa38 | 0.631 | 0.391 | 0.388 |
| Txndc15 | 0.631 | 0.454 | 0.453 |
| Ccdc50 | 0.631 | 0.492 | 0.445 |
| Mef2c | 0.630 | 1.067 | 0.584 |
| Fgd5 | 0.630 | 0.019 | 0.110 |
| Ccni | 0.630 | 0.605 | 0.591 |
| Cuedc2 | 0.630 | 0.592 | 0.532 |
| Ctnnbip1 | 0.630 | 0.326 | 0.392 |
| Mdh1 | 0.629 | 0.768 | 0.651 |
| Pak2 | 0.629 | 0.506 | 0.451 |
| Rab21 | 0.628 | 0.487 | 0.410 |
| Mrpl51 | 0.628 | 0.527 | 0.508 |
| Impad1 | 0.627 | 0.590 | 0.458 |
| Lrp10 | 0.627 | 0.452 | 0.430 |
| Ttc14 | 0.627 | 0.532 | 0.474 |
| Slc39a7 | 0.626 | 0.513 | 0.673 |
| AY036118 | 0.626 | 0.654 | 0.557 |
| Luc7l3 | 0.626 | 0.583 | 0.522 |
| Itgb4 | 0.626 | 0.019 | 0.014 |
| Setd5 | 0.625 | 0.554 | 0.602 |
| Ssrp1 | 0.625 | 0.515 | 0.571 |
| Tax1bp3 | 0.625 | 0.553 | 0.652 |
| Tnrc6c | 0.624 | 0.766 | 0.736 |
| Fam104a | 0.624 | 0.707 | 0.664 |
| P3h4 | 0.623 | 0.540 | 0.717 |
| Pdia5 | 0.623 | 0.149 | 0.213 |
| Adamts10 | 0.623 | 0.345 | 0.486 |
| Cdc42bpa | 0.623 | 0.393 | 0.297 |
| Afdn | 0.623 | 0.204 | 0.294 |
| Mprip | 0.623 | 0.875 | 0.773 |
| Dazap1 | 0.623 | 0.528 | 0.597 |
| Copb1 | 0.623 | 0.644 | 0.669 |
| Tmem208 | 0.622 | 0.505 | 0.526 |
| Zfp644 | 0.622 | 0.695 | 0.614 |
| Fam32a | 0.622 | 0.488 | 0.473 |
| Snrpf | 0.622 | 0.628 | 0.584 |
| U2surp | 0.622 | 0.636 | 0.583 |
| Ntan1 | 0.622 | 0.697 | 0.709 |
| Pcnx | 0.622 | 0.257 | 0.177 |
| Crtap | 0.622 | 0.558 | 0.880 |
| Ssna1 | 0.621 | 0.402 | 0.471 |
| Ginm1 | 0.621 | 0.565 | 0.412 |
| Stx5a | 0.621 | 0.500 | 0.543 |
| Map2k3 | 0.621 | 0.606 | 0.414 |
| Bola2 | 0.621 | 0.412 | 0.447 |
| Vat1 | 0.621 | 0.458 | 0.637 |
| Tram1 | 0.620 | 0.503 | 0.614 |

| | | | |
|---|---|---|---|
| Hcfc1r1 | 0.620 | 1.026 | 0.815 |
| Ghitm | 0.620 | 0.359 | 0.407 |
| Stxbp6 | 0.620 | 0.185 | 0.360 |
| Amotl1 | 0.620 | 0.520 | 0.513 |
| Hdac1 | 0.619 | 0.485 | 0.404 |
| Tgm2 | 0.619 | 0.221 | 0.163 |
| Polr2e | 0.618 | 0.400 | 0.577 |
| Ubl3 | 0.618 | 0.403 | 0.318 |
| Hspd1 | 0.618 | 0.775 | 0.733 |
| Sgms1 | 0.617 | 0.220 | 0.248 |
| Atp6v1a | 0.617 | 0.415 | 0.310 |
| Acbd3 | 0.617 | 0.551 | 0.552 |
| Glo1 | 0.615 | 0.571 | 0.531 |
| Bmyc | 0.615 | 0.497 | 0.511 |
| Psmd3 | 0.615 | 0.537 | 0.500 |
| Cops4 | 0.615 | 0.581 | 0.627 |
| Mkl2 | 0.614 | 0.171 | 0.148 |
| Nav1 | 0.614 | 0.663 | 0.619 |
| P4ha2 | 0.614 | 0.230 | 0.518 |
| Tm9sf2 | 0.614 | 0.498 | 0.442 |
| Thoc2 | 0.613 | 0.592 | 0.579 |
| Crebbp | 0.613 | 0.666 | 0.672 |
| Usf2 | 0.613 | 0.426 | 0.385 |
| Tubb2a | 0.613 | 0.427 | 0.656 |
| Hmg20b | 0.612 | 0.605 | 0.475 |
| Nmt2 | 0.612 | 0.294 | 0.259 |
| Npr1 | 0.612 | 0.153 | 0.138 |
| Blmh | 0.612 | 0.654 | 0.598 |
| Serp1 | 0.612 | 0.568 | 0.681 |
| Cast | 0.612 | 0.544 | 0.624 |
| Gps1 | 0.612 | 0.480 | 0.506 |
| Evi5 | 0.611 | 0.472 | 0.485 |
| Hbp1 | 0.611 | 0.541 | 0.519 |
| Polr2k | 0.611 | 0.503 | 0.447 |
| Zcchc6 | 0.611 | 0.437 | 0.483 |
| Adam15 | 0.611 | 0.219 | 0.307 |
| Tmed4 | 0.610 | 0.412 | 0.394 |
| Dcaf8 | 0.610 | 0.530 | 0.501 |
| Prpf38b | 0.609 | 0.621 | 0.468 |
| Wls | 0.609 | 0.707 | 0.633 |
| Tgfbr2 | 0.609 | 0.281 | 0.456 |
| Uqcc3 | 0.609 | 0.422 | 0.430 |
| Snx6 | 0.609 | 0.482 | 0.613 |
| Derl1 | 0.609 | 0.429 | 0.511 |
| Socs3 | 0.608 | 1.783 | 1.603 |
| Srsf7 | 0.608 | 0.623 | 0.543 |
| Enpp6 | 0.608 | 0.010 | 0.004 |
| Yy1 | 0.608 | 0.523 | 0.502 |

| | | | |
|---|---|---|---|
| 2900026A02F | 0.607 | 0.207 | 0.201 |
| G3bp1 | 0.607 | 0.576 | 0.555 |
| Zfp36 | 0.606 | 1.817 | 1.541 |
| Ei24 | 0.606 | 0.556 | 0.486 |
| Nap1l4 | 0.606 | 0.425 | 0.437 |
| Baz2b | 0.606 | 0.515 | 0.493 |
| Pcmt1 | 0.605 | 0.569 | 0.546 |
| Tead1 | 0.605 | 0.714 | 0.533 |
| Mest | 0.605 | 0.271 | 1.156 |
| Wac | 0.605 | 0.597 | 0.579 |
| Cuta | 0.605 | 0.550 | 0.594 |
| Plxna2 | 0.605 | 0.158 | 0.134 |
| Mob2 | 0.605 | 0.351 | 0.265 |
| Chic2 | 0.605 | 0.550 | 0.648 |
| Limch1 | 0.604 | 0.232 | 0.060 |
| Gtf2i | 0.604 | 0.552 | 0.555 |
| Vapb | 0.604 | 0.434 | 0.559 |
| Rab34 | 0.604 | 0.731 | 0.931 |
| Tle1 | 0.603 | 0.790 | 0.371 |
| Abcf1 | 0.603 | 0.476 | 0.472 |
| Cenpx | 0.603 | 0.364 | 0.410 |
| Pdgfa | 0.602 | 0.701 | 0.272 |
| BC005537 | 0.602 | 0.661 | 0.671 |
| Taok1 | 0.602 | 0.512 | 0.502 |
| Tspan4 | 0.602 | 0.394 | 0.550 |
| Anapc16 | 0.602 | 0.527 | 0.496 |
| Sptlc2 | 0.602 | 0.232 | 0.263 |
| Mpzl1 | 0.601 | 0.460 | 0.586 |
| Plscr3 | 0.601 | 0.368 | 0.434 |
| Tmem160 | 0.601 | 0.460 | 0.391 |
| Btbd7 | 0.601 | 0.499 | 0.413 |
| Pxdn | 0.601 | 0.577 | 0.570 |
| Mpc1 | 0.601 | 0.760 | 0.598 |
| Elmo1 | 0.600 | 0.129 | 0.087 |
| Comt | 0.600 | 0.590 | 0.662 |
| Brd4 | 0.600 | 0.613 | 0.646 |
| Aebp1 | 0.599 | 0.785 | 1.314 |
| Jtb | 0.599 | 0.496 | 0.531 |
| Nudt4 | 0.599 | 1.054 | 0.553 |
| Vps36 | 0.599 | 0.326 | 0.297 |
| Olfml2a | 0.599 | 0.259 | 0.122 |
| Anxa7 | 0.599 | 0.486 | 0.453 |
| Naxe | 0.598 | 0.489 | 0.595 |
| Strap | 0.598 | 0.558 | 0.558 |
| Bcas2 | 0.598 | 0.568 | 0.597 |
| Utrn | 0.597 | 0.521 | 0.339 |
| Mrps16 | 0.597 | 0.538 | 0.597 |
| Magoh | 0.597 | 0.664 | 0.571 |

| | | | |
|---|---|---|---|
| Cnn2 | 0.597 | 1.189 | 0.720 |
| Tsc22d3 | 0.596 | 0.333 | 0.495 |
| Pin1 | 0.596 | 0.501 | 0.435 |
| Hspa4 | 0.596 | 0.614 | 0.490 |
| Spon1 | 0.595 | 0.445 | 0.270 |
| Dnajc5 | 0.595 | 0.437 | 0.436 |
| Ggnbp2 | 0.595 | 0.639 | 0.609 |
| Spag7 | 0.595 | 0.499 | 0.454 |
| Aco2 | 0.594 | 0.489 | 0.398 |
| Ube2k | 0.594 | 0.608 | 0.571 |
| Impdh2 | 0.593 | 0.556 | 0.660 |
| Jak1 | 0.592 | 0.534 | 0.716 |
| Tshz2 | 0.592 | 0.775 | 1.253 |
| Pum1 | 0.592 | 0.615 | 0.586 |
| Ahnak2 | 0.592 | 0.842 | 0.664 |
| Sarnp | 0.592 | 0.668 | 0.607 |
| Ubxn6 | 0.592 | 0.541 | 0.473 |
| Man2b1 | 0.592 | 0.387 | 0.516 |
| Commd1 | 0.592 | 0.564 | 0.582 |
| Dnajb11 | 0.591 | 0.428 | 0.509 |
| Hey2 | 0.591 | 0.149 | 0.049 |
| Stt3a | 0.591 | 0.519 | 0.545 |
| Ctcf | 0.591 | 0.513 | 0.540 |
| Cggbp1 | 0.591 | 0.534 | 0.519 |
| Mvb12a | 0.590 | 0.359 | 0.396 |
| Pkp4 | 0.590 | 0.399 | 0.300 |
| Hnrnpl | 0.590 | 0.676 | 0.563 |
| Wbp1l | 0.590 | 0.444 | 0.371 |
| Pafah1b2 | 0.589 | 0.442 | 0.479 |
| Arpc5l | 0.589 | 0.567 | 0.449 |
| Ing1 | 0.589 | 0.531 | 0.511 |
| Itga9 | 0.589 | 0.765 | 0.275 |
| Yipf4 | 0.589 | 0.746 | 0.593 |
| Rp9 | 0.589 | 0.706 | 0.753 |
| Bub3 | 0.589 | 0.466 | 0.415 |
| Phf20l1 | 0.589 | 0.536 | 0.427 |
| Cyfip1 | 0.588 | 0.464 | 0.569 |
| Pdcd6ip | 0.588 | 0.455 | 0.436 |
| Eif3b | 0.588 | 0.481 | 0.484 |
| Prrc2b | 0.588 | 0.644 | 0.571 |
| Clec2d | 0.587 | 0.250 | 0.342 |
| Rb1cc1 | 0.587 | 0.606 | 0.566 |
| Sh3glb2 | 0.587 | 0.288 | 0.248 |
| Dusp3 | 0.586 | 0.324 | 0.300 |
| Col18a1 | 0.586 | 0.135 | 0.139 |
| Unc50 | 0.586 | 0.498 | 0.485 |
| Art4 | 0.586 | 0.282 | 0.280 |
| Wwp1 | 0.586 | 0.305 | 0.254 |

| | | | |
|---|---|---|---|
| Mob4 | 0.586 | 0.706 | 0.552 |
| Chrac1 | 0.585 | 0.580 | 0.518 |
| Tm6sf1 | 0.585 | 0.067 | 0.018 |
| Tbl1x | 0.584 | 0.694 | 0.686 |
| Rap1a | 0.583 | 0.694 | 0.741 |
| Wdr45b | 0.583 | 0.395 | 0.430 |
| Chchd1 | 0.583 | 0.501 | 0.558 |
| Cript | 0.583 | 0.492 | 0.401 |
| Sec22b | 0.583 | 0.445 | 0.612 |
| Chmp2b | 0.582 | 0.623 | 0.513 |
| Ttyh3 | 0.582 | 0.196 | 0.333 |
| Sap30l | 0.582 | 0.472 | 0.573 |
| Mtch2 | 0.582 | 0.557 | 0.515 |
| Sash1 | 0.582 | 0.303 | 0.513 |
| Rcan1 | 0.582 | 0.340 | 0.226 |
| Sppl3 | 0.581 | 0.486 | 0.499 |
| Coa3 | 0.581 | 0.530 | 0.587 |
| Smpd3 | 0.581 | 0.025 | 0.126 |
| Ptpn14 | 0.581 | 0.320 | 0.247 |
| Lamb1 | 0.581 | 0.178 | 0.752 |
| Psmb7 | 0.580 | 0.566 | 0.503 |
| Mrpl23 | 0.580 | 0.634 | 0.555 |
| Copz1 | 0.580 | 0.406 | 0.505 |
| Pfdn2 | 0.580 | 0.614 | 0.537 |
| Nmt1 | 0.579 | 0.443 | 0.395 |
| Smc6 | 0.579 | 0.675 | 0.724 |
| Emc6 | 0.579 | 0.456 | 0.404 |
| Os9 | 0.578 | 0.383 | 0.501 |
| Psmd12 | 0.578 | 0.616 | 0.514 |
| Xrn2 | 0.578 | 0.564 | 0.488 |
| Bmpr1a | 0.578 | 0.687 | 0.530 |
| Top2b | 0.578 | 0.574 | 0.509 |
| Ndn | 0.577 | 0.538 | 0.942 |
| Akt1s1 | 0.577 | 0.484 | 0.556 |
| Col14a1 | 0.577 | 0.796 | 1.952 |
| Gap43 | 0.577 | 0.039 | 0.155 |
| Tor1aip1 | 0.577 | 0.482 | 0.493 |
| Cdca7l | 0.577 | 0.010 | 0.012 |
| Hnrnpul2 | 0.576 | 0.480 | 0.525 |
| Cmpk1 | 0.575 | 0.484 | 0.487 |
| Kank1 | 0.575 | 0.760 | 0.434 |
| Arhgef1 | 0.575 | 0.319 | 0.311 |
| Pi16 | 0.575 | 0.051 | 0.652 |
| Mrpl18 | 0.575 | 0.494 | 0.457 |
| Fkbp4 | 0.574 | 0.605 | 0.607 |
| Pum2 | 0.573 | 0.516 | 0.461 |
| Sdha | 0.573 | 0.503 | 0.474 |
| Arhgef2 | 0.573 | 0.472 | 0.390 |

| | | | |
|---|---|---|---|
| Tfg | 0.573 | 0.608 | 0.605 |
| Prr13 | 0.572 | 0.260 | 0.349 |
| Cnih4 | 0.572 | 0.571 | 0.446 |
| Jmjd1c | 0.572 | 0.635 | 0.678 |
| Smc3 | 0.572 | 0.658 | 0.679 |
| Dennd5a | 0.571 | 0.367 | 0.398 |
| Rassf9 | 0.571 | 0.007 | 0.037 |
| Lyl1 | 0.571 | 0.005 | 0.006 |
| Cttnbp2nl | 0.571 | 0.156 | 0.157 |
| Pja2 | 0.570 | 0.675 | 0.564 |
| Sntb2 | 0.570 | 0.799 | 0.969 |
| Adrm1 | 0.570 | 0.513 | 0.443 |
| Hadha | 0.570 | 0.570 | 0.511 |
| Ppp2r5c | 0.569 | 0.497 | 0.419 |
| Ddx17 | 0.569 | 0.531 | 0.562 |
| Cops5 | 0.569 | 0.514 | 0.520 |
| Setd3 | 0.569 | 0.403 | 0.366 |
| Prpf4b | 0.568 | 0.560 | 0.538 |
| Ythdc1 | 0.568 | 0.536 | 0.510 |
| Nsmce4a | 0.568 | 0.733 | 0.544 |
| Qsox1 | 0.568 | 0.732 | 0.529 |
| Ctdnep1 | 0.568 | 0.448 | 0.498 |
| Smarca5 | 0.568 | 0.758 | 0.648 |
| Cr1l | 0.567 | 0.393 | 0.389 |
| Phf3 | 0.567 | 0.570 | 0.601 |
| Add3 | 0.567 | 0.306 | 0.782 |
| Ap2a2 | 0.567 | 0.435 | 0.715 |
| Polr2g | 0.567 | 0.533 | 0.466 |
| Fxr1 | 0.567 | 0.534 | 0.472 |
| Polr2c | 0.567 | 0.552 | 0.460 |
| Cul3 | 0.566 | 0.577 | 0.526 |
| 2-Mar | 0.566 | 0.683 | 0.679 |
| Ackr3 | 0.566 | 0.233 | 0.533 |
| Cdc26 | 0.566 | 0.541 | 0.627 |
| Chmp3 | 0.566 | 0.418 | 0.462 |
| Bad | 0.566 | 0.377 | 0.480 |
| Fam92a | 0.566 | 0.618 | 0.647 |
| Zfp207 | 0.565 | 0.455 | 0.428 |
| Myo1c | 0.565 | 0.501 | 0.327 |
| Tspan2 | 0.565 | 0.406 | 0.075 |
| Ubald2 | 0.564 | 0.582 | 0.429 |
| Uba1 | 0.564 | 0.463 | 0.483 |
| Rbbp7 | 0.564 | 0.787 | 0.785 |
| 5-Mar | 0.564 | 0.588 | 0.487 |
| Znhit1 | 0.564 | 0.358 | 0.404 |
| Birc6 | 0.563 | 0.609 | 0.538 |
| Vamp5 | 0.563 | 0.212 | 0.189 |
| Psme2 | 0.563 | 0.500 | 0.523 |

| | | | |
|---|---|---|---|
| Cers2 | 0.563 | 0.492 | 0.410 |
| Epn1 | 0.563 | 0.354 | 0.418 |
| Polr2i | 0.563 | 0.474 | 0.474 |
| Ppm1a | 0.563 | 0.690 | 0.734 |
| Uhrf2 | 0.563 | 0.394 | 0.473 |
| Lsm7 | 0.562 | 0.561 | 0.660 |
| Zfp148 | 0.562 | 0.513 | 0.569 |
| Pnkd | 0.561 | 0.446 | 0.366 |
| Tsg101 | 0.561 | 0.334 | 0.385 |
| Tmbim4 | 0.561 | 0.413 | 0.403 |
| Bud31 | 0.560 | 0.432 | 0.454 |
| Olfml2b | 0.560 | 1.268 | 0.892 |
| Armcx4 | 0.560 | 0.156 | 0.177 |
| Ap1s1 | 0.559 | 0.361 | 0.418 |
| Ubtf | 0.559 | 0.567 | 0.618 |
| Pdlim5 | 0.559 | 0.391 | 0.151 |
| Acaa1a | 0.559 | 0.477 | 0.550 |
| Pkn2 | 0.559 | 0.527 | 0.538 |
| Rasa1 | 0.559 | 0.359 | 0.358 |
| Jkamp | 0.559 | 0.539 | 0.462 |
| Erlec1 | 0.558 | 0.527 | 0.677 |
| Fam46a | 0.558 | 0.141 | 0.756 |
| Rab5a | 0.558 | 0.472 | 0.476 |
| Caprin1 | 0.557 | 0.456 | 0.462 |
| Lysmd2 | 0.557 | 0.261 | 0.454 |
| Pcnp | 0.557 | 0.550 | 0.539 |
| Sdf2 | 0.556 | 0.594 | 0.609 |
| Ubn2 | 0.556 | 0.539 | 0.472 |
| Dkk2 | 0.556 | 0.042 | 0.177 |
| Sltm | 0.555 | 0.547 | 0.443 |
| Ccdc59 | 0.555 | 0.475 | 0.450 |
| Bbip1 | 0.555 | 0.368 | 0.365 |
| Brd3 | 0.554 | 0.519 | 0.596 |
| Wdr1 | 0.554 | 0.574 | 0.417 |
| Sorbs1 | 0.554 | 0.473 | 0.169 |
| Myof | 0.554 | 0.408 | 0.380 |
| Mknk2 | 0.554 | 0.339 | 0.302 |
| Copg1 | 0.554 | 0.450 | 0.554 |
| Afap1l1 | 0.554 | 0.005 | 0.006 |
| Dlst | 0.553 | 0.398 | 0.406 |
| C1qbp | 0.553 | 0.510 | 0.516 |
| Rtcb | 0.553 | 0.421 | 0.453 |
| Golga4 | 0.553 | 0.457 | 0.417 |
| Tmem9 | 0.552 | 0.377 | 0.474 |
| Gabarapl1 | 0.552 | 0.363 | 0.392 |
| Cyb561 | 0.552 | 0.371 | 0.151 |
| Msl1 | 0.551 | 0.475 | 0.448 |
| Hhex | 0.551 | 0.005 | 0.004 |

| | | | |
|---|---|---|---|
| 2-Mar | 0.550 | 0.392 | 0.353 |
| Clcn3 | 0.550 | 0.146 | 0.154 |
| Rbbp4 | 0.550 | 0.589 | 0.554 |
| Plin3 | 0.549 | 0.539 | 0.669 |
| Zfpm1 | 0.549 | 0.177 | 0.170 |
| Pa2g4 | 0.549 | 0.593 | 0.536 |
| 7-Mar | 0.549 | 0.562 | 0.514 |
| Dctn4 | 0.549 | 0.593 | 0.484 |
| Gadd45b | 0.549 | 1.547 | 1.343 |
| Lrrc8a | 0.548 | 0.349 | 0.287 |
| Plp2 | 0.548 | 0.592 | 0.562 |
| Phf14 | 0.548 | 0.447 | 0.416 |
| Gars | 0.548 | 0.359 | 0.484 |
| Eif3j1 | 0.547 | 0.486 | 0.465 |
| Man1a2 | 0.547 | 0.676 | 0.580 |
| Gemin7 | 0.545 | 0.470 | 0.436 |
| Ogdh | 0.545 | 0.445 | 0.350 |
| Il6st | 0.545 | 0.529 | 0.705 |
| Echs1 | 0.545 | 0.633 | 0.587 |
| Fam114a2 | 0.545 | 0.409 | 0.357 |
| Arcn1 | 0.545 | 0.553 | 0.628 |
| Sgta | 0.544 | 0.472 | 0.538 |
| Emg1 | 0.544 | 0.494 | 0.466 |
| Spop | 0.544 | 0.579 | 0.512 |
| Gm16286 | 0.544 | 0.549 | 0.521 |
| Scoc | 0.543 | 0.426 | 0.354 |
| Traf7 | 0.543 | 0.350 | 0.410 |
| Yif1a | 0.543 | 0.426 | 0.555 |
| Ptpra | 0.543 | 0.576 | 0.624 |
| Smtnl2 | 0.542 | 0.057 | 0.221 |
| Mall | 0.542 | 0.006 | 0.005 |
| Mindy2 | 0.542 | 0.217 | 0.165 |
| Hsd17b10 | 0.542 | 0.780 | 0.684 |
| Sars | 0.541 | 0.526 | 0.522 |
| Pkig | 0.541 | 0.381 | 0.399 |
| Fyn | 0.541 | 0.221 | 0.528 |
| Lrrc8c | 0.541 | 0.123 | 0.122 |
| Wsb1 | 0.541 | 0.661 | 0.570 |
| Sqstm1 | 0.541 | 0.682 | 0.621 |
| Mrpl12 | 0.541 | 0.436 | 0.435 |
| Dlgap4 | 0.540 | 0.629 | 0.582 |
| Rbm42 | 0.539 | 0.449 | 0.429 |
| Csnk1g2 | 0.539 | 0.468 | 0.548 |
| Lhfp | 0.539 | 1.591 | 1.306 |
| Cdo1 | 0.539 | 0.534 | 0.616 |
| Lgals9 | 0.539 | 0.027 | 0.223 |
| Serpinf1 | 0.538 | 0.315 | 1.318 |
| Ddrgk1 | 0.538 | 0.653 | 0.642 |

| | | | |
|---|---|---|---|
| Glrx2 | 0.537 | 0.514 | 0.411 |
| Plcb4 | 0.536 | 0.197 | 0.149 |
| Ddx1 | 0.536 | 0.462 | 0.512 |
| Ppp2r2a | 0.536 | 0.467 | 0.490 |
| Hoxa5 | 0.536 | 0.362 | 0.228 |
| Mfap4 | 0.536 | 2.163 | 2.980 |
| Erp44 | 0.536 | 0.493 | 0.578 |
| Ndufv1 | 0.535 | 0.543 | 0.517 |
| Cop1 | 0.535 | 0.432 | 0.497 |
| Clstn1 | 0.535 | 0.460 | 0.453 |
| Xpnpep1 | 0.535 | 0.358 | 0.447 |
| Mrpl14 | 0.534 | 0.456 | 0.482 |
| Wbp4 | 0.534 | 0.475 | 0.451 |
| Dcn | 0.534 | 0.749 | 3.758 |
| Elf1 | 0.534 | 0.277 | 0.248 |
| Zc3h13 | 0.534 | 0.523 | 0.463 |
| Uaca | 0.533 | 0.323 | 0.291 |
| Ndufs8 | 0.533 | 0.420 | 0.402 |
| Smad6 | 0.533 | 0.230 | 0.117 |
| Tnrc6b | 0.533 | 0.650 | 0.600 |
| Nhsl2 | 0.533 | 0.146 | 0.156 |
| Snf8 | 0.532 | 0.422 | 0.402 |
| Tcaf1 | 0.532 | 0.514 | 0.490 |
| Ankrd46 | 0.532 | 0.180 | 0.217 |
| Mrpl43 | 0.531 | 0.451 | 0.395 |
| Zfand6 | 0.531 | 0.447 | 0.414 |
| Svbp | 0.531 | 0.574 | 0.648 |
| Kcmf1 | 0.531 | 0.559 | 0.520 |
| Smad4 | 0.530 | 0.526 | 0.555 |
| Uba2 | 0.530 | 0.687 | 0.479 |
| Aqp1 | 0.530 | 0.434 | 0.205 |
| Tead2 | 0.530 | 0.298 | 0.362 |
| Hadh | 0.530 | 0.564 | 0.501 |
| Syncrip | 0.530 | 0.599 | 0.609 |
| Magt1 | 0.530 | 0.394 | 0.433 |
| Ube2q1 | 0.529 | 0.356 | 0.363 |
| Sae1 | 0.529 | 0.340 | 0.387 |
| Clptm1l | 0.529 | 0.441 | 0.423 |
| Lars2 | 0.528 | 0.337 | 0.356 |
| Trpc4ap | 0.528 | 0.350 | 0.415 |
| Eci2 | 0.528 | 0.602 | 0.649 |
| Trim28 | 0.528 | 0.529 | 0.552 |
| Larp4b | 0.527 | 0.429 | 0.369 |
| Acvrl1 | 0.527 | 0.251 | 0.350 |
| Cxcl1 | 0.527 | 0.628 | 0.751 |
| Atp6v0d1 | 0.526 | 0.495 | 0.571 |
| Mkln1 | 0.526 | 0.597 | 0.503 |
| Grpel1 | 0.526 | 0.473 | 0.442 |

| | | | |
|---|---|---|---|
| Herc2 | 0.526 | 0.314 | 0.288 |
| Ptk2 | 0.526 | 0.252 | 0.284 |
| Thsd7a | 0.526 | 0.042 | 0.046 |
| Pold4 | 0.525 | 0.456 | 0.465 |
| Hspb1 | 0.525 | 1.666 | 0.580 |
| Itpripl2 | 0.524 | 0.347 | 0.438 |
| Lgmn | 0.524 | 0.444 | 0.361 |
| Trp53 | 0.524 | 0.415 | 0.352 |
| Clptm1 | 0.524 | 0.327 | 0.298 |
| Mesd | 0.524 | 0.399 | 0.459 |
| Hspa9 | 0.524 | 0.556 | 0.484 |
| Ociad1 | 0.524 | 0.534 | 0.467 |
| Gng12 | 0.523 | 0.615 | 0.540 |
| Tnks2 | 0.523 | 0.410 | 0.446 |
| Hmgb2 | 0.523 | 0.532 | 0.480 |
| Tmbim1 | 0.523 | 0.438 | 0.295 |
| Dnajc1 | 0.523 | 0.626 | 0.673 |
| Syngr2 | 0.522 | 0.403 | 0.250 |
| Vcl | 0.522 | 1.086 | 0.444 |
| Fam103a1 | 0.522 | 0.511 | 0.419 |
| Zmiz1 | 0.522 | 0.538 | 0.634 |
| Ndfip2 | 0.521 | 0.379 | 0.283 |
| Zfp36l2 | 0.521 | 0.795 | 0.723 |
| Rnf125 | 0.520 | 0.039 | 0.040 |
| Fnta | 0.520 | 0.492 | 0.504 |
| Antxr1 | 0.519 | 1.019 | 1.045 |
| Mrpl42 | 0.518 | 0.558 | 0.542 |
| Snrpb2 | 0.517 | 0.603 | 0.488 |
| Adss | 0.517 | 0.413 | 0.327 |
| Usp9x | 0.517 | 0.645 | 0.591 |
| Cnot6 | 0.517 | 0.351 | 0.426 |
| Pfkl | 0.517 | 0.305 | 0.193 |
| Dock6 | 0.516 | 0.185 | 0.158 |
| Zdhhc20 | 0.516 | 0.556 | 0.513 |
| Rab1b | 0.516 | 0.346 | 0.350 |
| Lsm6 | 0.516 | 0.493 | 0.503 |
| Pxn | 0.515 | 0.232 | 0.324 |
| Hes1 | 0.515 | 0.917 | 0.661 |
| Cttn | 0.515 | 0.471 | 0.485 |
| Ik | 0.515 | 0.491 | 0.437 |
| Ilf2 | 0.515 | 0.549 | 0.508 |
| Klhl6 | 0.515 | 0.006 | 0.011 |
| Ccdc88a | 0.515 | 0.644 | 0.586 |
| Sar1b | 0.514 | 0.466 | 0.386 |
| Dpysl2 | 0.514 | 0.445 | 0.647 |
| Nts | 0.514 | 0.010 | 0.013 |
| Cyhr1 | 0.513 | 0.587 | 0.509 |
| Gatad1 | 0.513 | 0.443 | 0.421 |

| | | | |
|---|---|---|---|
| Glrx5 | 0.513 | 0.460 | 0.434 |
| Mbtps1 | 0.513 | 0.534 | 0.601 |
| Pdcl3 | 0.512 | 0.663 | 0.449 |
| Hip1 | 0.512 | 0.523 | 0.252 |
| Aph1a | 0.512 | 0.416 | 0.385 |
| Sav1 | 0.512 | 0.240 | 0.270 |
| Dab2ip | 0.511 | 0.427 | 0.275 |
| Perp | 0.511 | 0.378 | 0.061 |
| Lsm4 | 0.511 | 0.532 | 0.563 |
| Fryl | 0.511 | 0.462 | 0.147 |
| Ppp1cb | 0.511 | 0.908 | 0.583 |
| Cacybp | 0.511 | 0.496 | 0.463 |
| Krit1 | 0.511 | 0.459 | 0.402 |
| Wsb2 | 0.510 | 0.447 | 0.396 |
| Chmp1a | 0.510 | 0.389 | 0.420 |
| Desi2 | 0.510 | 0.582 | 0.599 |
| R3hdm1 | 0.510 | 0.450 | 0.398 |
| Lama4 | 0.510 | 0.242 | 0.455 |
| Mrpl24 | 0.510 | 0.485 | 0.452 |
| Atp2a2 | 0.510 | 1.052 | 0.666 |
| Lage3 | 0.509 | 0.420 | 0.415 |
| Gusb | 0.509 | 0.342 | 0.458 |
| Egln2 | 0.509 | 0.453 | 0.437 |
| Cxcl16 | 0.509 | 0.045 | 0.097 |
| Trim24 | 0.509 | 0.291 | 0.285 |
| Ap2b1 | 0.508 | 0.265 | 0.367 |
| Srsf10 | 0.508 | 0.567 | 0.480 |
| Vasp | 0.507 | 0.614 | 0.300 |
| Rit1 | 0.507 | 0.389 | 0.339 |
| Bet1l | 0.507 | 0.388 | 0.529 |
| Fbxw2 | 0.506 | 0.471 | 0.446 |
| Ccdc47 | 0.506 | 0.437 | 0.352 |
| Agpat3 | 0.506 | 0.476 | 0.455 |
| P3h1 | 0.506 | 0.462 | 0.667 |
| Mrpl48 | 0.506 | 0.541 | 0.526 |
| Smarca4 | 0.505 | 0.364 | 0.335 |
| Rraga | 0.505 | 0.526 | 0.498 |
| Smim11 | 0.505 | 0.494 | 0.494 |
| Ptprj | 0.505 | 0.074 | 0.043 |
| Btbd1 | 0.504 | 0.352 | 0.444 |
| Oaz2 | 0.504 | 0.622 | 0.542 |
| Rnaseh2c | 0.504 | 0.382 | 0.418 |
| Lsm2 | 0.504 | 0.304 | 0.367 |
| Xist | 0.504 | 0.870 | 0.640 |
| Gatad2b | 0.504 | 0.580 | 0.509 |
| Irf2 | 0.503 | 0.330 | 0.307 |
| Klhl24 | 0.503 | 0.516 | 0.448 |
| Mrpl57 | 0.503 | 0.462 | 0.395 |

| | | | |
|---|---|---|---|
| Cbx5 | 0.503 | 0.459 | 0.528 |
| Hexb | 0.502 | 0.376 | 0.588 |
| Phax | 0.502 | 0.482 | 0.447 |
| Selenon | 0.502 | 0.258 | 0.486 |
| Cdipt | 0.502 | 0.278 | 0.250 |
| Emc4 | 0.502 | 0.403 | 0.304 |
| Hist1h1c | 0.502 | 0.414 | 0.197 |
| Mrps18a | 0.502 | 0.396 | 0.442 |
| Fam43a | 0.501 | 0.066 | 0.142 |
| Phldb2 | 0.501 | 0.299 | 0.327 |
| Mrpl13 | 0.501 | 0.454 | 0.469 |
| Timm17b | 0.501 | 0.384 | 0.454 |
| Dbnl | 0.501 | 0.250 | 0.263 |
| Rbfox2 | 0.501 | 0.774 | 0.649 |
| Trip12 | 0.501 | 0.420 | 0.460 |
| Uhrf1bp1l | 0.501 | 0.210 | 0.157 |
| Rpn1 | 0.500 | 0.455 | 0.586 |
| Sec14l1 | 0.500 | 0.295 | 0.407 |
| 4921524J17R | 0.500 | 0.286 | 0.223 |
| Celf1 | 0.500 | 0.491 | 0.381 |
| Elof1 | 0.500 | 0.481 | 0.519 |
| Mrps18c | 0.499 | 0.430 | 0.352 |
| Osbpl9 | 0.499 | 0.361 | 0.292 |
| Bola3 | 0.499 | 0.354 | 0.330 |
| Gstm1 | 0.499 | 0.557 | 0.679 |
| Siva1 | 0.499 | 0.724 | 0.778 |
| Pink1 | 0.498 | 0.413 | 0.477 |
| Lztfl1 | 0.498 | 0.325 | 0.355 |
| Furin | 0.498 | 0.437 | 0.319 |
| Gapvd1 | 0.498 | 0.446 | 0.426 |
| Itpr1 | 0.497 | 0.290 | 0.217 |
| Ssbp3 | 0.497 | 0.534 | 0.513 |
| Samm50 | 0.497 | 0.436 | 0.448 |
| Zcchc11 | 0.497 | 0.526 | 0.528 |
| Postn | 0.497 | 0.996 | 2.970 |
| Fgfr1op2 | 0.497 | 0.441 | 0.466 |
| Hif1a | 0.496 | 0.475 | 0.391 |
| Arl6ip4 | 0.496 | 0.412 | 0.373 |
| Stx7 | 0.495 | 0.338 | 0.275 |
| Galnt1 | 0.495 | 0.391 | 0.393 |
| Trappc2l | 0.495 | 0.438 | 0.498 |
| Ptprs | 0.494 | 0.655 | 0.819 |
| Acbd6 | 0.494 | 0.514 | 0.425 |
| Arhgap29 | 0.494 | 0.167 | 0.283 |
| Ankrd12 | 0.494 | 0.753 | 0.593 |
| Aph1b | 0.494 | 0.092 | 0.101 |
| Stx12 | 0.493 | 0.374 | 0.400 |
| Sbno1 | 0.493 | 0.574 | 0.464 |

| | | | |
|---|---|---|---|
| Fli1 | 0.493 | 0.062 | 0.049 |
| Irf2bpl | 0.493 | 0.818 | 0.634 |
| Trappc1 | 0.493 | 0.333 | 0.449 |
| Clip1 | 0.493 | 0.498 | 0.430 |
| Asah1 | 0.493 | 0.548 | 0.520 |
| Cstb | 0.493 | 0.740 | 0.621 |
| Acvr1 | 0.492 | 0.269 | 0.220 |
| Tmed5 | 0.492 | 0.452 | 0.395 |
| Inpp1 | 0.492 | 0.195 | 0.165 |
| Git2 | 0.491 | 0.264 | 0.307 |
| Txndc12 | 0.491 | 0.351 | 0.414 |
| Rest | 0.491 | 0.548 | 0.496 |
| Adcy4 | 0.491 | 0.072 | 0.070 |
| Anapc13 | 0.491 | 0.502 | 0.556 |
| Tmcc3 | 0.491 | 0.112 | 0.258 |
| Kif1b | 0.491 | 0.503 | 0.515 |
| Sel1l | 0.490 | 0.391 | 0.446 |
| Cdk11b | 0.490 | 0.445 | 0.444 |
| Tmf1 | 0.490 | 0.375 | 0.361 |
| Lipa | 0.490 | 0.193 | 0.340 |
| Ube2g2 | 0.490 | 0.186 | 0.272 |
| Uba5 | 0.490 | 0.378 | 0.465 |
| Mrps7 | 0.490 | 0.411 | 0.415 |
| Gkap1 | 0.490 | 0.549 | 0.460 |
| Sirt2 | 0.489 | 0.563 | 0.525 |
| Grina | 0.489 | 0.392 | 0.433 |
| Naxd | 0.489 | 0.426 | 0.486 |
| Pcm1 | 0.488 | 0.532 | 0.484 |
| 1110065P20R | 0.488 | 0.542 | 0.487 |
| Mapk1 | 0.488 | 0.408 | 0.424 |
| Srek1 | 0.488 | 0.550 | 0.515 |
| Fam135a | 0.487 | 0.126 | 0.128 |
| N4bp2l2 | 0.487 | 0.470 | 0.421 |
| Utp3 | 0.487 | 0.445 | 0.391 |
| Pak1ip1 | 0.486 | 0.379 | 0.305 |
| Fam114a1 | 0.486 | 0.407 | 0.643 |
| Cul1 | 0.486 | 0.525 | 0.504 |
| Dhx36 | 0.486 | 0.480 | 0.435 |
| Gde1 | 0.486 | 0.479 | 0.420 |
| Yaf2 | 0.486 | 0.397 | 0.324 |
| Tef | 0.486 | 0.282 | 0.230 |
| Zcchc17 | 0.486 | 0.363 | 0.328 |
| Cir1 | 0.486 | 0.443 | 0.433 |
| Cdc42ep5 | 0.485 | 0.426 | 0.559 |
| Ptp4a3 | 0.485 | 0.755 | 0.374 |
| Pbxip1 | 0.485 | 0.871 | 0.494 |
| Rlim | 0.485 | 0.488 | 0.574 |
| Gipc1 | 0.485 | 0.330 | 0.459 |

| | | | |
|---|---|---|---|
| Zbtb7a | 0.484 | 0.487 | 0.482 |
| Zbtb4 | 0.484 | 0.396 | 0.338 |
| Rsl1d1 | 0.484 | 0.556 | 0.507 |
| Cops8 | 0.484 | 0.434 | 0.422 |
| Swap70 | 0.484 | 0.176 | 0.153 |
| Mrps12 | 0.484 | 0.428 | 0.388 |
| Ilkap | 0.484 | 0.471 | 0.487 |
| 1110038F14F | 0.484 | 0.401 | 0.410 |
| Ncstn | 0.484 | 0.217 | 0.220 |
| Gpc6 | 0.483 | 1.171 | 0.924 |
| Huwe1 | 0.483 | 0.477 | 0.413 |
| Idh3g | 0.483 | 0.450 | 0.408 |
| Dnajc7 | 0.483 | 0.449 | 0.439 |
| Pdhb | 0.483 | 0.404 | 0.347 |
| Ube2a | 0.483 | 0.420 | 0.301 |
| Efhd2 | 0.483 | 0.370 | 0.219 |
| Nt5dc2 | 0.483 | 0.388 | 0.749 |
| Hpcal1 | 0.482 | 0.259 | 0.286 |
| Arl3 | 0.482 | 0.531 | 0.563 |
| Pnpla2 | 0.481 | 0.306 | 0.151 |
| Kdm5b | 0.481 | 0.650 | 0.676 |
| Ncor2 | 0.481 | 0.471 | 0.464 |
| Hypk | 0.481 | 0.379 | 0.486 |
| Tmem214 | 0.480 | 0.277 | 0.393 |
| Sfxn3 | 0.480 | 0.334 | 0.260 |
| Kdm2a | 0.480 | 0.464 | 0.437 |
| Wbp11 | 0.480 | 0.447 | 0.462 |
| Tm2d1 | 0.480 | 0.477 | 0.470 |
| Mga | 0.480 | 0.480 | 0.503 |
| Ap3s1 | 0.479 | 0.547 | 0.714 |
| Ubr5 | 0.479 | 0.474 | 0.374 |
| Epb41l2 | 0.479 | 0.342 | 0.676 |
| Fndc1 | 0.479 | 0.279 | 1.291 |
| Dicer1 | 0.479 | 0.341 | 0.303 |
| Gnai3 | 0.479 | 0.390 | 0.448 |
| Prpf19 | 0.478 | 0.445 | 0.471 |
| Hist1h2bc | 0.478 | 0.442 | 0.246 |
| Ppp4r3b | 0.478 | 0.398 | 0.348 |
| Myadm | 0.478 | 0.796 | 0.699 |
| Igf1r | 0.478 | 0.922 | 0.709 |
| Sf3b3 | 0.478 | 0.420 | 0.424 |
| Itfg1 | 0.478 | 0.364 | 0.341 |
| Cldnd1 | 0.477 | 0.680 | 0.579 |
| Selenot | 0.477 | 0.301 | 0.235 |
| Kras | 0.477 | 0.365 | 0.482 |
| Zrsr2 | 0.477 | 0.403 | 0.400 |
| Pdia4 | 0.477 | 0.354 | 0.366 |
| Cflar | 0.477 | 0.295 | 0.303 |

| | | | |
|---|---|---|---|
| Maea | 0.477 | 0.336 | 0.441 |
| Oxa1l | 0.476 | 0.321 | 0.305 |
| Dnmt3a | 0.476 | 0.522 | 0.527 |
| Gaa | 0.476 | 0.262 | 0.547 |
| Dpm2 | 0.476 | 0.354 | 0.408 |
| Immt | 0.476 | 0.468 | 0.407 |
| Shc1 | 0.476 | 0.460 | 0.425 |
| Mrpl28 | 0.476 | 0.399 | 0.433 |
| Prrc2a | 0.475 | 0.516 | 0.516 |
| Nudt21 | 0.475 | 0.491 | 0.471 |
| Commd7 | 0.475 | 0.326 | 0.353 |
| Rgs5 | 0.475 | 2.597 | 0.912 |
| Gadd45g | 0.475 | 1.081 | 0.642 |
| Dhx15 | 0.474 | 0.467 | 0.414 |
| Dennd4a | 0.474 | 0.194 | 0.237 |
| Sec63 | 0.474 | 0.450 | 0.471 |
| Pxdc1 | 0.474 | 0.231 | 0.330 |
| Gspt1 | 0.474 | 0.540 | 0.521 |
| Cdc16 | 0.474 | 0.394 | 0.560 |
| Ptp4a1 | 0.474 | 0.709 | 0.695 |
| Blvrb | 0.474 | 0.497 | 0.585 |
| Idh3b | 0.473 | 0.422 | 0.460 |
| Nudcd3 | 0.473 | 0.429 | 0.410 |
| Tspan31 | 0.473 | 0.433 | 0.496 |
| Eif4ebp1 | 0.472 | 0.656 | 0.757 |
| Krcc1 | 0.472 | 0.370 | 0.446 |
| Map4 | 0.472 | 0.482 | 0.490 |
| Pigyl | 0.471 | 0.458 | 0.405 |
| Moxd1 | 0.471 | 0.003 | 0.012 |
| Col23a1 | 0.471 | 0.039 | 0.080 |
| Dnpep | 0.470 | 0.368 | 0.474 |
| Cerk | 0.470 | 0.221 | 0.181 |
| Ikbip | 0.470 | 0.623 | 0.727 |
| Rassf1 | 0.470 | 0.727 | 0.687 |
| Mcrs1 | 0.470 | 0.440 | 0.377 |
| Rela | 0.470 | 0.349 | 0.361 |
| Tmx4 | 0.470 | 0.477 | 0.359 |
| Foxc2 | 0.470 | 0.359 | 0.210 |
| Csnk1d | 0.470 | 0.420 | 0.421 |
| Mark3 | 0.470 | 0.456 | 0.411 |
| Zmat2 | 0.469 | 0.415 | 0.382 |
| Brd1 | 0.469 | 0.499 | 0.448 |
| Appbp2 | 0.469 | 0.416 | 0.472 |
| Zranb2 | 0.469 | 0.446 | 0.441 |
| Gpr180 | 0.469 | 0.332 | 0.449 |
| Efna1 | 0.469 | 0.131 | 0.087 |
| Tcea1 | 0.469 | 0.458 | 0.444 |
| Usp47 | 0.469 | 0.457 | 0.388 |

| | | | |
|---|---|---|---|
| Hmgb3 | 0.469 | 0.316 | 0.208 |
| Srm | 0.468 | 0.361 | 0.600 |
| Snrnp48 | 0.468 | 0.372 | 0.290 |
| Nars | 0.468 | 0.466 | 0.484 |
| Fam129b | 0.468 | 0.366 | 0.229 |
| Maf1 | 0.468 | 0.409 | 0.383 |
| Nsfl1c | 0.468 | 0.402 | 0.402 |
| Arid1a | 0.468 | 0.516 | 0.460 |
| Keap1 | 0.468 | 0.415 | 0.444 |
| Pla2g12a | 0.467 | 0.246 | 0.254 |
| Mia3 | 0.467 | 0.438 | 0.535 |
| Oxct1 | 0.467 | 0.849 | 0.755 |
| Ccdc85b | 0.467 | 0.263 | 0.179 |
| Fbl | 0.466 | 0.471 | 0.501 |
| Sox7 | 0.466 | 0.005 | 0.009 |
| Snrk | 0.466 | 0.295 | 0.324 |
| Elavl1 | 0.466 | 0.491 | 0.471 |
| Phip | 0.466 | 0.477 | 0.495 |
| Mob1b | 0.466 | 0.190 | 0.154 |
| Rab8a | 0.466 | 0.340 | 0.378 |
| Ccdc124 | 0.466 | 0.346 | 0.334 |
| Nrd1 | 0.466 | 0.380 | 0.377 |
| Fam168a | 0.465 | 0.509 | 0.378 |
| Acaa2 | 0.465 | 0.564 | 0.613 |
| Ganab | 0.465 | 0.280 | 0.307 |
| Ptpn1 | 0.465 | 0.535 | 0.342 |
| Plbd2 | 0.465 | 0.236 | 0.381 |
| Kdm5a | 0.465 | 0.469 | 0.479 |
| Chd6 | 0.465 | 0.507 | 0.396 |
| Isl1 | 0.465 | 0.013 | 0.037 |
| Nos1ap | 0.464 | 0.087 | 0.049 |
| Scarb2 | 0.464 | 0.550 | 0.512 |
| Aip | 0.464 | 0.420 | 0.389 |
| Mrpl4 | 0.464 | 0.367 | 0.281 |
| Tmem238 | 0.464 | 0.167 | 0.118 |
| Ufd1 | 0.464 | 0.348 | 0.276 |
| Mrps34 | 0.464 | 0.401 | 0.338 |
| Rbm7 | 0.464 | 0.588 | 0.453 |
| Ltbp1 | 0.464 | 1.668 | 0.923 |
| Rragc | 0.464 | 0.432 | 0.412 |
| Midn | 0.463 | 0.618 | 0.428 |
| Strbp | 0.463 | 0.311 | 0.189 |
| Rnps1 | 0.463 | 0.403 | 0.402 |
| Bag6 | 0.462 | 0.418 | 0.405 |
| Lamtor3 | 0.462 | 0.483 | 0.428 |
| Nudcd2 | 0.462 | 0.289 | 0.342 |
| Ctnnd1 | 0.462 | 0.271 | 0.206 |
| Stk24 | 0.462 | 0.344 | 0.277 |

| | | | |
|---|---|---|---|
| Trim44 | 0.461 | 0.433 | 0.381 |
| Fcgrt | 0.461 | 0.862 | 1.264 |
| Fbln1 | 0.461 | 0.290 | 1.905 |
| Mdp1 | 0.461 | 0.247 | 0.249 |
| Chtop | 0.461 | 0.412 | 0.392 |
| Grb2 | 0.460 | 0.361 | 0.335 |
| Fam210b | 0.460 | 0.121 | 0.042 |
| Syne2 | 0.460 | 0.352 | 0.342 |
| Sema3f | 0.459 | 0.201 | 0.144 |
| 1110008P14F | 0.459 | 0.319 | 0.261 |
| Jam2 | 0.458 | 0.086 | 0.154 |
| Dnajc19 | 0.458 | 0.441 | 0.421 |
| Herpud1 | 0.458 | 0.397 | 0.496 |
| Pmp22 | 0.457 | 1.027 | 1.575 |
| Camk1 | 0.457 | 0.650 | 0.603 |
| Vmp1 | 0.457 | 0.508 | 0.526 |
| Slmap | 0.457 | 0.783 | 0.360 |
| Mrpl11 | 0.457 | 0.394 | 0.374 |
| Tmed7 | 0.457 | 0.465 | 0.411 |
| Usp7 | 0.456 | 0.397 | 0.370 |
| Clint1 | 0.456 | 0.430 | 0.491 |
| Sod2 | 0.456 | 0.459 | 0.401 |
| Llph | 0.456 | 0.403 | 0.457 |
| Fibp | 0.456 | 0.366 | 0.321 |
| Rab13 | 0.456 | 0.390 | 0.496 |
| Ddx3y | 0.456 | 0.386 | 0.414 |
| Taf1d | 0.456 | 0.582 | 0.516 |
| Klf9 | 0.456 | 1.086 | 0.980 |
| Wdfy3 | 0.455 | 0.277 | 0.227 |
| Clcn4 | 0.455 | 0.481 | 0.490 |
| Mrps5 | 0.455 | 0.285 | 0.295 |
| Tmem106b | 0.454 | 0.397 | 0.401 |
| Sorbs2 | 0.454 | 0.225 | 0.040 |
| Lsm12 | 0.454 | 0.430 | 0.437 |
| Pcif1 | 0.454 | 0.394 | 0.303 |
| Mrps17 | 0.454 | 0.474 | 0.480 |
| Znrf1 | 0.454 | 0.379 | 0.289 |
| Znrd1 | 0.453 | 0.385 | 0.415 |
| Ythdf2 | 0.453 | 0.306 | 0.302 |
| Zeb1 | 0.453 | 0.550 | 0.626 |
| Podxl | 0.453 | 0.007 | 0.005 |
| Ppp1r37 | 0.453 | 0.292 | 0.290 |
| Ptn | 0.453 | 1.326 | 2.313 |
| Atp6v1d | 0.453 | 0.331 | 0.318 |
| Hint2 | 0.453 | 0.351 | 0.276 |
| Evl | 0.453 | 0.296 | 0.359 |
| Serf1 | 0.453 | 0.485 | 0.809 |
| Ube2j2 | 0.452 | 0.384 | 0.358 |

| | | | |
|---|---|---|---|
| Galk1 | 0.452 | 0.444 | 0.446 |
| Mfng | 0.452 | 0.003 | 0.003 |
| Ranbp2 | 0.452 | 0.444 | 0.351 |
| Eif4ebp2 | 0.452 | 0.360 | 0.358 |
| St6galnac3 | 0.452 | 0.005 | 0.009 |
| Dusp11 | 0.452 | 0.492 | 0.396 |
| Ccm2l | 0.451 | 0.003 | 0.003 |
| Kmt2a | 0.451 | 0.570 | 0.477 |
| Alkbh5 | 0.450 | 0.463 | 0.450 |
| Ano6 | 0.450 | 0.325 | 0.258 |
| Rnpepl1 | 0.450 | 0.295 | 0.300 |
| Fchsd2 | 0.450 | 0.323 | 0.227 |
| Ddx46 | 0.450 | 0.468 | 0.373 |
| Cops3 | 0.449 | 0.437 | 0.438 |
| Hspa12b | 0.449 | 0.038 | 0.085 |
| Dap3 | 0.449 | 0.393 | 0.411 |
| 11-Sep | 0.449 | 0.384 | 0.711 |
| Gnb4 | 0.449 | 0.814 | 0.470 |
| Bcl7b | 0.449 | 0.299 | 0.224 |
| Bcl10 | 0.448 | 0.485 | 0.516 |
| Wapl | 0.448 | 0.484 | 0.380 |
| Nsd3 | 0.448 | 0.487 | 0.481 |
| Chst12 | 0.447 | 0.224 | 0.327 |
| Slc39a1 | 0.447 | 0.604 | 0.576 |
| Tsc22d4 | 0.447 | 0.682 | 0.532 |
| Irf2bp2 | 0.447 | 0.677 | 0.518 |
| Tvp23b | 0.447 | 0.303 | 0.365 |
| Zfpl1 | 0.447 | 0.293 | 0.329 |
| Rchy1 | 0.447 | 0.487 | 0.416 |
| Lamc1 | 0.446 | 0.693 | 0.558 |
| BC005624 | 0.446 | 0.351 | 0.345 |
| Rad23b | 0.446 | 0.428 | 0.422 |
| Pacsin2 | 0.446 | 0.395 | 0.341 |
| Ctdsp1 | 0.446 | 0.398 | 0.424 |
| Crk | 0.445 | 0.362 | 0.365 |
| Cdc5l | 0.445 | 0.428 | 0.416 |
| Fam120a | 0.445 | 0.425 | 0.429 |
| Slc25a39 | 0.445 | 0.325 | 0.377 |
| Trappc3 | 0.445 | 0.377 | 0.391 |
| Fkbp11 | 0.445 | 0.215 | 0.507 |
| Pgp | 0.445 | 0.307 | 0.286 |
| 2210016L21F | 0.445 | 0.374 | 0.342 |
| Rsf1 | 0.445 | 0.451 | 0.387 |
| Plxna4 | 0.444 | 0.087 | 0.044 |
| Ccny | 0.444 | 0.247 | 0.253 |
| Babam1 | 0.444 | 0.278 | 0.291 |
| Fip1l1 | 0.444 | 0.421 | 0.480 |
| Efemp2 | 0.444 | 0.693 | 0.986 |

| | | | |
|---|---|---|---|
| Tmed1 | 0.444 | 0.441 | 0.504 |
| Prkag1 | 0.444 | 0.296 | 0.317 |
| Med29 | 0.444 | 0.314 | 0.300 |
| Manbal | 0.444 | 0.448 | 0.412 |
| Brd8 | 0.444 | 0.371 | 0.379 |
| Dnttip1 | 0.444 | 0.363 | 0.344 |
| Gimap6 | 0.444 | 0.013 | 0.019 |
| Cyth3 | 0.443 | 0.471 | 0.655 |
| Rad21 | 0.443 | 0.328 | 0.335 |
| Scamp2 | 0.443 | 0.351 | 0.408 |
| Atf3 | 0.443 | 1.205 | 1.244 |
| Stk25 | 0.443 | 0.348 | 0.365 |
| 2810474O19l | 0.442 | 0.514 | 0.486 |
| Stk16 | 0.442 | 0.354 | 0.361 |
| Utp11 | 0.442 | 0.467 | 0.423 |
| Sbno2 | 0.442 | 1.123 | 0.659 |
| Necap2 | 0.441 | 0.216 | 0.227 |
| Bcl9l | 0.441 | 0.555 | 0.417 |
| Numa1 | 0.441 | 0.391 | 0.372 |
| Pfdn6 | 0.441 | 0.482 | 0.469 |
| Mapkapk2 | 0.441 | 0.354 | 0.313 |
| Vbp1 | 0.441 | 0.359 | 0.350 |
| Pdgfd | 0.440 | 0.127 | 0.123 |
| Casp4 | 0.439 | 0.131 | 0.122 |
| Mrps26 | 0.439 | 0.356 | 0.361 |
| Triobp | 0.439 | 0.263 | 0.404 |
| Rab24 | 0.439 | 0.398 | 0.414 |
| Dctn6 | 0.438 | 0.550 | 0.345 |
| Nr3c1 | 0.438 | 0.657 | 0.684 |
| Entpd1 | 0.438 | 0.230 | 0.034 |
| Scamp3 | 0.438 | 0.330 | 0.340 |
| Zeb2 | 0.438 | 0.532 | 0.910 |
| Lamb2 | 0.438 | 0.933 | 0.645 |
| Crlf2 | 0.437 | 0.379 | 0.317 |
| Ctbp2 | 0.437 | 0.247 | 0.225 |
| Stag1 | 0.437 | 0.416 | 0.489 |
| Cers5 | 0.437 | 0.375 | 0.337 |
| Eif2s1 | 0.437 | 0.360 | 0.353 |
| 0610037L13F | 0.437 | 0.352 | 0.408 |
| BC004004 | 0.436 | 0.313 | 0.380 |
| P4ha1 | 0.436 | 0.249 | 0.440 |
| Vezf1 | 0.436 | 0.542 | 0.439 |
| Zfp260 | 0.436 | 0.348 | 0.470 |
| Glyr1 | 0.436 | 0.365 | 0.358 |
| Tmem259 | 0.436 | 0.416 | 0.370 |
| Dgcr2 | 0.436 | 0.265 | 0.253 |
| Cyth2 | 0.436 | 0.442 | 0.352 |
| Dctn1 | 0.436 | 0.371 | 0.399 |

| | | | |
|---|---|---|---|
| Ctps | 0.435 | 0.234 | 0.167 |
| Vps35 | 0.435 | 0.512 | 0.522 |
| Rab27b | 0.434 | 0.141 | 0.046 |
| Npepl1 | 0.434 | 0.323 | 0.417 |
| Bfar | 0.434 | 0.395 | 0.554 |
| Brd9 | 0.434 | 0.357 | 0.346 |
| Eri3 | 0.434 | 0.387 | 0.518 |
| Immp1l | 0.434 | 0.453 | 0.425 |
| Rbck1 | 0.434 | 0.260 | 0.294 |
| Scfd1 | 0.434 | 0.395 | 0.417 |
| Smarcb1 | 0.434 | 0.457 | 0.430 |
| Taf6l | 0.434 | 0.293 | 0.328 |
| Igfbp7 | 0.434 | 2.404 | 2.061 |
| Mllt3 | 0.432 | 0.099 | 0.120 |
| Med8 | 0.432 | 0.308 | 0.323 |
| Cdk9 | 0.432 | 0.437 | 0.375 |
| Pitpnb | 0.432 | 0.387 | 0.401 |
| Vti1b | 0.432 | 0.322 | 0.336 |
| Hdac2 | 0.432 | 0.403 | 0.399 |
| Sdhd | 0.432 | 0.366 | 0.324 |
| Pld3 | 0.432 | 0.244 | 0.373 |
| Rbms2 | 0.432 | 0.342 | 0.405 |
| Pdzk1ip1 | 0.432 | 0.003 | 0.001 |
| Tia1 | 0.432 | 0.421 | 0.349 |
| Slc7a1 | 0.431 | 0.213 | 0.179 |
| Rbm22 | 0.431 | 0.386 | 0.333 |
| Med10 | 0.431 | 0.349 | 0.387 |
| Csnk2a2 | 0.431 | 0.402 | 0.371 |
| Lsm14a | 0.431 | 0.350 | 0.337 |
| Ube2e1 | 0.431 | 0.483 | 0.419 |
| Mtx2 | 0.430 | 0.340 | 0.404 |
| Msi2 | 0.430 | 0.496 | 0.394 |
| Pdha1 | 0.429 | 0.473 | 0.470 |
| Ppp1r13b | 0.429 | 0.088 | 0.075 |
| Gatsl3 | 0.429 | 0.109 | 0.264 |
| Alyref | 0.429 | 0.355 | 0.342 |
| Rtf2 | 0.429 | 0.333 | 0.321 |
| Fopnl | 0.429 | 0.338 | 0.347 |
| Arap3 | 0.428 | 0.005 | 0.008 |
| Knop1 | 0.428 | 0.344 | 0.438 |
| Snx4 | 0.428 | 0.459 | 0.431 |
| Dock4 | 0.428 | 0.062 | 0.123 |
| Agrn | 0.428 | 0.490 | 0.207 |
| Cnot6l | 0.428 | 0.386 | 0.340 |
| Bod1 | 0.428 | 0.556 | 0.491 |
| Slc35b1 | 0.427 | 0.322 | 0.272 |
| Abhd16a | 0.427 | 0.330 | 0.379 |
| Srpk2 | 0.427 | 0.502 | 0.432 |

| | | | |
|---|---|---|---|
| Tusc3 | 0.427 | 0.520 | 0.565 |
| Nsrp1 | 0.427 | 0.393 | 0.341 |
| Hnrnph2 | 0.427 | 0.400 | 0.342 |
| Itsn2 | 0.426 | 0.227 | 0.196 |
| Tlk1 | 0.426 | 0.347 | 0.292 |
| Fam96a | 0.426 | 0.323 | 0.388 |
| Gadd45gip1 | 0.426 | 0.408 | 0.381 |
| Hipk1 | 0.426 | 0.379 | 0.246 |
| Chchd7 | 0.426 | 0.353 | 0.376 |
| Ggh | 0.426 | 0.338 | 0.394 |
| Ndufa9 | 0.426 | 0.381 | 0.359 |
| Wtip | 0.426 | 0.636 | 0.458 |
| Eea1 | 0.425 | 0.556 | 0.451 |
| Wipi1 | 0.425 | 0.504 | 0.880 |
| Tgfb2 | 0.425 | 0.803 | 0.505 |
| Capza1 | 0.425 | 0.321 | 0.341 |
| Arih1 | 0.425 | 0.382 | 0.363 |
| Tjp1 | 0.425 | 0.269 | 0.203 |
| Cops2 | 0.425 | 0.460 | 0.428 |
| Asph | 0.425 | 0.424 | 0.288 |
| S100a1 | 0.425 | 0.400 | 0.221 |
| Epha4 | 0.424 | 0.229 | 0.149 |
| Kxd1 | 0.424 | 0.408 | 0.374 |
| Mcts1 | 0.424 | 0.357 | 0.373 |
| Rasl11b | 0.424 | 0.438 | 0.265 |
| Eif2b2 | 0.424 | 0.214 | 0.229 |
| Ubr4 | 0.424 | 0.371 | 0.346 |
| Fam96b | 0.424 | 0.403 | 0.323 |
| Snrpa | 0.424 | 0.346 | 0.375 |
| Tmem128 | 0.423 | 0.261 | 0.227 |
| Tcf7l2 | 0.423 | 0.559 | 0.429 |
| Arfgap2 | 0.423 | 0.331 | 0.312 |
| Akr1b3 | 0.423 | 0.371 | 0.380 |
| Lsm8 | 0.423 | 0.433 | 0.350 |
| Igbp1 | 0.422 | 0.366 | 0.346 |
| Lxn | 0.422 | 0.349 | 0.293 |
| Med13l | 0.422 | 0.797 | 0.736 |
| Oxr1 | 0.422 | 0.177 | 0.227 |
| Tmem263 | 0.422 | 0.379 | 0.471 |
| Upf3a | 0.421 | 0.404 | 0.380 |
| Yme1l1 | 0.421 | 0.412 | 0.414 |
| Tnfsf12 | 0.421 | 0.335 | 0.529 |
| Apc | 0.421 | 0.256 | 0.246 |
| Fkbp2 | 0.421 | 0.481 | 0.508 |
| Sox18 | 0.421 | 0.002 | 0.003 |
| Ptch1 | 0.420 | 0.206 | 0.238 |
| Tomm40 | 0.420 | 0.359 | 0.319 |
| Mat2b | 0.420 | 0.423 | 0.423 |

| | | | |
|---|---|---|---|
| Eif4e | 0.420 | 0.391 | 0.360 |
| Arl2 | 0.420 | 0.332 | 0.280 |
| Vgll4 | 0.420 | 0.545 | 0.423 |
| Insig2 | 0.420 | 0.365 | 0.354 |
| Rgl2 | 0.419 | 0.162 | 0.150 |
| Slco3a1 | 0.419 | 0.292 | 0.196 |
| Baz1b | 0.419 | 0.372 | 0.399 |
| Malsu1 | 0.419 | 0.377 | 0.381 |
| Churc1 | 0.418 | 0.426 | 0.350 |
| Ndufaf8 | 0.418 | 0.299 | 0.303 |
| Stk11 | 0.418 | 0.393 | 0.317 |
| Elf2 | 0.418 | 0.372 | 0.380 |
| Camta1 | 0.418 | 0.388 | 0.267 |
| Parl | 0.418 | 0.343 | 0.284 |
| Ltbr | 0.418 | 0.348 | 0.341 |
| 1810022K09F | 0.417 | 0.399 | 0.417 |
| Actr1a | 0.417 | 0.271 | 0.297 |
| Spr | 0.417 | 0.259 | 0.276 |
| Snap23 | 0.417 | 0.293 | 0.275 |
| Becn1 | 0.417 | 0.282 | 0.265 |
| Fam189a2 | 0.416 | 0.066 | 0.047 |
| Nfkbiz | 0.416 | 1.082 | 0.929 |
| Ppm1g | 0.416 | 0.366 | 0.331 |
| Kpnb1 | 0.416 | 0.347 | 0.430 |
| Slc38a10 | 0.416 | 0.261 | 0.437 |
| Zc3h14 | 0.416 | 0.430 | 0.391 |
| Atp6ap2 | 0.416 | 0.490 | 0.406 |
| Yipf1 | 0.416 | 0.299 | 0.329 |
| Hadhb | 0.415 | 0.466 | 0.349 |
| Gas6 | 0.415 | 0.713 | 0.890 |
| Wdr61 | 0.415 | 0.355 | 0.376 |
| Wbp1 | 0.415 | 0.211 | 0.216 |
| Retreg2 | 0.415 | 0.270 | 0.355 |
| Naglu | 0.415 | 0.136 | 0.182 |
| Mtus1 | 0.415 | 0.474 | 0.360 |
| Rbms3 | 0.415 | 0.764 | 0.930 |
| Kdm1a | 0.415 | 0.440 | 0.430 |
| Sqor | 0.415 | 0.068 | 0.187 |
| Pde4b | 0.415 | 0.557 | 0.256 |
| Dapk3 | 0.415 | 0.408 | 0.366 |
| Cited2 | 0.414 | 0.279 | 0.317 |
| Anapc2 | 0.414 | 0.319 | 0.360 |
| Snx2 | 0.414 | 0.416 | 0.470 |
| Smchd1 | 0.414 | 0.468 | 0.411 |
| Rbm6 | 0.414 | 0.342 | 0.348 |
| Snapc5 | 0.413 | 0.307 | 0.323 |
| Exosc4 | 0.413 | 0.293 | 0.292 |
| Safb | 0.413 | 0.380 | 0.382 |

| | | | |
|---|---|---|---|
| 2310039H08I | 0.413 | 0.366 | 0.471 |
| 2010111I01R | 0.413 | 0.542 | 0.215 |
| Twf1 | 0.413 | 0.365 | 0.433 |
| Mocs2 | 0.413 | 0.362 | 0.434 |
| Cdk5rap3 | 0.413 | 0.379 | 0.446 |
| Glod4 | 0.413 | 0.303 | 0.336 |
| Sec11c | 0.413 | 0.261 | 0.156 |
| Sf1 | 0.413 | 0.416 | 0.405 |
| Glb1 | 0.412 | 0.161 | 0.277 |
| Pdrg1 | 0.412 | 0.246 | 0.308 |
| Itga1 | 0.412 | 0.370 | 0.211 |
| Tor1b | 0.412 | 0.286 | 0.296 |
| Zfp638 | 0.412 | 0.388 | 0.390 |
| Taok2 | 0.412 | 0.154 | 0.193 |
| Gtpbp4 | 0.412 | 0.397 | 0.351 |
| Armcx3 | 0.412 | 0.374 | 0.531 |
| BC005561 | 0.411 | 0.424 | 0.347 |
| Ahcyl1 | 0.411 | 0.415 | 0.436 |
| St3gal1 | 0.411 | 0.165 | 0.149 |
| Nhp2 | 0.410 | 0.319 | 0.337 |
| Leprotl1 | 0.410 | 0.362 | 0.300 |
| Tspan14 | 0.410 | 0.066 | 0.097 |
| Eif2a | 0.410 | 0.456 | 0.387 |
| Dync1li1 | 0.410 | 0.363 | 0.272 |
| H1fx | 0.409 | 0.447 | 0.472 |
| Ola1 | 0.409 | 0.466 | 0.405 |
| Wasl | 0.409 | 0.439 | 0.438 |
| Ubl4a | 0.409 | 0.254 | 0.310 |
| Rnf215 | 0.409 | 0.259 | 0.257 |
| Actr1b | 0.409 | 0.389 | 0.325 |
| Ptgs1 | 0.409 | 0.083 | 0.086 |
| Riok3 | 0.408 | 0.321 | 0.255 |
| Ahsa1 | 0.408 | 0.434 | 0.426 |
| Cpsf6 | 0.408 | 0.374 | 0.358 |
| Med21 | 0.408 | 0.418 | 0.430 |
| Ebna1bp2 | 0.408 | 0.359 | 0.313 |
| Ddx54 | 0.408 | 0.302 | 0.299 |
| Ppp6c | 0.408 | 0.310 | 0.273 |
| Rbm17 | 0.408 | 0.443 | 0.427 |
| Atp2b4 | 0.408 | 0.522 | 0.362 |
| Gpatch8 | 0.408 | 0.432 | 0.398 |
| Bckdk | 0.407 | 0.258 | 0.288 |
| Ext1 | 0.407 | 0.602 | 0.508 |
| Mien1 | 0.407 | 0.307 | 0.338 |
| Glt8d1 | 0.407 | 0.398 | 0.536 |
| Tpd52 | 0.407 | 0.157 | 0.069 |
| Psmd14 | 0.406 | 0.375 | 0.373 |
| Esf1 | 0.406 | 0.413 | 0.340 |

| | | | |
|---|---|---|---|
| Cnot3 | 0.406 | 0.362 | 0.400 |
| Eif4g3 | 0.406 | 0.406 | 0.411 |
| Osgep | 0.406 | 0.372 | 0.404 |
| Etf1 | 0.405 | 0.432 | 0.420 |
| Prkacb | 0.405 | 0.514 | 0.372 |
| Man2a1 | 0.405 | 0.717 | 0.607 |
| Dguok | 0.405 | 0.364 | 0.424 |
| Nelfb | 0.405 | 0.262 | 0.298 |
| Bbx | 0.405 | 0.371 | 0.377 |
| Acat1 | 0.405 | 0.489 | 0.600 |
| Ube2z | 0.404 | 0.318 | 0.352 |
| Actr2 | 0.404 | 0.388 | 0.317 |
| Ddx42 | 0.404 | 0.352 | 0.348 |
| Chd8 | 0.404 | 0.386 | 0.367 |
| Dda1 | 0.404 | 0.343 | 0.328 |
| Csnk2a1 | 0.404 | 0.376 | 0.346 |
| Nop58 | 0.404 | 0.454 | 0.561 |
| Zmym5 | 0.404 | 0.389 | 0.379 |
| Eapp | 0.403 | 0.366 | 0.342 |
| Mrpl21 | 0.403 | 0.404 | 0.364 |
| Fam76a | 0.403 | 0.452 | 0.406 |
| Adgre5 | 0.403 | 0.284 | 0.100 |
| Tmem205 | 0.403 | 0.245 | 0.258 |
| Smco4 | 0.403 | 0.219 | 0.269 |
| Nsmce1 | 0.403 | 0.386 | 0.426 |
| Mgat4b | 0.403 | 0.241 | 0.200 |
| Upf3b | 0.403 | 0.375 | 0.203 |
| Rai14 | 0.403 | 0.178 | 0.294 |
| Hnrnpul1 | 0.402 | 0.399 | 0.425 |
| Ing4 | 0.402 | 0.353 | 0.369 |
| Prpsap1 | 0.402 | 0.344 | 0.288 |
| Arhgef15 | 0.402 | 0.004 | 0.006 |
| Xiap | 0.402 | 0.326 | 0.310 |
| Zfp771 | 0.401 | 0.592 | 0.517 |
| Nr2f6 | 0.401 | 0.445 | 0.354 |
| Rnf19a | 0.401 | 0.289 | 0.238 |
| Ric8a | 0.401 | 0.262 | 0.346 |
| Trip6 | 0.401 | 0.452 | 0.569 |
| Tab2 | 0.401 | 0.401 | 0.354 |
| Brd7 | 0.401 | 0.522 | 0.481 |
| Ankrd13a | 0.400 | 0.197 | 0.230 |
| Arl4a | 0.400 | 0.541 | 0.501 |
| Zfp326 | 0.400 | 0.370 | 0.349 |
| Naa50 | 0.400 | 0.421 | 0.429 |
| Trappc5 | 0.400 | 0.307 | 0.293 |
| Cops7a | 0.400 | 0.257 | 0.285 |
| Ube2v2 | 0.400 | 0.464 | 0.395 |
| Tbc1d20 | 0.400 | 0.321 | 0.344 |

| | | | |
|---|---|---|---|
| Git1 | 0.399 | 0.184 | 0.225 |
| Zcchc7 | 0.399 | 0.353 | 0.330 |
| Spata6 | 0.399 | 0.143 | 0.156 |
| Tpp2 | 0.399 | 0.379 | 0.379 |
| Nbeal1 | 0.399 | 0.486 | 0.442 |
| Rabl6 | 0.399 | 0.327 | 0.366 |
| Ppp2r2d | 0.399 | 0.422 | 0.443 |
| Tmem184b | 0.399 | 0.189 | 0.219 |
| Ppt2 | 0.399 | 0.301 | 0.221 |
| Ss18 | 0.399 | 0.264 | 0.255 |
| Ak2 | 0.399 | 0.418 | 0.334 |
| Smu1 | 0.399 | 0.375 | 0.331 |
| Pigk | 0.398 | 0.280 | 0.309 |
| Anapc4 | 0.398 | 0.319 | 0.315 |
| Ppil2 | 0.398 | 0.307 | 0.325 |
| Npepps | 0.398 | 0.429 | 0.465 |
| Angel2 | 0.398 | 0.333 | 0.324 |
| Supt5 | 0.397 | 0.388 | 0.389 |
| Fam8a1 | 0.397 | 0.320 | 0.295 |
| Arid2 | 0.397 | 0.388 | 0.346 |
| Fcho2 | 0.397 | 0.336 | 0.298 |
| Smpd1 | 0.397 | 0.292 | 0.261 |
| Zranb1 | 0.397 | 0.511 | 0.469 |
| Slirp | 0.397 | 0.341 | 0.335 |
| Pet100 | 0.397 | 0.254 | 0.248 |
| Sbds | 0.397 | 0.356 | 0.348 |
| St3gal4 | 0.397 | 0.208 | 0.306 |
| Ptprm | 0.397 | 0.105 | 0.073 |
| Scarf2 | 0.397 | 0.721 | 0.693 |
| Eif1a | 0.397 | 0.634 | 0.607 |
| Ccnt1 | 0.397 | 0.380 | 0.354 |
| Trp53i13 | 0.397 | 0.319 | 0.409 |
| Fam174a | 0.397 | 0.386 | 0.297 |
| Dynll2 | 0.397 | 0.310 | 0.308 |
| Angptl2 | 0.396 | 0.841 | 0.731 |
| Mpdu1 | 0.396 | 0.294 | 0.308 |
| Dkk3 | 0.396 | 0.855 | 1.010 |
| Srsf1 | 0.396 | 0.343 | 0.333 |
| Stx16 | 0.396 | 0.316 | 0.369 |
| Usp3 | 0.396 | 0.292 | 0.239 |
| Rtl8a | 0.395 | 0.443 | 0.451 |
| Adk | 0.395 | 0.252 | 0.208 |
| Tardbp | 0.395 | 0.336 | 0.365 |
| Mrpl36 | 0.395 | 0.293 | 0.276 |
| Uap1 | 0.395 | 0.332 | 0.362 |
| Otub1 | 0.395 | 0.379 | 0.358 |
| Pnpla8 | 0.394 | 0.424 | 0.413 |
| Mrpl15 | 0.394 | 0.371 | 0.380 |

| | | | |
|---|---|---|---|
| Wtap | 0.394 | 0.299 | 0.312 |
| Glce | 0.394 | 0.078 | 0.081 |
| Pkn1 | 0.394 | 0.191 | 0.185 |
| Paxx | 0.394 | 0.279 | 0.355 |
| Cisd2 | 0.394 | 0.429 | 0.435 |
| Smc4 | 0.394 | 0.370 | 0.291 |
| Arhgef28 | 0.393 | 0.058 | 0.162 |
| Cdc42bpb | 0.393 | 0.260 | 0.242 |
| Slc35b2 | 0.393 | 0.304 | 0.387 |
| Aak1 | 0.393 | 0.245 | 0.317 |
| Hmox2 | 0.393 | 0.323 | 0.327 |
| Dpagt1 | 0.393 | 0.108 | 0.133 |
| Nt5c3 | 0.393 | 0.178 | 0.155 |
| Tmem189 | 0.393 | 0.238 | 0.179 |
| Rc3h2 | 0.393 | 0.326 | 0.297 |
| Stt3b | 0.392 | 0.479 | 0.507 |
| Atg101 | 0.392 | 0.271 | 0.295 |
| 2310011J03R | 0.392 | 0.247 | 0.277 |
| Itch | 0.392 | 0.391 | 0.353 |
| Nr1h2 | 0.392 | 0.326 | 0.281 |
| Gstm5 | 0.392 | 0.506 | 0.750 |
| Dnttip2 | 0.392 | 0.431 | 0.444 |
| Emc8 | 0.392 | 0.639 | 0.382 |
| Unc5b | 0.392 | 0.242 | 0.223 |
| Kdelc2 | 0.391 | 0.256 | 0.391 |
| Lsm1 | 0.391 | 0.370 | 0.339 |
| 9930111J21R | 0.391 | 0.031 | 0.017 |
| Zdhhc3 | 0.391 | 0.253 | 0.348 |
| R3hdm4 | 0.391 | 0.485 | 0.506 |
| Mrpl27 | 0.391 | 0.369 | 0.337 |
| Rbm28 | 0.391 | 0.277 | 0.290 |
| Supt16 | 0.391 | 0.386 | 0.370 |
| Tbc1d17 | 0.391 | 0.316 | 0.349 |
| Gpbp1 | 0.390 | 0.390 | 0.450 |
| Nasp | 0.390 | 0.495 | 0.447 |
| Sema6a | 0.390 | 0.192 | 0.153 |
| Arfgap1 | 0.390 | 0.290 | 0.430 |
| Etnk1 | 0.390 | 0.205 | 0.245 |
| Ccs | 0.390 | 0.372 | 0.362 |
| Coro1b | 0.389 | 0.328 | 0.419 |
| Rab22a | 0.389 | 0.347 | 0.424 |
| Isyna1 | 0.389 | 0.409 | 0.178 |
| Qars | 0.389 | 0.340 | 0.303 |
| Mea1 | 0.389 | 0.330 | 0.340 |
| Setd2 | 0.389 | 0.400 | 0.367 |
| Pam16 | 0.388 | 0.369 | 0.325 |
| Cmtm7 | 0.388 | 0.342 | 0.379 |
| Tmem65 | 0.388 | 0.367 | 0.359 |

| | | | |
|---|---|---|---|
| Foxo1 | 0.388 | 0.322 | 0.297 |
| Kmt5b | 0.388 | 0.342 | 0.296 |
| Anp32e | 0.388 | 0.560 | 0.323 |
| Efcab14 | 0.387 | 0.284 | 0.303 |
| Yeats4 | 0.387 | 0.347 | 0.305 |
| Emc2 | 0.387 | 0.376 | 0.407 |
| Fam160b1 | 0.387 | 0.295 | 0.294 |
| Dpy30 | 0.387 | 0.377 | 0.346 |
| Gm26532 | 0.387 | 0.902 | 0.575 |
| Fbxo9 | 0.387 | 0.306 | 0.256 |
| Smagp | 0.387 | 0.009 | 0.009 |
| Pigx | 0.386 | 0.373 | 0.401 |
| Fez1 | 0.386 | 0.470 | 0.366 |
| Mgat1 | 0.386 | 0.354 | 0.418 |
| Nfkb1 | 0.386 | 0.328 | 0.329 |
| Gps2 | 0.386 | 0.314 | 0.299 |
| Map3k1 | 0.386 | 0.166 | 0.100 |
| Rnase4 | 0.386 | 0.270 | 0.768 |
| Vamp2 | 0.385 | 0.425 | 0.425 |
| Cebpz | 0.385 | 0.392 | 0.396 |
| Tmem57 | 0.385 | 0.337 | 0.335 |
| Chd2 | 0.385 | 0.404 | 0.386 |
| Txnl4a | 0.385 | 0.361 | 0.388 |
| Dtymk | 0.385 | 0.399 | 0.463 |
| Commd4 | 0.385 | 0.298 | 0.264 |
| Ly6e | 0.385 | 0.054 | 0.313 |
| Uba7 | 0.385 | 0.151 | 0.164 |
| B4gat1 | 0.385 | 0.268 | 0.307 |
| Prrc1 | 0.384 | 0.374 | 0.450 |
| U2af2 | 0.384 | 0.333 | 0.329 |
| Edem2 | 0.384 | 0.135 | 0.182 |
| Gpr107 | 0.384 | 0.305 | 0.325 |
| Pde3b | 0.384 | 0.106 | 0.054 |
| Dusp23 | 0.384 | 0.107 | 0.192 |
| Tnrc18 | 0.384 | 0.514 | 0.431 |
| Pnrc2 | 0.383 | 0.314 | 0.332 |
| Ncoa4 | 0.383 | 0.367 | 0.214 |
| Ctdspl | 0.383 | 0.498 | 0.343 |
| Ntn4 | 0.383 | 0.011 | 0.008 |
| Hjurp | 0.382 | 0.352 | 0.449 |
| Lpar1 | 0.382 | 0.414 | 0.838 |
| Itgb1bp1 | 0.382 | 0.278 | 0.324 |
| Ccz1 | 0.382 | 0.286 | 0.275 |
| Mrpl32 | 0.382 | 0.397 | 0.311 |
| Scyl1 | 0.382 | 0.293 | 0.367 |
| Mmp16 | 0.382 | 0.331 | 0.386 |
| Arf3 | 0.382 | 0.282 | 0.252 |
| Ap2a1 | 0.381 | 0.367 | 0.330 |

| | | | |
|---|---|---|---|
| Vta1 | 0.381 | 0.272 | 0.234 |
| Smtn | 0.381 | 1.009 | 0.427 |
| Amfr | 0.381 | 0.316 | 0.261 |
| Ifnar2 | 0.381 | 0.266 | 0.332 |
| Jagn1 | 0.381 | 0.277 | 0.240 |
| Tmem55b | 0.380 | 0.259 | 0.299 |
| Sec23a | 0.380 | 0.371 | 0.426 |
| Elovl5 | 0.380 | 0.264 | 0.208 |
| Zfyve21 | 0.380 | 0.433 | 0.486 |
| Abi1 | 0.380 | 0.422 | 0.323 |
| Fyttd1 | 0.380 | 0.453 | 0.354 |
| Rhbdf1 | 0.380 | 0.258 | 0.314 |
| Flii | 0.380 | 0.282 | 0.280 |
| Tcf3 | 0.379 | 0.301 | 0.377 |
| Tmem248 | 0.379 | 0.219 | 0.247 |
| Usp15 | 0.379 | 0.307 | 0.238 |
| Araf | 0.379 | 0.361 | 0.364 |
| 1700123O20I | 0.379 | 0.330 | 0.312 |
| Sp3 | 0.379 | 0.407 | 0.379 |
| Tiprl | 0.378 | 0.444 | 0.383 |
| Dgcr6 | 0.378 | 0.340 | 0.331 |
| Peli1 | 0.378 | 0.372 | 0.335 |
| Pcbp4 | 0.378 | 0.314 | 0.404 |
| Stx8 | 0.378 | 0.313 | 0.337 |
| Ago2 | 0.378 | 0.545 | 0.524 |
| Tnpo3 | 0.378 | 0.325 | 0.322 |
| Med25 | 0.378 | 0.371 | 0.378 |
| Mdfi | 0.378 | 0.082 | 0.188 |
| Drg1 | 0.378 | 0.337 | 0.314 |
| Vps4b | 0.378 | 0.338 | 0.340 |
| Mrps15 | 0.377 | 0.396 | 0.414 |
| Csnk1g3 | 0.377 | 0.350 | 0.347 |
| Fam173a | 0.377 | 0.317 | 0.302 |
| Atl2 | 0.377 | 0.207 | 0.139 |
| Txlna | 0.377 | 0.314 | 0.373 |
| Ppa1 | 0.377 | 0.273 | 0.322 |
| Clns1a | 0.377 | 0.296 | 0.322 |
| Dbn1 | 0.376 | 0.426 | 0.652 |
| Emp2 | 0.376 | 0.672 | 0.376 |
| Smurf2 | 0.376 | 0.261 | 0.345 |
| Tfdp2 | 0.376 | 0.296 | 0.243 |
| Dpm1 | 0.375 | 0.360 | 0.361 |
| Lin7c | 0.375 | 0.289 | 0.211 |
| Tspan18 | 0.375 | 0.129 | 0.136 |
| Crlf3 | 0.375 | 0.119 | 0.132 |
| Ccdc186 | 0.375 | 0.332 | 0.356 |
| Cdkn1a | 0.375 | 0.715 | 0.441 |
| Nptn | 0.375 | 0.849 | 0.639 |

| | | | |
|---|---|---|---|
| Hoxa4 | 0.375 | 0.244 | 0.155 |
| Sdhc | 0.375 | 0.441 | 0.388 |
| Raph1 | 0.375 | 0.488 | 0.730 |
| Grcc10 | 0.375 | 0.315 | 0.316 |
| Creld1 | 0.375 | 0.154 | 0.163 |
| Psmd9 | 0.375 | 0.275 | 0.294 |
| Commd8 | 0.375 | 0.331 | 0.311 |
| Fdx1 | 0.375 | 0.340 | 0.280 |
| Fzd6 | 0.375 | 0.060 | 0.051 |
| Specc1 | 0.374 | 0.364 | 0.156 |
| Acot7 | 0.374 | 0.348 | 0.231 |
| Avpi1 | 0.374 | 0.206 | 0.095 |
| Gm12840 | 0.374 | 1.353 | 0.854 |
| Tet3 | 0.374 | 0.406 | 0.313 |
| Nub1 | 0.374 | 0.302 | 0.299 |
| Tspan9 | 0.374 | 0.174 | 0.170 |
| Cpne3 | 0.374 | 0.424 | 0.339 |
| Nck1 | 0.373 | 0.163 | 0.211 |
| Pde4d | 0.373 | 0.454 | 0.154 |
| Mrps6 | 0.373 | 0.516 | 0.346 |
| Cdk16 | 0.373 | 0.252 | 0.441 |
| Pdlim4 | 0.373 | 0.452 | 0.264 |
| Klc1 | 0.373 | 0.348 | 0.352 |
| Mettl1 | 0.373 | 0.276 | 0.474 |
| Arl16 | 0.373 | 0.162 | 0.153 |
| Lats2 | 0.373 | 0.283 | 0.277 |
| Ppp6r3 | 0.373 | 0.330 | 0.302 |
| Tm9sf4 | 0.373 | 0.285 | 0.396 |
| Zfand3 | 0.372 | 0.331 | 0.364 |
| Smo | 0.372 | 0.468 | 0.514 |
| Pkhd1l1 | 0.372 | 0.003 | 0.005 |
| Cnot1 | 0.372 | 0.340 | 0.345 |
| Fam133b | 0.372 | 0.388 | 0.338 |
| Golph3 | 0.372 | 0.362 | 0.388 |
| Larp7 | 0.372 | 0.330 | 0.356 |
| Spg21 | 0.372 | 0.410 | 0.408 |
| Pycr2 | 0.372 | 0.238 | 0.218 |
| Zbtb38 | 0.372 | 0.570 | 0.369 |
| Glul | 0.372 | 0.400 | 0.388 |
| Maz | 0.372 | 0.443 | 0.430 |
| Mark2 | 0.371 | 0.361 | 0.306 |
| Rsbn1l | 0.371 | 0.396 | 0.295 |
| Naa15 | 0.371 | 0.392 | 0.329 |
| Mrpl2 | 0.371 | 0.293 | 0.293 |
| Bcl6b | 0.371 | 0.004 | 0.005 |
| Helz | 0.371 | 0.440 | 0.407 |
| Tor1a | 0.371 | 0.315 | 0.300 |
| Map3k11 | 0.371 | 0.056 | 0.048 |

| | | | |
|---|---|---|---|
| Actl6a | 0.371 | 0.272 | 0.247 |
| Tpgs1 | 0.370 | 0.339 | 0.277 |
| Gcnt1 | 0.370 | 0.008 | 0.011 |
| Anxa11 | 0.370 | 0.415 | 0.263 |
| Tm7sf3 | 0.370 | 0.301 | 0.513 |
| Sh2b3 | 0.370 | 0.130 | 0.117 |
| Dnajb4 | 0.370 | 0.622 | 0.564 |
| Timm22 | 0.370 | 0.295 | 0.303 |
| Mgea5 | 0.369 | 0.328 | 0.318 |
| Cnep1r1 | 0.369 | 0.287 | 0.315 |
| Ppid | 0.369 | 0.373 | 0.356 |
| Cab39 | 0.369 | 0.394 | 0.274 |
| Kif26a | 0.369 | 0.006 | 0.027 |
| Agpat4 | 0.368 | 0.358 | 0.412 |
| Nans | 0.368 | 0.252 | 0.373 |
| Lsm5 | 0.368 | 0.386 | 0.318 |
| Rbm26 | 0.368 | 0.385 | 0.369 |
| Snd1 | 0.368 | 0.348 | 0.305 |
| Sidt2 | 0.368 | 0.489 | 0.439 |
| Ptpa | 0.368 | 0.374 | 0.341 |
| Esyt1 | 0.368 | 0.343 | 0.296 |
| Adgrl2 | 0.368 | 0.603 | 0.379 |
| Tmx3 | 0.368 | 0.233 | 0.201 |
| Rbmxl1 | 0.368 | 0.466 | 0.428 |
| Acot9 | 0.367 | 0.288 | 0.312 |
| Nedd9 | 0.367 | 0.522 | 0.401 |
| Phb | 0.367 | 0.324 | 0.334 |
| Nek7 | 0.367 | 0.376 | 0.323 |
| Mrps23 | 0.367 | 0.274 | 0.270 |
| Armcx2 | 0.367 | 0.357 | 0.417 |
| Sec24d | 0.367 | 0.450 | 0.477 |
| Capg | 0.367 | 0.329 | 0.286 |
| Gramd1a | 0.367 | 0.321 | 0.289 |
| Ubr2 | 0.367 | 0.366 | 0.328 |
| Acadvl | 0.367 | 0.375 | 0.300 |
| AI837181 | 0.366 | 0.285 | 0.286 |
| Cox17 | 0.366 | 0.247 | 0.178 |
| Rsph9 | 0.366 | 0.118 | 0.075 |
| Igf1 | 0.366 | 0.391 | 1.433 |
| Elp5 | 0.366 | 0.278 | 0.316 |
| Rrp7a | 0.366 | 0.228 | 0.257 |
| Fbxo3 | 0.366 | 0.325 | 0.281 |
| Lamc3 | 0.366 | 0.026 | 0.014 |
| Papss1 | 0.366 | 0.419 | 0.414 |
| Creb3 | 0.366 | 0.337 | 0.328 |
| Cd82 | 0.366 | 0.135 | 0.049 |
| Mrps36 | 0.365 | 0.452 | 0.435 |
| Stard3nl | 0.365 | 0.349 | 0.367 |

| | | | |
|---|---|---|---|
| Trpm7 | 0.365 | 0.447 | 0.454 |
| Mtfr1l | 0.365 | 0.374 | 0.417 |
| Wipi2 | 0.365 | 0.287 | 0.274 |
| Twistnb | 0.365 | 0.333 | 0.304 |
| Nbr1 | 0.365 | 0.267 | 0.271 |
| Cks1b | 0.365 | 0.448 | 0.527 |
| Zc3h7a | 0.365 | 0.275 | 0.277 |
| Arl8a | 0.364 | 0.441 | 0.423 |
| Ebp | 0.364 | 0.218 | 0.251 |
| Snrpa1 | 0.364 | 0.309 | 0.311 |
| Ptpn2 | 0.364 | 0.343 | 0.247 |
| Fmc1 | 0.364 | 0.344 | 0.327 |
| Trnau1ap | 0.364 | 0.295 | 0.284 |
| Btbd3 | 0.364 | 0.153 | 0.133 |
| Spint2 | 0.364 | 0.310 | 0.067 |
| Rabgap1 | 0.364 | 0.330 | 0.300 |
| Tm2d3 | 0.364 | 0.338 | 0.308 |
| Herc1 | 0.364 | 0.343 | 0.288 |
| Ppp4c | 0.363 | 0.273 | 0.293 |
| Srpk1 | 0.363 | 0.333 | 0.323 |
| Gm10073 | 0.363 | 0.345 | 0.396 |
| Plekho1 | 0.363 | 0.484 | 0.190 |
| Atg3 | 0.363 | 0.415 | 0.372 |
| Mfsd6 | 0.363 | 0.154 | 0.092 |
| Esyt2 | 0.363 | 0.573 | 0.348 |
| Sucla2 | 0.362 | 0.438 | 0.330 |
| Mapk1ip1l | 0.362 | 0.343 | 0.342 |
| Prpf8 | 0.362 | 0.297 | 0.302 |
| Mob1a | 0.362 | 0.285 | 0.281 |
| Exoc3l4 | 0.362 | 0.027 | 0.041 |
| Ehmt1 | 0.362 | 0.321 | 0.301 |
| Htatsf1 | 0.361 | 0.433 | 0.436 |
| Gcc2 | 0.361 | 0.211 | 0.257 |
| Gm26917 | 0.361 | 0.346 | 0.305 |
| Gosr2 | 0.361 | 0.184 | 0.228 |
| Fam193a | 0.361 | 0.366 | 0.369 |
| Nufip2 | 0.361 | 0.461 | 0.458 |
| Arhgef7 | 0.361 | 0.253 | 0.195 |
| Cib1 | 0.361 | 0.161 | 0.166 |
| Cpne8 | 0.360 | 0.031 | 0.084 |
| Hprt | 0.360 | 0.256 | 0.225 |
| Fto | 0.360 | 0.379 | 0.406 |
| Tmem123 | 0.360 | 0.414 | 0.232 |
| Atp2c1 | 0.360 | 0.170 | 0.183 |
| Tmem86a | 0.360 | 0.145 | 0.174 |
| Chmp6 | 0.360 | 0.216 | 0.236 |
| Rnf4 | 0.360 | 0.304 | 0.317 |
| Tor2a | 0.360 | 0.305 | 0.412 |

| | | | |
|---|---|---|---|
| Ppfibp1 | 0.360 | 0.319 | 0.298 |
| Clpp | 0.360 | 0.350 | 0.370 |
| Gm47283 | 0.359 | 0.453 | 0.374 |
| Hmbs | 0.359 | 0.279 | 0.283 |
| Piezo1 | 0.359 | 0.137 | 0.190 |
| Psip1 | 0.359 | 0.718 | 0.472 |
| Usp22 | 0.359 | 0.293 | 0.346 |
| Chpf2 | 0.358 | 0.238 | 0.299 |
| Pear1 | 0.358 | 0.157 | 0.193 |
| Col4a1 | 0.358 | 1.932 | 1.724 |
| Chsy1 | 0.358 | 0.315 | 0.290 |
| Thoc3 | 0.358 | 0.264 | 0.297 |
| Mtmr2 | 0.358 | 0.189 | 0.178 |
| Sertad2 | 0.358 | 0.432 | 0.369 |
| Taf11 | 0.357 | 0.357 | 0.326 |
| Srp68 | 0.357 | 0.272 | 0.332 |
| Alg5 | 0.357 | 0.393 | 0.413 |
| Luc7l | 0.357 | 0.311 | 0.299 |
| Erc1 | 0.357 | 0.340 | 0.270 |
| Egln1 | 0.357 | 0.434 | 0.439 |
| Tcerg1 | 0.357 | 0.370 | 0.323 |
| Dnajc21 | 0.357 | 0.339 | 0.313 |
| Cdk2ap2 | 0.357 | 0.315 | 0.251 |
| Pin4 | 0.356 | 0.315 | 0.281 |
| Lrrc42 | 0.356 | 0.318 | 0.364 |
| Supt6 | 0.356 | 0.338 | 0.304 |
| Prkch | 0.356 | 0.006 | 0.010 |
| 4833420G17l | 0.356 | 0.255 | 0.270 |
| Lap3 | 0.356 | 0.372 | 0.381 |
| Otulin | 0.356 | 0.222 | 0.231 |
| Tfpi | 0.355 | 0.162 | 0.373 |
| Tanc1 | 0.355 | 0.298 | 0.167 |
| Pdxdc1 | 0.355 | 0.368 | 0.422 |
| Ip6k1 | 0.355 | 0.365 | 0.341 |
| Gtf2e2 | 0.354 | 0.380 | 0.318 |
| Lrrc32 | 0.354 | 0.058 | 0.128 |
| Smim12 | 0.354 | 0.228 | 0.173 |
| Ap3d1 | 0.354 | 0.317 | 0.383 |
| Fry | 0.354 | 0.511 | 0.206 |
| Get4 | 0.354 | 0.331 | 0.293 |
| Ppp3cb | 0.354 | 0.394 | 0.388 |
| Zfp106 | 0.354 | 0.356 | 0.392 |
| Wwp2 | 0.354 | 0.767 | 0.215 |
| Apex1 | 0.354 | 0.354 | 0.329 |
| R3hdm2 | 0.354 | 0.343 | 0.326 |
| Gstm2 | 0.354 | 0.338 | 0.500 |
| Ino80b | 0.354 | 0.232 | 0.229 |
| 2510002D24l | 0.354 | 0.267 | 0.186 |

| | | | |
|---|---|---|---|
| Topors | 0.354 | 0.443 | 0.396 |
| Fam53a | 0.354 | 0.260 | 0.202 |
| Ppme1 | 0.353 | 0.279 | 0.159 |
| Pds5a | 0.353 | 0.320 | 0.283 |
| Sdc2 | 0.353 | 0.730 | 0.816 |
| Irak1 | 0.353 | 0.307 | 0.342 |
| Atxn1 | 0.353 | 0.306 | 0.245 |
| Safb2 | 0.353 | 0.337 | 0.355 |
| Rars | 0.353 | 0.343 | 0.328 |
| Mfap3 | 0.353 | 0.204 | 0.208 |
| Wdr89 | 0.353 | 0.343 | 0.360 |
| Zfp358 | 0.352 | 0.176 | 0.180 |
| Gcn1l1 | 0.352 | 0.270 | 0.279 |
| Nemf | 0.352 | 0.350 | 0.288 |
| Stx6 | 0.352 | 0.245 | 0.181 |
| Tob1 | 0.351 | 0.493 | 0.284 |
| Kitl | 0.351 | 0.203 | 0.199 |
| Chp1 | 0.351 | 0.379 | 0.293 |
| Mettl23 | 0.351 | 0.346 | 0.392 |
| Brix1 | 0.351 | 0.319 | 0.307 |
| Sh3gl1 | 0.351 | 0.251 | 0.352 |
| Pkdcc | 0.351 | 0.899 | 0.735 |
| Tmem115 | 0.351 | 0.257 | 0.268 |
| Synj2bp | 0.351 | 0.283 | 0.244 |
| Pon3 | 0.350 | 0.054 | 0.079 |
| Pcolce2 | 0.350 | 0.330 | 0.372 |
| Trim27 | 0.350 | 0.276 | 0.334 |
| Lbr | 0.350 | 0.206 | 0.155 |
| H2-Q4 | 0.350 | 0.088 | 0.090 |
| Gbf1 | 0.350 | 0.271 | 0.317 |
| Qdpr | 0.350 | 0.251 | 0.262 |
| AI314180 | 0.350 | 0.356 | 0.333 |
| Gmps | 0.350 | 0.375 | 0.353 |
| Cfap20 | 0.350 | 0.441 | 0.469 |
| Stag2 | 0.350 | 0.354 | 0.383 |
| Erf | 0.350 | 0.358 | 0.262 |
| Msrb3 | 0.349 | 0.516 | 0.214 |
| Gjc1 | 0.349 | 0.493 | 0.380 |
| Pop5 | 0.349 | 0.296 | 0.186 |
| Gtpbp2 | 0.349 | 0.238 | 0.195 |
| Unc13b | 0.349 | 0.023 | 0.027 |
| Dram2 | 0.349 | 0.198 | 0.222 |
| Hid1 | 0.349 | 0.042 | 0.018 |
| Zfp664 | 0.349 | 0.234 | 0.194 |
| Bag5 | 0.348 | 0.344 | 0.446 |
| Acadm | 0.348 | 0.613 | 0.413 |
| Itpr3 | 0.348 | 0.030 | 0.035 |
| Tsc22d2 | 0.348 | 0.431 | 0.382 |

| | | | |
|---|---|---|---|
| Efr3a | 0.348 | 0.424 | 0.371 |
| Zmynd8 | 0.348 | 0.252 | 0.303 |
| Dock9 | 0.348 | 0.055 | 0.082 |
| Zfp266 | 0.348 | 0.284 | 0.301 |
| Slu7 | 0.348 | 0.276 | 0.264 |
| Nab1 | 0.348 | 0.407 | 0.440 |
| Ogfr | 0.348 | 0.290 | 0.349 |
| Commd6 | 0.348 | 0.316 | 0.282 |
| Arfrp1 | 0.348 | 0.258 | 0.256 |
| Mum1 | 0.348 | 0.207 | 0.230 |
| Rab3gap1 | 0.348 | 0.273 | 0.230 |
| Sf3a3 | 0.347 | 0.310 | 0.300 |
| Shisa4 | 0.347 | 0.707 | 0.406 |
| Smim3 | 0.347 | 0.301 | 0.247 |
| Mpnd | 0.347 | 0.373 | 0.366 |
| Prrg2 | 0.347 | 0.103 | 0.121 |
| Ldlrad4 | 0.347 | 0.284 | 0.257 |
| Bhlhe40 | 0.347 | 0.672 | 0.470 |
| Dnajb1 | 0.347 | 0.460 | 0.594 |
| Zdhhc12 | 0.347 | 0.128 | 0.066 |
| Snx9 | 0.347 | 0.265 | 0.319 |
| Slbp | 0.347 | 0.409 | 0.435 |
| Tmem5 | 0.347 | 0.403 | 0.437 |
| Nagk | 0.347 | 0.301 | 0.385 |
| Tmcc1 | 0.346 | 0.391 | 0.365 |
| Nfu1 | 0.346 | 0.329 | 0.322 |
| Kdm6b | 0.346 | 0.588 | 0.543 |
| Urod | 0.345 | 0.272 | 0.254 |
| Abi3 | 0.345 | 0.009 | 0.007 |
| Dld | 0.345 | 0.392 | 0.317 |
| Ep400 | 0.345 | 0.315 | 0.311 |
| Lyar | 0.345 | 0.298 | 0.309 |
| BC003331 | 0.345 | 0.318 | 0.353 |
| Cdip1 | 0.345 | 0.329 | 0.338 |
| Klhl5 | 0.345 | 0.096 | 0.151 |
| Scrn1 | 0.344 | 0.091 | 0.161 |
| Zfp622 | 0.344 | 0.311 | 0.315 |
| Tmem141 | 0.344 | 0.195 | 0.095 |
| Aida | 0.344 | 0.241 | 0.369 |
| Nabp2 | 0.344 | 0.305 | 0.314 |
| Frg1 | 0.344 | 0.304 | 0.298 |
| Rnf20 | 0.344 | 0.285 | 0.274 |
| Kmt2c | 0.344 | 0.410 | 0.393 |
| Kctd2 | 0.344 | 0.209 | 0.246 |
| Api5 | 0.344 | 0.296 | 0.243 |
| Chchd3 | 0.344 | 0.301 | 0.268 |
| Ripor1 | 0.344 | 0.196 | 0.164 |
| Impdh1 | 0.344 | 0.140 | 0.170 |

| | | | |
|---|---|---|---|
| Hba-a1 | 0.344 | 0.324 | 0.366 |
| Eva1a | 0.344 | 0.040 | 0.018 |
| Lmbrd1 | 0.343 | 0.303 | 0.246 |
| Atf7ip | 0.343 | 0.270 | 0.288 |
| Ube4b | 0.343 | 0.404 | 0.373 |
| Tox4 | 0.343 | 0.311 | 0.275 |
| Rfc1 | 0.343 | 0.383 | 0.472 |
| Hdgfl3 | 0.343 | 0.432 | 0.440 |
| Cdc42se1 | 0.343 | 0.330 | 0.252 |
| Snx5 | 0.343 | 0.257 | 0.384 |
| Tnpo1 | 0.342 | 0.307 | 0.344 |
| Leng8 | 0.342 | 0.372 | 0.299 |
| Ptpn11 | 0.342 | 0.401 | 0.420 |
| Rnf6 | 0.342 | 0.345 | 0.325 |
| Cdc37l1 | 0.342 | 0.412 | 0.330 |
| Rgl1 | 0.342 | 0.039 | 0.282 |
| Zcchc14 | 0.342 | 0.298 | 0.222 |
| Cul5 | 0.342 | 0.389 | 0.379 |
| Selp | 0.342 | 0.006 | 0.002 |
| Dmac1 | 0.341 | 0.295 | 0.346 |
| Akirin2 | 0.341 | 0.359 | 0.363 |
| Usp14 | 0.341 | 0.370 | 0.287 |
| Cnpy3 | 0.341 | 0.254 | 0.288 |
| S100a4 | 0.341 | 0.247 | 0.181 |
| Dhrs7 | 0.341 | 0.345 | 0.475 |
| Ube2g1 | 0.341 | 0.295 | 0.229 |
| Ythdf1 | 0.341 | 0.314 | 0.347 |
| Ephb4 | 0.341 | 0.135 | 0.158 |
| Hnrnph3 | 0.340 | 0.364 | 0.349 |
| Lims2 | 0.340 | 0.204 | 0.104 |
| Ugp2 | 0.340 | 0.444 | 0.678 |
| Tns2 | 0.340 | 0.240 | 0.269 |
| Ephx1 | 0.340 | 0.181 | 0.174 |
| Aga | 0.340 | 0.273 | 0.313 |
| Limd2 | 0.340 | 0.220 | 0.240 |
| Mta1 | 0.340 | 0.335 | 0.320 |
| Mcmbp | 0.339 | 0.327 | 0.320 |
| Casp6 | 0.339 | 0.188 | 0.143 |
| Gtf3a | 0.339 | 0.290 | 0.344 |
| Raf1 | 0.339 | 0.389 | 0.289 |
| Nfatc1 | 0.339 | 0.132 | 0.122 |
| Hdac3 | 0.339 | 0.282 | 0.302 |
| Stoml2 | 0.339 | 0.360 | 0.378 |
| Smndc1 | 0.339 | 0.306 | 0.230 |
| Emd | 0.339 | 0.515 | 0.356 |
| Usp16 | 0.339 | 0.343 | 0.387 |
| Rere | 0.339 | 0.327 | 0.319 |
| Fndc3a | 0.338 | 0.381 | 0.389 |

| | | | |
|---|---|---|---|
| Smg6 | 0.338 | 0.311 | 0.277 |
| Arhgap17 | 0.338 | 0.269 | 0.257 |
| Far1 | 0.338 | 0.322 | 0.291 |
| 2310009B15R | 0.338 | 0.244 | 0.248 |
| Klhdc3 | 0.338 | 0.332 | 0.285 |
| Mrpl35 | 0.338 | 0.356 | 0.313 |
| Zswim8 | 0.338 | 0.376 | 0.350 |
| Riok1 | 0.338 | 0.294 | 0.289 |
| Kansl1 | 0.338 | 0.328 | 0.271 |
| Arfgef1 | 0.337 | 0.331 | 0.359 |
| Suds3 | 0.337 | 0.296 | 0.340 |
| Rnf130 | 0.337 | 0.307 | 0.337 |
| Gmds | 0.337 | 0.166 | 0.168 |
| 1500011K16R | 0.337 | 0.207 | 0.152 |
| Lrp6 | 0.337 | 0.476 | 0.510 |
| Borcs7 | 0.337 | 0.392 | 0.334 |
| Frmd4b | 0.337 | 0.307 | 0.098 |
| Faf2 | 0.337 | 0.352 | 0.377 |
| Rnf126 | 0.336 | 0.275 | 0.280 |
| Airn | 0.336 | 0.316 | 0.482 |
| Tspan13 | 0.336 | 0.016 | 0.020 |
| Myo18a | 0.336 | 0.284 | 0.161 |
| Arrb1 | 0.336 | 0.066 | 0.093 |
| Hipk3 | 0.336 | 0.341 | 0.314 |
| Mkrn1 | 0.336 | 0.274 | 0.223 |
| Gsk3a | 0.336 | 0.326 | 0.321 |
| Calml4 | 0.335 | 0.076 | 0.235 |
| Cdc34 | 0.335 | 0.187 | 0.176 |
| Hk1 | 0.335 | 0.193 | 0.182 |
| Bcr | 0.335 | 0.225 | 0.124 |
| Amigo2 | 0.335 | 0.158 | 0.061 |
| Mif4gd | 0.335 | 0.251 | 0.246 |
| Nt5c | 0.335 | 0.379 | 0.342 |
| Stip1 | 0.335 | 0.322 | 0.237 |
| Zfp280d | 0.335 | 0.330 | 0.274 |
| Mgst1 | 0.335 | 0.344 | 0.495 |
| Gdi1 | 0.335 | 0.308 | 0.286 |
| Mbtd1 | 0.335 | 0.326 | 0.347 |
| Col4a5 | 0.334 | 0.615 | 0.446 |
| Sft2d2 | 0.334 | 0.151 | 0.082 |
| Zkscan3 | 0.334 | 0.282 | 0.296 |
| Parvb | 0.334 | 0.006 | 0.012 |
| Naa10 | 0.334 | 0.316 | 0.311 |
| Sdhaf4 | 0.334 | 0.205 | 0.263 |
| Commd2 | 0.334 | 0.287 | 0.267 |
| Fcf1 | 0.334 | 0.289 | 0.270 |
| Casp3 | 0.333 | 0.265 | 0.233 |
| Srpx2 | 0.333 | 0.104 | 0.272 |

| | | | |
|---|---|---|---|
| Dnajb9 | 0.333 | 0.391 | 0.408 |
| Atf2 | 0.333 | 0.294 | 0.291 |
| Afg3l1 | 0.333 | 0.248 | 0.289 |
| Ikbkb | 0.333 | 0.176 | 0.187 |
| Ylpm1 | 0.333 | 0.310 | 0.305 |
| Snapin | 0.333 | 0.294 | 0.310 |
| Rilpl2 | 0.333 | 0.237 | 0.206 |
| Ccnt2 | 0.332 | 0.284 | 0.267 |
| Mfsd10 | 0.332 | 0.414 | 0.378 |
| 6330403K07F | 0.332 | 0.385 | 0.422 |
| Mphosph8 | 0.332 | 0.332 | 0.253 |
| Prex2 | 0.332 | 0.086 | 0.048 |
| Acox1 | 0.332 | 0.257 | 0.177 |
| Cs | 0.332 | 0.319 | 0.238 |
| Tial1 | 0.332 | 0.344 | 0.349 |
| Cat | 0.332 | 0.268 | 0.329 |
| Rab5b | 0.332 | 0.267 | 0.266 |
| Kpna4 | 0.331 | 0.354 | 0.304 |
| Ppt1 | 0.331 | 0.342 | 0.359 |
| Tmem222 | 0.331 | 0.287 | 0.282 |
| Spin1 | 0.331 | 0.294 | 0.273 |
| Tug1 | 0.331 | 0.297 | 0.372 |
| Dohh | 0.331 | 0.277 | 0.262 |
| Rnf14 | 0.331 | 0.283 | 0.285 |
| 2610001J05R | 0.330 | 0.317 | 0.326 |
| Armcx1 | 0.330 | 0.220 | 0.276 |
| Erbin | 0.330 | 0.399 | 0.355 |
| Amotl2 | 0.330 | 0.164 | 0.262 |
| Agfg1 | 0.330 | 0.236 | 0.288 |
| H2afx | 0.330 | 0.281 | 0.294 |
| Trappc6a | 0.330 | 0.200 | 0.242 |
| Stim2 | 0.330 | 0.148 | 0.212 |
| Gm1673 | 0.329 | 0.143 | 0.357 |
| Plxdc2 | 0.329 | 0.569 | 0.902 |
| Golga2 | 0.329 | 0.258 | 0.303 |
| Emsy | 0.329 | 0.338 | 0.334 |
| Slc35b3 | 0.329 | 0.151 | 0.130 |
| Prdx3 | 0.329 | 0.230 | 0.245 |
| Pim3 | 0.329 | 0.262 | 0.210 |
| Uso1 | 0.329 | 0.309 | 0.319 |
| Cnot2 | 0.328 | 0.304 | 0.299 |
| Dtx3 | 0.328 | 0.512 | 0.504 |
| Camk4 | 0.328 | 0.017 | 0.037 |
| Mysm1 | 0.328 | 0.337 | 0.328 |
| Derl2 | 0.328 | 0.277 | 0.318 |
| Commd9 | 0.328 | 0.185 | 0.242 |
| Rhoq | 0.328 | 0.522 | 0.540 |
| Aggf1 | 0.328 | 0.331 | 0.248 |

| | | | |
|---|---|---|---|
| Ptpn12 | 0.328 | 0.213 | 0.206 |
| Man1a | 0.327 | 0.145 | 0.218 |
| Slc48a1 | 0.327 | 0.415 | 0.235 |
| Fscn1 | 0.327 | 0.083 | 0.519 |
| Dnm2 | 0.327 | 0.216 | 0.240 |
| Tmem33 | 0.327 | 0.307 | 0.248 |
| Mau2 | 0.327 | 0.314 | 0.278 |
| Wdr43 | 0.327 | 0.307 | 0.298 |
| Cdk17 | 0.327 | 0.160 | 0.086 |
| 0610010K14F | 0.327 | 0.232 | 0.291 |
| Ckb | 0.327 | 1.455 | 0.934 |
| Pcyox1 | 0.327 | 0.225 | 0.245 |
| Mnat1 | 0.326 | 0.288 | 0.262 |
| Uri1 | 0.326 | 0.388 | 0.336 |
| Flot1 | 0.326 | 0.301 | 0.289 |
| Praf2 | 0.326 | 0.271 | 0.416 |
| Iah1 | 0.326 | 0.313 | 0.495 |
| Pabpn1 | 0.326 | 0.415 | 0.362 |
| Atp6v0a1 | 0.326 | 0.219 | 0.178 |
| Tlnrd1 | 0.326 | 0.117 | 0.190 |
| Abhd8 | 0.326 | 0.182 | 0.194 |
| Smad5 | 0.326 | 0.262 | 0.305 |
| Rnf149 | 0.326 | 0.342 | 0.288 |
| Bet1 | 0.326 | 0.277 | 0.334 |
| Fam91a1 | 0.326 | 0.260 | 0.252 |
| Smim15 | 0.325 | 0.300 | 0.355 |
| Igsf3 | 0.325 | 0.712 | 0.728 |
| Gon4l | 0.325 | 0.262 | 0.292 |
| Arhgap1 | 0.325 | 0.302 | 0.285 |
| Megf6 | 0.325 | 0.165 | 0.370 |
| Tbx1 | 0.325 | 0.012 | 0.021 |
| Abhd17b | 0.325 | 0.206 | 0.205 |
| Ttc1 | 0.325 | 0.260 | 0.243 |
| Elk4 | 0.325 | 0.230 | 0.220 |
| Sfswap | 0.325 | 0.358 | 0.304 |
| Med31 | 0.325 | 0.286 | 0.235 |
| Dhrs1 | 0.325 | 0.264 | 0.265 |
| Fam168b | 0.325 | 0.324 | 0.340 |
| 4933434E20F | 0.325 | 0.222 | 0.246 |
| Tomm70a | 0.325 | 0.266 | 0.258 |
| Tnks1bp1 | 0.324 | 0.303 | 0.311 |
| Notch4 | 0.324 | 0.029 | 0.022 |
| Ttc19 | 0.324 | 0.489 | 0.508 |
| Lifr | 0.324 | 0.228 | 0.293 |
| Sike1 | 0.324 | 0.262 | 0.243 |
| Dnajc15 | 0.324 | 0.239 | 0.187 |
| Cdk12 | 0.324 | 0.353 | 0.338 |
| Zcchc9 | 0.324 | 0.348 | 0.313 |

| | | | |
|---|---|---|---|
| Klf10 | 0.324 | 0.372 | 0.340 |
| Ccdc6 | 0.324 | 0.428 | 0.358 |
| Smap2 | 0.324 | 0.182 | 0.176 |
| Ddx39 | 0.324 | 0.140 | 0.126 |
| Cpt1a | 0.324 | 0.285 | 0.227 |
| Cebpzos | 0.324 | 0.171 | 0.226 |
| Igf2bp3 | 0.323 | 0.496 | 0.468 |
| Ankrd10 | 0.323 | 0.319 | 0.256 |
| Ssbp1 | 0.323 | 0.360 | 0.332 |
| Grsf1 | 0.323 | 0.264 | 0.255 |
| Stx4a | 0.323 | 0.362 | 0.407 |
| Rnaset2b | 0.323 | 0.185 | 0.150 |
| Exoc7 | 0.322 | 0.312 | 0.297 |
| Dnlz | 0.322 | 0.309 | 0.341 |
| Rictor | 0.322 | 0.310 | 0.257 |
| Tmem64 | 0.322 | 0.223 | 0.319 |
| Smarcc2 | 0.322 | 0.271 | 0.313 |
| Gga1 | 0.322 | 0.204 | 0.236 |
| Aamdc | 0.322 | 0.316 | 0.314 |
| Smim19 | 0.322 | 0.321 | 0.372 |
| Tmem44 | 0.322 | 0.057 | 0.088 |
| Prnp | 0.322 | 0.333 | 0.320 |
| Mrpl22 | 0.321 | 0.279 | 0.238 |
| Ormdl3 | 0.321 | 0.390 | 0.470 |
| Slc25a51 | 0.321 | 0.272 | 0.270 |
| Arf2 | 0.321 | 0.214 | 0.216 |
| Xpa | 0.321 | 0.341 | 0.322 |
| Igf2bp2 | 0.321 | 0.264 | 0.351 |
| Supt20 | 0.321 | 0.289 | 0.271 |
| Tal1 | 0.321 | 0.002 | 0.002 |
| Mon2 | 0.321 | 0.318 | 0.346 |
| Gnpda2 | 0.321 | 0.247 | 0.261 |
| 4-Sep | 0.321 | 0.083 | 0.041 |
| Mmadhc | 0.320 | 0.363 | 0.321 |
| Etfa | 0.320 | 0.378 | 0.350 |
| Inhba | 0.320 | 0.942 | 0.469 |
| Snrnp40 | 0.320 | 0.259 | 0.260 |
| Eif2s3y | 0.320 | 0.272 | 0.250 |
| Washc1 | 0.320 | 0.282 | 0.282 |
| Cox20 | 0.320 | 0.358 | 0.282 |
| Srprb | 0.320 | 0.223 | 0.245 |
| Bak1 | 0.320 | 0.233 | 0.217 |
| Strn4 | 0.320 | 0.315 | 0.298 |
| Jup | 0.319 | 0.259 | 0.134 |
| Rc3h1 | 0.319 | 0.263 | 0.235 |
| Fbxl3 | 0.319 | 0.291 | 0.255 |
| Rnaset2a | 0.319 | 0.205 | 0.164 |
| Larp4 | 0.319 | 0.319 | 0.249 |

| | | | |
|---|---|---|---|
| Ctgf | 0.319 | 1.069 | 0.368 |
| Mtx1 | 0.319 | 0.256 | 0.225 |
| Bcl2l1 | 0.319 | 0.254 | 0.204 |
| Kdsr | 0.319 | 0.260 | 0.213 |
| Myo1d | 0.319 | 0.226 | 0.091 |
| Klhl35 | 0.319 | 0.007 | 0.009 |
| Arhgef40 | 0.319 | 0.420 | 0.525 |
| Exosc7 | 0.319 | 0.210 | 0.246 |
| Rnf115 | 0.318 | 0.257 | 0.199 |
| Pik3c2a | 0.318 | 0.294 | 0.223 |
| Atg5 | 0.318 | 0.255 | 0.225 |
| Mosmo | 0.318 | 0.327 | 0.280 |
| Rsrc1 | 0.318 | 0.296 | 0.293 |
| Dnajc2 | 0.318 | 0.264 | 0.251 |
| Zfp467 | 0.318 | 0.261 | 0.139 |
| Flot2 | 0.318 | 0.329 | 0.316 |
| Snhg12 | 0.318 | 0.448 | 0.487 |
| Armc1 | 0.318 | 0.335 | 0.342 |
| Mib1 | 0.318 | 0.281 | 0.245 |
| Pmpcb | 0.318 | 0.341 | 0.276 |
| Ywhag | 0.318 | 0.348 | 0.310 |
| Mlec | 0.317 | 0.514 | 0.627 |
| Ppil4 | 0.317 | 0.287 | 0.235 |
| Nkap | 0.317 | 0.207 | 0.189 |
| Pck2 | 0.317 | 0.241 | 0.285 |
| Arfip2 | 0.317 | 0.279 | 0.370 |
| Mfsd1 | 0.317 | 0.208 | 0.180 |
| Chd1 | 0.317 | 0.317 | 0.252 |
| Bvht | 0.317 | 0.039 | 0.069 |
| Hfe | 0.317 | 0.043 | 0.075 |
| Svil | 0.316 | 0.519 | 0.792 |
| Lta4h | 0.316 | 0.246 | 0.251 |
| Fmnl3 | 0.316 | 0.207 | 0.129 |
| Dync1li2 | 0.316 | 0.262 | 0.314 |
| Oaf | 0.316 | 0.245 | 0.486 |
| Csnk1g1 | 0.316 | 0.294 | 0.276 |
| Cic | 0.316 | 0.395 | 0.407 |
| 2410002F23R | 0.316 | 0.270 | 0.311 |
| Uggt1 | 0.315 | 0.172 | 0.193 |
| Zzz3 | 0.315 | 0.275 | 0.282 |
| Nomo1 | 0.315 | 0.149 | 0.129 |
| Mrpl38 | 0.315 | 0.230 | 0.242 |
| Mrpl55 | 0.315 | 0.298 | 0.261 |
| Col6a3 | 0.315 | 0.863 | 1.859 |
| Tspyl1 | 0.315 | 0.362 | 0.311 |
| 2700081O15R | 0.315 | 0.297 | 0.415 |
| Mapk14 | 0.314 | 0.171 | 0.189 |
| Tex264 | 0.314 | 0.351 | 0.407 |

| | | | |
|---|---|---|---|
| Sh3pxd2a | 0.314 | 0.636 | 0.683 |
| Anxa1 | 0.314 | 0.611 | 0.670 |
| Psme4 | 0.314 | 0.330 | 0.300 |
| Aspscr1 | 0.314 | 0.316 | 0.291 |
| Kat6b | 0.314 | 0.361 | 0.371 |
| Eif2s3x | 0.314 | 0.313 | 0.210 |
| Prkra | 0.314 | 0.296 | 0.277 |
| Micu1 | 0.314 | 0.140 | 0.199 |
| Aebp2 | 0.314 | 0.319 | 0.276 |
| Sipa1 | 0.313 | 0.118 | 0.178 |
| Crtc3 | 0.313 | 0.614 | 0.495 |
| Sbf2 | 0.313 | 0.326 | 0.354 |
| Pfdn4 | 0.313 | 0.329 | 0.260 |
| Pip5k1a | 0.313 | 0.301 | 0.251 |
| Kcnn4 | 0.313 | 0.003 | 0.004 |
| Tmem55a | 0.313 | 0.321 | 0.270 |
| Ggta1 | 0.313 | 0.179 | 0.170 |
| Dmwd | 0.313 | 0.317 | 0.277 |
| Gbp7 | 0.312 | 0.059 | 0.114 |
| Pmvk | 0.312 | 0.259 | 0.182 |
| Pard3 | 0.312 | 0.287 | 0.341 |
| Cox19 | 0.312 | 0.301 | 0.250 |
| Mllt10 | 0.312 | 0.301 | 0.277 |
| Ccdc107 | 0.312 | 0.755 | 0.335 |
| Trio | 0.312 | 0.453 | 0.679 |
| Sirpa | 0.312 | 0.087 | 0.177 |
| 6-Sep | 0.312 | 0.221 | 0.158 |
| Cdc123 | 0.312 | 0.296 | 0.328 |
| Atp11a | 0.312 | 0.128 | 0.175 |
| Pdlim2 | 0.312 | 0.393 | 0.926 |
| Ifrd1 | 0.311 | 0.619 | 0.546 |
| Gatad2a | 0.311 | 0.218 | 0.253 |
| Ankrd40 | 0.311 | 0.266 | 0.285 |
| Spty2d1 | 0.311 | 0.298 | 0.361 |
| Diaph1 | 0.311 | 0.192 | 0.192 |
| Phf21a | 0.311 | 0.301 | 0.316 |
| Ralgapa1 | 0.311 | 0.260 | 0.220 |
| Smim27 | 0.311 | 0.235 | 0.288 |
| 4833439L19F | 0.311 | 0.267 | 0.212 |
| Tssc4 | 0.311 | 0.286 | 0.277 |
| Slc39a13 | 0.311 | 0.258 | 0.404 |
| Smarcc1 | 0.311 | 0.421 | 0.432 |
| Mrto4 | 0.311 | 0.296 | 0.266 |
| Mcur1 | 0.311 | 0.272 | 0.197 |
| Tmem245 | 0.311 | 0.290 | 0.230 |
| Ufl1 | 0.311 | 0.296 | 0.287 |
| Ufsp2 | 0.311 | 0.282 | 0.338 |
| Zfp787 | 0.311 | 0.238 | 0.201 |

| | | | |
|---|---|---|---|
| Mpped2 | 0.311 | 0.464 | 0.287 |
| Mia2 | 0.311 | 0.358 | 0.331 |
| Prrt4 | 0.311 | 0.002 | 0.006 |
| Osbpl5 | 0.311 | 0.225 | 0.256 |
| Sigmar1 | 0.310 | 0.201 | 0.276 |
| B3gat3 | 0.310 | 0.267 | 0.331 |
| Eif2b5 | 0.310 | 0.222 | 0.219 |
| Atxn2 | 0.310 | 0.363 | 0.322 |
| Pdzd8 | 0.310 | 0.259 | 0.214 |
| Pop7 | 0.309 | 0.213 | 0.179 |
| Arl8b | 0.309 | 0.324 | 0.296 |
| Fastk | 0.309 | 0.271 | 0.334 |
| Snrnp200 | 0.309 | 0.256 | 0.239 |
| Trmt1 | 0.309 | 0.278 | 0.258 |
| Gnl1 | 0.309 | 0.342 | 0.390 |
| Asap1 | 0.309 | 0.417 | 0.473 |
| Cdc40 | 0.309 | 0.321 | 0.273 |
| Rae1 | 0.309 | 0.301 | 0.279 |
| Nr4a1 | 0.309 | 1.482 | 0.955 |
| Cbl | 0.309 | 0.250 | 0.181 |
| Arid4a | 0.309 | 0.365 | 0.312 |
| Nbea | 0.309 | 0.276 | 0.260 |
| Gpat4 | 0.308 | 0.180 | 0.247 |
| Nudt9 | 0.308 | 0.322 | 0.273 |
| Cdk13 | 0.308 | 0.299 | 0.287 |
| Sdc3 | 0.308 | 0.240 | 0.185 |
| 2510039O18I | 0.308 | 0.213 | 0.176 |
| Fgl2 | 0.308 | 0.769 | 0.378 |
| Ppp3r1 | 0.308 | 0.290 | 0.253 |
| Mzt1 | 0.308 | 0.254 | 0.236 |
| Igfbp3 | 0.308 | 0.063 | 0.402 |
| Cul4a | 0.308 | 0.254 | 0.245 |
| Zmpste24 | 0.308 | 0.232 | 0.187 |
| Mcee | 0.308 | 0.298 | 0.284 |
| Adam17 | 0.308 | 0.119 | 0.147 |
| Slc38a1 | 0.308 | 0.546 | 0.198 |
| Gmppb | 0.308 | 0.171 | 0.258 |
| Ugcg | 0.308 | 0.154 | 0.140 |
| Tgs1 | 0.308 | 0.197 | 0.205 |
| Ctnnbl1 | 0.307 | 0.211 | 0.223 |
| Card10 | 0.307 | 0.009 | 0.042 |
| Anpep | 0.307 | 0.022 | 0.353 |
| Exosc5 | 0.307 | 0.175 | 0.199 |
| Pcbd2 | 0.307 | 0.250 | 0.292 |
| Kansl2 | 0.307 | 0.236 | 0.225 |
| Pbx2 | 0.307 | 0.270 | 0.245 |
| Inpp5k | 0.307 | 0.220 | 0.158 |
| Ube3c | 0.307 | 0.197 | 0.215 |

| | | | |
|---|---|---|---|
| Polr3d | 0.307 | 0.248 | 0.350 |
| Rcor1 | 0.307 | 0.240 | 0.223 |
| Clk4 | 0.307 | 0.358 | 0.294 |
| Smad1 | 0.306 | 0.361 | 0.437 |
| Samd4b | 0.306 | 0.324 | 0.403 |
| Rapgef1 | 0.306 | 0.142 | 0.176 |
| Hivep2 | 0.306 | 0.336 | 0.445 |
| Sgce | 0.306 | 0.350 | 0.535 |
| Rabggtb | 0.306 | 0.267 | 0.202 |
| Rspry1 | 0.306 | 0.283 | 0.296 |
| Zfp330 | 0.306 | 0.317 | 0.321 |
| Gtf2b | 0.306 | 0.322 | 0.297 |
| Vps41 | 0.306 | 0.256 | 0.291 |
| Nrk | 0.306 | 0.280 | 1.114 |
| Ier5 | 0.306 | 0.716 | 0.530 |
| Med4 | 0.306 | 0.266 | 0.229 |
| Tmem132a | 0.305 | 0.541 | 0.262 |
| Ddx18 | 0.305 | 0.268 | 0.268 |
| M6pr | 0.305 | 0.259 | 0.208 |
| Lgals8 | 0.305 | 0.197 | 0.181 |
| Exoc3 | 0.305 | 0.337 | 0.358 |
| Lemd2 | 0.305 | 0.321 | 0.279 |
| Zfp871 | 0.305 | 0.248 | 0.254 |
| Chn1 | 0.305 | 0.115 | 0.063 |
| Marf1 | 0.305 | 0.282 | 0.254 |
| Fas | 0.305 | 0.138 | 0.145 |
| Tacc2 | 0.305 | 0.435 | 0.338 |
| Rpp25l | 0.305 | 0.199 | 0.201 |
| Tsnax | 0.305 | 0.283 | 0.294 |
| Rnpep | 0.305 | 0.263 | 0.210 |
| Cd59a | 0.305 | 0.125 | 0.063 |
| Xrn1 | 0.305 | 0.310 | 0.337 |
| Ube2e2 | 0.305 | 0.292 | 0.445 |
| Akr7a5 | 0.304 | 0.274 | 0.331 |
| Rab28 | 0.304 | 0.317 | 0.327 |
| Faap20 | 0.304 | 0.241 | 0.243 |
| Gm2000 | 0.304 | 0.268 | 0.329 |
| Akirin1 | 0.304 | 0.194 | 0.201 |
| Mpg | 0.304 | 0.225 | 0.294 |
| Crispld1 | 0.304 | 0.402 | 0.147 |
| Pdgfb | 0.304 | 0.014 | 0.004 |
| Eef2k | 0.304 | 0.586 | 0.378 |
| Pcdh7 | 0.304 | 1.137 | 0.803 |
| Srrt | 0.304 | 0.316 | 0.310 |
| Med13 | 0.304 | 0.378 | 0.345 |
| Nr2c2 | 0.304 | 0.277 | 0.246 |
| Klk8 | 0.304 | 0.012 | 0.025 |
| Gfpt1 | 0.304 | 0.257 | 0.364 |

| | | | |
|---|---|---|---|
| Hspb11 | 0.303 | 0.385 | 0.337 |
| Pgrmc2 | 0.303 | 0.281 | 0.264 |
| Stox2 | 0.303 | 0.247 | 0.239 |
| Mrps28 | 0.303 | 0.261 | 0.247 |
| Naa20 | 0.303 | 0.339 | 0.332 |
| 2810403A07F | 0.303 | 0.248 | 0.197 |
| Anapc1 | 0.303 | 0.223 | 0.237 |
| Naa35 | 0.303 | 0.272 | 0.282 |
| Fam50a | 0.303 | 0.279 | 0.268 |
| Dnajc25 | 0.302 | 0.317 | 0.411 |
| Ensa | 0.302 | 0.285 | 0.302 |
| Nosip | 0.302 | 0.208 | 0.188 |
| Sdhaf1 | 0.302 | 0.209 | 0.216 |
| Klhl13 | 0.302 | 0.171 | 0.454 |
| Spats2 | 0.302 | 0.152 | 0.198 |
| Surf1 | 0.302 | 0.269 | 0.281 |
| Hddc2 | 0.301 | 0.225 | 0.234 |
| Wipf3 | 0.301 | 0.002 | 0.009 |
| Usp1 | 0.301 | 0.248 | 0.271 |
| Oser1 | 0.301 | 0.238 | 0.239 |
| Ptprk | 0.301 | 0.318 | 0.302 |
| Ypel2 | 0.301 | 0.278 | 0.094 |
| Poldip2 | 0.301 | 0.333 | 0.378 |
| Slc35a4 | 0.301 | 0.222 | 0.233 |
| Gns | 0.301 | 0.248 | 0.349 |
| Dhx9 | 0.301 | 0.330 | 0.304 |
| Ncln | 0.300 | 0.250 | 0.326 |
| Hagh | 0.300 | 0.236 | 0.155 |
| Fbxo11 | 0.300 | 0.311 | 0.304 |
| Ccdc127 | 0.300 | 0.283 | 0.334 |
| Zfp637 | 0.300 | 0.364 | 0.348 |
| Ist1 | 0.300 | 0.235 | 0.216 |
| Zbed3 | 0.300 | 0.249 | 0.217 |
| Dmd | 0.300 | 0.389 | 0.155 |
| Gpaa1 | 0.300 | 0.188 | 0.210 |
| Exoc5 | 0.300 | 0.310 | 0.309 |
| Josd2 | 0.300 | 0.346 | 0.355 |
| Rpp21 | 0.300 | 0.269 | 0.287 |
| Smarcd1 | 0.300 | 0.259 | 0.192 |
| Cisd1 | 0.300 | 0.295 | 0.173 |
| Rab35 | 0.299 | 0.246 | 0.238 |
| Gtf3c2 | 0.299 | 0.287 | 0.302 |
| Tmtc2 | 0.299 | 0.259 | 0.127 |
| Vma21 | 0.299 | 0.289 | 0.310 |
| Washc2 | 0.299 | 0.300 | 0.339 |
| Ergic1 | 0.299 | 0.296 | 0.415 |
| Qrich1 | 0.299 | 0.334 | 0.294 |
| Rbm4b | 0.299 | 0.454 | 0.433 |

| | | | |
|---|---|---|---|
| Zfp560 | 0.299 | 0.269 | 0.245 |
| Ykt6 | 0.299 | 0.204 | 0.227 |
| Por | 0.299 | 0.581 | 0.405 |
| Sox11 | 0.298 | 0.089 | 0.109 |
| Rab9 | 0.298 | 0.383 | 0.397 |
| Snap47 | 0.298 | 0.241 | 0.311 |
| Isca2 | 0.298 | 0.239 | 0.213 |
| B4galt4 | 0.298 | 0.079 | 0.098 |
| Gmfb | 0.298 | 0.166 | 0.266 |
| Pex2 | 0.298 | 0.215 | 0.200 |
| Stard4 | 0.298 | 0.066 | 0.102 |
| Tmem63b | 0.298 | 0.237 | 0.180 |
| Pde6d | 0.298 | 0.235 | 0.288 |
| Cnppd1 | 0.298 | 0.220 | 0.223 |
| Trim8 | 0.298 | 0.216 | 0.215 |
| Taf15 | 0.298 | 0.273 | 0.267 |
| Acly | 0.298 | 0.230 | 0.258 |
| Brwd1 | 0.298 | 0.289 | 0.273 |
| Abca1 | 0.298 | 0.197 | 0.285 |
| Rtl8b | 0.297 | 0.269 | 0.285 |
| Cmtm6 | 0.297 | 0.177 | 0.132 |
| Rbpj | 0.297 | 0.389 | 0.522 |
| Hspa14 | 0.297 | 0.239 | 0.228 |
| Ube2j1 | 0.297 | 0.266 | 0.294 |
| Spns1 | 0.297 | 0.247 | 0.266 |
| Dpp3 | 0.297 | 0.239 | 0.258 |
| Impa1 | 0.297 | 0.462 | 0.333 |
| Ldb1 | 0.296 | 0.286 | 0.233 |
| Med1 | 0.296 | 0.288 | 0.229 |
| Cnot7 | 0.296 | 0.242 | 0.205 |
| Urm1 | 0.296 | 0.229 | 0.232 |
| Minpp1 | 0.296 | 0.238 | 0.229 |
| Eci1 | 0.296 | 0.262 | 0.228 |
| Erg28 | 0.296 | 0.295 | 0.427 |
| Tpst2 | 0.296 | 0.183 | 0.229 |
| Ppp2r5a | 0.296 | 0.242 | 0.273 |
| Lsm3 | 0.296 | 0.372 | 0.384 |
| Rbm18 | 0.295 | 0.337 | 0.309 |
| Xpr1 | 0.295 | 0.293 | 0.168 |
| Creb1 | 0.295 | 0.222 | 0.233 |
| Slc31a2 | 0.295 | 0.141 | 0.201 |
| Mta2 | 0.295 | 0.248 | 0.221 |
| Shoc2 | 0.295 | 0.199 | 0.193 |
| Hibadh | 0.295 | 0.314 | 0.282 |
| Bccip | 0.295 | 0.351 | 0.330 |
| Sft2d1 | 0.295 | 0.148 | 0.154 |
| Actg2 | 0.295 | 1.678 | 0.444 |
| Hint3 | 0.295 | 0.196 | 0.191 |

| | | | |
|---|---|---|---|
| Stau1 | 0.295 | 0.277 | 0.261 |
| Mef2d | 0.295 | 0.506 | 0.253 |
| Ccdc25 | 0.295 | 0.190 | 0.237 |
| Ltv1 | 0.295 | 0.199 | 0.273 |
| Hars | 0.294 | 0.276 | 0.268 |
| Pcf11 | 0.294 | 0.397 | 0.395 |
| Cds2 | 0.294 | 0.170 | 0.111 |
| Alkbh1 | 0.294 | 0.244 | 0.260 |
| Gpx8 | 0.294 | 0.729 | 1.043 |
| Acads | 0.294 | 0.288 | 0.214 |
| Msrb1 | 0.294 | 0.446 | 0.339 |
| Hspa4l | 0.294 | 0.078 | 0.040 |
| Gmpr2 | 0.294 | 0.182 | 0.154 |
| Usp53 | 0.294 | 0.273 | 0.134 |
| Mafg | 0.294 | 0.318 | 0.347 |
| Tmem126a | 0.294 | 0.350 | 0.351 |
| Sf3b4 | 0.294 | 0.298 | 0.252 |
| Ncaph2 | 0.294 | 0.225 | 0.278 |
| Cryzl1 | 0.294 | 0.259 | 0.240 |
| Tmem11 | 0.294 | 0.310 | 0.305 |
| Slc25a1 | 0.293 | 0.212 | 0.200 |
| Caskin2 | 0.293 | 0.094 | 0.065 |
| Prkd3 | 0.293 | 0.270 | 0.360 |
| Atp6v1c1 | 0.293 | 0.216 | 0.185 |
| Larp1 | 0.293 | 0.346 | 0.306 |
| Ino80d | 0.293 | 0.281 | 0.275 |
| B230118H07I | 0.293 | 0.220 | 0.200 |
| Nop56 | 0.293 | 0.339 | 0.325 |
| Paip1 | 0.292 | 0.265 | 0.205 |
| Itgb5 | 0.292 | 0.872 | 0.919 |
| Loxl1 | 0.292 | 1.261 | 1.719 |
| Phpt1 | 0.292 | 0.257 | 0.262 |
| 2410004B18F | 0.292 | 0.185 | 0.218 |
| Nrarp | 0.292 | 0.091 | 0.020 |
| Rgs19 | 0.292 | 0.119 | 0.150 |
| Nkiras2 | 0.292 | 0.209 | 0.181 |
| Acbd5 | 0.291 | 0.279 | 0.235 |
| Prkaca | 0.291 | 0.325 | 0.291 |
| Sec22a | 0.291 | 0.226 | 0.205 |
| Ralgapa2 | 0.291 | 0.105 | 0.046 |
| Crem | 0.291 | 0.260 | 0.164 |
| Cnot4 | 0.291 | 0.329 | 0.310 |
| Fam192a | 0.291 | 0.243 | 0.221 |
| Fut11 | 0.291 | 0.233 | 0.340 |
| Arfgap3 | 0.291 | 0.260 | 0.339 |
| Snhg6 | 0.291 | 0.385 | 0.330 |
| Hsd17b4 | 0.290 | 0.307 | 0.229 |
| Tecpr1 | 0.290 | 0.162 | 0.155 |

| | | | |
|---|---|---|---|
| Ogfod3 | 0.290 | 0.127 | 0.150 |
| Lnpep | 0.290 | 0.342 | 0.299 |
| Rapgef2 | 0.290 | 0.144 | 0.151 |
| Ppp1r18 | 0.290 | 0.319 | 0.241 |
| Ndfip1 | 0.289 | 0.221 | 0.198 |
| Nt5c3b | 0.289 | 0.312 | 0.320 |
| Med15 | 0.289 | 0.242 | 0.208 |
| Atf7 | 0.289 | 0.367 | 0.351 |
| Ecm2 | 0.289 | 0.202 | 0.276 |
| Hes6 | 0.289 | 0.218 | 0.131 |
| Bscl2 | 0.289 | 0.247 | 0.248 |
| Nipal2 | 0.289 | 0.119 | 0.100 |
| Naga | 0.289 | 0.201 | 0.240 |
| Swt1 | 0.289 | 0.335 | 0.256 |
| Nifk | 0.289 | 0.227 | 0.240 |
| Gpr108 | 0.289 | 0.257 | 0.259 |
| Prelid3b | 0.289 | 0.245 | 0.256 |
| Ruvbl1 | 0.288 | 0.319 | 0.303 |
| St6gal1 | 0.288 | 0.150 | 0.073 |
| Crebrf | 0.288 | 0.311 | 0.268 |
| Bzw2 | 0.288 | 0.313 | 0.313 |
| Cant1 | 0.288 | 0.184 | 0.209 |
| Peli2 | 0.288 | 0.123 | 0.160 |
| Plekhj1 | 0.288 | 0.293 | 0.408 |
| Polr2l | 0.288 | 0.216 | 0.186 |
| Celf2 | 0.288 | 0.690 | 1.284 |
| Map3k3 | 0.287 | 0.167 | 0.210 |
| Jmjd6 | 0.287 | 0.329 | 0.298 |
| Arhgap28 | 0.287 | 0.036 | 0.243 |
| Fam208a | 0.287 | 0.296 | 0.292 |
| Rhog | 0.287 | 0.203 | 0.243 |
| Ppp1r10 | 0.287 | 0.430 | 0.455 |
| Pex19 | 0.287 | 0.228 | 0.224 |
| Gnl3 | 0.287 | 0.398 | 0.443 |
| Vamp4 | 0.287 | 0.201 | 0.261 |
| Yap1 | 0.287 | 0.350 | 0.265 |
| Dchs1 | 0.286 | 0.290 | 0.350 |
| Ap1m1 | 0.286 | 0.249 | 0.231 |
| Faf1 | 0.286 | 0.264 | 0.214 |
| Kif2a | 0.286 | 0.308 | 0.339 |
| Gpbp1l1 | 0.286 | 0.237 | 0.164 |
| Slc16a3 | 0.286 | 0.086 | 0.160 |
| Zfp445 | 0.286 | 0.274 | 0.297 |
| Rfxap | 0.286 | 0.250 | 0.225 |
| Gdpd5 | 0.286 | 0.056 | 0.041 |
| Szrd1 | 0.286 | 0.257 | 0.262 |
| Cdk7 | 0.286 | 0.243 | 0.281 |
| Trafd1 | 0.286 | 0.585 | 0.529 |

| | | | |
|---|---|---|---|
| 2810004N23I | 0.286 | 0.259 | 0.253 |
| 4930453N24I | 0.286 | 0.287 | 0.282 |
| Timm50 | 0.286 | 0.223 | 0.248 |
| Eif1ad | 0.286 | 0.189 | 0.193 |
| Ahctf1 | 0.286 | 0.193 | 0.205 |
| Golm1 | 0.286 | 0.032 | 0.126 |
| Ak3 | 0.286 | 0.432 | 0.392 |
| Yes1 | 0.286 | 0.177 | 0.128 |
| 2610507B11I | 0.286 | 0.227 | 0.226 |
| Chtf8 | 0.286 | 0.219 | 0.241 |
| Kbtbd2 | 0.286 | 0.162 | 0.157 |
| Nudt3 | 0.285 | 0.219 | 0.207 |
| Asna1 | 0.285 | 0.235 | 0.202 |
| Borcs8 | 0.285 | 0.287 | 0.226 |
| Csk | 0.285 | 0.335 | 0.336 |
| Rbmx | 0.285 | 0.202 | 0.200 |
| Preb | 0.285 | 0.168 | 0.183 |
| Psmg4 | 0.285 | 0.313 | 0.259 |
| Fez2 | 0.285 | 0.182 | 0.201 |
| Sox13 | 0.285 | 0.024 | 0.022 |
| Max | 0.285 | 0.318 | 0.282 |
| Sncaip | 0.285 | 0.170 | 0.209 |
| Spen | 0.285 | 0.340 | 0.247 |
| Thap11 | 0.284 | 0.216 | 0.239 |
| Atf1 | 0.284 | 0.338 | 0.309 |
| Atp13a2 | 0.284 | 0.128 | 0.133 |
| Cnot8 | 0.284 | 0.200 | 0.204 |
| Nek6 | 0.284 | 0.147 | 0.196 |
| Wdr18 | 0.284 | 0.240 | 0.219 |
| Ccdc34 | 0.284 | 0.345 | 0.422 |
| Appl1 | 0.284 | 0.363 | 0.403 |
| Dnajb14 | 0.284 | 0.290 | 0.294 |
| Ascc3 | 0.284 | 0.288 | 0.272 |
| Nme2 | 0.284 | 0.186 | 0.230 |
| Clock | 0.284 | 0.350 | 0.292 |
| Nadk | 0.284 | 0.214 | 0.237 |
| Zfp68 | 0.284 | 0.265 | 0.288 |
| Grk6 | 0.284 | 0.191 | 0.205 |
| Tor1aip2 | 0.284 | 0.278 | 0.262 |
| Ireb2 | 0.283 | 0.253 | 0.247 |
| Scmh1 | 0.283 | 0.229 | 0.241 |
| Carhsp1 | 0.283 | 0.200 | 0.329 |
| Zzef1 | 0.283 | 0.225 | 0.222 |
| 4932438A13I | 0.283 | 0.288 | 0.265 |
| St3gal2 | 0.283 | 0.292 | 0.293 |
| Fam204a | 0.283 | 0.283 | 0.239 |
| Tsta3 | 0.283 | 0.204 | 0.209 |
| Cd40 | 0.283 | 0.003 | 0.011 |

| | | | |
|---|---|---|---|
| Kars | 0.283 | 0.231 | 0.256 |
| Yipf6 | 0.283 | 0.147 | 0.147 |
| Mrps10 | 0.283 | 0.219 | 0.234 |
| Ssbp2 | 0.283 | 0.724 | 0.698 |
| Dut | 0.283 | 0.289 | 0.318 |
| Hspa13 | 0.283 | 0.351 | 0.334 |
| 1110059E24F | 0.283 | 0.249 | 0.242 |
| Tmtc4 | 0.283 | 0.079 | 0.063 |
| Zfp516 | 0.283 | 0.310 | 0.308 |
| Mrpl41 | 0.282 | 0.340 | 0.297 |
| Tmem9b | 0.282 | 0.184 | 0.204 |
| Pbdc1 | 0.282 | 0.250 | 0.271 |
| Narf | 0.282 | 0.151 | 0.184 |
| Arl5a | 0.282 | 0.247 | 0.281 |
| Mrpl3 | 0.282 | 0.236 | 0.234 |
| Trabd | 0.282 | 0.231 | 0.225 |
| Mapkap1 | 0.282 | 0.247 | 0.250 |
| Mpp1 | 0.282 | 0.196 | 0.209 |
| Itgav | 0.282 | 0.491 | 0.429 |
| Gigyf2 | 0.282 | 0.283 | 0.276 |
| Ubac2 | 0.282 | 0.226 | 0.222 |
| Ap3b1 | 0.282 | 0.300 | 0.364 |
| Vps72 | 0.282 | 0.272 | 0.245 |
| Psen2 | 0.282 | 0.201 | 0.223 |
| Ube2w | 0.281 | 0.312 | 0.307 |
| Ripk1 | 0.281 | 0.441 | 0.551 |
| Ankhd1 | 0.281 | 0.295 | 0.265 |
| Asxl2 | 0.281 | 0.233 | 0.197 |
| Pik3r1 | 0.281 | 0.466 | 0.511 |
| Lypla2 | 0.281 | 0.125 | 0.137 |
| Stk38 | 0.281 | 0.225 | 0.387 |
| Cryab | 0.281 | 1.038 | 0.739 |
| Ncoa7 | 0.281 | 0.338 | 0.303 |
| Ptrhd1 | 0.281 | 0.209 | 0.207 |
| Hmgcl | 0.281 | 0.173 | 0.218 |
| Rfc2 | 0.281 | 0.206 | 0.280 |
| Ppp1r7 | 0.281 | 0.232 | 0.226 |
| Setbp1 | 0.281 | 0.380 | 0.359 |
| Sfxn1 | 0.281 | 0.257 | 0.481 |
| Fam212a | 0.280 | 0.100 | 0.132 |
| Pole3 | 0.280 | 0.193 | 0.229 |
| Gar1 | 0.280 | 0.198 | 0.216 |
| Ccdc32 | 0.280 | 0.196 | 0.207 |
| Hikeshi | 0.280 | 0.290 | 0.339 |
| Yae1d1 | 0.280 | 0.325 | 0.232 |
| Col6a1 | 0.280 | 1.809 | 2.434 |
| Lrrc41 | 0.280 | 0.235 | 0.258 |
| Mrpl40 | 0.280 | 0.181 | 0.235 |

| | | | |
|---|---|---|---|
| Rnf167 | 0.280 | 0.332 | 0.307 |
| Asf1a | 0.280 | 0.287 | 0.231 |
| Apopt1 | 0.280 | 0.257 | 0.256 |
| Lzts2 | 0.280 | 0.552 | 0.481 |
| Pipox | 0.280 | 0.054 | 0.009 |
| Tmem219 | 0.280 | 0.280 | 0.349 |
| Gas2l1 | 0.280 | 0.257 | 0.243 |
| Ndufaf3 | 0.279 | 0.189 | 0.149 |
| Sdcbp2 | 0.279 | 0.012 | 0.008 |
| Rcsd1 | 0.279 | 0.512 | 0.280 |
| Limd1 | 0.279 | 0.465 | 0.434 |
| Nipa2 | 0.279 | 0.327 | 0.285 |
| Abhd14a | 0.279 | 0.306 | 0.397 |
| Lyrm2 | 0.279 | 0.179 | 0.200 |
| Nus1 | 0.279 | 0.217 | 0.185 |
| Camsap2 | 0.279 | 0.199 | 0.167 |
| Adgrl1 | 0.279 | 0.458 | 0.356 |
| Prkag2 | 0.279 | 0.175 | 0.104 |
| Wdr82 | 0.279 | 0.205 | 0.227 |
| Cuedc1 | 0.278 | 0.409 | 0.411 |
| Azi2 | 0.278 | 0.274 | 0.249 |
| Atxn2l | 0.278 | 0.298 | 0.253 |
| Jmy | 0.278 | 0.265 | 0.197 |
| Ubxn2a | 0.278 | 0.169 | 0.150 |
| Trove2 | 0.278 | 0.233 | 0.211 |
| D5Ertd579e | 0.278 | 0.217 | 0.204 |
| Acadsb | 0.278 | 0.317 | 0.344 |
| Cacfd1 | 0.278 | 0.230 | 0.269 |
| Yrdc | 0.278 | 0.224 | 0.141 |
| Sec23ip | 0.278 | 0.179 | 0.211 |
| Plekhg5 | 0.278 | 0.115 | 0.064 |
| Akap12 | 0.278 | 0.466 | 1.691 |
| Lmf2 | 0.278 | 0.199 | 0.214 |
| Cpsf2 | 0.278 | 0.228 | 0.228 |
| Chordc1 | 0.277 | 0.195 | 0.190 |
| Ddx27 | 0.277 | 0.253 | 0.211 |
| Zfp277 | 0.277 | 0.179 | 0.211 |
| 1110059G10I | 0.277 | 0.235 | 0.203 |
| Gyg | 0.277 | 0.320 | 0.226 |
| Diablo | 0.277 | 0.312 | 0.299 |
| Hsd17b12 | 0.277 | 0.351 | 0.307 |
| Ubp1 | 0.277 | 0.276 | 0.237 |
| Cxxc1 | 0.277 | 0.287 | 0.325 |
| Tmem60 | 0.277 | 0.257 | 0.263 |
| Ddx21 | 0.277 | 0.238 | 0.238 |
| Nae1 | 0.277 | 0.351 | 0.294 |
| Pdgfrb | 0.277 | 0.660 | 0.588 |
| Bloc1s5 | 0.277 | 0.086 | 0.086 |

| | | | |
|---|---|---|---|
| Cmtm8 | 0.276 | 0.029 | 0.017 |
| Pkd2 | 0.276 | 0.402 | 0.487 |
| Nol12 | 0.276 | 0.255 | 0.321 |
| Tarbp2 | 0.276 | 0.189 | 0.193 |
| Srek1ip1 | 0.276 | 0.260 | 0.206 |
| Alas1 | 0.276 | 0.156 | 0.162 |
| Gimap5 | 0.276 | 0.002 | 0.003 |
| Ivns1abp | 0.276 | 0.155 | 0.170 |
| Agpat5 | 0.276 | 0.335 | 0.241 |
| Slc44a1 | 0.276 | 0.159 | 0.280 |
| Isca1 | 0.276 | 0.163 | 0.206 |
| Pan3 | 0.276 | 0.238 | 0.216 |
| Cfap36 | 0.276 | 0.289 | 0.251 |
| Ift27 | 0.276 | 0.354 | 0.495 |
| Tmem159 | 0.276 | 0.301 | 0.250 |
| Rhou | 0.276 | 0.352 | 0.511 |
| Rsl24d1 | 0.276 | 0.277 | 0.252 |
| Tbc1d23 | 0.276 | 0.249 | 0.308 |
| Spryd3 | 0.276 | 0.158 | 0.148 |
| Cby1 | 0.276 | 0.196 | 0.208 |
| Vps4a | 0.276 | 0.256 | 0.251 |
| Gtf2h1 | 0.276 | 0.230 | 0.221 |
| Abhd4 | 0.275 | 0.159 | 0.176 |
| Caml | 0.275 | 0.281 | 0.270 |
| Chd9 | 0.275 | 0.264 | 0.345 |
| Dhx30 | 0.275 | 0.294 | 0.257 |
| Timp1 | 0.275 | 0.342 | 0.667 |
| Ets1 | 0.275 | 0.248 | 0.161 |
| Casp12 | 0.275 | 0.233 | 0.237 |
| Tmem203 | 0.275 | 0.210 | 0.169 |
| 2610524H06I | 0.275 | 0.286 | 0.312 |
| Tipin | 0.275 | 0.287 | 0.326 |
| Bace1 | 0.275 | 0.617 | 0.562 |
| Ubqln1 | 0.275 | 0.230 | 0.219 |
| Scaf1 | 0.275 | 0.278 | 0.292 |
| Syt11 | 0.275 | 0.475 | 0.350 |
| Rad23a | 0.275 | 0.232 | 0.234 |
| Ep300 | 0.275 | 0.291 | 0.228 |
| Gpc4 | 0.275 | 0.323 | 0.152 |
| Fuca2 | 0.274 | 0.082 | 0.182 |
| Cdc42se2 | 0.274 | 0.182 | 0.145 |
| Mindy1 | 0.274 | 0.148 | 0.144 |
| Gna13 | 0.274 | 0.287 | 0.264 |
| Pdzd11 | 0.274 | 0.218 | 0.270 |
| Fgfrl1 | 0.274 | 0.302 | 0.169 |
| Smg1 | 0.274 | 0.316 | 0.251 |
| Srsf4 | 0.274 | 0.261 | 0.225 |
| Ggps1 | 0.274 | 0.256 | 0.240 |

| | | | |
|---|---|---|---|
| Ormdl2 | 0.274 | 0.069 | 0.100 |
| Pomgnt1 | 0.274 | 0.281 | 0.204 |
| Diaph2 | 0.274 | 0.271 | 0.253 |
| Man1c1 | 0.274 | 0.039 | 0.124 |
| Zfp580 | 0.274 | 0.307 | 0.269 |
| Slc25a36 | 0.274 | 0.293 | 0.275 |
| Ulk1 | 0.274 | 0.262 | 0.183 |
| Ubr3 | 0.274 | 0.273 | 0.218 |
| Ehbp1l1 | 0.274 | 0.475 | 0.264 |
| Decr1 | 0.274 | 0.281 | 0.325 |
| Virma | 0.273 | 0.283 | 0.271 |
| Tmem131 | 0.273 | 0.354 | 0.336 |
| Rnf216 | 0.273 | 0.221 | 0.242 |
| Lypla1 | 0.273 | 0.186 | 0.150 |
| Twsg1 | 0.273 | 0.537 | 0.487 |
| Polr2h | 0.273 | 0.253 | 0.277 |
| Ccdc115 | 0.273 | 0.189 | 0.157 |
| Usp4 | 0.273 | 0.263 | 0.254 |
| Ube2f | 0.273 | 0.226 | 0.218 |
| Txnrd1 | 0.273 | 0.295 | 0.273 |
| Cdr2 | 0.272 | 0.241 | 0.237 |
| Ilf3 | 0.272 | 0.283 | 0.244 |
| Twf2 | 0.272 | 0.131 | 0.167 |
| Abraxas2 | 0.272 | 0.236 | 0.243 |
| Eftud2 | 0.272 | 0.197 | 0.201 |
| Cdk6 | 0.272 | 0.412 | 0.249 |
| Ss18l2 | 0.272 | 0.269 | 0.277 |
| Dapk1 | 0.272 | 0.188 | 0.367 |
| D17Wsu92e | 0.272 | 0.250 | 0.209 |
| Thap12 | 0.272 | 0.317 | 0.285 |
| Prpf6 | 0.272 | 0.230 | 0.206 |
| Fam3a | 0.272 | 0.226 | 0.255 |
| Scpep1 | 0.271 | 0.534 | 0.564 |
| Frzb | 0.271 | 0.220 | 0.061 |
| Fbxo18 | 0.271 | 0.248 | 0.334 |
| Fars2 | 0.271 | 0.178 | 0.187 |
| 1600012H06I | 0.271 | 0.209 | 0.228 |
| Cetn2 | 0.271 | 0.252 | 0.256 |
| Gga2 | 0.271 | 0.226 | 0.343 |
| Kidins220 | 0.271 | 0.326 | 0.296 |
| Pop4 | 0.271 | 0.179 | 0.229 |
| Ift46 | 0.271 | 0.309 | 0.302 |
| Mdm2 | 0.271 | 0.318 | 0.259 |
| Herpud2 | 0.271 | 0.311 | 0.307 |
| Yeats2 | 0.271 | 0.198 | 0.244 |
| Cwf19l2 | 0.270 | 0.226 | 0.222 |
| Cdc42ep1 | 0.270 | 0.067 | 0.116 |
| Slc30a9 | 0.270 | 0.242 | 0.202 |

| | | | |
|---|---|---|---|
| Kat6a | 0.270 | 0.220 | 0.216 |
| Fam216a | 0.270 | 0.223 | 0.179 |
| Stk19 | 0.270 | 0.209 | 0.200 |
| Plekha3 | 0.270 | 0.195 | 0.151 |
| Trim26 | 0.270 | 0.214 | 0.215 |
| Ccm2 | 0.270 | 0.261 | 0.350 |
| Pom121 | 0.270 | 0.213 | 0.214 |
| Plpp5 | 0.270 | 0.193 | 0.269 |
| Lrp5 | 0.270 | 0.174 | 0.083 |
| Adprh | 0.269 | 0.297 | 0.443 |
| Akap8l | 0.269 | 0.265 | 0.262 |
| Borcs6 | 0.269 | 0.249 | 0.235 |
| Nudt16l1 | 0.269 | 0.194 | 0.201 |
| Sde2 | 0.269 | 0.342 | 0.330 |
| Taf7 | 0.269 | 0.288 | 0.327 |
| Kmt5a | 0.269 | 0.426 | 0.294 |
| Taf3 | 0.269 | 0.247 | 0.219 |
| Ctr9 | 0.269 | 0.294 | 0.261 |
| Mxra7 | 0.269 | 0.533 | 0.658 |
| Gle1 | 0.269 | 0.227 | 0.239 |
| Cul7 | 0.269 | 0.212 | 0.347 |
| Il10rb | 0.269 | 0.160 | 0.175 |
| Dcaf7 | 0.268 | 0.145 | 0.160 |
| Smurf1 | 0.268 | 0.137 | 0.129 |
| Bdp1 | 0.268 | 0.247 | 0.243 |
| Rapgef6 | 0.268 | 0.356 | 0.348 |
| Brox | 0.268 | 0.157 | 0.161 |
| Abca3 | 0.268 | 0.080 | 0.075 |
| Fam126a | 0.268 | 0.159 | 0.184 |
| Epc2 | 0.268 | 0.318 | 0.314 |
| Dtnbp1 | 0.268 | 0.370 | 0.445 |
| Fnip2 | 0.268 | 0.089 | 0.049 |
| Triap1 | 0.268 | 0.205 | 0.182 |
| Tob2 | 0.268 | 0.598 | 0.294 |
| Nsun2 | 0.268 | 0.204 | 0.257 |
| Suz12 | 0.267 | 0.275 | 0.207 |
| Dopey2 | 0.267 | 0.100 | 0.089 |
| Slc35a2 | 0.267 | 0.191 | 0.244 |
| Tusc2 | 0.267 | 0.130 | 0.146 |
| Ppfia1 | 0.267 | 0.246 | 0.221 |
| Eogt | 0.267 | 0.070 | 0.034 |
| Aup1 | 0.267 | 0.273 | 0.285 |
| BC029722 | 0.267 | 0.142 | 0.151 |
| Sec24c | 0.267 | 0.172 | 0.161 |
| Ehmt2 | 0.267 | 0.239 | 0.315 |
| Sntg2 | 0.267 | 0.254 | 0.131 |
| Bckdha | 0.267 | 0.282 | 0.251 |
| D16Ertd472e | 0.267 | 0.242 | 0.115 |

| | | | |
|---|---|---|---|
| Sdccag3 | 0.267 | 0.181 | 0.183 |
| Dpp8 | 0.267 | 0.192 | 0.204 |
| Hnrnpll | 0.267 | 0.344 | 0.270 |
| Sun1 | 0.266 | 0.194 | 0.170 |
| Irf3 | 0.266 | 0.308 | 0.266 |
| Laptm4b | 0.266 | 0.282 | 0.311 |
| Ap3m1 | 0.266 | 0.172 | 0.150 |
| 2200002D01I | 0.266 | 0.089 | 0.024 |
| Kifap3 | 0.266 | 0.271 | 0.384 |
| Babam2 | 0.266 | 0.190 | 0.227 |
| Plat | 0.266 | 0.132 | 0.383 |
| N4bp2 | 0.266 | 0.233 | 0.312 |
| Mybbp1a | 0.266 | 0.272 | 0.257 |
| Eif2ak2 | 0.266 | 0.177 | 0.176 |
| Prkar2a | 0.266 | 0.295 | 0.227 |
| Lsm10 | 0.266 | 0.283 | 0.273 |
| Ldb2 | 0.266 | 0.092 | 0.091 |
| Vkorc1l1 | 0.265 | 0.200 | 0.211 |
| Atad2b | 0.265 | 0.264 | 0.230 |
| Spats2l | 0.265 | 0.743 | 0.701 |
| Fgfr3 | 0.265 | 0.034 | 0.011 |
| Nectin3 | 0.265 | 0.295 | 0.334 |
| Wipf1 | 0.265 | 0.126 | 0.210 |
| Nsd2 | 0.265 | 0.225 | 0.189 |
| Lpar6 | 0.265 | 0.122 | 0.077 |
| Spsb3 | 0.265 | 0.203 | 0.188 |
| Pabpc4 | 0.264 | 0.280 | 0.277 |
| Ldah | 0.264 | 0.215 | 0.204 |
| Ero1l | 0.264 | 0.163 | 0.131 |
| Tmub2 | 0.264 | 0.222 | 0.234 |
| Gtf2ird1 | 0.264 | 0.198 | 0.188 |
| Atp6v1h | 0.264 | 0.253 | 0.192 |
| BC003965 | 0.264 | 0.221 | 0.204 |
| Hcfc1 | 0.264 | 0.218 | 0.208 |
| Smn1 | 0.264 | 0.231 | 0.225 |
| Sertad1 | 0.264 | 0.646 | 0.561 |
| Per1 | 0.264 | 0.333 | 0.277 |
| Rprd2 | 0.264 | 0.274 | 0.242 |
| Inafm1 | 0.264 | 0.277 | 0.286 |
| Sft2d3 | 0.264 | 0.150 | 0.147 |
| Zfp950 | 0.264 | 0.226 | 0.316 |
| Arid1b | 0.264 | 0.319 | 0.283 |
| Etl4 | 0.264 | 0.139 | 0.150 |
| Tnks | 0.263 | 0.299 | 0.275 |
| Ccnyl1 | 0.263 | 0.078 | 0.076 |
| Ipo11 | 0.263 | 0.062 | 0.081 |
| Afap1l2 | 0.263 | 0.101 | 0.106 |
| Timm44 | 0.263 | 0.276 | 0.252 |

| | | | |
|---|---|---|---|
| Fh1 | 0.263 | 0.357 | 0.264 |
| Hba-a2 | 0.263 | 0.239 | 0.275 |
| Btg3 | 0.263 | 0.234 | 0.152 |
| Ak6 | 0.263 | 0.260 | 0.214 |
| Zbtb8os | 0.263 | 0.231 | 0.262 |
| Iws1 | 0.263 | 0.250 | 0.249 |
| Xpo7 | 0.263 | 0.178 | 0.205 |
| Polr3k | 0.263 | 0.196 | 0.192 |
| Frmd6 | 0.263 | 0.334 | 0.651 |
| Snx11 | 0.263 | 0.059 | 0.051 |
| Gigyf1 | 0.263 | 0.277 | 0.242 |
| Dcaf13 | 0.263 | 0.218 | 0.202 |
| Cxxc5 | 0.262 | 0.314 | 0.389 |
| Clk3 | 0.262 | 0.243 | 0.273 |
| Coq10b | 0.262 | 0.549 | 0.478 |
| Pik3ca | 0.262 | 0.378 | 0.356 |
| Gtf3c6 | 0.262 | 0.208 | 0.230 |
| Cspp1 | 0.262 | 0.258 | 0.229 |
| Dnajc30 | 0.262 | 0.232 | 0.235 |
| Med9 | 0.262 | 0.222 | 0.236 |
| Atg12 | 0.262 | 0.196 | 0.165 |
| Setd1b | 0.262 | 0.291 | 0.239 |
| Trrap | 0.262 | 0.252 | 0.235 |
| Chn2 | 0.262 | 0.074 | 0.072 |
| Kif1c | 0.262 | 0.247 | 0.198 |
| Tbkbp1 | 0.262 | 0.033 | 0.070 |
| Plxnb2 | 0.262 | 0.367 | 0.459 |
| Usp50 | 0.262 | 0.143 | 0.133 |
| 6-Mar | 0.261 | 0.206 | 0.185 |
| Creld2 | 0.261 | 0.182 | 0.253 |
| Dpf2 | 0.261 | 0.220 | 0.245 |
| Ppp4r3a | 0.261 | 0.291 | 0.227 |
| Msantd4 | 0.261 | 0.189 | 0.173 |
| Stk3 | 0.261 | 0.230 | 0.275 |
| Hoxb4 | 0.261 | 0.500 | 0.349 |
| Kdm7a | 0.261 | 0.290 | 0.308 |
| Ddt | 0.261 | 0.237 | 0.229 |
| Rhobtb3 | 0.261 | 0.328 | 0.449 |
| Fbxo22 | 0.261 | 0.238 | 0.211 |
| Sp1 | 0.261 | 0.257 | 0.263 |
| Zfp451 | 0.261 | 0.245 | 0.290 |
| Prkab1 | 0.261 | 0.084 | 0.095 |
| Hbs1l | 0.261 | 0.217 | 0.211 |
| Maff | 0.261 | 0.343 | 0.300 |
| Lcmt1 | 0.261 | 0.241 | 0.232 |
| Pgd | 0.261 | 0.243 | 0.253 |
| Dnm1l | 0.261 | 0.249 | 0.262 |
| Ccndbp1 | 0.261 | 0.241 | 0.199 |

| | | | |
|---|---|---|---|
| Ahsa2 | 0.261 | 0.204 | 0.196 |
| Cept1 | 0.260 | 0.187 | 0.219 |
| Zfhx3 | 0.260 | 1.214 | 0.629 |
| Cyb5b | 0.260 | 0.193 | 0.170 |
| Hmgn5 | 0.260 | 0.284 | 0.150 |
| Wdr33 | 0.260 | 0.235 | 0.226 |
| Slc31a1 | 0.260 | 0.153 | 0.191 |
| Fbxw8 | 0.260 | 0.176 | 0.253 |
| Tmem98 | 0.260 | 0.184 | 0.159 |
| Zmat5 | 0.260 | 0.234 | 0.226 |
| Btbd2 | 0.260 | 0.237 | 0.269 |
| Cldn15 | 0.260 | 0.007 | 0.015 |
| Mvp | 0.260 | 0.204 | 0.202 |
| Usp8 | 0.260 | 0.272 | 0.261 |
| Sharpin | 0.260 | 0.173 | 0.174 |
| Emb | 0.260 | 0.080 | 0.035 |
| Wdr6 | 0.260 | 0.262 | 0.218 |
| Nova2 | 0.260 | 0.004 | 0.005 |
| Adgra3 | 0.260 | 0.171 | 0.173 |
| Mfsd14a | 0.260 | 0.230 | 0.180 |
| Zfyve9 | 0.259 | 0.152 | 0.168 |
| Imp4 | 0.259 | 0.170 | 0.165 |
| Mdm4 | 0.259 | 0.265 | 0.253 |
| Arfip1 | 0.259 | 0.214 | 0.219 |
| Pias1 | 0.259 | 0.381 | 0.272 |
| Phc3 | 0.259 | 0.240 | 0.201 |
| Asxl1 | 0.259 | 0.189 | 0.164 |
| Sost | 0.259 | 0.643 | 0.157 |
| N4bp1 | 0.259 | 0.179 | 0.215 |
| Mphosph6 | 0.259 | 0.218 | 0.220 |
| Hmg20a | 0.259 | 0.224 | 0.229 |
| Abcd3 | 0.258 | 0.300 | 0.280 |
| Lias | 0.258 | 0.238 | 0.244 |
| Gper1 | 0.258 | 0.043 | 0.014 |
| Klhl9 | 0.258 | 0.249 | 0.271 |
| Enpp4 | 0.258 | 0.207 | 0.089 |
| Rara | 0.258 | 0.271 | 0.291 |
| Mrpl44 | 0.258 | 0.117 | 0.094 |
| Asnsd1 | 0.258 | 0.191 | 0.332 |
| Mylip | 0.258 | 0.195 | 0.195 |
| Lztr1 | 0.258 | 0.235 | 0.260 |
| Lysmd3 | 0.258 | 0.273 | 0.327 |
| Ccdc174 | 0.258 | 0.271 | 0.238 |
| Atp13a3 | 0.258 | 0.299 | 0.258 |
| Hax1 | 0.258 | 0.225 | 0.233 |
| Serpinb9 | 0.258 | 0.169 | 0.113 |
| Mpdz | 0.258 | 0.239 | 0.297 |
| Tmx2 | 0.257 | 0.143 | 0.135 |

| | | | |
|---|---|---|---|
| Tmem47 | 0.257 | 0.248 | 0.147 |
| Ubap2 | 0.257 | 0.236 | 0.253 |
| Lyst | 0.257 | 0.133 | 0.133 |
| Psmb10 | 0.257 | 0.122 | 0.188 |
| Sssca1 | 0.257 | 0.239 | 0.234 |
| AC149090.1 | 0.257 | 0.227 | 0.217 |
| Aig1 | 0.257 | 0.406 | 0.421 |
| Pdcl | 0.257 | 0.166 | 0.137 |
| Fech | 0.257 | 0.216 | 0.205 |
| Arf6 | 0.257 | 0.278 | 0.329 |
| Ccdc102a | 0.257 | 0.280 | 0.423 |
| Tlk2 | 0.256 | 0.253 | 0.229 |
| Prpf38a | 0.256 | 0.264 | 0.222 |
| Akap8 | 0.256 | 0.232 | 0.208 |
| Upf1 | 0.256 | 0.197 | 0.189 |
| Clasp1 | 0.256 | 0.247 | 0.235 |
| Arhgap44 | 0.256 | 0.156 | 0.116 |
| CAAA0111838 | 0.256 | 0.223 | 0.222 |
| Ppm1f | 0.256 | 0.056 | 0.061 |
| Rexo1 | 0.256 | 0.256 | 0.241 |
| Ngrn | 0.256 | 0.160 | 0.155 |
| Tmem70 | 0.256 | 0.255 | 0.222 |
| Rnf139 | 0.256 | 0.216 | 0.191 |
| Mfap1b | 0.256 | 0.194 | 0.183 |
| Dock1 | 0.256 | 0.226 | 0.227 |
| Grk2 | 0.256 | 0.194 | 0.154 |
| Zmiz2 | 0.256 | 0.160 | 0.110 |
| Vps26a | 0.255 | 0.225 | 0.188 |
| Fbxw5 | 0.255 | 0.213 | 0.188 |
| Ino80e | 0.255 | 0.207 | 0.237 |
| Kif26b | 0.255 | 0.105 | 0.021 |
| Pkn3 | 0.255 | 0.004 | 0.006 |
| Nelfe | 0.255 | 0.231 | 0.296 |
| Dennd2a | 0.255 | 0.163 | 0.310 |
| Wdr70 | 0.255 | 0.184 | 0.190 |
| Poldip3 | 0.255 | 0.299 | 0.277 |
| 5031425E22R | 0.255 | 0.226 | 0.226 |
| Slc12a4 | 0.255 | 0.236 | 0.259 |
| Pcyt1a | 0.255 | 0.175 | 0.214 |
| Prorsd1 | 0.255 | 0.265 | 0.293 |
| Cd99l2 | 0.255 | 0.228 | 0.213 |
| Phf20 | 0.255 | 0.266 | 0.289 |
| Phf10 | 0.255 | 0.287 | 0.279 |
| Plekha1 | 0.255 | 0.264 | 0.173 |
| Wfdc1 | 0.255 | 0.229 | 0.195 |
| Trnt1 | 0.255 | 0.231 | 0.232 |
| Stx18 | 0.254 | 0.168 | 0.205 |
| Maip1 | 0.254 | 0.261 | 0.224 |

| | | | |
|---|---|---|---|
| Saysd1 | 0.254 | 0.178 | 0.183 |
| Rce1 | 0.254 | 0.255 | 0.217 |
| Itpa | 0.254 | 0.191 | 0.192 |
| Chd7 | 0.254 | 0.040 | 0.014 |
| Specc1l | 0.254 | 0.303 | 0.315 |
| N6amt1 | 0.254 | 0.361 | 0.325 |
| Cog8 | 0.254 | 0.164 | 0.169 |
| Fam220a | 0.254 | 0.343 | 0.382 |
| Rassf3 | 0.254 | 0.184 | 0.056 |
| Cep57 | 0.254 | 0.218 | 0.161 |
| Proser1 | 0.254 | 0.163 | 0.150 |
| Cxcl12 | 0.254 | 1.022 | 1.359 |
| Fam102b | 0.253 | 0.350 | 0.627 |
| Tnfaip2 | 0.253 | 0.010 | 0.071 |
| Zfp395 | 0.253 | 0.369 | 0.338 |
| Sh3bp4 | 0.253 | 0.135 | 0.158 |
| Rnf220 | 0.253 | 0.191 | 0.208 |
| Gskip | 0.253 | 0.193 | 0.187 |
| Acsl5 | 0.253 | 0.105 | 0.152 |
| Pigs | 0.253 | 0.192 | 0.245 |
| Secisbp2l | 0.253 | 0.208 | 0.189 |
| Dcaf11 | 0.253 | 0.199 | 0.148 |
| Sdc4 | 0.253 | 1.109 | 0.512 |
| Plekhg2 | 0.253 | 0.420 | 0.386 |
| Pih1d1 | 0.253 | 0.204 | 0.294 |
| Ipo7 | 0.253 | 0.251 | 0.223 |
| Abce1 | 0.253 | 0.248 | 0.246 |
| Cbr1 | 0.252 | 0.239 | 0.333 |
| Mrpl50 | 0.252 | 0.262 | 0.271 |
| Hpf1 | 0.252 | 0.292 | 0.285 |
| Tmem100 | 0.252 | 0.015 | 0.239 |
| Tmc6 | 0.252 | 0.131 | 0.082 |
| Epm2aip1 | 0.252 | 0.215 | 0.216 |
| Boc | 0.252 | 0.409 | 0.776 |
| Pias2 | 0.252 | 0.196 | 0.188 |
| Ndst1 | 0.252 | 0.255 | 0.238 |
| Ngdn | 0.252 | 0.248 | 0.209 |
| Plcg1 | 0.252 | 0.165 | 0.173 |
| Mxi1 | 0.252 | 0.259 | 0.304 |
| Pias3 | 0.251 | 0.164 | 0.189 |
| Tesk1 | 0.251 | 0.329 | 0.278 |
| Rabep1 | 0.251 | 0.252 | 0.250 |
| Frmd4a | 0.251 | 0.275 | 0.273 |
| Plekhf2 | 0.251 | 0.154 | 0.117 |
| Psmf1 | 0.251 | 0.149 | 0.157 |
| Slc16a2 | 0.251 | 0.240 | 0.575 |
| Nectin2 | 0.251 | 0.220 | 0.225 |
| Mthfd2l | 0.251 | 0.308 | 0.204 |

| | | | |
|---|---|---|---|
| Fam118b | 0.251 | 0.071 | 0.075 |
| Chka | 0.251 | 0.500 | 0.326 |
| Polb | 0.251 | 0.189 | 0.189 |
| Rap2b | 0.251 | 0.119 | 0.135 |
| Polr1c | 0.251 | 0.217 | 0.205 |
| Atp2a3 | 0.251 | 0.009 | 0.009 |
| Rcor3 | 0.251 | 0.293 | 0.269 |
| Smim20 | 0.251 | 0.175 | 0.218 |
| Idh1 | 0.251 | 0.239 | 0.392 |
| Tdg | 0.251 | 0.201 | 0.196 |
| Zmym4 | 0.251 | 0.233 | 0.234 |
| Zdhhc6 | 0.251 | 0.194 | 0.207 |
| Ide | 0.251 | 0.209 | 0.172 |
| Irf2bp1 | 0.250 | 0.241 | 0.213 |
| Pdpk1 | 0.250 | 0.274 | 0.254 |
| Washc3 | 0.250 | 0.216 | 0.244 |
| Fmr1 | 0.250 | 0.330 | 0.341 |
| Npm3 | 0.250 | 0.419 | 0.358 |
| Uvrag | 0.250 | 0.188 | 0.185 |
| Stat1 | 0.250 | 0.099 | 0.094 |
| Mrpl49 | 0.250 | 0.137 | 0.142 |
| Pafah1b3 | 0.250 | 0.269 | 0.430 |
| Polr2d | 0.250 | 0.209 | 0.192 |
| Dlc1 | 0.250 | 0.485 | 0.623 |
| Fam76b | 0.250 | 0.221 | 0.183 |
| Tbc1d15 | 0.250 | 0.260 | 0.247 |
| Chmp7 | 0.250 | 0.180 | 0.200 |
| Cnot10 | 0.250 | 0.284 | 0.243 |
| Mrpl9 | 0.250 | 0.230 | 0.195 |
| Napg | 0.250 | 0.260 | 0.224 |
| Cdkn1b | 0.250 | 0.301 | 0.209 |
| Mapk8 | 0.250 | 0.214 | 0.221 |
| Atrn | 0.249 | 0.185 | 0.156 |
| Fkbp14 | 0.249 | 0.198 | 0.388 |
| Rif1 | 0.249 | 0.272 | 0.219 |
| Gcsh | 0.249 | 0.264 | 0.297 |
| Itsn1 | 0.249 | 0.156 | 0.219 |
| Usp19 | 0.249 | 0.277 | 0.223 |
| Spcs3 | 0.249 | 0.273 | 0.187 |
| Ddx49 | 0.249 | 0.180 | 0.168 |
| Trib2 | 0.249 | 0.216 | 0.221 |
| Tars | 0.249 | 0.233 | 0.283 |
| Hacd4 | 0.248 | 0.247 | 0.106 |
| Cyb5r4 | 0.248 | 0.147 | 0.158 |
| Col6a2 | 0.248 | 1.490 | 2.154 |
| Gm17018 | 0.248 | 0.198 | 0.171 |
| Rnf114 | 0.248 | 0.137 | 0.101 |
| Apobec3 | 0.248 | 0.121 | 0.086 |

| | | | |
|---|---|---|---|
| Zdhhc4 | 0.248 | 0.220 | 0.204 |
| Pmm2 | 0.248 | 0.161 | 0.164 |
| Ppp1r9b | 0.248 | 0.226 | 0.224 |
| Kin | 0.248 | 0.229 | 0.209 |
| Oard1 | 0.248 | 0.168 | 0.157 |
| Zhx1 | 0.248 | 0.166 | 0.162 |
| Camk2g | 0.248 | 0.429 | 0.221 |
| Fxr2 | 0.248 | 0.238 | 0.226 |
| Acap2 | 0.248 | 0.231 | 0.281 |
| Cand1 | 0.247 | 0.280 | 0.284 |
| Ammecr1l | 0.247 | 0.243 | 0.263 |
| Ddx52 | 0.247 | 0.223 | 0.212 |
| Daam1 | 0.247 | 0.703 | 0.873 |
| Nfyb | 0.247 | 0.239 | 0.243 |
| Lrch3 | 0.247 | 0.226 | 0.188 |
| Rilpl1 | 0.247 | 0.211 | 0.223 |
| Rbm27 | 0.247 | 0.255 | 0.236 |
| Cep170 | 0.247 | 0.192 | 0.292 |
| Taz | 0.247 | 0.200 | 0.195 |
| Rxrb | 0.247 | 0.218 | 0.236 |
| Snx7 | 0.247 | 0.196 | 0.244 |
| Dcaf12 | 0.247 | 0.185 | 0.200 |
| Fbrsl1 | 0.247 | 0.174 | 0.126 |
| Slain2 | 0.247 | 0.248 | 0.227 |
| Rlf | 0.247 | 0.278 | 0.263 |
| Dvl1 | 0.247 | 0.162 | 0.162 |
| Smarcad1 | 0.246 | 0.174 | 0.196 |
| Gstp1 | 0.246 | 0.313 | 0.306 |
| Erlin2 | 0.246 | 0.121 | 0.177 |
| Tbx18 | 0.246 | 0.186 | 0.198 |
| Synrg | 0.246 | 0.212 | 0.167 |
| Ets2 | 0.246 | 0.460 | 0.252 |
| Prdm16 | 0.246 | 0.292 | 0.065 |
| Zdhhc16 | 0.246 | 0.152 | 0.185 |
| Mustn1 | 0.246 | 0.740 | 0.188 |
| Lama5 | 0.246 | 0.425 | 0.069 |
| Bud23 | 0.246 | 0.176 | 0.180 |
| Zfp652 | 0.245 | 0.248 | 0.278 |
| Capn6 | 0.245 | 0.227 | 0.342 |
| Tbcc | 0.245 | 0.182 | 0.201 |
| Pex11b | 0.245 | 0.203 | 0.209 |
| Siah1a | 0.245 | 0.280 | 0.255 |
| Rmnd5a | 0.245 | 0.226 | 0.177 |
| Map2k5 | 0.245 | 0.174 | 0.192 |
| Atp6v1b2 | 0.245 | 0.166 | 0.176 |
| Nipsnap2 | 0.245 | 0.188 | 0.189 |
| Ash2l | 0.245 | 0.238 | 0.242 |
| Pttg1 | 0.245 | 0.307 | 0.354 |

| | | | |
|---|---|---|---|
| Baz2a | 0.245 | 0.209 | 0.155 |
| Cdc73 | 0.245 | 0.259 | 0.234 |
| Rprd1b | 0.245 | 0.169 | 0.162 |
| Pex3 | 0.245 | 0.171 | 0.199 |
| Acyp1 | 0.245 | 0.158 | 0.155 |
| Bloc1s1 | 0.245 | 0.267 | 0.316 |
| Pank2 | 0.244 | 0.196 | 0.189 |
| Orai3 | 0.244 | 0.253 | 0.254 |
| Gt(ROSA)26So | 0.244 | 0.298 | 0.218 |
| Rnf166 | 0.244 | 0.188 | 0.175 |
| Pgap2 | 0.244 | 0.183 | 0.236 |
| Cnpy4 | 0.244 | 0.163 | 0.208 |
| Cebpg | 0.244 | 0.205 | 0.300 |
| Soat1 | 0.244 | 0.207 | 0.268 |
| Ncbp2 | 0.244 | 0.298 | 0.330 |
| Sart1 | 0.244 | 0.219 | 0.201 |
| Fkbp5 | 0.244 | 0.446 | 0.303 |
| Ptdss1 | 0.244 | 0.176 | 0.199 |
| 2300009A05F | 0.244 | 0.211 | 0.156 |
| Cish | 0.244 | 0.273 | 0.239 |
| Med19 | 0.244 | 0.141 | 0.128 |
| Trim33 | 0.244 | 0.232 | 0.207 |
| Stxbp3 | 0.244 | 0.250 | 0.199 |
| Gngt2 | 0.244 | 0.011 | 0.028 |
| Arl4d | 0.244 | 0.075 | 0.053 |
| Phf23 | 0.244 | 0.281 | 0.277 |
| Tufm | 0.243 | 0.162 | 0.178 |
| Lix1l | 0.243 | 0.383 | 0.431 |
| Srgap2 | 0.243 | 0.221 | 0.267 |
| Fam181b | 0.243 | 0.015 | 0.014 |
| Abcf2 | 0.243 | 0.201 | 0.192 |
| Snn | 0.243 | 0.109 | 0.105 |
| Dhrs4 | 0.243 | 0.270 | 0.276 |
| Otud5 | 0.243 | 0.196 | 0.191 |
| Cdkn2d | 0.243 | 0.229 | 0.182 |
| Slc30a5 | 0.243 | 0.231 | 0.239 |
| Vamp7 | 0.243 | 0.177 | 0.180 |
| Metap1 | 0.243 | 0.191 | 0.133 |
| Ptdss2 | 0.243 | 0.245 | 0.279 |
| Dcun1d1 | 0.243 | 0.266 | 0.211 |
| Arnt | 0.243 | 0.241 | 0.243 |
| Zfp511 | 0.243 | 0.189 | 0.167 |
| Mapre2 | 0.243 | 0.247 | 0.267 |
| Gtf3c1 | 0.243 | 0.192 | 0.184 |
| Ajuba | 0.243 | 0.107 | 0.083 |
| Fam172a | 0.243 | 0.275 | 0.322 |
| Pcolce | 0.243 | 0.385 | 1.805 |
| Dcps | 0.243 | 0.220 | 0.213 |

| | | | |
|---|---|---|---|
| Gamt | 0.243 | 0.426 | 0.411 |
| Atn1 | 0.243 | 0.260 | 0.206 |
| 4930402H24I | 0.242 | 0.213 | 0.257 |
| Zc3h7b | 0.242 | 0.219 | 0.192 |
| Xpo1 | 0.242 | 0.235 | 0.221 |
| Ppp5c | 0.242 | 0.231 | 0.215 |
| Pycard | 0.242 | 0.232 | 0.296 |
| Cep164 | 0.242 | 0.224 | 0.241 |
| 2310033P09I | 0.242 | 0.140 | 0.130 |
| Ick | 0.242 | 0.212 | 0.239 |
| Tmem45a | 0.241 | 0.272 | 0.417 |
| Fat1 | 0.241 | 0.403 | 0.260 |
| Abhd12 | 0.241 | 0.214 | 0.235 |
| Dip2b | 0.241 | 0.177 | 0.154 |
| Dhps | 0.241 | 0.229 | 0.200 |
| Osr1 | 0.241 | 0.819 | 1.029 |
| Prpf39 | 0.241 | 0.260 | 0.234 |
| Fdps | 0.241 | 0.334 | 0.347 |
| Mrps9 | 0.241 | 0.261 | 0.228 |
| Znhit2 | 0.241 | 0.112 | 0.123 |
| Tbpl1 | 0.241 | 0.218 | 0.204 |
| Pfn2 | 0.241 | 0.300 | 0.345 |
| Cpsf7 | 0.241 | 0.231 | 0.214 |
| Nfatc3 | 0.241 | 0.192 | 0.190 |
| Farsa | 0.241 | 0.216 | 0.199 |
| Lrrfip2 | 0.241 | 0.359 | 0.313 |
| AC160336.1 | 0.240 | 0.276 | 0.312 |
| Acd | 0.240 | 0.188 | 0.197 |
| Nudt5 | 0.240 | 0.160 | 0.172 |
| Dkc1 | 0.240 | 0.186 | 0.196 |
| Rfc3 | 0.240 | 0.212 | 0.188 |
| Coq7 | 0.240 | 0.263 | 0.263 |
| Fzd1 | 0.240 | 0.486 | 0.310 |
| Stim1 | 0.240 | 0.233 | 0.270 |
| Lars | 0.240 | 0.263 | 0.233 |
| Pacs1 | 0.240 | 0.140 | 0.151 |
| Itpkb | 0.240 | 0.225 | 0.165 |
| Eid2 | 0.240 | 0.284 | 0.291 |
| Mrps25 | 0.240 | 0.218 | 0.233 |
| Nsf | 0.240 | 0.142 | 0.141 |
| Nolc1 | 0.239 | 0.233 | 0.255 |
| Hexim1 | 0.239 | 0.334 | 0.252 |
| G6pc3 | 0.239 | 0.161 | 0.137 |
| Cep350 | 0.239 | 0.211 | 0.196 |
| Tle4 | 0.239 | 0.152 | 0.077 |
| Tmem183a | 0.239 | 0.241 | 0.194 |
| Ypel5 | 0.239 | 0.229 | 0.198 |
| Slc45a4 | 0.239 | 0.081 | 0.109 |

| | | | |
|---|---|---|---|
| Ranbp9 | 0.239 | 0.250 | 0.192 |
| Zfp521 | 0.239 | 0.030 | 0.127 |
| Mrpl45 | 0.239 | 0.196 | 0.194 |
| Mak16 | 0.239 | 0.175 | 0.165 |
| Mfap1a | 0.239 | 0.172 | 0.184 |
| Golga3 | 0.239 | 0.199 | 0.236 |
| Bex4 | 0.239 | 0.283 | 0.041 |
| Dido1 | 0.238 | 0.231 | 0.231 |
| Spg20 | 0.238 | 0.185 | 0.232 |
| Sec24b | 0.238 | 0.207 | 0.216 |
| Otud7b | 0.238 | 0.259 | 0.261 |
| Cwc25 | 0.238 | 0.180 | 0.210 |
| Sppl2a | 0.238 | 0.218 | 0.243 |
| Tmem101 | 0.238 | 0.178 | 0.183 |
| Ddr1 | 0.238 | 0.325 | 0.173 |
| Hs2st1 | 0.238 | 0.351 | 0.395 |
| Cbr2 | 0.238 | 0.230 | 0.033 |
| Cmtm3 | 0.238 | 0.239 | 0.644 |
| Anapc7 | 0.238 | 0.332 | 0.220 |
| Nip7 | 0.238 | 0.201 | 0.206 |
| Ddx47 | 0.238 | 0.226 | 0.210 |
| Rsbn1 | 0.238 | 0.200 | 0.160 |
| Pqbp1 | 0.238 | 0.252 | 0.254 |
| Mrpl46 | 0.238 | 0.186 | 0.153 |
| Cry2 | 0.238 | 0.223 | 0.243 |
| Dusp8 | 0.238 | 0.329 | 0.238 |
| Sms | 0.238 | 0.180 | 0.178 |
| Ap4s1 | 0.238 | 0.237 | 0.212 |
| Selenbp1 | 0.237 | 0.220 | 0.395 |
| Ttc4 | 0.237 | 0.204 | 0.193 |
| Eef2kmt | 0.237 | 0.157 | 0.206 |
| Taf12 | 0.237 | 0.156 | 0.167 |
| Got2 | 0.237 | 0.310 | 0.264 |
| Dennd6a | 0.237 | 0.171 | 0.159 |
| Apip | 0.237 | 0.256 | 0.233 |
| Ciao1 | 0.237 | 0.170 | 0.146 |
| Pes1 | 0.237 | 0.159 | 0.169 |
| B3galnt1 | 0.237 | 0.218 | 0.165 |
| Map2k4 | 0.237 | 0.204 | 0.174 |
| Fubp3 | 0.237 | 0.255 | 0.210 |
| Phactr4 | 0.237 | 0.166 | 0.136 |
| Ddx19a | 0.237 | 0.182 | 0.177 |
| Sh3pxd2b | 0.237 | 0.387 | 0.451 |
| Rnf38 | 0.237 | 0.146 | 0.111 |
| Nsmce3 | 0.237 | 0.279 | 0.236 |
| Tmem230 | 0.237 | 0.218 | 0.271 |
| Golt1b | 0.236 | 0.266 | 0.345 |
| Mtf2 | 0.236 | 0.196 | 0.165 |

| | | | |
|---|---|---|---|
| Mgrn1 | 0.236 | 0.261 | 0.230 |
| Cxcl2 | 0.236 | 0.170 | 0.134 |
| Lrif1 | 0.236 | 0.141 | 0.174 |
| Gem | 0.236 | 1.524 | 0.769 |
| Tcta | 0.236 | 0.148 | 0.194 |
| Zyg11b | 0.236 | 0.203 | 0.182 |
| Mrps11 | 0.236 | 0.195 | 0.195 |
| Rnf13 | 0.236 | 0.232 | 0.257 |
| Tbce | 0.236 | 0.209 | 0.182 |
| Dym | 0.236 | 0.111 | 0.154 |
| Pxk | 0.235 | 0.164 | 0.171 |
| Pik3c2b | 0.235 | 0.040 | 0.035 |
| Irf1 | 0.235 | 0.839 | 0.762 |
| Fam193b | 0.235 | 0.183 | 0.196 |
| Gm26699 | 0.235 | 0.282 | 0.232 |
| Plod2 | 0.235 | 0.458 | 0.568 |
| Kansl3 | 0.235 | 0.230 | 0.247 |
| Epb41l4aos | 0.235 | 0.306 | 0.290 |
| Cpxm1 | 0.235 | 0.261 | 0.811 |
| Spg7 | 0.235 | 0.186 | 0.168 |
| Adgra2 | 0.235 | 0.213 | 0.242 |
| Cd2bp2 | 0.235 | 0.158 | 0.178 |
| Hdgfl2 | 0.235 | 0.285 | 0.270 |
| Dnal4 | 0.235 | 0.206 | 0.201 |
| Ptprf | 0.234 | 0.313 | 0.257 |
| Nmd3 | 0.234 | 0.252 | 0.183 |
| Gabbr1 | 0.234 | 0.244 | 0.238 |
| Dhx40 | 0.234 | 0.237 | 0.199 |
| Arrdc3 | 0.234 | 0.199 | 0.221 |
| Cap1 | 0.234 | 0.348 | 0.245 |
| Ubl7 | 0.234 | 0.165 | 0.179 |
| Atxn7 | 0.234 | 0.266 | 0.201 |
| Sf3a1 | 0.234 | 0.177 | 0.183 |
| Casd1 | 0.234 | 0.173 | 0.163 |
| Bin1 | 0.234 | 0.276 | 0.414 |
| Cpne2 | 0.234 | 0.220 | 0.074 |
| Ptpn18 | 0.233 | 0.024 | 0.026 |
| Plaa | 0.233 | 0.233 | 0.192 |
| Fbxw11 | 0.233 | 0.211 | 0.172 |
| Hoxa3 | 0.233 | 0.307 | 0.268 |
| Tram2 | 0.233 | 0.164 | 0.167 |
| Ppp2r5e | 0.233 | 0.224 | 0.219 |
| Ppa2 | 0.233 | 0.242 | 0.229 |
| Pds5b | 0.233 | 0.285 | 0.300 |
| Slc25a17 | 0.233 | 0.311 | 0.261 |
| Las1l | 0.233 | 0.216 | 0.192 |
| Eif4enif1 | 0.233 | 0.242 | 0.237 |
| Kctd20 | 0.233 | 0.192 | 0.250 |

| | | | |
|---|---|---|---|
| Glipr2 | 0.233 | 0.189 | 0.328 |
| Ntmt1 | 0.233 | 0.260 | 0.264 |
| Ate1 | 0.233 | 0.245 | 0.184 |
| Ap1g1 | 0.232 | 0.149 | 0.157 |
| Amz2 | 0.232 | 0.170 | 0.258 |
| Taf13 | 0.232 | 0.235 | 0.196 |
| Arhgef10 | 0.232 | 0.081 | 0.153 |
| Pros1 | 0.232 | 0.215 | 0.237 |
| Rhot1 | 0.232 | 0.237 | 0.204 |
| Braf | 0.232 | 0.325 | 0.239 |
| Vars | 0.232 | 0.205 | 0.219 |
| Sap30bp | 0.232 | 0.212 | 0.179 |
| Gba | 0.232 | 0.088 | 0.166 |
| Exoc4 | 0.232 | 0.198 | 0.218 |
| Vegfa | 0.232 | 0.116 | 0.202 |
| Ndufs1 | 0.232 | 0.256 | 0.218 |
| Hist3h2a | 0.232 | 0.199 | 0.165 |
| Tet2 | 0.232 | 0.254 | 0.185 |
| Man2a2 | 0.232 | 0.219 | 0.233 |
| Katnbl1 | 0.232 | 0.159 | 0.143 |
| 1300002E11F | 0.232 | 0.160 | 0.130 |
| Mbd1 | 0.232 | 0.121 | 0.209 |
| Mcm6 | 0.231 | 0.241 | 0.267 |
| Zfp945 | 0.231 | 0.222 | 0.263 |
| Gna12 | 0.231 | 0.252 | 0.214 |
| Plpbp | 0.231 | 0.230 | 0.222 |
| Idnk | 0.231 | 0.223 | 0.212 |
| Rnf146 | 0.231 | 0.262 | 0.226 |
| Smad2 | 0.231 | 0.188 | 0.181 |
| Lrrcc1 | 0.231 | 0.190 | 0.200 |
| Elp3 | 0.231 | 0.161 | 0.176 |
| Kctd17 | 0.231 | 0.238 | 0.327 |
| Smarcd2 | 0.231 | 0.153 | 0.312 |
| Unc119 | 0.231 | 0.363 | 0.232 |
| Ppp2r3c | 0.231 | 0.160 | 0.159 |
| Dcaf10 | 0.231 | 0.212 | 0.219 |
| Fbn2 | 0.230 | 1.132 | 1.471 |
| Sumf2 | 0.230 | 0.139 | 0.214 |
| Thap7 | 0.230 | 0.166 | 0.172 |
| Ubfd1 | 0.230 | 0.198 | 0.288 |
| Rwdd4a | 0.230 | 0.164 | 0.147 |
| Ublcp1 | 0.230 | 0.246 | 0.210 |
| Ubr1 | 0.230 | 0.213 | 0.194 |
| 9930021J03R | 0.230 | 0.317 | 0.329 |
| Dctn5 | 0.230 | 0.164 | 0.199 |
| Nt5e | 0.230 | 0.205 | 0.044 |
| Upf2 | 0.230 | 0.246 | 0.216 |
| Ttc5 | 0.230 | 0.187 | 0.163 |

| | | | |
|---|---|---|---|
| Zfp608 | 0.230 | 0.103 | 0.167 |
| Sf3a2 | 0.230 | 0.184 | 0.184 |
| Numb | 0.230 | 0.102 | 0.107 |
| Hsd17b11 | 0.229 | 0.320 | 0.363 |
| Gm2a | 0.229 | 0.298 | 0.291 |
| Zfp422 | 0.229 | 0.203 | 0.187 |
| Ascc1 | 0.229 | 0.192 | 0.199 |
| Med30 | 0.229 | 0.234 | 0.174 |
| Senp2 | 0.229 | 0.290 | 0.257 |
| Epc1 | 0.229 | 0.237 | 0.200 |
| Bcar1 | 0.229 | 0.182 | 0.166 |
| Myh11 | 0.229 | 1.778 | 0.328 |
| Pnkp | 0.229 | 0.131 | 0.128 |
| Fam171a2 | 0.229 | 0.204 | 0.250 |
| Vps53 | 0.229 | 0.160 | 0.146 |
| Fam69a | 0.229 | 0.160 | 0.217 |
| Syne1 | 0.229 | 0.725 | 0.286 |
| Ddx41 | 0.229 | 0.200 | 0.186 |
| Tmem120a | 0.229 | 0.322 | 0.226 |
| Timm10 | 0.229 | 0.192 | 0.176 |
| H2-Ke6 | 0.229 | 0.171 | 0.139 |
| Setx | 0.228 | 0.167 | 0.142 |
| Ift43 | 0.228 | 0.548 | 0.420 |
| Tln2 | 0.228 | 0.078 | 0.159 |
| Ubqln2 | 0.228 | 0.184 | 0.177 |
| Grhpr | 0.228 | 0.287 | 0.211 |
| Arl15 | 0.228 | 0.050 | 0.081 |
| Birc2 | 0.228 | 0.398 | 0.295 |
| Zc3h18 | 0.228 | 0.239 | 0.231 |
| Setd7 | 0.228 | 0.226 | 0.221 |
| Pole4 | 0.228 | 0.288 | 0.295 |
| Fnbp4 | 0.228 | 0.228 | 0.218 |
| Zc2hc1a | 0.228 | 0.376 | 0.266 |
| Cgrrf1 | 0.228 | 0.167 | 0.163 |
| Arl5b | 0.228 | 0.253 | 0.291 |
| Nsmce2 | 0.228 | 0.273 | 0.298 |
| Etv6 | 0.228 | 0.190 | 0.113 |
| Mt3 | 0.228 | 0.004 | 0.005 |
| Osbpl8 | 0.228 | 0.216 | 0.177 |
| Dbt | 0.227 | 0.130 | 0.064 |
| Tsr3 | 0.227 | 0.204 | 0.189 |
| Rnf121 | 0.227 | 0.159 | 0.190 |
| Ndrg3 | 0.227 | 0.169 | 0.198 |
| Casc3 | 0.227 | 0.209 | 0.180 |
| Neu1 | 0.227 | 0.184 | 0.170 |
| Rnmt | 0.227 | 0.241 | 0.213 |
| Thbs2 | 0.227 | 0.526 | 1.017 |
| Cd44 | 0.227 | 0.263 | 0.392 |

| | | | |
|---|---|---|---|
| Mpv17 | 0.227 | 0.229 | 0.250 |
| Ciapin1 | 0.227 | 0.148 | 0.132 |
| Orc6 | 0.227 | 0.235 | 0.247 |
| Usp5 | 0.227 | 0.199 | 0.210 |
| Casp8ap2 | 0.227 | 0.231 | 0.243 |
| Nfyc | 0.227 | 0.197 | 0.170 |
| 9030624J02R | 0.227 | 0.199 | 0.219 |
| Galnt11 | 0.227 | 0.228 | 0.209 |
| Serhl | 0.227 | 0.183 | 0.172 |
| Pld1 | 0.227 | 0.164 | 0.172 |
| Plgrkt | 0.227 | 0.379 | 0.328 |
| Taok3 | 0.227 | 0.229 | 0.343 |
| Pex6 | 0.227 | 0.143 | 0.195 |
| Ube4a | 0.227 | 0.157 | 0.168 |
| Bop1 | 0.226 | 0.106 | 0.080 |
| Gm48898 | 0.226 | 0.001 | 0.002 |
| Rbm33 | 0.226 | 0.254 | 0.256 |
| Zfp24 | 0.226 | 0.203 | 0.171 |
| 4930523C07R | 0.226 | 0.923 | 0.406 |
| Stom | 0.226 | 0.590 | 0.231 |
| St6galnac4 | 0.226 | 0.196 | 0.191 |
| Eef1e1 | 0.226 | 0.113 | 0.087 |
| Sdhaf2 | 0.226 | 0.167 | 0.192 |
| Robo1 | 0.226 | 0.047 | 0.096 |
| Mtss1 | 0.226 | 0.270 | 0.100 |
| Sacm1l | 0.226 | 0.215 | 0.175 |
| Tmem185a | 0.226 | 0.131 | 0.139 |
| E2f4 | 0.226 | 0.149 | 0.164 |
| Rnft1 | 0.226 | 0.320 | 0.341 |
| Flywch1 | 0.226 | 0.171 | 0.233 |
| Mtmr6 | 0.226 | 0.232 | 0.217 |
| Kdm5c | 0.226 | 0.210 | 0.194 |
| Rcbtb2 | 0.226 | 0.306 | 0.282 |
| Pak4 | 0.226 | 0.062 | 0.061 |
| Pef1 | 0.226 | 0.170 | 0.167 |
| Tmem250-ps | 0.226 | 0.183 | 0.159 |
| Map3k7 | 0.226 | 0.181 | 0.187 |
| Senp7 | 0.226 | 0.259 | 0.271 |
| Sec16a | 0.226 | 0.189 | 0.214 |
| Rab3d | 0.226 | 0.155 | 0.198 |
| Nr4a2 | 0.225 | 1.099 | 0.475 |
| Shtn1 | 0.225 | 0.004 | 0.009 |
| Plin2 | 0.225 | 0.250 | 0.270 |
| Fasn | 0.225 | 0.191 | 0.256 |
| Fbxl5 | 0.225 | 0.184 | 0.190 |
| Anxa4 | 0.225 | 0.202 | 0.226 |
| Uvssa | 0.225 | 0.246 | 0.287 |
| Dalrd3 | 0.225 | 0.148 | 0.127 |

| | | | |
|---|---|---|---|
| Thap3 | 0.225 | 0.289 | 0.297 |
| Hacd1 | 0.225 | 0.804 | 0.364 |
| 0610009B22R | 0.225 | 0.197 | 0.176 |
| Epn2 | 0.224 | 0.238 | 0.208 |
| Akap11 | 0.224 | 0.256 | 0.222 |
| Aasdhppt | 0.224 | 0.224 | 0.187 |
| Btbd9 | 0.224 | 0.158 | 0.176 |
| Snx12 | 0.224 | 0.191 | 0.206 |
| Nfx1 | 0.224 | 0.237 | 0.235 |
| Trappc13 | 0.224 | 0.229 | 0.173 |
| Plpp2 | 0.224 | 0.095 | 0.043 |
| Tbl1xr1 | 0.224 | 0.225 | 0.190 |
| Dram1 | 0.224 | 0.054 | 0.078 |
| Ctsc | 0.224 | 0.219 | 0.084 |
| Dcakd | 0.224 | 0.233 | 0.306 |
| Rangrf | 0.223 | 0.126 | 0.108 |
| Tnfrsf12a | 0.223 | 0.639 | 0.315 |
| Agtrap | 0.223 | 0.126 | 0.230 |
| Ezr | 0.223 | 0.096 | 0.072 |
| Ints3 | 0.223 | 0.110 | 0.155 |
| Ifnar1 | 0.223 | 0.204 | 0.216 |
| Ppp2r3d | 0.223 | 0.214 | 0.219 |
| Mecp2 | 0.223 | 0.253 | 0.189 |
| Afap1 | 0.223 | 0.351 | 0.361 |
| Dsp | 0.223 | 0.200 | 0.035 |
| Dhrs3 | 0.223 | 0.450 | 0.382 |
| Mplkip | 0.223 | 0.218 | 0.280 |
| Dmxl1 | 0.223 | 0.230 | 0.197 |
| Npc1 | 0.223 | 0.291 | 0.171 |
| Trim25 | 0.223 | 0.051 | 0.054 |
| Rnf111 | 0.223 | 0.187 | 0.178 |
| Eml4 | 0.223 | 0.288 | 0.282 |
| BC017643 | 0.223 | 0.125 | 0.183 |
| Ssh1 | 0.223 | 0.199 | 0.177 |
| Pcgf3 | 0.222 | 0.187 | 0.191 |
| Gimap4 | 0.222 | 0.002 | 0.002 |
| Atrnl1 | 0.222 | 0.135 | 0.133 |
| Ythdf3 | 0.222 | 0.251 | 0.220 |
| Dirc2 | 0.222 | 0.264 | 0.257 |
| Cln5 | 0.222 | 0.130 | 0.215 |
| Slc1a4 | 0.222 | 0.159 | 0.133 |
| Gfod1 | 0.222 | 0.079 | 0.036 |
| Tspan5 | 0.222 | 0.186 | 0.227 |
| Myo9a | 0.222 | 0.291 | 0.330 |
| Btd | 0.222 | 0.186 | 0.228 |
| Ulk2 | 0.222 | 0.200 | 0.242 |
| Atp6v0a2 | 0.221 | 0.195 | 0.203 |
| Apmap | 0.221 | 0.139 | 0.101 |

| | | | |
|---|---|---|---|
| Rnf181 | 0.221 | 0.141 | 0.147 |
| Gak | 0.221 | 0.184 | 0.174 |
| Cog4 | 0.221 | 0.174 | 0.192 |
| Zfp322a | 0.221 | 0.190 | 0.201 |
| Ncbp3 | 0.221 | 0.197 | 0.184 |
| Mgat5 | 0.221 | 0.114 | 0.072 |
| Pgm2 | 0.221 | 0.111 | 0.246 |
| Brap | 0.221 | 0.170 | 0.143 |
| Plek2 | 0.221 | 0.002 | 0.001 |
| Smpd2 | 0.221 | 0.203 | 0.171 |
| Tbc1d10b | 0.221 | 0.165 | 0.174 |
| Snx21 | 0.221 | 0.213 | 0.252 |
| Ppp6r1 | 0.221 | 0.144 | 0.112 |
| Map3k6 | 0.221 | 0.073 | 0.076 |
| Tmem168 | 0.221 | 0.216 | 0.188 |
| Obsl1 | 0.221 | 0.230 | 0.407 |
| Zfp639 | 0.221 | 0.267 | 0.264 |
| Vps13a | 0.221 | 0.099 | 0.074 |
| Crbn | 0.221 | 0.191 | 0.158 |
| Vps13d | 0.221 | 0.187 | 0.167 |
| Hscb | 0.220 | 0.089 | 0.113 |
| Eloa | 0.220 | 0.183 | 0.206 |
| Tpst1 | 0.220 | 0.306 | 0.455 |
| Nipsnap3b | 0.220 | 0.180 | 0.178 |
| Washc4 | 0.220 | 0.247 | 0.247 |
| Rnf145 | 0.220 | 0.203 | 0.235 |
| Man1b1 | 0.220 | 0.145 | 0.191 |
| Idh3a | 0.220 | 0.228 | 0.175 |
| Acot13 | 0.220 | 0.209 | 0.235 |
| Prdm2 | 0.220 | 0.212 | 0.235 |
| Cnn1 | 0.220 | 1.423 | 0.333 |
| Tgfbr1 | 0.220 | 0.360 | 0.302 |
| Zfp329 | 0.220 | 0.158 | 0.158 |
| Brms1 | 0.220 | 0.081 | 0.085 |
| Hgsnat | 0.220 | 0.168 | 0.179 |
| Prpf31 | 0.220 | 0.189 | 0.151 |
| Alkbh3 | 0.219 | 0.199 | 0.169 |
| Fbxw9 | 0.219 | 0.143 | 0.165 |
| Ube2d1 | 0.219 | 0.199 | 0.192 |
| Exosc1 | 0.219 | 0.158 | 0.183 |
| Spaca6 | 0.219 | 0.290 | 0.256 |
| Hilpda | 0.219 | 0.210 | 0.181 |
| Drg2 | 0.219 | 0.180 | 0.182 |
| Serpina3g | 0.219 | 0.016 | 0.091 |
| Cndp2 | 0.219 | 0.158 | 0.159 |
| Nek9 | 0.219 | 0.193 | 0.193 |
| Osbp | 0.219 | 0.193 | 0.166 |
| Cdadc1 | 0.219 | 0.220 | 0.194 |

| | | | |
|---|---|---|---|
| Tollip | 0.219 | 0.130 | 0.124 |
| Il15 | 0.219 | 0.080 | 0.026 |
| Ap4m1 | 0.219 | 0.194 | 0.219 |
| Mpst | 0.218 | 0.206 | 0.222 |
| Fam58b | 0.218 | 0.156 | 0.162 |
| Nadk2 | 0.218 | 0.118 | 0.132 |
| Tpcn1 | 0.218 | 0.068 | 0.168 |
| Atxn7l3 | 0.218 | 0.207 | 0.178 |
| Sec24a | 0.218 | 0.225 | 0.279 |
| Polg | 0.218 | 0.204 | 0.182 |
| Trim41 | 0.218 | 0.197 | 0.138 |
| Rnaseh2b | 0.217 | 0.191 | 0.277 |
| Ercc1 | 0.217 | 0.260 | 0.260 |
| Abl1 | 0.217 | 0.329 | 0.336 |
| Papd4 | 0.217 | 0.217 | 0.256 |
| Pum3 | 0.217 | 0.194 | 0.238 |
| Dab2 | 0.217 | 0.270 | 0.546 |
| Clasp2 | 0.217 | 0.175 | 0.163 |
| Zfp579 | 0.217 | 0.150 | 0.117 |
| Peak1 | 0.217 | 0.382 | 0.454 |
| D10Jhu81e | 0.217 | 0.194 | 0.207 |
| Bcl2 | 0.217 | 0.335 | 0.341 |
| Lonp1 | 0.217 | 0.210 | 0.192 |
| Carnmt1 | 0.217 | 0.249 | 0.309 |
| Stxbp1 | 0.217 | 0.185 | 0.159 |
| Tshz1 | 0.217 | 0.373 | 0.296 |
| Gtf2f1 | 0.217 | 0.222 | 0.204 |
| Zfp362 | 0.217 | 0.216 | 0.168 |
| Zdhhc8 | 0.217 | 0.249 | 0.205 |
| Wdr45 | 0.217 | 0.192 | 0.203 |
| Ppp2cb | 0.216 | 0.244 | 0.184 |
| Aqr | 0.216 | 0.174 | 0.166 |
| Mindy3 | 0.216 | 0.233 | 0.157 |
| Ctdspl2 | 0.216 | 0.231 | 0.226 |
| Thumpd1 | 0.216 | 0.188 | 0.177 |
| Txnrd3 | 0.216 | 0.137 | 0.093 |
| Serpine1 | 0.216 | 0.545 | 0.343 |
| Otud6b | 0.216 | 0.163 | 0.132 |
| Tmtc1 | 0.216 | 0.135 | 0.149 |
| Syde1 | 0.216 | 0.240 | 0.220 |
| Lmo1 | 0.216 | 0.274 | 0.101 |
| Retreg3 | 0.216 | 0.151 | 0.166 |
| Zfp131 | 0.216 | 0.330 | 0.255 |
| Tppp3 | 0.216 | 0.272 | 0.652 |
| Slc37a3 | 0.216 | 0.148 | 0.240 |
| Tmem161a | 0.216 | 0.203 | 0.176 |
| Bbc3 | 0.216 | 0.132 | 0.147 |
| 2210016F16F | 0.215 | 0.278 | 0.276 |

| | | | |
|---|---|---|---|
| Rnf138 | 0.215 | 0.254 | 0.197 |
| Snx1 | 0.215 | 0.211 | 0.257 |
| Msl3 | 0.215 | 0.229 | 0.259 |
| Pdcd7 | 0.215 | 0.195 | 0.176 |
| 1110012L19F | 0.215 | 0.169 | 0.176 |
| Cyp20a1 | 0.215 | 0.253 | 0.413 |
| Dnajc14 | 0.215 | 0.201 | 0.222 |
| Aplp1 | 0.215 | 0.212 | 0.142 |
| Phactr1 | 0.215 | 0.382 | 0.407 |
| Rufy1 | 0.215 | 0.119 | 0.109 |
| Timm9 | 0.215 | 0.218 | 0.221 |
| Ecd | 0.214 | 0.190 | 0.168 |
| Ddit4 | 0.214 | 0.258 | 0.127 |
| Zbtb1 | 0.214 | 0.395 | 0.290 |
| Pla2g16 | 0.214 | 0.109 | 0.051 |
| Ndrg2 | 0.214 | 0.915 | 0.625 |
| Pepd | 0.214 | 0.164 | 0.140 |
| Exosc10 | 0.214 | 0.180 | 0.189 |
| Slc35a1 | 0.214 | 0.195 | 0.205 |
| Cmc1 | 0.214 | 0.182 | 0.197 |
| Dnajc4 | 0.214 | 0.181 | 0.163 |
| Pcsk7 | 0.214 | 0.159 | 0.132 |
| Kank3 | 0.214 | 0.039 | 0.041 |
| Stam2 | 0.214 | 0.170 | 0.169 |
| Heatr5a | 0.214 | 0.147 | 0.210 |
| Rev3l | 0.214 | 0.284 | 0.335 |
| Susd6 | 0.214 | 0.159 | 0.157 |
| Smg7 | 0.214 | 0.163 | 0.139 |
| Ocrl | 0.214 | 0.172 | 0.157 |
| Mrps30 | 0.214 | 0.239 | 0.234 |
| Tomm34 | 0.213 | 0.181 | 0.165 |
| Sult1a1 | 0.213 | 0.126 | 0.293 |
| Fads1 | 0.213 | 0.232 | 0.274 |
| Tmod2 | 0.213 | 0.070 | 0.256 |
| Ppie | 0.213 | 0.221 | 0.199 |
| Fam171a1 | 0.213 | 0.138 | 0.253 |
| Vrk3 | 0.213 | 0.121 | 0.161 |
| Ptprg | 0.213 | 0.142 | 0.147 |
| Sympk | 0.213 | 0.181 | 0.192 |
| Micu2 | 0.213 | 0.181 | 0.212 |
| Gab2 | 0.213 | 0.191 | 0.160 |
| Ifngr2 | 0.213 | 0.382 | 0.370 |
| Rtca | 0.213 | 0.174 | 0.193 |
| Prr12 | 0.213 | 0.228 | 0.209 |
| Rftn1 | 0.213 | 0.034 | 0.137 |
| Slc4a2 | 0.213 | 0.199 | 0.188 |
| Ankmy2 | 0.212 | 0.188 | 0.185 |
| Wnk4 | 0.212 | 0.008 | 0.021 |

| | | | |
|---|---|---|---|
| Golga5 | 0.212 | 0.182 | 0.188 |
| Lsg1 | 0.212 | 0.186 | 0.187 |
| Cables1 | 0.212 | 0.168 | 0.128 |
| Tsfm | 0.212 | 0.169 | 0.171 |
| Trps1 | 0.212 | 0.465 | 0.400 |
| Tigd2 | 0.212 | 0.217 | 0.240 |
| Snhg20 | 0.212 | 0.080 | 0.083 |
| Rora | 0.212 | 0.199 | 0.218 |
| Tmem192 | 0.212 | 0.137 | 0.157 |
| Ackr4 | 0.212 | 0.001 | 0.022 |
| Pyurf | 0.212 | 0.151 | 0.160 |
| Pthlh | 0.212 | 0.027 | 0.065 |
| Calcoco1 | 0.212 | 0.261 | 0.208 |
| Ppm1l | 0.212 | 0.309 | 0.056 |
| B3gnt8 | 0.212 | 0.072 | 0.110 |
| Tmem110 | 0.212 | 0.272 | 0.212 |
| Slc12a2 | 0.212 | 0.597 | 0.253 |
| Wars | 0.212 | 0.105 | 0.123 |
| C330007P06F | 0.212 | 0.172 | 0.171 |
| Mrps35 | 0.211 | 0.233 | 0.214 |
| Cdk8 | 0.211 | 0.215 | 0.220 |
| Faim | 0.211 | 0.121 | 0.117 |
| Exosc8 | 0.211 | 0.191 | 0.231 |
| Hipk2 | 0.211 | 0.423 | 0.203 |
| Ing2 | 0.211 | 0.233 | 0.214 |
| Smim26 | 0.211 | 0.206 | 0.139 |
| Jarid2 | 0.211 | 0.158 | 0.165 |
| Scn1b | 0.211 | 0.132 | 0.191 |
| Smyd2 | 0.211 | 0.194 | 0.230 |
| H2-T22 | 0.211 | 0.132 | 0.218 |
| Ccnh | 0.211 | 0.276 | 0.293 |
| Lpar4 | 0.211 | 0.066 | 0.320 |
| Col4a3bp | 0.211 | 0.167 | 0.266 |
| Arhgap23 | 0.210 | 0.157 | 0.140 |
| Trpv4 | 0.210 | 0.005 | 0.006 |
| L3mbtl3 | 0.210 | 0.141 | 0.135 |
| Nubp2 | 0.210 | 0.127 | 0.114 |
| Dlk1 | 0.210 | 0.393 | 2.390 |
| Kat7 | 0.210 | 0.177 | 0.164 |
| Cda | 0.210 | 0.002 | 0.004 |
| Igsf8 | 0.210 | 0.411 | 0.333 |
| Prr14 | 0.210 | 0.206 | 0.162 |
| Tmem87b | 0.210 | 0.301 | 0.245 |
| Pcmtd2 | 0.210 | 0.177 | 0.155 |
| Ptprd | 0.210 | 0.222 | 0.430 |
| Atg4b | 0.210 | 0.171 | 0.176 |
| Rras2 | 0.210 | 0.195 | 0.204 |
| Psmg2 | 0.210 | 0.213 | 0.195 |

| | | | |
|---|---|---|---|
| Ubac1 | 0.209 | 0.173 | 0.201 |
| Zfp397 | 0.209 | 0.196 | 0.213 |
| Klhl2 | 0.209 | 0.096 | 0.112 |
| Mcm3 | 0.209 | 0.116 | 0.176 |
| Cd55 | 0.209 | 0.197 | 0.287 |
| Ankra2 | 0.209 | 0.208 | 0.238 |
| Tfe3 | 0.209 | 0.256 | 0.251 |
| Dlg1 | 0.209 | 0.171 | 0.171 |
| Myo6 | 0.209 | 0.130 | 0.148 |
| Dnajc13 | 0.209 | 0.167 | 0.183 |
| Wdr83 | 0.209 | 0.147 | 0.132 |
| Cadps2 | 0.209 | 0.005 | 0.008 |
| Xab2 | 0.209 | 0.134 | 0.136 |
| Map4k2 | 0.209 | 0.054 | 0.049 |
| Brcc3 | 0.209 | 0.174 | 0.138 |
| Il3ra | 0.209 | 0.058 | 0.148 |
| Gxylt1 | 0.209 | 0.147 | 0.186 |
| Dars | 0.209 | 0.239 | 0.223 |
| Rufy3 | 0.208 | 0.132 | 0.133 |
| Abcf3 | 0.208 | 0.110 | 0.143 |
| Crebzf | 0.208 | 0.164 | 0.188 |
| Nrbf2 | 0.208 | 0.250 | 0.228 |
| Ppp2r3a | 0.208 | 0.234 | 0.259 |
| Cars2 | 0.208 | 0.163 | 0.166 |
| Slc25a46 | 0.208 | 0.190 | 0.185 |
| Qser1 | 0.208 | 0.228 | 0.251 |
| Tmem29 | 0.207 | 0.183 | 0.212 |
| Mospd1 | 0.207 | 0.189 | 0.155 |
| Mpv17l2 | 0.207 | 0.230 | 0.188 |
| Rad50 | 0.207 | 0.208 | 0.251 |
| Pdzd2 | 0.207 | 0.058 | 0.064 |
| Ten1 | 0.207 | 0.230 | 0.320 |
| 9130401M01 | 0.207 | 0.278 | 0.301 |
| Opa1 | 0.207 | 0.176 | 0.166 |
| Scaf8 | 0.207 | 0.202 | 0.168 |
| Mpp5 | 0.207 | 0.144 | 0.139 |
| Usp48 | 0.207 | 0.214 | 0.251 |
| Mycbp | 0.207 | 0.158 | 0.141 |
| Mapre3 | 0.207 | 0.148 | 0.137 |
| Nfkbil1 | 0.207 | 0.199 | 0.205 |
| Cabin1 | 0.207 | 0.331 | 0.281 |
| Gpr146 | 0.206 | 0.191 | 0.077 |
| Slc30a7 | 0.206 | 0.143 | 0.161 |
| Ipo5 | 0.206 | 0.288 | 0.290 |
| Fosl2 | 0.206 | 0.489 | 0.375 |
| Atad1 | 0.206 | 0.253 | 0.186 |
| Tmem242 | 0.206 | 0.197 | 0.210 |
| Adgrg6 | 0.206 | 0.044 | 0.016 |

| | | | |
|---|---|---|---|
| Ppil1 | 0.206 | 0.136 | 0.131 |
| Sertad4 | 0.206 | 0.129 | 0.246 |
| Lyve1 | 0.206 | 0.048 | 0.019 |
| Map4k3 | 0.206 | 0.143 | 0.138 |
| Pxmp4 | 0.206 | 0.178 | 0.179 |
| Phrf1 | 0.206 | 0.193 | 0.178 |
| Taf1 | 0.206 | 0.214 | 0.208 |
| 1500011B03F | 0.206 | 0.079 | 0.090 |
| Lum | 0.206 | 0.402 | 2.360 |
| Rabep2 | 0.206 | 0.110 | 0.105 |
| Nat9 | 0.206 | 0.126 | 0.141 |
| Fam49b | 0.206 | 0.219 | 0.150 |
| Mzt2 | 0.206 | 0.252 | 0.243 |
| Hmgcs1 | 0.206 | 0.312 | 0.444 |
| Ccdc90b | 0.206 | 0.228 | 0.252 |
| Qpct | 0.205 | 0.492 | 0.875 |
| Tcp11l2 | 0.205 | 0.258 | 0.271 |
| Pank3 | 0.205 | 0.168 | 0.167 |
| Maml2 | 0.205 | 0.226 | 0.199 |
| Suco | 0.205 | 0.212 | 0.180 |
| Arhgef5 | 0.205 | 0.155 | 0.042 |
| Meaf6 | 0.205 | 0.233 | 0.230 |
| Prkd2 | 0.205 | 0.075 | 0.090 |
| Bloc1s2 | 0.205 | 0.108 | 0.164 |
| Uba3 | 0.205 | 0.203 | 0.180 |
| Tmem246 | 0.205 | 0.302 | 0.200 |
| Btrc | 0.205 | 0.160 | 0.171 |
| Atp8b1 | 0.205 | 0.157 | 0.076 |
| Mum1l1 | 0.205 | 0.324 | 0.194 |
| Rab4a | 0.205 | 0.210 | 0.136 |
| Sort1 | 0.205 | 0.100 | 0.087 |
| Mfsd5 | 0.204 | 0.148 | 0.184 |
| Sema4d | 0.204 | 0.012 | 0.006 |
| Stk40 | 0.204 | 0.313 | 0.208 |
| Letm1 | 0.204 | 0.203 | 0.177 |
| Carm1 | 0.204 | 0.189 | 0.270 |
| Senp5 | 0.204 | 0.165 | 0.159 |
| Heatr6 | 0.204 | 0.197 | 0.228 |
| Kpna1 | 0.204 | 0.207 | 0.139 |
| Top3b | 0.204 | 0.154 | 0.170 |
| Eps15l1 | 0.204 | 0.151 | 0.146 |
| Vps37a | 0.204 | 0.198 | 0.149 |
| 2610008E11F | 0.204 | 0.116 | 0.116 |
| Atp11b | 0.204 | 0.244 | 0.173 |
| Ccpg1 | 0.204 | 0.155 | 0.185 |
| Avl9 | 0.204 | 0.223 | 0.157 |
| Cachd1 | 0.204 | 0.154 | 0.133 |
| Wdr13 | 0.203 | 0.167 | 0.147 |

| | | | |
|---|---|---|---|
| Tpgs2 | 0.203 | 0.140 | 0.192 |
| Zkscan1 | 0.203 | 0.195 | 0.175 |
| Samd14 | 0.203 | 0.219 | 0.295 |
| Sik3 | 0.203 | 0.244 | 0.247 |
| Nacc2 | 0.203 | 0.302 | 0.138 |
| Anapc10 | 0.203 | 0.191 | 0.178 |
| Tmem41b | 0.203 | 0.223 | 0.253 |
| Phkb | 0.203 | 0.172 | 0.163 |
| Sh2b1 | 0.203 | 0.153 | 0.177 |
| Mical1 | 0.203 | 0.323 | 0.237 |
| Xpo6 | 0.203 | 0.198 | 0.192 |
| Fam136a | 0.203 | 0.142 | 0.160 |
| Me1 | 0.203 | 0.246 | 0.269 |
| Haghl | 0.203 | 0.214 | 0.188 |
| Nkx2-3 | 0.203 | 0.002 | 0.001 |
| Stxbp5 | 0.203 | 0.089 | 0.094 |
| Bsdc1 | 0.203 | 0.156 | 0.139 |
| Kif13a | 0.202 | 0.158 | 0.145 |
| Vmac | 0.202 | 0.130 | 0.134 |
| Gclc | 0.202 | 0.281 | 0.206 |
| Lmnb1 | 0.202 | 0.119 | 0.141 |
| Uchl5 | 0.202 | 0.188 | 0.140 |
| Topbp1 | 0.202 | 0.133 | 0.166 |
| Stambp | 0.202 | 0.124 | 0.149 |
| Nbl1 | 0.202 | 0.861 | 0.905 |
| Tyms | 0.202 | 0.113 | 0.139 |
| Spast | 0.202 | 0.195 | 0.194 |
| Mitd1 | 0.202 | 0.099 | 0.109 |
| Dcaf17 | 0.202 | 0.131 | 0.139 |
| 1110032A03R | 0.202 | 0.169 | 0.191 |
| Auh | 0.202 | 0.363 | 0.281 |
| Mink1 | 0.202 | 0.134 | 0.111 |
| Rap1gds1 | 0.202 | 0.176 | 0.165 |
| Mical3 | 0.202 | 0.255 | 0.232 |
| Mmgt2 | 0.202 | 0.183 | 0.264 |
| Mphosph10 | 0.201 | 0.188 | 0.161 |
| Cdkn2aipnl | 0.201 | 0.091 | 0.091 |
| Pcdh19 | 0.201 | 0.012 | 0.230 |
| Gpc3 | 0.201 | 1.149 | 1.761 |
| Adamts2 | 0.201 | 1.312 | 1.507 |
| Smim8 | 0.201 | 0.180 | 0.191 |
| Kdm6a | 0.201 | 0.180 | 0.143 |
| Phf12 | 0.201 | 0.234 | 0.181 |
| Ppm1m | 0.201 | 0.136 | 0.162 |
| Rmnd5b | 0.201 | 0.161 | 0.153 |
| Sppl2b | 0.201 | 0.109 | 0.124 |
| 2510009E07R | 0.201 | 0.295 | 0.210 |
| Usp25 | 0.201 | 0.178 | 0.168 |

| | | | |
|---|---|---|---|
| Msmo1 | 0.201 | 0.194 | 0.251 |
| Nup50 | 0.201 | 0.161 | 0.160 |
| Eif2d | 0.201 | 0.125 | 0.138 |
| Med6 | 0.201 | 0.163 | 0.150 |
| Zfp800 | 0.201 | 0.189 | 0.177 |
| Prom1 | 0.201 | 0.003 | 0.004 |
| Myg1 | 0.200 | 0.187 | 0.198 |
| Poglut1 | 0.200 | 0.164 | 0.158 |
| Myef2 | 0.200 | 0.174 | 0.177 |
| Ago3 | 0.200 | 0.166 | 0.157 |
| Inpp5a | 0.200 | 0.377 | 0.226 |
| Pex14 | 0.200 | 0.170 | 0.187 |
| Klhdc10 | 0.200 | 0.155 | 0.211 |
| Plcd1 | 0.200 | 0.218 | 0.165 |
| Csrnp1 | 0.200 | 0.596 | 0.373 |
| Zc3hav1 | 0.200 | 0.055 | 0.112 |
| Nxnl2 | 0.200 | 0.020 | 0.021 |
| Zfp318 | 0.200 | 0.222 | 0.247 |
| Slc39a10 | 0.200 | 0.084 | 0.114 |
| Gopc | 0.200 | 0.146 | 0.131 |
| Sertad3 | 0.200 | 0.213 | 0.195 |
| Nubp1 | 0.200 | 0.157 | 0.116 |
| Bach1 | 0.200 | 0.314 | 0.291 |
| Ipo4 | 0.200 | 0.176 | 0.248 |
| Hspa1a | 0.200 | 0.313 | 0.192 |
| Dennd3 | 0.199 | 0.033 | 0.029 |
| Rhno1 | 0.199 | 0.157 | 0.187 |
| Eif2b4 | 0.199 | 0.196 | 0.179 |
| Psmb8 | 0.199 | 0.036 | 0.056 |
| Tada3 | 0.199 | 0.204 | 0.220 |
| Pcgf2 | 0.199 | 0.196 | 0.208 |
| Coq2 | 0.199 | 0.210 | 0.194 |
| Bloc1s4 | 0.199 | 0.157 | 0.174 |
| Txndc11 | 0.199 | 0.234 | 0.217 |
| Cd276 | 0.199 | 0.281 | 0.288 |
| Rnf26 | 0.199 | 0.199 | 0.231 |
| Pgs1 | 0.199 | 0.189 | 0.151 |
| Ints6 | 0.199 | 0.240 | 0.236 |
| Ctsf | 0.199 | 0.209 | 0.309 |
| Lpcat1 | 0.199 | 0.115 | 0.171 |
| Thap2 | 0.199 | 0.215 | 0.220 |
| Leo1 | 0.199 | 0.194 | 0.199 |
| Aldh7a1 | 0.199 | 0.388 | 0.380 |
| Zbtb22 | 0.199 | 0.168 | 0.134 |
| Pdcd11 | 0.199 | 0.175 | 0.193 |
| Cebpb | 0.199 | 1.438 | 0.891 |
| Ttc33 | 0.199 | 0.165 | 0.154 |
| Cldn7 | 0.199 | 0.017 | 0.027 |

| | | | |
|---|---|---|---|
| Gprin3 | 0.199 | 0.043 | 0.012 |
| St7l | 0.199 | 0.173 | 0.172 |
| Dstyk | 0.199 | 0.226 | 0.207 |
| Map4k5 | 0.198 | 0.237 | 0.251 |
| B4galt3 | 0.198 | 0.165 | 0.169 |
| Apaf1 | 0.198 | 0.100 | 0.140 |
| Mrpl16 | 0.198 | 0.185 | 0.163 |
| Ppm1b | 0.198 | 0.300 | 0.234 |
| Mrps31 | 0.198 | 0.185 | 0.142 |
| Parp1 | 0.198 | 0.208 | 0.200 |
| Zfand2b | 0.198 | 0.180 | 0.168 |
| Pim1 | 0.198 | 0.365 | 0.395 |
| Ifi35 | 0.198 | 0.157 | 0.143 |
| Eps15 | 0.198 | 0.208 | 0.256 |
| Map3k10 | 0.198 | 0.201 | 0.179 |
| Bnip3 | 0.198 | 0.325 | 0.195 |
| Reps1 | 0.198 | 0.213 | 0.246 |
| Gsr | 0.198 | 0.319 | 0.245 |
| Atf6b | 0.198 | 0.200 | 0.153 |
| Abcg2 | 0.198 | 0.080 | 0.183 |
| Slc35f5 | 0.197 | 0.180 | 0.289 |
| Farp1 | 0.197 | 0.264 | 0.266 |
| Scarf1 | 0.197 | 0.003 | 0.002 |
| Stx17 | 0.197 | 0.179 | 0.187 |
| Lonp2 | 0.197 | 0.220 | 0.212 |
| Hdhd2 | 0.197 | 0.168 | 0.173 |
| Dok4 | 0.197 | 0.058 | 0.047 |
| Gas1 | 0.197 | 0.490 | 1.321 |
| Nrf1 | 0.197 | 0.164 | 0.175 |
| Ndufaf4 | 0.197 | 0.149 | 0.146 |
| Mfsd14b | 0.197 | 0.182 | 0.170 |
| Pmpca | 0.197 | 0.143 | 0.141 |
| Agap3 | 0.197 | 0.197 | 0.245 |
| Nol11 | 0.197 | 0.204 | 0.190 |
| Uhmk1 | 0.197 | 0.088 | 0.083 |
| Nup214 | 0.197 | 0.167 | 0.154 |
| Fbxw7 | 0.197 | 0.170 | 0.124 |
| Dusp12 | 0.197 | 0.164 | 0.189 |
| Pard3b | 0.197 | 0.176 | 0.245 |
| Ptbp2 | 0.197 | 0.284 | 0.212 |
| Mettl26 | 0.197 | 0.219 | 0.171 |
| Alg14 | 0.197 | 0.152 | 0.133 |
| Ddit3 | 0.197 | 0.428 | 0.453 |
| Snx27 | 0.196 | 0.182 | 0.218 |
| Rnf141 | 0.196 | 0.069 | 0.062 |
| Hbb-bt | 0.196 | 0.164 | 0.191 |
| 1810013L24R | 0.196 | 0.181 | 0.169 |
| Trp53inp1 | 0.196 | 0.551 | 0.251 |

| | | | |
|---|---|---|---|
| Parp8 | 0.196 | 0.112 | 0.104 |
| Sh3bp5 | 0.196 | 0.022 | 0.280 |
| Jmjd8 | 0.196 | 0.106 | 0.118 |
| Pelp1 | 0.196 | 0.168 | 0.175 |
| Bola1 | 0.196 | 0.179 | 0.202 |
| Zfp830 | 0.196 | 0.187 | 0.160 |
| Scamp1 | 0.196 | 0.120 | 0.091 |
| B9d2 | 0.196 | 0.175 | 0.145 |
| Eif2b1 | 0.196 | 0.169 | 0.163 |
| Tmem43 | 0.196 | 0.242 | 0.242 |
| Tmem63a | 0.196 | 0.140 | 0.101 |
| Ldhb | 0.195 | 0.982 | 0.423 |
| Pigq | 0.195 | 0.101 | 0.102 |
| Plcg2 | 0.195 | 0.006 | 0.003 |
| Slc35e4 | 0.195 | 0.182 | 0.081 |
| Xrcc1 | 0.195 | 0.164 | 0.230 |
| Rpain | 0.195 | 0.220 | 0.172 |
| Rarg | 0.195 | 0.199 | 0.245 |
| Cyb561d2 | 0.195 | 0.139 | 0.131 |
| Zfx | 0.195 | 0.226 | 0.217 |
| Fnip1 | 0.195 | 0.287 | 0.324 |
| Nsmf | 0.195 | 0.212 | 0.233 |
| Cbfb | 0.195 | 0.185 | 0.285 |
| Cdkn2aip | 0.195 | 0.269 | 0.242 |
| Vangl1 | 0.195 | 0.092 | 0.145 |
| Wdr48 | 0.195 | 0.150 | 0.159 |
| Ltn1 | 0.195 | 0.191 | 0.172 |
| Paf1 | 0.195 | 0.241 | 0.186 |
| Tmem237 | 0.195 | 0.197 | 0.222 |
| 2310022B05F | 0.194 | 0.515 | 0.501 |
| Sephs1 | 0.194 | 0.198 | 0.198 |
| Ddi2 | 0.194 | 0.244 | 0.173 |
| Rnf41 | 0.194 | 0.109 | 0.100 |
| Pofut1 | 0.194 | 0.138 | 0.227 |
| Mapk9 | 0.194 | 0.146 | 0.164 |
| Skiv2l | 0.194 | 0.165 | 0.159 |
| Erap1 | 0.194 | 0.086 | 0.063 |
| AU019823 | 0.194 | 0.165 | 0.182 |
| Rbm10 | 0.194 | 0.164 | 0.164 |
| Mtpap | 0.194 | 0.175 | 0.172 |
| Uimc1 | 0.194 | 0.191 | 0.161 |
| Ppan | 0.194 | 0.142 | 0.162 |
| Rnf103 | 0.194 | 0.234 | 0.185 |
| Emc1 | 0.194 | 0.124 | 0.111 |
| Abhd14b | 0.194 | 0.175 | 0.211 |
| Uxs1 | 0.194 | 0.149 | 0.143 |
| Slc4a7 | 0.194 | 0.232 | 0.157 |
| Apool | 0.194 | 0.186 | 0.180 |

| | | | |
|---|---|---|---|
| Farsb | 0.194 | 0.156 | 0.162 |
| Mtf1 | 0.194 | 0.113 | 0.094 |
| Ube2l6 | 0.193 | 0.098 | 0.174 |
| Heca | 0.193 | 0.239 | 0.245 |
| Bphl | 0.193 | 0.188 | 0.184 |
| Ilvbl | 0.193 | 0.279 | 0.251 |
| Kat5 | 0.193 | 0.162 | 0.149 |
| Ccng2 | 0.193 | 0.174 | 0.184 |
| Iffo2 | 0.193 | 0.088 | 0.096 |
| Trim56 | 0.193 | 0.141 | 0.143 |
| Rnf214 | 0.193 | 0.228 | 0.193 |
| Clpx | 0.193 | 0.252 | 0.224 |
| Abcc5 | 0.192 | 0.279 | 0.252 |
| Ralgds | 0.192 | 0.171 | 0.254 |
| Rtkn | 0.192 | 0.080 | 0.140 |
| Ctso | 0.192 | 0.170 | 0.242 |
| Pou2f1 | 0.192 | 0.134 | 0.111 |
| Atp7a | 0.192 | 0.104 | 0.120 |
| Ebf1 | 0.192 | 0.407 | 1.175 |
| Rbfa | 0.192 | 0.273 | 0.267 |
| Dot1l | 0.192 | 0.248 | 0.220 |
| Noc2l | 0.192 | 0.150 | 0.211 |
| Ints11 | 0.192 | 0.118 | 0.146 |
| C2cd2 | 0.192 | 0.165 | 0.205 |
| Fam120b | 0.192 | 0.139 | 0.158 |
| Clip3 | 0.191 | 0.399 | 0.430 |
| Dll1 | 0.191 | 0.009 | 0.005 |
| Rrp15 | 0.191 | 0.206 | 0.207 |
| A430005L14R | 0.191 | 0.158 | 0.170 |
| Cntln | 0.191 | 0.218 | 0.271 |
| Iars | 0.191 | 0.143 | 0.178 |
| Nf1 | 0.191 | 0.269 | 0.256 |
| Hmbox1 | 0.191 | 0.164 | 0.130 |
| Ankrd50 | 0.191 | 0.148 | 0.157 |
| Iars2 | 0.191 | 0.201 | 0.162 |
| Ltbp2 | 0.191 | 0.846 | 0.779 |
| Pdgfrl | 0.191 | 0.800 | 0.711 |
| Osmr | 0.190 | 0.328 | 0.301 |
| Bicd2 | 0.190 | 0.244 | 0.254 |
| Cdc23 | 0.190 | 0.189 | 0.202 |
| Fam234a | 0.190 | 0.174 | 0.172 |
| Prr15 | 0.190 | 0.009 | 0.006 |
| Homer3 | 0.190 | 0.116 | 0.161 |
| Fam214b | 0.190 | 0.151 | 0.098 |
| Chfr | 0.190 | 0.175 | 0.148 |
| Cfap97 | 0.190 | 0.159 | 0.166 |
| Fkrp | 0.190 | 0.158 | 0.111 |
| Ier5l | 0.190 | 0.688 | 0.407 |

| | | | |
|---|---|---|---|
| Rab33b | 0.190 | 0.140 | 0.156 |
| Khsrp | 0.190 | 0.226 | 0.202 |
| Ezh1 | 0.190 | 0.099 | 0.141 |
| Zfp410 | 0.190 | 0.168 | 0.135 |
| Kctd5 | 0.190 | 0.109 | 0.122 |
| Zdhhc7 | 0.190 | 0.105 | 0.114 |
| Rin2 | 0.190 | 0.341 | 0.244 |
| Dctpp1 | 0.190 | 0.173 | 0.209 |
| Gpcpd1 | 0.190 | 0.210 | 0.205 |
| Numbl | 0.189 | 0.268 | 0.390 |
| Med7 | 0.189 | 0.192 | 0.188 |
| Tmem50b | 0.189 | 0.173 | 0.116 |
| Amdhd2 | 0.189 | 0.142 | 0.145 |
| Nrbp2 | 0.189 | 0.254 | 0.174 |
| Fbrs | 0.189 | 0.157 | 0.153 |
| Fam207a | 0.189 | 0.159 | 0.157 |
| Srebf1 | 0.189 | 0.162 | 0.132 |
| Mtap | 0.189 | 0.204 | 0.201 |
| Aars | 0.189 | 0.217 | 0.236 |
| Abhd5 | 0.189 | 0.142 | 0.190 |
| Zfp598 | 0.189 | 0.138 | 0.135 |
| Samd5 | 0.189 | 0.164 | 0.144 |
| Bambi | 0.189 | 0.541 | 0.392 |
| Gins4 | 0.189 | 0.134 | 0.171 |
| Nlk | 0.189 | 0.096 | 0.066 |
| Seh1l | 0.189 | 0.178 | 0.159 |
| Slc5a3 | 0.189 | 0.367 | 0.200 |
| Aimp2 | 0.188 | 0.207 | 0.179 |
| Clcc1 | 0.188 | 0.241 | 0.226 |
| Stat6 | 0.188 | 0.146 | 0.172 |
| Serpine2 | 0.188 | 1.594 | 0.553 |
| Unc45a | 0.188 | 0.295 | 0.185 |
| Gbp9 | 0.188 | 0.009 | 0.015 |
| Fyco1 | 0.188 | 0.216 | 0.176 |
| Timm8a1 | 0.188 | 0.184 | 0.195 |
| Trip10 | 0.188 | 0.193 | 0.228 |
| Sesn3 | 0.188 | 0.528 | 0.277 |
| Sun2 | 0.188 | 0.448 | 0.153 |
| Ints13 | 0.188 | 0.153 | 0.130 |
| Stard7 | 0.188 | 0.149 | 0.172 |
| Slc25a16 | 0.188 | 0.161 | 0.125 |
| Zfp651 | 0.188 | 0.347 | 0.387 |
| Il4ra | 0.187 | 0.043 | 0.096 |
| Mcu | 0.187 | 0.091 | 0.137 |
| Zfp740 | 0.187 | 0.237 | 0.212 |
| Nelfa | 0.187 | 0.160 | 0.169 |
| Pias4 | 0.187 | 0.178 | 0.166 |
| Zdhhc21 | 0.187 | 0.143 | 0.135 |

| | | | |
|---|---|---|---|
| Secisbp2 | 0.187 | 0.110 | 0.127 |
| Dyrk1a | 0.187 | 0.249 | 0.191 |
| 5730455P16F | 0.187 | 0.173 | 0.171 |
| Fbxl17 | 0.187 | 0.174 | 0.188 |
| Wdr44 | 0.187 | 0.114 | 0.128 |
| Cars | 0.187 | 0.164 | 0.212 |
| Per3 | 0.187 | 0.122 | 0.143 |
| Psmg3 | 0.187 | 0.131 | 0.124 |
| Naa60 | 0.187 | 0.084 | 0.122 |
| Nr2c2ap | 0.187 | 0.136 | 0.113 |
| Fbxo6 | 0.187 | 0.105 | 0.103 |
| Fzr1 | 0.187 | 0.143 | 0.152 |
| Epdr1 | 0.187 | 0.142 | 0.104 |
| Tmem42 | 0.187 | 0.130 | 0.158 |
| Psmd10 | 0.187 | 0.169 | 0.153 |
| Gfer | 0.187 | 0.134 | 0.157 |
| F2rl1 | 0.187 | 0.068 | 0.008 |
| Ubr7 | 0.187 | 0.120 | 0.113 |
| Hoxb2 | 0.187 | 0.408 | 0.255 |
| Snx24 | 0.187 | 0.165 | 0.174 |
| Hmgcr | 0.187 | 0.154 | 0.217 |
| Pcnx3 | 0.187 | 0.158 | 0.155 |
| Mad2l1bp | 0.187 | 0.112 | 0.092 |
| Coq9 | 0.187 | 0.176 | 0.166 |
| Prmt2 | 0.187 | 0.161 | 0.227 |
| Pgm3 | 0.187 | 0.158 | 0.288 |
| Iigp1 | 0.187 | 0.180 | 0.202 |
| Rasl11a | 0.187 | 0.713 | 0.353 |
| Fem1b | 0.187 | 0.177 | 0.183 |
| Cdk1 | 0.187 | 0.097 | 0.099 |
| Pik3ip1 | 0.186 | 0.282 | 0.241 |
| Smarcd3 | 0.186 | 0.420 | 0.230 |
| Bcat2 | 0.186 | 0.178 | 0.172 |
| Acp2 | 0.186 | 0.167 | 0.174 |
| Cpsf3 | 0.186 | 0.154 | 0.193 |
| Scnm1 | 0.186 | 0.177 | 0.183 |
| Gm15222 | 0.186 | 0.025 | 0.030 |
| Sox5 | 0.186 | 0.037 | 0.020 |
| Brf1 | 0.186 | 0.143 | 0.142 |
| Fam149b | 0.186 | 0.165 | 0.169 |
| Mospd3 | 0.186 | 0.145 | 0.179 |
| Cep63 | 0.186 | 0.146 | 0.146 |
| C1galt1c1 | 0.186 | 0.123 | 0.136 |
| Surf2 | 0.186 | 0.195 | 0.161 |
| Slfn2 | 0.186 | 0.012 | 0.013 |
| Cep250 | 0.186 | 0.104 | 0.103 |
| Rapgef3 | 0.186 | 0.016 | 0.040 |
| Pisd | 0.186 | 0.171 | 0.182 |

| | | | |
|---|---|---|---|
| Gm8186 | 0.186 | 0.141 | 0.172 |
| Smc5 | 0.186 | 0.214 | 0.219 |
| 0610030E20F | 0.186 | 0.169 | 0.204 |
| Dlat | 0.185 | 0.214 | 0.180 |
| Errfi1 | 0.185 | 0.467 | 0.505 |
| Pccb | 0.185 | 0.176 | 0.191 |
| Arid5a | 0.185 | 0.488 | 0.437 |
| Nmi | 0.185 | 0.102 | 0.116 |
| Mark4 | 0.185 | 0.136 | 0.176 |
| Gdpd1 | 0.185 | 0.117 | 0.074 |
| Bloc1s6 | 0.185 | 0.114 | 0.134 |
| Gripap1 | 0.185 | 0.159 | 0.168 |
| Nxt1 | 0.185 | 0.143 | 0.130 |
| Vcpip1 | 0.185 | 0.202 | 0.210 |
| Dcp2 | 0.185 | 0.099 | 0.115 |
| Trap1 | 0.185 | 0.178 | 0.173 |
| Rnf217 | 0.185 | 0.212 | 0.170 |
| Fbxo8 | 0.185 | 0.161 | 0.133 |
| Atl3 | 0.185 | 0.206 | 0.153 |
| Hat1 | 0.185 | 0.219 | 0.200 |
| Kank2 | 0.185 | 0.383 | 0.477 |
| Msl2 | 0.185 | 0.161 | 0.145 |
| Pcnt | 0.185 | 0.194 | 0.127 |
| Med16 | 0.185 | 0.192 | 0.191 |
| Bag3 | 0.185 | 0.434 | 0.410 |
| Vegfb | 0.184 | 0.289 | 0.348 |
| Tjp2 | 0.184 | 0.055 | 0.052 |
| Mrps18b | 0.184 | 0.189 | 0.182 |
| Nup88 | 0.184 | 0.189 | 0.151 |
| Dgkz | 0.184 | 0.129 | 0.091 |
| Tbc1d9b | 0.184 | 0.154 | 0.211 |
| Casc4 | 0.184 | 0.338 | 0.310 |
| Ppp1r15b | 0.184 | 0.161 | 0.174 |
| Paxbp1 | 0.184 | 0.195 | 0.160 |
| Itfg2 | 0.184 | 0.168 | 0.161 |
| Commd10 | 0.184 | 0.138 | 0.173 |
| Usp33 | 0.184 | 0.189 | 0.199 |
| Gata3 | 0.184 | 0.038 | 0.016 |
| Exosc9 | 0.184 | 0.156 | 0.115 |
| Lmo7 | 0.184 | 0.066 | 0.034 |
| Mospd2 | 0.184 | 0.127 | 0.146 |
| Orc4 | 0.184 | 0.223 | 0.180 |
| Atad2 | 0.184 | 0.198 | 0.218 |
| Cd320 | 0.184 | 0.153 | 0.218 |
| Scai | 0.184 | 0.185 | 0.189 |
| Klhl7 | 0.184 | 0.186 | 0.160 |
| Epb41 | 0.184 | 0.123 | 0.160 |
| Ube2q2 | 0.184 | 0.173 | 0.150 |

| | | | |
|---|---|---|---|
| Uap1l1 | 0.184 | 0.138 | 0.185 |
| Slc25a28 | 0.184 | 0.180 | 0.181 |
| Sema3g | 0.184 | 0.003 | 0.006 |
| B4galt7 | 0.184 | 0.106 | 0.134 |
| Ints6l | 0.184 | 0.249 | 0.156 |
| Cenpa | 0.184 | 0.109 | 0.090 |
| Gprc5a | 0.183 | 0.022 | 0.015 |
| Trip4 | 0.183 | 0.195 | 0.192 |
| Rrn3 | 0.183 | 0.180 | 0.140 |
| Dmtf1 | 0.183 | 0.167 | 0.185 |
| Kif3a | 0.183 | 0.275 | 0.330 |
| Adsl | 0.183 | 0.154 | 0.145 |
| Nav2 | 0.183 | 0.696 | 0.198 |
| Rint1 | 0.183 | 0.159 | 0.186 |
| Gnl3l | 0.183 | 0.175 | 0.151 |
| Trak2 | 0.183 | 0.126 | 0.111 |
| Gm5617 | 0.183 | 0.138 | 0.147 |
| Pgpep1 | 0.183 | 0.168 | 0.170 |
| Ppp2r5b | 0.183 | 0.111 | 0.146 |
| Zpr1 | 0.183 | 0.178 | 0.212 |
| Hdac4 | 0.183 | 0.289 | 0.250 |
| Alcam | 0.183 | 0.646 | 0.431 |
| Slc43a1 | 0.183 | 0.097 | 0.119 |
| Irf9 | 0.183 | 0.113 | 0.141 |
| Tmem39a | 0.183 | 0.178 | 0.215 |
| Sdk1 | 0.183 | 0.034 | 0.163 |
| Tfeb | 0.183 | 0.100 | 0.067 |
| Nat6 | 0.183 | 0.135 | 0.135 |
| Ddx56 | 0.182 | 0.136 | 0.166 |
| Fam129a | 0.182 | 0.539 | 0.621 |
| Hebp1 | 0.182 | 0.225 | 0.320 |
| Tmem126b | 0.182 | 0.150 | 0.136 |
| Syap1 | 0.182 | 0.206 | 0.192 |
| Wdr37 | 0.182 | 0.090 | 0.091 |
| Pstk | 0.182 | 0.203 | 0.181 |
| Gnpat | 0.182 | 0.142 | 0.139 |
| Sele | 0.182 | 0.037 | 0.092 |
| Tmem167b | 0.182 | 0.171 | 0.186 |
| Ndufaf2 | 0.182 | 0.233 | 0.226 |
| Ice1 | 0.182 | 0.193 | 0.202 |
| Tctex1d2 | 0.182 | 0.136 | 0.142 |
| Rdh14 | 0.182 | 0.261 | 0.284 |
| Kctd12b | 0.182 | 0.053 | 0.082 |
| Pja1 | 0.182 | 0.270 | 0.385 |
| Rgmb | 0.182 | 0.145 | 0.125 |
| Mboat7 | 0.182 | 0.243 | 0.221 |
| Dmtn | 0.182 | 0.009 | 0.015 |
| Desi1 | 0.182 | 0.121 | 0.146 |

| | | | |
|---|---|---|---|
| Ing3 | 0.182 | 0.174 | 0.174 |
| Col4a2 | 0.182 | 1.406 | 1.177 |
| Magohb | 0.182 | 0.206 | 0.173 |
| Ikzf5 | 0.181 | 0.172 | 0.155 |
| Pcdh9 | 0.181 | 0.084 | 0.231 |
| Tbk1 | 0.181 | 0.135 | 0.120 |
| Uckl1 | 0.181 | 0.156 | 0.101 |
| Apba1 | 0.181 | 0.021 | 0.065 |
| Ctnnal1 | 0.181 | 0.088 | 0.070 |
| Lancl1 | 0.181 | 0.230 | 0.252 |
| Rusc1 | 0.181 | 0.104 | 0.101 |
| Gabpb1 | 0.181 | 0.098 | 0.080 |
| Ankfy1 | 0.181 | 0.127 | 0.160 |
| Rab11fip2 | 0.181 | 0.279 | 0.300 |
| Spopl | 0.181 | 0.164 | 0.183 |
| Ino80 | 0.181 | 0.169 | 0.157 |
| Snrpn | 0.181 | 0.122 | 0.100 |
| Terf2ip | 0.181 | 0.142 | 0.132 |
| Cacul1 | 0.181 | 0.141 | 0.125 |
| Coasy | 0.181 | 0.130 | 0.140 |
| Gstz1 | 0.181 | 0.141 | 0.167 |
| Pde1a | 0.181 | 0.748 | 0.385 |
| 1810030O07I | 0.181 | 0.117 | 0.132 |
| Tubb2b | 0.181 | 0.102 | 0.409 |
| Tgif1 | 0.181 | 0.175 | 0.143 |
| Rpf1 | 0.181 | 0.171 | 0.145 |
| Magix | 0.181 | 0.001 | 0.001 |
| Dexi | 0.180 | 0.259 | 0.285 |
| Chchd5 | 0.180 | 0.121 | 0.143 |
| Mafk | 0.180 | 0.302 | 0.182 |
| Nom1 | 0.180 | 0.204 | 0.189 |
| Ibtk | 0.180 | 0.146 | 0.219 |
| Slf2 | 0.180 | 0.145 | 0.174 |
| Nin | 0.180 | 0.136 | 0.156 |
| Mtmr3 | 0.180 | 0.174 | 0.173 |
| Ptcd2 | 0.180 | 0.131 | 0.152 |
| Polr2b | 0.180 | 0.177 | 0.178 |
| Tbc1d10a | 0.180 | 0.152 | 0.195 |
| Eva1c | 0.180 | 0.158 | 0.067 |
| Rabif | 0.180 | 0.160 | 0.170 |
| Tubgcp2 | 0.180 | 0.168 | 0.194 |
| Dmap1 | 0.180 | 0.190 | 0.162 |
| Rab2b | 0.179 | 0.168 | 0.159 |
| Plscr4 | 0.179 | 0.048 | 0.183 |
| Ncoa2 | 0.179 | 0.214 | 0.244 |
| Fam122a | 0.179 | 0.188 | 0.191 |
| Osbpl2 | 0.179 | 0.161 | 0.173 |
| Parp2 | 0.179 | 0.139 | 0.150 |

| | | | |
|---|---|---|---|
| Nfs1 | 0.179 | 0.140 | 0.128 |
| Pvr | 0.179 | 0.281 | 0.145 |
| Med20 | 0.179 | 0.107 | 0.118 |
| 1010001N08I | 0.179 | 0.085 | 0.059 |
| Msx1 | 0.179 | 0.027 | 0.029 |
| Tmem138 | 0.179 | 0.134 | 0.149 |
| Tfam | 0.179 | 0.175 | 0.186 |
| Nradd | 0.179 | 0.213 | 0.118 |
| Kmt2b | 0.179 | 0.197 | 0.163 |
| Arl13b | 0.179 | 0.249 | 0.221 |
| Ppp4r1 | 0.179 | 0.132 | 0.130 |
| Bms1 | 0.179 | 0.187 | 0.192 |
| Bin3 | 0.178 | 0.086 | 0.134 |
| Aldh18a1 | 0.178 | 0.166 | 0.242 |
| Cdc42ep2 | 0.178 | 0.068 | 0.472 |
| Gatd1 | 0.178 | 0.251 | 0.369 |
| Lsm14b | 0.178 | 0.138 | 0.122 |
| Dennd1b | 0.178 | 0.123 | 0.136 |
| Crot | 0.178 | 0.177 | 0.170 |
| Spred1 | 0.178 | 0.253 | 0.287 |
| Golph3l | 0.178 | 0.126 | 0.139 |
| Plrg1 | 0.178 | 0.140 | 0.148 |
| Ankib1 | 0.178 | 0.145 | 0.161 |
| Nop14 | 0.178 | 0.243 | 0.214 |
| Ppp1r14a | 0.178 | 0.780 | 0.279 |
| Pogz | 0.178 | 0.154 | 0.130 |
| Gid8 | 0.178 | 0.163 | 0.153 |
| Actr8 | 0.178 | 0.147 | 0.147 |
| Arap1 | 0.178 | 0.148 | 0.476 |
| Thbs3 | 0.178 | 0.385 | 0.819 |
| Mrpl19 | 0.178 | 0.177 | 0.162 |
| Gale | 0.178 | 0.040 | 0.026 |
| Gmppa | 0.178 | 0.195 | 0.271 |
| Wdr5 | 0.178 | 0.100 | 0.108 |
| Ddx23 | 0.177 | 0.147 | 0.122 |
| Zfp688 | 0.177 | 0.090 | 0.099 |
| Tmem178 | 0.177 | 0.012 | 0.046 |
| Sufu | 0.177 | 0.104 | 0.109 |
| Pcna | 0.177 | 0.190 | 0.183 |
| Gria3 | 0.177 | 0.047 | 0.136 |
| Atg9a | 0.177 | 0.177 | 0.181 |
| Fnbp1 | 0.177 | 0.206 | 0.160 |
| Arl6ip6 | 0.177 | 0.184 | 0.188 |
| Zmym2 | 0.177 | 0.175 | 0.151 |
| Ckap5 | 0.177 | 0.211 | 0.181 |
| Pspc1 | 0.177 | 0.175 | 0.166 |
| Ugdh | 0.177 | 0.348 | 0.871 |
| Erich1 | 0.177 | 0.080 | 0.084 |

| | | | |
|---|---|---|---|
| Adh1 | 0.177 | 0.017 | 0.205 |
| Erg | 0.177 | 0.069 | 0.025 |
| 2310035C23F | 0.177 | 0.199 | 0.161 |
| Phtf1 | 0.176 | 0.140 | 0.181 |
| Ednrb | 0.176 | 0.038 | 0.024 |
| Gstt2 | 0.176 | 0.041 | 0.046 |
| Deaf1 | 0.176 | 0.177 | 0.251 |
| Cyyr1 | 0.176 | 0.006 | 0.002 |
| Pwwp2a | 0.176 | 0.212 | 0.191 |
| Rcc2 | 0.176 | 0.145 | 0.118 |
| Polr3gl | 0.176 | 0.210 | 0.219 |
| Cherp | 0.176 | 0.169 | 0.156 |
| Alad | 0.176 | 0.182 | 0.149 |
| Lbh | 0.176 | 0.451 | 0.545 |
| Sorbs3 | 0.176 | 0.100 | 0.121 |
| Rrp8 | 0.176 | 0.193 | 0.202 |
| Ap3s2 | 0.176 | 0.117 | 0.145 |
| Sesn1 | 0.176 | 0.320 | 0.252 |
| Plekho2 | 0.176 | 0.224 | 0.223 |
| Trp53rka | 0.176 | 0.091 | 0.105 |
| Hsdl1 | 0.176 | 0.074 | 0.077 |
| Med24 | 0.176 | 0.079 | 0.093 |
| Vps52 | 0.176 | 0.122 | 0.124 |
| Gm26771 | 0.176 | 0.241 | 0.167 |
| Lactb | 0.176 | 0.117 | 0.096 |
| Cdk5 | 0.175 | 0.168 | 0.170 |
| Cdr2l | 0.175 | 0.083 | 0.178 |
| Sec23b | 0.175 | 0.144 | 0.146 |
| Chid1 | 0.175 | 0.134 | 0.255 |
| Cyld | 0.175 | 0.153 | 0.114 |
| Gnl2 | 0.175 | 0.248 | 0.217 |
| Tfdp1 | 0.175 | 0.159 | 0.189 |
| Sumf1 | 0.175 | 0.180 | 0.154 |
| Wiz | 0.175 | 0.175 | 0.148 |
| Mta3 | 0.175 | 0.145 | 0.145 |
| Zdhhc5 | 0.175 | 0.167 | 0.164 |
| Exoc2 | 0.175 | 0.084 | 0.084 |
| Elp2 | 0.175 | 0.167 | 0.172 |
| Mogs | 0.175 | 0.151 | 0.260 |
| Adcy6 | 0.175 | 0.163 | 0.121 |
| Tspan12 | 0.175 | 0.042 | 0.055 |
| Gm9493 | 0.175 | 0.143 | 0.162 |
| 0610009O20I | 0.175 | 0.131 | 0.148 |
| Cnot11 | 0.175 | 0.175 | 0.157 |
| Aftph | 0.175 | 0.170 | 0.114 |
| Samd4 | 0.175 | 0.647 | 0.406 |
| Dus3l | 0.175 | 0.164 | 0.135 |
| Chpt1 | 0.175 | 0.436 | 0.287 |

| | | | |
|---|---|---|---|
| Sirt7 | 0.174 | 0.135 | 0.117 |
| Ngly1 | 0.174 | 0.236 | 0.221 |
| Ccdc71l | 0.174 | 0.205 | 0.161 |
| Ivd | 0.174 | 0.139 | 0.085 |
| Stxbp2 | 0.174 | 0.044 | 0.014 |
| Igfbp2 | 0.174 | 0.763 | 0.145 |
| Trappc12 | 0.174 | 0.121 | 0.144 |
| Tox2 | 0.174 | 0.016 | 0.014 |
| Vhl | 0.174 | 0.136 | 0.124 |
| Prmt5 | 0.174 | 0.158 | 0.159 |
| Nup98 | 0.174 | 0.262 | 0.255 |
| Fance | 0.174 | 0.289 | 0.248 |
| Sipa1l1 | 0.174 | 0.160 | 0.124 |
| AW549877 | 0.174 | 0.180 | 0.202 |
| Mt2 | 0.174 | 0.872 | 0.539 |
| Synj1 | 0.174 | 0.127 | 0.123 |
| Lhfpl2 | 0.174 | 0.109 | 0.613 |
| Kpna3 | 0.174 | 0.194 | 0.175 |
| Baiap2 | 0.174 | 0.217 | 0.326 |
| Kdm2b | 0.174 | 0.178 | 0.168 |
| Rasa2 | 0.174 | 0.294 | 0.235 |
| Sec61a2 | 0.173 | 0.209 | 0.239 |
| Gtpbp6 | 0.173 | 0.126 | 0.120 |
| Tcp11l1 | 0.173 | 0.063 | 0.050 |
| Ccdc58 | 0.173 | 0.149 | 0.157 |
| Atp9a | 0.173 | 0.279 | 0.399 |
| Arih2 | 0.173 | 0.194 | 0.168 |
| Hsf1 | 0.173 | 0.146 | 0.123 |
| Pygb | 0.173 | 0.342 | 0.329 |
| Rfx7 | 0.173 | 0.179 | 0.146 |
| Zfc3h1 | 0.173 | 0.174 | 0.161 |
| Dopey1 | 0.173 | 0.159 | 0.152 |
| Sccpdh | 0.173 | 0.178 | 0.173 |
| Dlg4 | 0.173 | 0.229 | 0.279 |
| P2rx4 | 0.173 | 0.230 | 0.203 |
| Abhd11 | 0.173 | 0.195 | 0.288 |
| Rab8b | 0.173 | 0.103 | 0.121 |
| Amd1 | 0.173 | 0.141 | 0.143 |
| 5730480H06I | 0.172 | 0.136 | 0.127 |
| Dtd1 | 0.172 | 0.171 | 0.169 |
| Guf1 | 0.172 | 0.195 | 0.251 |
| Cdk14 | 0.172 | 0.212 | 0.348 |
| Nes | 0.172 | 0.117 | 0.106 |
| Nvl | 0.172 | 0.179 | 0.186 |
| Alkbh7 | 0.172 | 0.172 | 0.155 |
| Tysnd1 | 0.172 | 0.128 | 0.133 |
| Pick1 | 0.172 | 0.132 | 0.114 |
| Fzd2 | 0.172 | 0.640 | 0.733 |

| | | | |
|---|---|---|---|
| Dscr3 | 0.172 | 0.154 | 0.152 |
| Lig1 | 0.172 | 0.147 | 0.189 |
| Gpd2 | 0.172 | 0.120 | 0.089 |
| Psen1 | 0.172 | 0.174 | 0.204 |
| Slc26a2 | 0.172 | 0.095 | 0.125 |
| Galt | 0.172 | 0.142 | 0.179 |
| Togaram1 | 0.171 | 0.142 | 0.136 |
| Dcaf6 | 0.171 | 0.188 | 0.142 |
| Cdc25b | 0.171 | 0.118 | 0.089 |
| Edc3 | 0.171 | 0.099 | 0.117 |
| Ccdc9 | 0.171 | 0.162 | 0.156 |
| Stn1 | 0.171 | 0.091 | 0.145 |
| Foxp4 | 0.171 | 0.294 | 0.201 |
| Jam3 | 0.171 | 0.592 | 0.399 |
| Tsc2 | 0.171 | 0.142 | 0.155 |
| Map3k2 | 0.171 | 0.244 | 0.232 |
| Mylk | 0.171 | 1.263 | 0.323 |
| R3hcc1 | 0.171 | 0.224 | 0.208 |
| Lman2l | 0.171 | 0.143 | 0.104 |
| Sos2 | 0.171 | 0.148 | 0.151 |
| Dusp16 | 0.171 | 0.182 | 0.204 |
| Gpalpp1 | 0.171 | 0.097 | 0.113 |
| Rnf2 | 0.171 | 0.229 | 0.277 |
| Gna14 | 0.171 | 0.212 | 0.062 |
| Tmem127 | 0.171 | 0.179 | 0.214 |
| Ipo9 | 0.171 | 0.153 | 0.194 |
| Metrnl | 0.171 | 0.224 | 0.295 |
| AU022252 | 0.171 | 0.114 | 0.123 |
| Blzf1 | 0.171 | 0.149 | 0.147 |
| Zcchc24 | 0.171 | 0.380 | 0.472 |
| Samd9l | 0.171 | 0.046 | 0.066 |
| Map2k7 | 0.171 | 0.191 | 0.186 |
| Srebf2 | 0.170 | 0.277 | 0.240 |
| Mtg2 | 0.170 | 0.118 | 0.108 |
| Eif2ak1 | 0.170 | 0.149 | 0.147 |
| Stx1a | 0.170 | 0.193 | 0.112 |
| Vti1a | 0.170 | 0.116 | 0.112 |
| Stard3 | 0.170 | 0.142 | 0.136 |
| Med22 | 0.170 | 0.105 | 0.099 |
| Ppp2r5d | 0.170 | 0.139 | 0.170 |
| Cox11 | 0.170 | 0.120 | 0.108 |
| Sptlc1 | 0.170 | 0.131 | 0.128 |
| Gan | 0.170 | 0.125 | 0.165 |
| Enc1 | 0.170 | 0.052 | 0.247 |
| Chkb | 0.170 | 0.191 | 0.208 |
| Lrrc47 | 0.170 | 0.120 | 0.122 |
| Mri1 | 0.170 | 0.167 | 0.194 |
| Mmp23 | 0.170 | 0.846 | 1.290 |

| | | | |
|---|---|---|---|
| Casp8 | 0.170 | 0.115 | 0.109 |
| Rnf168 | 0.169 | 0.133 | 0.121 |
| Nudt14 | 0.169 | 0.062 | 0.057 |
| Slc35c2 | 0.169 | 0.142 | 0.185 |
| Nfrkb | 0.169 | 0.165 | 0.135 |
| Ftsj3 | 0.169 | 0.152 | 0.151 |
| Jak2 | 0.169 | 0.155 | 0.108 |
| Bcl9 | 0.169 | 0.252 | 0.205 |
| Ift74 | 0.169 | 0.250 | 0.251 |
| Slc25a23 | 0.169 | 0.256 | 0.135 |
| Atf6 | 0.169 | 0.138 | 0.116 |
| Mtor | 0.169 | 0.154 | 0.097 |
| Zer1 | 0.169 | 0.192 | 0.178 |
| Dpy19l1 | 0.169 | 0.169 | 0.252 |
| Nelfcd | 0.169 | 0.130 | 0.133 |
| Ppcs | 0.169 | 0.083 | 0.083 |
| Atp9b | 0.169 | 0.129 | 0.136 |
| Alkbh6 | 0.169 | 0.118 | 0.089 |
| Uck2 | 0.168 | 0.093 | 0.049 |
| Nsmaf | 0.168 | 0.181 | 0.199 |
| Agps | 0.168 | 0.138 | 0.146 |
| Rnf44 | 0.168 | 0.123 | 0.091 |
| Hmgxb4 | 0.168 | 0.168 | 0.161 |
| B3glct | 0.168 | 0.259 | 0.206 |
| Pusl1 | 0.168 | 0.104 | 0.161 |
| Psme3 | 0.168 | 0.124 | 0.142 |
| Pcgf5 | 0.168 | 0.203 | 0.055 |
| Katna1 | 0.168 | 0.163 | 0.166 |
| Gpatch11 | 0.168 | 0.144 | 0.155 |
| Usp10 | 0.168 | 0.160 | 0.154 |
| Chuk | 0.168 | 0.174 | 0.129 |
| Lyrm4 | 0.168 | 0.081 | 0.064 |
| Nacc1 | 0.168 | 0.147 | 0.102 |
| Pex16 | 0.168 | 0.140 | 0.124 |
| Edem1 | 0.168 | 0.120 | 0.126 |
| Pex13 | 0.168 | 0.238 | 0.210 |
| Chmp1b | 0.168 | 0.150 | 0.131 |
| Naa16 | 0.168 | 0.140 | 0.116 |
| Ethe1 | 0.167 | 0.253 | 0.172 |
| Plekha5 | 0.167 | 0.230 | 0.360 |
| Pycrl | 0.167 | 0.139 | 0.132 |
| Usp39 | 0.167 | 0.127 | 0.118 |
| Rpap3 | 0.167 | 0.131 | 0.156 |
| Psmd5 | 0.167 | 0.147 | 0.133 |
| Xxylt1 | 0.167 | 0.107 | 0.100 |
| Usp12 | 0.167 | 0.144 | 0.107 |
| Nt5c2 | 0.167 | 0.177 | 0.169 |
| Coch | 0.167 | 0.005 | 0.007 |

| | | | |
|---|---|---|---|
| Apeh | 0.167 | 0.116 | 0.129 |
| Fkbp15 | 0.167 | 0.132 | 0.139 |
| Rad17 | 0.167 | 0.199 | 0.166 |
| Dxo | 0.167 | 0.138 | 0.164 |
| Iqsec1 | 0.167 | 0.297 | 0.248 |
| Sac3d1 | 0.167 | 0.154 | 0.146 |
| Ranbp3 | 0.167 | 0.168 | 0.166 |
| Ano10 | 0.167 | 0.194 | 0.178 |
| Serpinb1a | 0.167 | 0.043 | 0.076 |
| Ctbs | 0.167 | 0.121 | 0.170 |
| 3830406C13F | 0.167 | 0.172 | 0.178 |
| Rrs1 | 0.167 | 0.187 | 0.200 |
| Chm | 0.167 | 0.154 | 0.138 |
| Gabpb2 | 0.167 | 0.167 | 0.172 |
| Xrcc4 | 0.167 | 0.140 | 0.084 |
| Adprhl2 | 0.167 | 0.151 | 0.192 |
| Dhx38 | 0.167 | 0.138 | 0.136 |
| Myrip | 0.166 | 0.023 | 0.007 |
| Usp32 | 0.166 | 0.107 | 0.111 |
| Lpgat1 | 0.166 | 0.298 | 0.316 |
| Slc41a1 | 0.166 | 0.189 | 0.151 |
| Sin3a | 0.166 | 0.170 | 0.146 |
| Cog5 | 0.166 | 0.190 | 0.161 |
| Smim13 | 0.166 | 0.179 | 0.161 |
| Nfkbib | 0.166 | 0.147 | 0.124 |
| Dusp6 | 0.166 | 0.107 | 0.097 |
| Bcap29 | 0.166 | 0.122 | 0.100 |
| Gm37494 | 0.166 | 0.141 | 0.188 |
| Serping1 | 0.166 | 0.375 | 1.418 |
| Crkl | 0.166 | 0.142 | 0.143 |
| Ciz1 | 0.166 | 0.161 | 0.168 |
| Nde1 | 0.166 | 0.141 | 0.116 |
| Meis1 | 0.166 | 0.553 | 0.613 |
| Sbf1 | 0.166 | 0.134 | 0.147 |
| Fam241a | 0.166 | 0.278 | 0.055 |
| Tfb2m | 0.166 | 0.153 | 0.140 |
| Zfp281 | 0.166 | 0.331 | 0.180 |
| Mlycd | 0.166 | 0.150 | 0.148 |
| Dph6 | 0.166 | 0.124 | 0.106 |
| Vipas39 | 0.166 | 0.113 | 0.129 |
| Plekhm2 | 0.166 | 0.219 | 0.182 |
| Axin1 | 0.166 | 0.202 | 0.210 |
| Aktip | 0.166 | 0.149 | 0.148 |
| Prep | 0.166 | 0.108 | 0.090 |
| Herc4 | 0.166 | 0.195 | 0.120 |
| Prkcd | 0.166 | 0.291 | 0.235 |
| Mrpl37 | 0.165 | 0.134 | 0.120 |
| Trappc10 | 0.165 | 0.152 | 0.142 |

| | | | |
|---|---|---|---|
| Kdm3a | 0.165 | 0.143 | 0.147 |
| Enah | 0.165 | 0.743 | 0.635 |
| Wdr11 | 0.165 | 0.111 | 0.106 |
| Arsa | 0.165 | 0.059 | 0.138 |
| Slc12a6 | 0.165 | 0.153 | 0.137 |
| Gsto1 | 0.165 | 0.268 | 0.245 |
| Patz1 | 0.165 | 0.133 | 0.115 |
| St6galnac6 | 0.165 | 0.203 | 0.222 |
| Hsph1 | 0.165 | 0.196 | 0.159 |
| Ptgfrn | 0.165 | 0.165 | 0.392 |
| Pts | 0.165 | 0.140 | 0.170 |
| Bap1 | 0.165 | 0.129 | 0.134 |
| Mbip | 0.165 | 0.158 | 0.154 |
| Mier3 | 0.165 | 0.204 | 0.234 |
| Mamstr | 0.165 | 0.071 | 0.076 |
| Gnptab | 0.165 | 0.099 | 0.102 |
| Phf2 | 0.165 | 0.166 | 0.161 |
| Gulp1 | 0.165 | 0.207 | 0.235 |
| Vezt | 0.165 | 0.161 | 0.155 |
| Nampt | 0.165 | 0.186 | 0.177 |
| Grik5 | 0.164 | 0.176 | 0.143 |
| Exoc6b | 0.164 | 0.183 | 0.171 |
| Fbxl6 | 0.164 | 0.175 | 0.176 |
| Cp | 0.164 | 0.027 | 0.157 |
| Fam98a | 0.164 | 0.091 | 0.129 |
| Mars | 0.164 | 0.164 | 0.183 |
| Pf4 | 0.164 | 0.053 | 0.064 |
| Men1 | 0.164 | 0.113 | 0.137 |
| Trak1 | 0.164 | 0.232 | 0.226 |
| Slc35e1 | 0.164 | 0.091 | 0.126 |
| Creb3l1 | 0.164 | 0.197 | 0.564 |
| 1110004E09F | 0.164 | 0.159 | 0.155 |
| Mboat2 | 0.164 | 0.183 | 0.121 |
| Hivep1 | 0.164 | 0.168 | 0.133 |
| Ruvbl2 | 0.163 | 0.116 | 0.138 |
| Tubg1 | 0.163 | 0.120 | 0.117 |
| Shroom4 | 0.163 | 0.038 | 0.018 |
| Dll4 | 0.163 | 0.006 | 0.002 |
| Grk5 | 0.163 | 0.025 | 0.044 |
| Clpb | 0.163 | 0.173 | 0.144 |
| Cenpf | 0.163 | 0.048 | 0.047 |
| Gtf2a1 | 0.163 | 0.130 | 0.101 |
| Rrm1 | 0.163 | 0.121 | 0.179 |
| Phkg2 | 0.163 | 0.170 | 0.177 |
| Dhx8 | 0.163 | 0.128 | 0.123 |
| Larp1b | 0.163 | 0.204 | 0.070 |
| Cd3eap | 0.163 | 0.152 | 0.143 |
| Ahcyl2 | 0.163 | 0.230 | 0.138 |

| | | | |
|---|---|---|---|
| Cdyl | 0.163 | 0.152 | 0.237 |
| Vegfc | 0.163 | 0.030 | 0.095 |
| Lrig3 | 0.163 | 0.401 | 0.263 |
| Slc12a7 | 0.163 | 0.059 | 0.016 |
| Ift22 | 0.163 | 0.128 | 0.151 |
| Ago1 | 0.163 | 0.132 | 0.102 |
| Snhg9 | 0.163 | 0.119 | 0.129 |
| Rbsn | 0.163 | 0.124 | 0.151 |
| Map3k20 | 0.163 | 0.483 | 0.215 |
| Mrpl39 | 0.163 | 0.154 | 0.135 |
| Parn | 0.163 | 0.160 | 0.159 |
| Map2k1 | 0.163 | 0.166 | 0.140 |
| Nnmt | 0.163 | 0.136 | 0.351 |
| Ddhd2 | 0.162 | 0.205 | 0.213 |
| Zufsp | 0.162 | 0.139 | 0.164 |
| Ids | 0.162 | 0.133 | 0.124 |
| Ap1s2 | 0.162 | 0.107 | 0.107 |
| Trmt1l | 0.162 | 0.123 | 0.125 |
| Mcm7 | 0.162 | 0.143 | 0.188 |
| Nt5m | 0.162 | 0.106 | 0.137 |
| Dnajc11 | 0.162 | 0.140 | 0.139 |
| Shmt2 | 0.162 | 0.141 | 0.156 |
| Heph | 0.162 | 0.087 | 0.171 |
| Gpn1 | 0.162 | 0.128 | 0.135 |
| Lgalsl | 0.162 | 0.340 | 0.248 |
| Fam98c | 0.162 | 0.142 | 0.102 |
| Ddx10 | 0.162 | 0.172 | 0.150 |
| Gm15050 | 0.162 | 0.151 | 0.137 |
| Slc25a32 | 0.162 | 0.118 | 0.108 |
| B3gnt3 | 0.162 | 0.001 | 0.002 |
| Slc4a4 | 0.162 | 0.134 | 0.237 |
| Ccnk | 0.162 | 0.118 | 0.126 |
| Dnajb12 | 0.161 | 0.128 | 0.132 |
| Pi4k2a | 0.161 | 0.208 | 0.197 |
| Dr1 | 0.161 | 0.133 | 0.133 |
| Sh3kbp1 | 0.161 | 0.102 | 0.298 |
| Ranbp10 | 0.161 | 0.118 | 0.115 |
| Mid1ip1 | 0.161 | 0.226 | 0.158 |
| Zcchc8 | 0.161 | 0.160 | 0.128 |
| Zfp655 | 0.161 | 0.172 | 0.198 |
| Usp40 | 0.161 | 0.148 | 0.181 |
| Exoc1 | 0.161 | 0.131 | 0.122 |
| Htt | 0.161 | 0.151 | 0.159 |
| Dus1l | 0.161 | 0.124 | 0.112 |
| Rrm2b | 0.161 | 0.107 | 0.095 |
| Daxx | 0.161 | 0.148 | 0.150 |
| Thyn1 | 0.161 | 0.193 | 0.140 |
| Tnip1 | 0.161 | 0.110 | 0.128 |

| | | | |
|---|---|---|---|
| Fam219b | 0.161 | 0.096 | 0.110 |
| Lyrm9 | 0.161 | 0.021 | 0.019 |
| Aar2 | 0.161 | 0.093 | 0.082 |
| Skiv2l2 | 0.160 | 0.187 | 0.170 |
| Kif1bp | 0.160 | 0.173 | 0.169 |
| Ptpn9 | 0.160 | 0.226 | 0.225 |
| Mvb12b | 0.160 | 0.126 | 0.084 |
| Hey1 | 0.160 | 0.127 | 0.098 |
| Rxra | 0.160 | 0.200 | 0.193 |
| Ubxn7 | 0.160 | 0.161 | 0.144 |
| Nit1 | 0.160 | 0.191 | 0.156 |
| Rabgap1l | 0.160 | 0.183 | 0.169 |
| Fads3 | 0.160 | 0.243 | 0.193 |
| Rcbtb1 | 0.160 | 0.110 | 0.110 |
| Tmem106c | 0.160 | 0.092 | 0.212 |
| Utp14a | 0.160 | 0.193 | 0.173 |
| Slc9a1 | 0.160 | 0.121 | 0.121 |
| Mettl6 | 0.160 | 0.153 | 0.158 |
| Lig3 | 0.160 | 0.103 | 0.108 |
| Rasgrf2 | 0.160 | 0.017 | 0.003 |
| Hltf | 0.159 | 0.116 | 0.116 |
| Atg16l1 | 0.159 | 0.123 | 0.109 |
| Fem1c | 0.159 | 0.136 | 0.132 |
| Pithd1 | 0.159 | 0.138 | 0.123 |
| Socs4 | 0.159 | 0.151 | 0.147 |
| Slc20a1 | 0.159 | 0.091 | 0.113 |
| Slc2a1 | 0.159 | 0.132 | 0.071 |
| Ttc23 | 0.159 | 0.126 | 0.105 |
| Sugp1 | 0.159 | 0.125 | 0.116 |
| Rnf185 | 0.159 | 0.102 | 0.097 |
| Zfp512b | 0.159 | 0.169 | 0.147 |
| Zbtb16 | 0.159 | 0.371 | 0.239 |
| Nkain4 | 0.159 | 0.122 | 0.240 |
| Rbm34 | 0.159 | 0.156 | 0.123 |
| Mtif2 | 0.159 | 0.130 | 0.135 |
| Mlx | 0.159 | 0.146 | 0.121 |
| Atp13a1 | 0.159 | 0.134 | 0.135 |
| Mfsd11 | 0.159 | 0.165 | 0.161 |
| Gnb5 | 0.159 | 0.158 | 0.216 |
| Tbc1d8 | 0.159 | 0.058 | 0.018 |
| Eef1akmt1 | 0.159 | 0.236 | 0.230 |
| Ncoa6 | 0.159 | 0.170 | 0.153 |
| Gpr182 | 0.159 | 0.001 | 0.001 |
| Prkrip1 | 0.159 | 0.127 | 0.109 |
| Stard9 | 0.158 | 0.104 | 0.219 |
| Gpatch2 | 0.158 | 0.152 | 0.138 |
| Tuba1c | 0.158 | 0.424 | 0.234 |
| Klhl22 | 0.158 | 0.175 | 0.250 |

| | | | |
|---|---|---|---|
| Gfod2 | 0.158 | 0.069 | 0.064 |
| Blvra | 0.158 | 0.160 | 0.142 |
| Fbxl14 | 0.158 | 0.125 | 0.100 |
| Upp2 | 0.158 | 0.079 | 0.084 |
| Rap2a | 0.158 | 0.145 | 0.414 |
| Cc2d2a | 0.158 | 0.166 | 0.209 |
| Smim4 | 0.158 | 0.188 | 0.183 |
| Tmem119 | 0.158 | 0.217 | 0.611 |
| Wipf2 | 0.158 | 0.124 | 0.102 |
| Lcorl | 0.158 | 0.117 | 0.171 |
| Ttc13 | 0.158 | 0.120 | 0.142 |
| Snta1 | 0.158 | 0.389 | 0.370 |
| Rufy2 | 0.158 | 0.136 | 0.186 |
| Pno1 | 0.158 | 0.140 | 0.126 |
| Eed | 0.158 | 0.159 | 0.147 |
| Zfp62 | 0.158 | 0.142 | 0.174 |
| Zfp532 | 0.158 | 0.168 | 0.153 |
| Rdm1 | 0.158 | 0.147 | 0.167 |
| Arrdc1 | 0.158 | 0.080 | 0.057 |
| Usp46 | 0.157 | 0.196 | 0.216 |
| Glis2 | 0.157 | 0.173 | 0.300 |
| Mex3a | 0.157 | 0.199 | 0.175 |
| Sdad1 | 0.157 | 0.156 | 0.147 |
| Mex3c | 0.157 | 0.231 | 0.207 |
| Zfpm2 | 0.157 | 0.176 | 0.276 |
| Palm | 0.157 | 0.221 | 0.399 |
| Btf3l4 | 0.157 | 0.177 | 0.153 |
| Abt1 | 0.157 | 0.133 | 0.102 |
| Fam117b | 0.157 | 0.136 | 0.138 |
| Ldlrap1 | 0.157 | 0.076 | 0.109 |
| Zcchc3 | 0.157 | 0.214 | 0.210 |
| Cmc2 | 0.157 | 0.129 | 0.152 |
| Zfp865 | 0.157 | 0.124 | 0.132 |
| Rpp14 | 0.157 | 0.147 | 0.152 |
| Drosha | 0.157 | 0.119 | 0.149 |
| Golga1 | 0.157 | 0.133 | 0.137 |
| Mettl27 | 0.157 | 0.056 | 0.063 |
| Ptprr | 0.157 | 0.002 | 0.003 |
| Snx15 | 0.157 | 0.107 | 0.108 |
| Sec14l2 | 0.157 | 0.015 | 0.009 |
| Sgcb | 0.157 | 0.375 | 0.269 |
| 2010320M18 | 0.157 | 0.246 | 0.173 |
| Myd88 | 0.157 | 0.206 | 0.269 |
| Selenoi | 0.157 | 0.222 | 0.179 |
| Kat2b | 0.157 | 0.203 | 0.139 |
| Tmem198b | 0.157 | 0.108 | 0.170 |
| Dusp7 | 0.157 | 0.172 | 0.166 |
| Uxt | 0.157 | 0.117 | 0.120 |

| | | | |
|---|---|---|---|
| Ubald1 | 0.157 | 0.194 | 0.164 |
| Ankrd13c | 0.157 | 0.117 | 0.109 |
| Josd1 | 0.157 | 0.157 | 0.159 |
| Fam126b | 0.157 | 0.152 | 0.133 |
| Thoc1 | 0.157 | 0.136 | 0.132 |
| Vps16 | 0.156 | 0.112 | 0.123 |
| Nit2 | 0.156 | 0.253 | 0.230 |
| Tbcd | 0.156 | 0.089 | 0.087 |
| Cdon | 0.156 | 0.234 | 0.381 |
| Slc38a9 | 0.156 | 0.125 | 0.129 |
| Inppl1 | 0.156 | 0.258 | 0.173 |
| Asb8 | 0.156 | 0.156 | 0.120 |
| Foxj3 | 0.156 | 0.153 | 0.152 |
| Pikfyve | 0.156 | 0.133 | 0.132 |
| Coa6 | 0.156 | 0.135 | 0.124 |
| Shroom3 | 0.156 | 0.264 | 0.060 |
| Il2rg | 0.156 | 0.008 | 0.005 |
| Stoml1 | 0.156 | 0.143 | 0.163 |
| Fbxl20 | 0.156 | 0.181 | 0.144 |
| Med12 | 0.156 | 0.133 | 0.146 |
| Asl | 0.156 | 0.157 | 0.102 |
| Smc2 | 0.156 | 0.105 | 0.144 |
| Ssh2 | 0.156 | 0.121 | 0.182 |
| Ubtd1 | 0.156 | 0.115 | 0.103 |
| Tbc1d12 | 0.156 | 0.129 | 0.181 |
| Lrrk1 | 0.156 | 0.149 | 0.269 |
| Stap2 | 0.156 | 0.003 | 0.004 |
| Nipsnap1 | 0.156 | 0.095 | 0.070 |
| Osbpl3 | 0.156 | 0.183 | 0.084 |
| Hip1r | 0.156 | 0.108 | 0.031 |
| Zfp423 | 0.155 | 0.057 | 0.153 |
| Plaur | 0.155 | 0.090 | 0.092 |
| Abr | 0.155 | 0.195 | 0.092 |
| Casp2 | 0.155 | 0.072 | 0.076 |
| Zfp212 | 0.155 | 0.151 | 0.136 |
| Nudt18 | 0.155 | 0.127 | 0.133 |
| Dyrk2 | 0.155 | 0.253 | 0.284 |
| Ophn1 | 0.155 | 0.100 | 0.158 |
| Tiparp | 0.155 | 0.367 | 0.367 |
| Slit2 | 0.155 | 0.262 | 0.429 |
| 8-Mar | 0.155 | 0.105 | 0.146 |
| Tada1 | 0.155 | 0.103 | 0.113 |
| Camkk2 | 0.155 | 0.221 | 0.280 |
| Armc8 | 0.155 | 0.137 | 0.123 |
| Mapk12 | 0.155 | 0.007 | 0.028 |
| Fam222b | 0.155 | 0.118 | 0.105 |
| Ring1 | 0.155 | 0.225 | 0.218 |
| Bysl | 0.155 | 0.088 | 0.094 |

| | | | |
|---|---|---|---|
| Fundc1 | 0.155 | 0.152 | 0.176 |
| Strip1 | 0.155 | 0.099 | 0.101 |
| Tmem140 | 0.155 | 0.067 | 0.140 |
| Coa5 | 0.155 | 0.140 | 0.171 |
| Fam174b | 0.155 | 0.079 | 0.058 |
| Zc3hc1 | 0.155 | 0.139 | 0.135 |
| C1galt1 | 0.155 | 0.212 | 0.140 |
| Ankrd39 | 0.155 | 0.151 | 0.162 |
| Timm29 | 0.155 | 0.135 | 0.137 |
| Anks1 | 0.154 | 0.111 | 0.067 |
| Pcyt2 | 0.154 | 0.144 | 0.143 |
| Pitpnc1 | 0.154 | 0.090 | 0.129 |
| Ttll7 | 0.154 | 0.223 | 0.051 |
| Stx3 | 0.154 | 0.045 | 0.028 |
| Tsen15 | 0.154 | 0.184 | 0.196 |
| D10Wsu102e | 0.154 | 0.356 | 0.178 |
| Mpp6 | 0.154 | 0.807 | 0.852 |
| Mthfsd | 0.154 | 0.110 | 0.141 |
| Otud4 | 0.154 | 0.142 | 0.145 |
| Timm21 | 0.154 | 0.168 | 0.182 |
| Gm26802 | 0.154 | 0.543 | 0.475 |
| Parg | 0.154 | 0.160 | 0.126 |
| Zfp746 | 0.154 | 0.142 | 0.143 |
| Wdr77 | 0.154 | 0.131 | 0.164 |
| Ndufaf7 | 0.153 | 0.145 | 0.148 |
| Polr3c | 0.153 | 0.117 | 0.110 |
| Myo9b | 0.153 | 0.221 | 0.190 |
| Cdh11 | 0.153 | 0.561 | 0.851 |
| Slc25a25 | 0.153 | 0.231 | 0.132 |
| Pmm1 | 0.153 | 0.337 | 0.406 |
| Map3k4 | 0.153 | 0.149 | 0.123 |
| Cpeb4 | 0.153 | 0.160 | 0.123 |
| Gnptg | 0.153 | 0.113 | 0.127 |
| Rab10os | 0.153 | 0.105 | 0.120 |
| Acp6 | 0.153 | 0.063 | 0.039 |
| Mfn2 | 0.153 | 0.124 | 0.126 |
| Galnt10 | 0.153 | 0.091 | 0.103 |
| Dusp22 | 0.153 | 0.083 | 0.093 |
| Mlxip | 0.153 | 0.170 | 0.131 |
| Armc5 | 0.153 | 0.127 | 0.113 |
| Mepce | 0.153 | 0.179 | 0.152 |
| Tubgcp3 | 0.153 | 0.107 | 0.095 |
| Cyb5r1 | 0.153 | 0.129 | 0.106 |
| Kdr | 0.153 | 0.012 | 0.014 |
| Cinp | 0.153 | 0.124 | 0.124 |
| Mettl16 | 0.153 | 0.131 | 0.158 |
| Trappc2 | 0.153 | 0.074 | 0.063 |
| Csf1 | 0.153 | 0.152 | 0.337 |

| | | | |
|---|---|---|---|
| Ap1ar | 0.153 | 0.171 | 0.128 |
| Efemp1 | 0.153 | 0.019 | 0.093 |
| Kalrn | 0.153 | 0.280 | 0.125 |
| Stat5b | 0.152 | 0.113 | 0.120 |
| Klhdc8b | 0.152 | 0.442 | 0.215 |
| Nploc4 | 0.152 | 0.119 | 0.106 |
| Cwc27 | 0.152 | 0.154 | 0.133 |
| Ptrh2 | 0.152 | 0.116 | 0.138 |
| Zbtb44 | 0.152 | 0.185 | 0.170 |
| Agap1 | 0.152 | 0.205 | 0.259 |
| Tprkb | 0.152 | 0.205 | 0.201 |
| Msh3 | 0.152 | 0.120 | 0.114 |
| 10-Sep | 0.152 | 0.090 | 0.084 |
| Afg3l2 | 0.152 | 0.172 | 0.144 |
| Snapc3 | 0.152 | 0.133 | 0.152 |
| Phc1 | 0.152 | 0.155 | 0.124 |
| Alg10b | 0.152 | 0.110 | 0.111 |
| Cln6 | 0.152 | 0.088 | 0.118 |
| Mob3c | 0.152 | 0.147 | 0.199 |
| Ccdc8 | 0.152 | 0.186 | 0.279 |
| Msantd2 | 0.152 | 0.184 | 0.152 |
| Dynlt1f | 0.152 | 0.134 | 0.235 |
| Kdelc1 | 0.152 | 0.143 | 0.164 |
| Chst14 | 0.152 | 0.145 | 0.218 |
| Crcp | 0.152 | 0.106 | 0.104 |
| Tceal1 | 0.151 | 0.174 | 0.146 |
| Nob1 | 0.151 | 0.164 | 0.125 |
| Zfp821 | 0.151 | 0.190 | 0.145 |
| Ezh2 | 0.151 | 0.180 | 0.229 |
| Ccdc86 | 0.151 | 0.137 | 0.136 |
| Amn1 | 0.151 | 0.165 | 0.157 |
| Vps9d1 | 0.151 | 0.133 | 0.135 |
| Mrpl10 | 0.151 | 0.122 | 0.129 |
| Stard13 | 0.151 | 0.091 | 0.064 |
| Ccdc71 | 0.151 | 0.104 | 0.121 |
| Efnb1 | 0.151 | 0.254 | 0.417 |
| Odf2 | 0.151 | 0.199 | 0.203 |
| Msra | 0.151 | 0.130 | 0.072 |
| Armc10 | 0.151 | 0.120 | 0.092 |
| Trim2 | 0.151 | 0.056 | 0.117 |
| Abcb10 | 0.151 | 0.050 | 0.049 |
| Ppp1r35 | 0.151 | 0.125 | 0.115 |
| Rreb1 | 0.151 | 0.116 | 0.104 |
| Atad3a | 0.151 | 0.157 | 0.162 |
| Uck1 | 0.151 | 0.104 | 0.134 |
| Lmod1 | 0.151 | 1.346 | 0.324 |
| Slc17a5 | 0.151 | 0.124 | 0.133 |
| Col4a4 | 0.151 | 0.025 | 0.056 |

| | | | |
|---|---|---|---|
| Psmg1 | 0.151 | 0.179 | 0.180 |
| Ercc5 | 0.151 | 0.169 | 0.168 |
| Nicn1 | 0.151 | 0.176 | 0.164 |
| Tex10 | 0.151 | 0.161 | 0.175 |
| 2610002M06 | 0.151 | 0.151 | 0.172 |
| Dnajc24 | 0.151 | 0.123 | 0.113 |
| Dgcr14 | 0.151 | 0.130 | 0.109 |
| Stx2 | 0.151 | 0.098 | 0.150 |
| Sh3rf1 | 0.150 | 0.135 | 0.112 |
| Ankrd49 | 0.150 | 0.091 | 0.111 |
| Dolk | 0.150 | 0.136 | 0.160 |
| Nynrin | 0.150 | 0.158 | 0.190 |
| Ift52 | 0.150 | 0.113 | 0.131 |
| Rgs12 | 0.150 | 0.103 | 0.128 |
| Pggt1b | 0.150 | 0.089 | 0.080 |
| Coq6 | 0.150 | 0.105 | 0.104 |
| Cand2 | 0.150 | 0.124 | 0.130 |
| Suclg2 | 0.150 | 0.175 | 0.197 |
| Nck2 | 0.150 | 0.088 | 0.055 |
| Slc36a4 | 0.150 | 0.137 | 0.145 |
| Rnf34 | 0.150 | 0.193 | 0.155 |
| Morc2a | 0.150 | 0.175 | 0.143 |
| 3110040N11I | 0.150 | 0.082 | 0.090 |
| Fgd1 | 0.150 | 0.176 | 0.193 |
| Slc25a30 | 0.150 | 0.087 | 0.046 |
| Necap1 | 0.150 | 0.087 | 0.104 |
| Zdhhc18 | 0.150 | 0.096 | 0.081 |
| Gm13889 | 0.150 | 1.034 | 0.187 |
| Zfp593 | 0.150 | 0.176 | 0.124 |
| Sirt3 | 0.149 | 0.174 | 0.166 |
| Nup62 | 0.149 | 0.138 | 0.171 |
| Samd8 | 0.149 | 0.156 | 0.106 |
| Dlgap1 | 0.149 | 0.039 | 0.020 |
| Trf | 0.149 | 0.026 | 0.036 |
| Foxk2 | 0.149 | 0.125 | 0.112 |
| Fbxo28 | 0.149 | 0.142 | 0.126 |
| Ccdc88c | 0.149 | 0.008 | 0.012 |
| Utp18 | 0.149 | 0.161 | 0.122 |
| Ankrd33b | 0.149 | 0.010 | 0.011 |
| Ift122 | 0.149 | 0.048 | 0.066 |
| Ogfod2 | 0.149 | 0.075 | 0.091 |
| Fbxo31 | 0.149 | 0.281 | 0.201 |
| Taf2 | 0.149 | 0.174 | 0.130 |
| Tmem164 | 0.149 | 0.133 | 0.125 |
| Zfp512 | 0.149 | 0.121 | 0.141 |
| Rpa1 | 0.149 | 0.118 | 0.113 |
| D430042O09I | 0.149 | 0.129 | 0.175 |
| Gch1 | 0.149 | 0.068 | 0.033 |

| | | | |
|---|---|---|---|
| Zfp1 | 0.149 | 0.163 | 0.150 |
| Msh2 | 0.149 | 0.114 | 0.123 |
| Mrc2 | 0.149 | 0.289 | 0.496 |
| Pold3 | 0.149 | 0.108 | 0.099 |
| 5031439G07I | 0.149 | 0.101 | 0.181 |
| Capn7 | 0.149 | 0.128 | 0.135 |
| Plcb3 | 0.149 | 0.155 | 0.106 |
| Mbd6 | 0.149 | 0.175 | 0.139 |
| Tatdn1 | 0.149 | 0.148 | 0.157 |
| Tmco3 | 0.149 | 0.203 | 0.204 |
| Cul2 | 0.148 | 0.165 | 0.138 |
| Kdm4a | 0.148 | 0.135 | 0.142 |
| Hs6st1 | 0.148 | 0.072 | 0.065 |
| Aifm1 | 0.148 | 0.161 | 0.121 |
| Impact | 0.148 | 0.183 | 0.180 |
| Prmt7 | 0.148 | 0.176 | 0.191 |
| Dnajc9 | 0.148 | 0.131 | 0.134 |
| Zscan21 | 0.148 | 0.123 | 0.129 |
| Nab2 | 0.148 | 0.270 | 0.225 |
| Pip5k1c | 0.148 | 0.275 | 0.219 |
| Abhd13 | 0.148 | 0.108 | 0.083 |
| Znhit6 | 0.148 | 0.149 | 0.161 |
| Cep83 | 0.148 | 0.165 | 0.191 |
| Nhsl1 | 0.148 | 0.060 | 0.118 |
| Ptgr2 | 0.148 | 0.083 | 0.085 |
| Trp53bp1 | 0.148 | 0.165 | 0.133 |
| Adam9 | 0.148 | 0.271 | 0.180 |
| Gart | 0.148 | 0.165 | 0.153 |
| Arhgap21 | 0.148 | 0.138 | 0.301 |
| Csgalnact2 | 0.148 | 0.212 | 0.221 |
| Pradc1 | 0.147 | 0.072 | 0.101 |
| Ints14 | 0.147 | 0.130 | 0.112 |
| Gpr89 | 0.147 | 0.107 | 0.084 |
| Crnkl1 | 0.147 | 0.139 | 0.121 |
| Micall1 | 0.147 | 0.247 | 0.125 |
| Nfatc4 | 0.147 | 0.181 | 0.258 |
| Slc39a3 | 0.147 | 0.119 | 0.120 |
| Ppil3 | 0.147 | 0.149 | 0.161 |
| Pnpla6 | 0.147 | 0.075 | 0.062 |
| Acyp2 | 0.147 | 0.136 | 0.137 |
| Aen | 0.147 | 0.102 | 0.098 |
| Ankrd16 | 0.147 | 0.142 | 0.130 |
| Wdr12 | 0.147 | 0.138 | 0.135 |
| Sigirr | 0.147 | 0.002 | 0.002 |
| Rph3al | 0.147 | 0.005 | 0.003 |
| Llgl1 | 0.147 | 0.076 | 0.119 |
| Nfil3 | 0.147 | 0.719 | 0.397 |
| Slc16a13 | 0.147 | 0.004 | 0.002 |

| | | | |
|---|---|---|---|
| Habp4 | 0.147 | 0.170 | 0.170 |
| Hibch | 0.147 | 0.124 | 0.102 |
| Lmf1 | 0.147 | 0.095 | 0.120 |
| Synpo2 | 0.147 | 0.627 | 0.128 |
| Atpaf1 | 0.147 | 0.100 | 0.090 |
| Whamm | 0.147 | 0.144 | 0.104 |
| Yars | 0.147 | 0.148 | 0.156 |
| Sardh | 0.146 | 0.148 | 0.149 |
| Rpa3 | 0.146 | 0.140 | 0.112 |
| Pkia | 0.146 | 0.200 | 0.172 |
| Zscan26 | 0.146 | 0.151 | 0.137 |
| Tbc1d22a | 0.146 | 0.064 | 0.068 |
| Prdm11 | 0.146 | 0.174 | 0.113 |
| Aph1c | 0.146 | 0.023 | 0.017 |
| Fiz1 | 0.146 | 0.153 | 0.142 |
| Apbb1 | 0.146 | 0.166 | 0.137 |
| Tmem18 | 0.146 | 0.126 | 0.086 |
| Mtss1l | 0.146 | 0.262 | 0.112 |
| Akap5 | 0.146 | 0.086 | 0.013 |
| Pigc | 0.146 | 0.134 | 0.144 |
| Pogk | 0.146 | 0.149 | 0.149 |
| Shprh | 0.146 | 0.156 | 0.139 |
| Limk2 | 0.146 | 0.106 | 0.074 |
| Zfp398 | 0.146 | 0.147 | 0.129 |
| Ica1 | 0.146 | 0.041 | 0.014 |
| Zfp386 | 0.146 | 0.168 | 0.196 |
| Fes | 0.146 | 0.021 | 0.055 |
| Insr | 0.146 | 0.259 | 0.160 |
| Endog | 0.146 | 0.150 | 0.175 |
| Slc39a9 | 0.146 | 0.108 | 0.102 |
| Brip1os | 0.145 | 0.128 | 0.171 |
| Shkbp1 | 0.145 | 0.087 | 0.060 |
| Angptl4 | 0.145 | 0.180 | 0.349 |
| F8a | 0.145 | 0.082 | 0.055 |
| Exosc3 | 0.145 | 0.139 | 0.142 |
| Nop16 | 0.145 | 0.152 | 0.172 |
| Htra2 | 0.145 | 0.170 | 0.299 |
| Copg2 | 0.145 | 0.159 | 0.122 |
| Bcorl1 | 0.145 | 0.097 | 0.118 |
| Fmo5 | 0.145 | 0.047 | 0.084 |
| Bok | 0.145 | 0.138 | 0.213 |
| Pde8a | 0.145 | 0.010 | 0.029 |
| Prkx | 0.145 | 0.130 | 0.111 |
| Gm7694 | 0.145 | 0.027 | 0.011 |
| Cdc42ep4 | 0.145 | 0.178 | 0.241 |
| Zfyve27 | 0.145 | 0.118 | 0.098 |
| Tfip11 | 0.145 | 0.093 | 0.091 |
| Sgk3 | 0.145 | 0.055 | 0.036 |

| | | | |
|---|---|---|---|
| Tsc1 | 0.145 | 0.160 | 0.153 |
| Khdrbs3 | 0.145 | 0.329 | 0.144 |
| Gpn3 | 0.145 | 0.126 | 0.106 |
| Zfp157 | 0.145 | 0.142 | 0.137 |
| Prickle1 | 0.145 | 0.183 | 0.358 |
| Gypc | 0.145 | 0.166 | 0.149 |
| Srd5a3 | 0.145 | 0.138 | 0.179 |
| Rnf170 | 0.145 | 0.150 | 0.167 |
| Prickle2 | 0.145 | 0.083 | 0.116 |
| Dedd | 0.145 | 0.122 | 0.112 |
| C87436 | 0.145 | 0.155 | 0.148 |
| Leng1 | 0.145 | 0.176 | 0.158 |
| Akt3 | 0.145 | 0.129 | 0.121 |
| Cog2 | 0.144 | 0.123 | 0.143 |
| Tmem199 | 0.144 | 0.089 | 0.064 |
| Snx8 | 0.144 | 0.064 | 0.097 |
| Cdk19 | 0.144 | 0.102 | 0.088 |
| Sh3bp5l | 0.144 | 0.087 | 0.094 |
| Trim11 | 0.144 | 0.105 | 0.107 |
| Phf1 | 0.144 | 0.117 | 0.109 |
| Sp3os | 0.144 | 0.136 | 0.118 |
| Aldh6a1 | 0.144 | 0.210 | 0.138 |
| Cask | 0.144 | 0.200 | 0.228 |
| Bmt2 | 0.144 | 0.185 | 0.162 |
| Rnf40 | 0.144 | 0.106 | 0.097 |
| Acad8 | 0.144 | 0.122 | 0.113 |
| Ncoa1 | 0.144 | 0.231 | 0.230 |
| Sc5d | 0.144 | 0.120 | 0.142 |
| Lsp1 | 0.144 | 0.616 | 0.743 |
| Samhd1 | 0.144 | 0.103 | 0.158 |
| Prpf18 | 0.144 | 0.147 | 0.137 |
| Arhgap35 | 0.144 | 0.187 | 0.166 |
| Pycr1 | 0.144 | 0.084 | 0.287 |
| Gsta4 | 0.144 | 0.011 | 0.013 |
| Atp8b2 | 0.144 | 0.117 | 0.098 |
| Plcd3 | 0.144 | 0.058 | 0.040 |
| Axl | 0.144 | 0.601 | 0.982 |
| Zc3h4 | 0.144 | 0.128 | 0.118 |
| Alg1 | 0.144 | 0.138 | 0.144 |
| Etfdh | 0.144 | 0.148 | 0.133 |
| BC028528 | 0.144 | 0.265 | 0.219 |
| Ppp1r9a | 0.144 | 0.127 | 0.230 |
| Rexo4 | 0.144 | 0.130 | 0.142 |
| Trappc11 | 0.143 | 0.098 | 0.078 |
| Vps26b | 0.143 | 0.148 | 0.177 |
| Mtmr11 | 0.143 | 0.136 | 0.131 |
| Rab40c | 0.143 | 0.097 | 0.111 |
| Rap2c | 0.143 | 0.144 | 0.140 |

| | | | |
|---|---|---|---|
| Slc24a5 | 0.143 | 0.176 | 0.155 |
| Sema6d | 0.143 | 0.031 | 0.098 |
| Fut8 | 0.143 | 0.216 | 0.250 |
| Rangap1 | 0.143 | 0.096 | 0.103 |
| Mamld1 | 0.143 | 0.105 | 0.078 |
| Tcirg1 | 0.143 | 0.096 | 0.097 |
| Med17 | 0.143 | 0.113 | 0.106 |
| Fam45a | 0.143 | 0.119 | 0.156 |
| Scarb1 | 0.143 | 0.108 | 0.104 |
| G0s2 | 0.143 | 0.100 | 0.197 |
| Ctu2 | 0.143 | 0.116 | 0.100 |
| Atg13 | 0.143 | 0.117 | 0.096 |
| Lats1 | 0.143 | 0.219 | 0.203 |
| Gpatch1 | 0.142 | 0.116 | 0.111 |
| Blcap | 0.142 | 0.092 | 0.147 |
| Tusc1 | 0.142 | 0.171 | 0.143 |
| Lin37 | 0.142 | 0.123 | 0.111 |
| Smyd3 | 0.142 | 0.059 | 0.067 |
| Fam118a | 0.142 | 0.237 | 0.135 |
| Tnnt2 | 0.142 | 0.568 | 0.205 |
| Adgrg1 | 0.142 | 0.005 | 0.003 |
| Pwp1 | 0.142 | 0.137 | 0.133 |
| Zfp646 | 0.142 | 0.128 | 0.108 |
| Dnmt1 | 0.142 | 0.132 | 0.130 |
| Arl4c | 0.142 | 0.042 | 0.058 |
| Ube3b | 0.142 | 0.111 | 0.131 |
| Rasal2 | 0.142 | 0.106 | 0.083 |
| Nod1 | 0.142 | 0.039 | 0.041 |
| Ints7 | 0.142 | 0.145 | 0.117 |
| Ttf1 | 0.142 | 0.104 | 0.105 |
| Zcchc10 | 0.142 | 0.147 | 0.123 |
| Nudt19 | 0.142 | 0.214 | 0.202 |
| Prcc | 0.142 | 0.127 | 0.120 |
| Xpo4 | 0.142 | 0.154 | 0.147 |
| Tbc1d24 | 0.142 | 0.095 | 0.115 |
| Ncoa5 | 0.142 | 0.108 | 0.120 |
| E030030I06Ri | 0.142 | 0.040 | 0.050 |
| Prr5 | 0.142 | 0.024 | 0.183 |
| Bc1 | 0.142 | 0.110 | 0.130 |
| Vps11 | 0.142 | 0.134 | 0.115 |
| Hoxb3 | 0.142 | 0.310 | 0.238 |
| Rybp | 0.142 | 0.166 | 0.186 |
| Ahdc1 | 0.142 | 0.233 | 0.190 |
| Net1 | 0.142 | 0.358 | 0.089 |
| Adamts5 | 0.141 | 0.318 | 0.637 |
| Dnajc18 | 0.141 | 0.194 | 0.184 |
| Fra10ac1 | 0.141 | 0.127 | 0.109 |
| Pik3c3 | 0.141 | 0.090 | 0.093 |

| | | | |
|---|---|---|---|
| Rbm14 | 0.141 | 0.089 | 0.096 |
| Rbbp9 | 0.141 | 0.157 | 0.147 |
| Banp | 0.141 | 0.112 | 0.137 |
| Uchl3 | 0.141 | 0.141 | 0.179 |
| Tmem80 | 0.141 | 0.107 | 0.116 |
| Kti12 | 0.141 | 0.077 | 0.081 |
| Rnpc3 | 0.141 | 0.136 | 0.123 |
| Dhdds | 0.141 | 0.116 | 0.121 |
| Khk | 0.141 | 0.122 | 0.153 |
| Rnaseh2a | 0.141 | 0.123 | 0.129 |
| Ctps2 | 0.141 | 0.221 | 0.209 |
| Oxsr1 | 0.141 | 0.100 | 0.094 |
| Dmac2 | 0.141 | 0.078 | 0.062 |
| Crat | 0.141 | 0.132 | 0.158 |
| Tmem251 | 0.141 | 0.158 | 0.116 |
| Tma16 | 0.141 | 0.119 | 0.132 |
| Pgam5 | 0.141 | 0.090 | 0.101 |
| Hif3a | 0.141 | 0.158 | 0.158 |
| Gm49336 | 0.141 | 0.117 | 0.128 |
| Edc4 | 0.141 | 0.127 | 0.130 |
| Tdp2 | 0.141 | 0.191 | 0.217 |
| Slc25a20 | 0.141 | 0.096 | 0.075 |
| Anapc15 | 0.140 | 0.114 | 0.096 |
| Usp24 | 0.140 | 0.107 | 0.112 |
| Trappc9 | 0.140 | 0.110 | 0.102 |
| Dgcr8 | 0.140 | 0.118 | 0.134 |
| Mut | 0.140 | 0.153 | 0.135 |
| Zfp263 | 0.140 | 0.115 | 0.117 |
| Terf2 | 0.140 | 0.154 | 0.142 |
| Cse1l | 0.140 | 0.119 | 0.106 |
| B9d1 | 0.140 | 0.137 | 0.173 |
| Snapc2 | 0.140 | 0.127 | 0.101 |
| Srf | 0.140 | 0.204 | 0.146 |
| Plscr1 | 0.140 | 0.056 | 0.084 |
| Slc39a6 | 0.140 | 0.257 | 0.313 |
| Memo1 | 0.140 | 0.164 | 0.163 |
| Ap1b1 | 0.140 | 0.111 | 0.100 |
| Hdac5 | 0.140 | 0.322 | 0.259 |
| Mfsd4a | 0.140 | 0.079 | 0.078 |
| Bhlhb9 | 0.140 | 0.197 | 0.178 |
| Polr1a | 0.140 | 0.130 | 0.124 |
| Raet1e | 0.140 | 0.003 | 0.004 |
| Mcat | 0.140 | 0.120 | 0.133 |
| Adarb1 | 0.140 | 0.329 | 0.143 |
| Pknox2 | 0.140 | 0.049 | 0.069 |
| Enpp5 | 0.140 | 0.255 | 0.170 |
| Socs6 | 0.140 | 0.151 | 0.132 |
| Wdtc1 | 0.139 | 0.128 | 0.120 |

| | | | |
|---|---|---|---|
| Pik3r4 | 0.139 | 0.112 | 0.128 |
| Abcd4 | 0.139 | 0.112 | 0.113 |
| Naprt | 0.139 | 0.118 | 0.127 |
| Klhl4 | 0.139 | 0.002 | 0.002 |
| Crls1 | 0.139 | 0.160 | 0.160 |
| Coq5 | 0.139 | 0.122 | 0.133 |
| Cramp1l | 0.139 | 0.163 | 0.150 |
| Naf1 | 0.139 | 0.134 | 0.102 |
| D3Ertd254e | 0.139 | 0.129 | 0.139 |
| Smg5 | 0.139 | 0.139 | 0.113 |
| Cul4b | 0.139 | 0.159 | 0.154 |
| Prps1 | 0.139 | 0.140 | 0.106 |
| Zfp523 | 0.139 | 0.127 | 0.122 |
| Fdft1 | 0.139 | 0.224 | 0.176 |
| Jpt2 | 0.139 | 0.064 | 0.093 |
| Msrb2 | 0.139 | 0.104 | 0.103 |
| Prpf4 | 0.139 | 0.130 | 0.143 |
| Pex7 | 0.139 | 0.139 | 0.124 |
| 3110002H16I | 0.139 | 0.081 | 0.046 |
| Fbxl15 | 0.139 | 0.133 | 0.117 |
| Snapc1 | 0.139 | 0.113 | 0.100 |
| Hmox1 | 0.139 | 0.124 | 0.141 |
| Gtf3c3 | 0.139 | 0.107 | 0.103 |
| Tmem181a | 0.139 | 0.069 | 0.082 |
| Wdr73 | 0.139 | 0.069 | 0.058 |
| Mrgbp | 0.139 | 0.080 | 0.079 |
| Scaper | 0.138 | 0.179 | 0.159 |
| Clca3a2 | 0.138 | 1.196 | 0.577 |
| Ostm1 | 0.138 | 0.064 | 0.061 |
| Npnt | 0.138 | 0.221 | 0.034 |
| Coa7 | 0.138 | 0.093 | 0.082 |
| Rhbdf2 | 0.138 | 0.040 | 0.016 |
| Fbxo33 | 0.138 | 0.212 | 0.176 |
| Glmn | 0.138 | 0.148 | 0.171 |
| Camta2 | 0.138 | 0.098 | 0.078 |
| Mrs2 | 0.138 | 0.074 | 0.066 |
| 3110062M04 | 0.138 | 0.061 | 0.069 |
| Bcl2l13 | 0.138 | 0.097 | 0.093 |
| Dis3l2 | 0.138 | 0.077 | 0.088 |
| S100pbp | 0.138 | 0.123 | 0.115 |
| Bnip1 | 0.138 | 0.130 | 0.130 |
| Mansc1 | 0.138 | 0.033 | 0.021 |
| Ppm1d | 0.138 | 0.161 | 0.134 |
| Gk5 | 0.138 | 0.108 | 0.138 |
| Shb | 0.138 | 0.113 | 0.069 |
| Phgdh | 0.138 | 0.370 | 0.349 |
| Mrrf | 0.138 | 0.156 | 0.144 |
| Tox | 0.138 | 0.011 | 0.025 |

| | | | |
|---|---|---|---|
| Sdf2l1 | 0.138 | 0.028 | 0.053 |
| Usf1 | 0.138 | 0.120 | 0.107 |
| Orc3 | 0.138 | 0.122 | 0.139 |
| Cdc14a | 0.137 | 0.072 | 0.054 |
| Bcl11a | 0.137 | 0.300 | 0.089 |
| Prr3 | 0.137 | 0.110 | 0.124 |
| Cog6 | 0.137 | 0.104 | 0.121 |
| Cstf3 | 0.137 | 0.197 | 0.165 |
| Bcl2l12 | 0.137 | 0.156 | 0.136 |
| Camk2n1 | 0.137 | 0.400 | 0.579 |
| Dpt | 0.137 | 0.288 | 1.951 |
| 9-Sep | 0.137 | 0.289 | 0.182 |
| Mapk8ip3 | 0.137 | 0.158 | 0.151 |
| Mad2l2 | 0.137 | 0.053 | 0.062 |
| Fam160a2 | 0.137 | 0.101 | 0.093 |
| Gtf2h4 | 0.137 | 0.116 | 0.118 |
| Crybg3 | 0.137 | 0.192 | 0.130 |
| Trim12a | 0.137 | 0.084 | 0.132 |
| Ddx58 | 0.137 | 0.104 | 0.100 |
| Sgf29 | 0.137 | 0.118 | 0.124 |
| Nop9 | 0.137 | 0.125 | 0.110 |
| Tmem88 | 0.137 | 0.108 | 0.137 |
| Cab39l | 0.137 | 0.081 | 0.129 |
| Slc4a1ap | 0.137 | 0.112 | 0.101 |
| Ikbkap | 0.137 | 0.126 | 0.160 |
| Thoc6 | 0.137 | 0.135 | 0.106 |
| Jade1 | 0.137 | 0.168 | 0.156 |
| St5 | 0.136 | 0.212 | 0.182 |
| Cdc27 | 0.136 | 0.143 | 0.152 |
| Ddx55 | 0.136 | 0.104 | 0.095 |
| Cox6b2 | 0.136 | 0.172 | 0.173 |
| Cpox | 0.136 | 0.117 | 0.150 |
| Hacd3 | 0.136 | 0.119 | 0.094 |
| Ly96 | 0.136 | 0.087 | 0.105 |
| Coq10a | 0.136 | 0.129 | 0.104 |
| Aatf | 0.136 | 0.131 | 0.138 |
| Cep68 | 0.136 | 0.091 | 0.083 |
| St7 | 0.136 | 0.146 | 0.168 |
| Eml1 | 0.136 | 0.264 | 0.282 |
| Kmt2d | 0.136 | 0.168 | 0.133 |
| 1190002N15I | 0.136 | 0.213 | 0.130 |
| Gbp3 | 0.136 | 0.013 | 0.045 |
| Olfm1 | 0.136 | 0.113 | 0.263 |
| Dgkh | 0.136 | 0.347 | 0.165 |
| Rpgrip1 | 0.136 | 0.149 | 0.120 |
| Cdh3 | 0.136 | 0.176 | 0.057 |
| Acbd4 | 0.136 | 0.146 | 0.132 |
| Akt2 | 0.136 | 0.291 | 0.444 |

| | | | |
|---|---|---|---|
| Ncdn | 0.136 | 0.114 | 0.117 |
| Rnf8 | 0.136 | 0.129 | 0.130 |
| Slc7a7 | 0.136 | 0.047 | 0.058 |
| Rbbp8 | 0.136 | 0.148 | 0.119 |
| Hand2 | 0.135 | 0.131 | 0.185 |
| Apold1 | 0.135 | 0.007 | 0.005 |
| Sipa1l2 | 0.135 | 0.261 | 0.182 |
| Corin | 0.135 | 0.168 | 0.055 |
| Fam13b | 0.135 | 0.133 | 0.120 |
| Egln3 | 0.135 | 0.673 | 0.151 |
| Fam217b | 0.135 | 0.117 | 0.052 |
| Mbd5 | 0.135 | 0.173 | 0.144 |
| Dpp9 | 0.135 | 0.152 | 0.186 |
| Ip6k2 | 0.135 | 0.180 | 0.154 |
| D030056L22F | 0.135 | 0.089 | 0.108 |
| Pdhx | 0.135 | 0.110 | 0.095 |
| Rpa2 | 0.135 | 0.125 | 0.173 |
| 4931406C07F | 0.135 | 0.138 | 0.160 |
| A130010J15R | 0.135 | 0.052 | 0.055 |
| Lzic | 0.135 | 0.126 | 0.108 |
| Filip1 | 0.135 | 0.414 | 0.122 |
| Lfng | 0.135 | 0.046 | 0.052 |
| Gm20186 | 0.135 | 0.255 | 0.117 |
| H2-Q7 | 0.135 | 0.003 | 0.003 |
| Vps37b | 0.135 | 0.273 | 0.097 |
| Mirg | 0.135 | 0.119 | 0.377 |
| Dip2a | 0.135 | 0.103 | 0.113 |
| Slc25a44 | 0.135 | 0.107 | 0.105 |
| Mthfr | 0.135 | 0.074 | 0.088 |
| Pip4k2c | 0.135 | 0.148 | 0.111 |
| Slc39a14 | 0.135 | 0.211 | 0.147 |
| Kif13b | 0.135 | 0.106 | 0.097 |
| Tceal3 | 0.135 | 0.237 | 0.103 |
| Ppih | 0.135 | 0.115 | 0.136 |
| Gm16104 | 0.134 | 0.002 | 0.003 |
| Ralgapb | 0.134 | 0.104 | 0.090 |
| Zfp110 | 0.134 | 0.097 | 0.122 |
| Cdpf1 | 0.134 | 0.130 | 0.134 |
| Dvl3 | 0.134 | 0.230 | 0.182 |
| Slc25a38 | 0.134 | 0.101 | 0.085 |
| Mkrn2 | 0.134 | 0.132 | 0.132 |
| Tubgcp5 | 0.134 | 0.123 | 0.114 |
| Ly6g6d | 0.134 | 0.006 | 0.002 |
| Esco1 | 0.134 | 0.130 | 0.097 |
| Trappc8 | 0.134 | 0.119 | 0.111 |
| Wisp1 | 0.134 | 0.761 | 0.565 |
| Mvk | 0.134 | 0.123 | 0.158 |
| Zfp64 | 0.134 | 0.097 | 0.108 |

| | | | |
|---|---|---|---|
| Rgs2 | 0.134 | 0.366 | 0.127 |
| Capn15 | 0.134 | 0.118 | 0.130 |
| Vps13b | 0.134 | 0.110 | 0.148 |
| Mnt | 0.134 | 0.194 | 0.166 |
| Camsap1 | 0.134 | 0.223 | 0.189 |
| Kirrel | 0.134 | 0.122 | 0.113 |
| Plekhg3 | 0.134 | 0.515 | 0.324 |
| Wdfy2 | 0.134 | 0.115 | 0.162 |
| Haus2 | 0.134 | 0.102 | 0.096 |
| Slc9a8 | 0.134 | 0.098 | 0.108 |
| Ppox | 0.134 | 0.131 | 0.121 |
| Tle3 | 0.134 | 0.135 | 0.215 |
| Slc25a35 | 0.134 | 0.009 | 0.008 |
| Rabepk | 0.134 | 0.137 | 0.159 |
| Inpp5d | 0.134 | 0.002 | 0.003 |
| Frs2 | 0.133 | 0.208 | 0.192 |
| Uspl1 | 0.133 | 0.088 | 0.089 |
| Ppp1r21 | 0.133 | 0.109 | 0.114 |
| Itga6 | 0.133 | 0.044 | 0.038 |
| Wdr20 | 0.133 | 0.155 | 0.132 |
| 1700021F05F | 0.133 | 0.193 | 0.199 |
| Pdcd2 | 0.133 | 0.164 | 0.152 |
| Clk2 | 0.133 | 0.137 | 0.136 |
| Tsr1 | 0.133 | 0.109 | 0.125 |
| Dnaja3 | 0.133 | 0.080 | 0.101 |
| Dync1i1 | 0.133 | 0.008 | 0.015 |
| Prkab2 | 0.133 | 0.207 | 0.236 |
| Slc6a8 | 0.133 | 0.233 | 0.208 |
| Vps54 | 0.133 | 0.132 | 0.103 |
| Srcap | 0.133 | 0.128 | 0.105 |
| Qpctl | 0.133 | 0.112 | 0.090 |
| Xpot | 0.133 | 0.128 | 0.134 |
| Gtf2h2 | 0.133 | 0.114 | 0.114 |
| Rhod | 0.133 | 0.113 | 0.112 |
| Cdc14b | 0.133 | 0.152 | 0.117 |
| Ubap1 | 0.133 | 0.131 | 0.137 |
| Smg9 | 0.133 | 0.125 | 0.121 |
| Zfp979 | 0.133 | 0.014 | 0.021 |
| Trim3 | 0.133 | 0.150 | 0.147 |
| Ggcx | 0.133 | 0.100 | 0.171 |
| Usf3 | 0.133 | 0.102 | 0.075 |
| 2410089E03F | 0.133 | 0.186 | 0.189 |
| Abl2 | 0.133 | 0.104 | 0.094 |
| Myo10 | 0.133 | 0.143 | 0.122 |
| Gpatch4 | 0.133 | 0.093 | 0.085 |
| Usp36 | 0.133 | 0.168 | 0.170 |
| Tfpt | 0.133 | 0.131 | 0.122 |
| 2700062C07F | 0.133 | 0.127 | 0.127 |

| | | | |
|---|---|---|---|
| Cbx4 | 0.132 | 0.190 | 0.140 |
| Kctd3 | 0.132 | 0.124 | 0.128 |
| Sdr39u1 | 0.132 | 0.099 | 0.071 |
| Scyl2 | 0.132 | 0.077 | 0.079 |
| Phldb1 | 0.132 | 0.186 | 0.318 |
| Akr1e1 | 0.132 | 0.132 | 0.181 |
| Arhgap24 | 0.132 | 0.162 | 0.185 |
| Ube2o | 0.132 | 0.200 | 0.162 |
| Gprasp1 | 0.132 | 0.186 | 0.237 |
| Ly6a | 0.132 | 0.026 | 0.466 |
| Trib1 | 0.132 | 0.339 | 0.285 |
| Gse1 | 0.132 | 0.061 | 0.098 |
| Zfp282 | 0.132 | 0.115 | 0.128 |
| Vps50 | 0.132 | 0.154 | 0.122 |
| Snx10 | 0.132 | 0.075 | 0.115 |
| Sp110 | 0.132 | 0.025 | 0.041 |
| Rpp30 | 0.132 | 0.124 | 0.127 |
| Neto2 | 0.132 | 0.023 | 0.014 |
| Fbxo34 | 0.132 | 0.086 | 0.089 |
| Gabpa | 0.132 | 0.134 | 0.120 |
| Large1 | 0.132 | 0.250 | 0.235 |
| Klc4 | 0.132 | 0.091 | 0.072 |
| Ttl | 0.132 | 0.123 | 0.148 |
| Vsig10 | 0.131 | 0.090 | 0.094 |
| Pskh1 | 0.131 | 0.122 | 0.154 |
| E2f5 | 0.131 | 0.122 | 0.117 |
| Mex3d | 0.131 | 0.161 | 0.134 |
| Tab1 | 0.131 | 0.075 | 0.075 |
| Galk2 | 0.131 | 0.121 | 0.191 |
| Atp11c | 0.131 | 0.087 | 0.067 |
| Pld2 | 0.131 | 0.125 | 0.110 |
| Ints8 | 0.131 | 0.103 | 0.090 |
| Acot8 | 0.131 | 0.074 | 0.054 |
| Pibf1 | 0.131 | 0.137 | 0.133 |
| Megf8 | 0.131 | 0.094 | 0.111 |
| Top2a | 0.131 | 0.102 | 0.113 |
| Zbtb11os1 | 0.131 | 0.077 | 0.092 |
| Ints1 | 0.131 | 0.124 | 0.121 |
| Snap29 | 0.131 | 0.075 | 0.087 |
| Pir | 0.131 | 0.072 | 0.122 |
| Arrdc4 | 0.131 | 0.205 | 0.201 |
| Trmt6 | 0.131 | 0.128 | 0.115 |
| Klhdc4 | 0.131 | 0.108 | 0.092 |
| Krr1 | 0.131 | 0.147 | 0.113 |
| Reck | 0.131 | 0.147 | 0.192 |
| Rab11fip5 | 0.131 | 0.073 | 0.109 |
| Trmt13 | 0.131 | 0.091 | 0.101 |
| Snx13 | 0.131 | 0.104 | 0.085 |

| | | | |
|---|---|---|---|
| Morn2 | 0.131 | 0.061 | 0.082 |
| Phf11d | 0.131 | 0.047 | 0.102 |
| Rdh10 | 0.131 | 0.080 | 0.079 |
| Pdcd2l | 0.131 | 0.120 | 0.109 |
| 6030458C11I | 0.131 | 0.092 | 0.097 |
| Eif2b3 | 0.130 | 0.131 | 0.152 |
| Fam20c | 0.130 | 0.193 | 0.228 |
| Nfxl1 | 0.130 | 0.139 | 0.107 |
| Aspn | 0.130 | 0.790 | 1.844 |
| Rab27a | 0.130 | 0.051 | 0.007 |
| Mras | 0.130 | 0.170 | 0.113 |
| Fam189b | 0.130 | 0.160 | 0.188 |
| 1810055G02I | 0.130 | 0.219 | 0.279 |
| Adprm | 0.130 | 0.125 | 0.105 |
| Tjap1 | 0.130 | 0.122 | 0.143 |
| Appl2 | 0.130 | 0.113 | 0.153 |
| Agpat1 | 0.130 | 0.097 | 0.089 |
| Zfp524 | 0.130 | 0.100 | 0.111 |
| Shf | 0.130 | 0.085 | 0.110 |
| Unk | 0.130 | 0.111 | 0.104 |
| Ampd2 | 0.130 | 0.087 | 0.111 |
| Kcne3 | 0.130 | 0.002 | 0.003 |
| Slc9a3r1 | 0.130 | 0.186 | 0.089 |
| Ifrd2 | 0.130 | 0.108 | 0.110 |
| Zbtb11 | 0.130 | 0.129 | 0.124 |
| Iffo1 | 0.130 | 0.116 | 0.111 |
| Ncbp1 | 0.130 | 0.119 | 0.125 |
| Plcb1 | 0.130 | 0.154 | 0.128 |
| Rrnad1 | 0.130 | 0.108 | 0.100 |
| Tor3a | 0.130 | 0.094 | 0.085 |
| Agk | 0.129 | 0.111 | 0.113 |
| Ccrl2 | 0.129 | 0.038 | 0.024 |
| Tulp3 | 0.129 | 0.127 | 0.151 |
| Mettl7a1 | 0.129 | 0.082 | 0.063 |
| Alkbh8 | 0.129 | 0.135 | 0.115 |
| Vwa5a | 0.129 | 0.136 | 0.185 |
| Dennd4c | 0.129 | 0.117 | 0.113 |
| Yipf2 | 0.129 | 0.054 | 0.040 |
| Aff1 | 0.129 | 0.123 | 0.072 |
| Pik3r3 | 0.129 | 0.031 | 0.017 |
| Zbtb37 | 0.129 | 0.106 | 0.142 |
| Mafb | 0.129 | 0.236 | 0.265 |
| Taf6 | 0.129 | 0.114 | 0.124 |
| Rai1 | 0.129 | 0.139 | 0.133 |
| Ctxn1 | 0.129 | 0.347 | 0.176 |
| Mtmr14 | 0.129 | 0.111 | 0.112 |
| Wdr75 | 0.129 | 0.124 | 0.117 |
| Mterf3 | 0.129 | 0.136 | 0.103 |

| | | | |
|---|---|---|---|
| Atxn3 | 0.129 | 0.092 | 0.059 |
| 1700025G04I | 0.129 | 0.637 | 0.525 |
| Spsb2 | 0.129 | 0.231 | 0.165 |
| Ptpn23 | 0.129 | 0.135 | 0.132 |
| Nup54 | 0.129 | 0.139 | 0.134 |
| Aptx | 0.129 | 0.123 | 0.126 |
| Kat8 | 0.129 | 0.112 | 0.108 |
| Hyal1 | 0.129 | 0.098 | 0.092 |
| Gas8 | 0.129 | 0.143 | 0.152 |
| Samd1 | 0.129 | 0.219 | 0.249 |
| Hbegf | 0.129 | 0.183 | 0.065 |
| Cryzl2 | 0.129 | 0.074 | 0.105 |
| Usp6nl | 0.129 | 0.086 | 0.053 |
| Fbxo38 | 0.129 | 0.129 | 0.115 |
| Gpr137 | 0.129 | 0.079 | 0.089 |
| Armcx5 | 0.128 | 0.136 | 0.120 |
| Mrpl1 | 0.128 | 0.143 | 0.132 |
| Senp1 | 0.128 | 0.121 | 0.123 |
| Pabpc4l | 0.128 | 0.087 | 0.116 |
| Ankrd37 | 0.128 | 0.051 | 0.032 |
| Tmem243 | 0.128 | 0.118 | 0.097 |
| Sil1 | 0.128 | 0.121 | 0.167 |
| Sema4c | 0.128 | 0.137 | 0.145 |
| Trmt2a | 0.128 | 0.104 | 0.120 |
| Ccng1 | 0.128 | 0.205 | 0.279 |
| Slc25a24 | 0.128 | 0.128 | 0.171 |
| Sik2 | 0.128 | 0.157 | 0.114 |
| Cbll1 | 0.128 | 0.131 | 0.108 |
| Zfp46 | 0.128 | 0.171 | 0.154 |
| Fuom | 0.128 | 0.151 | 0.160 |
| Haus4 | 0.128 | 0.104 | 0.119 |
| 1810043G02I | 0.128 | 0.111 | 0.127 |
| Mettl5 | 0.128 | 0.136 | 0.122 |
| Pbx3 | 0.128 | 0.220 | 0.143 |
| Plekhh3 | 0.128 | 0.140 | 0.142 |
| Mkks | 0.128 | 0.069 | 0.073 |
| C1qtnf7 | 0.128 | 0.092 | 0.404 |
| Paip2b | 0.128 | 0.104 | 0.111 |
| Klc2 | 0.128 | 0.091 | 0.058 |
| Nup160 | 0.128 | 0.050 | 0.058 |
| Hif1an | 0.128 | 0.118 | 0.116 |
| Ube2c | 0.128 | 0.101 | 0.102 |
| Kpna6 | 0.128 | 0.147 | 0.108 |
| Vps33a | 0.128 | 0.074 | 0.073 |
| Fam173b | 0.127 | 0.085 | 0.105 |
| Tet1 | 0.127 | 0.166 | 0.144 |
| Rev1 | 0.127 | 0.107 | 0.112 |
| Agl | 0.127 | 0.238 | 0.145 |

| | | | |
|---|---|---|---|
| Rbm15b | 0.127 | 0.141 | 0.121 |
| Metap1d | 0.127 | 0.144 | 0.148 |
| Rrp36 | 0.127 | 0.082 | 0.087 |
| Taf5l | 0.127 | 0.105 | 0.096 |
| Eef1akmt2 | 0.127 | 0.126 | 0.125 |
| Tex2 | 0.127 | 0.073 | 0.055 |
| Nf2 | 0.127 | 0.104 | 0.109 |
| Zfp9 | 0.127 | 0.153 | 0.262 |
| Papd5 | 0.127 | 0.118 | 0.101 |
| Wdr46 | 0.127 | 0.113 | 0.105 |
| Dhx16 | 0.127 | 0.093 | 0.089 |
| Btbd10 | 0.127 | 0.144 | 0.134 |
| Mul1 | 0.127 | 0.112 | 0.116 |
| Zfp428 | 0.127 | 0.086 | 0.109 |
| Tmem260 | 0.127 | 0.099 | 0.112 |
| Atxn1l | 0.127 | 0.113 | 0.095 |
| Tbck | 0.127 | 0.105 | 0.171 |
| Nckap5l | 0.127 | 0.089 | 0.068 |
| Lpin2 | 0.127 | 0.101 | 0.141 |
| Kansl1l | 0.127 | 0.187 | 0.149 |
| Ppwd1 | 0.127 | 0.118 | 0.113 |
| Sik1 | 0.127 | 0.298 | 0.149 |
| Wrb | 0.126 | 0.206 | 0.107 |
| Tbcel | 0.126 | 0.157 | 0.160 |
| Sh3d19 | 0.126 | 0.130 | 0.313 |
| Pwwp2b | 0.126 | 0.046 | 0.042 |
| Gm26632 | 0.126 | 0.286 | 0.076 |
| Ldlr | 0.126 | 0.254 | 0.368 |
| Utp23 | 0.126 | 0.092 | 0.091 |
| Ankrd54 | 0.126 | 0.104 | 0.119 |
| Coro2b | 0.126 | 0.053 | 0.070 |
| Zfp414 | 0.126 | 0.143 | 0.153 |
| Xylt2 | 0.126 | 0.137 | 0.251 |
| Xaf1 | 0.126 | 0.038 | 0.069 |
| Dcaf1 | 0.126 | 0.114 | 0.105 |
| Cd302 | 0.126 | 0.227 | 0.457 |
| Abcc1 | 0.126 | 0.216 | 0.134 |
| Amer1 | 0.126 | 0.244 | 0.287 |
| Arl6 | 0.126 | 0.110 | 0.117 |
| C1qtnf2 | 0.126 | 0.312 | 0.784 |
| Coro7 | 0.126 | 0.059 | 0.081 |
| Phlda1 | 0.126 | 0.783 | 0.930 |
| Tnfsf10 | 0.126 | 0.001 | 0.001 |
| Cd248 | 0.126 | 0.753 | 0.884 |
| Cks2 | 0.126 | 0.131 | 0.111 |
| Tnpo2 | 0.125 | 0.123 | 0.110 |
| Kat14 | 0.125 | 0.109 | 0.135 |
| Unc119b | 0.125 | 0.084 | 0.086 |

| | | | |
|---|---|---|---|
| Xrcc6 | 0.125 | 0.098 | 0.100 |
| Fam98b | 0.125 | 0.125 | 0.117 |
| Ppip5k2 | 0.125 | 0.079 | 0.077 |
| Mms19 | 0.125 | 0.137 | 0.124 |
| Bmi1 | 0.125 | 0.149 | 0.111 |
| Nufip1 | 0.125 | 0.119 | 0.138 |
| Chchd4 | 0.125 | 0.098 | 0.069 |
| Isg15 | 0.125 | 0.071 | 0.068 |
| Mettl2 | 0.125 | 0.101 | 0.133 |
| Acsl3 | 0.125 | 0.160 | 0.168 |
| Insig1 | 0.125 | 0.179 | 0.246 |
| Opa3 | 0.125 | 0.134 | 0.121 |
| Thap4 | 0.125 | 0.123 | 0.139 |
| Zfp672 | 0.125 | 0.102 | 0.096 |
| Fbxo30 | 0.125 | 0.423 | 0.175 |
| Znhit3 | 0.125 | 0.135 | 0.134 |
| Rb1 | 0.125 | 0.087 | 0.035 |
| Racgap1 | 0.125 | 0.035 | 0.037 |
| Hspb8 | 0.125 | 0.293 | 0.119 |
| Eml3 | 0.125 | 0.090 | 0.112 |
| Lpin1 | 0.124 | 0.196 | 0.123 |
| Jdp2 | 0.124 | 0.140 | 0.294 |
| Acvr1b | 0.124 | 0.089 | 0.070 |
| Lgr4 | 0.124 | 0.203 | 0.122 |
| Dpy19l4 | 0.124 | 0.117 | 0.103 |
| Glrx | 0.124 | 0.207 | 0.148 |
| Gab1 | 0.124 | 0.212 | 0.223 |
| Tmem252 | 0.124 | 0.002 | 0.001 |
| Tmem97 | 0.124 | 0.105 | 0.216 |
| Ftl1-ps1 | 0.124 | 0.089 | 0.115 |
| Pde12 | 0.124 | 0.211 | 0.239 |
| Kdm3b | 0.124 | 0.142 | 0.115 |
| Rad54l2 | 0.124 | 0.126 | 0.090 |
| Sco2 | 0.124 | 0.104 | 0.107 |
| Rhpn2 | 0.124 | 0.009 | 0.010 |
| Nudcd1 | 0.124 | 0.087 | 0.108 |
| Il17rc | 0.124 | 0.176 | 0.156 |
| Ift57 | 0.124 | 0.153 | 0.145 |
| G2e3 | 0.124 | 0.122 | 0.111 |
| Scd1 | 0.124 | 0.155 | 0.134 |
| Mfn1 | 0.124 | 0.120 | 0.106 |
| Notch2 | 0.124 | 0.333 | 0.288 |
| Gtf2f2 | 0.124 | 0.090 | 0.083 |
| Fam110a | 0.124 | 0.072 | 0.083 |
| Selenoo | 0.124 | 0.111 | 0.104 |
| Gid4 | 0.123 | 0.168 | 0.151 |
| Tbc1d1 | 0.123 | 0.150 | 0.077 |
| Zfp30 | 0.123 | 0.163 | 0.146 |

| | | | |
|---|---|---|---|
| Tgfbrap1 | 0.123 | 0.091 | 0.079 |
| Fam219a | 0.123 | 0.046 | 0.046 |
| Rab3a | 0.123 | 0.224 | 0.166 |
| Rmdn1 | 0.123 | 0.126 | 0.125 |
| Ccar2 | 0.123 | 0.070 | 0.076 |
| Gm15417 | 0.123 | 0.181 | 0.081 |
| Tro | 0.123 | 0.213 | 0.284 |
| Pfkfb3 | 0.123 | 0.051 | 0.090 |
| Ado | 0.123 | 0.159 | 0.163 |
| Cwc22 | 0.123 | 0.153 | 0.115 |
| Atp1b1 | 0.123 | 0.809 | 0.709 |
| Msto1 | 0.123 | 0.073 | 0.073 |
| Clasrp | 0.123 | 0.141 | 0.132 |
| Frat2 | 0.123 | 0.068 | 0.097 |
| Nhlrc2 | 0.123 | 0.135 | 0.129 |
| Izumo4 | 0.123 | 0.135 | 0.143 |
| Zmat3 | 0.123 | 0.091 | 0.127 |
| Trub2 | 0.123 | 0.077 | 0.080 |
| Gpank1 | 0.123 | 0.089 | 0.065 |
| Dcaf5 | 0.123 | 0.139 | 0.117 |
| AU040320 | 0.123 | 0.057 | 0.067 |
| Acot1 | 0.123 | 0.202 | 0.258 |
| Gin1 | 0.123 | 0.113 | 0.098 |
| Cplx2 | 0.122 | 0.086 | 0.092 |
| Dtd2 | 0.122 | 0.141 | 0.140 |
| Med27 | 0.122 | 0.103 | 0.085 |
| E4f1 | 0.122 | 0.114 | 0.116 |
| Med11 | 0.122 | 0.051 | 0.057 |
| Ndst2 | 0.122 | 0.076 | 0.094 |
| Ucp2 | 0.122 | 0.044 | 0.100 |
| Urgcp | 0.122 | 0.122 | 0.119 |
| 5033406O09I | 0.122 | 0.001 | 0.006 |
| Lancl2 | 0.122 | 0.097 | 0.095 |
| Arsj | 0.122 | 0.011 | 0.068 |
| Spata7 | 0.122 | 0.121 | 0.085 |
| Cers6 | 0.122 | 0.045 | 0.076 |
| Rptor | 0.122 | 0.116 | 0.104 |
| Zfp609 | 0.122 | 0.132 | 0.146 |
| Pyroxd1 | 0.122 | 0.101 | 0.097 |
| Rnf213 | 0.122 | 0.054 | 0.073 |
| Gramd3 | 0.122 | 0.118 | 0.177 |
| Ebpl | 0.122 | 0.188 | 0.298 |
| Morc3 | 0.122 | 0.146 | 0.160 |
| Yod1 | 0.122 | 0.095 | 0.082 |
| Fam208b | 0.122 | 0.127 | 0.142 |
| Stat2 | 0.122 | 0.142 | 0.175 |
| Polr3f | 0.122 | 0.104 | 0.088 |
| Tbc1d14 | 0.122 | 0.102 | 0.119 |

| | | | |
|---|---|---|---|
| Rfng | 0.122 | 0.108 | 0.091 |
| Pi4ka | 0.122 | 0.145 | 0.145 |
| Alg3 | 0.122 | 0.077 | 0.074 |
| Ubxn8 | 0.122 | 0.110 | 0.096 |
| Nol8 | 0.122 | 0.098 | 0.079 |
| Prkaa1 | 0.121 | 0.119 | 0.092 |
| C1qtnf1 | 0.121 | 0.124 | 0.103 |
| Ttc37 | 0.121 | 0.111 | 0.125 |
| Wdsub1 | 0.121 | 0.147 | 0.143 |
| Traf2 | 0.121 | 0.068 | 0.073 |
| Thumpd3 | 0.121 | 0.076 | 0.082 |
| Zfp592 | 0.121 | 0.129 | 0.155 |
| Zfand1 | 0.121 | 0.123 | 0.118 |
| Frmd8 | 0.121 | 0.097 | 0.076 |
| Rasgef1b | 0.121 | 0.029 | 0.054 |
| Psme2b | 0.121 | 0.013 | 0.008 |
| Terf1 | 0.121 | 0.123 | 0.111 |
| Dennd1a | 0.121 | 0.128 | 0.139 |
| Tpra1 | 0.121 | 0.070 | 0.066 |
| Tradd | 0.121 | 0.103 | 0.141 |
| Prpf40b | 0.121 | 0.089 | 0.070 |
| Nfya | 0.121 | 0.068 | 0.073 |
| Ccdc68 | 0.121 | 0.081 | 0.036 |
| Exoc6 | 0.121 | 0.035 | 0.014 |
| Tes | 0.121 | 0.123 | 0.017 |
| Fbxl22 | 0.121 | 0.574 | 0.127 |
| Arhgap6 | 0.121 | 0.154 | 0.152 |
| Casp7 | 0.121 | 0.061 | 0.056 |
| Dlg5 | 0.121 | 0.134 | 0.157 |
| B3galt6 | 0.121 | 0.068 | 0.093 |
| Acer3 | 0.121 | 0.147 | 0.109 |
| Etv3 | 0.121 | 0.115 | 0.108 |
| Atf5 | 0.121 | 0.224 | 0.565 |
| Fxyd1 | 0.121 | 1.154 | 0.870 |
| Reep2 | 0.121 | 0.122 | 0.050 |
| Cstf2t | 0.121 | 0.071 | 0.073 |
| Cabp1 | 0.120 | 0.011 | 0.004 |
| Zfp444 | 0.120 | 0.106 | 0.086 |
| Fbxw17 | 0.120 | 0.077 | 0.086 |
| Pdk2 | 0.120 | 0.108 | 0.110 |
| Tapt1 | 0.120 | 0.141 | 0.089 |
| Znrf2 | 0.120 | 0.064 | 0.051 |
| Foxo3 | 0.120 | 0.248 | 0.266 |
| Lpcat2 | 0.120 | 0.038 | 0.014 |
| Cenpc1 | 0.120 | 0.119 | 0.111 |
| Micu3 | 0.120 | 0.208 | 0.131 |
| Sobp | 0.120 | 0.258 | 0.213 |
| Fam57a | 0.120 | 0.026 | 0.045 |

| | | | |
|---|---|---|---|
| Hectd4 | 0.120 | 0.097 | 0.098 |
| Hcfc2 | 0.120 | 0.113 | 0.137 |
| Npr2 | 0.120 | 0.186 | 0.189 |
| Rbl2 | 0.120 | 0.149 | 0.135 |
| Ak1 | 0.120 | 0.152 | 0.141 |
| Rnf169 | 0.120 | 0.122 | 0.098 |
| Agtpbp1 | 0.120 | 0.139 | 0.129 |
| Cntrl | 0.120 | 0.125 | 0.119 |
| Thop1 | 0.120 | 0.118 | 0.137 |
| Fam213a | 0.120 | 0.046 | 0.058 |
| Atg10 | 0.120 | 0.100 | 0.115 |
| Pxylp1 | 0.120 | 0.055 | 0.047 |
| Pdk1 | 0.120 | 0.083 | 0.090 |
| Slc10a7 | 0.120 | 0.087 | 0.109 |
| Dgat1 | 0.120 | 0.084 | 0.084 |
| Nbas | 0.120 | 0.087 | 0.125 |
| E130309D02F | 0.120 | 0.084 | 0.086 |
| B3gnt9 | 0.120 | 0.336 | 0.438 |
| Abi2 | 0.120 | 0.255 | 0.238 |
| Arhgap8 | 0.119 | 0.004 | 0.005 |
| Taf1b | 0.119 | 0.112 | 0.079 |
| Tcof1 | 0.119 | 0.167 | 0.188 |
| Nup153 | 0.119 | 0.137 | 0.115 |
| Tomm6 | 0.119 | 0.112 | 0.113 |
| Gsdmd | 0.119 | 0.033 | 0.079 |
| Ttc17 | 0.119 | 0.117 | 0.104 |
| Tgif2 | 0.119 | 0.152 | 0.119 |
| Rpe | 0.119 | 0.098 | 0.093 |
| Ccnjl | 0.119 | 0.050 | 0.044 |
| Cep162 | 0.119 | 0.076 | 0.086 |
| Tmem19 | 0.119 | 0.136 | 0.149 |
| Rpf2 | 0.119 | 0.122 | 0.108 |
| Nudt2 | 0.119 | 0.087 | 0.126 |
| Elmsan1 | 0.119 | 0.128 | 0.143 |
| Fam53c | 0.119 | 0.149 | 0.163 |
| Scaf4 | 0.119 | 0.117 | 0.097 |
| Pigm | 0.119 | 0.063 | 0.073 |
| Pfkp | 0.119 | 0.440 | 0.200 |
| Prelid2 | 0.119 | 0.103 | 0.073 |
| E2f6 | 0.119 | 0.102 | 0.079 |
| Trp53inp2 | 0.119 | 0.142 | 0.114 |
| Adpgk | 0.119 | 0.089 | 0.107 |
| Sox12 | 0.119 | 0.201 | 0.150 |
| 8-Sep | 0.119 | 0.254 | 0.429 |
| Dcxr | 0.119 | 0.111 | 0.069 |
| Pip4k2a | 0.119 | 0.101 | 0.103 |
| Rassf8 | 0.119 | 0.261 | 0.305 |
| Igf2bp1 | 0.119 | 0.054 | 0.102 |

| | | | |
|---|---|---|---|
| Vasn | 0.119 | 0.475 | 0.370 |
| Slc35b4 | 0.118 | 0.103 | 0.115 |
| Miga1 | 0.118 | 0.090 | 0.083 |
| Slc7a6os | 0.118 | 0.181 | 0.143 |
| Polr3h | 0.118 | 0.106 | 0.109 |
| Stc1 | 0.118 | 0.140 | 0.248 |
| Cln8 | 0.118 | 0.080 | 0.107 |
| Zfp275 | 0.118 | 0.117 | 0.116 |
| Antxr2 | 0.118 | 0.163 | 0.195 |
| Ing5 | 0.118 | 0.123 | 0.122 |
| Cdk2 | 0.118 | 0.113 | 0.095 |
| Ralgps2 | 0.118 | 0.044 | 0.167 |
| Mapk6 | 0.118 | 0.154 | 0.155 |
| Vps51 | 0.118 | 0.062 | 0.048 |
| Rbm43 | 0.118 | 0.078 | 0.072 |
| Mynn | 0.118 | 0.133 | 0.122 |
| Cdkal1 | 0.118 | 0.122 | 0.116 |
| Pitrm1 | 0.118 | 0.194 | 0.220 |
| Vangl2 | 0.118 | 0.093 | 0.091 |
| Mbd4 | 0.118 | 0.040 | 0.051 |
| Dync2h1 | 0.118 | 0.153 | 0.152 |
| 1110032F04F | 0.118 | 0.001 | 0.001 |
| Cdc25a | 0.118 | 0.115 | 0.109 |
| Man2c1 | 0.118 | 0.119 | 0.134 |
| Zfp26 | 0.118 | 0.123 | 0.104 |
| Tbc1d8b | 0.118 | 0.090 | 0.104 |
| Tars2 | 0.118 | 0.076 | 0.069 |
| Ecsit | 0.117 | 0.176 | 0.146 |
| Srrd | 0.117 | 0.100 | 0.092 |
| Capn1 | 0.117 | 0.117 | 0.204 |
| Hspbp1 | 0.117 | 0.112 | 0.097 |
| Fam160b2 | 0.117 | 0.086 | 0.090 |
| Ppp1r16a | 0.117 | 0.094 | 0.118 |
| Syvn1 | 0.117 | 0.101 | 0.101 |
| Zfp553 | 0.117 | 0.091 | 0.086 |
| Galnt7 | 0.117 | 0.044 | 0.068 |
| Aven | 0.117 | 0.098 | 0.117 |
| Abtb1 | 0.117 | 0.113 | 0.110 |
| Tmtc3 | 0.117 | 0.107 | 0.078 |
| Cog7 | 0.117 | 0.089 | 0.089 |
| Rabl2 | 0.117 | 0.094 | 0.101 |
| Cradd | 0.117 | 0.089 | 0.070 |
| Gm4117 | 0.117 | 0.137 | 0.075 |
| Gemin2 | 0.117 | 0.073 | 0.064 |
| Rbm12 | 0.117 | 0.103 | 0.117 |
| Gm26782 | 0.117 | 0.138 | 0.160 |
| Cpt2 | 0.117 | 0.112 | 0.141 |
| Stam | 0.117 | 0.155 | 0.179 |

| | | | |
|---|---|---|---|
| Ank | 0.117 | 0.148 | 0.093 |
| Asb3 | 0.117 | 0.131 | 0.146 |
| Etfrf1 | 0.117 | 0.117 | 0.089 |
| Gzf1 | 0.117 | 0.077 | 0.079 |
| Slc35a3 | 0.117 | 0.067 | 0.055 |
| Dip2c | 0.117 | 0.187 | 0.152 |
| Chad | 0.117 | 0.009 | 0.005 |
| Pex5 | 0.116 | 0.136 | 0.131 |
| Tmem135 | 0.116 | 0.088 | 0.120 |
| Zfp768 | 0.116 | 0.103 | 0.118 |
| Pomt2 | 0.116 | 0.064 | 0.079 |
| Cnnm3 | 0.116 | 0.099 | 0.105 |
| Fam107a | 0.116 | 0.055 | 0.033 |
| Triqk | 0.116 | 0.141 | 0.131 |
| Hs3st1 | 0.116 | 0.005 | 0.076 |
| Naa25 | 0.116 | 0.107 | 0.078 |
| Enoph1 | 0.116 | 0.077 | 0.093 |
| Cstf2 | 0.116 | 0.080 | 0.075 |
| Trim12c | 0.116 | 0.065 | 0.092 |
| Ceacam1 | 0.116 | 0.001 | 0.002 |
| Tbc1d7 | 0.116 | 0.132 | 0.165 |
| Enho | 0.116 | 0.022 | 0.051 |
| Apoo | 0.116 | 0.118 | 0.105 |
| Als2cl | 0.116 | 0.010 | 0.004 |
| Riok2 | 0.116 | 0.109 | 0.107 |
| Scrib | 0.116 | 0.092 | 0.089 |
| Toe1 | 0.116 | 0.154 | 0.141 |
| Cluap1 | 0.116 | 0.136 | 0.139 |
| Capn10 | 0.116 | 0.067 | 0.082 |
| Ntng1 | 0.116 | 0.019 | 0.034 |
| Ino80c | 0.116 | 0.097 | 0.097 |
| Ercc3 | 0.116 | 0.091 | 0.077 |
| 9430038I01R | 0.116 | 0.096 | 0.127 |
| Wdr74 | 0.116 | 0.110 | 0.099 |
| Eipr1 | 0.116 | 0.067 | 0.053 |
| Tbc1d19 | 0.116 | 0.106 | 0.106 |
| Cep120 | 0.116 | 0.118 | 0.101 |
| Nedd1 | 0.116 | 0.090 | 0.097 |
| Relb | 0.116 | 0.171 | 0.166 |
| Kptn | 0.115 | 0.106 | 0.098 |
| Cep85l | 0.115 | 0.107 | 0.140 |
| Washc5 | 0.115 | 0.105 | 0.132 |
| Fbxl12 | 0.115 | 0.078 | 0.074 |
| Sncg | 0.115 | 0.022 | 0.008 |
| Cep295 | 0.115 | 0.107 | 0.099 |
| Wee1 | 0.115 | 0.112 | 0.150 |
| Slc25a12 | 0.115 | 0.129 | 0.133 |
| Soga1 | 0.115 | 0.097 | 0.121 |

| | | | |
|---|---|---|---|
| Zkscan17 | 0.115 | 0.130 | 0.102 |
| Parp9 | 0.115 | 0.070 | 0.060 |
| Endod1 | 0.115 | 0.300 | 0.115 |
| Cbr3 | 0.115 | 0.048 | 0.131 |
| H2-Q6 | 0.115 | 0.002 | 0.001 |
| Cttnbp2 | 0.115 | 0.059 | 0.057 |
| Pus1 | 0.115 | 0.101 | 0.112 |
| Saal1 | 0.115 | 0.126 | 0.148 |
| Rad51d | 0.115 | 0.086 | 0.078 |
| Zdhhc17 | 0.115 | 0.110 | 0.101 |
| Wdfy1 | 0.115 | 0.327 | 0.164 |
| Rarres2 | 0.115 | 0.853 | 1.225 |
| Pold2 | 0.115 | 0.135 | 0.152 |
| 4930503L19F | 0.115 | 0.082 | 0.058 |
| Ppp1r12b | 0.115 | 0.459 | 0.260 |
| Gm30025 | 0.115 | 0.104 | 0.154 |
| Mki67 | 0.115 | 0.063 | 0.070 |
| Mapk7 | 0.115 | 0.101 | 0.161 |
| Ubqln4 | 0.115 | 0.093 | 0.096 |
| Ocel1 | 0.115 | 0.069 | 0.068 |
| Gstm7 | 0.115 | 0.072 | 0.032 |
| Cutc | 0.115 | 0.086 | 0.099 |
| Vps37c | 0.115 | 0.075 | 0.069 |
| Zfp407 | 0.115 | 0.078 | 0.078 |
| Hmces | 0.115 | 0.120 | 0.096 |
| Bach2 | 0.114 | 0.073 | 0.173 |
| Ptges2 | 0.114 | 0.103 | 0.101 |
| Zfp703 | 0.114 | 0.311 | 0.311 |
| Aldh3a2 | 0.114 | 0.107 | 0.112 |
| Atg2a | 0.114 | 0.110 | 0.107 |
| Pptc7 | 0.114 | 0.147 | 0.097 |
| Sephs2 | 0.114 | 0.112 | 0.097 |
| Bcor | 0.114 | 0.279 | 0.174 |
| Nr1d2 | 0.114 | 0.117 | 0.098 |
| Aldh9a1 | 0.114 | 0.057 | 0.072 |
| Casp9 | 0.114 | 0.082 | 0.085 |
| Bicd1 | 0.114 | 0.369 | 0.164 |
| Xrcc5 | 0.114 | 0.125 | 0.137 |
| Thoc5 | 0.114 | 0.105 | 0.090 |
| Fer | 0.114 | 0.126 | 0.171 |
| Osbpl11 | 0.114 | 0.133 | 0.124 |
| Pcca | 0.114 | 0.077 | 0.065 |
| Unc93b1 | 0.114 | 0.030 | 0.053 |
| Tmem218 | 0.114 | 0.155 | 0.191 |
| Cxcl5 | 0.114 | 0.006 | 0.012 |
| Fzd3 | 0.114 | 0.110 | 0.086 |
| Kmt5c | 0.114 | 0.069 | 0.089 |
| Rprm | 0.114 | 0.003 | 0.002 |

| | | | |
|---|---|---|---|
| Usp42 | 0.114 | 0.110 | 0.138 |
| Alg2 | 0.114 | 0.094 | 0.118 |
| Gimap9 | 0.114 | 0.006 | 0.016 |
| Cx3cl1 | 0.114 | 0.027 | 0.006 |
| Tbc1d5 | 0.114 | 0.089 | 0.114 |
| Fbxo45 | 0.114 | 0.075 | 0.080 |
| Spns2 | 0.113 | 0.002 | 0.003 |
| Mroh1 | 0.113 | 0.115 | 0.073 |
| Med14 | 0.113 | 0.114 | 0.085 |
| Plekha8 | 0.113 | 0.054 | 0.060 |
| Tmem150a | 0.113 | 0.159 | 0.174 |
| Ankle2 | 0.113 | 0.143 | 0.143 |
| Telo2 | 0.113 | 0.147 | 0.163 |
| Tubgcp4 | 0.113 | 0.127 | 0.136 |
| Dhx32 | 0.113 | 0.175 | 0.118 |
| Celsr2 | 0.113 | 0.065 | 0.015 |
| Gm5914 | 0.113 | 0.168 | 0.112 |
| B4galt6 | 0.113 | 0.205 | 0.113 |
| Abat | 0.113 | 0.140 | 0.100 |
| Clmp | 0.113 | 0.558 | 0.718 |
| Tubb6 | 0.113 | 0.372 | 0.583 |
| Ppp6r2 | 0.113 | 0.109 | 0.103 |
| Tbl3 | 0.113 | 0.095 | 0.094 |
| Hist3h2ba | 0.113 | 0.070 | 0.047 |
| Lrpprc | 0.113 | 0.128 | 0.119 |
| Loxl2 | 0.113 | 0.413 | 0.443 |
| Hacd2 | 0.113 | 0.233 | 0.134 |
| Atat1 | 0.113 | 0.163 | 0.176 |
| Gmeb1 | 0.113 | 0.100 | 0.107 |
| Myo1e | 0.113 | 0.207 | 0.054 |
| Eef1akmt4 | 0.113 | 0.049 | 0.069 |
| Sirt1 | 0.113 | 0.174 | 0.142 |
| Impa2 | 0.113 | 0.153 | 0.082 |
| Sos1 | 0.113 | 0.140 | 0.128 |
| Gtf3c4 | 0.113 | 0.108 | 0.166 |
| Map3k12 | 0.113 | 0.141 | 0.170 |
| 1700096K18F | 0.113 | 0.076 | 0.083 |
| Tnk2 | 0.113 | 0.130 | 0.080 |
| Miip | 0.113 | 0.084 | 0.058 |
| 2810001G20I | 0.113 | 0.109 | 0.109 |
| Klhl20 | 0.113 | 0.072 | 0.071 |
| Atic | 0.113 | 0.178 | 0.168 |
| Psph | 0.113 | 0.101 | 0.091 |
| Zfyve16 | 0.113 | 0.108 | 0.128 |
| Cacna1a | 0.112 | 0.051 | 0.097 |
| Pmf1 | 0.112 | 0.110 | 0.089 |
| Mis18a | 0.112 | 0.047 | 0.062 |
| Tap1 | 0.112 | 0.055 | 0.070 |

| | | | |
|---|---|---|---|
| Ipo8 | 0.112 | 0.095 | 0.089 |
| Birc5 | 0.112 | 0.057 | 0.075 |
| Pard6a | 0.112 | 0.079 | 0.084 |
| Uggt2 | 0.112 | 0.107 | 0.096 |
| Itgb3bp | 0.112 | 0.082 | 0.110 |
| Epha5 | 0.112 | 0.012 | 0.008 |
| Papss2 | 0.112 | 0.103 | 0.163 |
| Rpusd4 | 0.112 | 0.057 | 0.059 |
| Abhd2 | 0.112 | 0.202 | 0.062 |
| Lrp1 | 0.112 | 1.071 | 1.350 |
| Hps4 | 0.112 | 0.089 | 0.100 |
| Scap | 0.112 | 0.116 | 0.121 |
| Adam19 | 0.112 | 0.261 | 0.218 |
| Hars2 | 0.112 | 0.096 | 0.086 |
| Brms1l | 0.112 | 0.094 | 0.098 |
| Armcx6 | 0.112 | 0.119 | 0.177 |
| Dhx57 | 0.112 | 0.076 | 0.089 |
| Hbb-y | 0.112 | 0.000 | 0.002 |
| Ric1 | 0.112 | 0.146 | 0.125 |
| Cdca4 | 0.112 | 0.120 | 0.106 |
| Klhl28 | 0.112 | 0.105 | 0.123 |
| Cry1 | 0.112 | 0.131 | 0.131 |
| Tom1l2 | 0.112 | 0.135 | 0.104 |
| Zfp120 | 0.112 | 0.070 | 0.099 |
| Ppp1r8 | 0.111 | 0.098 | 0.094 |
| Pi4k2b | 0.111 | 0.129 | 0.183 |
| Rab6b | 0.111 | 0.019 | 0.080 |
| Tnip2 | 0.111 | 0.110 | 0.099 |
| Ap5m1 | 0.111 | 0.087 | 0.079 |
| Caap1 | 0.111 | 0.112 | 0.099 |
| Anks3 | 0.111 | 0.137 | 0.182 |
| Ebag9 | 0.111 | 0.143 | 0.144 |
| AI987944 | 0.111 | 0.105 | 0.106 |
| Gmcl1 | 0.111 | 0.107 | 0.110 |
| Ascc2 | 0.111 | 0.075 | 0.090 |
| Mterf4 | 0.111 | 0.104 | 0.101 |
| Sord | 0.111 | 0.161 | 0.177 |
| Tst | 0.111 | 0.188 | 0.118 |
| Lrrc14 | 0.111 | 0.103 | 0.108 |
| Pcid2 | 0.111 | 0.081 | 0.073 |
| 5730409E04F | 0.111 | 0.073 | 0.093 |
| Pgghg | 0.111 | 0.120 | 0.166 |
| B230354K17F | 0.111 | 0.091 | 0.158 |
| 2310061I04R | 0.111 | 0.079 | 0.080 |
| Fxn | 0.111 | 0.140 | 0.150 |
| Tcim | 0.111 | 0.109 | 0.087 |
| Rusc2 | 0.110 | 0.113 | 0.079 |
| Rrp1b | 0.110 | 0.108 | 0.112 |

| | | | |
|---|---|---|---|
| Edem3 | 0.110 | 0.115 | 0.078 |
| Dnase1l1 | 0.110 | 0.057 | 0.073 |
| Osbpl1a | 0.110 | 0.234 | 0.353 |
| Rcl1 | 0.110 | 0.087 | 0.093 |
| Wdcp | 0.110 | 0.080 | 0.078 |
| Sart3 | 0.110 | 0.120 | 0.114 |
| Klhl3 | 0.110 | 0.028 | 0.019 |
| Utp4 | 0.110 | 0.114 | 0.095 |
| Pacs2 | 0.110 | 0.105 | 0.184 |
| Acot2 | 0.110 | 0.111 | 0.119 |
| Ptk2b | 0.110 | 0.003 | 0.006 |
| Itga3 | 0.110 | 0.080 | 0.013 |
| Surf6 | 0.110 | 0.066 | 0.056 |
| Ces2g | 0.110 | 0.069 | 0.139 |
| Wrn | 0.110 | 0.116 | 0.104 |
| Hoxa2 | 0.110 | 0.137 | 0.108 |
| Asah2 | 0.110 | 0.034 | 0.037 |
| Zw10 | 0.110 | 0.083 | 0.075 |
| Pde1b | 0.110 | 0.017 | 0.016 |
| Nsg1 | 0.110 | 0.128 | 0.213 |
| Phykpl | 0.109 | 0.100 | 0.118 |
| Oraov1 | 0.109 | 0.064 | 0.074 |
| Spock2 | 0.109 | 0.042 | 0.208 |
| Rnf113a2 | 0.109 | 0.105 | 0.088 |
| Atg7 | 0.109 | 0.128 | 0.117 |
| Mthfsl | 0.109 | 0.137 | 0.113 |
| Elmo2 | 0.109 | 0.132 | 0.153 |
| Ppp1r16b | 0.109 | 0.003 | 0.001 |
| Ift172 | 0.109 | 0.057 | 0.079 |
| Ttc9c | 0.109 | 0.091 | 0.093 |
| Zfp820 | 0.109 | 0.049 | 0.084 |
| Wrnip1 | 0.109 | 0.172 | 0.136 |
| Actr6 | 0.109 | 0.074 | 0.064 |
| Kri1 | 0.109 | 0.078 | 0.067 |
| Tmem107 | 0.109 | 0.107 | 0.256 |
| Fam20b | 0.109 | 0.119 | 0.107 |
| Hist1h2ap | 0.109 | 0.080 | 0.065 |
| Mllt6 | 0.109 | 0.106 | 0.079 |
| Zfp219 | 0.109 | 0.139 | 0.094 |
| Cmc4 | 0.109 | 0.113 | 0.116 |
| Zfp57 | 0.109 | 0.059 | 0.099 |
| Ccdc82 | 0.109 | 0.144 | 0.124 |
| Rab3ip | 0.109 | 0.155 | 0.086 |
| Zgpat | 0.108 | 0.080 | 0.074 |
| Dsel | 0.108 | 0.173 | 0.286 |
| Rhbdd2 | 0.108 | 0.093 | 0.087 |
| Pan2 | 0.108 | 0.072 | 0.082 |
| Tyw1 | 0.108 | 0.109 | 0.105 |

| | | | |
|---|---|---|---|
| Fbxo21 | 0.108 | 0.119 | 0.115 |
| Mtfr1 | 0.108 | 0.105 | 0.067 |
| Trim37 | 0.108 | 0.083 | 0.076 |
| Thsd1 | 0.108 | 0.007 | 0.013 |
| Rbmx2 | 0.108 | 0.113 | 0.091 |
| Fam53b | 0.108 | 0.077 | 0.066 |
| Papd7 | 0.108 | 0.194 | 0.154 |
| Prr7 | 0.108 | 0.090 | 0.101 |
| Atxn7l1 | 0.108 | 0.065 | 0.058 |
| Vopp1 | 0.108 | 0.078 | 0.081 |
| Mios | 0.108 | 0.132 | 0.145 |
| Tirap | 0.108 | 0.063 | 0.075 |
| Pgm1 | 0.108 | 0.071 | 0.083 |
| Gtdc1 | 0.108 | 0.082 | 0.088 |
| Zfp507 | 0.108 | 0.083 | 0.090 |
| Snip1 | 0.108 | 0.087 | 0.086 |
| Plscr2 | 0.108 | 0.009 | 0.019 |
| Pde7a | 0.108 | 0.131 | 0.103 |
| Cep290 | 0.108 | 0.111 | 0.116 |
| Cep19 | 0.108 | 0.086 | 0.108 |
| Dnajb2 | 0.108 | 0.146 | 0.152 |
| Stk38l | 0.108 | 0.310 | 0.110 |
| Slc20a2 | 0.107 | 0.212 | 0.128 |
| Wdr3 | 0.107 | 0.087 | 0.093 |
| Slc7a6 | 0.107 | 0.140 | 0.157 |
| Icmt | 0.107 | 0.082 | 0.070 |
| Klf15 | 0.107 | 0.170 | 0.094 |
| Trim62 | 0.107 | 0.033 | 0.021 |
| Ccdc43 | 0.107 | 0.097 | 0.080 |
| Samd12 | 0.107 | 0.009 | 0.025 |
| Mus81 | 0.107 | 0.116 | 0.093 |
| Arhgap10 | 0.107 | 0.293 | 0.392 |
| Vps13c | 0.107 | 0.070 | 0.073 |
| Madd | 0.107 | 0.031 | 0.020 |
| Txndc16 | 0.107 | 0.109 | 0.101 |
| Hira | 0.107 | 0.104 | 0.089 |
| Brpf1 | 0.107 | 0.138 | 0.109 |
| Pald1 | 0.107 | 0.074 | 0.036 |
| Rnf219 | 0.107 | 0.100 | 0.090 |
| Agbl3 | 0.107 | 0.106 | 0.095 |
| Parp4 | 0.107 | 0.083 | 0.080 |
| Akap2 | 0.107 | 0.211 | 0.113 |
| Sspn | 0.107 | 0.424 | 0.433 |
| Arl14ep | 0.107 | 0.103 | 0.104 |
| Phf13 | 0.107 | 0.162 | 0.147 |
| Strn | 0.107 | 0.101 | 0.100 |
| Rpap1 | 0.107 | 0.074 | 0.071 |
| Zfp503 | 0.107 | 0.333 | 0.372 |

| | | | |
|---|---|---|---|
| Nfkb2 | 0.107 | 0.105 | 0.108 |
| Arhgap12 | 0.107 | 0.118 | 0.149 |
| Ccdc137 | 0.107 | 0.063 | 0.074 |
| Dact3 | 0.107 | 0.478 | 0.450 |
| Mcm3ap | 0.107 | 0.134 | 0.120 |
| Cln3 | 0.107 | 0.082 | 0.071 |
| Spsb1 | 0.107 | 0.307 | 0.262 |
| Rab23 | 0.107 | 0.112 | 0.181 |
| Pfkm | 0.107 | 0.136 | 0.107 |
| Bag4 | 0.106 | 0.079 | 0.092 |
| Prkca | 0.106 | 0.184 | 0.167 |
| Ambra1 | 0.106 | 0.092 | 0.087 |
| Ncoa3 | 0.106 | 0.133 | 0.116 |
| Lrrc49 | 0.106 | 0.075 | 0.098 |
| Lppos | 0.106 | 0.080 | 0.135 |
| Slc2a10 | 0.106 | 0.059 | 0.072 |
| AI413582 | 0.106 | 0.074 | 0.119 |
| Slc30a1 | 0.106 | 0.063 | 0.054 |
| Klf11 | 0.106 | 0.130 | 0.127 |
| Fam210a | 0.106 | 0.122 | 0.121 |
| Zfhx2 | 0.106 | 0.023 | 0.016 |
| Ctc1 | 0.106 | 0.090 | 0.096 |
| Prdm5 | 0.106 | 0.165 | 0.205 |
| Chd1l | 0.106 | 0.077 | 0.064 |
| Acad9 | 0.106 | 0.091 | 0.106 |
| Tspyl2 | 0.106 | 0.156 | 0.140 |
| Ptpn4 | 0.106 | 0.051 | 0.031 |
| Mertk | 0.106 | 0.159 | 0.058 |
| Zfp606 | 0.106 | 0.079 | 0.085 |
| Extl3 | 0.106 | 0.196 | 0.124 |
| Rell1 | 0.106 | 0.086 | 0.089 |
| D230025D16I | 0.106 | 0.119 | 0.115 |
| Cpped1 | 0.106 | 0.137 | 0.117 |
| Snx16 | 0.106 | 0.109 | 0.108 |
| Aco1 | 0.106 | 0.158 | 0.160 |
| Tatdn2 | 0.106 | 0.131 | 0.089 |
| Ocln | 0.106 | 0.000 | 0.000 |
| Setd1a | 0.106 | 0.117 | 0.105 |
| Ccdc97 | 0.106 | 0.093 | 0.089 |
| Gatc | 0.106 | 0.063 | 0.073 |
| Ctif | 0.106 | 0.113 | 0.100 |
| Ccnb2 | 0.106 | 0.043 | 0.047 |
| Scrn2 | 0.106 | 0.070 | 0.063 |
| Tyw5 | 0.106 | 0.114 | 0.087 |
| Asb7 | 0.106 | 0.118 | 0.107 |
| Rtp4 | 0.106 | 0.015 | 0.037 |
| Pygo2 | 0.106 | 0.125 | 0.116 |
| Gm9844 | 0.105 | 0.036 | 0.071 |

| | | | |
|---|---|---|---|
| Ppat | 0.105 | 0.070 | 0.056 |
| Iqcb1 | 0.105 | 0.176 | 0.121 |
| Cpsf4 | 0.105 | 0.055 | 0.066 |
| Eaf1 | 0.105 | 0.055 | 0.044 |
| Nhlrc3 | 0.105 | 0.048 | 0.059 |
| Wdr53 | 0.105 | 0.086 | 0.078 |
| Adar | 0.105 | 0.064 | 0.059 |
| Spout1 | 0.105 | 0.097 | 0.125 |
| Nexn | 0.105 | 0.863 | 0.299 |
| Dgkd | 0.105 | 0.177 | 0.114 |
| Cmklr1 | 0.105 | 0.078 | 0.086 |
| B630019A10F | 0.105 | 0.035 | 0.010 |
| Cercam | 0.105 | 0.298 | 0.404 |
| Mbtps2 | 0.105 | 0.100 | 0.091 |
| Maml1 | 0.105 | 0.122 | 0.099 |
| Tmem185b | 0.105 | 0.086 | 0.072 |
| Tmem129 | 0.105 | 0.083 | 0.109 |
| Abcd1 | 0.105 | 0.086 | 0.075 |
| Ammecr1 | 0.105 | 0.153 | 0.159 |
| Dera | 0.105 | 0.087 | 0.106 |
| Fbxo25 | 0.105 | 0.098 | 0.109 |
| Stbd1 | 0.105 | 0.577 | 0.380 |
| Basp1 | 0.105 | 0.408 | 0.683 |
| Mid1 | 0.105 | 0.108 | 0.156 |
| Dusp18 | 0.105 | 0.077 | 0.099 |
| Slc25a40 | 0.105 | 0.062 | 0.045 |
| Trmt10c | 0.105 | 0.096 | 0.085 |
| CT025619.1 | 0.105 | 0.146 | 0.251 |
| Inpp5b | 0.105 | 0.083 | 0.067 |
| Fam26e | 0.105 | 0.098 | 0.081 |
| Gnpnat1 | 0.105 | 0.155 | 0.176 |
| Atoh8 | 0.104 | 0.210 | 0.110 |
| Parp12 | 0.104 | 0.086 | 0.094 |
| Atg4d | 0.104 | 0.111 | 0.085 |
| Crybb3 | 0.104 | 0.055 | 0.047 |
| Cotl1 | 0.104 | 0.076 | 0.288 |
| Gm13056 | 0.104 | 0.015 | 0.024 |
| Katnb1 | 0.104 | 0.100 | 0.088 |
| Wdr60 | 0.104 | 0.155 | 0.164 |
| Rad1 | 0.104 | 0.088 | 0.104 |
| S100b | 0.104 | 0.046 | 0.155 |
| Snx14 | 0.104 | 0.132 | 0.114 |
| Naa30 | 0.104 | 0.129 | 0.112 |
| Nhs | 0.104 | 0.343 | 0.242 |
| Csf2ra | 0.104 | 0.111 | 0.180 |
| Fbxo32 | 0.104 | 0.398 | 0.127 |
| Mindy4 | 0.104 | 0.041 | 0.038 |
| Wdr36 | 0.104 | 0.084 | 0.077 |

| | | | |
|---|---|---|---|
| Gucd1 | 0.104 | 0.117 | 0.137 |
| Pinx1 | 0.104 | 0.094 | 0.090 |
| Dvl2 | 0.104 | 0.147 | 0.175 |
| Dlx5 | 0.104 | 0.071 | 0.053 |
| Bcl2l11 | 0.104 | 0.146 | 0.121 |
| Rin1 | 0.104 | 0.035 | 0.011 |
| Nkiras1 | 0.104 | 0.108 | 0.125 |
| Cbr4 | 0.104 | 0.116 | 0.126 |
| Spata13 | 0.104 | 0.074 | 0.013 |
| Hspa2 | 0.104 | 0.369 | 0.164 |
| Nagpa | 0.104 | 0.081 | 0.085 |
| Ccdc77 | 0.103 | 0.064 | 0.081 |
| B230307C23F | 0.103 | 0.059 | 0.073 |
| Dph5 | 0.103 | 0.098 | 0.082 |
| Tmem206 | 0.103 | 0.024 | 0.025 |
| Akap7 | 0.103 | 0.113 | 0.116 |
| Dzip3 | 0.103 | 0.118 | 0.127 |
| Cyp4f16 | 0.103 | 0.096 | 0.091 |
| Foxn2 | 0.103 | 0.107 | 0.127 |
| Gorasp1 | 0.103 | 0.078 | 0.114 |
| P2ry6 | 0.103 | 0.021 | 0.013 |
| Uqcc1 | 0.103 | 0.092 | 0.096 |
| Chil5 | 0.103 | 0.001 | 0.000 |
| Ahi1 | 0.103 | 0.151 | 0.141 |
| Trib3 | 0.103 | 0.029 | 0.026 |
| Rabl3 | 0.103 | 0.069 | 0.076 |
| Ercc6l2 | 0.103 | 0.099 | 0.129 |
| Gpsm1 | 0.103 | 0.182 | 0.202 |
| Gm20604 | 0.103 | 0.078 | 0.075 |
| Txlng | 0.103 | 0.085 | 0.087 |
| 1500009L16F | 0.103 | 0.355 | 0.308 |
| Pml | 0.103 | 0.077 | 0.099 |
| Aldh3b1 | 0.103 | 0.005 | 0.009 |
| Prmt3 | 0.103 | 0.081 | 0.112 |
| Rnaseh1 | 0.103 | 0.093 | 0.096 |
| Rrp9 | 0.103 | 0.103 | 0.093 |
| Klhl18 | 0.103 | 0.086 | 0.098 |
| Ptk7 | 0.103 | 0.069 | 0.095 |
| Clcn5 | 0.103 | 0.112 | 0.367 |
| Gm17745 | 0.103 | 0.001 | 0.000 |
| Fahd1 | 0.103 | 0.092 | 0.084 |
| Tmem38b | 0.103 | 0.270 | 0.170 |
| Zbtb10 | 0.103 | 0.122 | 0.129 |
| Mcm5 | 0.103 | 0.061 | 0.077 |
| Sestd1 | 0.103 | 0.129 | 0.126 |
| Zdhhc9 | 0.103 | 0.104 | 0.157 |
| Hyou1 | 0.103 | 0.070 | 0.069 |
| Tmem87a | 0.103 | 0.100 | 0.077 |

| | | | |
|---|---|---|---|
| Faap100 | 0.103 | 0.108 | 0.125 |
| Cactin | 0.103 | 0.099 | 0.101 |
| C2cd3 | 0.103 | 0.066 | 0.062 |
| Mtm1 | 0.102 | 0.070 | 0.052 |
| Orc2 | 0.102 | 0.117 | 0.105 |
| Baz1a | 0.102 | 0.070 | 0.047 |
| Mpi | 0.102 | 0.070 | 0.058 |
| Emc9 | 0.102 | 0.079 | 0.065 |
| Plekhb2 | 0.102 | 0.025 | 0.022 |
| St6galnac2 | 0.102 | 0.018 | 0.047 |
| Gon7 | 0.102 | 0.070 | 0.055 |
| Cep104 | 0.102 | 0.098 | 0.085 |
| Mis12 | 0.102 | 0.059 | 0.058 |
| Zfp827 | 0.102 | 0.111 | 0.109 |
| Mrps22 | 0.102 | 0.132 | 0.105 |
| Lrrc45 | 0.102 | 0.111 | 0.107 |
| Tgds | 0.102 | 0.094 | 0.112 |
| C030006K11F | 0.102 | 0.059 | 0.039 |
| Fancl | 0.102 | 0.078 | 0.078 |
| Capn5 | 0.102 | 0.022 | 0.059 |
| Dok2 | 0.102 | 0.006 | 0.006 |
| Pcyox1l | 0.102 | 0.107 | 0.041 |
| Sgms2 | 0.102 | 0.081 | 0.071 |
| Npat | 0.102 | 0.114 | 0.112 |
| Atr | 0.102 | 0.107 | 0.101 |
| Rftn2 | 0.102 | 0.178 | 0.185 |
| Dffb | 0.102 | 0.091 | 0.088 |
| Nudt22 | 0.102 | 0.111 | 0.085 |
| Xpc | 0.102 | 0.124 | 0.085 |
| C1qtnf4 | 0.102 | 0.194 | 0.179 |
| 2610037D02I | 0.102 | 0.138 | 0.139 |
| Igf2 | 0.102 | 0.039 | 0.155 |
| Def8 | 0.102 | 0.088 | 0.100 |
| Ints5.1 | 0.102 | 0.097 | 0.118 |
| Lmbr1l | 0.102 | 0.096 | 0.099 |
| Zfp90 | 0.102 | 0.133 | 0.142 |
| Cygb | 0.102 | 0.240 | 1.215 |
| Ccl27a | 0.102 | 0.225 | 0.167 |
| Sirt6 | 0.102 | 0.077 | 0.078 |
| Isy1 | 0.102 | 0.144 | 0.148 |
| St3gal6 | 0.102 | 0.100 | 0.071 |
| AW554918 | 0.102 | 0.051 | 0.050 |
| Arl10 | 0.102 | 0.046 | 0.029 |
| P2ry2 | 0.102 | 0.004 | 0.019 |
| Fktn | 0.102 | 0.118 | 0.124 |
| Tom1l1 | 0.102 | 0.060 | 0.048 |
| Cep44 | 0.102 | 0.111 | 0.104 |
| Zfyve26 | 0.102 | 0.083 | 0.086 |

| | | | |
|---|---|---|---|
| Gbp5 | 0.101 | 0.010 | 0.016 |
| Tcea3 | 0.101 | 0.502 | 0.416 |
| Traf6 | 0.101 | 0.060 | 0.046 |
| Rin3 | 0.101 | 0.138 | 0.085 |
| Entpd7 | 0.101 | 0.103 | 0.090 |
| Usp37 | 0.101 | 0.105 | 0.112 |
| Ints9 | 0.101 | 0.114 | 0.099 |
| Prdm4 | 0.101 | 0.075 | 0.086 |
| Nudt13 | 0.101 | 0.092 | 0.082 |
| Slc35d1 | 0.101 | 0.132 | 0.079 |
| Hccs | 0.101 | 0.096 | 0.099 |
| Rft1 | 0.101 | 0.114 | 0.106 |
| Cnp | 0.101 | 0.084 | 0.095 |
| Sugp2 | 0.101 | 0.093 | 0.092 |
| 2610044O15I | 0.101 | 0.091 | 0.092 |
| Lrrc57 | 0.101 | 0.053 | 0.049 |
| Rhot2 | 0.101 | 0.069 | 0.059 |
| Nbn | 0.101 | 0.116 | 0.119 |
| Itga2b | 0.101 | 0.043 | 0.083 |
| Ccdc61 | 0.101 | 0.058 | 0.061 |
| Fbxo42 | 0.101 | 0.092 | 0.087 |
| Nprl2 | 0.101 | 0.064 | 0.062 |
| Fzd8 | 0.101 | 0.127 | 0.125 |
| Kctd6 | 0.101 | 0.130 | 0.102 |
| Adamts12 | 0.101 | 0.191 | 0.386 |
| Adamts9 | 0.101 | 0.519 | 0.414 |
| Lmbrd2 | 0.101 | 0.093 | 0.069 |
| Cluh | 0.100 | 0.073 | 0.073 |
| Kdm4c | 0.100 | 0.082 | 0.071 |
| Cox18 | 0.100 | 0.095 | 0.095 |
| Slc52a2 | 0.100 | 0.069 | 0.059 |
| Ccdc22 | 0.100 | 0.103 | 0.089 |
| Trim65 | 0.100 | 0.045 | 0.032 |
| Mthfd2 | 0.100 | 0.202 | 0.146 |
| Trim32 | 0.100 | 0.105 | 0.132 |
| Kdm4b | 0.100 | 0.072 | 0.081 |
| Htra3 | 0.100 | 0.177 | 1.256 |
| Abtb2 | 0.100 | 0.162 | 0.092 |
| Alg9 | 0.100 | 0.066 | 0.077 |
| Grpel2 | 0.100 | 0.115 | 0.104 |
| Klhl12 | 0.100 | 0.101 | 0.098 |
| 3110082I17R | 0.100 | 0.070 | 0.068 |
| Zadh2 | 0.100 | 0.133 | 0.150 |
| Maml3 | 0.100 | 0.242 | 0.165 |
| Acap3 | 0.100 | 0.094 | 0.118 |
| Fam124a | 0.100 | 0.030 | 0.158 |
| Tdrd3 | 0.100 | 0.150 | 0.181 |
| Rasa3 | 0.100 | 0.193 | 0.158 |

| | | | |
|---|---|---|---|
| Fam213b | 0.100 | 0.090 | 0.164 |
| Zfp354c | 0.100 | 0.224 | 0.302 |
| Parp6 | 0.100 | 0.079 | 0.092 |
| Paxip1 | 0.100 | 0.096 | 0.087 |
| Ksr1 | 0.100 | 0.102 | 0.113 |
| Dimt1 | 0.100 | 0.114 | 0.118 |
| Strada | 0.100 | 0.075 | 0.060 |
| Dock11 | 0.100 | 0.111 | 0.140 |
| Abhd18 | 0.100 | 0.088 | 0.079 |
| Dnah2os | 0.100 | 0.003 | 0.011 |
| Adck5 | 0.100 | 0.111 | 0.118 |
| Nup85 | 0.100 | 0.093 | 0.119 |
| Txnrd2 | 0.099 | 0.081 | 0.067 |
| Tpbg | 0.099 | 0.149 | 0.179 |
| Ttc32 | 0.099 | 0.083 | 0.064 |
| Them4 | 0.099 | 0.082 | 0.105 |
| Fkbp1b | 0.099 | 0.089 | 0.105 |
| Rhobtb1 | 0.099 | 0.236 | 0.225 |
| Eri1 | 0.099 | 0.082 | 0.105 |
| Ankzf1 | 0.099 | 0.078 | 0.093 |
| Mapk1ip1 | 0.099 | 0.141 | 0.108 |
| Trim39 | 0.099 | 0.064 | 0.071 |
| H6pd | 0.099 | 0.129 | 0.211 |
| Bend5 | 0.099 | 0.224 | 0.197 |
| Gm26518 | 0.099 | 0.189 | 0.166 |
| Fem1a | 0.099 | 0.080 | 0.082 |
| Tm9sf1 | 0.099 | 0.085 | 0.091 |
| Ercc2 | 0.099 | 0.051 | 0.045 |
| Ogn | 0.099 | 0.724 | 1.485 |
| Gmnn | 0.099 | 0.096 | 0.085 |
| Btbd6 | 0.099 | 0.126 | 0.075 |
| Slc8b1 | 0.099 | 0.026 | 0.065 |
| Lrch2 | 0.099 | 0.121 | 0.122 |
| Mcc | 0.099 | 0.069 | 0.144 |
| Mavs | 0.099 | 0.069 | 0.064 |
| Ptcd3 | 0.099 | 0.101 | 0.088 |
| Gm14305 | 0.099 | 0.095 | 0.095 |
| Ccdc60 | 0.099 | 0.099 | 0.021 |
| Epb41l4a | 0.099 | 0.006 | 0.009 |
| Trpm4 | 0.099 | 0.160 | 0.130 |
| Stat5a | 0.098 | 0.088 | 0.064 |
| Cpt1c | 0.098 | 0.140 | 0.200 |
| Tcf20 | 0.098 | 0.179 | 0.110 |
| Cdk10 | 0.098 | 0.117 | 0.142 |
| Entpd5 | 0.098 | 0.057 | 0.109 |
| Gorab | 0.098 | 0.108 | 0.118 |
| Mtfmt | 0.098 | 0.076 | 0.068 |
| Tmem41a | 0.098 | 0.090 | 0.106 |

| | | | |
|---|---|---|---|
| Tril | 0.098 | 0.054 | 0.159 |
| Arhgef17 | 0.098 | 0.393 | 0.184 |
| Lsm11 | 0.098 | 0.125 | 0.068 |
| Fadd | 0.098 | 0.048 | 0.056 |
| Heatr3 | 0.098 | 0.120 | 0.138 |
| F10 | 0.098 | 0.001 | 0.000 |
| Snx19 | 0.098 | 0.069 | 0.073 |
| Krt10 | 0.098 | 0.087 | 0.130 |
| Hsdl2 | 0.098 | 0.126 | 0.099 |
| 2310057M21 | 0.098 | 0.099 | 0.089 |
| Fbxo7 | 0.098 | 0.075 | 0.077 |
| Rmdn2 | 0.098 | 0.078 | 0.099 |
| Ccdc167 | 0.098 | 0.063 | 0.077 |
| Malt1 | 0.098 | 0.109 | 0.066 |
| Dcaf15 | 0.098 | 0.084 | 0.082 |
| Cyth1 | 0.098 | 0.088 | 0.075 |
| Plekhn1 | 0.098 | 0.102 | 0.088 |
| Srp54a | 0.098 | 0.128 | 0.132 |
| Noc4l | 0.098 | 0.093 | 0.102 |
| 0610010F05F | 0.098 | 0.117 | 0.088 |
| Synj2 | 0.098 | 0.130 | 0.083 |
| Rnf19b | 0.098 | 0.230 | 0.120 |
| Srbd1 | 0.098 | 0.061 | 0.076 |
| Scyl3 | 0.098 | 0.069 | 0.060 |
| Zfp101 | 0.098 | 0.103 | 0.086 |
| Rars2 | 0.098 | 0.091 | 0.093 |
| Spata2 | 0.097 | 0.094 | 0.073 |
| Arhgef18 | 0.097 | 0.078 | 0.078 |
| Slc39a11 | 0.097 | 0.029 | 0.067 |
| Prune2 | 0.097 | 0.030 | 0.006 |
| Foxj2 | 0.097 | 0.159 | 0.103 |
| Mcm4 | 0.097 | 0.110 | 0.157 |
| Eid2b | 0.097 | 0.070 | 0.072 |
| Phospho2 | 0.097 | 0.083 | 0.081 |
| Ndufa4l2 | 0.097 | 0.194 | 0.414 |
| Gm26526 | 0.097 | 0.080 | 0.061 |
| Panx1 | 0.097 | 0.081 | 0.165 |
| Ttll1 | 0.097 | 0.119 | 0.102 |
| Mtmr10 | 0.097 | 0.068 | 0.070 |
| Ipo13 | 0.097 | 0.079 | 0.073 |
| Cul9 | 0.097 | 0.072 | 0.082 |
| Mdn1 | 0.097 | 0.079 | 0.072 |
| Taf8 | 0.097 | 0.077 | 0.066 |
| Pphln1 | 0.097 | 0.076 | 0.070 |
| Zfp236 | 0.097 | 0.067 | 0.074 |
| Nt5dc1 | 0.097 | 0.078 | 0.054 |
| Ripk2 | 0.097 | 0.098 | 0.053 |
| Def6 | 0.097 | 0.014 | 0.029 |

| | | | |
|---|---|---|---|
| Srgap1 | 0.097 | 0.020 | 0.124 |
| Sgcd | 0.097 | 0.221 | 0.136 |
| Phf8 | 0.097 | 0.114 | 0.106 |
| Ifitm10 | 0.096 | 0.119 | 0.070 |
| Nrde2 | 0.096 | 0.086 | 0.074 |
| Itih5 | 0.096 | 0.079 | 0.878 |
| Zc3h3 | 0.096 | 0.057 | 0.063 |
| Gfm1 | 0.096 | 0.111 | 0.084 |
| Uty | 0.096 | 0.085 | 0.089 |
| Elp6 | 0.096 | 0.054 | 0.066 |
| Gne | 0.096 | 0.060 | 0.077 |
| Celsr1 | 0.096 | 0.010 | 0.003 |
| Phyh | 0.096 | 0.186 | 0.174 |
| Gm20696 | 0.096 | 0.091 | 0.068 |
| Nphp1 | 0.096 | 0.108 | 0.098 |
| Ric8b | 0.096 | 0.090 | 0.076 |
| Gpr137b | 0.096 | 0.090 | 0.091 |
| Med23 | 0.096 | 0.087 | 0.083 |
| Haus7 | 0.096 | 0.110 | 0.080 |
| Zdhhc1 | 0.096 | 0.091 | 0.146 |
| Mgll | 0.096 | 0.382 | 0.292 |
| Lrba | 0.096 | 0.133 | 0.084 |
| Parp14 | 0.096 | 0.020 | 0.035 |
| Tango2 | 0.096 | 0.120 | 0.171 |
| Gosr1 | 0.096 | 0.089 | 0.075 |
| Qsox2 | 0.096 | 0.078 | 0.063 |
| Acox3 | 0.096 | 0.090 | 0.099 |
| Slc22a15 | 0.096 | 0.062 | 0.114 |
| Lcor | 0.096 | 0.094 | 0.089 |
| Otud1 | 0.096 | 0.180 | 0.122 |
| Clip2 | 0.096 | 0.083 | 0.104 |
| Pdk3 | 0.096 | 0.135 | 0.134 |
| Tstd2 | 0.096 | 0.080 | 0.092 |
| Zbtb45 | 0.096 | 0.057 | 0.072 |
| R3hcc1l | 0.096 | 0.063 | 0.081 |
| Gphn | 0.096 | 0.132 | 0.114 |
| Nol6 | 0.096 | 0.058 | 0.071 |
| D930016D06I | 0.095 | 0.096 | 0.077 |
| Ccdc157 | 0.095 | 0.068 | 0.086 |
| Ftsj1 | 0.095 | 0.100 | 0.109 |
| Isoc2b | 0.095 | 0.054 | 0.077 |
| Mcf2l | 0.095 | 0.011 | 0.045 |
| Kif16b | 0.095 | 0.197 | 0.093 |
| Rcc1l | 0.095 | 0.088 | 0.098 |
| Slc30a4 | 0.095 | 0.100 | 0.090 |
| Khnyn | 0.095 | 0.100 | 0.107 |
| Rnf25 | 0.095 | 0.097 | 0.106 |
| Fmo2 | 0.095 | 0.161 | 0.755 |

| | | | |
|---|---|---|---|
| Clec3b | 0.095 | 0.075 | 1.144 |
| Zfp462 | 0.095 | 0.058 | 0.137 |
| AW209491 | 0.095 | 0.063 | 0.053 |
| Ccnc | 0.095 | 0.124 | 0.104 |
| Usp38 | 0.095 | 0.092 | 0.082 |
| Hinfp | 0.095 | 0.070 | 0.085 |
| Hace1 | 0.095 | 0.095 | 0.081 |
| Trmt10b | 0.095 | 0.079 | 0.088 |
| Bag2 | 0.095 | 0.172 | 0.105 |
| Socs7 | 0.095 | 0.118 | 0.098 |
| Wrap73 | 0.095 | 0.094 | 0.112 |
| Nrm | 0.095 | 0.080 | 0.099 |
| Phf6 | 0.095 | 0.068 | 0.067 |
| Gli3 | 0.095 | 0.121 | 0.197 |
| Scly | 0.095 | 0.046 | 0.054 |
| Atg14 | 0.095 | 0.090 | 0.080 |
| Map1s | 0.095 | 0.126 | 0.150 |
| Hoxb3os | 0.094 | 0.119 | 0.058 |
| Brf2 | 0.094 | 0.078 | 0.063 |
| Parm1 | 0.094 | 0.807 | 0.263 |
| Isoc1 | 0.094 | 0.177 | 0.298 |
| Sclt1 | 0.094 | 0.101 | 0.158 |
| Unc45b | 0.094 | 0.004 | 0.008 |
| Sp4 | 0.094 | 0.091 | 0.056 |
| Cad | 0.094 | 0.072 | 0.126 |
| Cog1 | 0.094 | 0.092 | 0.097 |
| Rab3gap2 | 0.094 | 0.120 | 0.105 |
| Mocs1 | 0.094 | 0.049 | 0.053 |
| Cep192 | 0.094 | 0.100 | 0.113 |
| Zfp160 | 0.094 | 0.075 | 0.080 |
| Tmem216 | 0.094 | 0.086 | 0.092 |
| 2610301B20R | 0.094 | 0.084 | 0.102 |
| AC165953.2 | 0.094 | 0.130 | 0.123 |
| Acaca | 0.094 | 0.134 | 0.148 |
| Ppm1k | 0.094 | 0.067 | 0.059 |
| Ercc4 | 0.094 | 0.082 | 0.073 |
| Mapk8ip1 | 0.094 | 0.129 | 0.220 |
| Ctdp1 | 0.094 | 0.079 | 0.059 |
| Senp3 | 0.094 | 0.107 | 0.080 |
| Fam109a | 0.094 | 0.097 | 0.100 |
| Sphk2 | 0.094 | 0.100 | 0.083 |
| 2810006K23R | 0.094 | 0.050 | 0.077 |
| Neurl4 | 0.094 | 0.072 | 0.061 |
| Flcn | 0.094 | 0.099 | 0.109 |
| 4632415L05R | 0.094 | 0.074 | 0.081 |
| P2ry1 | 0.094 | 0.002 | 0.006 |
| 5430416N02R | 0.094 | 0.117 | 0.115 |
| Klhl26 | 0.093 | 0.120 | 0.093 |

| | | | |
|---|---|---|---|
| Tmem175 | 0.093 | 0.096 | 0.125 |
| Acvr2b | 0.093 | 0.137 | 0.077 |
| Mxd1 | 0.093 | 0.143 | 0.117 |
| Tmem68 | 0.093 | 0.089 | 0.078 |
| Stk4 | 0.093 | 0.119 | 0.099 |
| Phlpp1 | 0.093 | 0.084 | 0.084 |
| Patl1 | 0.093 | 0.083 | 0.073 |
| Irgq | 0.093 | 0.105 | 0.098 |
| Ptpre | 0.093 | 0.007 | 0.004 |
| Ushbp1 | 0.093 | 0.008 | 0.008 |
| Polk | 0.093 | 0.056 | 0.131 |
| AI467606 | 0.093 | 0.001 | 0.001 |
| Snrnp25 | 0.093 | 0.070 | 0.081 |
| AC161165.6 | 0.093 | 0.040 | 0.044 |
| Ggact | 0.093 | 0.050 | 0.016 |
| Dock7 | 0.093 | 0.311 | 0.363 |
| Adcy3 | 0.093 | 0.108 | 0.125 |
| Sh3tc1 | 0.093 | 0.002 | 0.004 |
| Actr5 | 0.093 | 0.060 | 0.059 |
| Coq4 | 0.093 | 0.068 | 0.069 |
| Zfp513 | 0.093 | 0.088 | 0.076 |
| Zfp568 | 0.093 | 0.097 | 0.080 |
| Trp53i11 | 0.093 | 0.046 | 0.024 |
| Msantd3 | 0.093 | 0.135 | 0.128 |
| Ppp2r1b | 0.093 | 0.089 | 0.068 |
| Syt15 | 0.093 | 0.046 | 0.011 |
| Zfyve19 | 0.093 | 0.089 | 0.095 |
| Cbfa2t2 | 0.093 | 0.137 | 0.136 |
| Tns3 | 0.093 | 0.233 | 0.310 |
| Zbtb7c | 0.093 | 0.090 | 0.035 |
| Fuz | 0.093 | 0.124 | 0.123 |
| Tgfbi | 0.093 | 0.614 | 0.506 |
| Zmynd19 | 0.093 | 0.080 | 0.113 |
| Rgp1 | 0.093 | 0.070 | 0.079 |
| B4galt2 | 0.093 | 0.170 | 0.211 |
| Rcan2 | 0.093 | 0.818 | 0.295 |
| Setdb1 | 0.093 | 0.116 | 0.111 |
| Mknk1 | 0.093 | 0.072 | 0.086 |
| Gpsm3 | 0.093 | 0.013 | 0.069 |
| Ror2 | 0.093 | 0.150 | 0.161 |
| Zbtb14 | 0.093 | 0.062 | 0.071 |
| Ankrd24 | 0.093 | 0.097 | 0.102 |
| Ephb3 | 0.093 | 0.142 | 0.252 |
| Bpnt1 | 0.093 | 0.092 | 0.047 |
| Slc1a5 | 0.092 | 0.194 | 0.365 |
| Mtif3 | 0.092 | 0.075 | 0.066 |
| Slc37a4 | 0.092 | 0.055 | 0.064 |
| Chaf1b | 0.092 | 0.051 | 0.047 |

| | | | |
|---|---|---|---|
| Maoa | 0.092 | 0.157 | 0.120 |
| Vps45 | 0.092 | 0.090 | 0.085 |
| Slc30a6 | 0.092 | 0.078 | 0.085 |
| Acat2 | 0.092 | 0.106 | 0.117 |
| Met | 0.092 | 0.024 | 0.058 |
| Samd10 | 0.092 | 0.011 | 0.007 |
| Serinc5 | 0.092 | 0.070 | 0.085 |
| Kifc3 | 0.092 | 0.056 | 0.068 |
| Mtmr9 | 0.092 | 0.063 | 0.072 |
| N4bp3 | 0.092 | 0.022 | 0.014 |
| Gramd1b | 0.092 | 0.015 | 0.028 |
| Prune1 | 0.092 | 0.088 | 0.081 |
| Nif3l1 | 0.092 | 0.078 | 0.093 |
| Fntb | 0.092 | 0.067 | 0.078 |
| Gm15867 | 0.092 | 0.162 | 0.091 |
| Gramd4 | 0.092 | 0.026 | 0.030 |
| Pi15 | 0.092 | 0.642 | 0.634 |
| Tbc1d4 | 0.092 | 0.079 | 0.076 |
| Ppp4r4 | 0.092 | 0.004 | 0.009 |
| Gdap2 | 0.092 | 0.087 | 0.111 |
| Nup188 | 0.092 | 0.087 | 0.098 |
| Kbtbd3 | 0.092 | 0.068 | 0.064 |
| Kif3b | 0.092 | 0.072 | 0.072 |
| Cox4i2 | 0.092 | 0.586 | 0.272 |
| 1190007I07R | 0.092 | 0.076 | 0.067 |
| Arhgap32 | 0.092 | 0.098 | 0.159 |
| Cc2d1b | 0.092 | 0.095 | 0.100 |
| Umps | 0.091 | 0.142 | 0.124 |
| Bicral | 0.091 | 0.109 | 0.094 |
| Layn | 0.091 | 0.081 | 0.064 |
| Mrpl47 | 0.091 | 0.066 | 0.070 |
| Eml2 | 0.091 | 0.141 | 0.051 |
| Zfp687 | 0.091 | 0.124 | 0.118 |
| Chrnb1 | 0.091 | 0.027 | 0.022 |
| Sp2 | 0.091 | 0.103 | 0.077 |
| Fam60a | 0.091 | 0.047 | 0.060 |
| Ccdc84 | 0.091 | 0.076 | 0.059 |
| Dhrs7b | 0.091 | 0.131 | 0.129 |
| Tex30 | 0.091 | 0.100 | 0.072 |
| Fat4 | 0.091 | 0.249 | 0.171 |
| Traf5 | 0.091 | 0.255 | 0.172 |
| Zfp385a | 0.091 | 0.103 | 0.334 |
| Abhd6 | 0.091 | 0.053 | 0.046 |
| Zfp846 | 0.091 | 0.084 | 0.084 |
| Slc33a1 | 0.091 | 0.089 | 0.081 |
| Mrps27 | 0.091 | 0.059 | 0.066 |
| Mrgpre | 0.091 | 0.007 | 0.020 |
| Crtc1 | 0.091 | 0.077 | 0.068 |

| | | | |
|---|---|---|---|
| Colec12 | 0.091 | 0.416 | 0.493 |
| Ccdc112 | 0.091 | 0.162 | 0.124 |
| Elp4 | 0.091 | 0.108 | 0.101 |
| Fzd9 | 0.091 | 0.034 | 0.007 |
| Zhx3 | 0.091 | 0.124 | 0.122 |
| Paqr8 | 0.091 | 0.034 | 0.024 |
| Smarca1 | 0.091 | 0.277 | 0.326 |
| Gm26735 | 0.091 | 0.138 | 0.077 |
| Cstf1 | 0.091 | 0.075 | 0.073 |
| Slc25a37 | 0.091 | 0.070 | 0.054 |
| Reep4 | 0.091 | 0.077 | 0.104 |
| Bicc1 | 0.091 | 0.326 | 1.398 |
| Prrx1 | 0.091 | 0.650 | 0.998 |
| Rnf5 | 0.091 | 0.105 | 0.084 |
| Tmem14a | 0.091 | 0.087 | 0.103 |
| Hirip3 | 0.091 | 0.096 | 0.103 |
| Serpini1 | 0.091 | 0.239 | 0.038 |
| Galnt4 | 0.091 | 0.065 | 0.083 |
| Rpap2 | 0.090 | 0.103 | 0.104 |
| Zbtb5 | 0.090 | 0.065 | 0.074 |
| Zxdc | 0.090 | 0.105 | 0.100 |
| Slc9a6 | 0.090 | 0.137 | 0.111 |
| Vps39 | 0.090 | 0.068 | 0.069 |
| 1600002K03F | 0.090 | 0.077 | 0.070 |
| Ccdc91 | 0.090 | 0.071 | 0.107 |
| Rmi1 | 0.090 | 0.089 | 0.096 |
| Chchd6 | 0.090 | 0.090 | 0.134 |
| Uba6 | 0.090 | 0.109 | 0.131 |
| Zbtb41 | 0.090 | 0.099 | 0.091 |
| Pcnx4 | 0.090 | 0.101 | 0.084 |
| 1600020E01F | 0.090 | 0.098 | 0.108 |
| Pgf | 0.090 | 0.043 | 0.111 |
| Tnxb | 0.090 | 0.261 | 1.224 |
| Ap4b1 | 0.090 | 0.065 | 0.080 |
| Oma1 | 0.090 | 0.061 | 0.048 |
| Plekhg1 | 0.090 | 0.011 | 0.079 |
| Ap3m2 | 0.090 | 0.013 | 0.036 |
| Rbm48 | 0.090 | 0.090 | 0.070 |
| Usp45 | 0.090 | 0.096 | 0.077 |
| Slf1 | 0.090 | 0.114 | 0.108 |
| Vps8 | 0.090 | 0.098 | 0.076 |
| Tmub1 | 0.090 | 0.078 | 0.062 |
| Brpf3 | 0.090 | 0.058 | 0.060 |
| Siah2 | 0.090 | 0.118 | 0.070 |
| Mettl21a | 0.090 | 0.095 | 0.137 |
| Zfp938 | 0.090 | 0.082 | 0.078 |
| Sdccag8 | 0.090 | 0.074 | 0.121 |
| Egr3 | 0.090 | 0.393 | 0.356 |

| | | | |
|---|---|---|---|
| Gm10762 | 0.090 | 0.055 | 0.047 |
| Svip | 0.090 | 0.040 | 0.023 |
| Sfxn2 | 0.090 | 0.105 | 0.132 |
| Cog3 | 0.089 | 0.070 | 0.066 |
| Gm12216 | 0.089 | 0.326 | 0.342 |
| Akap1 | 0.089 | 0.093 | 0.064 |
| Rnf122 | 0.089 | 0.281 | 0.345 |
| Gm8797 | 0.089 | 0.089 | 0.101 |
| Accs | 0.089 | 0.068 | 0.062 |
| Pafah2 | 0.089 | 0.050 | 0.059 |
| Tmem173 | 0.089 | 0.014 | 0.066 |
| Sall2 | 0.089 | 0.071 | 0.067 |
| Arfgef3 | 0.089 | 0.003 | 0.006 |
| Heyl | 0.089 | 0.338 | 0.238 |
| Styx | 0.089 | 0.101 | 0.095 |
| Slc6a6 | 0.089 | 0.405 | 0.162 |
| Nectin1 | 0.089 | 0.037 | 0.061 |
| Cptp | 0.089 | 0.041 | 0.043 |
| Haus1 | 0.089 | 0.089 | 0.110 |
| Zfp617 | 0.089 | 0.066 | 0.075 |
| Zfp868 | 0.089 | 0.068 | 0.064 |
| Rida | 0.089 | 0.083 | 0.066 |
| Pitpnm2 | 0.089 | 0.261 | 0.094 |
| Foxk1 | 0.089 | 0.092 | 0.062 |
| Smtnl1 | 0.089 | 0.000 | 0.001 |
| Chst15 | 0.089 | 0.057 | 0.032 |
| Kcnt2 | 0.089 | 0.004 | 0.044 |
| Sirt5 | 0.089 | 0.086 | 0.092 |
| Phka2 | 0.089 | 0.129 | 0.113 |
| Ccl25 | 0.088 | 0.073 | 0.095 |
| Tap2 | 0.088 | 0.057 | 0.060 |
| Podxl2 | 0.088 | 0.316 | 0.439 |
| Zfp931 | 0.088 | 0.075 | 0.069 |
| Asb6 | 0.088 | 0.075 | 0.054 |
| Wdr41 | 0.088 | 0.079 | 0.088 |
| Mccc1 | 0.088 | 0.106 | 0.094 |
| Mecr | 0.088 | 0.086 | 0.075 |
| Col8a1 | 0.088 | 0.054 | 0.098 |
| Mov10 | 0.088 | 0.048 | 0.094 |
| Gstt3 | 0.088 | 0.057 | 0.107 |
| Mad1l1 | 0.088 | 0.059 | 0.069 |
| Spryd4 | 0.088 | 0.056 | 0.064 |
| H2-M3 | 0.088 | 0.016 | 0.046 |
| Smyd5 | 0.088 | 0.046 | 0.068 |
| Fastkd2 | 0.088 | 0.065 | 0.060 |
| Akip1 | 0.088 | 0.081 | 0.105 |
| C2cd2l | 0.088 | 0.061 | 0.040 |
| Optn | 0.088 | 0.093 | 0.081 |

| | | | |
|---|---|---|---|
| Mkl1 | 0.088 | 0.109 | 0.135 |
| Kcnj2 | 0.088 | 0.001 | 0.001 |
| Gm20342 | 0.088 | 0.144 | 0.125 |
| Cyp51 | 0.088 | 0.229 | 0.237 |
| Slc35f6 | 0.088 | 0.090 | 0.082 |
| Snx33 | 0.088 | 0.073 | 0.090 |
| Ddr2 | 0.088 | 0.517 | 0.629 |
| Ganc | 0.088 | 0.085 | 0.075 |
| C130074G19I | 0.088 | 0.002 | 0.003 |
| Rnf31 | 0.088 | 0.092 | 0.111 |
| Nceh1 | 0.088 | 0.022 | 0.020 |
| Trmu | 0.088 | 0.090 | 0.107 |
| Mast2 | 0.088 | 0.129 | 0.132 |
| Hexdc | 0.087 | 0.073 | 0.059 |
| Pip4k2b | 0.087 | 0.075 | 0.112 |
| Rprd1a | 0.087 | 0.131 | 0.121 |
| Zfp949 | 0.087 | 0.079 | 0.058 |
| Elf4 | 0.087 | 0.068 | 0.152 |
| Nmnat2 | 0.087 | 0.021 | 0.008 |
| Lypd6 | 0.087 | 0.002 | 0.001 |
| Siae | 0.087 | 0.087 | 0.107 |
| Apex2 | 0.087 | 0.073 | 0.067 |
| Zswim7 | 0.087 | 0.152 | 0.138 |
| Traf4 | 0.087 | 0.129 | 0.216 |
| Zfp942 | 0.087 | 0.083 | 0.098 |
| Zmym3 | 0.087 | 0.124 | 0.102 |
| Baalc | 0.087 | 0.035 | 0.070 |
| Esrra | 0.087 | 0.066 | 0.069 |
| Fbxo4 | 0.087 | 0.127 | 0.144 |
| Slc27a3 | 0.087 | 0.058 | 0.128 |
| Zfp788 | 0.087 | 0.073 | 0.068 |
| Spire1 | 0.087 | 0.147 | 0.091 |
| Rfesd | 0.087 | 0.079 | 0.086 |
| Stk10 | 0.087 | 0.023 | 0.012 |
| Mr1 | 0.087 | 0.074 | 0.120 |
| Dusp19 | 0.087 | 0.075 | 0.085 |
| Zfp335 | 0.087 | 0.061 | 0.052 |
| Bcas3 | 0.087 | 0.114 | 0.073 |
| Cyp2j6 | 0.087 | 0.072 | 0.129 |
| Rassf10 | 0.087 | 0.009 | 0.003 |
| Slc27a4 | 0.087 | 0.087 | 0.097 |
| Slc8a1 | 0.087 | 0.050 | 0.026 |
| Arhgef25 | 0.086 | 0.570 | 0.571 |
| Brwd3 | 0.086 | 0.097 | 0.089 |
| Aaas | 0.086 | 0.040 | 0.054 |
| Cacnb3 | 0.086 | 0.287 | 0.326 |
| Aacs | 0.086 | 0.064 | 0.079 |
| Slx1b | 0.086 | 0.074 | 0.073 |

| | | | |
|---|---|---|---|
| Cables2 | 0.086 | 0.093 | 0.097 |
| Trmt10a | 0.086 | 0.050 | 0.076 |
| Rmnd1 | 0.086 | 0.093 | 0.112 |
| Ap5s1 | 0.086 | 0.063 | 0.085 |
| Fxyd6 | 0.086 | 0.157 | 0.571 |
| Hmga2 | 0.086 | 0.007 | 0.012 |
| Rfxank | 0.086 | 0.061 | 0.071 |
| Nup93 | 0.086 | 0.112 | 0.088 |
| Arntl | 0.086 | 0.089 | 0.079 |
| Wdr4 | 0.086 | 0.063 | 0.078 |
| Inip | 0.086 | 0.046 | 0.049 |
| Dhfr | 0.086 | 0.060 | 0.082 |
| Pik3r2 | 0.086 | 0.203 | 0.301 |
| Pla2g15 | 0.086 | 0.078 | 0.096 |
| Kiz | 0.086 | 0.114 | 0.139 |
| Znfx1 | 0.086 | 0.086 | 0.068 |
| D1Ertd622e | 0.086 | 0.157 | 0.057 |
| Zfp654 | 0.086 | 0.085 | 0.073 |
| Vrk1 | 0.086 | 0.074 | 0.072 |
| 1700037H04I | 0.086 | 0.062 | 0.077 |
| Hdac8 | 0.086 | 0.081 | 0.074 |
| Rfx1 | 0.086 | 0.062 | 0.060 |
| Cpeb2 | 0.086 | 0.047 | 0.045 |
| Dcp1b | 0.086 | 0.075 | 0.056 |
| Lrwd1 | 0.086 | 0.079 | 0.090 |
| Gm16740 | 0.086 | 0.058 | 0.055 |
| 4931414P19I | 0.086 | 0.020 | 0.021 |
| Tc2n | 0.086 | 0.030 | 0.014 |
| Zfand2a | 0.086 | 0.063 | 0.052 |
| Ell | 0.085 | 0.105 | 0.089 |
| Dis3 | 0.085 | 0.106 | 0.113 |
| Ikzf2 | 0.085 | 0.080 | 0.091 |
| Irgm1 | 0.085 | 0.063 | 0.052 |
| Kctd18 | 0.085 | 0.085 | 0.089 |
| Lyplal1 | 0.085 | 0.053 | 0.065 |
| Ctsk | 0.085 | 0.227 | 0.641 |
| Pclaf | 0.085 | 0.053 | 0.074 |
| Pprc1 | 0.085 | 0.095 | 0.130 |
| Narfl | 0.085 | 0.077 | 0.072 |
| Cpsf1 | 0.085 | 0.074 | 0.078 |
| Ttc39b | 0.085 | 0.132 | 0.054 |
| Galnt15 | 0.085 | 0.128 | 0.134 |
| Utp6 | 0.085 | 0.095 | 0.107 |
| Nsun4 | 0.085 | 0.069 | 0.070 |
| E130308A19I | 0.085 | 0.069 | 0.066 |
| Mpp7 | 0.085 | 0.074 | 0.030 |
| Ncapd3 | 0.085 | 0.097 | 0.085 |
| Zfp142 | 0.085 | 0.094 | 0.105 |

| | | | |
|---|---|---|---|
| Pigu | 0.085 | 0.086 | 0.081 |
| Spon2 | 0.085 | 0.182 | 0.334 |
| Bves | 0.085 | 0.148 | 0.018 |
| Zfp948 | 0.085 | 0.064 | 0.091 |
| Kbtbd4 | 0.085 | 0.061 | 0.057 |
| Arhgap18 | 0.085 | 0.022 | 0.019 |
| Tank | 0.085 | 0.150 | 0.109 |
| Isg20l2 | 0.085 | 0.081 | 0.085 |
| Tmem161b | 0.085 | 0.063 | 0.051 |
| Acsl4 | 0.085 | 0.065 | 0.047 |
| Nudt7 | 0.085 | 0.040 | 0.061 |
| Rbm15 | 0.085 | 0.093 | 0.079 |
| Rnf135 | 0.084 | 0.037 | 0.049 |
| Kctd9 | 0.084 | 0.049 | 0.085 |
| Myo1b | 0.084 | 0.133 | 0.087 |
| Agpat2 | 0.084 | 0.237 | 0.210 |
| Helz2 | 0.084 | 0.033 | 0.038 |
| Cacna2d1 | 0.084 | 0.434 | 0.315 |
| Adamtsl4 | 0.084 | 0.191 | 0.264 |
| Dcun1d2 | 0.084 | 0.048 | 0.057 |
| Gm16536 | 0.084 | 0.058 | 0.055 |
| Prpf3 | 0.084 | 0.075 | 0.073 |
| Armt1 | 0.084 | 0.064 | 0.077 |
| Rpia | 0.084 | 0.089 | 0.102 |
| Mief1 | 0.084 | 0.082 | 0.076 |
| Mettl3 | 0.084 | 0.076 | 0.079 |
| Zbtb24 | 0.084 | 0.086 | 0.076 |
| Cxcl15 | 0.084 | 0.128 | 0.053 |
| Nme6 | 0.084 | 0.069 | 0.059 |
| 1110051M20 | 0.084 | 0.087 | 0.105 |
| Srr | 0.084 | 0.071 | 0.116 |
| Trp53bp2 | 0.084 | 0.145 | 0.128 |
| Hist1h1e | 0.084 | 0.064 | 0.059 |
| Gcc1 | 0.084 | 0.219 | 0.222 |
| Erbb2 | 0.084 | 0.074 | 0.101 |
| Endov | 0.084 | 0.067 | 0.071 |
| Ptcd1 | 0.084 | 0.064 | 0.069 |
| Tyk2 | 0.084 | 0.059 | 0.056 |
| Mtr | 0.084 | 0.096 | 0.080 |
| Zfyve1 | 0.084 | 0.080 | 0.074 |
| Smad3 | 0.084 | 0.096 | 0.151 |
| Spin4 | 0.084 | 0.060 | 0.070 |
| Hoxaas3 | 0.084 | 0.070 | 0.063 |
| Wdr92 | 0.084 | 0.064 | 0.068 |
| Ccdc149 | 0.084 | 0.036 | 0.038 |
| Nudt16 | 0.084 | 0.076 | 0.092 |
| Chpf | 0.084 | 0.433 | 0.635 |
| Usp54 | 0.084 | 0.086 | 0.055 |

| | | | |
|---|---|---|---|
| Arfgef2 | 0.084 | 0.060 | 0.055 |
| Oasl2 | 0.084 | 0.034 | 0.056 |
| Slit3 | 0.083 | 0.655 | 0.620 |
| Plk3 | 0.083 | 0.052 | 0.075 |
| Col16a1 | 0.083 | 0.319 | 1.021 |
| Disp1 | 0.083 | 0.075 | 0.141 |
| Paqr4 | 0.083 | 0.072 | 0.073 |
| Ubtd2 | 0.083 | 0.109 | 0.236 |
| Slc15a4 | 0.083 | 0.057 | 0.064 |
| Dtx3l | 0.083 | 0.034 | 0.036 |
| Tada2b | 0.083 | 0.100 | 0.088 |
| Nudt1 | 0.083 | 0.062 | 0.082 |
| Tgfbr3 | 0.083 | 0.167 | 0.348 |
| Mcm2 | 0.083 | 0.059 | 0.084 |
| Pex11g | 0.083 | 0.039 | 0.040 |
| Rabgef1 | 0.083 | 0.130 | 0.134 |
| Btaf1 | 0.083 | 0.099 | 0.076 |
| Ankrd27 | 0.083 | 0.122 | 0.172 |
| Alg11 | 0.083 | 0.092 | 0.103 |
| Dclk1 | 0.083 | 0.375 | 0.532 |
| Dubr | 0.083 | 0.121 | 0.168 |
| Ccdc66 | 0.083 | 0.103 | 0.108 |
| Nppc | 0.083 | 0.009 | 0.009 |
| Tbc1d22b | 0.083 | 0.067 | 0.058 |
| Gtpbp3 | 0.083 | 0.068 | 0.072 |
| Manea | 0.083 | 0.105 | 0.161 |
| Aldh16a1 | 0.083 | 0.094 | 0.114 |
| Ccdc141 | 0.083 | 0.269 | 0.096 |
| Tmem120b | 0.083 | 0.027 | 0.057 |
| Dtx2 | 0.083 | 0.104 | 0.118 |
| Lgals3bp | 0.083 | 0.036 | 0.137 |
| Fgfbp3 | 0.083 | 0.053 | 0.058 |
| Tapbpl | 0.083 | 0.045 | 0.040 |
| Dhodh | 0.083 | 0.073 | 0.075 |
| Vrk2 | 0.083 | 0.067 | 0.082 |
| Mto1 | 0.083 | 0.063 | 0.062 |
| Hrct1 | 0.083 | 0.041 | 0.037 |
| Cc2d1a | 0.083 | 0.076 | 0.091 |
| Fig4 | 0.083 | 0.078 | 0.072 |
| Tomm40l | 0.083 | 0.067 | 0.059 |
| Tmem39b | 0.083 | 0.100 | 0.104 |
| Pknox1 | 0.083 | 0.104 | 0.167 |
| Nol4l | 0.082 | 0.031 | 0.034 |
| Zfp74 | 0.082 | 0.090 | 0.095 |
| Zfp810 | 0.082 | 0.067 | 0.053 |
| Ttll12 | 0.082 | 0.056 | 0.089 |
| Pcdhga12 | 0.082 | 0.070 | 0.077 |
| Abhd17c | 0.082 | 0.054 | 0.085 |

| | | | |
|---|---|---|---|
| Exoc8 | 0.082 | 0.037 | 0.039 |
| 1810006J02R | 0.082 | 0.002 | 0.001 |
| Stk35 | 0.082 | 0.076 | 0.072 |
| Smpd4 | 0.082 | 0.070 | 0.065 |
| Prkcz | 0.082 | 0.037 | 0.045 |
| Mtmr1 | 0.082 | 0.093 | 0.083 |
| Rom1 | 0.082 | 0.083 | 0.067 |
| Ofd1 | 0.082 | 0.100 | 0.094 |
| Acsl1 | 0.082 | 0.258 | 0.091 |
| Pigh | 0.082 | 0.057 | 0.063 |
| Supt7l | 0.082 | 0.062 | 0.061 |
| Cep41 | 0.082 | 0.044 | 0.041 |
| 2700097O09I | 0.082 | 0.077 | 0.077 |
| Atrip | 0.082 | 0.083 | 0.077 |
| Trmt11 | 0.082 | 0.080 | 0.079 |
| Cox10 | 0.082 | 0.075 | 0.066 |
| Tbp | 0.082 | 0.062 | 0.064 |
| Sdsl | 0.082 | 0.012 | 0.005 |
| Abhd10 | 0.082 | 0.111 | 0.094 |
| 1110008L16F | 0.082 | 0.043 | 0.040 |
| Elk1 | 0.082 | 0.107 | 0.107 |
| Zfp84 | 0.082 | 0.049 | 0.051 |
| H2afy2 | 0.082 | 0.275 | 0.250 |
| 2810402E24F | 0.082 | 0.055 | 0.058 |
| Tfcp2 | 0.082 | 0.102 | 0.102 |
| Pus10 | 0.082 | 0.068 | 0.060 |
| Zfp738 | 0.082 | 0.068 | 0.074 |
| Gm26870 | 0.082 | 0.069 | 0.060 |
| Slc22a17 | 0.081 | 0.164 | 0.196 |
| Pot1a | 0.081 | 0.061 | 0.076 |
| Sh3d21 | 0.081 | 0.111 | 0.068 |
| Slfn5 | 0.081 | 0.007 | 0.086 |
| Gpm6b | 0.081 | 0.234 | 0.343 |
| Pla2g6 | 0.081 | 0.028 | 0.044 |
| Zbtb43 | 0.081 | 0.087 | 0.080 |
| Dnajc17 | 0.081 | 0.106 | 0.087 |
| Tbx20 | 0.081 | 0.067 | 0.065 |
| Ccdc117 | 0.081 | 0.094 | 0.069 |
| Dcun1d3 | 0.081 | 0.073 | 0.098 |
| Mettl14 | 0.081 | 0.054 | 0.044 |
| Zbtb33 | 0.081 | 0.060 | 0.063 |
| Kdm5d | 0.081 | 0.051 | 0.068 |
| Pgm2l1 | 0.081 | 0.062 | 0.070 |
| Cadm1 | 0.081 | 0.024 | 0.038 |
| Trim16 | 0.081 | 0.074 | 0.088 |
| Ptpn21 | 0.081 | 0.149 | 0.142 |
| Slc27a1 | 0.081 | 0.261 | 0.159 |
| Nxpe4 | 0.081 | 0.013 | 0.023 |

| | | | |
|---|---|---|---|
| Lipt2 | 0.081 | 0.061 | 0.072 |
| Sgsh | 0.081 | 0.031 | 0.038 |
| Gata4 | 0.081 | 0.020 | 0.058 |
| Polr3e | 0.081 | 0.078 | 0.067 |
| AI480526 | 0.081 | 0.042 | 0.034 |
| Champ1 | 0.081 | 0.090 | 0.107 |
| Kcnj8 | 0.081 | 1.098 | 0.511 |
| Fgfr1op | 0.081 | 0.098 | 0.084 |
| Mtus2 | 0.081 | 0.580 | 0.226 |
| Lrig2 | 0.081 | 0.090 | 0.079 |
| Slfn9 | 0.081 | 0.025 | 0.031 |
| Zfp518a | 0.081 | 0.066 | 0.075 |
| Dhx29 | 0.081 | 0.094 | 0.106 |
| Rcan3 | 0.081 | 0.058 | 0.090 |
| Mvd | 0.081 | 0.051 | 0.080 |
| Gstt1 | 0.081 | 0.140 | 0.254 |
| Riox1 | 0.081 | 0.056 | 0.045 |
| Ntn1 | 0.081 | 0.132 | 0.203 |
| Decr2 | 0.081 | 0.065 | 0.070 |
| Snx30 | 0.081 | 0.075 | 0.110 |
| Ddit4l | 0.081 | 0.039 | 0.025 |
| Rmdn3 | 0.081 | 0.086 | 0.085 |
| Pigf | 0.081 | 0.072 | 0.061 |
| Lin54 | 0.081 | 0.075 | 0.061 |
| Deptor | 0.081 | 0.071 | 0.111 |
| Tmem204 | 0.081 | 0.026 | 0.080 |
| Bik | 0.081 | 0.182 | 0.110 |
| Stau2 | 0.081 | 0.100 | 0.108 |
| Zfp384 | 0.081 | 0.063 | 0.052 |
| Hpse | 0.081 | 0.010 | 0.036 |
| Ankrd1 | 0.080 | 0.027 | 0.010 |
| Negr1 | 0.080 | 0.016 | 0.021 |
| C2cd5 | 0.080 | 0.084 | 0.088 |
| Asrgl1 | 0.080 | 0.107 | 0.109 |
| Cyp39a1 | 0.080 | 0.134 | 0.053 |
| Dgka | 0.080 | 0.054 | 0.044 |
| Pym1 | 0.080 | 0.084 | 0.074 |
| Igip | 0.080 | 0.059 | 0.064 |
| Dse | 0.080 | 0.218 | 0.236 |
| Tmem255b | 0.080 | 0.001 | 0.000 |
| Mid2 | 0.080 | 0.088 | 0.074 |
| 3110043O21I | 0.080 | 0.069 | 0.050 |
| Gadd45a | 0.080 | 0.100 | 0.073 |
| Agtr2 | 0.080 | 0.064 | 0.709 |
| Supt3 | 0.080 | 0.079 | 0.097 |
| Ang | 0.080 | 0.044 | 0.147 |
| Zfp251 | 0.080 | 0.122 | 0.113 |
| Fbxl19 | 0.080 | 0.046 | 0.045 |

| | | | |
|---|---|---|---|
| Cops7b | 0.080 | 0.068 | 0.068 |
| Ghdc | 0.080 | 0.086 | 0.071 |
| 1110034G24I | 0.080 | 0.055 | 0.059 |
| Tatdn3 | 0.080 | 0.069 | 0.055 |
| Rfwd3 | 0.080 | 0.059 | 0.060 |
| Zbtb2 | 0.080 | 0.080 | 0.068 |
| Mdfic | 0.080 | 0.097 | 0.156 |
| Abcb8 | 0.080 | 0.064 | 0.082 |
| B4galt5 | 0.080 | 0.079 | 0.084 |
| Slc39a8 | 0.080 | 0.015 | 0.017 |
| Smox | 0.080 | 0.107 | 0.143 |
| Mob3a | 0.080 | 0.092 | 0.105 |
| Adck1 | 0.080 | 0.054 | 0.054 |
| Prex1 | 0.080 | 0.021 | 0.028 |
| Col27a1 | 0.080 | 0.414 | 0.496 |
| Zfp113 | 0.080 | 0.070 | 0.071 |
| Fkbpl | 0.080 | 0.068 | 0.055 |
| Kctd11 | 0.080 | 0.063 | 0.139 |
| Mcoln1 | 0.080 | 0.052 | 0.066 |
| Rnd1 | 0.080 | 0.132 | 0.072 |
| Ttc8 | 0.080 | 0.113 | 0.098 |
| Nkd1 | 0.080 | 0.392 | 0.183 |
| Lipo3 | 0.080 | 0.094 | 0.080 |
| Chst1 | 0.079 | 0.017 | 0.036 |
| Zfp87 | 0.079 | 0.050 | 0.051 |
| Qtrt1 | 0.079 | 0.099 | 0.104 |
| Pdik1l | 0.079 | 0.050 | 0.057 |
| Mthfd1 | 0.079 | 0.094 | 0.057 |
| Ints12 | 0.079 | 0.071 | 0.079 |
| Tnfrsf23 | 0.079 | 0.040 | 0.066 |
| Hoxa10 | 0.079 | 0.000 | 0.000 |
| Sbsn | 0.079 | 0.099 | 0.123 |
| Obscn | 0.079 | 0.002 | 0.002 |
| Invs | 0.079 | 0.070 | 0.077 |
| Chst11 | 0.079 | 0.100 | 0.045 |
| Prr14l | 0.079 | 0.080 | 0.083 |
| Prickle3 | 0.079 | 0.074 | 0.073 |
| Vac14 | 0.079 | 0.064 | 0.053 |
| Gcdh | 0.079 | 0.096 | 0.089 |
| Ell2 | 0.079 | 0.177 | 0.062 |
| Vwc2 | 0.079 | 0.032 | 0.117 |
| Spryd7 | 0.079 | 0.096 | 0.111 |
| A230050P20I | 0.079 | 0.024 | 0.062 |
| Scube2 | 0.079 | 0.002 | 0.015 |
| Nat10 | 0.079 | 0.069 | 0.075 |
| Alkbh2 | 0.079 | 0.041 | 0.050 |
| Natd1 | 0.079 | 0.100 | 0.094 |
| Id4 | 0.079 | 0.509 | 0.223 |

| | | | |
|---|---|---|---|
| Aagab | 0.079 | 0.042 | 0.041 |
| Amot | 0.079 | 0.029 | 0.042 |
| Rbak | 0.079 | 0.096 | 0.076 |
| Zfp777 | 0.079 | 0.094 | 0.096 |
| Cnot9 | 0.079 | 0.075 | 0.074 |
| Coq8b | 0.079 | 0.090 | 0.122 |
| Elovl7 | 0.079 | 0.013 | 0.006 |
| Stk39 | 0.078 | 0.225 | 0.303 |
| Ndufaf1 | 0.078 | 0.098 | 0.101 |
| A630072M18 | 0.078 | 0.069 | 0.068 |
| Myl4 | 0.078 | 0.080 | 0.095 |
| Abcg1 | 0.078 | 0.009 | 0.004 |
| Nov | 0.078 | 0.613 | 0.539 |
| Gtf2h3 | 0.078 | 0.075 | 0.074 |
| Il13ra1 | 0.078 | 0.192 | 0.174 |
| Gcnt2 | 0.078 | 0.172 | 0.041 |
| H2-T23 | 0.078 | 0.048 | 0.067 |
| Dedd2 | 0.078 | 0.057 | 0.033 |
| Ift81 | 0.078 | 0.201 | 0.212 |
| Cenpw | 0.078 | 0.104 | 0.079 |
| Hist2h2be | 0.078 | 0.018 | 0.009 |
| Clcf1 | 0.078 | 0.096 | 0.053 |
| Rngtt | 0.078 | 0.087 | 0.081 |
| Hps5 | 0.078 | 0.058 | 0.067 |
| Naa40 | 0.078 | 0.061 | 0.042 |
| Tceanc2 | 0.078 | 0.093 | 0.096 |
| Ikbkg | 0.078 | 0.060 | 0.063 |
| Col9a1 | 0.078 | 0.057 | 0.047 |
| Tpm3-rs7 | 0.078 | 0.033 | 0.036 |
| Ndufaf6 | 0.078 | 0.062 | 0.065 |
| Adi1 | 0.078 | 0.061 | 0.061 |
| Exoc3l | 0.078 | 0.001 | 0.001 |
| Scara3 | 0.078 | 0.222 | 0.491 |
| Bend7 | 0.077 | 0.024 | 0.015 |
| Zrsr1 | 0.077 | 0.065 | 0.057 |
| Bmf | 0.077 | 0.245 | 0.168 |
| Prr5l | 0.077 | 0.029 | 0.020 |
| Map3k8 | 0.077 | 0.256 | 0.255 |
| Elovl6 | 0.077 | 0.068 | 0.100 |
| Tmcc2 | 0.077 | 0.057 | 0.074 |
| Extl2 | 0.077 | 0.144 | 0.146 |
| Ppif | 0.077 | 0.050 | 0.062 |
| Usp21 | 0.077 | 0.085 | 0.073 |
| Nek1 | 0.077 | 0.076 | 0.103 |
| Smarcal1 | 0.077 | 0.072 | 0.065 |
| Arhgef11 | 0.077 | 0.099 | 0.083 |
| Ikbke | 0.077 | 0.046 | 0.054 |
| Ctns | 0.077 | 0.053 | 0.058 |

| | | | |
|---|---|---|---|
| Fam107b | 0.077 | 0.121 | 0.101 |
| Grrp1 | 0.077 | 0.001 | 0.001 |
| Stradb | 0.077 | 0.055 | 0.048 |
| Hoxa1 | 0.077 | 0.031 | 0.030 |
| Ankrd9 | 0.077 | 0.074 | 0.041 |
| Plag1 | 0.077 | 0.040 | 0.074 |
| Map2k6 | 0.077 | 0.041 | 0.035 |
| Bicra | 0.077 | 0.063 | 0.055 |
| Cldn34c1 | 0.077 | 0.064 | 0.062 |
| Nup155 | 0.077 | 0.066 | 0.068 |
| G430095P16I | 0.077 | 0.022 | 0.017 |
| Ntpcr | 0.077 | 0.140 | 0.176 |
| Cyth4 | 0.077 | 0.004 | 0.005 |
| Crtc2 | 0.077 | 0.098 | 0.069 |
| Zfp944 | 0.077 | 0.069 | 0.081 |
| Dpy19l3 | 0.077 | 0.052 | 0.035 |
| Eif2ak3 | 0.077 | 0.045 | 0.077 |
| Spag1 | 0.076 | 0.014 | 0.014 |
| Tmem170b | 0.076 | 0.056 | 0.048 |
| Cox16 | 0.076 | 0.057 | 0.068 |
| Gucy1a1 | 0.076 | 0.581 | 0.388 |
| Exosc2 | 0.076 | 0.062 | 0.060 |
| Slc25a19 | 0.076 | 0.080 | 0.077 |
| Ints4 | 0.076 | 0.063 | 0.062 |
| Gss | 0.076 | 0.087 | 0.059 |
| Clcn7 | 0.076 | 0.075 | 0.072 |
| Atm | 0.076 | 0.067 | 0.079 |
| Slx4ip | 0.076 | 0.031 | 0.054 |
| Lgr6 | 0.076 | 0.910 | 0.135 |
| Zmynd15 | 0.076 | 0.007 | 0.007 |
| Lemd3 | 0.076 | 0.104 | 0.078 |
| Hacl1 | 0.076 | 0.069 | 0.066 |
| Rnd3 | 0.076 | 0.363 | 0.392 |
| St6galnac5 | 0.076 | 0.002 | 0.007 |
| Il15ra | 0.076 | 0.035 | 0.021 |
| Iqgap2 | 0.076 | 0.056 | 0.089 |
| Ddhd1 | 0.076 | 0.119 | 0.107 |
| Efl1 | 0.076 | 0.100 | 0.093 |
| Magi1 | 0.076 | 0.126 | 0.096 |
| Galnt18 | 0.076 | 0.100 | 0.096 |
| Fam117a | 0.076 | 0.043 | 0.074 |
| Heatr5b | 0.076 | 0.070 | 0.057 |
| Zfp180 | 0.076 | 0.069 | 0.063 |
| Tmem209 | 0.076 | 0.055 | 0.055 |
| Mmgt1 | 0.076 | 0.093 | 0.080 |
| Zfp213 | 0.076 | 0.043 | 0.051 |
| Qrsl1 | 0.076 | 0.043 | 0.040 |
| Ppm1h | 0.075 | 0.025 | 0.020 |

| | | | |
|---|---|---|---|
| Ints10 | 0.075 | 0.079 | 0.087 |
| 1810032O08I | 0.075 | 0.112 | 0.084 |
| Slc10a3 | 0.075 | 0.035 | 0.053 |
| Phf7 | 0.075 | 0.057 | 0.062 |
| Pla2g4a | 0.075 | 0.224 | 0.118 |
| Hspa1b | 0.075 | 0.098 | 0.053 |
| Ern1 | 0.075 | 0.049 | 0.041 |
| Arid3b | 0.075 | 0.038 | 0.017 |
| Gprc5b | 0.075 | 0.152 | 0.248 |
| Kcnn3 | 0.075 | 0.022 | 0.029 |
| Alg13 | 0.075 | 0.102 | 0.077 |
| Acsf3 | 0.075 | 0.037 | 0.044 |
| Gys1 | 0.075 | 0.084 | 0.091 |
| Macrod2 | 0.075 | 0.090 | 0.077 |
| Cyp26b1 | 0.075 | 0.066 | 0.023 |
| Bcdin3d | 0.075 | 0.062 | 0.071 |
| Vps18 | 0.075 | 0.071 | 0.071 |
| Zfp809 | 0.075 | 0.075 | 0.086 |
| Prkce | 0.075 | 0.162 | 0.161 |
| Ifi47 | 0.075 | 0.018 | 0.016 |
| Zcchc2 | 0.075 | 0.073 | 0.058 |
| Wscd1 | 0.075 | 0.002 | 0.005 |
| Atpaf2 | 0.075 | 0.079 | 0.074 |
| Mllt1 | 0.075 | 0.075 | 0.081 |
| Hus1 | 0.075 | 0.043 | 0.033 |
| Tmc4 | 0.075 | 0.063 | 0.074 |
| Wwox | 0.075 | 0.079 | 0.082 |
| G6pdx | 0.075 | 0.058 | 0.073 |
| Apba3 | 0.075 | 0.081 | 0.091 |
| Wfs1 | 0.075 | 0.070 | 0.065 |
| Galns | 0.074 | 0.044 | 0.075 |
| Lnpk | 0.074 | 0.147 | 0.158 |
| Atg16l2 | 0.074 | 0.087 | 0.086 |
| Cgref1 | 0.074 | 0.052 | 0.144 |
| Trub1 | 0.074 | 0.117 | 0.101 |
| Six5 | 0.074 | 0.088 | 0.107 |
| Wdr24 | 0.074 | 0.049 | 0.050 |
| Nnat | 0.074 | 0.036 | 0.049 |
| Dhcr7 | 0.074 | 0.061 | 0.101 |
| Islr | 0.074 | 0.711 | 1.266 |
| B430010I23R | 0.074 | 0.099 | 0.073 |
| Mrps2 | 0.074 | 0.069 | 0.064 |
| Nipal3 | 0.074 | 0.019 | 0.006 |
| Cnksr3 | 0.074 | 0.130 | 0.085 |
| Utp20 | 0.074 | 0.088 | 0.077 |
| Coprs | 0.074 | 0.072 | 0.077 |
| Acad11 | 0.074 | 0.027 | 0.027 |
| Dync2li1 | 0.074 | 0.151 | 0.186 |

| | | | |
|---|---|---|---|
| Nr1d1 | 0.074 | 0.095 | 0.042 |
| Mcm9 | 0.074 | 0.064 | 0.055 |
| Rab11fip3 | 0.074 | 0.068 | 0.063 |
| Cd74 | 0.074 | 0.027 | 0.015 |
| Zbtb18 | 0.074 | 0.027 | 0.014 |
| Nup35 | 0.074 | 0.059 | 0.062 |
| Birc3 | 0.074 | 0.115 | 0.164 |
| Utp15 | 0.074 | 0.073 | 0.067 |
| Acot11 | 0.074 | 0.060 | 0.040 |
| Tyro3 | 0.074 | 0.198 | 0.135 |
| Irak4 | 0.074 | 0.042 | 0.053 |
| Hps1 | 0.074 | 0.065 | 0.080 |
| Amd2 | 0.074 | 0.083 | 0.086 |
| Mrm2 | 0.074 | 0.049 | 0.050 |
| Hoxa9 | 0.074 | 0.000 | 0.000 |
| Prrg1 | 0.074 | 0.062 | 0.093 |
| Ipmk | 0.074 | 0.066 | 0.071 |
| Gucy1b1 | 0.074 | 0.466 | 0.287 |
| Sgpl1 | 0.074 | 0.143 | 0.148 |
| Serpinb8 | 0.074 | 0.031 | 0.034 |
| Gpr173 | 0.074 | 0.069 | 0.082 |
| Vsig2 | 0.074 | 0.003 | 0.001 |
| Zfp653 | 0.074 | 0.090 | 0.072 |
| Polr3a | 0.073 | 0.055 | 0.042 |
| Cspg4 | 0.073 | 0.827 | 0.266 |
| Nsun6 | 0.073 | 0.058 | 0.077 |
| Nme7 | 0.073 | 0.093 | 0.108 |
| Aste1 | 0.073 | 0.076 | 0.076 |
| Plxna1 | 0.073 | 0.102 | 0.103 |
| Rad52 | 0.073 | 0.101 | 0.082 |
| Dbr1 | 0.073 | 0.072 | 0.064 |
| Echdc1 | 0.073 | 0.057 | 0.057 |
| Haus3 | 0.073 | 0.066 | 0.078 |
| Acsf2 | 0.073 | 0.057 | 0.080 |
| 2010315B03F | 0.073 | 0.048 | 0.053 |
| Vps33b | 0.073 | 0.051 | 0.047 |
| Zcchc4 | 0.073 | 0.064 | 0.060 |
| Ints2 | 0.073 | 0.099 | 0.082 |
| Rgma | 0.073 | 0.103 | 0.134 |
| Pigv | 0.073 | 0.054 | 0.050 |
| Grp | 0.073 | 0.041 | 0.007 |
| Mlh1 | 0.073 | 0.059 | 0.066 |
| Rtl4 | 0.073 | 0.001 | 0.016 |
| Itga8 | 0.073 | 0.853 | 0.173 |
| Me2 | 0.073 | 0.058 | 0.067 |
| Fhl2 | 0.073 | 0.149 | 0.162 |
| Tmem94 | 0.073 | 0.053 | 0.047 |
| Dolpp1 | 0.073 | 0.057 | 0.061 |

| | | | |
|---|---|---|---|
| Dbf4 | 0.073 | 0.054 | 0.046 |
| Fbxo17 | 0.073 | 0.147 | 0.145 |
| Ociad2 | 0.073 | 0.055 | 0.010 |
| Dcp1a | 0.073 | 0.082 | 0.069 |
| Fam167b | 0.073 | 0.001 | 0.001 |
| Zfp764 | 0.073 | 0.035 | 0.049 |
| Vit | 0.073 | 0.009 | 0.181 |
| Zfp81 | 0.073 | 0.058 | 0.064 |
| Olfml3 | 0.073 | 0.105 | 0.828 |
| Cklf | 0.073 | 0.098 | 0.109 |
| Zfp667 | 0.073 | 0.053 | 0.061 |
| Magee1 | 0.073 | 0.107 | 0.101 |
| Cfi | 0.073 | 0.000 | 0.002 |
| Bclaf3 | 0.073 | 0.139 | 0.116 |
| Gpatch2l | 0.072 | 0.085 | 0.081 |
| Myl7 | 0.072 | 0.049 | 0.070 |
| Pnpt1 | 0.072 | 0.062 | 0.052 |
| Tti2 | 0.072 | 0.056 | 0.059 |
| Crabp2 | 0.072 | 0.044 | 0.047 |
| Pi4kb | 0.072 | 0.075 | 0.084 |
| Cd1d1 | 0.072 | 0.054 | 0.041 |
| Mtmr4 | 0.072 | 0.076 | 0.073 |
| Pign | 0.072 | 0.071 | 0.068 |
| Ccdc93 | 0.072 | 0.065 | 0.079 |
| Sfi1 | 0.072 | 0.056 | 0.069 |
| Rfc5 | 0.072 | 0.073 | 0.093 |
| Prim2 | 0.072 | 0.065 | 0.095 |
| Tom1 | 0.072 | 0.080 | 0.113 |
| Csgalnact1 | 0.072 | 0.086 | 0.036 |
| 2-Sep | 0.072 | 0.066 | 0.075 |
| Nup37 | 0.072 | 0.071 | 0.078 |
| Mlst8 | 0.072 | 0.035 | 0.043 |
| Mre11a | 0.072 | 0.054 | 0.066 |
| Fam110b | 0.072 | 0.349 | 0.437 |
| Thap1 | 0.072 | 0.062 | 0.049 |
| Gpd1l | 0.072 | 0.075 | 0.080 |
| Arhgap27 | 0.072 | 0.004 | 0.008 |
| Tmem40 | 0.072 | 0.002 | 0.004 |
| Timeless | 0.072 | 0.022 | 0.046 |
| Ccdc181 | 0.072 | 0.033 | 0.021 |
| Spata24 | 0.072 | 0.080 | 0.059 |
| Nedd4l | 0.072 | 0.041 | 0.054 |
| Gatsl2 | 0.072 | 0.133 | 0.145 |
| Slc11a2 | 0.072 | 0.075 | 0.059 |
| Magi3 | 0.072 | 0.053 | 0.114 |
| Dnase2a | 0.072 | 0.056 | 0.067 |
| Zfp319 | 0.071 | 0.066 | 0.067 |
| Mtg1 | 0.071 | 0.051 | 0.046 |

| | | | |
|---|---|---|---|
| Eif2ak4 | 0.071 | 0.094 | 0.068 |
| Tbl2 | 0.071 | 0.081 | 0.100 |
| Tram1l1 | 0.071 | 0.071 | 0.065 |
| Acacb | 0.071 | 0.010 | 0.009 |
| 1-Sep | 0.071 | 0.048 | 0.050 |
| Fermt3 | 0.071 | 0.005 | 0.003 |
| Arsk | 0.071 | 0.097 | 0.110 |
| Ogfod1 | 0.071 | 0.048 | 0.044 |
| Elac2 | 0.071 | 0.073 | 0.062 |
| Zscan12 | 0.071 | 0.062 | 0.071 |
| Zfp574 | 0.071 | 0.078 | 0.075 |
| Acad12 | 0.071 | 0.013 | 0.024 |
| Zfp729a | 0.071 | 0.051 | 0.053 |
| Adat2 | 0.071 | 0.047 | 0.052 |
| Kcnh1 | 0.071 | 0.000 | 0.001 |
| Neurl2 | 0.071 | 0.203 | 0.080 |
| Cryz | 0.071 | 0.082 | 0.083 |
| Lmbr1 | 0.071 | 0.059 | 0.097 |
| Gpkow | 0.071 | 0.069 | 0.066 |
| Slc35d2 | 0.071 | 0.034 | 0.039 |
| Plk4 | 0.071 | 0.064 | 0.071 |
| Tnni1 | 0.071 | 0.014 | 0.015 |
| Zfp707 | 0.071 | 0.087 | 0.056 |
| Erlin1 | 0.071 | 0.042 | 0.068 |
| Pde7b | 0.071 | 0.008 | 0.005 |
| Col4a6 | 0.071 | 0.298 | 0.168 |
| Plekha2 | 0.071 | 0.102 | 0.088 |
| Bcl7a | 0.071 | 0.172 | 0.107 |
| Pdxk | 0.071 | 0.054 | 0.051 |
| Kat2a | 0.071 | 0.082 | 0.084 |
| Vstm5 | 0.071 | 0.007 | 0.010 |
| Epha7 | 0.071 | 0.098 | 0.225 |
| Hmgxb3 | 0.071 | 0.071 | 0.070 |
| Alox5ap | 0.071 | 0.015 | 0.020 |
| Hist1h4i | 0.071 | 0.032 | 0.028 |
| Clic5 | 0.071 | 0.005 | 0.003 |
| Hspb2 | 0.071 | 0.605 | 0.200 |
| Polr1e | 0.071 | 0.053 | 0.041 |
| Tacc3 | 0.070 | 0.040 | 0.048 |
| Fbxl18 | 0.070 | 0.073 | 0.098 |
| Hspb6 | 0.070 | 0.333 | 0.071 |
| Tsr2 | 0.070 | 0.095 | 0.108 |
| Gemin8 | 0.070 | 0.056 | 0.048 |
| Idi1 | 0.070 | 0.118 | 0.130 |
| Gtpbp8 | 0.070 | 0.059 | 0.089 |
| Ahr | 0.070 | 0.018 | 0.013 |
| Flrt2 | 0.070 | 0.240 | 0.540 |
| Fbxo36 | 0.070 | 0.033 | 0.045 |

| | | | |
|---|---|---|---|
| Bcl2l2 | 0.070 | 0.086 | 0.075 |
| Mtrf1l | 0.070 | 0.083 | 0.087 |
| Slc25a34 | 0.070 | 0.053 | 0.012 |
| Gfm2 | 0.070 | 0.041 | 0.039 |
| Shq1 | 0.070 | 0.048 | 0.040 |
| Diras2 | 0.070 | 0.025 | 0.064 |
| Arhgef9 | 0.070 | 0.063 | 0.072 |
| Eya3 | 0.070 | 0.056 | 0.056 |
| Slc35g3 | 0.070 | 0.000 | 0.000 |
| Dyrk1b | 0.070 | 0.043 | 0.048 |
| Tlr4 | 0.070 | 0.026 | 0.045 |
| Zfp943 | 0.070 | 0.067 | 0.077 |
| Oplah | 0.070 | 0.059 | 0.084 |
| Mill2 | 0.070 | 0.032 | 0.116 |
| Borcs5 | 0.070 | 0.077 | 0.075 |
| Iqce | 0.070 | 0.058 | 0.092 |
| Zfp628 | 0.070 | 0.080 | 0.082 |
| Mettl13 | 0.070 | 0.048 | 0.060 |
| Dhx33 | 0.070 | 0.091 | 0.133 |
| Pex12 | 0.070 | 0.053 | 0.071 |
| Jrkl | 0.070 | 0.067 | 0.061 |
| Fpgs | 0.070 | 0.067 | 0.098 |
| Fgd4 | 0.070 | 0.058 | 0.119 |
| Slc23a2 | 0.070 | 0.058 | 0.054 |
| Bmp2k | 0.070 | 0.100 | 0.095 |
| Pde4dip | 0.070 | 0.190 | 0.165 |
| Tctn1 | 0.070 | 0.053 | 0.077 |
| Taf5 | 0.070 | 0.076 | 0.079 |
| Kif23 | 0.070 | 0.039 | 0.032 |
| Pus3 | 0.070 | 0.047 | 0.048 |
| Spef1 | 0.070 | 0.037 | 0.040 |
| Snai2 | 0.070 | 0.162 | 0.204 |
| Hs1bp3 | 0.069 | 0.054 | 0.059 |
| Pde5a | 0.069 | 0.648 | 0.497 |
| Tmem186 | 0.069 | 0.025 | 0.030 |
| 2900060B14F | 0.069 | 0.059 | 0.050 |
| Zfp770 | 0.069 | 0.061 | 0.061 |
| Stxbp4 | 0.069 | 0.080 | 0.088 |
| Tbc1d13 | 0.069 | 0.035 | 0.053 |
| Ror1 | 0.069 | 0.537 | 0.289 |
| Lrrn1 | 0.069 | 0.090 | 0.080 |
| Dcaf4 | 0.069 | 0.064 | 0.103 |
| Nubpl | 0.069 | 0.043 | 0.041 |
| Slc2a8 | 0.069 | 0.089 | 0.088 |
| Rasa4 | 0.069 | 0.023 | 0.072 |
| Map1a | 0.069 | 0.343 | 0.225 |
| Rab31 | 0.069 | 0.145 | 0.140 |
| Fndc4 | 0.069 | 0.139 | 0.212 |

| | | | |
|---|---|---|---|
| Atp2b2 | 0.069 | 0.001 | 0.000 |
| Prmt6 | 0.069 | 0.045 | 0.039 |
| Zfp618 | 0.069 | 0.031 | 0.126 |
| Leng9 | 0.069 | 0.096 | 0.116 |
| Ly6c1 | 0.069 | 0.008 | 0.185 |
| Slc35c1 | 0.069 | 0.033 | 0.034 |
| Coa4 | 0.069 | 0.028 | 0.036 |
| Zfp825 | 0.069 | 0.066 | 0.074 |
| Rab29 | 0.069 | 0.045 | 0.062 |
| Nox4 | 0.069 | 0.024 | 0.056 |
| Arxes2 | 0.069 | 0.118 | 0.160 |
| Akap10 | 0.069 | 0.052 | 0.046 |
| Ramp1 | 0.069 | 0.833 | 0.228 |
| Pms2 | 0.069 | 0.068 | 0.072 |
| Hps3 | 0.069 | 0.056 | 0.048 |
| Vav2 | 0.069 | 0.032 | 0.081 |
| Cdan1 | 0.069 | 0.053 | 0.056 |
| Letmd1 | 0.069 | 0.105 | 0.104 |
| Tada2a | 0.068 | 0.092 | 0.126 |
| Slc18b1 | 0.068 | 0.056 | 0.050 |
| Sh3bgr | 0.068 | 0.668 | 0.086 |
| Tk2 | 0.068 | 0.059 | 0.074 |
| Rap1gap2 | 0.068 | 0.061 | 0.017 |
| Dnm3 | 0.068 | 0.132 | 0.119 |
| E2f3 | 0.068 | 0.061 | 0.054 |
| Coil | 0.068 | 0.066 | 0.061 |
| Haus8 | 0.068 | 0.092 | 0.064 |
| Dapp1 | 0.068 | 0.082 | 0.077 |
| Acvr2a | 0.068 | 0.070 | 0.108 |
| Cacna1g | 0.068 | 0.171 | 0.208 |
| Plekhm1 | 0.068 | 0.047 | 0.050 |
| Inpp5f | 0.068 | 0.146 | 0.117 |
| Rabggta | 0.068 | 0.035 | 0.035 |
| Slc25a45 | 0.068 | 0.015 | 0.008 |
| Dpp7 | 0.068 | 0.060 | 0.100 |
| Zdhhc2 | 0.068 | 0.115 | 0.042 |
| Hectd3 | 0.068 | 0.058 | 0.058 |
| Map7d3 | 0.068 | 0.095 | 0.136 |
| Gm45051 | 0.068 | 0.037 | 0.033 |
| Prmt9 | 0.068 | 0.084 | 0.085 |
| Mtmr12 | 0.068 | 0.063 | 0.048 |
| Bri3bp | 0.068 | 0.157 | 0.158 |
| Krba1 | 0.068 | 0.089 | 0.071 |
| Flnc | 0.068 | 0.255 | 0.293 |
| Gbp4 | 0.068 | 0.001 | 0.001 |
| Zkscan8 | 0.068 | 0.046 | 0.062 |
| B3galt4 | 0.068 | 0.057 | 0.054 |
| Irs2 | 0.068 | 0.563 | 0.450 |

| | | | |
|---|---|---|---|
| Repin1 | 0.068 | 0.045 | 0.054 |
| Zfp994 | 0.067 | 0.041 | 0.054 |
| P3h2 | 0.067 | 0.075 | 0.087 |
| Sap130 | 0.067 | 0.067 | 0.068 |
| Wdr47 | 0.067 | 0.061 | 0.047 |
| Zbtb34 | 0.067 | 0.065 | 0.075 |
| Pex1 | 0.067 | 0.062 | 0.054 |
| Orc5 | 0.067 | 0.047 | 0.052 |
| Slc35a5 | 0.067 | 0.087 | 0.063 |
| Psat1 | 0.067 | 0.048 | 0.045 |
| Gm32036 | 0.067 | 0.018 | 0.006 |
| Cmtm4 | 0.067 | 0.093 | 0.071 |
| Lrrc8d | 0.067 | 0.031 | 0.044 |
| Cep89 | 0.067 | 0.071 | 0.072 |
| Slc25a10 | 0.067 | 0.040 | 0.055 |
| Ldlrad3 | 0.067 | 0.220 | 0.118 |
| Tnfaip3 | 0.067 | 0.088 | 0.073 |
| Chst2 | 0.067 | 0.321 | 0.410 |
| Taf9b | 0.067 | 0.038 | 0.030 |
| Fam199x | 0.067 | 0.072 | 0.061 |
| Ank2 | 0.067 | 0.241 | 0.692 |
| Cdca8 | 0.067 | 0.038 | 0.043 |
| Slc12a9 | 0.067 | 0.072 | 0.077 |
| Sipa1l3 | 0.067 | 0.066 | 0.079 |
| Lrrc28 | 0.067 | 0.085 | 0.062 |
| Nsun5 | 0.067 | 0.050 | 0.049 |
| Nup133 | 0.067 | 0.073 | 0.086 |
| Src | 0.067 | 0.103 | 0.111 |
| Idua | 0.067 | 0.052 | 0.060 |
| Arhgef19 | 0.067 | 0.074 | 0.067 |
| Spg11 | 0.067 | 0.067 | 0.078 |
| Ctf1 | 0.067 | 0.092 | 0.064 |
| Clybl | 0.067 | 0.048 | 0.070 |
| Sprtn | 0.067 | 0.056 | 0.044 |
| Klhl42 | 0.067 | 0.085 | 0.078 |
| Chek2 | 0.067 | 0.024 | 0.025 |
| Gm49359 | 0.067 | 0.059 | 0.074 |
| Cyp4b1 | 0.067 | 0.001 | 0.001 |
| Purg | 0.067 | 0.109 | 0.133 |
| Prokr2 | 0.067 | 0.000 | 0.000 |
| Cchcr1 | 0.067 | 0.073 | 0.069 |
| Pop1 | 0.067 | 0.031 | 0.037 |
| Nr2c1 | 0.067 | 0.067 | 0.069 |
| 2900076A07F | 0.067 | 0.047 | 0.049 |
| Calhm2 | 0.067 | 0.087 | 0.121 |
| Il17rd | 0.067 | 0.043 | 0.128 |
| Lef1 | 0.067 | 0.002 | 0.000 |
| Dffa | 0.067 | 0.055 | 0.061 |

| | | | |
|---|---|---|---|
| Tango6 | 0.067 | 0.172 | 0.092 |
| Tlcd2 | 0.067 | 0.046 | 0.125 |
| Nop2 | 0.067 | 0.076 | 0.075 |
| Mlkl | 0.067 | 0.004 | 0.013 |
| Des | 0.067 | 0.567 | 0.154 |
| Paox | 0.067 | 0.091 | 0.103 |
| Papolg | 0.067 | 0.040 | 0.035 |
| Lefty1 | 0.067 | 0.041 | 0.069 |
| Cdh4 | 0.067 | 0.071 | 0.097 |
| Letm2 | 0.067 | 0.084 | 0.110 |
| Trim23 | 0.066 | 0.074 | 0.073 |
| Fhod1 | 0.066 | 0.079 | 0.067 |
| 2700049A03R | 0.066 | 0.083 | 0.068 |
| Eps8 | 0.066 | 0.033 | 0.256 |
| Gm9856 | 0.066 | 0.060 | 0.059 |
| Gpr4 | 0.066 | 0.003 | 0.004 |
| Zfp959 | 0.066 | 0.056 | 0.055 |
| Fbxw4 | 0.066 | 0.098 | 0.092 |
| Rubcn | 0.066 | 0.086 | 0.078 |
| Flt4 | 0.066 | 0.002 | 0.003 |
| Ccdc114 | 0.066 | 0.030 | 0.037 |
| Nle1 | 0.066 | 0.063 | 0.067 |
| Mllt11 | 0.066 | 0.101 | 0.085 |
| Ntrk3 | 0.066 | 0.660 | 0.709 |
| Fam71a | 0.066 | 0.143 | 0.163 |
| Cnrip1 | 0.066 | 0.146 | 0.308 |
| Gmeb2 | 0.066 | 0.062 | 0.050 |
| Abca13 | 0.066 | 0.000 | 0.001 |
| S1pr2 | 0.066 | 0.092 | 0.316 |
| Iqsec2 | 0.066 | 0.066 | 0.097 |
| Zfp235 | 0.066 | 0.041 | 0.043 |
| Gtf2e1 | 0.066 | 0.033 | 0.045 |
| Slc4a3 | 0.066 | 0.169 | 0.134 |
| Tbrg4 | 0.066 | 0.048 | 0.045 |
| Stard10 | 0.066 | 0.065 | 0.048 |
| Tnfrsf26 | 0.066 | 0.000 | 0.001 |
| Sqle | 0.066 | 0.177 | 0.194 |
| Rhbdd1 | 0.066 | 0.031 | 0.024 |
| Fam19a5 | 0.066 | 0.082 | 0.134 |
| Noa1 | 0.066 | 0.081 | 0.069 |
| Ankrd44 | 0.066 | 0.338 | 0.207 |
| Zbtb12 | 0.066 | 0.081 | 0.075 |
| Gtf3c5 | 0.066 | 0.056 | 0.054 |
| Adck2 | 0.066 | 0.050 | 0.064 |
| B3galnt2 | 0.066 | 0.119 | 0.079 |
| Gm21988 | 0.066 | 0.039 | 0.043 |
| Rpusd1 | 0.066 | 0.037 | 0.065 |
| Bend4 | 0.066 | 0.024 | 0.016 |

| | | | |
|---|---|---|---|
| Spry1 | 0.066 | 0.193 | 0.389 |
| Btbd19 | 0.066 | 0.024 | 0.033 |
| Zbtb48 | 0.065 | 0.039 | 0.037 |
| Abi3bp | 0.065 | 0.686 | 0.759 |
| Plekha4 | 0.065 | 0.139 | 0.152 |
| Enox2 | 0.065 | 0.027 | 0.028 |
| Irak2 | 0.065 | 0.118 | 0.029 |
| Atg4a | 0.065 | 0.082 | 0.079 |
| Taf4 | 0.065 | 0.064 | 0.042 |
| Nudt17 | 0.065 | 0.019 | 0.024 |
| Spc24 | 0.065 | 0.068 | 0.078 |
| Zfp932 | 0.065 | 0.066 | 0.067 |
| Pdgfra | 0.065 | 0.173 | 0.799 |
| Ccdc130 | 0.065 | 0.061 | 0.054 |
| Zfp51 | 0.065 | 0.037 | 0.037 |
| Zfp711 | 0.065 | 0.033 | 0.033 |
| Zfp952 | 0.065 | 0.044 | 0.053 |
| Ythdc2 | 0.065 | 0.089 | 0.079 |
| Pdss2 | 0.065 | 0.066 | 0.071 |
| Xndc1 | 0.065 | 0.056 | 0.074 |
| Ifi27l2a | 0.065 | 0.043 | 0.136 |
| Zfp146 | 0.065 | 0.063 | 0.080 |
| Cacnb4 | 0.065 | 0.022 | 0.015 |
| Nrgn | 0.065 | 0.377 | 0.040 |
| Timp4 | 0.065 | 0.390 | 0.061 |
| Cdc20 | 0.065 | 0.038 | 0.034 |
| Gpr153 | 0.065 | 0.286 | 0.501 |
| Spindoc | 0.065 | 0.089 | 0.079 |
| Cnnm2 | 0.065 | 0.088 | 0.042 |
| Nsun3 | 0.065 | 0.028 | 0.034 |
| Pqlc3 | 0.065 | 0.053 | 0.070 |
| Gpn2 | 0.065 | 0.042 | 0.040 |
| Ttc7b | 0.065 | 0.141 | 0.082 |
| Hhip | 0.065 | 0.013 | 0.057 |
| Eif5a2 | 0.065 | 0.054 | 0.045 |
| B830012L14F | 0.065 | 0.051 | 0.199 |
| Rfc4 | 0.065 | 0.063 | 0.101 |
| Gbe1 | 0.065 | 0.095 | 0.085 |
| Vipr1 | 0.065 | 0.028 | 0.006 |
| Tmem201 | 0.065 | 0.070 | 0.071 |
| Clp1 | 0.065 | 0.066 | 0.053 |
| Nqo2 | 0.065 | 0.065 | 0.099 |
| Mapkbp1 | 0.064 | 0.083 | 0.079 |
| Zkscan14 | 0.064 | 0.039 | 0.036 |
| Atp8a1 | 0.064 | 0.026 | 0.026 |
| C1qtnf12 | 0.064 | 0.333 | 0.245 |
| Zfp317 | 0.064 | 0.074 | 0.064 |
| Gins2 | 0.064 | 0.077 | 0.110 |

| | | | |
|---|---|---|---|
| Hunk | 0.064 | 0.087 | 0.105 |
| Cmss1 | 0.064 | 0.072 | 0.086 |
| Slc29a3 | 0.064 | 0.081 | 0.104 |
| Trdmt1 | 0.064 | 0.050 | 0.033 |
| Zfp668 | 0.064 | 0.045 | 0.051 |
| Zbtb8a | 0.064 | 0.096 | 0.109 |
| Dtl | 0.064 | 0.085 | 0.132 |
| Dmpk | 0.064 | 0.527 | 0.230 |
| Ippk | 0.064 | 0.051 | 0.052 |
| Hmgcs2 | 0.064 | 0.279 | 0.138 |
| Ift80 | 0.064 | 0.063 | 0.053 |
| Ccdc125 | 0.064 | 0.010 | 0.006 |
| Fam20a | 0.064 | 0.103 | 0.115 |
| B630019K06F | 0.064 | 0.045 | 0.041 |
| Plagl2 | 0.064 | 0.072 | 0.057 |
| Npr3 | 0.064 | 0.157 | 0.130 |
| Gp1ba | 0.064 | 0.000 | 0.001 |
| Lockd | 0.064 | 0.043 | 0.041 |
| Zscan29 | 0.064 | 0.069 | 0.069 |
| Ermp1 | 0.064 | 0.103 | 0.077 |
| Ano1 | 0.064 | 0.494 | 0.109 |
| Chst7 | 0.064 | 0.012 | 0.015 |
| Ehd3 | 0.064 | 0.064 | 0.034 |
| Arhgef3 | 0.064 | 0.011 | 0.047 |
| Dbndd2 | 0.064 | 0.057 | 0.031 |
| Dpep1 | 0.064 | 0.336 | 0.875 |
| Rad9a | 0.064 | 0.033 | 0.040 |
| L3mbtl2 | 0.064 | 0.034 | 0.028 |
| Zbtb17 | 0.064 | 0.060 | 0.048 |
| Dupd1 | 0.063 | 0.002 | 0.002 |
| Pcdhb17 | 0.063 | 0.053 | 0.049 |
| Gtpbp1 | 0.063 | 0.062 | 0.048 |
| Cep170b | 0.063 | 0.153 | 0.112 |
| Glis3 | 0.063 | 0.017 | 0.032 |
| Ermard | 0.063 | 0.060 | 0.049 |
| Adgrl4 | 0.063 | 0.002 | 0.001 |
| Klhdc1 | 0.063 | 0.056 | 0.045 |
| Rhbdd3 | 0.063 | 0.076 | 0.097 |
| Xpnpep3 | 0.063 | 0.053 | 0.061 |
| Cdkn2c | 0.063 | 0.087 | 0.048 |
| Zbtb46 | 0.063 | 0.037 | 0.041 |
| Flad1 | 0.063 | 0.049 | 0.055 |
| Pax9 | 0.063 | 0.032 | 0.015 |
| Poc1b | 0.063 | 0.068 | 0.094 |
| Wfdc2 | 0.063 | 0.002 | 0.004 |
| Gm26724 | 0.063 | 0.074 | 0.058 |
| Dtnb | 0.063 | 0.075 | 0.077 |
| Wdr91 | 0.063 | 0.063 | 0.060 |

| | | | |
|---|---|---|---|
| Rnf144b | 0.063 | 0.018 | 0.003 |
| Mthfs | 0.063 | 0.088 | 0.077 |
| BC017158 | 0.063 | 0.049 | 0.060 |
| Grwd1 | 0.063 | 0.067 | 0.073 |
| Fbxo10 | 0.063 | 0.055 | 0.060 |
| Dnaja4 | 0.063 | 0.055 | 0.038 |
| Ficd | 0.063 | 0.049 | 0.051 |
| Zfp866 | 0.063 | 0.053 | 0.066 |
| Nptxr | 0.063 | 0.109 | 0.098 |
| Pde9a | 0.063 | 0.041 | 0.042 |
| Zfp623 | 0.063 | 0.044 | 0.052 |
| Chic1 | 0.063 | 0.089 | 0.074 |
| Crebl2 | 0.062 | 0.045 | 0.039 |
| Smug1 | 0.062 | 0.044 | 0.037 |
| Slc25a26 | 0.062 | 0.037 | 0.037 |
| Col12a1 | 0.062 | 0.260 | 0.742 |
| 2510046G10I | 0.062 | 0.031 | 0.028 |
| Zfp629 | 0.062 | 0.052 | 0.051 |
| Pdgfc | 0.062 | 0.252 | 0.107 |
| Zfp143 | 0.062 | 0.059 | 0.049 |
| Ticam1 | 0.062 | 0.057 | 0.071 |
| Gnai1 | 0.062 | 0.247 | 0.128 |
| Lysmd1 | 0.062 | 0.049 | 0.048 |
| Asb13 | 0.062 | 0.078 | 0.056 |
| Zc3h6 | 0.062 | 0.086 | 0.097 |
| Tbccd1 | 0.062 | 0.058 | 0.055 |
| Fahd2a | 0.062 | 0.059 | 0.027 |
| Ccdc94 | 0.062 | 0.040 | 0.036 |
| Rdh11 | 0.062 | 0.106 | 0.086 |
| Lgals7 | 0.062 | 0.266 | 0.194 |
| Ephb6 | 0.062 | 0.122 | 0.055 |
| Trim13 | 0.062 | 0.037 | 0.029 |
| Prpsap2 | 0.062 | 0.076 | 0.072 |
| Zfp119a | 0.062 | 0.043 | 0.052 |
| Sap18b | 0.062 | 0.055 | 0.053 |
| Sdhaf3 | 0.062 | 0.085 | 0.080 |
| P2rx7 | 0.062 | 0.034 | 0.011 |
| Fancm | 0.062 | 0.066 | 0.048 |
| Cyp2d22 | 0.062 | 0.077 | 0.180 |
| Zfp518b | 0.062 | 0.050 | 0.078 |
| Trp53rkb | 0.062 | 0.038 | 0.037 |
| Dbp | 0.062 | 0.125 | 0.107 |
| Gpm6a | 0.062 | 0.006 | 0.007 |
| Gm46430 | 0.062 | 0.056 | 0.076 |
| Tmem82 | 0.062 | 0.067 | 0.018 |
| Tlr2 | 0.062 | 0.066 | 0.063 |
| Pold1 | 0.062 | 0.050 | 0.065 |
| Ssu2 | 0.062 | 0.002 | 0.003 |

| | | | |
|---|---|---|---|
| Zfp760 | 0.062 | 0.037 | 0.048 |
| Alg12 | 0.062 | 0.038 | 0.039 |
| Osbpl7 | 0.062 | 0.042 | 0.040 |
| Ankrd28 | 0.062 | 0.062 | 0.061 |
| Rnf123 | 0.062 | 0.088 | 0.064 |
| Sacs | 0.062 | 0.155 | 0.038 |
| Arpin | 0.062 | 0.062 | 0.087 |
| Zfp780b | 0.061 | 0.063 | 0.066 |
| Klf16 | 0.061 | 0.082 | 0.084 |
| St3gal3 | 0.061 | 0.041 | 0.044 |
| Odf2l | 0.061 | 0.096 | 0.079 |
| Phlpp2 | 0.061 | 0.097 | 0.102 |
| Fstl3 | 0.061 | 0.435 | 0.220 |
| Ogfrl1 | 0.061 | 0.138 | 0.091 |
| Prkd1 | 0.061 | 0.048 | 0.033 |
| Tdrd7 | 0.061 | 0.049 | 0.047 |
| Rnd2 | 0.061 | 0.080 | 0.102 |
| Carf | 0.061 | 0.050 | 0.066 |
| Nrxn3 | 0.061 | 0.001 | 0.007 |
| Dus2 | 0.061 | 0.061 | 0.061 |
| Etaa1 | 0.061 | 0.083 | 0.108 |
| Pacsin3 | 0.061 | 0.177 | 0.112 |
| Colec11 | 0.061 | 0.038 | 0.047 |
| Tamm41 | 0.061 | 0.050 | 0.059 |
| Lysmd4 | 0.061 | 0.049 | 0.049 |
| Tctn3 | 0.061 | 0.032 | 0.040 |
| Pou6f1 | 0.061 | 0.069 | 0.040 |
| Pak7 | 0.061 | 0.065 | 0.010 |
| Slc41a3 | 0.061 | 0.247 | 0.180 |
| Mlh3 | 0.061 | 0.054 | 0.059 |
| Snx25 | 0.061 | 0.062 | 0.041 |
| Ankrd26 | 0.061 | 0.088 | 0.095 |
| Myocd | 0.061 | 0.278 | 0.045 |
| Trpc5 | 0.061 | 0.001 | 0.004 |
| Gm9917 | 0.061 | 0.013 | 0.013 |
| Tmem51 | 0.061 | 0.066 | 0.045 |
| Camsap3 | 0.061 | 0.014 | 0.024 |
| Batf | 0.061 | 0.013 | 0.017 |
| Inafm2 | 0.061 | 0.139 | 0.037 |
| Zfp37 | 0.061 | 0.071 | 0.067 |
| Riox2 | 0.061 | 0.056 | 0.057 |
| Tbc1d16 | 0.061 | 0.098 | 0.176 |
| Epha2 | 0.061 | 0.011 | 0.013 |
| Cnst | 0.061 | 0.061 | 0.056 |
| Kcnk6 | 0.061 | 0.018 | 0.048 |
| Trmt2b | 0.061 | 0.081 | 0.078 |
| Ak4 | 0.061 | 0.008 | 0.009 |
| Sap30 | 0.061 | 0.052 | 0.035 |

| | | | |
|---|---|---|---|
| Stat4 | 0.061 | 0.006 | 0.003 |
| Gla | 0.061 | 0.060 | 0.043 |
| Tmem229b | 0.061 | 0.011 | 0.019 |
| Prkg1 | 0.061 | 0.272 | 0.140 |
| Zfp369 | 0.061 | 0.058 | 0.050 |
| Prkci | 0.061 | 0.080 | 0.087 |
| Gm26853 | 0.061 | 0.066 | 0.073 |
| Plekha6 | 0.060 | 0.007 | 0.012 |
| Gm14296 | 0.060 | 0.054 | 0.065 |
| Wars2 | 0.060 | 0.049 | 0.063 |
| Pdpr | 0.060 | 0.047 | 0.042 |
| Rel | 0.060 | 0.113 | 0.091 |
| Asb4 | 0.060 | 0.022 | 0.083 |
| Rspo3 | 0.060 | 0.503 | 0.206 |
| Prss35 | 0.060 | 0.242 | 0.515 |
| Csdc2 | 0.060 | 0.193 | 0.073 |
| Psmb9 | 0.060 | 0.011 | 0.014 |
| Elmod3 | 0.060 | 0.066 | 0.070 |
| Sema5a | 0.060 | 0.348 | 0.439 |
| Tfap4 | 0.060 | 0.045 | 0.054 |
| Slc4a8 | 0.060 | 0.013 | 0.009 |
| Fzd5 | 0.060 | 0.112 | 0.054 |
| Tdp1 | 0.060 | 0.067 | 0.073 |
| Got1 | 0.060 | 0.041 | 0.049 |
| Zc3hav1l | 0.060 | 0.051 | 0.093 |
| Tcf7 | 0.060 | 0.019 | 0.004 |
| Shank3 | 0.060 | 0.004 | 0.009 |
| Zbed4 | 0.060 | 0.027 | 0.035 |
| Polrmt | 0.060 | 0.049 | 0.042 |
| Trim34a | 0.060 | 0.017 | 0.027 |
| Scfd2 | 0.060 | 0.033 | 0.059 |
| Ska2 | 0.060 | 0.047 | 0.056 |
| Tsen54 | 0.060 | 0.066 | 0.050 |
| Parp3 | 0.060 | 0.232 | 0.249 |
| Sat2 | 0.060 | 0.106 | 0.098 |
| Fbxl4 | 0.060 | 0.065 | 0.061 |
| Slc26a11 | 0.060 | 0.052 | 0.064 |
| Gm17501 | 0.060 | 0.058 | 0.039 |
| Tsen2 | 0.060 | 0.064 | 0.111 |
| Myc | 0.060 | 0.148 | 0.161 |
| Hsf2 | 0.060 | 0.098 | 0.083 |
| Gm20300 | 0.060 | 0.044 | 0.048 |
| Cdh26 | 0.060 | 0.003 | 0.006 |
| Il1r1 | 0.060 | 0.219 | 0.215 |
| Ccdc136 | 0.060 | 0.163 | 0.092 |
| Pcdhgc4 | 0.060 | 0.049 | 0.064 |
| Cbwd1 | 0.060 | 0.058 | 0.052 |
| Inf2 | 0.059 | 0.093 | 0.056 |

| | | | |
|---|---|---|---|
| Fgf12 | 0.059 | 0.003 | 0.005 |
| Cd38 | 0.059 | 0.002 | 0.005 |
| Mib2 | 0.059 | 0.105 | 0.123 |
| Sfmbt1 | 0.059 | 0.107 | 0.095 |
| Ttpal | 0.059 | 0.080 | 0.080 |
| Pamr1 | 0.059 | 0.356 | 0.400 |
| Zdhhc13 | 0.059 | 0.042 | 0.039 |
| Zkscan6 | 0.059 | 0.066 | 0.054 |
| Map3k14 | 0.059 | 0.125 | 0.113 |
| Nup205 | 0.059 | 0.058 | 0.051 |
| Dis3l | 0.059 | 0.042 | 0.046 |
| Ift88 | 0.059 | 0.066 | 0.068 |
| Gemin5 | 0.059 | 0.059 | 0.061 |
| Slc25a14 | 0.059 | 0.065 | 0.062 |
| Rp2 | 0.059 | 0.069 | 0.056 |
| Spin2c | 0.059 | 0.018 | 0.022 |
| Irf7 | 0.059 | 0.023 | 0.042 |
| Tle2 | 0.059 | 0.074 | 0.120 |
| Aoc3 | 0.059 | 0.681 | 0.331 |
| Ifih1 | 0.059 | 0.020 | 0.031 |
| Fam111a | 0.059 | 0.037 | 0.122 |
| Ccdc191 | 0.059 | 0.050 | 0.068 |
| Zfp334 | 0.059 | 0.057 | 0.058 |
| Abca5 | 0.059 | 0.068 | 0.084 |
| Elmod2 | 0.059 | 0.077 | 0.085 |
| Zfp12 | 0.059 | 0.050 | 0.052 |
| Cenpe | 0.059 | 0.035 | 0.032 |
| Oxld1 | 0.059 | 0.051 | 0.059 |
| Carmil1 | 0.059 | 0.109 | 0.128 |
| Tmem104 | 0.059 | 0.028 | 0.038 |
| Cdca3 | 0.059 | 0.037 | 0.046 |
| Mfsd8 | 0.059 | 0.042 | 0.045 |
| Tefm | 0.059 | 0.051 | 0.050 |
| Cxxc4 | 0.059 | 0.012 | 0.026 |
| Hdac6 | 0.059 | 0.082 | 0.070 |
| Smim1 | 0.059 | 0.202 | 0.527 |
| Dach1 | 0.059 | 0.014 | 0.002 |
| Katnal1 | 0.059 | 0.080 | 0.091 |
| Tbc1d9 | 0.059 | 0.046 | 0.025 |
| Rem1 | 0.058 | 0.116 | 0.116 |
| Gbp2 | 0.058 | 0.020 | 0.046 |
| Arhgef26 | 0.058 | 0.081 | 0.084 |
| Rgs10 | 0.058 | 0.234 | 0.169 |
| 1700052K11F | 0.058 | 0.054 | 0.044 |
| Lipe | 0.058 | 0.040 | 0.042 |
| Arsb | 0.058 | 0.100 | 0.172 |
| Tmem241 | 0.058 | 0.037 | 0.057 |
| Arel1 | 0.058 | 0.042 | 0.048 |

| | | | |
|---|---|---|---|
| Mturn | 0.058 | 0.031 | 0.078 |
| Ehbp1 | 0.058 | 0.131 | 0.174 |
| Wnt4 | 0.058 | 0.042 | 0.052 |
| D17H6S53E | 0.058 | 0.058 | 0.051 |
| Utp14b | 0.058 | 0.091 | 0.117 |
| Zfp276 | 0.058 | 0.051 | 0.049 |
| Foxred1 | 0.058 | 0.062 | 0.063 |
| Nln | 0.058 | 0.093 | 0.086 |
| Rasgrp3 | 0.058 | 0.006 | 0.005 |
| Inpp4a | 0.058 | 0.104 | 0.084 |
| Meox1 | 0.058 | 0.026 | 0.171 |
| Rab3il1 | 0.058 | 0.056 | 0.255 |
| Angpt2 | 0.058 | 0.613 | 0.553 |
| Tgfb3 | 0.058 | 0.322 | 0.524 |
| Dysf | 0.058 | 0.130 | 0.023 |
| Iqcc | 0.058 | 0.054 | 0.063 |
| Nsdhl | 0.058 | 0.074 | 0.092 |
| Sox6 | 0.058 | 0.015 | 0.008 |
| Pwp2 | 0.058 | 0.037 | 0.037 |
| Efna4 | 0.058 | 0.101 | 0.087 |
| Cep85 | 0.058 | 0.045 | 0.040 |
| Tlr3 | 0.058 | 0.014 | 0.032 |
| Tbc1d2 | 0.058 | 0.005 | 0.003 |
| Megf9 | 0.058 | 0.048 | 0.063 |
| Pomt1 | 0.058 | 0.061 | 0.074 |
| Synm | 0.058 | 0.077 | 0.023 |
| Cisd3 | 0.058 | 0.048 | 0.034 |
| Cdk5rap2 | 0.058 | 0.060 | 0.118 |
| Notch3 | 0.058 | 0.395 | 0.304 |
| Tspan32 | 0.058 | 0.006 | 0.011 |
| Ppard | 0.058 | 0.081 | 0.074 |
| Parp10 | 0.058 | 0.020 | 0.022 |
| Polg2 | 0.058 | 0.132 | 0.132 |
| Snx32 | 0.058 | 0.040 | 0.034 |
| Gm10036 | 0.058 | 0.045 | 0.062 |
| Harbi1 | 0.058 | 0.047 | 0.047 |
| Vps37d | 0.058 | 0.139 | 0.124 |
| Tmem35b | 0.058 | 0.042 | 0.029 |
| Bcs1l | 0.058 | 0.067 | 0.077 |
| Icosl | 0.058 | 0.004 | 0.024 |
| Cenpm | 0.058 | 0.028 | 0.040 |
| Atg2b | 0.058 | 0.065 | 0.067 |
| Fitm2 | 0.058 | 0.056 | 0.044 |
| Zfp697 | 0.058 | 0.033 | 0.043 |
| Yars2 | 0.058 | 0.057 | 0.062 |
| Mmp11 | 0.058 | 0.193 | 0.187 |
| Jade2 | 0.058 | 0.083 | 0.024 |
| Zxdb | 0.058 | 0.072 | 0.052 |

| | | | |
|---|---|---|---|
| Klhl15 | 0.057 | 0.076 | 0.072 |
| Tnfrsf9 | 0.057 | 0.001 | 0.000 |
| Unkl | 0.057 | 0.066 | 0.063 |
| Slc25a53 | 0.057 | 0.037 | 0.042 |
| Cox15 | 0.057 | 0.039 | 0.036 |
| 0610040B10R | 0.057 | 0.030 | 0.033 |
| Ptgds | 0.057 | 0.004 | 0.004 |
| Naaladl2 | 0.057 | 0.027 | 0.067 |
| Lrig1 | 0.057 | 0.191 | 0.101 |
| Sema6c | 0.057 | 0.101 | 0.185 |
| Gm15261 | 0.057 | 0.145 | 0.101 |
| Usb1 | 0.057 | 0.067 | 0.075 |
| Zfp597 | 0.057 | 0.045 | 0.049 |
| Traf3 | 0.057 | 0.062 | 0.066 |
| Mex3b | 0.057 | 0.111 | 0.143 |
| Ano8 | 0.057 | 0.060 | 0.062 |
| Dgke | 0.057 | 0.059 | 0.023 |
| Mdc1 | 0.057 | 0.065 | 0.069 |
| Gmpr | 0.057 | 0.094 | 0.034 |
| Prrg3 | 0.057 | 0.117 | 0.145 |
| Zfp383 | 0.057 | 0.056 | 0.103 |
| Galc | 0.057 | 0.033 | 0.040 |
| Zfp934 | 0.057 | 0.056 | 0.061 |
| Fibin | 0.057 | 0.592 | 0.835 |
| Top1mt | 0.057 | 0.037 | 0.041 |
| Ampd3 | 0.057 | 0.006 | 0.032 |
| Zfp346 | 0.057 | 0.052 | 0.044 |
| Inpp5e | 0.057 | 0.053 | 0.057 |
| Il20rb | 0.057 | 0.020 | 0.023 |
| Alg6 | 0.057 | 0.066 | 0.106 |
| Mpz | 0.057 | 0.129 | 0.097 |
| Lin9 | 0.057 | 0.047 | 0.040 |
| Zfp472 | 0.057 | 0.029 | 0.036 |
| Klhl21 | 0.057 | 0.131 | 0.073 |
| Fam13a | 0.057 | 0.062 | 0.103 |
| Tifa | 0.057 | 0.065 | 0.110 |
| Gm6277 | 0.057 | 0.082 | 0.062 |
| 2310001H17R | 0.057 | 0.012 | 0.017 |
| Spidr | 0.057 | 0.102 | 0.135 |
| Prc1 | 0.057 | 0.049 | 0.050 |
| Zbtb21 | 0.057 | 0.099 | 0.100 |
| Exd2 | 0.057 | 0.063 | 0.055 |
| 4833403J16R | 0.057 | 0.000 | 0.001 |
| Slc25a13 | 0.056 | 0.022 | 0.014 |
| Sowahc | 0.056 | 0.090 | 0.080 |
| Jag2 | 0.056 | 0.005 | 0.002 |
| Herc6 | 0.056 | 0.036 | 0.032 |
| Gm10138 | 0.056 | 0.038 | 0.044 |

| | | | |
|---|---|---|---|
| Il27ra | 0.056 | 0.007 | 0.021 |
| Gpx3 | 0.056 | 0.063 | 0.677 |
| Nprl3 | 0.056 | 0.039 | 0.033 |
| Vldlr | 0.056 | 0.500 | 0.646 |
| Dcbld2 | 0.056 | 0.119 | 0.224 |
| Nim1k | 0.056 | 0.064 | 0.084 |
| Msl3l2 | 0.056 | 0.088 | 0.072 |
| Homer1 | 0.056 | 0.110 | 0.106 |
| AI597479 | 0.056 | 0.056 | 0.066 |
| Fgf13 | 0.056 | 0.067 | 0.040 |
| Tmem17 | 0.056 | 0.075 | 0.070 |
| Dok1 | 0.056 | 0.067 | 0.112 |
| Ccdc73 | 0.056 | 0.037 | 0.043 |
| Fign | 0.056 | 0.021 | 0.047 |
| Trmt61a | 0.056 | 0.045 | 0.045 |
| Snupn | 0.056 | 0.066 | 0.068 |
| Plpp6 | 0.056 | 0.051 | 0.039 |
| 9930038B18R | 0.056 | 0.000 | 0.000 |
| Ranbp6 | 0.056 | 0.055 | 0.052 |
| Pcdhgc5 | 0.056 | 0.037 | 0.040 |
| Nol9 | 0.056 | 0.046 | 0.052 |
| Fen1 | 0.056 | 0.046 | 0.052 |
| Mcts2 | 0.056 | 0.107 | 0.071 |
| Nphp3 | 0.056 | 0.059 | 0.057 |
| Epb41l5 | 0.056 | 0.067 | 0.035 |
| Fgf18 | 0.056 | 0.062 | 0.292 |
| Slc38a6 | 0.056 | 0.039 | 0.087 |
| Cipc | 0.056 | 0.049 | 0.042 |
| Fancc | 0.056 | 0.029 | 0.037 |
| Pus7 | 0.055 | 0.099 | 0.074 |
| Amacr | 0.055 | 0.050 | 0.074 |
| Pnpo | 0.055 | 0.071 | 0.034 |
| Bcl3 | 0.055 | 0.095 | 0.079 |
| Pcdh10 | 0.055 | 0.025 | 0.034 |
| Hdac10 | 0.055 | 0.040 | 0.059 |
| Gdf15 | 0.055 | 0.054 | 0.038 |
| Rpusd3 | 0.055 | 0.038 | 0.030 |
| Mief2 | 0.055 | 0.041 | 0.043 |
| Srfbp1 | 0.055 | 0.051 | 0.045 |
| Schip1 | 0.055 | 0.084 | 0.012 |
| Adra2a | 0.055 | 0.002 | 0.003 |
| Zfp954 | 0.055 | 0.048 | 0.051 |
| Mphosph9 | 0.055 | 0.058 | 0.044 |
| Tagap1 | 0.055 | 0.039 | 0.040 |
| Inpp4b | 0.055 | 0.078 | 0.021 |
| Depdc5 | 0.055 | 0.052 | 0.058 |
| Chaf1a | 0.055 | 0.034 | 0.063 |
| Platr25 | 0.055 | 0.053 | 0.065 |

| | | | |
|---|---|---|---|
| Comtd1 | 0.055 | 0.071 | 0.081 |
| Adam33 | 0.055 | 0.117 | 0.209 |
| Szt2 | 0.055 | 0.049 | 0.041 |
| Traf3ip2 | 0.055 | 0.049 | 0.056 |
| Zmym1 | 0.055 | 0.048 | 0.059 |
| Prps2 | 0.055 | 0.085 | 0.048 |
| Tmem67 | 0.055 | 0.057 | 0.046 |
| Pter | 0.055 | 0.079 | 0.078 |
| Zfp715 | 0.055 | 0.059 | 0.047 |
| Vav3 | 0.055 | 0.117 | 0.107 |
| Fbxo46 | 0.055 | 0.040 | 0.037 |
| Gtpbp10 | 0.055 | 0.049 | 0.043 |
| Dgkk | 0.055 | 0.022 | 0.012 |
| Kremen1 | 0.055 | 0.076 | 0.079 |
| Nid2 | 0.055 | 0.235 | 0.354 |
| Gm38399 | 0.055 | 0.023 | 0.028 |
| Zkscan5 | 0.055 | 0.057 | 0.047 |
| Akr1b10 | 0.055 | 0.059 | 0.045 |
| Zfp747 | 0.055 | 0.041 | 0.039 |
| Nlrx1 | 0.055 | 0.068 | 0.075 |
| Klhl36 | 0.054 | 0.039 | 0.051 |
| Gm43672 | 0.054 | 0.054 | 0.037 |
| Myo5a | 0.054 | 0.123 | 0.121 |
| Guca1a | 0.054 | 0.026 | 0.081 |
| Ptar1 | 0.054 | 0.038 | 0.046 |
| Dcun1d4 | 0.054 | 0.053 | 0.049 |
| Gstk1 | 0.054 | 0.030 | 0.031 |
| Fam206a | 0.054 | 0.048 | 0.052 |
| 6720427I07R | 0.054 | 0.071 | 0.073 |
| Smg8 | 0.054 | 0.048 | 0.042 |
| Ccnd1 | 0.054 | 0.038 | 0.210 |
| Ercc8 | 0.054 | 0.045 | 0.042 |
| B3gnt5 | 0.054 | 0.050 | 0.022 |
| Tor4a | 0.054 | 0.013 | 0.006 |
| Skp2 | 0.054 | 0.047 | 0.056 |
| Gm6563 | 0.054 | 0.056 | 0.048 |
| Sgsm2 | 0.054 | 0.068 | 0.104 |
| Trmt12 | 0.054 | 0.034 | 0.033 |
| E130307A14F | 0.054 | 0.071 | 0.074 |
| Tmsb15b1 | 0.054 | 0.062 | 0.040 |
| Ssx2ip | 0.054 | 0.068 | 0.104 |
| Fam109b | 0.054 | 0.053 | 0.112 |
| Slc7a2 | 0.054 | 0.359 | 0.113 |
| Poc5 | 0.054 | 0.062 | 0.065 |
| Pola2 | 0.054 | 0.033 | 0.073 |
| Nqo1 | 0.054 | 0.127 | 0.036 |
| Cyp4f13 | 0.054 | 0.055 | 0.076 |
| Tmem131l | 0.054 | 0.125 | 0.139 |

| | | | |
|---|---|---|---|
| Nup43 | 0.054 | 0.061 | 0.065 |
| Kyat3 | 0.054 | 0.055 | 0.039 |
| Uhrf1 | 0.054 | 0.025 | 0.037 |
| Vars2 | 0.054 | 0.039 | 0.042 |
| Slc35e2 | 0.054 | 0.056 | 0.056 |
| Socs5 | 0.054 | 0.087 | 0.120 |
| Pdp1 | 0.054 | 0.101 | 0.078 |
| Cystm1 | 0.054 | 0.026 | 0.020 |
| Tmem38a | 0.054 | 0.083 | 0.110 |
| Cdkl5 | 0.054 | 0.152 | 0.077 |
| Atp6v0e2 | 0.054 | 0.059 | 0.040 |
| As3mt | 0.054 | 0.019 | 0.025 |
| Esr1 | 0.054 | 0.010 | 0.015 |
| Gcfc2 | 0.054 | 0.058 | 0.058 |
| Faap24 | 0.054 | 0.042 | 0.048 |
| Limk1 | 0.054 | 0.055 | 0.109 |
| Gspt2 | 0.054 | 0.046 | 0.045 |
| Lrrc1 | 0.053 | 0.050 | 0.070 |
| Agr2 | 0.053 | 0.000 | 0.001 |
| Trpc1 | 0.053 | 0.177 | 0.115 |
| L3hypdh | 0.053 | 0.101 | 0.238 |
| Rundc1 | 0.053 | 0.059 | 0.057 |
| Mia | 0.053 | 0.173 | 0.040 |
| Irak1bp1 | 0.053 | 0.202 | 0.190 |
| Igsf10 | 0.053 | 0.074 | 0.254 |
| Hlcs | 0.053 | 0.032 | 0.057 |
| Thumpd2 | 0.053 | 0.065 | 0.063 |
| Lrrc75a | 0.053 | 0.167 | 0.259 |
| Zswim1 | 0.053 | 0.035 | 0.031 |
| Trim30a | 0.053 | 0.003 | 0.015 |
| Spn | 0.053 | 0.001 | 0.001 |
| Lrrc3b | 0.053 | 0.005 | 0.001 |
| Ndc1 | 0.053 | 0.049 | 0.051 |
| Lrrc8b | 0.053 | 0.010 | 0.023 |
| Eefsec | 0.053 | 0.064 | 0.058 |
| Map6 | 0.053 | 0.111 | 0.074 |
| Dlx6 | 0.053 | 0.042 | 0.029 |
| Fbf1 | 0.053 | 0.054 | 0.055 |
| Bckdhb | 0.053 | 0.086 | 0.128 |
| Cblb | 0.053 | 0.125 | 0.131 |
| 9-Mar | 0.053 | 0.062 | 0.052 |
| Ypel1 | 0.053 | 0.048 | 0.078 |
| Zfp13 | 0.053 | 0.053 | 0.054 |
| Cep57l1 | 0.053 | 0.062 | 0.059 |
| Matn2 | 0.053 | 0.161 | 1.031 |
| Nipal4 | 0.053 | 0.037 | 0.005 |
| Mdga1 | 0.053 | 0.086 | 0.076 |
| A930004D18I | 0.053 | 0.040 | 0.050 |

| | | | |
|---|---|---|---|
| Sphk1 | 0.053 | 0.145 | 0.095 |
| Tpd52l1 | 0.053 | 0.068 | 0.044 |
| Nthl1 | 0.053 | 0.088 | 0.080 |
| Adnp2 | 0.053 | 0.065 | 0.067 |
| Med26 | 0.053 | 0.051 | 0.054 |
| Tenm4 | 0.053 | 0.081 | 0.041 |
| Msh6 | 0.053 | 0.073 | 0.068 |
| Slc46a3 | 0.053 | 0.010 | 0.006 |
| Rpgr | 0.053 | 0.038 | 0.036 |
| Eif4e3 | 0.053 | 0.095 | 0.081 |
| Zfp710 | 0.053 | 0.067 | 0.055 |
| Nemp1 | 0.053 | 0.052 | 0.042 |
| Hykk | 0.052 | 0.036 | 0.056 |
| Ung | 0.052 | 0.023 | 0.038 |
| Cep97 | 0.052 | 0.078 | 0.070 |
| Fastkd3 | 0.052 | 0.042 | 0.046 |
| C77080 | 0.052 | 0.015 | 0.009 |
| Per2 | 0.052 | 0.136 | 0.094 |
| Zfp119b | 0.052 | 0.049 | 0.038 |
| Hoxa7 | 0.052 | 0.001 | 0.000 |
| Mon1a | 0.052 | 0.052 | 0.038 |
| Kctd15 | 0.052 | 0.144 | 0.116 |
| Lmnb2 | 0.052 | 0.031 | 0.037 |
| Diexf | 0.052 | 0.054 | 0.049 |
| Pex26 | 0.052 | 0.041 | 0.037 |
| Pitpnm3 | 0.052 | 0.006 | 0.001 |
| Tppp | 0.052 | 0.014 | 0.021 |
| Kantr | 0.052 | 0.053 | 0.044 |
| Arid3a | 0.052 | 0.111 | 0.048 |
| Itgbl1 | 0.052 | 0.081 | 0.159 |
| Dgkq | 0.052 | 0.057 | 0.052 |
| Tkfc | 0.052 | 0.048 | 0.060 |
| Zbtb6 | 0.052 | 0.048 | 0.047 |
| Spata1 | 0.052 | 0.046 | 0.045 |
| Syt7 | 0.052 | 0.003 | 0.003 |
| Tbc1d2b | 0.052 | 0.150 | 0.188 |
| Gm42517 | 0.052 | 0.003 | 0.004 |
| Jak3 | 0.052 | 0.008 | 0.016 |
| Tnc | 0.052 | 0.259 | 0.659 |
| Zbtb42 | 0.052 | 0.027 | 0.018 |
| Pde4a | 0.052 | 0.100 | 0.093 |
| Bivm | 0.052 | 0.042 | 0.040 |
| Gba2 | 0.052 | 0.050 | 0.053 |
| Znrd1as | 0.052 | 0.025 | 0.030 |
| Lclat1 | 0.052 | 0.094 | 0.081 |
| Peg12 | 0.052 | 0.028 | 0.062 |
| Taf1a | 0.052 | 0.042 | 0.039 |
| Stk11ip | 0.052 | 0.037 | 0.045 |

| | | | |
|---|---|---|---|
| Ercc6 | 0.052 | 0.039 | 0.038 |
| Wdr55 | 0.052 | 0.060 | 0.048 |
| Poll | 0.052 | 0.033 | 0.039 |
| E130311K13F | 0.052 | 0.053 | 0.083 |
| Nt5dc3 | 0.052 | 0.079 | 0.044 |
| 2310022A10F | 0.052 | 0.056 | 0.051 |
| Bahd1 | 0.052 | 0.044 | 0.037 |
| Nmb | 0.052 | 0.022 | 0.033 |
| Mgat4a | 0.052 | 0.103 | 0.128 |
| Ap4e1 | 0.052 | 0.081 | 0.070 |
| Wdr59 | 0.052 | 0.031 | 0.027 |
| Zfp53 | 0.052 | 0.042 | 0.049 |
| Mipep | 0.052 | 0.049 | 0.057 |
| Ncapd2 | 0.051 | 0.033 | 0.041 |
| Wfdc3 | 0.051 | 0.008 | 0.004 |
| Itgb3 | 0.051 | 0.177 | 0.111 |
| 1700008J07R | 0.051 | 0.022 | 0.034 |
| Nol10 | 0.051 | 0.046 | 0.043 |
| Ubxn2b | 0.051 | 0.075 | 0.079 |
| Nudt6 | 0.051 | 0.043 | 0.039 |
| Rnasel | 0.051 | 0.054 | 0.126 |
| 2900093K20F | 0.051 | 0.038 | 0.037 |
| Fsd1l | 0.051 | 0.033 | 0.048 |
| Polr3b | 0.051 | 0.044 | 0.042 |
| Ddx28 | 0.051 | 0.055 | 0.054 |
| Hdac9 | 0.051 | 0.061 | 0.029 |
| Grk3 | 0.051 | 0.175 | 0.095 |
| Rbbp5 | 0.051 | 0.044 | 0.041 |
| Foxo4 | 0.051 | 0.057 | 0.061 |
| Mettl8 | 0.051 | 0.028 | 0.037 |
| Gja1 | 0.051 | 0.268 | 0.166 |
| Bcar3 | 0.051 | 0.128 | 0.065 |
| Nmral1 | 0.051 | 0.060 | 0.057 |
| Dock10 | 0.051 | 0.051 | 0.188 |
| Ice2 | 0.051 | 0.052 | 0.059 |
| Sgpp1 | 0.051 | 0.090 | 0.154 |
| Klhdc8a | 0.051 | 0.001 | 0.000 |
| Gm30524 | 0.051 | 0.010 | 0.015 |
| Gm10638 | 0.051 | 0.046 | 0.100 |
| Arhgap42 | 0.051 | 0.056 | 0.105 |
| Tpmt | 0.051 | 0.076 | 0.067 |
| Mast3 | 0.051 | 0.048 | 0.038 |
| Zfp60 | 0.051 | 0.047 | 0.048 |
| Myom1 | 0.051 | 0.368 | 0.070 |
| Rbm19 | 0.051 | 0.038 | 0.051 |
| Napb | 0.051 | 0.037 | 0.030 |
| Mgme1 | 0.051 | 0.037 | 0.055 |
| Ogg1 | 0.051 | 0.039 | 0.040 |

| | | | |
|---|---|---|---|
| Gatb | 0.051 | 0.029 | 0.036 |
| Zc3h10 | 0.051 | 0.065 | 0.056 |
| Satb1 | 0.051 | 0.066 | 0.066 |
| Usp30 | 0.051 | 0.060 | 0.054 |
| Spred3 | 0.051 | 0.058 | 0.041 |
| Snapc4 | 0.051 | 0.060 | 0.066 |
| Tinf2 | 0.051 | 0.040 | 0.034 |
| Maf | 0.051 | 0.334 | 0.494 |
| Pitpnm1 | 0.051 | 0.068 | 0.029 |
| Lgals3 | 0.051 | 0.079 | 0.033 |
| Stard5 | 0.050 | 0.039 | 0.044 |
| Dph7 | 0.050 | 0.045 | 0.045 |
| Traf1 | 0.050 | 0.011 | 0.018 |
| Bud13 | 0.050 | 0.046 | 0.048 |
| Tab3 | 0.050 | 0.038 | 0.031 |
| Heatr1 | 0.050 | 0.038 | 0.044 |
| Fgf2 | 0.050 | 0.203 | 0.137 |
| Fam35a | 0.050 | 0.043 | 0.037 |
| Ttbk2 | 0.050 | 0.065 | 0.078 |
| Ttc27 | 0.050 | 0.048 | 0.045 |
| C030014I23R | 0.050 | 0.051 | 0.062 |
| Zfp867 | 0.050 | 0.046 | 0.051 |
| Zfp316 | 0.050 | 0.065 | 0.074 |
| Tchp | 0.050 | 0.050 | 0.053 |
| Parp11 | 0.050 | 0.046 | 0.052 |
| Tmed8 | 0.050 | 0.042 | 0.028 |
| Incenp | 0.050 | 0.041 | 0.047 |
| Spata5 | 0.050 | 0.075 | 0.069 |
| Mier2 | 0.050 | 0.069 | 0.079 |
| Ttc30b | 0.050 | 0.043 | 0.059 |
| Wdyhv1 | 0.050 | 0.073 | 0.068 |
| Osgepl1 | 0.050 | 0.060 | 0.062 |
| D11Wsu47e | 0.050 | 0.041 | 0.042 |
| Tpx2 | 0.050 | 0.042 | 0.038 |
| Nrtn | 0.050 | 0.383 | 0.348 |
| Ptgr1 | 0.050 | 0.136 | 0.148 |
| Hells | 0.050 | 0.042 | 0.060 |
| Ighmbp2 | 0.050 | 0.043 | 0.041 |
| Ddx51 | 0.050 | 0.029 | 0.029 |
| 5033428I22R | 0.050 | 0.028 | 0.059 |
| Lnx2 | 0.050 | 0.085 | 0.064 |
| Recql | 0.050 | 0.054 | 0.063 |
| Gm9889 | 0.050 | 0.003 | 0.004 |
| Fmn2 | 0.050 | 0.004 | 0.007 |
| Mon1b | 0.050 | 0.031 | 0.028 |
| Sp140 | 0.050 | 0.040 | 0.052 |
| Znrf3 | 0.050 | 0.077 | 0.037 |
| Rwdd2a | 0.050 | 0.020 | 0.019 |

| | | | |
|---|---|---|---|
| Tcf19 | 0.050 | 0.051 | 0.072 |
| Tnfrsf1b | 0.050 | 0.022 | 0.034 |
| Gm14295 | 0.050 | 0.060 | 0.064 |
| Pecr | 0.050 | 0.105 | 0.104 |
| E2f1 | 0.050 | 0.051 | 0.118 |
| Pomgnt2 | 0.050 | 0.052 | 0.061 |
| Gli2 | 0.050 | 0.081 | 0.104 |
| Prdm15 | 0.050 | 0.043 | 0.053 |
| Ripk3 | 0.050 | 0.030 | 0.038 |
| Rai2 | 0.049 | 0.013 | 0.026 |
| Mettl22 | 0.049 | 0.060 | 0.055 |
| Ip6k3 | 0.049 | 0.000 | 0.000 |
| Gm26601 | 0.049 | 0.044 | 0.045 |
| Ttf2 | 0.049 | 0.032 | 0.030 |
| Azin2 | 0.049 | 0.152 | 0.114 |
| Trim59 | 0.049 | 0.065 | 0.030 |
| Hs6st2 | 0.049 | 0.090 | 0.090 |
| Sh2d3c | 0.049 | 0.030 | 0.033 |
| Pglyrp1 | 0.049 | 0.009 | 0.013 |
| Fam122b | 0.049 | 0.031 | 0.055 |
| Dtwd1 | 0.049 | 0.065 | 0.103 |
| Mad2l1 | 0.049 | 0.031 | 0.031 |
| Hectd2 | 0.049 | 0.142 | 0.081 |
| Alyref2 | 0.049 | 0.054 | 0.069 |
| Knstrn | 0.049 | 0.035 | 0.029 |
| Fam214a | 0.049 | 0.092 | 0.080 |
| Alg8 | 0.049 | 0.033 | 0.042 |
| Diaph3 | 0.049 | 0.024 | 0.034 |
| Bbs2 | 0.049 | 0.040 | 0.044 |
| Ankrd52 | 0.049 | 0.059 | 0.048 |
| Tbc1d32 | 0.049 | 0.043 | 0.051 |
| Psd3 | 0.049 | 0.070 | 0.146 |
| Zbed5 | 0.049 | 0.045 | 0.053 |
| Dcaf12l1 | 0.049 | 0.015 | 0.031 |
| Zfp449 | 0.049 | 0.047 | 0.097 |
| Zfp280c | 0.049 | 0.048 | 0.042 |
| Rnf165 | 0.049 | 0.012 | 0.111 |
| 2700046G09I | 0.049 | 0.033 | 0.027 |
| Cbx8 | 0.049 | 0.049 | 0.048 |
| Ntf3 | 0.049 | 0.376 | 0.054 |
| Osgin2 | 0.049 | 0.082 | 0.066 |
| Trim68 | 0.049 | 0.035 | 0.036 |
| Zfp870 | 0.049 | 0.033 | 0.037 |
| Klhl25 | 0.049 | 0.052 | 0.043 |
| Lpl | 0.049 | 0.116 | 0.518 |
| Fam149a | 0.049 | 0.052 | 0.092 |
| Taf1c | 0.049 | 0.035 | 0.033 |
| Traf3ip1 | 0.049 | 0.050 | 0.041 |

| | | | |
|---|---|---|---|
| Dhrs13 | 0.049 | 0.052 | 0.061 |
| Eral1 | 0.048 | 0.037 | 0.032 |
| Cep135 | 0.048 | 0.036 | 0.043 |
| Zfp958 | 0.048 | 0.036 | 0.042 |
| Pole2 | 0.048 | 0.040 | 0.060 |
| Irgm2 | 0.048 | 0.022 | 0.036 |
| AU020206 | 0.048 | 0.055 | 0.062 |
| Sgsm3 | 0.048 | 0.034 | 0.041 |
| Ikzf4 | 0.048 | 0.301 | 0.203 |
| Nlgn2 | 0.048 | 0.129 | 0.187 |
| 4930428N03I | 0.048 | 0.000 | 0.001 |
| Pcgf1 | 0.048 | 0.048 | 0.059 |
| Gemin6 | 0.048 | 0.048 | 0.051 |
| Tnnt1 | 0.048 | 0.150 | 0.164 |
| Gm37305 | 0.048 | 0.029 | 0.038 |
| Zfp984 | 0.048 | 0.040 | 0.064 |
| Lrrc40 | 0.048 | 0.052 | 0.059 |
| Tstd3 | 0.048 | 0.035 | 0.035 |
| 3632451O06I | 0.048 | 0.075 | 0.327 |
| Pcbd1 | 0.048 | 0.032 | 0.030 |
| Zfp799 | 0.048 | 0.035 | 0.040 |
| Zfp729b | 0.048 | 0.046 | 0.046 |
| Ccdc126 | 0.048 | 0.053 | 0.048 |
| Inhbb | 0.048 | 0.020 | 0.009 |
| Rita1 | 0.048 | 0.054 | 0.073 |
| Gm17056 | 0.048 | 0.322 | 0.350 |
| Car5b | 0.048 | 0.024 | 0.017 |
| BC051226 | 0.048 | 0.041 | 0.041 |
| Abca2 | 0.048 | 0.061 | 0.053 |
| Plekhm3 | 0.048 | 0.064 | 0.049 |
| Tlcd1 | 0.048 | 0.129 | 0.171 |
| Zfp105 | 0.048 | 0.049 | 0.049 |
| Fam69b | 0.048 | 0.158 | 0.187 |
| Sema3c | 0.048 | 0.282 | 0.472 |
| Evc2 | 0.048 | 0.116 | 0.174 |
| Shroom2 | 0.048 | 0.012 | 0.011 |
| Ppcdc | 0.048 | 0.068 | 0.070 |
| Egfl8 | 0.048 | 0.027 | 0.029 |
| Phka1 | 0.048 | 0.053 | 0.043 |
| Dhrs11 | 0.048 | 0.046 | 0.031 |
| Zfp426 | 0.048 | 0.072 | 0.082 |
| 9330020H09I | 0.048 | 0.042 | 0.031 |
| 4930452B06F | 0.048 | 0.005 | 0.003 |
| Zfp40 | 0.048 | 0.042 | 0.051 |
| Primpol | 0.048 | 0.030 | 0.045 |
| Cep70 | 0.047 | 0.050 | 0.029 |
| S1pr4 | 0.047 | 0.000 | 0.000 |
| Nbeal2 | 0.047 | 0.005 | 0.003 |

| | | | |
|---|---|---|---|
| Tcaim | 0.047 | 0.039 | 0.031 |
| Nup107 | 0.047 | 0.039 | 0.044 |
| Slc36a1 | 0.047 | 0.027 | 0.043 |
| Vipr2 | 0.047 | 0.012 | 0.030 |
| Usp49 | 0.047 | 0.034 | 0.031 |
| Rab40b | 0.047 | 0.101 | 0.116 |
| Timmdc1 | 0.047 | 0.044 | 0.044 |
| Ebf4 | 0.047 | 0.080 | 0.060 |
| Ggct | 0.047 | 0.024 | 0.032 |
| Dusp28 | 0.047 | 0.044 | 0.047 |
| Mapkapk3 | 0.047 | 0.027 | 0.020 |
| Foxp2 | 0.047 | 0.552 | 0.528 |
| Brdt | 0.047 | 0.057 | 0.074 |
| Zfp992 | 0.047 | 0.031 | 0.056 |
| Sco1 | 0.047 | 0.033 | 0.035 |
| Mmaa | 0.047 | 0.060 | 0.054 |
| Hic2 | 0.047 | 0.051 | 0.056 |
| Trit1 | 0.047 | 0.049 | 0.049 |
| 4933427D14I | 0.047 | 0.041 | 0.034 |
| Lca5 | 0.047 | 0.055 | 0.083 |
| Twnk | 0.047 | 0.047 | 0.044 |
| Chac2 | 0.047 | 0.048 | 0.044 |
| Supv3l1 | 0.047 | 0.051 | 0.054 |
| Armc9 | 0.047 | 0.091 | 0.087 |
| Gm14964 | 0.047 | 0.029 | 0.020 |
| Mettl25 | 0.047 | 0.045 | 0.029 |
| Zfp976 | 0.047 | 0.036 | 0.037 |
| Nipa1 | 0.047 | 0.077 | 0.085 |
| Prim1 | 0.047 | 0.072 | 0.078 |
| Col25a1 | 0.047 | 0.028 | 0.010 |
| Mettl17 | 0.047 | 0.039 | 0.033 |
| Pcdhb22 | 0.047 | 0.039 | 0.041 |
| Lrfn4 | 0.047 | 0.121 | 0.073 |
| Atl1 | 0.047 | 0.039 | 0.044 |
| Mterf2 | 0.047 | 0.047 | 0.035 |
| AW146154 | 0.047 | 0.029 | 0.030 |
| Vgll3 | 0.047 | 0.197 | 0.110 |
| Tecpr2 | 0.047 | 0.047 | 0.047 |
| Rfx5 | 0.047 | 0.043 | 0.036 |
| Tspyl3 | 0.047 | 0.028 | 0.038 |
| Pramef8 | 0.047 | 0.057 | 0.085 |
| Ormdl1 | 0.046 | 0.059 | 0.041 |
| Pola1 | 0.046 | 0.044 | 0.054 |
| Trmt5 | 0.046 | 0.029 | 0.038 |
| Ston1 | 0.046 | 0.026 | 0.032 |
| Unc13d | 0.046 | 0.008 | 0.005 |
| Fam131a | 0.046 | 0.093 | 0.060 |
| Rogdi | 0.046 | 0.065 | 0.052 |

| | | | |
|---|---|---|---|
| Gprasp2 | 0.046 | 0.096 | 0.129 |
| Fgd6 | 0.046 | 0.032 | 0.028 |
| Dnmbp | 0.046 | 0.057 | 0.069 |
| Mcub | 0.046 | 0.051 | 0.042 |
| Mrm3 | 0.046 | 0.041 | 0.055 |
| Sox8 | 0.046 | 0.015 | 0.013 |
| Ccdc166 | 0.046 | 0.020 | 0.025 |
| Noc3l | 0.046 | 0.067 | 0.059 |
| 2810454H06I | 0.046 | 0.040 | 0.043 |
| Stk17b | 0.046 | 0.116 | 0.097 |
| Ddx20 | 0.046 | 0.051 | 0.052 |
| Fam229b | 0.046 | 0.049 | 0.063 |
| Gga3 | 0.046 | 0.037 | 0.046 |
| Tmem254a | 0.046 | 0.026 | 0.040 |
| Prkdc | 0.046 | 0.033 | 0.044 |
| Pced1b | 0.046 | 0.026 | 0.018 |
| Ccdc163 | 0.046 | 0.070 | 0.063 |
| Ereg | 0.046 | 0.054 | 0.019 |
| Tpk1 | 0.046 | 0.026 | 0.025 |
| Ddx11 | 0.046 | 0.012 | 0.014 |
| Ppip5k1 | 0.046 | 0.047 | 0.034 |
| Cdca7 | 0.046 | 0.055 | 0.054 |
| Rrad | 0.046 | 0.538 | 0.267 |
| Ap5z1 | 0.046 | 0.031 | 0.032 |
| Tubgcp6 | 0.046 | 0.032 | 0.039 |
| Halr1 | 0.046 | 0.001 | 0.003 |
| Ttc9 | 0.046 | 0.004 | 0.006 |
| Neo1 | 0.046 | 0.372 | 0.225 |
| Tarsl2 | 0.046 | 0.032 | 0.024 |
| Tti1 | 0.046 | 0.043 | 0.044 |
| Wnt11 | 0.046 | 0.005 | 0.053 |
| Prdm6 | 0.046 | 0.819 | 0.379 |
| Pgap1 | 0.046 | 0.167 | 0.079 |
| Rwdd2b | 0.046 | 0.025 | 0.025 |
| Fut10 | 0.046 | 0.077 | 0.099 |
| Cdkn3 | 0.046 | 0.024 | 0.023 |
| Cdh24 | 0.046 | 0.055 | 0.034 |
| Tepsin | 0.046 | 0.029 | 0.026 |
| Gstm4 | 0.046 | 0.043 | 0.065 |
| Shroom1 | 0.046 | 0.012 | 0.013 |
| Pik3cb | 0.046 | 0.022 | 0.008 |
| Lcp1 | 0.046 | 0.010 | 0.011 |
| Spred2 | 0.046 | 0.111 | 0.125 |
| Wdr76 | 0.046 | 0.021 | 0.017 |
| Rfx3 | 0.046 | 0.054 | 0.047 |
| Zfp429 | 0.045 | 0.029 | 0.038 |
| Fdxr | 0.045 | 0.040 | 0.102 |
| Hdhd3 | 0.045 | 0.048 | 0.054 |

| | | | |
|---|---|---|---|
| Atad5 | 0.045 | 0.035 | 0.038 |
| Atp10a | 0.045 | 0.153 | 0.099 |
| Spry2 | 0.045 | 0.229 | 0.164 |
| Snx29 | 0.045 | 0.058 | 0.065 |
| Hgh1 | 0.045 | 0.032 | 0.031 |
| Zhx2 | 0.045 | 0.027 | 0.039 |
| Mettl18 | 0.045 | 0.025 | 0.026 |
| 1700086O06I | 0.045 | 0.035 | 0.050 |
| Tpcn2 | 0.045 | 0.057 | 0.060 |
| Tcf15 | 0.045 | 0.001 | 0.002 |
| Igtp | 0.045 | 0.030 | 0.037 |
| A430105I19R | 0.045 | 0.049 | 0.011 |
| Olfm2 | 0.045 | 0.004 | 0.043 |
| Zfp286 | 0.045 | 0.058 | 0.060 |
| Sars2 | 0.045 | 0.029 | 0.034 |
| Gm35167 | 0.045 | 0.000 | 0.001 |
| Zswim9 | 0.045 | 0.050 | 0.057 |
| 4930522L14F | 0.045 | 0.037 | 0.029 |
| Dhx35 | 0.045 | 0.030 | 0.025 |
| Aldoc | 0.045 | 0.048 | 0.006 |
| Map7 | 0.045 | 0.031 | 0.016 |
| Llgl2 | 0.045 | 0.021 | 0.011 |
| Lyn | 0.045 | 0.018 | 0.023 |
| Lactb2 | 0.045 | 0.052 | 0.048 |
| Rarb | 0.045 | 0.066 | 0.102 |
| AC164424.1 | 0.045 | 0.039 | 0.042 |
| Rbm38 | 0.045 | 0.021 | 0.010 |
| Wdr81 | 0.045 | 0.039 | 0.044 |
| Zfp839 | 0.045 | 0.057 | 0.044 |
| D2hgdh | 0.045 | 0.034 | 0.046 |
| Pcdhb20 | 0.045 | 0.033 | 0.034 |
| Porcn | 0.045 | 0.064 | 0.049 |
| Zc3h11a | 0.045 | 0.036 | 0.038 |
| Efs | 0.045 | 0.260 | 0.392 |
| Mmab | 0.045 | 0.051 | 0.068 |
| Mfhas1 | 0.045 | 0.207 | 0.209 |
| Zfp975 | 0.045 | 0.037 | 0.042 |
| Pigo | 0.045 | 0.034 | 0.037 |
| Gm42715 | 0.045 | 0.039 | 0.041 |
| Coq8a | 0.045 | 0.085 | 0.071 |
| Slc38a7 | 0.045 | 0.056 | 0.060 |
| Zc3h12c | 0.045 | 0.043 | 0.035 |
| Wrap53 | 0.045 | 0.035 | 0.040 |
| Exo5 | 0.045 | 0.036 | 0.043 |
| Man2b2 | 0.044 | 0.023 | 0.052 |
| Fuk | 0.044 | 0.042 | 0.028 |
| Zbtb7b | 0.044 | 0.081 | 0.048 |
| 2410131K14F | 0.044 | 0.052 | 0.045 |

| | | | |
|---|---|---|---|
| Tbc1d30 | 0.044 | 0.003 | 0.001 |
| Fdxacb1 | 0.044 | 0.040 | 0.039 |
| Sergef | 0.044 | 0.077 | 0.078 |
| Rasgrp2 | 0.044 | 0.277 | 0.075 |
| Socs1 | 0.044 | 0.240 | 0.438 |
| Gm6710 | 0.044 | 0.076 | 0.085 |
| Ttc21b | 0.044 | 0.056 | 0.064 |
| Plppr2 | 0.044 | 0.031 | 0.052 |
| Ccdc148 | 0.044 | 0.010 | 0.008 |
| Krt80 | 0.044 | 0.007 | 0.009 |
| Tm6sf2 | 0.044 | 0.003 | 0.002 |
| Pxmp2 | 0.044 | 0.022 | 0.009 |
| Serp2 | 0.044 | 0.124 | 0.117 |
| Dzip1l | 0.044 | 0.161 | 0.112 |
| 3-Mar | 0.044 | 0.022 | 0.022 |
| Tspan11 | 0.044 | 0.036 | 0.307 |
| Kit | 0.044 | 0.010 | 0.012 |
| Dnajc16 | 0.044 | 0.034 | 0.027 |
| Gtf2ird2 | 0.044 | 0.040 | 0.050 |
| A530017D24I | 0.044 | 0.045 | 0.036 |
| Manba | 0.044 | 0.037 | 0.034 |
| Ctu1 | 0.044 | 0.021 | 0.030 |
| Armc6 | 0.044 | 0.043 | 0.063 |
| Mctp1 | 0.044 | 0.022 | 0.005 |
| Ago4 | 0.044 | 0.075 | 0.067 |
| Bid | 0.044 | 0.014 | 0.010 |
| Siah1b | 0.044 | 0.036 | 0.028 |
| Prr16 | 0.044 | 0.006 | 0.013 |
| Zfp748 | 0.044 | 0.023 | 0.030 |
| Fmo1 | 0.044 | 0.185 | 0.640 |
| Dnm3os | 0.044 | 0.344 | 0.600 |
| Plxnc1 | 0.044 | 0.003 | 0.011 |
| Wdr7 | 0.044 | 0.043 | 0.044 |
| A430033K04I | 0.044 | 0.028 | 0.032 |
| Anln | 0.044 | 0.060 | 0.033 |
| Rbp2 | 0.044 | 0.002 | 0.006 |
| Coq3 | 0.044 | 0.053 | 0.040 |
| Trac | 0.044 | 0.206 | 0.378 |
| Mrm1 | 0.044 | 0.025 | 0.034 |
| E2f2 | 0.044 | 0.004 | 0.005 |
| Zfp367 | 0.044 | 0.053 | 0.030 |
| Bora | 0.044 | 0.053 | 0.041 |
| Gm17354 | 0.044 | 0.052 | 0.051 |
| Entpd6 | 0.044 | 0.055 | 0.082 |
| Cfap43 | 0.044 | 0.065 | 0.057 |
| C1ra | 0.044 | 0.062 | 0.271 |
| 1600014C10I | 0.044 | 0.046 | 0.041 |
| Tut1 | 0.044 | 0.038 | 0.037 |

| | | | |
|---|---|---|---|
| Pigg | 0.044 | 0.024 | 0.025 |
| Gm13111 | 0.044 | 0.066 | 0.009 |
| Usp20 | 0.044 | 0.075 | 0.039 |
| Daglb | 0.044 | 0.047 | 0.115 |
| Zfp446 | 0.044 | 0.045 | 0.054 |
| Fbxo5 | 0.043 | 0.027 | 0.028 |
| Lrrc17 | 0.043 | 0.290 | 0.893 |
| Tmem143 | 0.043 | 0.039 | 0.041 |
| Ankrd13b | 0.043 | 0.083 | 0.080 |
| Miga2 | 0.043 | 0.049 | 0.061 |
| Atp1a2 | 0.043 | 0.484 | 0.521 |
| Eml5 | 0.043 | 0.029 | 0.014 |
| Cped1 | 0.043 | 0.239 | 0.444 |
| Pde1c | 0.043 | 0.003 | 0.005 |
| Bbs4 | 0.043 | 0.062 | 0.056 |
| Zfp248 | 0.043 | 0.037 | 0.036 |
| Zfp719 | 0.043 | 0.065 | 0.085 |
| Tsga10 | 0.043 | 0.058 | 0.057 |
| Nr2f2 | 0.043 | 0.326 | 0.468 |
| Mtrf1 | 0.043 | 0.051 | 0.043 |
| Stx11 | 0.043 | 0.045 | 0.057 |
| Oxsm | 0.043 | 0.018 | 0.027 |
| Abcc4 | 0.043 | 0.005 | 0.009 |
| Gdf10 | 0.043 | 0.310 | 0.726 |
| Mgmt | 0.043 | 0.072 | 0.099 |
| Speg | 0.043 | 0.490 | 0.123 |
| Cyb5d2 | 0.043 | 0.024 | 0.038 |
| Qtrt2 | 0.043 | 0.046 | 0.040 |
| Zbtb39 | 0.043 | 0.036 | 0.027 |
| Fam124b | 0.043 | 0.000 | 0.000 |
| Grap | 0.043 | 0.007 | 0.012 |
| Recql5 | 0.043 | 0.030 | 0.028 |
| Ulk3 | 0.043 | 0.039 | 0.035 |
| Donson | 0.043 | 0.027 | 0.035 |
| Tiam1 | 0.043 | 0.035 | 0.172 |
| Dhdh | 0.043 | 0.023 | 0.032 |
| Vwa8 | 0.043 | 0.067 | 0.037 |
| Tulp1 | 0.043 | 0.127 | 0.111 |
| Sox9 | 0.043 | 0.535 | 0.477 |
| Tspyl4 | 0.043 | 0.049 | 0.049 |
| Fam46c | 0.043 | 0.036 | 0.041 |
| Tcea2 | 0.043 | 0.026 | 0.025 |
| Rdh5 | 0.043 | 0.066 | 0.048 |
| Ckap2 | 0.043 | 0.033 | 0.035 |
| Prrx2 | 0.043 | 0.526 | 0.398 |
| Adal | 0.043 | 0.040 | 0.058 |
| Haus6 | 0.043 | 0.041 | 0.041 |
| Ankrd29 | 0.043 | 0.075 | 0.047 |

| | | | |
|---|---|---|---|
| Pced1a | 0.042 | 0.041 | 0.081 |
| Cdyl2 | 0.042 | 0.040 | 0.027 |
| Tnfrsf21 | 0.042 | 0.167 | 0.033 |
| Ccpg1os | 0.042 | 0.032 | 0.038 |
| Metrn | 0.042 | 0.071 | 0.072 |
| Hddc3 | 0.042 | 0.040 | 0.055 |
| Tenm3 | 0.042 | 0.518 | 0.383 |
| Tldc1 | 0.042 | 0.027 | 0.034 |
| Arhgap20 | 0.042 | 0.005 | 0.161 |
| Bloc1s3 | 0.042 | 0.028 | 0.023 |
| Zfp946 | 0.042 | 0.041 | 0.042 |
| Gstcd | 0.042 | 0.025 | 0.039 |
| Mblac1 | 0.042 | 0.028 | 0.029 |
| Klf12 | 0.042 | 0.074 | 0.069 |
| Zfp65 | 0.042 | 0.033 | 0.039 |
| Tprn | 0.042 | 0.019 | 0.015 |
| Mmd | 0.042 | 0.183 | 0.058 |
| Mb21d2 | 0.042 | 0.081 | 0.055 |
| Grk4 | 0.042 | 0.058 | 0.047 |
| Chml | 0.042 | 0.057 | 0.032 |
| Rab43 | 0.042 | 0.075 | 0.071 |
| Helb | 0.042 | 0.067 | 0.070 |
| Zmat1 | 0.042 | 0.062 | 0.068 |
| Trim46 | 0.042 | 0.026 | 0.033 |
| Eya1 | 0.042 | 0.001 | 0.001 |
| Plxna4os1 | 0.042 | 0.000 | 0.001 |
| Gm28523 | 0.042 | 0.019 | 0.019 |
| Igsf9b | 0.042 | 0.327 | 0.051 |
| Zfp442 | 0.042 | 0.034 | 0.054 |
| 4732440D04I | 0.042 | 0.045 | 0.055 |
| Blm | 0.042 | 0.032 | 0.040 |
| Hspb7 | 0.042 | 0.430 | 0.119 |
| Mtx3 | 0.042 | 0.035 | 0.042 |
| Zbtb26 | 0.042 | 0.024 | 0.024 |
| Nepro | 0.042 | 0.038 | 0.041 |
| Etfbkmt | 0.042 | 0.055 | 0.062 |
| Popdc3 | 0.042 | 0.032 | 0.004 |
| Gm16124 | 0.042 | 0.061 | 0.035 |
| Zfp35 | 0.042 | 0.034 | 0.040 |
| Dennd4b | 0.042 | 0.025 | 0.044 |
| Zfp626 | 0.042 | 0.039 | 0.039 |
| Wnt5b | 0.042 | 0.184 | 0.287 |
| Fgfr2 | 0.042 | 0.394 | 0.114 |
| Fam71e1 | 0.042 | 0.025 | 0.023 |
| Fcer1g | 0.042 | 0.032 | 0.030 |
| 1810062G17I | 0.042 | 0.038 | 0.039 |
| Thrb | 0.042 | 0.267 | 0.100 |
| Syn2 | 0.041 | 0.014 | 0.005 |

| | | | |
|---|---|---|---|
| Pkp2 | 0.041 | 0.001 | 0.001 |
| Tac2 | 0.041 | 0.003 | 0.002 |
| Dcdc2a | 0.041 | 0.001 | 0.005 |
| Ppl | 0.041 | 0.012 | 0.021 |
| Mccc2 | 0.041 | 0.088 | 0.070 |
| Ddx59 | 0.041 | 0.036 | 0.036 |
| Plp1 | 0.041 | 0.038 | 0.036 |
| Cdk5rap1 | 0.041 | 0.036 | 0.033 |
| Card6 | 0.041 | 0.013 | 0.012 |
| Bche | 0.041 | 0.261 | 0.083 |
| Slc25a22 | 0.041 | 0.049 | 0.041 |
| Slc25a29 | 0.041 | 0.024 | 0.027 |
| Ncs1 | 0.041 | 0.151 | 0.128 |
| Pomk | 0.041 | 0.041 | 0.068 |
| Gabre | 0.041 | 0.111 | 0.096 |
| Gm26890 | 0.041 | 0.053 | 0.049 |
| Pcdh1 | 0.041 | 0.003 | 0.003 |
| Gab3 | 0.041 | 0.024 | 0.045 |
| Ckap2l | 0.041 | 0.015 | 0.015 |
| Pth1r | 0.041 | 0.277 | 0.392 |
| Mthfd1l | 0.041 | 0.052 | 0.092 |
| Bex1 | 0.041 | 0.018 | 0.018 |
| Naaa | 0.041 | 0.091 | 0.166 |
| Dusp2 | 0.041 | 0.029 | 0.014 |
| Gdf6 | 0.041 | 0.004 | 0.026 |
| Rab26os | 0.041 | 0.056 | 0.045 |
| Nmrk1 | 0.041 | 0.041 | 0.062 |
| Rcor2 | 0.041 | 0.042 | 0.033 |
| Slc35f1 | 0.041 | 0.015 | 0.004 |
| BC037034 | 0.041 | 0.032 | 0.039 |
| Lrrk2 | 0.041 | 0.070 | 0.059 |
| Zscan2 | 0.041 | 0.034 | 0.029 |
| Srxn1 | 0.041 | 0.053 | 0.093 |
| Prepl | 0.041 | 0.039 | 0.032 |
| Elac1 | 0.041 | 0.029 | 0.030 |
| Tex9 | 0.041 | 0.069 | 0.061 |
| Thsd4 | 0.041 | 0.587 | 0.127 |
| BC034090 | 0.041 | 0.040 | 0.041 |
| Fam13c | 0.041 | 0.280 | 0.089 |
| Cit | 0.041 | 0.011 | 0.017 |
| Usp2 | 0.041 | 0.138 | 0.063 |
| Rad18 | 0.041 | 0.052 | 0.054 |
| Ift140 | 0.041 | 0.045 | 0.051 |
| Rffl | 0.041 | 0.049 | 0.033 |
| Acan | 0.041 | 0.636 | 0.079 |
| Sfmbt2 | 0.041 | 0.020 | 0.014 |
| Galnt16 | 0.041 | 0.257 | 0.462 |
| Polr1b | 0.041 | 0.039 | 0.035 |

| | | | |
|---|---|---|---|
| P4htm | 0.041 | 0.120 | 0.118 |
| Rapgef4 | 0.041 | 0.012 | 0.008 |
| Agfg2 | 0.041 | 0.108 | 0.085 |
| Cwf19l1 | 0.041 | 0.027 | 0.027 |
| Scrn3 | 0.041 | 0.054 | 0.053 |
| Thnsl1 | 0.041 | 0.022 | 0.029 |
| Gdap10 | 0.041 | 0.058 | 0.064 |
| Zfp408 | 0.041 | 0.034 | 0.033 |
| Lynx1 | 0.041 | 0.027 | 0.063 |
| Rtel1 | 0.041 | 0.034 | 0.036 |
| Fastkd5 | 0.041 | 0.025 | 0.026 |
| Trim45 | 0.041 | 0.048 | 0.063 |
| Alms1 | 0.041 | 0.062 | 0.059 |
| Col11a1 | 0.041 | 0.038 | 0.199 |
| Alkbh4 | 0.041 | 0.025 | 0.022 |
| C1qa | 0.040 | 0.052 | 0.052 |
| Lcmt2 | 0.040 | 0.062 | 0.071 |
| Gm26778 | 0.040 | 0.006 | 0.015 |
| Ulbp1 | 0.040 | 0.004 | 0.008 |
| Wdr5b | 0.040 | 0.030 | 0.030 |
| Pqlc2 | 0.040 | 0.036 | 0.036 |
| Ttc26 | 0.040 | 0.023 | 0.027 |
| Rfx2 | 0.040 | 0.040 | 0.036 |
| Zfp995 | 0.040 | 0.034 | 0.050 |
| Arhgef10l | 0.040 | 0.101 | 0.094 |
| Als2 | 0.040 | 0.031 | 0.031 |
| Lrrc20 | 0.040 | 0.032 | 0.036 |
| Zfp61 | 0.040 | 0.071 | 0.070 |
| Tnfrsf11a | 0.040 | 0.010 | 0.002 |
| Fpgt | 0.040 | 0.027 | 0.030 |
| Itpk1 | 0.040 | 0.107 | 0.040 |
| Rbp7 | 0.040 | 0.003 | 0.004 |
| Evc | 0.040 | 0.086 | 0.157 |
| Itga4 | 0.040 | 0.187 | 0.032 |
| Usp31 | 0.040 | 0.034 | 0.030 |
| St8sia2 | 0.040 | 0.082 | 0.141 |
| 2210408I21R | 0.040 | 0.044 | 0.050 |
| Magi2 | 0.040 | 0.097 | 0.103 |
| Snrnp35 | 0.040 | 0.033 | 0.048 |
| Hmga1 | 0.040 | 0.040 | 0.020 |
| Orai2 | 0.040 | 0.024 | 0.019 |
| Rab32 | 0.040 | 0.068 | 0.081 |
| Tmem154 | 0.040 | 0.015 | 0.007 |
| Tmem267 | 0.040 | 0.049 | 0.062 |
| Fam171b | 0.040 | 0.163 | 0.208 |
| Dnmt3b | 0.040 | 0.025 | 0.024 |
| C1qtnf9 | 0.040 | 0.001 | 0.002 |
| U2af1l4 | 0.040 | 0.040 | 0.031 |

| | | | |
|---|---|---|---|
| Epg5 | 0.040 | 0.036 | 0.030 |
| Ddx31 | 0.040 | 0.024 | 0.031 |
| Htatip2 | 0.040 | 0.029 | 0.039 |
| Gm11266 | 0.040 | 0.030 | 0.045 |
| 1700017B05F | 0.040 | 0.176 | 0.139 |
| Thnsl2 | 0.040 | 0.051 | 0.077 |
| Rab30 | 0.040 | 0.043 | 0.096 |
| Pcdhb14 | 0.040 | 0.027 | 0.028 |
| Suv39h1 | 0.040 | 0.057 | 0.055 |
| Klhl8 | 0.040 | 0.046 | 0.088 |
| Cdk20 | 0.040 | 0.030 | 0.064 |
| Trim6 | 0.040 | 0.013 | 0.020 |
| Morc4 | 0.039 | 0.065 | 0.144 |
| Zfp111 | 0.039 | 0.038 | 0.047 |
| Sass6 | 0.039 | 0.030 | 0.031 |
| Hlx | 0.039 | 0.042 | 0.085 |
| Cbx7 | 0.039 | 0.023 | 0.014 |
| 1700030K09F | 0.039 | 0.038 | 0.046 |
| Dars2 | 0.039 | 0.026 | 0.026 |
| Abca8a | 0.039 | 0.037 | 0.517 |
| Ndufaf5 | 0.039 | 0.040 | 0.045 |
| Espn | 0.039 | 0.012 | 0.018 |
| BC002059 | 0.039 | 0.029 | 0.034 |
| Tasp1 | 0.039 | 0.051 | 0.056 |
| Rbm12b2 | 0.039 | 0.040 | 0.040 |
| Lrrc75b | 0.039 | 0.115 | 0.033 |
| Bdnf | 0.039 | 0.272 | 0.122 |
| Rbl1 | 0.039 | 0.019 | 0.025 |
| Tubd1 | 0.039 | 0.018 | 0.027 |
| Gm16503 | 0.039 | 0.017 | 0.022 |
| Gpr161 | 0.039 | 0.067 | 0.070 |
| Ubiad1 | 0.039 | 0.034 | 0.031 |
| E430018J23R | 0.039 | 0.036 | 0.046 |
| Zscan22 | 0.039 | 0.020 | 0.022 |
| Pcdhb7 | 0.039 | 0.019 | 0.024 |
| Tigd5 | 0.039 | 0.024 | 0.029 |
| Slc19a1 | 0.039 | 0.076 | 0.057 |
| Clec11a | 0.039 | 0.050 | 0.143 |
| Kif3c | 0.039 | 0.048 | 0.034 |
| Wnt2 | 0.039 | 0.002 | 0.083 |
| Tnfrsf22 | 0.039 | 0.010 | 0.005 |
| Nrxn2 | 0.039 | 0.019 | 0.038 |
| Acy1 | 0.039 | 0.037 | 0.071 |
| Rnf24 | 0.039 | 0.062 | 0.075 |
| Mcph1 | 0.039 | 0.032 | 0.028 |
| Atp8a2 | 0.039 | 0.003 | 0.002 |
| Mtrr | 0.039 | 0.040 | 0.055 |
| Atmin | 0.039 | 0.040 | 0.043 |

| | | | |
|---|---|---|---|
| Fancf | 0.039 | 0.030 | 0.048 |
| Adnp | 0.039 | 0.042 | 0.035 |
| Uevld | 0.039 | 0.038 | 0.035 |
| Cdt1 | 0.039 | 0.027 | 0.038 |
| Zfp982 | 0.039 | 0.013 | 0.016 |
| Tbc1d25 | 0.039 | 0.030 | 0.026 |
| Nckipsd | 0.039 | 0.030 | 0.039 |
| Igfbp6 | 0.039 | 0.066 | 0.301 |
| Lca5l | 0.039 | 0.079 | 0.075 |
| Gpsm2 | 0.039 | 0.137 | 0.214 |
| Ccdc138 | 0.039 | 0.060 | 0.078 |
| Kbtbd7 | 0.039 | 0.043 | 0.041 |
| 3010003L21F | 0.039 | 0.030 | 0.024 |
| Zfp758 | 0.039 | 0.042 | 0.045 |
| Fah | 0.039 | 0.039 | 0.033 |
| Nupl2 | 0.039 | 0.033 | 0.029 |
| Eno3 | 0.039 | 0.075 | 0.056 |
| Hist1h2ae | 0.039 | 0.037 | 0.027 |
| Tuft1 | 0.039 | 0.005 | 0.006 |
| Spa17 | 0.039 | 0.031 | 0.026 |
| Echdc2 | 0.039 | 0.256 | 0.194 |
| Lonrf1 | 0.039 | 0.083 | 0.119 |
| Ssh3 | 0.039 | 0.043 | 0.022 |
| Lins1 | 0.039 | 0.032 | 0.036 |
| Gnpda1 | 0.038 | 0.050 | 0.053 |
| Zfp69 | 0.038 | 0.003 | 0.006 |
| Zfp874b | 0.038 | 0.045 | 0.047 |
| Ptges3l | 0.038 | 0.203 | 0.095 |
| Smcr8 | 0.038 | 0.043 | 0.044 |
| Klhl23 | 0.038 | 0.189 | 0.105 |
| Pou5f2 | 0.038 | 0.046 | 0.051 |
| Ube2v1 | 0.038 | 0.032 | 0.030 |
| Evi5l | 0.038 | 0.045 | 0.075 |
| Lyrm1 | 0.038 | 0.037 | 0.029 |
| Galm | 0.038 | 0.052 | 0.081 |
| Akr1c14 | 0.038 | 0.032 | 0.094 |
| Zfp775 | 0.038 | 0.046 | 0.043 |
| Gsdme | 0.038 | 0.327 | 0.312 |
| Prox1 | 0.038 | 0.011 | 0.005 |
| Dclre1a | 0.038 | 0.064 | 0.067 |
| Spc25 | 0.038 | 0.024 | 0.033 |
| Tmem53 | 0.038 | 0.119 | 0.131 |
| C1qb | 0.038 | 0.054 | 0.054 |
| Sp100 | 0.038 | 0.005 | 0.026 |
| 9930012K11F | 0.038 | 0.006 | 0.015 |
| Gm14005 | 0.038 | 0.006 | 0.008 |
| Nat14 | 0.038 | 0.073 | 0.029 |
| Wdr34 | 0.038 | 0.067 | 0.077 |

| | | | |
|---|---|---|---|
| Disc1 | 0.038 | 0.015 | 0.027 |
| Veph1 | 0.038 | 0.000 | 0.002 |
| Lin52 | 0.038 | 0.037 | 0.030 |
| Adcy5 | 0.038 | 0.484 | 0.108 |
| Zfp933 | 0.038 | 0.035 | 0.034 |
| Actc1 | 0.038 | 0.047 | 0.046 |
| Tk1 | 0.038 | 0.027 | 0.030 |
| Upp1 | 0.038 | 0.002 | 0.001 |
| Flywch2 | 0.038 | 0.065 | 0.143 |
| Nxt2 | 0.038 | 0.095 | 0.057 |
| Clcn6 | 0.038 | 0.037 | 0.033 |
| Pclo | 0.038 | 0.053 | 0.021 |
| Mpv17l | 0.038 | 0.067 | 0.044 |
| Rac3 | 0.038 | 0.112 | 0.141 |
| 4732491K20F | 0.038 | 0.034 | 0.032 |
| Zfp182 | 0.038 | 0.039 | 0.042 |
| Cldn12 | 0.038 | 0.045 | 0.046 |
| Cngb1 | 0.038 | 0.009 | 0.011 |
| Srgap3 | 0.038 | 0.108 | 0.094 |
| Scn3b | 0.038 | 0.009 | 0.026 |
| Lcn2 | 0.038 | 0.031 | 0.027 |
| Msln | 0.038 | 0.018 | 0.024 |
| Pde10a | 0.038 | 0.348 | 0.077 |
| Ears2 | 0.038 | 0.059 | 0.047 |
| Hk2 | 0.038 | 0.260 | 0.273 |
| Robo2 | 0.038 | 0.007 | 0.072 |
| Stard8 | 0.038 | 0.040 | 0.035 |
| Plpp7 | 0.038 | 0.044 | 0.043 |
| Arhgap25 | 0.037 | 0.001 | 0.001 |
| Ccp110 | 0.037 | 0.053 | 0.051 |
| Mms22l | 0.037 | 0.023 | 0.033 |
| Armc7 | 0.037 | 0.020 | 0.017 |
| Runx1t1 | 0.037 | 0.228 | 0.360 |
| Tnnc1 | 0.037 | 0.138 | 0.040 |
| Zfp763 | 0.037 | 0.033 | 0.037 |
| Meox2 | 0.037 | 0.013 | 0.159 |
| Jph2 | 0.037 | 0.268 | 0.079 |
| Sytl4 | 0.037 | 0.014 | 0.021 |
| Pank4 | 0.037 | 0.039 | 0.039 |
| 9530036M11 | 0.037 | 0.003 | 0.003 |
| Pdzrn3 | 0.037 | 0.448 | 0.269 |
| Nat2 | 0.037 | 0.019 | 0.015 |
| Itpripl1 | 0.037 | 0.037 | 0.026 |
| Wdr35 | 0.037 | 0.094 | 0.109 |
| Plac9b | 0.037 | 0.044 | 0.108 |
| Dlg3 | 0.037 | 0.079 | 0.057 |
| Zmym6 | 0.037 | 0.028 | 0.041 |
| Rbm4 | 0.037 | 0.031 | 0.029 |

| | | | |
|---|---|---|---|
| Otud7a | 0.037 | 0.002 | 0.004 |
| Smoc2 | 0.037 | 0.304 | 0.766 |
| Vps25 | 0.037 | 0.022 | 0.020 |
| Pltp | 0.037 | 0.037 | 0.149 |
| Dck | 0.037 | 0.025 | 0.020 |
| Ppp3cc | 0.037 | 0.066 | 0.062 |
| Gm26624 | 0.037 | 0.007 | 0.010 |
| Atp5s | 0.037 | 0.035 | 0.040 |
| Ttll5 | 0.037 | 0.069 | 0.123 |
| Casz1 | 0.037 | 0.038 | 0.009 |
| Vwa1 | 0.037 | 0.167 | 0.053 |
| Has2 | 0.037 | 0.020 | 0.037 |
| Tmem170 | 0.037 | 0.024 | 0.020 |
| 2700038G22I | 0.037 | 0.034 | 0.040 |
| Rab4b | 0.037 | 0.033 | 0.031 |
| Prss23 | 0.037 | 0.147 | 0.331 |
| Arrdc2 | 0.037 | 0.047 | 0.039 |
| Sec22c | 0.037 | 0.026 | 0.037 |
| Zfp7 | 0.037 | 0.026 | 0.023 |
| Pemt | 0.037 | 0.020 | 0.026 |
| 1110002L01F | 0.037 | 0.031 | 0.030 |
| Dennd2c | 0.037 | 0.033 | 0.022 |
| H2-Ab1 | 0.037 | 0.013 | 0.005 |
| D6Wsu163e | 0.037 | 0.041 | 0.044 |
| Pfas | 0.037 | 0.039 | 0.036 |
| Oxnad1 | 0.037 | 0.109 | 0.174 |
| Cerkl | 0.037 | 0.007 | 0.004 |
| Chrna1os | 0.037 | 0.066 | 0.062 |
| Ralgps1 | 0.037 | 0.046 | 0.028 |
| Slc22a18 | 0.037 | 0.004 | 0.017 |
| Cmtr1 | 0.037 | 0.039 | 0.033 |
| BC022687 | 0.037 | 0.076 | 0.038 |
| Gm13212 | 0.037 | 0.018 | 0.041 |
| Xpo5 | 0.037 | 0.029 | 0.028 |
| Dynlt1c | 0.037 | 0.031 | 0.037 |
| Tspan17 | 0.036 | 0.083 | 0.069 |
| Cep95 | 0.036 | 0.041 | 0.063 |
| Smpd5 | 0.036 | 0.009 | 0.011 |
| Zc3h8 | 0.036 | 0.034 | 0.033 |
| Trmt44 | 0.036 | 0.026 | 0.021 |
| Bex2 | 0.036 | 0.061 | 0.036 |
| Acy3 | 0.036 | 0.039 | 0.045 |
| Rdh13 | 0.036 | 0.052 | 0.051 |
| Usp11 | 0.036 | 0.046 | 0.079 |
| Dand5 | 0.036 | 0.046 | 0.046 |
| Ccdc28a | 0.036 | 0.037 | 0.052 |
| Atg4c | 0.036 | 0.041 | 0.041 |
| Lmtk2 | 0.036 | 0.022 | 0.021 |

| | | | |
|---|---|---|---|
| Col4a3 | 0.036 | 0.010 | 0.010 |
| Spice1 | 0.036 | 0.049 | 0.074 |
| Hhat | 0.036 | 0.055 | 0.066 |
| Zc3h12a | 0.036 | 0.061 | 0.052 |
| Pcsk2 | 0.036 | 0.004 | 0.005 |
| Abcb7 | 0.036 | 0.028 | 0.030 |
| Robo4 | 0.036 | 0.001 | 0.001 |
| 4930481A15F | 0.036 | 0.021 | 0.026 |
| Dmxl2 | 0.036 | 0.100 | 0.057 |
| Lpcat4 | 0.036 | 0.018 | 0.008 |
| Cenpq | 0.036 | 0.066 | 0.045 |
| Frat1 | 0.036 | 0.012 | 0.018 |
| Lrp4 | 0.036 | 0.030 | 0.064 |
| Mmp15 | 0.036 | 0.020 | 0.025 |
| N4bp2l1 | 0.036 | 0.088 | 0.067 |
| Dnaaf5 | 0.036 | 0.025 | 0.018 |
| D3Ertd751e | 0.036 | 0.025 | 0.048 |
| Aff3 | 0.036 | 0.076 | 0.195 |
| 9930104L06F | 0.036 | 0.038 | 0.032 |
| Pcgf6 | 0.036 | 0.053 | 0.042 |
| Sepsecs | 0.036 | 0.035 | 0.039 |
| Edrf1 | 0.036 | 0.059 | 0.054 |
| Phtf2 | 0.036 | 0.200 | 0.062 |
| Slc25a15 | 0.036 | 0.032 | 0.048 |
| Kif21a | 0.036 | 0.025 | 0.027 |
| Cfap45 | 0.036 | 0.019 | 0.015 |
| Wdr19 | 0.036 | 0.039 | 0.040 |
| 4930455G09I | 0.036 | 0.001 | 0.001 |
| Map9 | 0.036 | 0.058 | 0.052 |
| E2f7 | 0.036 | 0.011 | 0.021 |
| Hivep3 | 0.036 | 0.037 | 0.120 |
| Abca7 | 0.035 | 0.034 | 0.047 |
| Marveld2 | 0.035 | 0.001 | 0.003 |
| Dnal1 | 0.035 | 0.050 | 0.044 |
| Pawr | 0.035 | 0.266 | 0.058 |
| Mypop | 0.035 | 0.070 | 0.080 |
| Nek4 | 0.035 | 0.070 | 0.061 |
| Rtn1 | 0.035 | 0.009 | 0.028 |
| Nnt | 0.035 | 0.036 | 0.032 |
| Zfp930 | 0.035 | 0.028 | 0.032 |
| Arxes1 | 0.035 | 0.093 | 0.126 |
| Rasd2 | 0.035 | 0.056 | 0.028 |
| Xkr8 | 0.035 | 0.025 | 0.036 |
| Adamtsl1 | 0.035 | 0.163 | 0.264 |
| Trpt1 | 0.035 | 0.043 | 0.033 |
| Aif1l | 0.035 | 0.005 | 0.014 |
| Rhobtb2 | 0.035 | 0.038 | 0.034 |
| AW551984 | 0.035 | 0.010 | 0.188 |

| | | | |
|---|---|---|---|
| Zbtb40 | 0.035 | 0.021 | 0.023 |
| Clspn | 0.035 | 0.032 | 0.038 |
| Rrm2 | 0.035 | 0.038 | 0.038 |
| Tmem184c | 0.035 | 0.131 | 0.192 |
| Crocc | 0.035 | 0.035 | 0.032 |
| Gm45871 | 0.035 | 0.031 | 0.036 |
| Fam167a | 0.035 | 0.013 | 0.035 |
| Itga2 | 0.035 | 0.030 | 0.010 |
| Tep1 | 0.035 | 0.052 | 0.061 |
| Suox | 0.035 | 0.040 | 0.034 |
| Smyd4 | 0.035 | 0.031 | 0.037 |
| Clca3a1 | 0.035 | 0.198 | 0.176 |
| Nemp2 | 0.035 | 0.026 | 0.038 |
| Igsf23 | 0.035 | 0.005 | 0.005 |
| Gm4285 | 0.035 | 0.030 | 0.049 |
| Zdhhc24 | 0.035 | 0.024 | 0.030 |
| Ttll4 | 0.035 | 0.035 | 0.044 |
| Plac8 | 0.035 | 0.169 | 0.596 |
| Slc2a13 | 0.035 | 0.084 | 0.067 |
| Lamb3 | 0.035 | 0.005 | 0.009 |
| 4930430F08R | 0.035 | 0.045 | 0.038 |
| Ccnb1 | 0.035 | 0.026 | 0.025 |
| Glb1l | 0.035 | 0.073 | 0.088 |
| Tnfrsf10b | 0.035 | 0.033 | 0.012 |
| Lurap1 | 0.035 | 0.033 | 0.007 |
| C920006O11I | 0.035 | 0.040 | 0.039 |
| Gm11423 | 0.035 | 0.019 | 0.024 |
| Wdr27 | 0.035 | 0.026 | 0.022 |
| Pard6g | 0.035 | 0.158 | 0.147 |
| Flrt3 | 0.034 | 0.303 | 0.123 |
| Ttc38 | 0.034 | 0.052 | 0.073 |
| Adamts4 | 0.034 | 0.148 | 0.186 |
| Nkapl | 0.034 | 0.019 | 0.015 |
| Pygl | 0.034 | 0.100 | 0.102 |
| Ppp1r3c | 0.034 | 0.068 | 0.076 |
| Rbm41 | 0.034 | 0.036 | 0.033 |
| Ttc12 | 0.034 | 0.024 | 0.021 |
| Tspyl5 | 0.034 | 0.039 | 0.037 |
| Pot1b | 0.034 | 0.024 | 0.021 |
| Ccna2 | 0.034 | 0.026 | 0.028 |
| Nars2 | 0.034 | 0.039 | 0.030 |
| Slc40a1 | 0.034 | 0.010 | 0.042 |
| 5033430I15R | 0.034 | 0.036 | 0.021 |
| Lig4 | 0.034 | 0.031 | 0.028 |
| D130040H23I | 0.034 | 0.022 | 0.030 |
| Cobll1 | 0.034 | 0.174 | 0.032 |
| Ramp3 | 0.034 | 0.005 | 0.005 |
| D830025C05I | 0.034 | 0.065 | 0.080 |

| | | | |
|---|---|---|---|
| Nr6a1 | 0.034 | 0.017 | 0.016 |
| Abcc9 | 0.034 | 0.543 | 0.447 |
| Paqr7 | 0.034 | 0.037 | 0.019 |
| Fam187b | 0.034 | 0.001 | 0.001 |
| Tmem191c | 0.034 | 0.053 | 0.032 |
| Aim2 | 0.034 | 0.001 | 0.002 |
| Ybey | 0.034 | 0.031 | 0.027 |
| Slc41a2 | 0.034 | 0.110 | 0.044 |
| Zfp217 | 0.034 | 0.017 | 0.053 |
| Mapk13 | 0.034 | 0.020 | 0.030 |
| Hist1h4d | 0.034 | 0.043 | 0.031 |
| Gm2415 | 0.034 | 0.031 | 0.031 |
| 9530082P21R | 0.034 | 0.013 | 0.018 |
| Dlx6os1 | 0.034 | 0.015 | 0.012 |
| 5930430L01R | 0.034 | 0.061 | 0.036 |
| Efcab2 | 0.034 | 0.022 | 0.039 |
| Olfr920 | 0.034 | 0.072 | 0.101 |
| Fam102a | 0.034 | 0.060 | 0.105 |
| Pex10 | 0.034 | 0.037 | 0.043 |
| Abraxas1 | 0.034 | 0.055 | 0.052 |
| 2700033N17R | 0.034 | 0.027 | 0.053 |
| Cenpo | 0.034 | 0.017 | 0.028 |
| Gm48529 | 0.034 | 0.044 | 0.012 |
| Bahcc1 | 0.034 | 0.029 | 0.036 |
| Lmln | 0.034 | 0.024 | 0.021 |
| 2610035D17R | 0.034 | 0.029 | 0.033 |
| Gm47101 | 0.034 | 0.034 | 0.013 |
| Dnajb5 | 0.034 | 0.044 | 0.025 |
| Tyrobp | 0.034 | 0.031 | 0.030 |
| Raet1d | 0.034 | 0.002 | 0.009 |
| Arrb2 | 0.034 | 0.032 | 0.045 |
| Haus5 | 0.034 | 0.022 | 0.028 |
| Tarbp1 | 0.034 | 0.017 | 0.018 |
| Pcbp3 | 0.034 | 0.028 | 0.029 |
| Gm5141 | 0.034 | 0.023 | 0.019 |
| Pmaip1 | 0.034 | 0.004 | 0.004 |
| Gm29107 | 0.033 | 0.023 | 0.010 |
| Tha1 | 0.033 | 0.013 | 0.016 |
| 1190005I06R | 0.033 | 0.284 | 0.046 |
| Dhcr24 | 0.033 | 0.051 | 0.073 |
| Rab38 | 0.033 | 0.012 | 0.008 |
| Gramd1c | 0.033 | 0.026 | 0.034 |
| Cep112 | 0.033 | 0.069 | 0.096 |
| Jmjd7 | 0.033 | 0.045 | 0.049 |
| Mcrip2 | 0.033 | 0.058 | 0.050 |
| Cdh2 | 0.033 | 0.156 | 0.143 |
| Kctd13 | 0.033 | 0.064 | 0.063 |
| Pus7l | 0.033 | 0.029 | 0.035 |

| | | | |
|---|---|---|---|
| Zfp874a | 0.033 | 0.036 | 0.044 |
| Ninl | 0.033 | 0.049 | 0.031 |
| Zfp229 | 0.033 | 0.030 | 0.039 |
| Wdr90 | 0.033 | 0.033 | 0.033 |
| Aldh5a1 | 0.033 | 0.026 | 0.028 |
| Gm17586 | 0.033 | 0.036 | 0.041 |
| Vamp1 | 0.033 | 0.016 | 0.018 |
| Intu | 0.033 | 0.016 | 0.019 |
| Hist1h4h | 0.033 | 0.042 | 0.018 |
| AC163032.1 | 0.033 | 0.036 | 0.020 |
| Pdp2 | 0.033 | 0.055 | 0.045 |
| C530008M17 | 0.033 | 0.058 | 0.025 |
| Prag1 | 0.033 | 0.081 | 0.057 |
| Col5a3 | 0.033 | 0.020 | 0.411 |
| Rbm45 | 0.033 | 0.026 | 0.026 |
| Slc16a9 | 0.033 | 0.027 | 0.034 |
| Mlph | 0.033 | 0.009 | 0.005 |
| 9330160F10F | 0.033 | 0.032 | 0.031 |
| Nap1l3 | 0.033 | 0.026 | 0.021 |
| Pex11a | 0.033 | 0.042 | 0.020 |
| Prkar2b | 0.033 | 0.066 | 0.051 |
| Cep131 | 0.033 | 0.037 | 0.049 |
| Noct | 0.033 | 0.065 | 0.048 |
| Snai1 | 0.033 | 0.175 | 0.222 |
| Gm12184 | 0.033 | 0.052 | 0.047 |
| Kif22 | 0.033 | 0.023 | 0.024 |
| Bbs9 | 0.033 | 0.045 | 0.042 |
| Xkr6 | 0.033 | 0.017 | 0.017 |
| Col6a6 | 0.033 | 0.169 | 0.697 |
| Mttp | 0.033 | 0.036 | 0.078 |
| Rasd1 | 0.033 | 0.090 | 0.028 |
| Zfp324 | 0.033 | 0.025 | 0.035 |
| Gm42047 | 0.033 | 0.020 | 0.022 |
| Col9a2 | 0.033 | 0.046 | 0.147 |
| Tiam2 | 0.033 | 0.045 | 0.076 |
| Lbp | 0.033 | 0.004 | 0.080 |
| Rpp38 | 0.033 | 0.039 | 0.048 |
| Poc1a | 0.033 | 0.028 | 0.039 |
| Swsap1 | 0.033 | 0.018 | 0.022 |
| Zdhhc15 | 0.033 | 0.030 | 0.041 |
| 4833418N02I | 0.033 | 0.035 | 0.038 |
| C030034I22R | 0.032 | 0.015 | 0.021 |
| Stk32b | 0.032 | 0.075 | 0.034 |
| Tead4 | 0.032 | 0.002 | 0.002 |
| Slc35e3 | 0.032 | 0.039 | 0.055 |
| Rbks | 0.032 | 0.005 | 0.005 |
| C230035I16R | 0.032 | 0.014 | 0.015 |
| Ugt8a | 0.032 | 0.003 | 0.003 |

| | | | |
|---|---|---|---|
| Agbl5 | 0.032 | 0.028 | 0.020 |
| Eri2 | 0.032 | 0.032 | 0.038 |
| Gm11361 | 0.032 | 0.030 | 0.030 |
| Renbp | 0.032 | 0.066 | 0.127 |
| Hmmr | 0.032 | 0.018 | 0.017 |
| Dapk2 | 0.032 | 0.023 | 0.018 |
| Adamtsl2 | 0.032 | 0.082 | 0.163 |
| Rcc1 | 0.032 | 0.036 | 0.053 |
| Rgs16 | 0.032 | 0.035 | 0.115 |
| Olfml1 | 0.032 | 0.089 | 0.181 |
| Hemk1 | 0.032 | 0.030 | 0.045 |
| Fjx1 | 0.032 | 0.267 | 0.109 |
| Gm9951 | 0.032 | 0.018 | 0.002 |
| 2810408A11F | 0.032 | 0.016 | 0.024 |
| Lck | 0.032 | 0.008 | 0.004 |
| Zfp566 | 0.032 | 0.029 | 0.034 |
| Dhx37 | 0.032 | 0.021 | 0.022 |
| Dph2 | 0.032 | 0.017 | 0.021 |
| Hook1 | 0.032 | 0.002 | 0.001 |
| Rnf113a1 | 0.032 | 0.029 | 0.024 |
| Zbtb25 | 0.032 | 0.066 | 0.078 |
| Gm34030 | 0.032 | 0.017 | 0.068 |
| Kirrel3 | 0.032 | 0.005 | 0.007 |
| E330009J07R | 0.032 | 0.022 | 0.021 |
| Fam234b | 0.032 | 0.031 | 0.039 |
| Zfp961 | 0.032 | 0.022 | 0.023 |
| Cenpj | 0.032 | 0.024 | 0.023 |
| Nfkbie | 0.032 | 0.021 | 0.028 |
| Tshz3 | 0.032 | 0.091 | 0.155 |
| Gm10419 | 0.032 | 0.002 | 0.002 |
| Jade3 | 0.032 | 0.019 | 0.012 |
| Usp35 | 0.032 | 0.035 | 0.034 |
| Pcdh17 | 0.032 | 0.007 | 0.018 |
| Gm20045 | 0.032 | 0.031 | 0.026 |
| 3222401L13F | 0.032 | 0.037 | 0.035 |
| Sema4b | 0.032 | 0.061 | 0.034 |
| Klhl11 | 0.032 | 0.030 | 0.025 |
| Cd27 | 0.032 | 0.009 | 0.018 |
| Efna2 | 0.032 | 0.071 | 0.117 |
| Btc | 0.032 | 0.709 | 0.201 |
| Mfsd12 | 0.032 | 0.041 | 0.047 |
| Nfatc2ip | 0.032 | 0.038 | 0.021 |
| Chd3os | 0.032 | 0.043 | 0.051 |
| Mettl24 | 0.032 | 0.040 | 0.014 |
| Fmnl2 | 0.032 | 0.096 | 0.153 |
| Tsku | 0.032 | 0.028 | 0.084 |
| Zscan18 | 0.032 | 0.026 | 0.025 |
| Gp9 | 0.032 | 0.003 | 0.002 |

| | | | |
|---|---|---|---|
| Scara5 | 0.032 | 0.129 | 0.358 |
| Neu3 | 0.032 | 0.027 | 0.020 |
| Exog | 0.032 | 0.026 | 0.036 |
| Arhgap39 | 0.032 | 0.042 | 0.041 |
| Ppp1r3b | 0.032 | 0.039 | 0.016 |
| Lrrc51 | 0.032 | 0.018 | 0.028 |
| Zfp991 | 0.032 | 0.024 | 0.037 |
| Tmem8b | 0.032 | 0.072 | 0.050 |
| Nudt11 | 0.032 | 0.026 | 0.016 |
| Hebp2 | 0.032 | 0.068 | 0.083 |
| Cenps | 0.032 | 0.019 | 0.036 |
| Plxnb1 | 0.031 | 0.086 | 0.030 |
| Txnl4b | 0.031 | 0.016 | 0.022 |
| Sh3bp1 | 0.031 | 0.107 | 0.139 |
| H2-Eb1 | 0.031 | 0.008 | 0.013 |
| Foxm1 | 0.031 | 0.018 | 0.020 |
| Tedc2 | 0.031 | 0.027 | 0.028 |
| Neil1 | 0.031 | 0.018 | 0.017 |
| Mfsd3 | 0.031 | 0.031 | 0.040 |
| Flvcr1 | 0.031 | 0.049 | 0.035 |
| Apbb1ip | 0.031 | 0.035 | 0.165 |
| Il1rap | 0.031 | 0.085 | 0.046 |
| Gcat | 0.031 | 0.051 | 0.082 |
| AA986860 | 0.031 | 0.007 | 0.009 |
| Slc18a2 | 0.031 | 0.003 | 0.003 |
| Fads2 | 0.031 | 0.032 | 0.027 |
| Chmp4c | 0.031 | 0.075 | 0.007 |
| Sorcs2 | 0.031 | 0.096 | 0.275 |
| Zfp937 | 0.031 | 0.020 | 0.032 |
| Zfp58 | 0.031 | 0.026 | 0.032 |
| Dus4l | 0.031 | 0.025 | 0.026 |
| Gdf11 | 0.031 | 0.047 | 0.057 |
| Trabd2b | 0.031 | 0.203 | 0.128 |
| Rassf7 | 0.031 | 0.016 | 0.007 |
| Htra4 | 0.031 | 0.042 | 0.016 |
| Dglucy | 0.031 | 0.010 | 0.009 |
| Gm5113 | 0.031 | 0.022 | 0.022 |
| Dnajc12 | 0.031 | 0.072 | 0.034 |
| Rgs7bp | 0.031 | 0.251 | 0.051 |
| Usp28 | 0.031 | 0.058 | 0.041 |
| Lrsam1 | 0.031 | 0.033 | 0.045 |
| Tspan8 | 0.031 | 0.001 | 0.001 |
| Hoxd8 | 0.031 | 0.001 | 0.006 |
| Bmper | 0.031 | 0.242 | 0.139 |
| 1700037C18F | 0.031 | 0.026 | 0.021 |
| Steap3 | 0.031 | 0.047 | 0.208 |
| Ncapg2 | 0.031 | 0.035 | 0.039 |
| Lama3 | 0.031 | 0.017 | 0.008 |

| | | | |
|---|---|---|---|
| Dtwd2 | 0.031 | 0.025 | 0.029 |
| Pid1 | 0.031 | 0.051 | 0.590 |
| Sbspon | 0.031 | 0.062 | 0.007 |
| Loxl3 | 0.031 | 0.134 | 0.209 |
| Cdkl3 | 0.031 | 0.037 | 0.050 |
| Nr1h3 | 0.031 | 0.018 | 0.026 |
| Galnt6 | 0.031 | 0.001 | 0.001 |
| Herc3 | 0.031 | 0.064 | 0.019 |
| Pfkfb2 | 0.031 | 0.018 | 0.018 |
| Ncald | 0.031 | 0.303 | 0.244 |
| Prss12 | 0.031 | 0.147 | 0.427 |
| Polm | 0.031 | 0.031 | 0.029 |
| Akap17b | 0.031 | 0.098 | 0.043 |
| Zfp27 | 0.031 | 0.036 | 0.046 |
| Exoc3l2 | 0.031 | 0.001 | 0.004 |
| Umad1 | 0.031 | 0.044 | 0.051 |
| Smim24 | 0.031 | 0.012 | 0.029 |
| Gipc2 | 0.031 | 0.052 | 0.013 |
| Zfp52 | 0.031 | 0.030 | 0.022 |
| Cenpk | 0.031 | 0.036 | 0.039 |
| Uros | 0.031 | 0.031 | 0.028 |
| Mrnip | 0.031 | 0.023 | 0.029 |
| Epha3 | 0.031 | 0.176 | 0.538 |
| Det1 | 0.031 | 0.040 | 0.043 |
| Pank1 | 0.031 | 0.064 | 0.044 |
| Ranbp17 | 0.031 | 0.033 | 0.023 |
| Ccdc15 | 0.031 | 0.018 | 0.029 |
| Cep295nl | 0.031 | 0.020 | 0.023 |
| Gm14326 | 0.031 | 0.023 | 0.025 |
| Thg1l | 0.030 | 0.039 | 0.031 |
| Wdpcp | 0.030 | 0.035 | 0.040 |
| Gins3 | 0.030 | 0.012 | 0.015 |
| Neurl3 | 0.030 | 0.009 | 0.017 |
| Aldh4a1 | 0.030 | 0.029 | 0.016 |
| Nat8f1 | 0.030 | 0.026 | 0.022 |
| Gm16867 | 0.030 | 0.014 | 0.013 |
| Slc25a33 | 0.030 | 0.046 | 0.030 |
| Casp1 | 0.030 | 0.002 | 0.001 |
| Unc5c | 0.030 | 0.235 | 0.103 |
| Aasdh | 0.030 | 0.023 | 0.025 |
| Tead3 | 0.030 | 0.219 | 0.134 |
| Mtcp1 | 0.030 | 0.020 | 0.023 |
| Zfp692 | 0.030 | 0.030 | 0.029 |
| Gm45902 | 0.030 | 0.019 | 0.022 |
| Tfpi2 | 0.030 | 0.006 | 0.076 |
| Ptpdc1 | 0.030 | 0.061 | 0.101 |
| Ddb2 | 0.030 | 0.040 | 0.033 |
| A430046D13I | 0.030 | 0.062 | 0.031 |

| | | | |
|---|---|---|---|
| Top3a | 0.030 | 0.026 | 0.031 |
| Otud3 | 0.030 | 0.031 | 0.033 |
| Kif11 | 0.030 | 0.019 | 0.020 |
| Tmem26 | 0.030 | 0.499 | 0.337 |
| Sned1 | 0.030 | 0.076 | 0.263 |
| Psrc1 | 0.030 | 0.016 | 0.016 |
| Rilp | 0.030 | 0.026 | 0.013 |
| Slc7a5 | 0.030 | 0.044 | 0.024 |
| Brca2 | 0.030 | 0.047 | 0.065 |
| Mocs3 | 0.030 | 0.040 | 0.042 |
| Dhx34 | 0.030 | 0.042 | 0.057 |
| Enpp2 | 0.030 | 0.168 | 0.187 |
| Ccnj | 0.030 | 0.020 | 0.017 |
| Cep76 | 0.030 | 0.045 | 0.028 |
| Kyat1 | 0.030 | 0.023 | 0.028 |
| Gm8113 | 0.030 | 0.011 | 0.007 |
| Gm4673 | 0.030 | 0.025 | 0.026 |
| Rtn2 | 0.030 | 0.114 | 0.126 |
| Olfr1033 | 0.030 | 0.150 | 0.101 |
| Pcdhb16 | 0.030 | 0.020 | 0.019 |
| Ret | 0.030 | 0.001 | 0.004 |
| Zfp595 | 0.030 | 0.022 | 0.028 |
| Aldh3a1 | 0.030 | 0.001 | 0.003 |
| Gm26669 | 0.030 | 0.128 | 0.139 |
| Fzd7 | 0.030 | 0.149 | 0.092 |
| Nat8l | 0.030 | 0.014 | 0.004 |
| Mal | 0.030 | 0.012 | 0.010 |
| Mipol1 | 0.030 | 0.043 | 0.042 |
| Enkd1 | 0.030 | 0.025 | 0.031 |
| Gm14403 | 0.030 | 0.018 | 0.030 |
| Morn4 | 0.030 | 0.024 | 0.016 |
| Prps1l3 | 0.030 | 0.018 | 0.023 |
| Engase | 0.030 | 0.016 | 0.033 |
| Ank3 | 0.029 | 0.040 | 0.173 |
| Ifit3 | 0.029 | 0.005 | 0.034 |
| Zfp963 | 0.029 | 0.020 | 0.024 |
| Frk | 0.029 | 0.104 | 0.058 |
| Slco1c1 | 0.029 | 0.027 | 0.010 |
| Ccdc51 | 0.029 | 0.029 | 0.022 |
| Rpgrip1l | 0.029 | 0.028 | 0.032 |
| Gm45353 | 0.029 | 0.006 | 0.007 |
| Gm4707 | 0.029 | 0.044 | 0.028 |
| Tnfaip8l1 | 0.029 | 0.007 | 0.007 |
| Zswim4 | 0.029 | 0.043 | 0.039 |
| Pou3f1 | 0.029 | 0.062 | 0.038 |
| Prodh | 0.029 | 0.012 | 0.006 |
| Prss36 | 0.029 | 0.028 | 0.042 |
| Ndrg4 | 0.029 | 0.086 | 0.105 |

| | | | |
|---|---|---|---|
| Ispd | 0.029 | 0.021 | 0.044 |
| Nutf2 | 0.029 | 0.031 | 0.030 |
| Zfhx4 | 0.029 | 0.020 | 0.156 |
| Ccdc134 | 0.029 | 0.050 | 0.052 |
| Fam177a | 0.029 | 0.038 | 0.023 |
| Lrrc56 | 0.029 | 0.019 | 0.024 |
| AC123870.1 | 0.029 | 0.024 | 0.021 |
| Flt3l | 0.029 | 0.031 | 0.065 |
| Ropn1l | 0.029 | 0.019 | 0.023 |
| Arhgef6 | 0.029 | 0.102 | 0.072 |
| Foxs1 | 0.029 | 0.182 | 0.203 |
| Fastkd1 | 0.029 | 0.026 | 0.022 |
| Mmp28 | 0.029 | 0.002 | 0.006 |
| Gm16845 | 0.029 | 0.027 | 0.021 |
| Rad51ap1 | 0.029 | 0.013 | 0.015 |
| Ccdc39 | 0.029 | 0.024 | 0.024 |
| C730034F03R | 0.029 | 0.041 | 0.051 |
| Mterf1a | 0.029 | 0.013 | 0.012 |
| Tgfa | 0.029 | 0.003 | 0.004 |
| AU041133 | 0.029 | 0.047 | 0.051 |
| Nrg1 | 0.029 | 0.003 | 0.015 |
| Abcb6 | 0.029 | 0.041 | 0.040 |
| Cenpl | 0.029 | 0.032 | 0.025 |
| Ptprz1 | 0.029 | 0.300 | 0.042 |
| Gm49085 | 0.029 | 0.052 | 0.037 |
| Erbb4 | 0.029 | 0.050 | 0.009 |
| Arsg | 0.029 | 0.044 | 0.053 |
| Gm10561 | 0.029 | 0.039 | 0.033 |
| Bbs5 | 0.029 | 0.043 | 0.038 |
| Angptl6 | 0.029 | 0.017 | 0.027 |
| Slc52a3 | 0.029 | 0.001 | 0.003 |
| Pctp | 0.029 | 0.003 | 0.006 |
| Tdrp | 0.029 | 0.010 | 0.008 |
| Uhrf1bp1 | 0.029 | 0.035 | 0.039 |
| Gm4631 | 0.029 | 0.034 | 0.041 |
| Serpina1e | 0.029 | 0.000 | 0.000 |
| Gm26881 | 0.029 | 0.018 | 0.035 |
| 2310068J16R | 0.029 | 0.031 | 0.022 |
| Pcdhgb4 | 0.029 | 0.031 | 0.030 |
| Skida1 | 0.029 | 0.023 | 0.022 |
| Prx | 0.029 | 0.042 | 0.038 |
| Gm12359 | 0.029 | 0.018 | 0.012 |
| Gm42670 | 0.029 | 0.011 | 0.012 |
| Crybg1 | 0.029 | 0.073 | 0.017 |
| 9430015G10I | 0.029 | 0.026 | 0.028 |
| Slc6a9 | 0.028 | 0.067 | 0.041 |
| Gas2 | 0.028 | 0.037 | 0.104 |
| Casq1 | 0.028 | 0.028 | 0.021 |

| | | | |
|---|---|---|---|
| Zfp974 | 0.028 | 0.020 | 0.023 |
| Rbm12b1 | 0.028 | 0.023 | 0.026 |
| Evpl | 0.028 | 0.010 | 0.008 |
| Nckap5 | 0.028 | 0.009 | 0.056 |
| Mycn | 0.028 | 0.002 | 0.001 |
| Ttpa | 0.028 | 0.011 | 0.006 |
| Upk3b | 0.028 | 0.018 | 0.030 |
| Sh3yl1 | 0.028 | 0.029 | 0.045 |
| Mecomos | 0.028 | 0.018 | 0.010 |
| Acss2 | 0.028 | 0.050 | 0.063 |
| Mpp2 | 0.028 | 0.062 | 0.027 |
| Bicdl1 | 0.028 | 0.033 | 0.042 |
| Cdc42bpg | 0.028 | 0.027 | 0.031 |
| A030001D20I | 0.028 | 0.009 | 0.011 |
| Syce2 | 0.028 | 0.031 | 0.050 |
| Dact1 | 0.028 | 0.235 | 0.461 |
| Cmpk2 | 0.028 | 0.015 | 0.030 |
| Chrd | 0.028 | 0.036 | 0.075 |
| Oxt | 0.028 | 0.003 | 0.004 |
| Adamts15 | 0.028 | 0.209 | 0.230 |
| Kif20a | 0.028 | 0.016 | 0.014 |
| Eepd1 | 0.028 | 0.043 | 0.046 |
| Cd36 | 0.028 | 0.009 | 0.005 |
| Tmco4 | 0.028 | 0.031 | 0.042 |
| Kif15 | 0.028 | 0.013 | 0.014 |
| Cryl1 | 0.028 | 0.064 | 0.087 |
| Gm16973 | 0.028 | 0.021 | 0.016 |
| Acrbp | 0.028 | 0.038 | 0.043 |
| Celf4 | 0.028 | 0.228 | 0.245 |
| Zcchc18 | 0.028 | 0.065 | 0.028 |
| Aox1 | 0.028 | 0.034 | 0.036 |
| BC024978 | 0.028 | 0.029 | 0.040 |
| B930036N10I | 0.028 | 0.028 | 0.042 |
| Ccne2 | 0.028 | 0.030 | 0.038 |
| Syn3 | 0.028 | 0.027 | 0.014 |
| Cyb5rl | 0.028 | 0.024 | 0.019 |
| 9130023H24I | 0.028 | 0.026 | 0.024 |
| Tmeff1 | 0.028 | 0.090 | 0.089 |
| Zfp647 | 0.028 | 0.028 | 0.033 |
| E430024I08Ri | 0.028 | 0.041 | 0.044 |
| Gm43086 | 0.028 | 0.000 | 0.001 |
| Gm10269 | 0.028 | 0.026 | 0.029 |
| Slc12a8 | 0.028 | 0.009 | 0.015 |
| Gm47547 | 0.028 | 0.011 | 0.005 |
| Tbxa2r | 0.028 | 0.213 | 0.231 |
| Cetn4 | 0.028 | 0.043 | 0.045 |
| Oscp1 | 0.028 | 0.066 | 0.072 |
| Gpr21 | 0.028 | 0.024 | 0.007 |

| | | | |
|---|---|---|---|
| Tmie | 0.028 | 0.006 | 0.002 |
| Trim21 | 0.028 | 0.018 | 0.021 |
| Prkcg | 0.028 | 0.054 | 0.047 |
| Gm1976 | 0.028 | 0.031 | 0.032 |
| Dclre1c | 0.028 | 0.018 | 0.019 |
| Lrmda | 0.028 | 0.001 | 0.006 |
| Gpr19 | 0.028 | 0.036 | 0.048 |
| Tubb3 | 0.028 | 0.007 | 0.011 |
| Gm11837 | 0.028 | 0.062 | 0.050 |
| Hdac11 | 0.028 | 0.128 | 0.069 |
| Epb41l3 | 0.028 | 0.161 | 0.447 |
| Ufsp1 | 0.028 | 0.029 | 0.012 |
| Dctd | 0.028 | 0.028 | 0.045 |
| Cenpv | 0.028 | 0.041 | 0.065 |
| Zfp661 | 0.028 | 0.005 | 0.006 |
| Slc43a2 | 0.028 | 0.014 | 0.007 |
| Ptpn3 | 0.028 | 0.057 | 0.041 |
| Lama2 | 0.028 | 0.355 | 0.663 |
| Fam120c | 0.028 | 0.030 | 0.024 |
| Kpna2 | 0.027 | 0.038 | 0.037 |
| Hoxb5 | 0.027 | 0.156 | 0.079 |
| Zfp575 | 0.027 | 0.029 | 0.095 |
| Adamts8 | 0.027 | 0.079 | 0.044 |
| Usp27x | 0.027 | 0.037 | 0.060 |
| F8 | 0.027 | 0.006 | 0.008 |
| 2410018L13F | 0.027 | 0.026 | 0.041 |
| Tnfrsf11b | 0.027 | 0.363 | 0.240 |
| Zfp202 | 0.027 | 0.012 | 0.016 |
| Akna | 0.027 | 0.016 | 0.028 |
| Cxadr | 0.027 | 0.013 | 0.007 |
| Rhebl1 | 0.027 | 0.034 | 0.031 |
| Parp16 | 0.027 | 0.024 | 0.024 |
| Podn | 0.027 | 0.085 | 0.116 |
| Simc1 | 0.027 | 0.068 | 0.061 |
| Commd5 | 0.027 | 0.018 | 0.016 |
| Ly75 | 0.027 | 0.031 | 0.003 |
| Gpatch3 | 0.027 | 0.018 | 0.016 |
| Xylt1 | 0.027 | 0.110 | 0.089 |
| Piga | 0.027 | 0.044 | 0.053 |
| Abcb1b | 0.027 | 0.062 | 0.038 |
| Tmem240 | 0.027 | 0.028 | 0.035 |
| Gm19426 | 0.027 | 0.024 | 0.039 |
| Vegfd | 0.027 | 0.118 | 0.418 |
| Zfp784 | 0.027 | 0.020 | 0.021 |
| Klc3 | 0.027 | 0.162 | 0.058 |
| Nusap1 | 0.027 | 0.026 | 0.028 |
| Dna2 | 0.027 | 0.018 | 0.014 |
| Fignl2 | 0.027 | 0.005 | 0.025 |

| | | | |
|---|---|---|---|
| Nek8 | 0.027 | 0.055 | 0.066 |
| Arv1 | 0.027 | 0.023 | 0.018 |
| Gm39469 | 0.027 | 0.016 | 0.017 |
| Mlf1 | 0.027 | 0.146 | 0.084 |
| Ager | 0.027 | 0.022 | 0.016 |
| Slc1a3 | 0.027 | 0.061 | 0.124 |
| Fxyd4 | 0.027 | 0.074 | 0.070 |
| Cnnm4 | 0.027 | 0.033 | 0.028 |
| Nmnat1 | 0.027 | 0.037 | 0.033 |
| C130013H08I | 0.027 | 0.013 | 0.013 |
| Ddx19b | 0.027 | 0.041 | 0.030 |
| Chst3 | 0.027 | 0.090 | 0.026 |
| 8030462N17I | 0.027 | 0.026 | 0.012 |
| Zfp28 | 0.027 | 0.034 | 0.034 |
| Gm21781 | 0.027 | 0.017 | 0.015 |
| Zfp280b | 0.027 | 0.025 | 0.024 |
| Adora2a | 0.027 | 0.024 | 0.064 |
| Ifi203 | 0.027 | 0.002 | 0.023 |
| Mis18bp1 | 0.027 | 0.015 | 0.018 |
| Zc4h2 | 0.027 | 0.035 | 0.033 |
| 1810044D09I | 0.027 | 0.010 | 0.017 |
| Gm43848 | 0.027 | 0.026 | 0.037 |
| Rnf152 | 0.027 | 0.052 | 0.056 |
| Hist1h2ac | 0.027 | 0.030 | 0.018 |
| Atp23 | 0.027 | 0.046 | 0.056 |
| Mtcl1 | 0.027 | 0.013 | 0.012 |
| 5033403F01F | 0.027 | 0.003 | 0.003 |
| Fam84b | 0.027 | 0.017 | 0.012 |
| Spata2l | 0.027 | 0.025 | 0.022 |
| Pdk4 | 0.027 | 0.152 | 0.070 |
| Gm38910 | 0.027 | 0.006 | 0.004 |
| Ubox5 | 0.027 | 0.017 | 0.018 |
| Kcnk3 | 0.027 | 0.411 | 0.043 |
| Rtl6 | 0.027 | 0.025 | 0.028 |
| Zdbf2 | 0.027 | 0.151 | 0.105 |
| Ifitm5 | 0.027 | 0.023 | 0.042 |
| Tnfaip6 | 0.027 | 0.157 | 0.307 |
| Sh3bp2 | 0.026 | 0.024 | 0.033 |
| Rsad1 | 0.026 | 0.028 | 0.036 |
| Unc13c | 0.026 | 0.043 | 0.021 |
| Hspbap1 | 0.026 | 0.028 | 0.028 |
| Fmn1 | 0.026 | 0.013 | 0.017 |
| Gm20632 | 0.026 | 0.052 | 0.023 |
| 1500015A07F | 0.026 | 0.016 | 0.018 |
| Pak3 | 0.026 | 0.148 | 0.093 |
| 1810024B03F | 0.026 | 0.036 | 0.036 |
| Il17d | 0.026 | 0.096 | 0.248 |
| Rrp12 | 0.026 | 0.023 | 0.024 |

| | | | |
|---|---|---|---|
| Ppp1r36 | 0.026 | 0.041 | 0.019 |
| Knl1 | 0.026 | 0.014 | 0.015 |
| Slc22a21 | 0.026 | 0.055 | 0.040 |
| Trim5 | 0.026 | 0.007 | 0.015 |
| Has3 | 0.026 | 0.046 | 0.007 |
| Usp29 | 0.026 | 0.037 | 0.089 |
| B3gntl1 | 0.026 | 0.022 | 0.022 |
| Lgi2 | 0.026 | 0.158 | 0.613 |
| Pnck | 0.026 | 0.090 | 0.047 |
| Lipt1 | 0.026 | 0.024 | 0.019 |
| Afg1l | 0.026 | 0.016 | 0.019 |
| Col15a1 | 0.026 | 0.128 | 0.178 |
| Dzip1 | 0.026 | 0.109 | 0.118 |
| Nanos3 | 0.026 | 0.029 | 0.024 |
| Aars2 | 0.026 | 0.033 | 0.036 |
| Aarsd1 | 0.026 | 0.017 | 0.016 |
| Adamtsl5 | 0.026 | 0.105 | 0.189 |
| Gm36033 | 0.026 | 0.029 | 0.018 |
| Gm26660 | 0.026 | 0.015 | 0.021 |
| Rbpms2 | 0.026 | 0.239 | 0.063 |
| Bean1 | 0.026 | 0.021 | 0.020 |
| Zfp354a | 0.026 | 0.027 | 0.032 |
| Brat1 | 0.026 | 0.031 | 0.038 |
| Tyw3 | 0.026 | 0.031 | 0.032 |
| Focad | 0.026 | 0.025 | 0.040 |
| Pnp2 | 0.026 | 0.007 | 0.020 |
| Pigb | 0.026 | 0.026 | 0.029 |
| Cxcl3 | 0.026 | 0.001 | 0.001 |
| 1810009A15F | 0.026 | 0.012 | 0.016 |
| Pcdhb21 | 0.026 | 0.025 | 0.025 |
| Urb1 | 0.026 | 0.040 | 0.036 |
| Spock3 | 0.026 | 0.013 | 0.025 |
| BC037039 | 0.026 | 0.021 | 0.025 |
| Rarres1 | 0.026 | 0.040 | 0.015 |
| Dclre1b | 0.026 | 0.026 | 0.030 |
| Mblac2 | 0.026 | 0.051 | 0.033 |
| Slc5a7 | 0.026 | 0.004 | 0.003 |
| Asns | 0.026 | 0.056 | 0.046 |
| Rnf150 | 0.026 | 0.182 | 0.154 |
| Ifit1 | 0.026 | 0.007 | 0.019 |
| Fndc10 | 0.026 | 0.066 | 0.123 |
| Cep126 | 0.026 | 0.024 | 0.014 |
| B230217C12F | 0.026 | 0.050 | 0.063 |
| Slc7a4 | 0.026 | 0.012 | 0.011 |
| Eda | 0.026 | 0.022 | 0.049 |
| Mars2 | 0.026 | 0.023 | 0.019 |
| Gm20707 | 0.026 | 0.014 | 0.027 |
| Pgap3 | 0.025 | 0.029 | 0.028 |

| | | | |
|---|---|---|---|
| Wnt5a | 0.025 | 0.074 | 0.306 |
| Gimap8 | 0.025 | 0.000 | 0.001 |
| Ppargc1a | 0.025 | 0.088 | 0.028 |
| Gpr45 | 0.025 | 0.009 | 0.008 |
| C1qc | 0.025 | 0.036 | 0.036 |
| Ppp1r3d | 0.025 | 0.068 | 0.031 |
| Cyp4v3 | 0.025 | 0.014 | 0.013 |
| Dnah14 | 0.025 | 0.001 | 0.001 |
| Adrb2 | 0.025 | 0.020 | 0.011 |
| A730063M14 | 0.025 | 0.021 | 0.017 |
| 3110009E18F | 0.025 | 0.040 | 0.042 |
| Slc46a1 | 0.025 | 0.030 | 0.037 |
| Tmem62 | 0.025 | 0.027 | 0.021 |
| Lst1 | 0.025 | 0.022 | 0.047 |
| Emilin2 | 0.025 | 0.007 | 0.193 |
| Radil | 0.025 | 0.005 | 0.038 |
| Hvcn1 | 0.025 | 0.010 | 0.075 |
| Setdb2 | 0.025 | 0.014 | 0.021 |
| Poli | 0.025 | 0.017 | 0.027 |
| Ovca2 | 0.025 | 0.010 | 0.016 |
| Zdhhc14 | 0.025 | 0.139 | 0.041 |
| Zeb2os | 0.025 | 0.039 | 0.070 |
| Slc29a4 | 0.025 | 0.002 | 0.001 |
| AL590144.2 | 0.025 | 0.004 | 0.011 |
| Gm34342 | 0.025 | 0.035 | 0.061 |
| Pdss1 | 0.025 | 0.033 | 0.035 |
| Ttc34 | 0.025 | 0.001 | 0.001 |
| Rorc | 0.025 | 0.002 | 0.000 |
| Mrvi1 | 0.025 | 0.283 | 0.045 |
| A330009N23I | 0.025 | 0.006 | 0.031 |
| Kcnb1 | 0.025 | 0.097 | 0.091 |
| Tsix | 0.025 | 0.052 | 0.035 |
| Syde2 | 0.025 | 0.112 | 0.061 |
| Pbk | 0.025 | 0.026 | 0.026 |
| Rexo5 | 0.025 | 0.017 | 0.024 |
| Ntrk2 | 0.025 | 0.138 | 0.539 |
| Rgs4 | 0.025 | 0.415 | 0.135 |
| Ttll3 | 0.025 | 0.044 | 0.055 |
| 1700088E04F | 0.025 | 0.019 | 0.018 |
| Oas1c | 0.025 | 0.008 | 0.005 |
| Atxn7l2 | 0.025 | 0.034 | 0.038 |
| Gm4876 | 0.025 | 0.031 | 0.008 |
| Nfatc2 | 0.025 | 0.060 | 0.121 |
| 1700084E18F | 0.025 | 0.016 | 0.014 |
| Six4 | 0.025 | 0.016 | 0.032 |
| Cd247 | 0.025 | 0.001 | 0.000 |
| 4933431E20F | 0.025 | 0.027 | 0.047 |
| 4933408B17F | 0.025 | 0.023 | 0.022 |

| | | | |
|---|---|---|---|
| Serinc2 | 0.025 | 0.027 | 0.026 |
| 4930579G24I | 0.025 | 0.026 | 0.029 |
| Entpd2 | 0.025 | 0.044 | 0.330 |
| Ipp | 0.025 | 0.027 | 0.014 |
| Rasgef1a | 0.025 | 0.006 | 0.009 |
| Vsir | 0.025 | 0.011 | 0.029 |
| Kank4 | 0.025 | 0.030 | 0.021 |
| Mark1 | 0.025 | 0.173 | 0.123 |
| Laptm5 | 0.025 | 0.012 | 0.013 |
| Cldn3 | 0.025 | 0.070 | 0.084 |
| F5 | 0.025 | 0.000 | 0.001 |
| Helq | 0.025 | 0.024 | 0.024 |
| Kcnk2 | 0.025 | 0.001 | 0.004 |
| Ccdc18 | 0.025 | 0.019 | 0.013 |
| Fbxl8 | 0.025 | 0.020 | 0.045 |
| Itgb8 | 0.025 | 0.092 | 0.057 |
| Myo19 | 0.025 | 0.021 | 0.025 |
| Cyb561d1 | 0.025 | 0.015 | 0.011 |
| Tigar | 0.025 | 0.017 | 0.009 |
| Hdhd5 | 0.025 | 0.019 | 0.017 |
| Aplnr | 0.025 | 0.003 | 0.003 |
| Tnni3 | 0.024 | 0.027 | 0.027 |
| Mtfp1 | 0.024 | 0.012 | 0.012 |
| B230369F24F | 0.024 | 0.045 | 0.041 |
| Dsn1 | 0.024 | 0.016 | 0.026 |
| Dusp10 | 0.024 | 0.168 | 0.165 |
| Lrrc61 | 0.024 | 0.023 | 0.020 |
| Sgip1 | 0.024 | 0.010 | 0.019 |
| 8430408G22I | 0.024 | 0.010 | 0.001 |
| Fyb | 0.024 | 0.046 | 0.028 |
| Pigl | 0.024 | 0.018 | 0.016 |
| Tsacc | 0.024 | 0.024 | 0.031 |
| Pcdhga6 | 0.024 | 0.028 | 0.028 |
| Fchsd1 | 0.024 | 0.022 | 0.026 |
| Gm30122 | 0.024 | 0.027 | 0.016 |
| Acaa1b | 0.024 | 0.002 | 0.008 |
| Adh7 | 0.024 | 0.001 | 0.003 |
| Pcdh12 | 0.024 | 0.000 | 0.001 |
| Tmem108 | 0.024 | 0.049 | 0.030 |
| Slc15a2 | 0.024 | 0.155 | 0.032 |
| Plce1 | 0.024 | 0.131 | 0.250 |
| Gm15441 | 0.024 | 0.014 | 0.014 |
| AI606181 | 0.024 | 0.027 | 0.024 |
| Gpihbp1 | 0.024 | 0.006 | 0.006 |
| Zfp41 | 0.024 | 0.039 | 0.037 |
| Clec16a | 0.024 | 0.041 | 0.033 |
| Wisp2 | 0.024 | 0.237 | 0.091 |
| Cpeb3 | 0.024 | 0.058 | 0.024 |

| | | | |
|---|---|---|---|
| Jpx | 0.024 | 0.020 | 0.021 |
| Foxo6 | 0.024 | 0.036 | 0.026 |
| Slc25a27 | 0.024 | 0.043 | 0.031 |
| Clip4 | 0.024 | 0.180 | 0.187 |
| Mndal | 0.024 | 0.003 | 0.028 |
| 9130221H12I | 0.024 | 0.025 | 0.029 |
| Epm2a | 0.024 | 0.028 | 0.015 |
| Gdpgp1 | 0.024 | 0.018 | 0.014 |
| Asxl3 | 0.024 | 0.015 | 0.047 |
| Rpp40 | 0.024 | 0.048 | 0.065 |
| Mks1 | 0.024 | 0.037 | 0.040 |
| Kif20b | 0.024 | 0.019 | 0.022 |
| D730003I15R | 0.024 | 0.028 | 0.042 |
| Rsph3a | 0.024 | 0.024 | 0.027 |
| Clec4e | 0.024 | 0.000 | 0.001 |
| Hps6 | 0.024 | 0.030 | 0.025 |
| Fabp3 | 0.024 | 0.235 | 0.077 |
| Dnajc27 | 0.024 | 0.033 | 0.034 |
| Ebf3 | 0.024 | 0.047 | 0.240 |
| Slc1a6 | 0.024 | 0.023 | 0.012 |
| Nkrf | 0.024 | 0.021 | 0.018 |
| Slc25a47 | 0.024 | 0.023 | 0.030 |
| Camk1d | 0.024 | 0.022 | 0.033 |
| Rimklb | 0.024 | 0.059 | 0.035 |
| Ebf2 | 0.024 | 0.015 | 0.498 |
| Ptgs2 | 0.024 | 0.188 | 0.043 |
| Rmst | 0.024 | 0.001 | 0.000 |
| Jmjd4 | 0.024 | 0.016 | 0.020 |
| Sbk1 | 0.024 | 0.026 | 0.032 |
| Med18 | 0.024 | 0.015 | 0.017 |
| Map3k7cl | 0.024 | 0.386 | 0.094 |
| Gm15706 | 0.024 | 0.025 | 0.034 |
| Pdpn | 0.024 | 0.048 | 0.290 |
| Krt79 | 0.024 | 0.019 | 0.013 |
| Ccdc85c | 0.024 | 0.010 | 0.010 |
| Efcab7 | 0.024 | 0.019 | 0.021 |
| Zfp605 | 0.024 | 0.025 | 0.028 |
| Ifitm1 | 0.024 | 0.029 | 0.104 |
| Vil1 | 0.024 | 0.000 | 0.000 |
| Itpkc | 0.024 | 0.065 | 0.077 |
| Lrp3 | 0.024 | 0.096 | 0.182 |
| Cd68 | 0.024 | 0.076 | 0.086 |
| Gm19412 | 0.024 | 0.028 | 0.037 |
| Bbs7 | 0.024 | 0.039 | 0.039 |
| Sesn2 | 0.024 | 0.040 | 0.031 |
| Napepld | 0.024 | 0.033 | 0.022 |
| Hexim2 | 0.024 | 0.028 | 0.022 |
| AC146911.1 | 0.024 | 0.019 | 0.016 |

| | | | |
|---|---|---|---|
| Morc1 | 0.023 | 0.001 | 0.001 |
| Gda | 0.023 | 0.003 | 0.015 |
| Ifit2 | 0.023 | 0.015 | 0.072 |
| P2ry10b | 0.023 | 0.003 | 0.014 |
| Fbxl12os | 0.023 | 0.016 | 0.017 |
| Rccd1 | 0.023 | 0.050 | 0.046 |
| Gmip | 0.023 | 0.037 | 0.052 |
| Slc22a5 | 0.023 | 0.014 | 0.012 |
| Rsph3b | 0.023 | 0.031 | 0.023 |
| Zfp335os | 0.023 | 0.008 | 0.005 |
| Twist2 | 0.023 | 0.116 | 0.545 |
| Psd | 0.023 | 0.345 | 0.221 |
| Tmem37 | 0.023 | 0.005 | 0.008 |
| Kctd1 | 0.023 | 0.214 | 0.065 |
| Foxf1 | 0.023 | 0.361 | 0.108 |
| Wdhd1 | 0.023 | 0.021 | 0.024 |
| Trip13 | 0.023 | 0.016 | 0.026 |
| Abcc10 | 0.023 | 0.023 | 0.021 |
| Lncpint | 0.023 | 0.054 | 0.021 |
| Tle6 | 0.023 | 0.024 | 0.041 |
| Ppp1r3e | 0.023 | 0.016 | 0.017 |
| Timm23 | 0.023 | 0.022 | 0.020 |
| E130102H24F | 0.023 | 0.055 | 0.065 |
| Sema7a | 0.023 | 0.001 | 0.002 |
| Nell1 | 0.023 | 0.296 | 0.073 |
| Vcpkmt | 0.023 | 0.032 | 0.035 |
| Slx4 | 0.023 | 0.027 | 0.033 |
| Prdm10 | 0.023 | 0.025 | 0.025 |
| Runx1 | 0.023 | 0.153 | 0.156 |
| Thada | 0.023 | 0.020 | 0.024 |
| 2510017J16R | 0.023 | 0.047 | 0.043 |
| Eps8l1 | 0.023 | 0.013 | 0.011 |
| C4b | 0.023 | 0.042 | 0.099 |
| Pars2 | 0.023 | 0.014 | 0.016 |
| Zfp39 | 0.023 | 0.024 | 0.035 |
| Sema3d | 0.023 | 0.073 | 0.308 |
| C330027C09F | 0.023 | 0.015 | 0.015 |
| Ccdc152 | 0.023 | 0.013 | 0.015 |
| Zfp458 | 0.023 | 0.016 | 0.018 |
| Nrros | 0.023 | 0.006 | 0.005 |
| Trim30d | 0.023 | 0.010 | 0.042 |
| Rsl1 | 0.023 | 0.017 | 0.023 |
| Sall1 | 0.023 | 0.000 | 0.000 |
| Rad54l | 0.023 | 0.014 | 0.012 |
| Gm6133 | 0.023 | 0.025 | 0.027 |
| Lekr1 | 0.023 | 0.022 | 0.036 |
| Gm15510 | 0.023 | 0.019 | 0.016 |
| Col24a1 | 0.023 | 0.198 | 0.160 |

| | | | |
|---|---|---|---|
| Pik3cg | 0.023 | 0.002 | 0.002 |
| Il6 | 0.023 | 0.044 | 0.156 |
| H2-Aa | 0.023 | 0.007 | 0.004 |
| Dixdc1 | 0.023 | 0.096 | 0.085 |
| Zfp54 | 0.023 | 0.007 | 0.012 |
| Ssc5d | 0.023 | 0.225 | 0.439 |
| Akr1c13 | 0.023 | 0.002 | 0.003 |
| Hdc | 0.023 | 0.003 | 0.006 |
| Trmo | 0.023 | 0.021 | 0.017 |
| Acp5 | 0.023 | 0.008 | 0.029 |
| C920021L13F | 0.023 | 0.034 | 0.031 |
| Jazf1 | 0.023 | 0.016 | 0.008 |
| Nudt12 | 0.023 | 0.005 | 0.007 |
| Gm9803 | 0.023 | 0.019 | 0.015 |
| Epha1 | 0.022 | 0.032 | 0.044 |
| Ss18l1 | 0.022 | 0.026 | 0.024 |
| Enpp3 | 0.022 | 0.039 | 0.186 |
| Gpr155 | 0.022 | 0.025 | 0.053 |
| C530044C16F | 0.022 | 0.022 | 0.028 |
| St14 | 0.022 | 0.003 | 0.003 |
| Arhgap27os1 | 0.022 | 0.000 | 0.001 |
| Slc38a4 | 0.022 | 0.160 | 0.376 |
| Gm15446 | 0.022 | 0.029 | 0.028 |
| Cps1 | 0.022 | 0.000 | 0.000 |
| Mbp | 0.022 | 0.044 | 0.034 |
| Usp18 | 0.022 | 0.010 | 0.011 |
| Npas2 | 0.022 | 0.034 | 0.025 |
| A130051J06R | 0.022 | 0.007 | 0.006 |
| Cenpp | 0.022 | 0.024 | 0.018 |
| Egfr | 0.022 | 0.167 | 0.407 |
| Hlf | 0.022 | 0.096 | 0.113 |
| Ccdc189 | 0.022 | 0.017 | 0.024 |
| Pdzd4 | 0.022 | 0.054 | 0.065 |
| Pygo1 | 0.022 | 0.155 | 0.150 |
| C330018D20I | 0.022 | 0.027 | 0.037 |
| Pcsk2os1 | 0.022 | 0.001 | 0.001 |
| 4933415A04F | 0.022 | 0.008 | 0.008 |
| Hdx | 0.022 | 0.015 | 0.018 |
| Gdpd3 | 0.022 | 0.011 | 0.014 |
| Tmem69 | 0.022 | 0.017 | 0.014 |
| Gm10602 | 0.022 | 0.023 | 0.014 |
| Strip2 | 0.022 | 0.041 | 0.014 |
| Mettl15 | 0.022 | 0.032 | 0.030 |
| Il33 | 0.022 | 0.055 | 0.439 |
| Gm10031 | 0.022 | 0.018 | 0.022 |
| AI838599 | 0.022 | 0.233 | 0.102 |
| E230032D23F | 0.022 | 0.000 | 0.000 |
| Bmp2 | 0.022 | 0.199 | 0.092 |

| | | | |
|---|---|---|---|
| Hapln1 | 0.022 | 0.030 | 0.044 |
| Dusp5 | 0.022 | 0.178 | 0.132 |
| Homez | 0.022 | 0.018 | 0.018 |
| Sgo2a | 0.022 | 0.024 | 0.015 |
| Tceanc | 0.022 | 0.022 | 0.025 |
| Lrg1 | 0.022 | 0.030 | 0.005 |
| Mapt | 0.022 | 0.039 | 0.031 |
| Tnfsf9 | 0.022 | 0.110 | 0.080 |
| Srpx | 0.022 | 0.168 | 0.478 |
| Tslp | 0.022 | 0.035 | 0.018 |
| Slc16a7 | 0.022 | 0.077 | 0.020 |
| Gm6213 | 0.022 | 0.002 | 0.007 |
| Cdc7 | 0.022 | 0.017 | 0.036 |
| Mss51 | 0.022 | 0.024 | 0.029 |
| Zfp873 | 0.022 | 0.038 | 0.037 |
| 1700086D15I | 0.022 | 0.004 | 0.003 |
| Plekhf1 | 0.022 | 0.091 | 0.142 |
| Mettl4 | 0.022 | 0.010 | 0.015 |
| Gm33104 | 0.022 | 0.000 | 0.000 |
| Ccdc173 | 0.022 | 0.031 | 0.026 |
| Zfp607b | 0.022 | 0.033 | 0.028 |
| Glt8d2 | 0.022 | 0.036 | 0.250 |
| Gnb1l | 0.022 | 0.017 | 0.021 |
| Zfr2 | 0.022 | 0.064 | 0.078 |
| CAAA0114733 | 0.022 | 0.016 | 0.025 |
| Garnl3 | 0.022 | 0.055 | 0.074 |
| Gm15545 | 0.022 | 0.022 | 0.027 |
| 1700066M21 | 0.022 | 0.020 | 0.018 |
| Gfpt2 | 0.022 | 0.028 | 0.320 |
| Fcor | 0.022 | 0.018 | 0.031 |
| Neurl1a | 0.022 | 0.074 | 0.036 |
| Adamts6 | 0.022 | 0.165 | 0.068 |
| Fgf11 | 0.022 | 0.015 | 0.029 |
| 1200007C13I | 0.022 | 0.172 | 0.123 |
| Aqp11 | 0.022 | 0.008 | 0.008 |
| Tmem177 | 0.022 | 0.015 | 0.013 |
| Mfsd13a | 0.022 | 0.005 | 0.003 |
| Nhej1 | 0.022 | 0.032 | 0.033 |
| Osbpl6 | 0.022 | 0.093 | 0.052 |
| Serpina3n | 0.022 | 0.030 | 0.171 |
| Fignl1 | 0.022 | 0.014 | 0.020 |
| Dnaaf3 | 0.022 | 0.010 | 0.019 |
| Kcnh2 | 0.022 | 0.097 | 0.019 |
| 6330418K02I | 0.022 | 0.023 | 0.024 |
| Zfp968 | 0.022 | 0.019 | 0.019 |
| Nfkbid | 0.022 | 0.007 | 0.005 |
| Lrp12 | 0.022 | 0.061 | 0.044 |
| Wnt9b | 0.022 | 0.000 | 0.000 |

| | | | |
|---|---|---|---|
| Colca2 | 0.022 | 0.012 | 0.009 |
| Amigo1 | 0.022 | 0.021 | 0.029 |
| Tnfrsf25 | 0.021 | 0.011 | 0.004 |
| Ahcy | 0.021 | 0.033 | 0.027 |
| Slc26a6 | 0.021 | 0.029 | 0.024 |
| Nrp2 | 0.021 | 0.073 | 0.166 |
| AC162302.1 | 0.021 | 0.027 | 0.021 |
| Milr1 | 0.021 | 0.066 | 0.080 |
| Tub | 0.021 | 0.219 | 0.077 |
| Ap1g2 | 0.021 | 0.014 | 0.017 |
| Gm17315 | 0.021 | 0.026 | 0.015 |
| Fam120aos | 0.021 | 0.021 | 0.019 |
| Zfp790 | 0.021 | 0.021 | 0.025 |
| Nptx1 | 0.021 | 0.015 | 0.016 |
| Scml2 | 0.021 | 0.015 | 0.019 |
| Cbs | 0.021 | 0.079 | 0.022 |
| Apoc1 | 0.021 | 0.003 | 0.011 |
| Pllp | 0.021 | 0.001 | 0.002 |
| Eml6 | 0.021 | 0.005 | 0.004 |
| Adat1 | 0.021 | 0.020 | 0.024 |
| Gm17092 | 0.021 | 0.024 | 0.024 |
| Sfxn4 | 0.021 | 0.017 | 0.029 |
| Nanos1 | 0.021 | 0.034 | 0.018 |
| Inha | 0.021 | 0.055 | 0.052 |
| Farp2 | 0.021 | 0.042 | 0.042 |
| Stac3 | 0.021 | 0.017 | 0.019 |
| Gm10399 | 0.021 | 0.012 | 0.012 |
| Fbln7 | 0.021 | 0.037 | 0.099 |
| 4930550C14F | 0.021 | 0.012 | 0.009 |
| Gm48933 | 0.021 | 0.028 | 0.040 |
| Ankrd6 | 0.021 | 0.006 | 0.016 |
| Snx20 | 0.021 | 0.003 | 0.006 |
| Cpz | 0.021 | 0.044 | 0.570 |
| Echdc3 | 0.021 | 0.042 | 0.036 |
| Zfp141 | 0.021 | 0.031 | 0.035 |
| Six2 | 0.021 | 0.078 | 0.053 |
| Penk | 0.021 | 0.087 | 0.583 |
| Lhpp | 0.021 | 0.021 | 0.024 |
| Etv5 | 0.021 | 0.129 | 0.059 |
| Phf11b | 0.021 | 0.001 | 0.007 |
| Tuba4a | 0.021 | 0.036 | 0.010 |
| Meis3 | 0.021 | 0.158 | 0.157 |
| D630045J12F | 0.021 | 0.032 | 0.045 |
| Gm17259 | 0.021 | 0.013 | 0.012 |
| Fam160a1 | 0.021 | 0.018 | 0.003 |
| Fam89b | 0.021 | 0.014 | 0.014 |
| Zfp341 | 0.021 | 0.013 | 0.017 |
| Dusp26 | 0.021 | 0.004 | 0.002 |

| | | | |
|---|---|---|---|
| 2610021A01F | 0.021 | 0.018 | 0.023 |
| Mapk11 | 0.021 | 0.004 | 0.009 |
| Lgals4 | 0.021 | 0.037 | 0.042 |
| Cbx2 | 0.021 | 0.017 | 0.024 |
| Rab19 | 0.021 | 0.004 | 0.005 |
| Msh5 | 0.021 | 0.008 | 0.015 |
| Srd5a1 | 0.021 | 0.036 | 0.040 |
| Car3 | 0.021 | 0.039 | 0.033 |
| Lyz2 | 0.021 | 0.031 | 0.040 |
| Trnp1 | 0.021 | 0.167 | 0.212 |
| Endou | 0.021 | 0.008 | 0.008 |
| Cd46 | 0.021 | 0.021 | 0.026 |
| Apol7d | 0.021 | 0.017 | 0.015 |
| Hsd17b7 | 0.021 | 0.065 | 0.073 |
| Patj | 0.021 | 0.031 | 0.028 |
| Ephb2 | 0.021 | 0.139 | 0.232 |
| Atp1b2 | 0.021 | 0.389 | 0.136 |
| Gldc | 0.021 | 0.002 | 0.005 |
| Aifm2 | 0.021 | 0.053 | 0.076 |
| 3110021N24F | 0.021 | 0.025 | 0.034 |
| Slc17a9 | 0.021 | 0.008 | 0.005 |
| Pcdhb19 | 0.021 | 0.028 | 0.026 |
| Zfp189 | 0.021 | 0.022 | 0.022 |
| Sugct | 0.021 | 0.030 | 0.030 |
| Arhgap11a | 0.021 | 0.020 | 0.020 |
| Zfp287 | 0.021 | 0.036 | 0.038 |
| Dcbld1 | 0.021 | 0.052 | 0.026 |
| Gm11273 | 0.021 | 0.014 | 0.014 |
| Hist2h2aa1 | 0.021 | 0.012 | 0.009 |
| Zfp2 | 0.021 | 0.031 | 0.024 |
| Tnfsfm13 | 0.021 | 0.014 | 0.012 |
| Mapkapk5 | 0.021 | 0.029 | 0.033 |
| Zfat | 0.021 | 0.016 | 0.018 |
| Serpinc1 | 0.021 | 0.016 | 0.014 |
| 1810021B22F | 0.021 | 0.017 | 0.020 |
| Hydin | 0.021 | 0.000 | 0.001 |
| Gm16486 | 0.021 | 0.057 | 0.096 |
| Taco1os | 0.021 | 0.021 | 0.034 |
| Nme4 | 0.020 | 0.031 | 0.096 |
| Kcnq1 | 0.020 | 0.000 | 0.001 |
| Il6ra | 0.020 | 0.029 | 0.050 |
| Mtbp | 0.020 | 0.030 | 0.027 |
| Nudt8.1 | 0.020 | 0.038 | 0.035 |
| Galnt17 | 0.020 | 0.117 | 0.373 |
| Tmem220 | 0.020 | 0.047 | 0.029 |
| 4632404H12F | 0.020 | 0.006 | 0.005 |
| Cdc6 | 0.020 | 0.006 | 0.010 |
| Gins1 | 0.020 | 0.019 | 0.021 |

| | | | |
|---|---|---|---|
| Sfn | 0.020 | 0.023 | 0.012 |
| Adamts7 | 0.020 | 0.021 | 0.054 |
| 2010109A12F | 0.020 | 0.014 | 0.019 |
| Sfxn5 | 0.020 | 0.030 | 0.034 |
| Sntb1 | 0.020 | 0.059 | 0.010 |
| Cd52 | 0.020 | 0.015 | 0.016 |
| Fam227a | 0.020 | 0.018 | 0.014 |
| Rab36 | 0.020 | 0.034 | 0.031 |
| 4931428F04F | 0.020 | 0.021 | 0.029 |
| Hook2 | 0.020 | 0.053 | 0.048 |
| Cdhr4 | 0.020 | 0.018 | 0.019 |
| Them6 | 0.020 | 0.006 | 0.007 |
| Aplf | 0.020 | 0.036 | 0.045 |
| Zfp709 | 0.020 | 0.018 | 0.020 |
| Psmc3ip | 0.020 | 0.046 | 0.046 |
| Prkcq | 0.020 | 0.003 | 0.002 |
| Kcnmb4 | 0.020 | 0.123 | 0.044 |
| Hpgds | 0.020 | 0.019 | 0.023 |
| Ism2 | 0.020 | 0.001 | 0.006 |
| Amigo3 | 0.020 | 0.016 | 0.033 |
| Fggy | 0.020 | 0.032 | 0.025 |
| Abcc12 | 0.020 | 0.000 | 0.000 |
| Ednra | 0.020 | 0.253 | 0.323 |
| Ctla2b | 0.020 | 0.000 | 0.001 |
| Vill | 0.020 | 0.016 | 0.016 |
| Wfdc18 | 0.020 | 0.033 | 0.163 |
| Garem1 | 0.020 | 0.135 | 0.089 |
| Pianp | 0.020 | 0.320 | 0.122 |
| Zfp689 | 0.020 | 0.016 | 0.023 |
| Pcdhb18 | 0.020 | 0.011 | 0.009 |
| Pkmyt1 | 0.020 | 0.022 | 0.051 |
| Rec8 | 0.020 | 0.002 | 0.002 |
| Gm6169 | 0.020 | 0.024 | 0.019 |
| Reep1 | 0.020 | 0.030 | 0.012 |
| Ptpru | 0.020 | 0.027 | 0.023 |
| AC154760.1 | 0.020 | 0.019 | 0.022 |
| Fbxo44 | 0.020 | 0.031 | 0.010 |
| Cntrob | 0.020 | 0.023 | 0.023 |
| Rad51 | 0.020 | 0.014 | 0.025 |
| Mfsd4b4 | 0.020 | 0.012 | 0.012 |
| Snhg10 | 0.020 | 0.033 | 0.024 |
| Urb2 | 0.020 | 0.018 | 0.015 |
| Ccl2 | 0.020 | 0.046 | 0.090 |
| Zfp971 | 0.020 | 0.017 | 0.015 |
| 1110019D14I | 0.020 | 0.014 | 0.017 |
| Zfp955a | 0.020 | 0.018 | 0.018 |
| Zswim3 | 0.020 | 0.008 | 0.015 |
| Il17ra | 0.020 | 0.026 | 0.048 |

| | | | |
|---|---|---|---|
| Mir17hg | 0.020 | 0.039 | 0.031 |
| Wnt16 | 0.020 | 0.021 | 0.002 |
| Zc3h12b | 0.020 | 0.032 | 0.033 |
| Nexmif | 0.020 | 0.003 | 0.012 |
| Shd | 0.020 | 0.045 | 0.026 |
| Shpk | 0.020 | 0.011 | 0.021 |
| Aldh1a7 | 0.020 | 0.001 | 0.003 |
| E530011L22R | 0.020 | 0.029 | 0.035 |
| Cep152 | 0.020 | 0.013 | 0.015 |
| Cd109 | 0.020 | 0.061 | 0.050 |
| Aurka | 0.020 | 0.012 | 0.011 |
| Rflna | 0.020 | 0.047 | 0.038 |
| Cdc45 | 0.020 | 0.018 | 0.023 |
| Arap2 | 0.020 | 0.300 | 0.086 |
| Aff2 | 0.020 | 0.067 | 0.070 |
| Plk1 | 0.020 | 0.012 | 0.013 |
| Clcn2 | 0.020 | 0.023 | 0.020 |
| Akap6 | 0.020 | 0.170 | 0.026 |
| Dnaaf2 | 0.020 | 0.021 | 0.021 |
| Cited4 | 0.020 | 0.013 | 0.004 |
| Chchd10 | 0.020 | 0.048 | 0.024 |
| Trpm6 | 0.020 | 0.001 | 0.001 |
| Efcc1 | 0.020 | 0.001 | 0.028 |
| Nuf2 | 0.020 | 0.010 | 0.013 |
| 4933406I18R | 0.020 | 0.005 | 0.003 |
| 1810010H24I | 0.020 | 0.046 | 0.180 |
| Dnajb3 | 0.020 | 0.016 | 0.028 |
| Mb21d1 | 0.020 | 0.008 | 0.017 |
| Lrrc29 | 0.020 | 0.011 | 0.011 |
| Irs1 | 0.020 | 0.093 | 0.150 |
| Cenph | 0.020 | 0.014 | 0.020 |
| Ar | 0.020 | 0.035 | 0.127 |
| Ankrd42 | 0.020 | 0.019 | 0.019 |
| Adamtsl3 | 0.020 | 0.093 | 0.119 |
| Creb3l4 | 0.020 | 0.012 | 0.014 |
| Pcdhga9 | 0.020 | 0.022 | 0.023 |
| 1700113A16F | 0.020 | 0.018 | 0.020 |
| Rgl3 | 0.020 | 0.015 | 0.006 |
| Aatk | 0.019 | 0.030 | 0.019 |
| Egr2 | 0.019 | 0.125 | 0.164 |
| Ifi44 | 0.019 | 0.003 | 0.002 |
| Slc5a6 | 0.019 | 0.013 | 0.020 |
| Gm37422 | 0.019 | 0.009 | 0.010 |
| Tspan33 | 0.019 | 0.007 | 0.004 |
| Aspm | 0.019 | 0.016 | 0.016 |
| Zfp551 | 0.019 | 0.021 | 0.018 |
| Aldh1a1 | 0.019 | 0.007 | 0.165 |
| D130020L05F | 0.019 | 0.031 | 0.035 |

| | | | |
|---|---|---|---|
| Cib2 | 0.019 | 0.121 | 0.046 |
| Cfp | 0.019 | 0.026 | 0.020 |
| Jrk | 0.019 | 0.015 | 0.017 |
| Zfp184 | 0.019 | 0.023 | 0.016 |
| Sult5a1 | 0.019 | 0.047 | 0.265 |
| Kdm1b | 0.019 | 0.031 | 0.029 |
| Slc35g2 | 0.019 | 0.006 | 0.004 |
| Zfp956 | 0.019 | 0.029 | 0.029 |
| Eya4 | 0.019 | 0.005 | 0.049 |
| Atp6v1g2 | 0.019 | 0.019 | 0.016 |
| Anxa8 | 0.019 | 0.015 | 0.013 |
| Nek3 | 0.019 | 0.018 | 0.022 |
| Fancg | 0.019 | 0.008 | 0.014 |
| Krt19 | 0.019 | 0.020 | 0.044 |
| Plekhd1 | 0.019 | 0.004 | 0.009 |
| Rab20 | 0.019 | 0.140 | 0.088 |
| Cmtr2 | 0.019 | 0.013 | 0.011 |
| Kif1a | 0.019 | 0.116 | 0.156 |
| Retsat | 0.019 | 0.039 | 0.040 |
| 1110035H17I | 0.019 | 0.023 | 0.014 |
| Rassf5 | 0.019 | 0.004 | 0.012 |
| Pou5f1 | 0.019 | 0.013 | 0.017 |
| Taco1 | 0.019 | 0.014 | 0.013 |
| Gas7 | 0.019 | 0.085 | 0.359 |
| Arvcf | 0.019 | 0.047 | 0.041 |
| Sh2d4b | 0.019 | 0.001 | 0.008 |
| Lad1 | 0.019 | 0.000 | 0.000 |
| Gm4262 | 0.019 | 0.004 | 0.014 |
| 9130019O22I | 0.019 | 0.011 | 0.016 |
| Ppp1r18os | 0.019 | 0.018 | 0.022 |
| Ncaph | 0.019 | 0.021 | 0.025 |
| Mxd3 | 0.019 | 0.013 | 0.017 |
| Xlr | 0.019 | 0.009 | 0.020 |
| Pdzrn4 | 0.019 | 0.012 | 0.002 |
| Coro1a | 0.019 | 0.018 | 0.019 |
| Gm4951 | 0.019 | 0.022 | 0.057 |
| Zgrf1 | 0.019 | 0.025 | 0.028 |
| Lor | 0.019 | 0.007 | 0.005 |
| Myh6 | 0.019 | 0.014 | 0.023 |
| Ephb1 | 0.019 | 0.021 | 0.011 |
| Ttn | 0.019 | 0.013 | 0.016 |
| Zfp607a | 0.019 | 0.030 | 0.024 |
| Hoxd3os1 | 0.019 | 0.000 | 0.001 |
| Zfp112 | 0.019 | 0.031 | 0.047 |
| Lrch1 | 0.019 | 0.088 | 0.066 |
| Plek | 0.019 | 0.011 | 0.013 |
| Erich2 | 0.019 | 0.008 | 0.004 |
| Vsig10l | 0.019 | 0.020 | 0.022 |

| | | | |
|---|---|---|---|
| Zbtb49 | 0.019 | 0.019 | 0.017 |
| Micall2 | 0.019 | 0.010 | 0.012 |
| D430019H16I | 0.019 | 0.034 | 0.063 |
| C230037L18F | 0.019 | 0.004 | 0.007 |
| Sema5b | 0.019 | 0.022 | 0.017 |
| Ildr2 | 0.019 | 0.009 | 0.061 |
| AL591952.3 | 0.019 | 0.024 | 0.026 |
| Gria4 | 0.019 | 0.006 | 0.026 |
| Gm14167 | 0.019 | 0.010 | 0.011 |
| Pcdhb5 | 0.019 | 0.010 | 0.010 |
| Fam81a | 0.019 | 0.138 | 0.015 |
| Kcnab1 | 0.019 | 0.134 | 0.039 |
| Fndc5 | 0.019 | 0.055 | 0.128 |
| Ndc80 | 0.019 | 0.011 | 0.013 |
| 1110046J04R | 0.019 | 0.075 | 0.070 |
| A930028N01I | 0.019 | 0.020 | 0.006 |
| Syngr1 | 0.019 | 0.004 | 0.004 |
| Ttc30a1 | 0.019 | 0.010 | 0.015 |
| K230015D01I | 0.019 | 0.019 | 0.014 |
| Zfp300 | 0.019 | 0.014 | 0.018 |
| Glycam1 | 0.019 | 0.000 | 0.000 |
| Mst1r | 0.019 | 0.016 | 0.050 |
| Rragd | 0.018 | 0.029 | 0.028 |
| Gm21887 | 0.018 | 0.028 | 0.019 |
| Asap3 | 0.018 | 0.022 | 0.067 |
| Map4k1 | 0.018 | 0.002 | 0.003 |
| Zfp85 | 0.018 | 0.011 | 0.017 |
| Rasl10a | 0.018 | 0.014 | 0.044 |
| Crispld2 | 0.018 | 0.455 | 0.205 |
| Acad10 | 0.018 | 0.005 | 0.009 |
| Gm4924 | 0.018 | 0.021 | 0.033 |
| Gpd1 | 0.018 | 0.018 | 0.009 |
| Ndor1 | 0.018 | 0.011 | 0.012 |
| Ripply3 | 0.018 | 0.002 | 0.004 |
| Grb14 | 0.018 | 0.161 | 0.093 |
| 1700097N02I | 0.018 | 0.000 | 0.000 |
| Epop | 0.018 | 0.023 | 0.014 |
| Brsk1 | 0.018 | 0.054 | 0.059 |
| Hap1 | 0.018 | 0.006 | 0.003 |
| Klf8 | 0.018 | 0.020 | 0.009 |
| Ptx3 | 0.018 | 0.063 | 0.179 |
| Zfp772 | 0.018 | 0.022 | 0.017 |
| Mn1 | 0.018 | 0.250 | 0.302 |
| Cldn11 | 0.018 | 0.003 | 0.021 |
| Cytip | 0.018 | 0.036 | 0.007 |
| AV099323 | 0.018 | 0.007 | 0.010 |
| BC052040 | 0.018 | 0.019 | 0.028 |
| Zfp691 | 0.018 | 0.018 | 0.023 |

| | | | |
|---|---|---|---|
| Adgrl3 | 0.018 | 0.082 | 0.112 |
| 1700048O20I | 0.018 | 0.020 | 0.027 |
| Ripor2 | 0.018 | 0.016 | 0.033 |
| Msi1 | 0.018 | 0.014 | 0.021 |
| Icam4 | 0.018 | 0.002 | 0.003 |
| Stox1 | 0.018 | 0.001 | 0.003 |
| Ccdc14 | 0.018 | 0.021 | 0.018 |
| Fam151b | 0.018 | 0.028 | 0.031 |
| Slc2a9 | 0.018 | 0.003 | 0.005 |
| Lurap1l | 0.018 | 0.061 | 0.164 |
| Zfp185 | 0.018 | 0.010 | 0.020 |
| Pcdh18 | 0.018 | 0.095 | 0.227 |
| Gsg1 | 0.018 | 0.001 | 0.002 |
| Ptrh1 | 0.018 | 0.018 | 0.018 |
| 1700109H08I | 0.018 | 0.020 | 0.017 |
| Pcdhga10 | 0.018 | 0.016 | 0.010 |
| Zfp612 | 0.018 | 0.054 | 0.021 |
| Asb1 | 0.018 | 0.050 | 0.054 |
| Fam49a | 0.018 | 0.020 | 0.016 |
| Cenpn | 0.018 | 0.017 | 0.020 |
| Zfp955b | 0.018 | 0.023 | 0.024 |
| Gm10643 | 0.018 | 0.013 | 0.016 |
| 6430573P05F | 0.018 | 0.014 | 0.015 |
| Bfsp1 | 0.018 | 0.014 | 0.047 |
| Ptger1 | 0.018 | 0.029 | 0.023 |
| Zcwpw1 | 0.018 | 0.007 | 0.020 |
| Gm17435 | 0.018 | 0.035 | 0.054 |
| 4930518I15R | 0.018 | 0.010 | 0.010 |
| Frs3 | 0.018 | 0.010 | 0.007 |
| 9030025P20F | 0.018 | 0.018 | 0.015 |
| Gm42595 | 0.018 | 0.015 | 0.014 |
| Zfp420 | 0.018 | 0.022 | 0.023 |
| Cnbd2 | 0.018 | 0.014 | 0.011 |
| Il16 | 0.018 | 0.033 | 0.019 |
| Iba57 | 0.018 | 0.008 | 0.014 |
| Phkg1 | 0.018 | 0.052 | 0.039 |
| Nacad | 0.018 | 0.098 | 0.029 |
| Proser3 | 0.018 | 0.031 | 0.024 |
| Mastl | 0.018 | 0.029 | 0.022 |
| Aldh1a3 | 0.018 | 0.010 | 0.089 |
| Fbxl2 | 0.018 | 0.036 | 0.012 |
| Enpp1 | 0.018 | 0.136 | 0.222 |
| Vstm4 | 0.018 | 0.101 | 0.445 |
| Pms1 | 0.018 | 0.012 | 0.018 |
| Gm19557 | 0.018 | 0.001 | 0.000 |
| Eme2 | 0.018 | 0.017 | 0.014 |
| Rab26 | 0.018 | 0.015 | 0.012 |
| Cecr2 | 0.018 | 0.003 | 0.008 |

| | | | |
|---|---|---|---|
| Sirt4 | 0.018 | 0.036 | 0.034 |
| Pdxp | 0.018 | 0.020 | 0.026 |
| Cadm3 | 0.018 | 0.004 | 0.057 |
| Zfp128 | 0.018 | 0.010 | 0.009 |
| Isoc2a | 0.018 | 0.023 | 0.025 |
| Dnm1 | 0.018 | 0.047 | 0.235 |
| A830010M20 | 0.018 | 0.020 | 0.027 |
| Smkr-ps | 0.018 | 0.009 | 0.011 |
| Gm340 | 0.018 | 0.018 | 0.018 |
| Gm17455 | 0.018 | 0.024 | 0.020 |
| Tnik | 0.018 | 0.225 | 0.121 |
| Ldb3 | 0.017 | 0.117 | 0.015 |
| Otub2 | 0.017 | 0.012 | 0.019 |
| Gm38393 | 0.017 | 0.008 | 0.008 |
| Gm12166 | 0.017 | 0.010 | 0.009 |
| Ppp2r2b | 0.017 | 0.021 | 0.063 |
| Zfp456 | 0.017 | 0.006 | 0.009 |
| Csf1r | 0.017 | 0.015 | 0.017 |
| Pm20d2 | 0.017 | 0.011 | 0.005 |
| Ftx | 0.017 | 0.026 | 0.029 |
| Adcy9 | 0.017 | 0.077 | 0.039 |
| Setd6 | 0.017 | 0.078 | 0.033 |
| Nod2 | 0.017 | 0.000 | 0.001 |
| Tmc7 | 0.017 | 0.004 | 0.006 |
| Epb41l1 | 0.017 | 0.095 | 0.116 |
| 2900089D17I | 0.017 | 0.052 | 0.028 |
| Jakmip1 | 0.017 | 0.000 | 0.000 |
| Eif3j2 | 0.017 | 0.032 | 0.042 |
| Cacna1c | 0.017 | 0.145 | 0.073 |
| Rnase1 | 0.017 | 0.000 | 0.000 |
| Chrna2 | 0.017 | 0.001 | 0.001 |
| Gmfg | 0.017 | 0.009 | 0.013 |
| Ly6h | 0.017 | 0.003 | 0.031 |
| St8sia6 | 0.017 | 0.001 | 0.001 |
| Pfkfb4 | 0.017 | 0.026 | 0.032 |
| Ren1 | 0.017 | 0.015 | 0.250 |
| Hist2h2ac | 0.017 | 0.012 | 0.009 |
| Pcx | 0.017 | 0.012 | 0.016 |
| 6430548M08 | 0.017 | 0.011 | 0.018 |
| Gm16638 | 0.017 | 0.024 | 0.022 |
| Lonrf3 | 0.017 | 0.004 | 0.002 |
| Frg2f1 | 0.017 | 0.026 | 0.039 |
| Clec1b | 0.017 | 0.001 | 0.000 |
| Kif4 | 0.017 | 0.010 | 0.012 |
| Frem2 | 0.017 | 0.035 | 0.004 |
| Greb1l | 0.017 | 0.013 | 0.021 |
| Gk | 0.017 | 0.071 | 0.029 |
| Kifc2 | 0.017 | 0.018 | 0.018 |

| | | | |
|---|---|---|---|
| Cacnb1 | 0.017 | 0.069 | 0.050 |
| Gm15952 | 0.017 | 0.016 | 0.022 |
| Tmem268 | 0.017 | 0.009 | 0.014 |
| Pag1 | 0.017 | 0.050 | 0.049 |
| Suv39h2 | 0.017 | 0.018 | 0.017 |
| Glrb | 0.017 | 0.008 | 0.015 |
| Rep15 | 0.017 | 0.020 | 0.028 |
| Gm26982 | 0.017 | 0.015 | 0.010 |
| Bbs1 | 0.017 | 0.020 | 0.017 |
| Hist4h4 | 0.017 | 0.025 | 0.021 |
| Tmod1 | 0.017 | 0.112 | 0.056 |
| 1810062O18I | 0.017 | 0.017 | 0.025 |
| Mmachc | 0.017 | 0.024 | 0.025 |
| Gm527 | 0.017 | 0.024 | 0.040 |
| Hopx | 0.017 | 0.028 | 0.061 |
| Stc2 | 0.017 | 0.156 | 0.086 |
| Dpf1 | 0.017 | 0.062 | 0.048 |
| Spry4 | 0.017 | 0.142 | 0.102 |
| Acpp | 0.017 | 0.000 | 0.001 |
| Pitx2 | 0.017 | 0.002 | 0.004 |
| Setmar | 0.017 | 0.016 | 0.016 |
| Myo7a | 0.017 | 0.008 | 0.016 |
| 1110006O24I | 0.017 | 0.011 | 0.012 |
| Ccdc62 | 0.017 | 0.016 | 0.021 |
| Pomc | 0.017 | 0.006 | 0.013 |
| Mdrl | 0.017 | 0.021 | 0.011 |
| Kcnc2 | 0.017 | 0.007 | 0.018 |
| AW011738 | 0.017 | 0.038 | 0.016 |
| Pcdhgc3 | 0.017 | 0.022 | 0.026 |
| Pole | 0.017 | 0.013 | 0.021 |
| Ooep | 0.017 | 0.024 | 0.005 |
| Scube3 | 0.017 | 0.198 | 0.041 |
| Adam12 | 0.017 | 0.010 | 0.207 |
| Gm10053 | 0.017 | 0.013 | 0.008 |
| Ccdc30 | 0.017 | 0.013 | 0.016 |
| Igdcc4 | 0.017 | 0.148 | 0.320 |
| C330011M18 | 0.017 | 0.009 | 0.011 |
| Hspa12a | 0.017 | 0.005 | 0.024 |
| Fam217a | 0.017 | 0.002 | 0.007 |
| Hnmt | 0.017 | 0.010 | 0.014 |
| Tceal5 | 0.017 | 0.056 | 0.085 |
| Mfsd9 | 0.017 | 0.007 | 0.015 |
| Fan1 | 0.017 | 0.015 | 0.011 |
| Bbs10 | 0.017 | 0.014 | 0.014 |
| Nupr1l | 0.017 | 0.009 | 0.008 |
| Cep55 | 0.017 | 0.005 | 0.003 |
| Unc5d | 0.017 | 0.004 | 0.002 |
| Kcnq4 | 0.017 | 0.023 | 0.093 |

| | | | |
|---|---|---|---|
| Asf1b | 0.017 | 0.008 | 0.014 |
| 2700046A07F | 0.017 | 0.008 | 0.003 |
| Klf5 | 0.017 | 0.032 | 0.037 |
| Gm15283 | 0.017 | 0.021 | 0.016 |
| Gm26514 | 0.017 | 0.004 | 0.006 |
| 4930539J05R | 0.017 | 0.021 | 0.021 |
| Pimreg | 0.017 | 0.012 | 0.017 |
| Gm28875 | 0.017 | 0.089 | 0.080 |
| Zfp72 | 0.016 | 0.014 | 0.019 |
| Zfp438 | 0.016 | 0.024 | 0.034 |
| Odf3b | 0.016 | 0.000 | 0.000 |
| D130017N08I | 0.016 | 0.010 | 0.016 |
| Tmem91 | 0.016 | 0.011 | 0.007 |
| Zfp599 | 0.016 | 0.012 | 0.007 |
| Chek1 | 0.016 | 0.023 | 0.025 |
| Senp8 | 0.016 | 0.020 | 0.036 |
| Cst6 | 0.016 | 0.005 | 0.004 |
| Gm14419 | 0.016 | 0.025 | 0.024 |
| Adamts14 | 0.016 | 0.048 | 0.024 |
| Nudt10 | 0.016 | 0.017 | 0.011 |
| C8g | 0.016 | 0.007 | 0.013 |
| Sh3gl2 | 0.016 | 0.011 | 0.017 |
| Rinl | 0.016 | 0.013 | 0.013 |
| 4933404O12I | 0.016 | 0.024 | 0.018 |
| Gm19531 | 0.016 | 0.023 | 0.039 |
| Fam184a | 0.016 | 0.011 | 0.011 |
| Carmil3 | 0.016 | 0.014 | 0.030 |
| Rhbdl2 | 0.016 | 0.002 | 0.001 |
| Lrrc15 | 0.016 | 0.002 | 0.013 |
| Carmn | 0.016 | 0.367 | 0.143 |
| Gabra1 | 0.016 | 0.158 | 0.188 |
| Gm29157 | 0.016 | 0.001 | 0.004 |
| Sag | 0.016 | 0.018 | 0.019 |
| Gm9774 | 0.016 | 0.014 | 0.014 |
| Glt28d2 | 0.016 | 0.011 | 0.003 |
| Zfp273 | 0.016 | 0.013 | 0.020 |
| Daam2 | 0.016 | 0.173 | 0.124 |
| Tmem212 | 0.016 | 0.008 | 0.002 |
| Etnk2 | 0.016 | 0.013 | 0.037 |
| Kcne4 | 0.016 | 0.137 | 0.189 |
| Map3k5 | 0.016 | 0.087 | 0.043 |
| Lbhd1 | 0.016 | 0.010 | 0.011 |
| Vash1 | 0.016 | 0.006 | 0.012 |
| Gm38394 | 0.016 | 0.023 | 0.021 |
| Hist1h2be | 0.016 | 0.011 | 0.010 |
| Nxpe2 | 0.016 | 0.004 | 0.002 |
| Ctnnd2 | 0.016 | 0.009 | 0.030 |
| Macrod1 | 0.016 | 0.047 | 0.025 |

| | | | |
|---|---|---|---|
| Pcdhga2 | 0.016 | 0.024 | 0.021 |
| Cyp2j9 | 0.016 | 0.007 | 0.014 |
| Spaar | 0.016 | 0.002 | 0.003 |
| Elmo3 | 0.016 | 0.011 | 0.006 |
| Krt1 | 0.016 | 0.012 | 0.004 |
| Zik1 | 0.016 | 0.022 | 0.018 |
| Opn3 | 0.016 | 0.030 | 0.017 |
| Fam83d | 0.016 | 0.050 | 0.022 |
| Hist1h1b | 0.016 | 0.021 | 0.015 |
| P2rx1 | 0.016 | 0.003 | 0.001 |
| Nespas | 0.016 | 0.012 | 0.032 |
| Zfp11 | 0.016 | 0.014 | 0.017 |
| Gpr85 | 0.016 | 0.005 | 0.047 |
| Ska1 | 0.016 | 0.011 | 0.009 |
| Cyp2r1 | 0.016 | 0.006 | 0.013 |
| Camk2n2 | 0.016 | 0.031 | 0.028 |
| Gm14327 | 0.016 | 0.016 | 0.020 |
| 4932422M17 | 0.016 | 0.022 | 0.021 |
| Aard | 0.016 | 0.030 | 0.055 |
| Ino80dos | 0.016 | 0.021 | 0.018 |
| Nadsyn1 | 0.016 | 0.011 | 0.014 |
| Zranb3 | 0.016 | 0.019 | 0.014 |
| Gm11713 | 0.016 | 0.012 | 0.016 |
| Rnf157 | 0.016 | 0.056 | 0.056 |
| Gm16192 | 0.016 | 0.007 | 0.006 |
| Grin2d | 0.016 | 0.018 | 0.037 |
| Scamp5 | 0.016 | 0.023 | 0.052 |
| Cck | 0.016 | 0.143 | 0.014 |
| Naif1 | 0.016 | 0.016 | 0.016 |
| Ceacam10 | 0.016 | 0.000 | 0.000 |
| Krtcap3 | 0.016 | 0.005 | 0.003 |
| Gm12758 | 0.016 | 0.009 | 0.010 |
| AC174678.1 | 0.016 | 0.007 | 0.011 |
| Tox3 | 0.016 | 0.046 | 0.027 |
| AC153140.2 | 0.016 | 0.010 | 0.008 |
| Lix1 | 0.016 | 0.005 | 0.013 |
| Spag4 | 0.016 | 0.012 | 0.013 |
| Nebl | 0.016 | 0.050 | 0.008 |
| Gdf7 | 0.016 | 0.000 | 0.000 |
| Taf4b | 0.016 | 0.023 | 0.023 |
| Map10 | 0.016 | 0.022 | 0.022 |
| Fam185a | 0.016 | 0.024 | 0.022 |
| C3 | 0.016 | 0.027 | 0.136 |
| Gm9780 | 0.016 | 0.015 | 0.040 |
| Arhgap26 | 0.016 | 0.017 | 0.028 |
| Gm45351 | 0.016 | 0.009 | 0.009 |
| Gm28557 | 0.016 | 0.009 | 0.011 |
| Cst9 | 0.016 | 0.008 | 0.009 |

| | | | |
|---|---|---|---|
| Dynlt1a | 0.016 | 0.019 | 0.026 |
| Haspin | 0.016 | 0.004 | 0.004 |
| Sema3b | 0.015 | 0.058 | 0.092 |
| Zfp3 | 0.015 | 0.011 | 0.011 |
| 4930513N10I | 0.015 | 0.012 | 0.011 |
| Fhl3 | 0.015 | 0.038 | 0.022 |
| 2610318N02I | 0.015 | 0.000 | 0.001 |
| Samd15 | 0.015 | 0.015 | 0.011 |
| Cep78 | 0.015 | 0.037 | 0.035 |
| Dennd6b | 0.015 | 0.023 | 0.019 |
| Etv1 | 0.015 | 0.076 | 0.050 |
| 5730596B20F | 0.015 | 0.014 | 0.009 |
| Sdk2 | 0.015 | 0.077 | 0.125 |
| Cma1 | 0.015 | 0.014 | 0.015 |
| Tedc1 | 0.015 | 0.007 | 0.008 |
| Ppm1n | 0.015 | 0.053 | 0.075 |
| 2700012I20R | 0.015 | 0.006 | 0.005 |
| Pcdhgb5 | 0.015 | 0.020 | 0.017 |
| Ccdc122 | 0.015 | 0.011 | 0.026 |
| Nabp1 | 0.015 | 0.037 | 0.044 |
| Cdh10 | 0.015 | 0.009 | 0.004 |
| Mcm10 | 0.015 | 0.009 | 0.012 |
| Espl1 | 0.015 | 0.005 | 0.007 |
| Arhgap33 | 0.015 | 0.019 | 0.033 |
| Smpdl3b | 0.015 | 0.006 | 0.011 |
| Gm5577 | 0.015 | 0.010 | 0.010 |
| Zscan20 | 0.015 | 0.012 | 0.013 |
| Trim7 | 0.015 | 0.060 | 0.028 |
| Alox12 | 0.015 | 0.011 | 0.012 |
| Gchfr | 0.015 | 0.001 | 0.001 |
| 4930515G01I | 0.015 | 0.006 | 0.006 |
| Fam57b | 0.015 | 0.005 | 0.001 |
| Hist1h1d | 0.015 | 0.016 | 0.007 |
| Gm10687 | 0.015 | 0.027 | 0.004 |
| Gm10353 | 0.015 | 0.016 | 0.013 |
| Gng3 | 0.015 | 0.039 | 0.028 |
| Nmnat3 | 0.015 | 0.013 | 0.009 |
| Nr4a3 | 0.015 | 0.190 | 0.084 |
| Pnpla7 | 0.015 | 0.029 | 0.046 |
| Zfp93 | 0.015 | 0.016 | 0.015 |
| Mrc1 | 0.015 | 0.015 | 0.017 |
| Tmem231 | 0.015 | 0.038 | 0.029 |
| Whrn | 0.015 | 0.061 | 0.038 |
| Pim2 | 0.015 | 0.013 | 0.006 |
| Pcyt1b | 0.015 | 0.008 | 0.015 |
| Apln | 0.015 | 0.001 | 0.008 |
| Gm45716 | 0.015 | 0.107 | 0.026 |
| Sgo1 | 0.015 | 0.014 | 0.016 |

| | | | |
|---|---|---|---|
| Larp6 | 0.015 | 0.068 | 0.077 |
| H2-Q10 | 0.015 | 0.006 | 0.007 |
| Slc19a2 | 0.015 | 0.027 | 0.027 |
| Iqgap3 | 0.015 | 0.008 | 0.009 |
| Zfp169 | 0.015 | 0.008 | 0.008 |
| Gpr137c | 0.015 | 0.039 | 0.036 |
| Zfp850 | 0.015 | 0.010 | 0.007 |
| Mzf1 | 0.015 | 0.015 | 0.015 |
| 6720489N17I | 0.015 | 0.029 | 0.040 |
| Gm16751 | 0.015 | 0.009 | 0.008 |
| Trpm3 | 0.015 | 0.037 | 0.023 |
| BC049715 | 0.015 | 0.012 | 0.012 |
| Gm47689 | 0.015 | 0.016 | 0.011 |
| Pcdhb4 | 0.015 | 0.005 | 0.005 |
| 5430403G16I | 0.015 | 0.041 | 0.023 |
| Srp54c | 0.015 | 0.017 | 0.014 |
| Gm12258 | 0.015 | 0.016 | 0.024 |
| Gm9913 | 0.015 | 0.013 | 0.020 |
| Il2 | 0.015 | 0.021 | 0.025 |
| Wdr25 | 0.015 | 0.013 | 0.011 |
| Tmem136 | 0.015 | 0.058 | 0.046 |
| Fam83g | 0.015 | 0.001 | 0.001 |
| Cbarp | 0.015 | 0.118 | 0.057 |
| Tfb1m | 0.015 | 0.020 | 0.030 |
| Map3k13 | 0.015 | 0.008 | 0.008 |
| Esco2 | 0.015 | 0.008 | 0.007 |
| Tnk1 | 0.015 | 0.002 | 0.007 |
| Palb2 | 0.015 | 0.010 | 0.007 |
| Gm48960 | 0.015 | 0.007 | 0.006 |
| Gng10 | 0.015 | 0.023 | 0.034 |
| Ckm | 0.015 | 0.100 | 0.023 |
| Dnah1 | 0.015 | 0.009 | 0.029 |
| Rtl3 | 0.015 | 0.027 | 0.390 |
| Duox2 | 0.015 | 0.000 | 0.000 |
| Tfrc | 0.015 | 0.022 | 0.009 |
| Gm32618 | 0.015 | 0.047 | 0.038 |
| Cep72 | 0.015 | 0.014 | 0.015 |
| Lrfn1 | 0.015 | 0.021 | 0.027 |
| Gatm | 0.015 | 0.017 | 0.019 |
| 5-Sep | 0.015 | 0.044 | 0.042 |
| Zfp947 | 0.015 | 0.013 | 0.019 |
| Paqr5 | 0.014 | 0.001 | 0.001 |
| Tbc1d31 | 0.014 | 0.024 | 0.033 |
| Cmbl | 0.014 | 0.019 | 0.057 |
| Slc16a1 | 0.014 | 0.041 | 0.045 |
| Zim1 | 0.014 | 0.050 | 0.285 |
| Cpne7 | 0.014 | 0.030 | 0.039 |
| Prdm9 | 0.014 | 0.006 | 0.004 |

| | | | |
|---|---|---|---|
| Vwa3a | 0.014 | 0.037 | 0.016 |
| Cep128 | 0.014 | 0.014 | 0.021 |
| Gm11627 | 0.014 | 0.146 | 0.155 |
| Gm14418 | 0.014 | 0.027 | 0.034 |
| Tchh | 0.014 | 0.013 | 0.027 |
| B130006D01I | 0.014 | 0.016 | 0.018 |
| Enkur | 0.014 | 0.007 | 0.017 |
| 1700019L13F | 0.014 | 0.024 | 0.020 |
| Gm49077 | 0.014 | 0.014 | 0.018 |
| Ncapg | 0.014 | 0.012 | 0.012 |
| Scn2a | 0.014 | 0.056 | 0.052 |
| Gm15972 | 0.014 | 0.006 | 0.008 |
| Csrnp2 | 0.014 | 0.047 | 0.040 |
| Shisa2 | 0.014 | 0.012 | 0.010 |
| Efhb | 0.014 | 0.013 | 0.023 |
| Tmem221 | 0.014 | 0.000 | 0.000 |
| Dpep2 | 0.014 | 0.002 | 0.001 |
| Zkscan7 | 0.014 | 0.017 | 0.023 |
| Gm45250 | 0.014 | 0.012 | 0.013 |
| Zfp960 | 0.014 | 0.008 | 0.009 |
| Sgtb | 0.014 | 0.029 | 0.026 |
| Mns1 | 0.014 | 0.012 | 0.012 |
| Adam11 | 0.014 | 0.045 | 0.023 |
| Prkaa2 | 0.014 | 0.059 | 0.033 |
| 1010001B22F | 0.014 | 0.011 | 0.008 |
| Dagla | 0.014 | 0.015 | 0.028 |
| Gm34552 | 0.014 | 0.007 | 0.003 |
| Gm16861 | 0.014 | 0.010 | 0.008 |
| Gm15675 | 0.014 | 0.029 | 0.034 |
| Nova1 | 0.014 | 0.004 | 0.018 |
| Gpr157 | 0.014 | 0.011 | 0.011 |
| Vsig8 | 0.014 | 0.003 | 0.002 |
| Efhd1 | 0.014 | 0.040 | 0.055 |
| Rad9b | 0.014 | 0.019 | 0.025 |
| Ppp1r13l | 0.014 | 0.020 | 0.020 |
| Dscc1 | 0.014 | 0.017 | 0.030 |
| Mob3b | 0.014 | 0.070 | 0.028 |
| 6230400D17I | 0.014 | 0.014 | 0.021 |
| Prcd | 0.014 | 0.027 | 0.092 |
| Gm16141 | 0.014 | 0.035 | 0.034 |
| Ltf | 0.014 | 0.000 | 0.000 |
| Gm15853 | 0.014 | 0.010 | 0.005 |
| BC055324 | 0.014 | 0.009 | 0.013 |
| Adamts17 | 0.014 | 0.033 | 0.093 |
| Noxo1 | 0.014 | 0.012 | 0.021 |
| Apobec2 | 0.014 | 0.012 | 0.005 |
| Klk10 | 0.014 | 0.000 | 0.001 |
| Gpc1 | 0.014 | 0.111 | 0.069 |

| | | | |
|---|---|---|---|
| Gm32688 | 0.014 | 0.000 | 0.000 |
| Gucy1a2 | 0.014 | 0.026 | 0.017 |
| Xlr3b | 0.014 | 0.020 | 0.019 |
| Slc25a42 | 0.014 | 0.022 | 0.026 |
| Pde6h | 0.014 | 0.002 | 0.001 |
| Sv2a | 0.014 | 0.149 | 0.035 |
| Csf2rb | 0.014 | 0.002 | 0.005 |
| Dock8 | 0.014 | 0.034 | 0.011 |
| Sema3a | 0.014 | 0.010 | 0.051 |
| Zfp869 | 0.014 | 0.010 | 0.009 |
| Gm14698 | 0.014 | 0.014 | 0.013 |
| Gm14325 | 0.014 | 0.011 | 0.016 |
| Gm47469 | 0.014 | 0.009 | 0.014 |
| Pvt1 | 0.014 | 0.048 | 0.022 |
| Zfp563 | 0.014 | 0.014 | 0.019 |
| AC151836.1 | 0.014 | 0.011 | 0.007 |
| Gm9938 | 0.014 | 0.013 | 0.016 |
| Cftr | 0.014 | 0.001 | 0.000 |
| Gm3435 | 0.014 | 0.005 | 0.007 |
| Medag | 0.014 | 0.020 | 0.288 |
| Sapcd1 | 0.014 | 0.003 | 0.018 |
| Gm43573 | 0.014 | 0.010 | 0.005 |
| Ggnbp1 | 0.014 | 0.054 | 0.039 |
| A630001G21I | 0.014 | 0.004 | 0.002 |
| Ddc | 0.014 | 0.015 | 0.026 |
| Zfp583 | 0.014 | 0.026 | 0.025 |
| AC129328.1 | 0.014 | 0.020 | 0.026 |
| Hic1 | 0.014 | 0.029 | 0.355 |
| Pcdhga7 | 0.014 | 0.016 | 0.013 |
| Depdc7 | 0.014 | 0.008 | 0.010 |
| Spaca9 | 0.014 | 0.004 | 0.004 |
| Atcayos | 0.014 | 0.177 | 0.030 |
| Tert | 0.014 | 0.013 | 0.013 |
| Ect2 | 0.014 | 0.011 | 0.012 |
| Zfp882 | 0.014 | 0.019 | 0.020 |
| Mkrn3 | 0.014 | 0.007 | 0.010 |
| 4930524J08R | 0.013 | 0.018 | 0.020 |
| Klhl29 | 0.013 | 0.024 | 0.040 |
| Peg10 | 0.013 | 0.016 | 0.053 |
| Zfp791 | 0.013 | 0.018 | 0.014 |
| Cdh6 | 0.013 | 0.159 | 0.025 |
| Tcaf2 | 0.013 | 0.006 | 0.026 |
| Clmn | 0.013 | 0.107 | 0.122 |
| Ccdc106 | 0.013 | 0.092 | 0.066 |
| A930033H14I | 0.013 | 0.028 | 0.029 |
| Omd | 0.013 | 0.011 | 0.052 |
| Ntm | 0.013 | 0.020 | 0.174 |
| Raver2 | 0.013 | 0.028 | 0.033 |

| | | | |
|---|---|---|---|
| Gprin2 | 0.013 | 0.016 | 0.011 |
| Zfp953 | 0.013 | 0.007 | 0.010 |
| Chrm2 | 0.013 | 0.063 | 0.092 |
| Osbpl10 | 0.013 | 0.023 | 0.017 |
| Angel1 | 0.013 | 0.008 | 0.015 |
| Tcf23 | 0.013 | 0.001 | 0.003 |
| Pidd1 | 0.013 | 0.007 | 0.008 |
| Lrfn3 | 0.013 | 0.042 | 0.040 |
| Gm20467 | 0.013 | 0.000 | 0.000 |
| Gm17655 | 0.013 | 0.005 | 0.008 |
| Eef1akmt3 | 0.013 | 0.011 | 0.040 |
| Ulk4 | 0.013 | 0.040 | 0.037 |
| Efr3b | 0.013 | 0.051 | 0.020 |
| Fxyd7 | 0.013 | 0.027 | 0.023 |
| Gm20498 | 0.013 | 0.014 | 0.016 |
| Retreg1 | 0.013 | 0.077 | 0.115 |
| Ptpn13 | 0.013 | 0.049 | 0.116 |
| Gpt | 0.013 | 0.010 | 0.010 |
| Carns1 | 0.013 | 0.051 | 0.023 |
| Fam89a | 0.013 | 0.004 | 0.001 |
| Ttk | 0.013 | 0.007 | 0.009 |
| Rttn | 0.013 | 0.015 | 0.013 |
| 3830403N18I | 0.013 | 0.004 | 0.014 |
| Vash2 | 0.013 | 0.050 | 0.079 |
| Arl5c | 0.013 | 0.029 | 0.019 |
| Hyal3 | 0.013 | 0.015 | 0.016 |
| Ccdc92 | 0.013 | 0.006 | 0.003 |
| Eda2r | 0.013 | 0.008 | 0.027 |
| 2610020C07I | 0.013 | 0.026 | 0.026 |
| Angptl1 | 0.013 | 0.160 | 0.379 |
| Wdr78 | 0.013 | 0.010 | 0.016 |
| Ccl7 | 0.013 | 0.020 | 0.090 |
| Elovl4 | 0.013 | 0.048 | 0.046 |
| Cybrd1 | 0.013 | 0.025 | 0.027 |
| Gm10282 | 0.013 | 0.009 | 0.011 |
| Pnmal2 | 0.013 | 0.054 | 0.079 |
| Ric3 | 0.013 | 0.018 | 0.005 |
| Cdc25c | 0.013 | 0.005 | 0.007 |
| Spaca1 | 0.013 | 0.006 | 0.015 |
| AI661453 | 0.013 | 0.001 | 0.008 |
| Gm38186 | 0.013 | 0.008 | 0.013 |
| 9530077C05I | 0.013 | 0.034 | 0.027 |
| Sorl1 | 0.013 | 0.133 | 0.052 |
| Rtn4ip1 | 0.013 | 0.011 | 0.012 |
| Zfp14 | 0.013 | 0.013 | 0.010 |
| Spdl1 | 0.013 | 0.006 | 0.007 |
| Ero1lb | 0.013 | 0.015 | 0.017 |
| Pbx4 | 0.013 | 0.016 | 0.024 |

| | | | |
|---|---|---|---|
| Kcnmb1 | 0.013 | 0.181 | 0.015 |
| Kif7 | 0.013 | 0.035 | 0.039 |
| Gng8 | 0.013 | 0.071 | 0.127 |
| Runx2 | 0.013 | 0.013 | 0.015 |
| B930082K07F | 0.013 | 0.009 | 0.010 |
| Cobl | 0.013 | 0.123 | 0.038 |
| Saa3 | 0.013 | 0.021 | 0.026 |
| E230001N04F | 0.013 | 0.039 | 0.031 |
| Slc16a4 | 0.013 | 0.018 | 0.010 |
| Rassf6 | 0.013 | 0.002 | 0.001 |
| Gm43062 | 0.013 | 0.018 | 0.014 |
| Gm14508 | 0.013 | 0.003 | 0.003 |
| Kif18a | 0.013 | 0.020 | 0.022 |
| Gm20075 | 0.013 | 0.008 | 0.008 |
| Plcl1 | 0.013 | 0.095 | 0.088 |
| Tmco6 | 0.013 | 0.019 | 0.030 |
| A330069E16F | 0.013 | 0.010 | 0.012 |
| Efhc1 | 0.013 | 0.009 | 0.023 |
| Abhd15 | 0.013 | 0.002 | 0.003 |
| 3110001I22R | 0.013 | 0.020 | 0.018 |
| Gm3636 | 0.013 | 0.012 | 0.011 |
| Fancb | 0.013 | 0.008 | 0.028 |
| Mkln1os | 0.013 | 0.011 | 0.005 |
| 2600006K01F | 0.013 | 0.038 | 0.038 |
| Ppp1r1a | 0.013 | 0.126 | 0.060 |
| Rnls | 0.013 | 0.004 | 0.004 |
| Serpina1b | 0.013 | 0.000 | 0.000 |
| Kcnk7 | 0.013 | 0.006 | 0.007 |
| Gm26910 | 0.013 | 0.013 | 0.011 |
| Gm31135 | 0.013 | 0.002 | 0.000 |
| Nwd1 | 0.013 | 0.003 | 0.002 |
| Dnajc28 | 0.013 | 0.006 | 0.008 |
| Tmprss6 | 0.013 | 0.001 | 0.006 |
| Tm4sf4 | 0.013 | 0.001 | 0.001 |
| Nostrin | 0.013 | 0.002 | 0.009 |
| A630089N07I | 0.013 | 0.028 | 0.030 |
| Cyb5d1 | 0.013 | 0.010 | 0.011 |
| Bend3 | 0.013 | 0.022 | 0.016 |
| Gm28529 | 0.013 | 0.005 | 0.008 |
| B230322F03F | 0.013 | 0.007 | 0.007 |
| Smad9 | 0.013 | 0.049 | 0.025 |
| Nuak2 | 0.013 | 0.008 | 0.046 |
| 4631405J19R | 0.013 | 0.084 | 0.010 |
| 6330403L08F | 0.013 | 0.030 | 0.025 |
| Rbp4 | 0.013 | 0.017 | 0.211 |
| Pask | 0.013 | 0.009 | 0.012 |
| Zfp951 | 0.013 | 0.012 | 0.016 |
| Gm14286 | 0.013 | 0.008 | 0.010 |

| | | | |
|---|---|---|---|
| Pigw | 0.013 | 0.011 | 0.010 |
| Dnase1l2 | 0.013 | 0.006 | 0.010 |
| Tmem151a | 0.012 | 0.063 | 0.037 |
| Ppp1r3f | 0.012 | 0.019 | 0.021 |
| Gli1 | 0.012 | 0.030 | 0.193 |
| Gm20655 | 0.012 | 0.004 | 0.002 |
| Tec | 0.012 | 0.005 | 0.004 |
| Spag5 | 0.012 | 0.009 | 0.010 |
| G730003C15I | 0.012 | 0.006 | 0.007 |
| Xrcc2 | 0.012 | 0.009 | 0.015 |
| Rab3b | 0.012 | 0.027 | 0.010 |
| Slc37a2 | 0.012 | 0.006 | 0.007 |
| F420014N23F | 0.012 | 0.013 | 0.012 |
| Homer2 | 0.012 | 0.013 | 0.041 |
| Prox2 | 0.012 | 0.009 | 0.010 |
| Galnt3 | 0.012 | 0.001 | 0.000 |
| Gm29666 | 0.012 | 0.011 | 0.011 |
| 4930532G15I | 0.012 | 0.030 | 0.023 |
| 6030408B16F | 0.012 | 0.025 | 0.165 |
| Gm14399 | 0.012 | 0.009 | 0.009 |
| F830016B08F | 0.012 | 0.007 | 0.037 |
| Dmkn | 0.012 | 0.006 | 0.007 |
| Gm21992 | 0.012 | 0.016 | 0.019 |
| Neurl1b | 0.012 | 0.060 | 0.035 |
| Tdrkh | 0.012 | 0.003 | 0.006 |
| Myrf | 0.012 | 0.082 | 0.063 |
| Sec16b | 0.012 | 0.017 | 0.068 |
| Adm | 0.012 | 0.040 | 0.300 |
| Gm31834 | 0.012 | 0.002 | 0.003 |
| Tcte2 | 0.012 | 0.003 | 0.007 |
| Gm29367 | 0.012 | 0.004 | 0.003 |
| Nfe2 | 0.012 | 0.001 | 0.000 |
| Tubb4a | 0.012 | 0.124 | 0.110 |
| Gm45509 | 0.012 | 0.016 | 0.014 |
| Spsb4 | 0.012 | 0.107 | 0.104 |
| C1rl | 0.012 | 0.012 | 0.015 |
| Il18bp | 0.012 | 0.008 | 0.008 |
| 6030445D17I | 0.012 | 0.003 | 0.010 |
| Yjefn3 | 0.012 | 0.023 | 0.032 |
| Camk2b | 0.012 | 0.030 | 0.029 |
| Wnt9a | 0.012 | 0.049 | 0.120 |
| Gm31597 | 0.012 | 0.001 | 0.003 |
| Rtl5 | 0.012 | 0.036 | 0.028 |
| Slc9a5 | 0.012 | 0.011 | 0.018 |
| Stard6 | 0.012 | 0.008 | 0.007 |
| Amt | 0.012 | 0.021 | 0.010 |
| BC048403 | 0.012 | 0.008 | 0.008 |
| Zfp619 | 0.012 | 0.012 | 0.013 |

| | | | |
|---|---|---|---|
| Trim9 | 0.012 | 0.024 | 0.007 |
| Batf3 | 0.012 | 0.009 | 0.084 |
| Gm4070 | 0.012 | 0.000 | 0.000 |
| Gm48855 | 0.012 | 0.023 | 0.018 |
| Pcp4 | 0.012 | 0.090 | 0.016 |
| Eno1b | 0.012 | 0.006 | 0.006 |
| A230072E10F | 0.012 | 0.004 | 0.002 |
| Fam221a | 0.012 | 0.034 | 0.028 |
| Fam105a | 0.012 | 0.009 | 0.016 |
| Gm49313 | 0.012 | 0.001 | 0.002 |
| Zwilch | 0.012 | 0.012 | 0.013 |
| Zfp526 | 0.012 | 0.012 | 0.013 |
| Gm11772 | 0.012 | 0.009 | 0.008 |
| Tubg2 | 0.012 | 0.023 | 0.023 |
| Gbp6 | 0.012 | 0.002 | 0.007 |
| Gm26615 | 0.012 | 0.071 | 0.081 |
| Myo1h | 0.012 | 0.020 | 0.022 |
| Rassf4 | 0.012 | 0.063 | 0.028 |
| Bub1b | 0.012 | 0.018 | 0.017 |
| 2900009J06R | 0.012 | 0.014 | 0.015 |
| Slc7a11 | 0.012 | 0.007 | 0.006 |
| Ccdc65 | 0.012 | 0.003 | 0.004 |
| Pygm | 0.012 | 0.123 | 0.055 |
| 1700056N10I | 0.012 | 0.010 | 0.014 |
| Gm20661 | 0.012 | 0.011 | 0.011 |
| 2010015M23 | 0.012 | 0.009 | 0.007 |
| Gm47802 | 0.012 | 0.016 | 0.011 |
| Mok | 0.012 | 0.010 | 0.010 |
| Defb42 | 0.012 | 0.011 | 0.008 |
| Rprml | 0.012 | 0.001 | 0.002 |
| Fam72a | 0.012 | 0.015 | 0.016 |
| 0610039K10F | 0.012 | 0.005 | 0.004 |
| Gm7467 | 0.012 | 0.011 | 0.012 |
| Ccne1 | 0.012 | 0.026 | 0.050 |
| Ube2t | 0.012 | 0.020 | 0.010 |
| 4930404N11I | 0.012 | 0.014 | 0.015 |
| Frmd5 | 0.012 | 0.006 | 0.007 |
| Usp13 | 0.012 | 0.050 | 0.029 |
| Gm4950 | 0.012 | 0.008 | 0.007 |
| Ctse | 0.012 | 0.000 | 0.000 |
| Cdca5 | 0.012 | 0.006 | 0.007 |
| Gm10260 | 0.012 | 0.009 | 0.008 |
| D630036H23I | 0.012 | 0.008 | 0.007 |
| Mdm1 | 0.012 | 0.016 | 0.018 |
| Plekha7 | 0.012 | 0.008 | 0.005 |
| Gm31105 | 0.012 | 0.021 | 0.020 |
| Kctd7 | 0.012 | 0.016 | 0.021 |
| Srms | 0.012 | 0.000 | 0.000 |

| | | | |
|---|---|---|---|
| Zfp94 | 0.012 | 0.017 | 0.021 |
| Angpt1 | 0.012 | 0.008 | 0.026 |
| Gm6598 | 0.012 | 0.008 | 0.010 |
| Ydjc | 0.012 | 0.014 | 0.015 |
| Gm26545 | 0.012 | 0.015 | 0.017 |
| Pld4 | 0.012 | 0.007 | 0.006 |
| Cdca2 | 0.012 | 0.008 | 0.009 |
| Il1b | 0.012 | 0.009 | 0.007 |
| Arntl2 | 0.012 | 0.036 | 0.020 |
| Bbs12 | 0.012 | 0.019 | 0.018 |
| Kcna6 | 0.012 | 0.012 | 0.024 |
| Gfra2 | 0.012 | 0.018 | 0.108 |
| Mamdc4 | 0.012 | 0.005 | 0.010 |
| Cntnap1 | 0.012 | 0.008 | 0.010 |
| Popdc2 | 0.012 | 0.047 | 0.016 |
| Nudt15 | 0.012 | 0.011 | 0.010 |
| Atg9b | 0.012 | 0.000 | 0.000 |
| E430024P14F | 0.012 | 0.006 | 0.008 |
| Gm10320 | 0.012 | 0.010 | 0.013 |
| Lsamp | 0.012 | 0.006 | 0.053 |
| 4930426I24R | 0.011 | 0.018 | 0.012 |
| Gm11730 | 0.011 | 0.000 | 0.001 |
| AW112010 | 0.011 | 0.001 | 0.005 |
| Calca | 0.011 | 0.006 | 0.010 |
| Gm3604 | 0.011 | 0.014 | 0.018 |
| Ppp1r26 | 0.011 | 0.026 | 0.013 |
| Gm27010 | 0.011 | 0.035 | 0.039 |
| Mybl2 | 0.011 | 0.007 | 0.008 |
| Gm47438 | 0.011 | 0.015 | 0.012 |
| Shcbp1 | 0.011 | 0.015 | 0.017 |
| Nav3 | 0.011 | 0.015 | 0.122 |
| Lacc1 | 0.011 | 0.028 | 0.048 |
| Slc2a3 | 0.011 | 0.020 | 0.015 |
| Lhx8 | 0.011 | 0.000 | 0.022 |
| 4930570G19I | 0.011 | 0.003 | 0.008 |
| Lrrtm1 | 0.011 | 0.004 | 0.006 |
| Fam212b | 0.011 | 0.023 | 0.007 |
| Lrtm2 | 0.011 | 0.004 | 0.000 |
| Kcng2 | 0.011 | 0.005 | 0.019 |
| Myo16 | 0.011 | 0.011 | 0.009 |
| Kif2c | 0.011 | 0.008 | 0.008 |
| Hcls1 | 0.011 | 0.010 | 0.007 |
| Pparg | 0.011 | 0.005 | 0.039 |
| 9130011E15F | 0.011 | 0.014 | 0.013 |
| Chtf18 | 0.011 | 0.005 | 0.012 |
| Neil3 | 0.011 | 0.012 | 0.013 |
| Gas2l3 | 0.011 | 0.030 | 0.020 |
| Tmem71 | 0.011 | 0.003 | 0.001 |

| | | | |
|---|---|---|---|
| Mpp3 | 0.011 | 0.008 | 0.008 |
| A430105J06R | 0.011 | 0.006 | 0.008 |
| Chadl | 0.011 | 0.027 | 0.041 |
| Hyls1 | 0.011 | 0.013 | 0.012 |
| Dok7 | 0.011 | 0.002 | 0.003 |
| Lrrc73 | 0.011 | 0.019 | 0.029 |
| Stx1b | 0.011 | 0.028 | 0.038 |
| 9330182L06F | 0.011 | 0.037 | 0.014 |
| Cd84 | 0.011 | 0.001 | 0.001 |
| Ccdc57 | 0.011 | 0.012 | 0.015 |
| Ankrd13d | 0.011 | 0.023 | 0.012 |
| Ypel4 | 0.011 | 0.037 | 0.065 |
| Gm16386 | 0.011 | 0.006 | 0.008 |
| Gm11496 | 0.011 | 0.000 | 0.000 |
| Dennd2d | 0.011 | 0.001 | 0.001 |
| Dusp4 | 0.011 | 0.012 | 0.004 |
| Gm49322 | 0.011 | 0.005 | 0.008 |
| Dhx58 | 0.011 | 0.004 | 0.007 |
| Pmel | 0.011 | 0.009 | 0.025 |
| Dtx4 | 0.011 | 0.021 | 0.130 |
| Pcdhgb6 | 0.011 | 0.009 | 0.008 |
| C130050O18I | 0.011 | 0.015 | 0.003 |
| Gm20721 | 0.011 | 0.021 | 0.023 |
| Zfp983 | 0.011 | 0.012 | 0.013 |
| Tnfrsf13b | 0.011 | 0.001 | 0.001 |
| Fst | 0.011 | 0.034 | 0.386 |
| 9130604C24F | 0.011 | 0.006 | 0.006 |
| AI429214 | 0.011 | 0.005 | 0.007 |
| Dtx1 | 0.011 | 0.019 | 0.043 |
| Acot6 | 0.011 | 0.016 | 0.028 |
| Gm48551 | 0.011 | 0.012 | 0.011 |
| Adam30 | 0.011 | 0.020 | 0.026 |
| Adcy7 | 0.011 | 0.151 | 0.167 |
| Caprin2 | 0.011 | 0.061 | 0.080 |
| 1700056E22F | 0.011 | 0.022 | 0.028 |
| Mppe1 | 0.011 | 0.008 | 0.010 |
| Tfap2b | 0.011 | 0.346 | 0.084 |
| Pla2g5 | 0.011 | 0.001 | 0.000 |
| Efcab11 | 0.011 | 0.005 | 0.007 |
| Sh2b2 | 0.011 | 0.013 | 0.021 |
| Ch25h | 0.011 | 0.004 | 0.013 |
| Plin4 | 0.011 | 0.167 | 0.028 |
| Ccno | 0.011 | 0.032 | 0.036 |
| Htr2a | 0.011 | 0.002 | 0.006 |
| Aldh1l2 | 0.011 | 0.078 | 0.067 |
| Pcdhga11 | 0.011 | 0.009 | 0.009 |
| Gm36445 | 0.011 | 0.009 | 0.008 |
| A930015D03I | 0.011 | 0.005 | 0.003 |

| | | | |
|---|---|---|---|
| Alas2 | 0.011 | 0.005 | 0.007 |
| E2f8 | 0.011 | 0.004 | 0.005 |
| 9930111J21R | 0.011 | 0.001 | 0.000 |
| Xrcc3 | 0.011 | 0.012 | 0.015 |
| Rundc3a | 0.011 | 0.008 | 0.008 |
| Gm10600 | 0.011 | 0.110 | 0.143 |
| Gm32856 | 0.011 | 0.003 | 0.005 |
| Gm45844 | 0.011 | 0.014 | 0.012 |
| Adra2c | 0.011 | 0.006 | 0.001 |
| Ppfibp2 | 0.011 | 0.089 | 0.066 |
| Susd2 | 0.011 | 0.102 | 0.035 |
| Rab37 | 0.011 | 0.001 | 0.001 |
| Lnx1 | 0.011 | 0.040 | 0.028 |
| Grem1 | 0.011 | 0.204 | 0.046 |
| 4930480K23F | 0.011 | 0.005 | 0.005 |
| Ston2 | 0.011 | 0.015 | 0.076 |
| Rgs6 | 0.011 | 0.039 | 0.041 |
| Rapgefl1 | 0.011 | 0.011 | 0.016 |
| L2hgdh | 0.011 | 0.026 | 0.017 |
| Slc24a3 | 0.011 | 0.088 | 0.085 |
| Cpeb1 | 0.011 | 0.016 | 0.033 |
| D730045B01I | 0.011 | 0.008 | 0.008 |
| Brca1 | 0.011 | 0.007 | 0.011 |
| Stk32a | 0.011 | 0.003 | 0.009 |
| Itgal | 0.011 | 0.000 | 0.003 |
| Krt8 | 0.011 | 0.010 | 0.006 |
| Col20a1 | 0.011 | 0.003 | 0.008 |
| Vpreb1 | 0.011 | 0.004 | 0.006 |
| AY074887 | 0.011 | 0.012 | 0.012 |
| Gm29650 | 0.011 | 0.002 | 0.003 |
| Prelp | 0.011 | 0.090 | 0.116 |
| Gnal | 0.011 | 0.016 | 0.012 |
| Gm12454 | 0.011 | 0.032 | 0.013 |
| Gm43714 | 0.011 | 0.006 | 0.009 |
| D630039A03I | 0.011 | 0.007 | 0.001 |
| F13a1 | 0.011 | 0.020 | 0.029 |
| Slc24a4 | 0.011 | 0.176 | 0.018 |
| Cited1 | 0.011 | 0.040 | 0.036 |
| Tube1 | 0.011 | 0.007 | 0.012 |
| Pcdhb9 | 0.011 | 0.012 | 0.016 |
| Ddias | 0.011 | 0.007 | 0.009 |
| Pelo | 0.011 | 0.005 | 0.006 |
| Casc1 | 0.011 | 0.015 | 0.020 |
| Spata48 | 0.011 | 0.006 | 0.007 |
| Gm9885 | 0.011 | 0.136 | 0.142 |
| Clec4g | 0.011 | 0.000 | 0.000 |
| Lrp11 | 0.011 | 0.007 | 0.004 |
| Kcnd1 | 0.011 | 0.013 | 0.032 |

| | | | |
|---|---|---|---|
| Optc | 0.011 | 0.118 | 0.017 |
| Dusp14 | 0.011 | 0.048 | 0.028 |
| Brip1 | 0.010 | 0.023 | 0.031 |
| Gm10825 | 0.010 | 0.000 | 0.000 |
| Gm5784 | 0.010 | 0.010 | 0.007 |
| Upb1 | 0.010 | 0.017 | 0.018 |
| Slc10a6 | 0.010 | 0.031 | 0.108 |
| Nlgn1 | 0.010 | 0.050 | 0.088 |
| Gm13715 | 0.010 | 0.002 | 0.003 |
| Wdr66 | 0.010 | 0.006 | 0.016 |
| Dcdc2b | 0.010 | 0.016 | 0.014 |
| Fgf1 | 0.010 | 0.036 | 0.020 |
| Gfra4 | 0.010 | 0.039 | 0.026 |
| D830026I12R | 0.010 | 0.001 | 0.000 |
| Gm43796 | 0.010 | 0.007 | 0.005 |
| Itga10 | 0.010 | 0.094 | 0.031 |
| Gm10642 | 0.010 | 0.002 | 0.002 |
| Gm10734 | 0.010 | 0.003 | 0.005 |
| Pbld1 | 0.010 | 0.007 | 0.008 |
| Slc39a4 | 0.010 | 0.000 | 0.001 |
| Lrrc27 | 0.010 | 0.010 | 0.010 |
| Cdk18 | 0.010 | 0.001 | 0.003 |
| Ccdc74a | 0.010 | 0.026 | 0.046 |
| Fam161a | 0.010 | 0.019 | 0.024 |
| Zfp658 | 0.010 | 0.011 | 0.016 |
| Ccsap | 0.010 | 0.017 | 0.012 |
| Rad51c | 0.010 | 0.017 | 0.015 |
| 5430431A17F | 0.010 | 0.021 | 0.008 |
| Rab42 | 0.010 | 0.010 | 0.017 |
| Tmeff2 | 0.010 | 0.011 | 0.209 |
| Gm28187 | 0.010 | 0.001 | 0.000 |
| Ccdc120 | 0.010 | 0.010 | 0.007 |
| Cacna2d3 | 0.010 | 0.001 | 0.003 |
| Gcnt4 | 0.010 | 0.016 | 0.006 |
| Wdr62 | 0.010 | 0.007 | 0.011 |
| Arnt2 | 0.010 | 0.026 | 0.033 |
| Gm10369 | 0.010 | 0.009 | 0.051 |
| Tspan15 | 0.010 | 0.015 | 0.011 |
| D830035M03 | 0.010 | 0.005 | 0.007 |
| Zfp97 | 0.010 | 0.012 | 0.018 |
| Celf5 | 0.010 | 0.005 | 0.007 |
| Eps8l2 | 0.010 | 0.002 | 0.001 |
| Stk36 | 0.010 | 0.012 | 0.014 |
| Bub1 | 0.010 | 0.006 | 0.007 |
| Gdf9 | 0.010 | 0.008 | 0.013 |
| A930029G22I | 0.010 | 0.006 | 0.012 |
| Cacnb2 | 0.010 | 0.196 | 0.065 |
| Kdm8 | 0.010 | 0.008 | 0.007 |

| | | | |
|---|---|---|---|
| Etohd2 | 0.010 | 0.011 | 0.011 |
| Fn3krp | 0.010 | 0.010 | 0.010 |
| Ak5 | 0.010 | 0.005 | 0.028 |
| Gm26609 | 0.010 | 0.023 | 0.015 |
| 4831440E17F | 0.010 | 0.013 | 0.011 |
| Gm26945 | 0.010 | 0.006 | 0.031 |
| 2700016F22F | 0.010 | 0.011 | 0.008 |
| Prr11 | 0.010 | 0.005 | 0.005 |
| Fancd2 | 0.010 | 0.005 | 0.004 |
| Has1 | 0.010 | 0.011 | 0.153 |
| Spi1 | 0.010 | 0.008 | 0.008 |
| Gm6793 | 0.010 | 0.014 | 0.013 |
| Zfp993 | 0.010 | 0.003 | 0.008 |
| Zc2hc1c | 0.010 | 0.011 | 0.012 |
| Vwa7 | 0.010 | 0.000 | 0.000 |
| Ap5b1 | 0.010 | 0.006 | 0.007 |
| B3galt1 | 0.010 | 0.059 | 0.053 |
| Gm26670 | 0.010 | 0.013 | 0.015 |
| Gm47204 | 0.010 | 0.024 | 0.037 |
| Gpr156 | 0.010 | 0.005 | 0.003 |
| Nr3c2 | 0.010 | 0.007 | 0.009 |
| Hk1os | 0.010 | 0.010 | 0.011 |
| Adh6a | 0.010 | 0.000 | 0.000 |
| Lypd2 | 0.010 | 0.002 | 0.025 |
| Tssk4 | 0.010 | 0.004 | 0.005 |
| Gm16794 | 0.010 | 0.010 | 0.013 |
| 9530034E10F | 0.010 | 0.014 | 0.009 |
| Mme | 0.010 | 0.132 | 0.146 |
| Fsbp | 0.010 | 0.003 | 0.003 |
| Teddm2 | 0.010 | 0.014 | 0.013 |
| Nr2f1 | 0.010 | 0.049 | 0.139 |
| Epyc | 0.010 | 0.008 | 0.188 |
| Gm26882 | 0.010 | 0.014 | 0.013 |
| Kbtbd11 | 0.010 | 0.005 | 0.007 |
| Gm19325 | 0.010 | 0.030 | 0.022 |
| Gm13620 | 0.010 | 0.009 | 0.013 |
| Lrrtm2 | 0.010 | 0.016 | 0.015 |
| Tmem144 | 0.010 | 0.005 | 0.005 |
| Syt1 | 0.010 | 0.149 | 0.075 |
| Hoga1 | 0.010 | 0.005 | 0.016 |
| Alpk3 | 0.010 | 0.004 | 0.004 |
| Slc2a12 | 0.010 | 0.019 | 0.018 |
| Gm15614 | 0.010 | 0.050 | 0.038 |
| Ttc7 | 0.010 | 0.017 | 0.020 |
| Etaa1os | 0.010 | 0.009 | 0.018 |
| 9130008F23F | 0.010 | 0.022 | 0.020 |
| Pik3r6 | 0.010 | 0.002 | 0.002 |
| Gm15402 | 0.010 | 0.008 | 0.009 |

| | | | |
|---|---|---|---|
| Rbm11 | 0.010 | 0.008 | 0.009 |
| Zfp677 | 0.010 | 0.014 | 0.021 |
| Gpt2 | 0.010 | 0.026 | 0.032 |
| Susd5 | 0.010 | 0.155 | 0.010 |
| 1700066B19F | 0.010 | 0.001 | 0.001 |
| Plekhh1 | 0.010 | 0.002 | 0.001 |
| Pdf | 0.010 | 0.004 | 0.004 |
| Gm13010 | 0.010 | 0.008 | 0.001 |
| Itih4 | 0.010 | 0.087 | 0.018 |
| Dlgap5 | 0.010 | 0.012 | 0.010 |
| Zfp712 | 0.010 | 0.007 | 0.010 |
| Lcat | 0.010 | 0.012 | 0.023 |
| Syk | 0.010 | 0.013 | 0.011 |
| Lgi3 | 0.010 | 0.045 | 0.034 |
| Prelid3a | 0.010 | 0.038 | 0.009 |
| Gpr27 | 0.010 | 0.081 | 0.119 |
| Selplg | 0.010 | 0.013 | 0.008 |
| Zfp78 | 0.010 | 0.006 | 0.007 |
| Baiap2l1 | 0.010 | 0.044 | 0.020 |
| Oas1a | 0.010 | 0.001 | 0.002 |
| Ccnf | 0.010 | 0.006 | 0.007 |
| Gnaz | 0.010 | 0.013 | 0.017 |
| Ptchd1 | 0.010 | 0.001 | 0.007 |
| Kcnma1 | 0.010 | 0.242 | 0.071 |
| Gdpd2 | 0.010 | 0.026 | 0.046 |
| Cxcl13 | 0.010 | 0.110 | 0.055 |
| Ppm1e | 0.010 | 0.083 | 0.024 |
| Gm26809 | 0.010 | 0.062 | 0.027 |
| Gm49207 | 0.010 | 0.011 | 0.011 |
| Mitf | 0.010 | 0.047 | 0.032 |
| Zfp365 | 0.010 | 0.041 | 0.035 |
| Usp51 | 0.010 | 0.009 | 0.007 |
| Aurkb | 0.010 | 0.014 | 0.015 |
| B3gnt2 | 0.010 | 0.014 | 0.005 |
| Gm15998 | 0.010 | 0.036 | 0.007 |
| Rwdd3 | 0.010 | 0.020 | 0.018 |
| Gm6288 | 0.010 | 0.002 | 0.002 |
| Pyroxd2 | 0.010 | 0.007 | 0.030 |
| Rsad2 | 0.010 | 0.003 | 0.003 |
| Trbc2 | 0.010 | 0.002 | 0.001 |
| Sgcg | 0.010 | 0.238 | 0.026 |
| Hist2h4 | 0.010 | 0.014 | 0.015 |
| Paqr3 | 0.010 | 0.012 | 0.011 |
| Asic1 | 0.010 | 0.002 | 0.014 |
| 1700041G16I | 0.010 | 0.003 | 0.003 |
| Gm28731 | 0.010 | 0.005 | 0.003 |
| Kctd14 | 0.010 | 0.003 | 0.006 |
| Dpyd | 0.010 | 0.015 | 0.036 |

| | | | |
|---|---|---|---|
| Glra1 | 0.010 | 0.017 | 0.018 |
| 9530026P05F | 0.010 | 0.038 | 0.029 |
| Gm19590 | 0.010 | 0.000 | 0.000 |
| 4930447C04F | 0.010 | 0.027 | 0.045 |
| 2810457G06I | 0.010 | 0.037 | 0.022 |
| Acta1 | 0.010 | 0.010 | 0.004 |
| Pcdhb3 | 0.010 | 0.007 | 0.009 |
| Mnd1 | 0.010 | 0.035 | 0.085 |
| Exo1 | 0.010 | 0.007 | 0.007 |
| Gata5 | 0.009 | 0.008 | 0.004 |
| H2-DMb1 | 0.009 | 0.015 | 0.013 |
| Acr | 0.009 | 0.000 | 0.001 |
| Ccser1 | 0.009 | 0.002 | 0.003 |
| Gm26733 | 0.009 | 0.016 | 0.011 |
| Gm14966 | 0.009 | 0.011 | 0.011 |
| Susd1 | 0.009 | 0.002 | 0.002 |
| Gm10516 | 0.009 | 0.005 | 0.005 |
| Hrc | 0.009 | 0.150 | 0.026 |
| Smim10l2a | 0.009 | 0.015 | 0.010 |
| 4930403P22F | 0.009 | 0.059 | 0.073 |
| Gm16152 | 0.009 | 0.032 | 0.071 |
| Zfp366 | 0.009 | 0.001 | 0.001 |
| Gpr35 | 0.009 | 0.001 | 0.004 |
| Efna3 | 0.009 | 0.024 | 0.011 |
| Igsf9 | 0.009 | 0.035 | 0.020 |
| Arsi | 0.009 | 0.028 | 0.201 |
| Degs2 | 0.009 | 0.001 | 0.001 |
| Wscd2 | 0.009 | 0.118 | 0.050 |
| Tekt2 | 0.009 | 0.035 | 0.021 |
| 1700001O22I | 0.009 | 0.007 | 0.005 |
| Chrm3 | 0.009 | 0.008 | 0.006 |
| Pih1d2 | 0.009 | 0.006 | 0.009 |
| Gm48914 | 0.009 | 0.016 | 0.010 |
| Cmya5 | 0.009 | 0.060 | 0.044 |
| A830035O19I | 0.009 | 0.008 | 0.009 |
| C2 | 0.009 | 0.007 | 0.074 |
| Ms4a6d | 0.009 | 0.004 | 0.006 |
| A430106G13I | 0.009 | 0.019 | 0.030 |
| Sh3bgrl2 | 0.009 | 0.002 | 0.007 |
| Glipr1 | 0.009 | 0.212 | 0.053 |
| Cenpu | 0.009 | 0.013 | 0.013 |
| Gm28874 | 0.009 | 0.006 | 0.009 |
| Sgsm1 | 0.009 | 0.003 | 0.004 |
| Proser2 | 0.009 | 0.019 | 0.011 |
| 2810405F17F | 0.009 | 0.005 | 0.010 |
| Kcnip3 | 0.009 | 0.015 | 0.037 |
| Zfp808 | 0.009 | 0.008 | 0.009 |
| C030037D09I | 0.009 | 0.008 | 0.015 |

| | | | |
|---|---|---|---|
| Il11 | 0.009 | 0.006 | 0.007 |
| Six3 | 0.009 | 0.000 | 0.000 |
| Lpar2 | 0.009 | 0.012 | 0.009 |
| Rims1 | 0.009 | 0.029 | 0.016 |
| Pcp4l1 | 0.009 | 0.068 | 0.012 |
| Arc | 0.009 | 0.141 | 0.042 |
| Oip5 | 0.009 | 0.004 | 0.006 |
| Ccl12 | 0.009 | 0.007 | 0.008 |
| Dclk2 | 0.009 | 0.022 | 0.016 |
| Tmem81 | 0.009 | 0.007 | 0.005 |
| Gm13205 | 0.009 | 0.017 | 0.007 |
| Gm7160 | 0.009 | 0.008 | 0.008 |
| Phtf1os | 0.009 | 0.008 | 0.015 |
| Syt12 | 0.009 | 0.018 | 0.006 |
| Rnf32 | 0.009 | 0.003 | 0.003 |
| Cdkl2 | 0.009 | 0.028 | 0.017 |
| Il1rl2 | 0.009 | 0.009 | 0.053 |
| Masp1 | 0.009 | 0.005 | 0.035 |
| Fhdc1 | 0.009 | 0.026 | 0.022 |
| Abcb9 | 0.009 | 0.024 | 0.010 |
| Chn1os3 | 0.009 | 0.019 | 0.016 |
| Gm26722 | 0.009 | 0.062 | 0.035 |
| Cux2 | 0.009 | 0.049 | 0.023 |
| Otogl | 0.009 | 0.049 | 0.008 |
| Kcnk10 | 0.009 | 0.000 | 0.000 |
| Immp2l | 0.009 | 0.012 | 0.011 |
| Cage1 | 0.009 | 0.008 | 0.008 |
| Hist1h3d | 0.009 | 0.015 | 0.008 |
| Ms4a7 | 0.009 | 0.012 | 0.009 |
| Gvin1 | 0.009 | 0.000 | 0.000 |
| Gm7324 | 0.009 | 0.018 | 0.020 |
| Isg20 | 0.009 | 0.007 | 0.013 |
| Piezo2 | 0.009 | 0.100 | 0.148 |
| Galnt5 | 0.009 | 0.002 | 0.015 |
| Tmem106a | 0.009 | 0.012 | 0.023 |
| Gm13713 | 0.009 | 0.001 | 0.002 |
| Slc15a3 | 0.009 | 0.006 | 0.003 |
| AC163040.1 | 0.009 | 0.005 | 0.005 |
| Gm44026 | 0.009 | 0.000 | 0.000 |
| Kazn | 0.009 | 0.030 | 0.081 |
| Gm16083 | 0.009 | 0.004 | 0.004 |
| Gm10658 | 0.009 | 0.005 | 0.005 |
| Cubn | 0.009 | 0.004 | 0.004 |
| Gm12355 | 0.009 | 0.012 | 0.011 |
| Ccdc17 | 0.009 | 0.006 | 0.006 |
| Slco1a4 | 0.009 | 0.001 | 0.003 |
| Mgat3 | 0.009 | 0.019 | 0.021 |
| Hcn4 | 0.009 | 0.002 | 0.002 |

| | | | |
|---|---|---|---|
| Gm10605 | 0.009 | 0.004 | 0.044 |
| Mrln | 0.009 | 0.013 | 0.009 |
| Zfp970 | 0.009 | 0.007 | 0.012 |
| Xkr5 | 0.009 | 0.005 | 0.003 |
| AC164883.2 | 0.009 | 0.003 | 0.002 |
| Adamts3 | 0.009 | 0.009 | 0.013 |
| 9530059O14I | 0.009 | 0.001 | 0.001 |
| Egf | 0.009 | 0.010 | 0.013 |
| Camkmt | 0.009 | 0.007 | 0.006 |
| Wfikkn2 | 0.009 | 0.035 | 0.057 |
| Gm30108 | 0.009 | 0.011 | 0.007 |
| Ly6g6e | 0.009 | 0.000 | 0.000 |
| Adgrg2 | 0.009 | 0.015 | 0.053 |
| Troap | 0.009 | 0.004 | 0.006 |
| Kcne1l | 0.009 | 0.111 | 0.127 |
| Gm27030 | 0.009 | 0.011 | 0.017 |
| Gm11696 | 0.009 | 0.008 | 0.006 |
| Gm10037 | 0.009 | 0.013 | 0.008 |
| Naalad2 | 0.009 | 0.018 | 0.130 |
| H2-DMa | 0.009 | 0.012 | 0.012 |
| Pfkfb1 | 0.009 | 0.008 | 0.008 |
| Plcl2 | 0.009 | 0.004 | 0.039 |
| Adgre1 | 0.009 | 0.006 | 0.005 |
| Car12 | 0.009 | 0.006 | 0.010 |
| 1700029J07R | 0.009 | 0.033 | 0.047 |
| Myl6b | 0.009 | 0.010 | 0.005 |
| Fam131c | 0.009 | 0.004 | 0.004 |
| Smim5 | 0.009 | 0.041 | 0.025 |
| Zdhhc23 | 0.009 | 0.033 | 0.014 |
| Grid1 | 0.009 | 0.000 | 0.001 |
| Ankef1 | 0.009 | 0.001 | 0.003 |
| Rbm3os | 0.009 | 0.003 | 0.007 |
| 1110020A21I | 0.009 | 0.009 | 0.008 |
| Glcci1 | 0.009 | 0.020 | 0.012 |
| 4833445I07R | 0.009 | 0.006 | 0.007 |
| Pgr | 0.009 | 0.034 | 0.016 |
| 2310030G06I | 0.009 | 0.050 | 0.021 |
| Gpr135 | 0.009 | 0.023 | 0.049 |
| A730011C13I | 0.009 | 0.006 | 0.003 |
| Rorb | 0.009 | 0.003 | 0.004 |
| Chac1 | 0.009 | 0.005 | 0.002 |
| C1s1 | 0.009 | 0.038 | 0.210 |
| Irak3 | 0.009 | 0.010 | 0.076 |
| Slc22a4 | 0.009 | 0.004 | 0.002 |
| Ccdc159 | 0.009 | 0.005 | 0.008 |
| Gm40841 | 0.009 | 0.007 | 0.011 |
| Slurp1 | 0.009 | 0.005 | 0.008 |
| Ifit3b | 0.009 | 0.002 | 0.007 |

| | | | |
|---|---|---|---|
| Cfap54 | 0.009 | 0.000 | 0.000 |
| Grin3a | 0.009 | 0.010 | 0.028 |
| Tmem56 | 0.009 | 0.004 | 0.004 |
| Pabpc1l | 0.009 | 0.004 | 0.004 |
| Lpin3 | 0.009 | 0.005 | 0.015 |
| Fbxo40 | 0.009 | 0.058 | 0.021 |
| Gm43254 | 0.009 | 0.009 | 0.011 |
| Gm26935 | 0.009 | 0.018 | 0.018 |
| Npy1r | 0.009 | 0.076 | 0.019 |
| Pla2g7 | 0.009 | 0.004 | 0.007 |
| 4930525G20I | 0.009 | 0.002 | 0.004 |
| Igf2os | 0.009 | 0.005 | 0.020 |
| Fgr | 0.009 | 0.007 | 0.003 |
| Depdc1a | 0.009 | 0.009 | 0.008 |
| Adam23 | 0.009 | 0.030 | 0.071 |
| Gm36660 | 0.009 | 0.004 | 0.001 |
| Mybpc3 | 0.009 | 0.009 | 0.012 |
| Rerg | 0.009 | 0.030 | 0.073 |
| Gm20878 | 0.009 | 0.020 | 0.019 |
| Spta1 | 0.008 | 0.000 | 0.000 |
| Frmd3 | 0.008 | 0.003 | 0.014 |
| Zfp940 | 0.008 | 0.014 | 0.009 |
| Tpbgl | 0.008 | 0.004 | 0.065 |
| Ddx60 | 0.008 | 0.003 | 0.010 |
| Scube1 | 0.008 | 0.059 | 0.061 |
| Mboat1 | 0.008 | 0.041 | 0.020 |
| Micalcl | 0.008 | 0.011 | 0.006 |
| Lag3 | 0.008 | 0.029 | 0.041 |
| C430049E01F | 0.008 | 0.000 | 0.000 |
| Cemip | 0.008 | 0.005 | 0.017 |
| Tmem150c | 0.008 | 0.001 | 0.004 |
| Hecw2 | 0.008 | 0.025 | 0.026 |
| Gm16070 | 0.008 | 0.011 | 0.002 |
| Loxl4 | 0.008 | 0.019 | 0.009 |
| Barx2 | 0.008 | 0.000 | 0.000 |
| Smc2os | 0.008 | 0.002 | 0.003 |
| 2010007H06I | 0.008 | 0.004 | 0.003 |
| Gm20522 | 0.008 | 0.011 | 0.011 |
| Pramef12 | 0.008 | 0.002 | 0.002 |
| Tonsl | 0.008 | 0.011 | 0.015 |
| Gm13594 | 0.008 | 0.016 | 0.004 |
| Gm15013 | 0.008 | 0.005 | 0.009 |
| Ppfia3 | 0.008 | 0.017 | 0.014 |
| Ccdc69 | 0.008 | 0.028 | 0.007 |
| Ccdc146 | 0.008 | 0.013 | 0.006 |
| F3 | 0.008 | 0.245 | 0.136 |
| Erp27 | 0.008 | 0.006 | 0.004 |
| Gm15892 | 0.008 | 0.026 | 0.024 |

| | | | |
|---|---|---|---|
| Shisa9 | 0.008 | 0.088 | 0.106 |
| Gm26847 | 0.008 | 0.003 | 0.004 |
| Rab39b | 0.008 | 0.025 | 0.023 |
| Ces2b | 0.008 | 0.000 | 0.000 |
| Smim6 | 0.008 | 0.002 | 0.001 |
| Pcdhga5 | 0.008 | 0.017 | 0.014 |
| Hist1h3e | 0.008 | 0.007 | 0.005 |
| A330035P11F | 0.008 | 0.006 | 0.005 |
| Ccl4 | 0.008 | 0.016 | 0.012 |
| Lyrm7 | 0.008 | 0.011 | 0.013 |
| Hmga1b | 0.008 | 0.007 | 0.003 |
| Gm16365 | 0.008 | 0.010 | 0.031 |
| Ablim2 | 0.008 | 0.001 | 0.003 |
| Slc9b1 | 0.008 | 0.001 | 0.001 |
| Adssl1 | 0.008 | 0.068 | 0.012 |
| Zfp469 | 0.008 | 0.020 | 0.248 |
| Mannr | 0.008 | 0.000 | 0.000 |
| Gm40604 | 0.008 | 0.000 | 0.000 |
| Cadm4 | 0.008 | 0.009 | 0.010 |
| Rbm24 | 0.008 | 0.211 | 0.054 |
| Rhbdl3 | 0.008 | 0.024 | 0.052 |
| Gm12802 | 0.008 | 0.014 | 0.010 |
| Tmc8 | 0.008 | 0.001 | 0.001 |
| Slc25a21 | 0.008 | 0.004 | 0.005 |
| Med12l | 0.008 | 0.026 | 0.012 |
| Kifc1 | 0.008 | 0.009 | 0.011 |
| Dgkg | 0.008 | 0.150 | 0.012 |
| Krt7 | 0.008 | 0.009 | 0.012 |
| Ehhadh | 0.008 | 0.011 | 0.007 |
| Gm28723 | 0.008 | 0.004 | 0.012 |
| Gm16133 | 0.008 | 0.022 | 0.011 |
| Rnf128 | 0.008 | 0.031 | 0.012 |
| B930094E09F | 0.008 | 0.003 | 0.005 |
| Ttc39a | 0.008 | 0.004 | 0.011 |
| 4930555A03F | 0.008 | 0.000 | 0.000 |
| Gm17477 | 0.008 | 0.009 | 0.009 |
| Gm28800 | 0.008 | 0.001 | 0.001 |
| AC122413.1 | 0.008 | 0.014 | 0.011 |
| Gm26510 | 0.008 | 0.000 | 0.000 |
| Gm39822 | 0.008 | 0.000 | 0.000 |
| Pcdh11x | 0.008 | 0.084 | 0.093 |
| Gm37469 | 0.008 | 0.000 | 0.000 |
| Rundc3b | 0.008 | 0.003 | 0.005 |
| Gm567 | 0.008 | 0.000 | 0.000 |
| 4931440F15F | 0.008 | 0.001 | 0.002 |
| Slc46a2 | 0.008 | 0.011 | 0.011 |
| Gm37233 | 0.008 | 0.014 | 0.030 |
| Gnao1 | 0.008 | 0.050 | 0.082 |

| | | | |
|---|---|---|---|
| Gm7334 | 0.008 | 0.010 | 0.006 |
| Tcp11 | 0.008 | 0.004 | 0.007 |
| Opcml | 0.008 | 0.002 | 0.056 |
| Gm26511 | 0.008 | 0.012 | 0.014 |
| Wnt7a | 0.008 | 0.000 | 0.000 |
| Gm20939 | 0.008 | 0.003 | 0.004 |
| Apobr | 0.008 | 0.012 | 0.016 |
| Gm49097 | 0.008 | 0.003 | 0.004 |
| Gm47906 | 0.008 | 0.000 | 0.000 |
| Gm48045 | 0.008 | 0.011 | 0.010 |
| Gm16897 | 0.008 | 0.001 | 0.001 |
| Tmem147os | 0.008 | 0.008 | 0.007 |
| Arhgap27os2 | 0.008 | 0.003 | 0.009 |
| Zfp108 | 0.008 | 0.009 | 0.009 |
| Ctss | 0.008 | 0.014 | 0.012 |
| Serpina1d | 0.008 | 0.000 | 0.000 |
| Acss1 | 0.008 | 0.026 | 0.008 |
| Zbtb3 | 0.008 | 0.008 | 0.007 |
| Matk | 0.008 | 0.011 | 0.011 |
| Wnk3 | 0.008 | 0.013 | 0.018 |
| 4430402I18R | 0.008 | 0.019 | 0.011 |
| Acap1 | 0.008 | 0.001 | 0.001 |
| Wnt10b | 0.008 | 0.001 | 0.030 |
| Bard1 | 0.008 | 0.005 | 0.005 |
| Fam161b | 0.008 | 0.009 | 0.009 |
| Ust | 0.008 | 0.057 | 0.081 |
| Gm10273 | 0.008 | 0.005 | 0.005 |
| B4galnt1 | 0.008 | 0.013 | 0.025 |
| Cds1 | 0.008 | 0.039 | 0.008 |
| Gm20219 | 0.008 | 0.014 | 0.009 |
| Arhgef16 | 0.008 | 0.003 | 0.002 |
| Gm16287 | 0.008 | 0.004 | 0.008 |
| Setd4 | 0.008 | 0.016 | 0.027 |
| Tmem255a | 0.008 | 0.003 | 0.011 |
| Parvaos | 0.008 | 0.009 | 0.012 |
| Ncmap | 0.008 | 0.008 | 0.006 |
| Ccdc3 | 0.008 | 0.040 | 0.027 |
| Srl | 0.008 | 0.011 | 0.008 |
| Gm45495 | 0.008 | 0.004 | 0.003 |
| Gm17669 | 0.008 | 0.006 | 0.004 |
| AC124739.2 | 0.008 | 0.009 | 0.006 |
| Apol9b | 0.008 | 0.003 | 0.003 |
| Astn2 | 0.008 | 0.159 | 0.039 |
| 9030612E09F | 0.008 | 0.015 | 0.012 |
| Rmi2 | 0.008 | 0.008 | 0.019 |
| Caskin1 | 0.008 | 0.004 | 0.003 |
| Sox10 | 0.008 | 0.005 | 0.006 |
| Rdh12 | 0.008 | 0.008 | 0.004 |

| | | | |
|---|---|---|---|
| Tcf21 | 0.008 | 0.022 | 0.221 |
| Klhdc9 | 0.008 | 0.008 | 0.008 |
| Bmp15 | 0.008 | 0.001 | 0.000 |
| Adam5 | 0.008 | 0.005 | 0.014 |
| Hist2h3c1 | 0.008 | 0.012 | 0.005 |
| Gm17146 | 0.008 | 0.003 | 0.002 |
| Dgki | 0.008 | 0.010 | 0.005 |
| Bcl6 | 0.008 | 0.050 | 0.036 |
| Star | 0.008 | 0.015 | 0.009 |
| Gm12976 | 0.008 | 0.003 | 0.002 |
| Gm3716 | 0.008 | 0.000 | 0.001 |
| 1700055D18I | 0.008 | 0.005 | 0.006 |
| Nfe2l3 | 0.008 | 0.011 | 0.016 |
| 2310040G24I | 0.008 | 0.015 | 0.011 |
| Car9 | 0.008 | 0.015 | 0.004 |
| Foxl1 | 0.008 | 0.025 | 0.013 |
| Fn3k | 0.008 | 0.008 | 0.003 |
| Gm28050 | 0.008 | 0.007 | 0.006 |
| Slc37a1 | 0.008 | 0.004 | 0.003 |
| Slco5a1 | 0.008 | 0.014 | 0.011 |
| Gpam | 0.008 | 0.004 | 0.006 |
| Tnfsf15 | 0.008 | 0.008 | 0.002 |
| Zbp1 | 0.008 | 0.019 | 0.019 |
| Zfp973 | 0.008 | 0.007 | 0.010 |
| Fcgr3 | 0.008 | 0.010 | 0.013 |
| Pik3cd | 0.008 | 0.022 | 0.025 |
| Polh | 0.008 | 0.005 | 0.007 |
| Defb25 | 0.008 | 0.016 | 0.020 |
| Syt17 | 0.008 | 0.002 | 0.003 |
| Gm43352 | 0.008 | 0.000 | 0.000 |
| Gm45076 | 0.008 | 0.009 | 0.001 |
| Slfn3 | 0.008 | 0.001 | 0.003 |
| Gm26759 | 0.008 | 0.010 | 0.007 |
| Gm14391 | 0.008 | 0.004 | 0.006 |
| Fap | 0.008 | 0.099 | 0.367 |
| Rad51b | 0.008 | 0.006 | 0.030 |
| Gm48623 | 0.008 | 0.008 | 0.006 |
| Kif9 | 0.008 | 0.003 | 0.005 |
| Art3 | 0.008 | 0.055 | 0.014 |
| Cacng4 | 0.008 | 0.018 | 0.021 |
| Sox6os | 0.008 | 0.004 | 0.001 |
| Actr3b | 0.008 | 0.040 | 0.043 |
| Cd14 | 0.008 | 0.016 | 0.015 |
| Sfrp4 | 0.008 | 0.006 | 0.024 |
| Gm48420 | 0.007 | 0.004 | 0.005 |
| H2-M5 | 0.007 | 0.005 | 0.011 |
| Zfp433 | 0.007 | 0.005 | 0.004 |
| Lss | 0.007 | 0.041 | 0.036 |

| | | | |
|---|---|---|---|
| Ciart | 0.007 | 0.014 | 0.008 |
| Mcm8 | 0.007 | 0.009 | 0.009 |
| AC163633.1 | 0.007 | 0.004 | 0.005 |
| Ldhd | 0.007 | 0.014 | 0.008 |
| 2310034G01I | 0.007 | 0.012 | 0.010 |
| Tpsb2 | 0.007 | 0.006 | 0.007 |
| Fmod | 0.007 | 0.044 | 0.142 |
| Upk1a | 0.007 | 0.005 | 0.004 |
| Rufy4 | 0.007 | 0.003 | 0.003 |
| Ltb | 0.007 | 0.002 | 0.002 |
| Melk | 0.007 | 0.009 | 0.010 |
| Hsd11b1 | 0.007 | 0.010 | 0.037 |
| Gbp8 | 0.007 | 0.000 | 0.000 |
| Crabp1 | 0.007 | 0.005 | 0.059 |
| Col8a2 | 0.007 | 0.076 | 0.129 |
| Pou2f2 | 0.007 | 0.003 | 0.003 |
| Paqr6 | 0.007 | 0.091 | 0.109 |
| Fras1 | 0.007 | 0.083 | 0.064 |
| Abca9 | 0.007 | 0.008 | 0.081 |
| Nme3 | 0.007 | 0.010 | 0.006 |
| Gm30648 | 0.007 | 0.000 | 0.003 |
| Mab21l2 | 0.007 | 0.221 | 0.078 |
| Usp43 | 0.007 | 0.001 | 0.002 |
| Syne3 | 0.007 | 0.025 | 0.031 |
| Fam198a | 0.007 | 0.020 | 0.096 |
| 1700102H20I | 0.007 | 0.010 | 0.005 |
| Lyzl4 | 0.007 | 0.005 | 0.000 |
| Ubash3b | 0.007 | 0.005 | 0.024 |
| Gm36839 | 0.007 | 0.014 | 0.012 |
| Ube2cbp | 0.007 | 0.010 | 0.011 |
| L1cam | 0.007 | 0.003 | 0.003 |
| Slc6a17 | 0.007 | 0.063 | 0.020 |
| Mcpt4 | 0.007 | 0.009 | 0.007 |
| Hsd3b7 | 0.007 | 0.003 | 0.004 |
| Gm15537 | 0.007 | 0.004 | 0.004 |
| Mpeg1 | 0.007 | 0.002 | 0.001 |
| Dpp4 | 0.007 | 0.003 | 0.040 |
| Cpm | 0.007 | 0.044 | 0.011 |
| Zfp59 | 0.007 | 0.008 | 0.007 |
| Dnaic1 | 0.007 | 0.000 | 0.001 |
| Gm2237 | 0.007 | 0.013 | 0.009 |
| Rad51ap2 | 0.007 | 0.023 | 0.008 |
| Pcdhb2 | 0.007 | 0.002 | 0.003 |
| Dmrta2 | 0.007 | 0.009 | 0.008 |
| Ptpn6 | 0.007 | 0.004 | 0.004 |
| Cnih2 | 0.007 | 0.020 | 0.038 |
| Gm3448 | 0.007 | 0.019 | 0.029 |
| Gm48796 | 0.007 | 0.007 | 0.006 |

| | | | |
|---|---|---|---|
| Gm38832 | 0.007 | 0.003 | 0.002 |
| Arhgef39 | 0.007 | 0.020 | 0.008 |
| Lzts3 | 0.007 | 0.014 | 0.012 |
| Oit3 | 0.007 | 0.001 | 0.002 |
| Cd300c2 | 0.007 | 0.004 | 0.004 |
| Rbm8a2 | 0.007 | 0.006 | 0.004 |
| Slc7a14 | 0.007 | 0.002 | 0.002 |
| Calb2 | 0.007 | 0.001 | 0.000 |
| Ppargc1b | 0.007 | 0.017 | 0.009 |
| Nipal1 | 0.007 | 0.012 | 0.005 |
| Apoa2 | 0.007 | 0.004 | 0.005 |
| Kctd21 | 0.007 | 0.007 | 0.014 |
| Gm15133 | 0.007 | 0.001 | 0.004 |
| Rhd | 0.007 | 0.001 | 0.010 |
| Nhlrc1 | 0.007 | 0.009 | 0.006 |
| Ssc4d | 0.007 | 0.004 | 0.006 |
| Gm26797 | 0.007 | 0.014 | 0.014 |
| Kcnmb4os2 | 0.007 | 0.016 | 0.005 |
| Gm17103 | 0.007 | 0.003 | 0.003 |
| 5830454E08R | 0.007 | 0.007 | 0.009 |
| Gm5086 | 0.007 | 0.000 | 0.005 |
| Gm20517 | 0.007 | 0.006 | 0.005 |
| Emid1 | 0.007 | 0.060 | 0.030 |
| Tnfsf13os | 0.007 | 0.011 | 0.006 |
| Plbd1 | 0.007 | 0.002 | 0.001 |
| Wwc1 | 0.007 | 0.032 | 0.033 |
| Alpl | 0.007 | 0.018 | 0.033 |
| Csrp3 | 0.007 | 0.021 | 0.012 |
| 9430060I03R | 0.007 | 0.011 | 0.012 |
| E030037K01R | 0.007 | 0.004 | 0.003 |
| Ccl21a | 0.007 | 0.005 | 0.006 |
| Rnase10 | 0.007 | 0.008 | 0.014 |
| Susd4 | 0.007 | 0.113 | 0.060 |
| Abcb1a | 0.007 | 0.007 | 0.004 |
| Gm10645 | 0.007 | 0.010 | 0.005 |
| Irf8 | 0.007 | 0.014 | 0.064 |
| Hpn | 0.007 | 0.001 | 0.002 |
| Zfp382 | 0.007 | 0.014 | 0.012 |
| Mypopos | 0.007 | 0.004 | 0.006 |
| Zkscan4 | 0.007 | 0.007 | 0.012 |
| Dmc1 | 0.007 | 0.022 | 0.018 |
| Col17a1 | 0.007 | 0.005 | 0.014 |
| Cilp2 | 0.007 | 0.009 | 0.006 |
| Cd59b | 0.007 | 0.019 | 0.019 |
| Tnfsf13b | 0.007 | 0.007 | 0.017 |
| Pcdhb8 | 0.007 | 0.006 | 0.007 |
| Pcdhb13 | 0.007 | 0.003 | 0.003 |
| Gm26830 | 0.007 | 0.002 | 0.003 |

| | | | |
|---|---|---|---|
| Wdr54 | 0.007 | 0.016 | 0.024 |
| Junos | 0.007 | 0.010 | 0.012 |
| Dio2 | 0.007 | 0.002 | 0.004 |
| Nat8f4 | 0.007 | 0.003 | 0.003 |
| Lsmem1 | 0.007 | 0.047 | 0.042 |
| Shmt1 | 0.007 | 0.012 | 0.011 |
| Syt10 | 0.007 | 0.001 | 0.000 |
| AK157302 | 0.007 | 0.003 | 0.005 |
| Gm42067 | 0.007 | 0.010 | 0.005 |
| Lypd1 | 0.007 | 0.013 | 0.078 |
| 2810433D01I | 0.007 | 0.032 | 0.093 |
| Il17b | 0.007 | 0.093 | 0.016 |
| Itga7 | 0.007 | 0.084 | 0.013 |
| Ptgfr | 0.007 | 0.001 | 0.022 |
| Tgm5 | 0.007 | 0.001 | 0.001 |
| Sdr42e1 | 0.007 | 0.002 | 0.004 |
| Tesk2 | 0.007 | 0.014 | 0.012 |
| Lama1 | 0.007 | 0.015 | 0.001 |
| Apc2 | 0.007 | 0.011 | 0.020 |
| Camkk1 | 0.007 | 0.022 | 0.055 |
| Hecw1 | 0.007 | 0.012 | 0.007 |
| Gm26647 | 0.007 | 0.000 | 0.000 |
| Egflam | 0.007 | 0.025 | 0.157 |
| Gm13524 | 0.007 | 0.008 | 0.008 |
| Plxna3 | 0.007 | 0.045 | 0.063 |
| Aoc2 | 0.007 | 0.009 | 0.006 |
| Stmn3 | 0.007 | 0.003 | 0.006 |
| Cgn | 0.007 | 0.001 | 0.001 |
| Fev | 0.007 | 0.000 | 0.000 |
| Gm26540 | 0.007 | 0.001 | 0.001 |
| Cdiptos | 0.007 | 0.002 | 0.008 |
| Tsx | 0.007 | 0.004 | 0.004 |
| Nlrc3 | 0.007 | 0.009 | 0.008 |
| Col7a1 | 0.007 | 0.006 | 0.006 |
| Cap2 | 0.007 | 0.173 | 0.057 |
| Tm7sf2 | 0.007 | 0.017 | 0.019 |
| Gm12981 | 0.007 | 0.012 | 0.008 |
| Gm44752 | 0.007 | 0.012 | 0.022 |
| 1700028J19R | 0.007 | 0.006 | 0.004 |
| Cox7a1 | 0.007 | 0.008 | 0.010 |
| Lrrn2 | 0.007 | 0.065 | 0.158 |
| Slc25a43 | 0.007 | 0.007 | 0.001 |
| B3gat2 | 0.007 | 0.005 | 0.004 |
| Mocos | 0.007 | 0.005 | 0.011 |
| Ndst3 | 0.007 | 0.035 | 0.024 |
| 2810021J22R | 0.007 | 0.026 | 0.028 |
| Nlgn3 | 0.007 | 0.035 | 0.042 |
| Gm43703 | 0.007 | 0.007 | 0.007 |

| | | | |
|---|---|---|---|
| Gm26583 | 0.007 | 0.012 | 0.013 |
| Gm42726 | 0.007 | 0.020 | 0.017 |
| Stambpl1 | 0.007 | 0.016 | 0.033 |
| 4921507P07F | 0.007 | 0.004 | 0.002 |
| Cacng7 | 0.007 | 0.086 | 0.099 |
| Gm45605 | 0.007 | 0.004 | 0.005 |
| Gtse1 | 0.007 | 0.004 | 0.005 |
| Raver1 | 0.007 | 0.007 | 0.007 |
| Gm9958 | 0.007 | 0.003 | 0.004 |
| Hcn2 | 0.007 | 0.003 | 0.006 |
| Abcg3 | 0.007 | 0.000 | 0.001 |
| Pick1.1 | 0.007 | 0.019 | 0.015 |
| Scg5 | 0.007 | 0.008 | 0.002 |
| Map2 | 0.007 | 0.044 | 0.031 |
| Zfp811 | 0.007 | 0.012 | 0.031 |
| Gm45890 | 0.007 | 0.003 | 0.004 |
| Ndnf | 0.007 | 0.031 | 0.081 |
| 9030624G23I | 0.007 | 0.007 | 0.011 |
| Ada | 0.007 | 0.010 | 0.017 |
| Msantd1 | 0.007 | 0.004 | 0.004 |
| Rab7b | 0.007 | 0.008 | 0.121 |
| Shc4 | 0.007 | 0.019 | 0.018 |
| Pak1 | 0.007 | 0.008 | 0.029 |
| Slc9a9 | 0.007 | 0.011 | 0.024 |
| Gm26534 | 0.007 | 0.006 | 0.007 |
| Fcrls | 0.007 | 0.009 | 0.010 |
| Gsap | 0.007 | 0.014 | 0.012 |
| Fhl4 | 0.007 | 0.003 | 0.007 |
| 9430065F17F | 0.007 | 0.005 | 0.008 |
| Nxpe3 | 0.007 | 0.016 | 0.012 |
| Serpina3i | 0.007 | 0.000 | 0.002 |
| Psma8 | 0.007 | 0.002 | 0.004 |
| Ttc39c | 0.007 | 0.039 | 0.045 |
| Gm14322 | 0.007 | 0.011 | 0.015 |
| Pappa | 0.007 | 0.118 | 0.217 |
| Wdr31 | 0.007 | 0.011 | 0.010 |
| Sostdc1 | 0.007 | 0.008 | 0.010 |
| 5033426O07I | 0.007 | 0.001 | 0.000 |
| Lrrn4 | 0.007 | 0.003 | 0.005 |
| Cd53 | 0.007 | 0.004 | 0.003 |
| 1700003F12F | 0.007 | 0.002 | 0.013 |
| Cyp27a1 | 0.007 | 0.005 | 0.041 |
| H60b | 0.007 | 0.000 | 0.000 |
| Gm27029 | 0.007 | 0.011 | 0.011 |
| Gm28809 | 0.007 | 0.007 | 0.004 |
| Abcc3 | 0.007 | 0.004 | 0.003 |
| Piwil2 | 0.007 | 0.006 | 0.006 |
| BC030867 | 0.007 | 0.002 | 0.003 |

| | | | |
|---|---|---|---|
| Gm26901 | 0.007 | 0.004 | 0.012 |
| Anks6 | 0.007 | 0.025 | 0.042 |
| Gm34006 | 0.007 | 0.005 | 0.012 |
| Wasf1 | 0.007 | 0.036 | 0.034 |
| Gm26664 | 0.007 | 0.007 | 0.005 |
| Slc45a3 | 0.007 | 0.057 | 0.051 |
| Reps2 | 0.007 | 0.005 | 0.015 |
| 4831440D22I | 0.006 | 0.005 | 0.004 |
| Cnksr1 | 0.006 | 0.012 | 0.002 |
| Gm15559 | 0.006 | 0.006 | 0.005 |
| Vdr | 0.006 | 0.003 | 0.004 |
| Papolb | 0.006 | 0.001 | 0.003 |
| Zfp759 | 0.006 | 0.005 | 0.008 |
| Apobec1 | 0.006 | 0.007 | 0.005 |
| Gm5860 | 0.006 | 0.007 | 0.007 |
| S100a8 | 0.006 | 0.007 | 0.003 |
| Rasl12 | 0.006 | 0.158 | 0.048 |
| Gstp2 | 0.006 | 0.008 | 0.008 |
| Pagr1a | 0.006 | 0.007 | 0.004 |
| 1700018L02F | 0.006 | 0.012 | 0.015 |
| Gfra1 | 0.006 | 0.040 | 0.144 |
| Gzmm | 0.006 | 0.006 | 0.007 |
| Tmem121 | 0.006 | 0.045 | 0.063 |
| D230017M19 | 0.006 | 0.029 | 0.012 |
| H3f3aos | 0.006 | 0.003 | 0.006 |
| Kcnk1 | 0.006 | 0.146 | 0.069 |
| Eid3 | 0.006 | 0.018 | 0.020 |
| A230057D06I | 0.006 | 0.015 | 0.039 |
| Jph1 | 0.006 | 0.061 | 0.025 |
| Zfp493 | 0.006 | 0.004 | 0.008 |
| Gm12999 | 0.006 | 0.001 | 0.000 |
| A730015C16F | 0.006 | 0.003 | 0.002 |
| Rec114 | 0.006 | 0.001 | 0.001 |
| Pcdhga4 | 0.006 | 0.005 | 0.005 |
| Morn1 | 0.006 | 0.015 | 0.018 |
| Tmem202 | 0.006 | 0.004 | 0.003 |
| Kntc1 | 0.006 | 0.006 | 0.006 |
| Eno2 | 0.006 | 0.056 | 0.045 |
| Gnat2 | 0.006 | 0.001 | 0.004 |
| Dok3 | 0.006 | 0.009 | 0.008 |
| Dnph1 | 0.006 | 0.005 | 0.007 |
| Ccl3 | 0.006 | 0.029 | 0.014 |
| Amz1 | 0.006 | 0.004 | 0.006 |
| Kiss1r | 0.006 | 0.003 | 0.003 |
| Pgam2 | 0.006 | 0.014 | 0.008 |
| Gm11131 | 0.006 | 0.002 | 0.003 |
| Klrg2 | 0.006 | 0.004 | 0.001 |
| Irx2 | 0.006 | 0.007 | 0.020 |

| | | | |
|---|---|---|---|
| Sapcd2 | 0.006 | 0.004 | 0.005 |
| Gm28151 | 0.006 | 0.003 | 0.005 |
| Lancl3 | 0.006 | 0.027 | 0.006 |
| Tnf | 0.006 | 0.005 | 0.005 |
| Trem2 | 0.006 | 0.005 | 0.004 |
| Gm48604 | 0.006 | 0.011 | 0.013 |
| 1110002O04I | 0.006 | 0.006 | 0.012 |
| Cacna2d2 | 0.006 | 0.002 | 0.006 |
| Gm48619 | 0.006 | 0.002 | 0.004 |
| Mylpf | 0.006 | 0.006 | 0.007 |
| Agmo | 0.006 | 0.000 | 0.003 |
| Gm15587 | 0.006 | 0.000 | 0.000 |
| Gm10244 | 0.006 | 0.002 | 0.003 |
| Tnni2 | 0.006 | 0.011 | 0.010 |
| 3300002I08R | 0.006 | 0.015 | 0.017 |
| Hist1h2bh | 0.006 | 0.004 | 0.002 |
| Fabp7 | 0.006 | 0.006 | 0.005 |
| Ccdc96 | 0.006 | 0.003 | 0.003 |
| Mcmdc2 | 0.006 | 0.009 | 0.005 |
| Fam228b | 0.006 | 0.009 | 0.019 |
| Aunip | 0.006 | 0.003 | 0.003 |
| Adgrg3 | 0.006 | 0.000 | 0.001 |
| Kcna2 | 0.006 | 0.042 | 0.016 |
| Gm46440 | 0.006 | 0.009 | 0.008 |
| Gm28707 | 0.006 | 0.007 | 0.008 |
| Fam155a | 0.006 | 0.000 | 0.001 |
| BC025920 | 0.006 | 0.004 | 0.007 |
| Phlda2 | 0.006 | 0.004 | 0.003 |
| Gm9991 | 0.006 | 0.000 | 0.002 |
| Glyctk | 0.006 | 0.004 | 0.005 |
| Gm46218 | 0.006 | 0.005 | 0.011 |
| Fbxo27 | 0.006 | 0.027 | 0.026 |
| Gm13112 | 0.006 | 0.000 | 0.000 |
| Gm30655 | 0.006 | 0.002 | 0.001 |
| Mmp17 | 0.006 | 0.064 | 0.009 |
| Hes7 | 0.006 | 0.007 | 0.011 |
| Gm14302 | 0.006 | 0.003 | 0.001 |
| BC043934 | 0.006 | 0.015 | 0.007 |
| S100a9 | 0.006 | 0.007 | 0.004 |
| C5ar2 | 0.006 | 0.001 | 0.001 |
| 4833413G10I | 0.006 | 0.009 | 0.005 |
| Nup210l | 0.006 | 0.019 | 0.027 |
| Gabra3 | 0.006 | 0.056 | 0.147 |
| Pcsk6 | 0.006 | 0.016 | 0.184 |
| Rtn4rl1 | 0.006 | 0.007 | 0.008 |
| Gm15728 | 0.006 | 0.012 | 0.015 |
| Cfap126 | 0.006 | 0.004 | 0.005 |
| Kcnrg | 0.006 | 0.003 | 0.002 |

| | | | |
|---|---|---|---|
| Gm43101 | 0.006 | 0.012 | 0.026 |
| Rac2 | 0.006 | 0.009 | 0.007 |
| Gm3839 | 0.006 | 0.005 | 0.005 |
| Osr2 | 0.006 | 0.026 | 0.232 |
| Gm6712 | 0.006 | 0.004 | 0.005 |
| Ryr2 | 0.006 | 0.141 | 0.065 |
| Cybb | 0.006 | 0.004 | 0.004 |
| Mrpl53 | 0.006 | 0.003 | 0.003 |
| Dcx | 0.006 | 0.008 | 0.009 |
| Snca | 0.006 | 0.004 | 0.003 |
| Gm35339 | 0.006 | 0.005 | 0.004 |
| Astn1 | 0.006 | 0.000 | 0.001 |
| Adgrb2 | 0.006 | 0.021 | 0.056 |
| Spink10 | 0.006 | 0.003 | 0.004 |
| Olfr267 | 0.006 | 0.014 | 0.009 |
| Aif1 | 0.006 | 0.013 | 0.013 |
| 9330179D12I | 0.006 | 0.007 | 0.007 |
| Lpar3 | 0.006 | 0.007 | 0.014 |
| Kcnj15 | 0.006 | 0.096 | 0.057 |
| Pcdhga8 | 0.006 | 0.007 | 0.005 |
| Tbx2 | 0.006 | 0.060 | 0.084 |
| Zfp455 | 0.006 | 0.006 | 0.008 |
| Rassf2 | 0.006 | 0.022 | 0.120 |
| A730046J19R | 0.006 | 0.006 | 0.005 |
| Appbp2os | 0.006 | 0.002 | 0.003 |
| Zfp454 | 0.006 | 0.018 | 0.020 |
| Gm36283 | 0.006 | 0.017 | 0.004 |
| Gm26571 | 0.006 | 0.003 | 0.002 |
| Rell2 | 0.006 | 0.002 | 0.001 |
| Gm4724 | 0.006 | 0.008 | 0.009 |
| Iglon5 | 0.006 | 0.039 | 0.047 |
| Fmnl1 | 0.006 | 0.001 | 0.004 |
| Rhoh | 0.006 | 0.001 | 0.001 |
| Nrn1 | 0.006 | 0.094 | 0.033 |
| Zfp966 | 0.006 | 0.007 | 0.010 |
| Ripor3 | 0.006 | 0.019 | 0.045 |
| D930048N14I | 0.006 | 0.004 | 0.002 |
| Tesc | 0.006 | 0.010 | 0.003 |
| Hist1h4j | 0.006 | 0.003 | 0.001 |
| Traip | 0.006 | 0.010 | 0.015 |
| Tnnt3 | 0.006 | 0.058 | 0.041 |
| Gm47767 | 0.006 | 0.005 | 0.003 |
| Akr1b8 | 0.006 | 0.021 | 0.072 |
| Matn4 | 0.006 | 0.048 | 0.037 |
| Kcnab3 | 0.006 | 0.008 | 0.009 |
| Cd5 | 0.006 | 0.000 | 0.000 |
| Fam83e | 0.006 | 0.014 | 0.011 |
| Xdh | 0.006 | 0.007 | 0.076 |

| | | | |
|---|---|---|---|
| Gm3550 | 0.006 | 0.005 | 0.007 |
| Tspan2os | 0.006 | 0.005 | 0.002 |
| Slc22a14 | 0.006 | 0.000 | 0.001 |
| Cspg5 | 0.006 | 0.024 | 0.047 |
| AC152827.1 | 0.006 | 0.003 | 0.004 |
| Ribc1 | 0.006 | 0.003 | 0.003 |
| Gm6225 | 0.006 | 0.009 | 0.003 |
| 3110070M22 | 0.006 | 0.018 | 0.015 |
| Scnn1a | 0.006 | 0.005 | 0.004 |
| Polr3g | 0.006 | 0.010 | 0.011 |
| Gm34934 | 0.006 | 0.032 | 0.033 |
| Padi4 | 0.006 | 0.000 | 0.001 |
| Ppfia4 | 0.006 | 0.022 | 0.021 |
| Hoxb5os | 0.006 | 0.076 | 0.036 |
| Gm26772 | 0.006 | 0.004 | 0.006 |
| Nek2 | 0.006 | 0.003 | 0.005 |
| Gm13470 | 0.006 | 0.008 | 0.023 |
| Mettl5os | 0.006 | 0.004 | 0.003 |
| Bdh1 | 0.006 | 0.064 | 0.007 |
| Olr1 | 0.006 | 0.000 | 0.000 |
| Ccdc110 | 0.006 | 0.007 | 0.002 |
| Clic3 | 0.006 | 0.005 | 0.006 |
| Ticrr | 0.006 | 0.006 | 0.005 |
| Edil3 | 0.006 | 0.001 | 0.002 |
| Ctnna3 | 0.006 | 0.084 | 0.016 |
| Cd83 | 0.006 | 0.011 | 0.038 |
| Gm5239 | 0.006 | 0.006 | 0.009 |
| Nrcam | 0.006 | 0.006 | 0.014 |
| Gm29394 | 0.006 | 0.009 | 0.007 |
| Bcat1 | 0.006 | 0.007 | 0.094 |
| Gm26617 | 0.006 | 0.003 | 0.001 |
| Hsf2bp | 0.006 | 0.004 | 0.002 |
| Brd3os | 0.006 | 0.029 | 0.041 |
| AC123724.1 | 0.006 | 0.008 | 0.006 |
| Gm26581 | 0.006 | 0.000 | 0.000 |
| Gm45767 | 0.006 | 0.019 | 0.018 |
| Camk2a | 0.006 | 0.002 | 0.009 |
| Zfp174 | 0.006 | 0.011 | 0.010 |
| Ifi207 | 0.006 | 0.005 | 0.029 |
| Map2k3os | 0.006 | 0.024 | 0.013 |
| Gm10382 | 0.006 | 0.007 | 0.004 |
| Doc2g | 0.006 | 0.014 | 0.009 |
| 4921504A21I | 0.006 | 0.010 | 0.014 |
| Dct | 0.006 | 0.007 | 0.007 |
| Apol8 | 0.006 | 0.019 | 0.006 |
| Foxq1 | 0.006 | 0.003 | 0.001 |
| Rsph1 | 0.006 | 0.008 | 0.018 |
| Gm26826 | 0.006 | 0.012 | 0.007 |

| | | | |
|---|---|---|---|
| Bmp3 | 0.006 | 0.038 | 0.205 |
| Gm7233 | 0.006 | 0.000 | 0.000 |
| Gm47512 | 0.006 | 0.007 | 0.007 |
| Uprt | 0.006 | 0.004 | 0.004 |
| Rnf112 | 0.006 | 0.020 | 0.021 |
| Chp2 | 0.006 | 0.000 | 0.000 |
| 6030443J06R | 0.006 | 0.007 | 0.008 |
| Mgst2 | 0.006 | 0.003 | 0.003 |
| Mir155hg | 0.006 | 0.010 | 0.010 |
| Tcfl5 | 0.006 | 0.003 | 0.009 |
| Comp | 0.006 | 0.006 | 0.014 |
| Proca1 | 0.006 | 0.022 | 0.047 |
| Lrrc55 | 0.006 | 0.001 | 0.016 |
| Gls2 | 0.006 | 0.003 | 0.005 |
| Ccdc171 | 0.006 | 0.022 | 0.023 |
| Gm6377 | 0.006 | 0.004 | 0.002 |
| Rp1 | 0.006 | 0.000 | 0.000 |
| Tnfrsf18 | 0.006 | 0.005 | 0.003 |
| Lrrc71 | 0.006 | 0.001 | 0.002 |
| Hrg | 0.006 | 0.000 | 0.000 |
| Fbxw18 | 0.006 | 0.000 | 0.000 |
| Epor | 0.006 | 0.010 | 0.008 |
| Spink2 | 0.006 | 0.009 | 0.149 |
| Cpne5 | 0.006 | 0.010 | 0.019 |
| Tbc1d10c | 0.006 | 0.004 | 0.007 |
| Cdk5r1 | 0.006 | 0.024 | 0.022 |
| Fcgr2b | 0.006 | 0.005 | 0.003 |
| Zfp941 | 0.006 | 0.001 | 0.001 |
| Man2c1os | 0.006 | 0.009 | 0.007 |
| Hipk4 | 0.006 | 0.012 | 0.006 |
| Pcsk4 | 0.006 | 0.008 | 0.004 |
| AC130815.2 | 0.006 | 0.006 | 0.023 |
| Lingo1 | 0.006 | 0.042 | 0.054 |
| Gm6566 | 0.006 | 0.007 | 0.007 |
| Tbx3 | 0.006 | 0.045 | 0.026 |
| Kif14 | 0.006 | 0.004 | 0.003 |
| Igsf6 | 0.006 | 0.001 | 0.002 |
| Lrrc26 | 0.006 | 0.000 | 0.000 |
| Gm26902 | 0.005 | 0.000 | 0.002 |
| Gm26720 | 0.005 | 0.012 | 0.008 |
| Nrg2 | 0.005 | 0.071 | 0.021 |
| 4932412D23I | 0.005 | 0.000 | 0.000 |
| Asphd2 | 0.005 | 0.011 | 0.057 |
| Eci3 | 0.005 | 0.006 | 0.016 |
| Zfp385b | 0.005 | 0.073 | 0.019 |
| Il13ra2 | 0.005 | 0.001 | 0.005 |
| Syt3 | 0.005 | 0.005 | 0.006 |
| 1500002C15I | 0.005 | 0.010 | 0.014 |

| | | | |
|---|---|---|---|
| Gm13305 | 0.005 | 0.035 | 0.037 |
| Nme5 | 0.005 | 0.011 | 0.011 |
| Gm30003 | 0.005 | 0.003 | 0.003 |
| Cys1 | 0.005 | 0.079 | 0.109 |
| Chrdl2 | 0.005 | 0.071 | 0.019 |
| Pgbd1 | 0.005 | 0.005 | 0.003 |
| Tnnc2 | 0.005 | 0.007 | 0.003 |
| Osbp2 | 0.005 | 0.003 | 0.002 |
| Ehf | 0.005 | 0.000 | 0.001 |
| 6430573F11F | 0.005 | 0.104 | 0.054 |
| Gca | 0.005 | 0.013 | 0.007 |
| Rgcc | 0.005 | 0.053 | 0.022 |
| Tmem35a | 0.005 | 0.051 | 0.034 |
| Rhof | 0.005 | 0.007 | 0.013 |
| 4930426L09F | 0.005 | 0.004 | 0.003 |
| Sstr1 | 0.005 | 0.000 | 0.001 |
| Rbfox1 | 0.005 | 0.002 | 0.007 |
| Gm49164 | 0.005 | 0.005 | 0.004 |
| Itgb2 | 0.005 | 0.001 | 0.001 |
| Zfp964 | 0.005 | 0.009 | 0.009 |
| 1700061G19I | 0.005 | 0.025 | 0.009 |
| Gabrb1 | 0.005 | 0.022 | 0.011 |
| Zfp114 | 0.005 | 0.020 | 0.018 |
| Tm4sf20 | 0.005 | 0.000 | 0.000 |
| Cav3 | 0.005 | 0.079 | 0.016 |
| Tmppe | 0.005 | 0.003 | 0.004 |
| Vip | 0.005 | 0.000 | 0.000 |
| Lrrtm3 | 0.005 | 0.006 | 0.004 |
| Cxcl10 | 0.005 | 0.052 | 0.070 |
| Epb41l4b | 0.005 | 0.018 | 0.017 |
| Sln | 0.005 | 0.007 | 0.011 |
| Sema6b | 0.005 | 0.005 | 0.004 |
| Ttc21a | 0.005 | 0.013 | 0.011 |
| Gm26558 | 0.005 | 0.007 | 0.008 |
| Slc6a20b | 0.005 | 0.013 | 0.015 |
| Gm9833 | 0.005 | 0.004 | 0.004 |
| Zscan25 | 0.005 | 0.004 | 0.004 |
| Anxa9 | 0.005 | 0.003 | 0.005 |
| Gm17227 | 0.005 | 0.007 | 0.004 |
| Ppil6 | 0.005 | 0.009 | 0.006 |
| Cyp26a1 | 0.005 | 0.000 | 0.002 |
| Rapsn | 0.005 | 0.015 | 0.006 |
| Synpo2l | 0.005 | 0.030 | 0.005 |
| Gm42957 | 0.005 | 0.019 | 0.014 |
| Ly6g5b | 0.005 | 0.004 | 0.001 |
| Crygn | 0.005 | 0.006 | 0.011 |
| Gm10371 | 0.005 | 0.041 | 0.024 |
| Lrrc4b | 0.005 | 0.037 | 0.041 |

| | | | |
|---|---|---|---|
| Gm26521 | 0.005 | 0.008 | 0.007 |
| Eme1 | 0.005 | 0.002 | 0.005 |
| Ovgp1 | 0.005 | 0.005 | 0.005 |
| Kcnj14 | 0.005 | 0.000 | 0.000 |
| Hist1h2bj | 0.005 | 0.002 | 0.001 |
| Colgalt2 | 0.005 | 0.004 | 0.009 |
| Abhd3 | 0.005 | 0.006 | 0.007 |
| 9230110C19F | 0.005 | 0.006 | 0.014 |
| Gm14410 | 0.005 | 0.007 | 0.008 |
| Ppp1r14c | 0.005 | 0.096 | 0.024 |
| Kifc5b | 0.005 | 0.002 | 0.003 |
| Hhipl1 | 0.005 | 0.045 | 0.088 |
| Thsd7b | 0.005 | 0.139 | 0.090 |
| Slc7a10 | 0.005 | 0.003 | 0.108 |
| Parpbp | 0.005 | 0.003 | 0.008 |
| A230087F16F | 0.005 | 0.008 | 0.012 |
| Sh3tc2 | 0.005 | 0.001 | 0.001 |
| Cth | 0.005 | 0.009 | 0.015 |
| Car11 | 0.005 | 0.060 | 0.040 |
| Tmem79 | 0.005 | 0.005 | 0.004 |
| 4930537H20I | 0.005 | 0.002 | 0.001 |
| Pcdhga1 | 0.005 | 0.004 | 0.005 |
| Rpusd2 | 0.005 | 0.004 | 0.004 |
| Mybl1 | 0.005 | 0.018 | 0.016 |
| Olfr482 | 0.005 | 0.003 | 0.002 |
| Rgs7 | 0.005 | 0.122 | 0.020 |
| Gm9008 | 0.005 | 0.002 | 0.002 |
| Eaf2 | 0.005 | 0.009 | 0.011 |
| Trpc6 | 0.005 | 0.000 | 0.000 |
| B230217O12I | 0.005 | 0.004 | 0.003 |
| Gm26637 | 0.005 | 0.001 | 0.001 |
| A230005M16 | 0.005 | 0.002 | 0.002 |
| Dgkb | 0.005 | 0.061 | 0.042 |
| Col13a1 | 0.005 | 0.005 | 0.016 |
| Axin2 | 0.005 | 0.017 | 0.016 |
| Arhgap45 | 0.005 | 0.021 | 0.055 |
| Cldn2 | 0.005 | 0.005 | 0.004 |
| Tnfrsf4 | 0.005 | 0.005 | 0.004 |
| Adam22 | 0.005 | 0.028 | 0.062 |
| P2rx6 | 0.005 | 0.012 | 0.009 |
| AA467197 | 0.005 | 0.001 | 0.003 |
| Park2 | 0.005 | 0.010 | 0.016 |
| Gm20633 | 0.005 | 0.005 | 0.004 |
| Pcdhga3 | 0.005 | 0.013 | 0.017 |
| Oasl1 | 0.005 | 0.001 | 0.003 |
| Gm43568 | 0.005 | 0.000 | 0.001 |
| Satb2 | 0.005 | 0.005 | 0.003 |
| Gm14221 | 0.005 | 0.004 | 0.005 |

| | | | |
|---|---|---|---|
| Fbxl21 | 0.005 | 0.005 | 0.004 |
| Batf2 | 0.005 | 0.000 | 0.001 |
| Gm16499.1 | 0.005 | 0.002 | 0.002 |
| Atp6v0c | 0.005 | 0.011 | 0.011 |
| Fbxw15 | 0.005 | 0.000 | 0.000 |
| Sh3gl3 | 0.005 | 0.002 | 0.009 |
| Dgkeos | 0.005 | 0.003 | 0.003 |
| 9430037G07I | 0.005 | 0.004 | 0.005 |
| Dpysl5 | 0.005 | 0.055 | 0.012 |
| Ptk6 | 0.005 | 0.000 | 0.002 |
| 6330562C20F | 0.005 | 0.015 | 0.024 |
| Mrap | 0.005 | 0.011 | 0.037 |
| Grip2 | 0.005 | 0.069 | 0.008 |
| Lrriq3 | 0.005 | 0.003 | 0.005 |
| Spata33 | 0.005 | 0.016 | 0.023 |
| Mafa | 0.005 | 0.009 | 0.007 |
| Rad54b | 0.005 | 0.005 | 0.006 |
| Gm15910 | 0.005 | 0.026 | 0.009 |
| Ctsw | 0.005 | 0.000 | 0.000 |
| Ddi1 | 0.005 | 0.002 | 0.001 |
| Gm44777 | 0.005 | 0.008 | 0.005 |
| Gm9465 | 0.005 | 0.004 | 0.003 |
| Pon1 | 0.005 | 0.000 | 0.000 |
| Npff | 0.005 | 0.006 | 0.007 |
| Zfp418 | 0.005 | 0.005 | 0.006 |
| Tmlhe | 0.005 | 0.011 | 0.012 |
| Gm4419 | 0.005 | 0.005 | 0.006 |
| Plb1 | 0.005 | 0.097 | 0.015 |
| A4galt | 0.005 | 0.105 | 0.035 |
| B230208H11I | 0.005 | 0.009 | 0.006 |
| Gm13091 | 0.005 | 0.002 | 0.002 |
| Doc2b | 0.005 | 0.002 | 0.005 |
| Prph | 0.005 | 0.010 | 0.009 |
| Gm13830 | 0.005 | 0.008 | 0.012 |
| Gm8251 | 0.005 | 0.014 | 0.008 |
| 6030419C18F | 0.005 | 0.041 | 0.154 |
| Ncf1 | 0.005 | 0.007 | 0.009 |
| Gng2 | 0.005 | 0.060 | 0.052 |
| Actn3 | 0.005 | 0.006 | 0.023 |
| Pate2 | 0.005 | 0.005 | 0.003 |
| Col9a3 | 0.005 | 0.096 | 0.028 |
| Gm26775 | 0.005 | 0.000 | 0.000 |
| Agtr1a | 0.005 | 0.018 | 0.065 |
| 4933402J07R | 0.005 | 0.000 | 0.001 |
| Gm26563 | 0.005 | 0.004 | 0.004 |
| Ablim3 | 0.005 | 0.001 | 0.001 |
| Fam78a | 0.005 | 0.001 | 0.000 |
| Gm26523 | 0.005 | 0.028 | 0.057 |

| | | | |
|---|---|---|---|
| Gm5093 | 0.005 | 0.003 | 0.005 |
| Gm3336 | 0.005 | 0.027 | 0.049 |
| Muc13 | 0.005 | 0.002 | 0.002 |
| Gm46447 | 0.005 | 0.008 | 0.006 |
| AI427809 | 0.005 | 0.004 | 0.065 |
| Cdkn2b | 0.005 | 0.008 | 0.023 |
| Fosl1 | 0.005 | 0.015 | 0.004 |
| Col11a2 | 0.005 | 0.006 | 0.010 |
| P2ry14 | 0.005 | 0.056 | 0.013 |
| Rnf225 | 0.005 | 0.011 | 0.013 |
| Mycl | 0.005 | 0.006 | 0.026 |
| Gm26566 | 0.005 | 0.005 | 0.004 |
| A2m | 0.005 | 0.000 | 0.000 |
| Arf4os | 0.005 | 0.005 | 0.009 |
| Serpinb9b | 0.005 | 0.000 | 0.000 |
| Svep1 | 0.005 | 0.007 | 0.204 |
| Hist1h2bg | 0.005 | 0.002 | 0.001 |
| Cpa2 | 0.005 | 0.007 | 0.012 |
| Cidea | 0.005 | 0.125 | 0.015 |
| Kirrel3os | 0.005 | 0.001 | 0.001 |
| Palm3 | 0.005 | 0.001 | 0.002 |
| Cfb | 0.005 | 0.001 | 0.017 |
| C1qtnf3 | 0.005 | 0.009 | 0.132 |
| Gm15408 | 0.005 | 0.003 | 0.003 |
| Prrt1 | 0.005 | 0.033 | 0.025 |
| Dhtkd1 | 0.005 | 0.011 | 0.010 |
| A530013C23R | 0.005 | 0.031 | 0.013 |
| Gm15541 | 0.005 | 0.005 | 0.004 |
| Serpina3f | 0.005 | 0.000 | 0.005 |
| A3galt2 | 0.005 | 0.006 | 0.007 |
| Trim14 | 0.005 | 0.001 | 0.000 |
| Pcdhb10 | 0.005 | 0.002 | 0.002 |
| Muc1 | 0.005 | 0.010 | 0.012 |
| Gm3854 | 0.005 | 0.008 | 0.008 |
| Cd72 | 0.005 | 0.011 | 0.004 |
| Apol9a | 0.005 | 0.012 | 0.009 |
| Riiad1 | 0.005 | 0.009 | 0.019 |
| Myo5b | 0.005 | 0.002 | 0.002 |
| Stmnd1 | 0.005 | 0.014 | 0.100 |
| BC049762 | 0.005 | 0.002 | 0.001 |
| Lepr | 0.005 | 0.002 | 0.030 |
| Gm11491 | 0.005 | 0.005 | 0.006 |
| Mylk3 | 0.005 | 0.006 | 0.018 |
| Lrrc36 | 0.005 | 0.003 | 0.015 |
| Shc2 | 0.005 | 0.032 | 0.105 |
| 4930590J08R | 0.005 | 0.015 | 0.013 |
| Gm26764 | 0.005 | 0.004 | 0.004 |
| Stk26 | 0.005 | 0.008 | 0.008 |

| | | | |
|---|---|---|---|
| Poln | 0.005 | 0.003 | 0.001 |
| Slamf1 | 0.005 | 0.003 | 0.005 |
| Olfr1034 | 0.005 | 0.026 | 0.016 |
| Ap1s3 | 0.005 | 0.002 | 0.001 |
| Gm4035 | 0.005 | 0.001 | 0.002 |
| Tssk6 | 0.005 | 0.005 | 0.005 |
| Gm26682 | 0.005 | 0.006 | 0.010 |
| Tes3-ps | 0.005 | 0.005 | 0.004 |
| Gm11110 | 0.005 | 0.023 | 0.012 |
| Slc2a2 | 0.005 | 0.001 | 0.001 |
| Slc16a8 | 0.005 | 0.004 | 0.006 |
| Gm14168 | 0.005 | 0.002 | 0.002 |
| Gm9888 | 0.005 | 0.004 | 0.004 |
| D5Ertd605e | 0.005 | 0.000 | 0.002 |
| Cdnf | 0.005 | 0.003 | 0.001 |
| Gm17529 | 0.005 | 0.010 | 0.009 |
| Lrrc25 | 0.005 | 0.004 | 0.004 |
| 9330104G04I | 0.005 | 0.011 | 0.011 |
| Stil | 0.005 | 0.009 | 0.012 |
| Cyp4f17 | 0.005 | 0.007 | 0.005 |
| Gm10392 | 0.005 | 0.002 | 0.002 |
| Gm9801 | 0.005 | 0.006 | 0.019 |
| A930024N18I | 0.005 | 0.001 | 0.000 |
| Dtna | 0.005 | 0.048 | 0.008 |
| Gm26737 | 0.005 | 0.004 | 0.007 |
| Taf9 | 0.005 | 0.003 | 0.002 |
| Phf19 | 0.005 | 0.008 | 0.014 |
| Hpgd | 0.005 | 0.024 | 0.058 |
| Gm43403 | 0.005 | 0.001 | 0.001 |
| Vmn1r90 | 0.005 | 0.004 | 0.010 |
| Lrrc4 | 0.005 | 0.009 | 0.015 |
| Adgrd1 | 0.005 | 0.004 | 0.125 |
| Dusp27 | 0.005 | 0.061 | 0.011 |
| Scml4 | 0.005 | 0.001 | 0.013 |
| 1700029I15R | 0.005 | 0.004 | 0.005 |
| Hoxb6 | 0.005 | 0.036 | 0.019 |
| 9330198I05R | 0.005 | 0.000 | 0.001 |
| Sh2d5 | 0.005 | 0.006 | 0.006 |
| Gm49083 | 0.005 | 0.002 | 0.001 |
| Gm15873 | 0.005 | 0.003 | 0.004 |
| Ccl9 | 0.005 | 0.007 | 0.013 |
| Slco2b1 | 0.005 | 0.002 | 0.006 |
| Ngfr | 0.005 | 0.005 | 0.041 |
| Cacng8 | 0.005 | 0.003 | 0.004 |
| Derl3 | 0.005 | 0.001 | 0.002 |
| 1810012K16F | 0.005 | 0.001 | 0.001 |
| Cr2 | 0.005 | 0.001 | 0.002 |
| Pbld2 | 0.005 | 0.002 | 0.003 |

| | | | |
|---|---|---|---|
| G630016G05I | 0.005 | 0.004 | 0.010 |
| BC024063 | 0.005 | 0.009 | 0.011 |
| Gm12089 | 0.005 | 0.008 | 0.008 |
| Aspg | 0.005 | 0.000 | 0.002 |
| Nup210 | 0.005 | 0.003 | 0.002 |
| Cd200r1 | 0.005 | 0.001 | 0.002 |
| Gm2379 | 0.005 | 0.001 | 0.005 |
| E130208F15R | 0.005 | 0.004 | 0.002 |
| Lmtk3 | 0.005 | 0.013 | 0.009 |
| Abca8b | 0.005 | 0.006 | 0.073 |
| Ppara | 0.005 | 0.014 | 0.013 |
| Gm42893 | 0.005 | 0.003 | 0.002 |
| Tvp23bos | 0.005 | 0.008 | 0.013 |
| Gen1 | 0.005 | 0.013 | 0.014 |
| Bnc2 | 0.005 | 0.017 | 0.041 |
| Slc16a10 | 0.005 | 0.029 | 0.036 |
| E030013I19Ri | 0.004 | 0.044 | 0.009 |
| Dynlt1b | 0.004 | 0.004 | 0.004 |
| 1700012C14F | 0.004 | 0.004 | 0.007 |
| Prokr1 | 0.004 | 0.005 | 0.041 |
| Gm34084 | 0.004 | 0.000 | 0.002 |
| Fam135b | 0.004 | 0.005 | 0.001 |
| Gm11716 | 0.004 | 0.001 | 0.000 |
| Mc1r | 0.004 | 0.019 | 0.023 |
| Klhl32 | 0.004 | 0.016 | 0.011 |
| Gm29994 | 0.004 | 0.007 | 0.006 |
| AU015791 | 0.004 | 0.000 | 0.000 |
| Kif5c | 0.004 | 0.065 | 0.020 |
| Fam78b | 0.004 | 0.015 | 0.006 |
| Gjb5 | 0.004 | 0.005 | 0.033 |
| 1700019D03I | 0.004 | 0.011 | 0.068 |
| Sytl5 | 0.004 | 0.005 | 0.005 |
| Pmepa1os | 0.004 | 0.003 | 0.003 |
| Tnfrsf14 | 0.004 | 0.001 | 0.001 |
| Gm7008 | 0.004 | 0.003 | 0.003 |
| Car7 | 0.004 | 0.002 | 0.001 |
| Gm15834 | 0.004 | 0.003 | 0.004 |
| Gm10827 | 0.004 | 0.003 | 0.004 |
| Cst13 | 0.004 | 0.000 | 0.000 |
| Cfap69 | 0.004 | 0.012 | 0.010 |
| Clec12b | 0.004 | 0.000 | 0.000 |
| Cenpi | 0.004 | 0.005 | 0.011 |
| Drc3 | 0.004 | 0.004 | 0.004 |
| Kcnd3 | 0.004 | 0.018 | 0.013 |
| 4933429H19I | 0.004 | 0.003 | 0.004 |
| B230398E01F | 0.004 | 0.004 | 0.009 |
| Slamf9 | 0.004 | 0.006 | 0.012 |
| Tnn | 0.004 | 0.105 | 0.022 |

| | | | |
|---|---|---|---|
| Hmgcll1 | 0.004 | 0.000 | 0.000 |
| Evi2a | 0.004 | 0.003 | 0.002 |
| Plekhh2 | 0.004 | 0.033 | 0.151 |
| Hist2h3c2 | 0.004 | 0.005 | 0.003 |
| Gm34961 | 0.004 | 0.008 | 0.010 |
| Gm9530 | 0.004 | 0.002 | 0.003 |
| Slfn8 | 0.004 | 0.004 | 0.009 |
| Gm20324 | 0.004 | 0.007 | 0.004 |
| CT025678.1 | 0.004 | 0.003 | 0.002 |
| Nkd2 | 0.004 | 0.011 | 0.015 |
| Gm10451 | 0.004 | 0.002 | 0.007 |
| A430078I02R | 0.004 | 0.004 | 0.004 |
| Arhgef4 | 0.004 | 0.012 | 0.008 |
| Mfsd13b | 0.004 | 0.000 | 0.001 |
| Pnoc | 0.004 | 0.245 | 0.060 |
| Kcp | 0.004 | 0.007 | 0.073 |
| A430110L20F | 0.004 | 0.002 | 0.004 |
| Adam8 | 0.004 | 0.007 | 0.012 |
| Gm26606 | 0.004 | 0.006 | 0.010 |
| Mapk15 | 0.004 | 0.001 | 0.001 |
| Gm45799 | 0.004 | 0.003 | 0.002 |
| Slc12a5 | 0.004 | 0.000 | 0.000 |
| Ms4a6b | 0.004 | 0.005 | 0.006 |
| Gm28535 | 0.004 | 0.003 | 0.005 |
| Fam222a | 0.004 | 0.007 | 0.007 |
| Adh6b | 0.004 | 0.000 | 0.001 |
| Itpka | 0.004 | 0.006 | 0.002 |
| Xkr4 | 0.004 | 0.010 | 0.049 |
| Gsta3 | 0.004 | 0.003 | 0.006 |
| Pld5 | 0.004 | 0.009 | 0.002 |
| Gm6634 | 0.004 | 0.002 | 0.001 |
| Ptch2 | 0.004 | 0.005 | 0.079 |
| 9230104L09F | 0.004 | 0.000 | 0.001 |
| Il7 | 0.004 | 0.001 | 0.003 |
| Gm17634 | 0.004 | 0.007 | 0.004 |
| P2rx5 | 0.004 | 0.020 | 0.013 |
| 1700125H20I | 0.004 | 0.000 | 0.001 |
| Pstpip1 | 0.004 | 0.004 | 0.009 |
| Amer2 | 0.004 | 0.072 | 0.010 |
| Rapgef3os2 | 0.004 | 0.001 | 0.004 |
| Fhod3 | 0.004 | 0.036 | 0.077 |
| Slc26a7 | 0.004 | 0.015 | 0.047 |
| Ikzf1 | 0.004 | 0.001 | 0.002 |
| Hsf5 | 0.004 | 0.005 | 0.005 |
| Eya2 | 0.004 | 0.019 | 0.063 |
| Gm26760 | 0.004 | 0.003 | 0.006 |
| Ngef | 0.004 | 0.007 | 0.013 |
| Ccl8 | 0.004 | 0.002 | 0.003 |

| | | | |
|---|---|---|---|
| Maob | 0.004 | 0.048 | 0.028 |
| Gm10501 | 0.004 | 0.005 | 0.005 |
| Gm46404 | 0.004 | 0.009 | 0.011 |
| Gm13075 | 0.004 | 0.002 | 0.003 |
| Crmp1 | 0.004 | 0.000 | 0.002 |
| Pcdhac2 | 0.004 | 0.002 | 0.001 |
| Adap1 | 0.004 | 0.074 | 0.018 |
| Rspo1 | 0.004 | 0.023 | 0.034 |
| Cd80 | 0.004 | 0.003 | 0.002 |
| B230303O12I | 0.004 | 0.011 | 0.010 |
| Npw | 0.004 | 0.001 | 0.001 |
| Cct6b | 0.004 | 0.003 | 0.003 |
| Cmah | 0.004 | 0.003 | 0.021 |
| Gm11827 | 0.004 | 0.025 | 0.016 |
| Pirb | 0.004 | 0.006 | 0.005 |
| Krt17 | 0.004 | 0.001 | 0.001 |
| Ngb | 0.004 | 0.005 | 0.003 |
| B230317F23F | 0.004 | 0.002 | 0.002 |
| A430090L17F | 0.004 | 0.001 | 0.001 |
| Atp6ap1l | 0.004 | 0.014 | 0.009 |
| Fcna | 0.004 | 0.003 | 0.005 |
| Esrp2 | 0.004 | 0.001 | 0.001 |
| Gm47123 | 0.004 | 0.011 | 0.012 |
| Gm19705 | 0.004 | 0.014 | 0.013 |
| Ccdc169 | 0.004 | 0.002 | 0.001 |
| Tigd3 | 0.004 | 0.002 | 0.004 |
| Adhfe1 | 0.004 | 0.024 | 0.019 |
| 2410080I02R | 0.004 | 0.018 | 0.030 |
| AC127341.3 | 0.004 | 0.018 | 0.014 |
| Lao1 | 0.004 | 0.000 | 0.000 |
| Gm26536 | 0.004 | 0.003 | 0.004 |
| Alpk1 | 0.004 | 0.012 | 0.015 |
| Gdf2 | 0.004 | 0.000 | 0.002 |
| Myl1 | 0.004 | 0.008 | 0.006 |
| Txlnb | 0.004 | 0.008 | 0.005 |
| Slc38a3 | 0.004 | 0.005 | 0.002 |
| Gm42997 | 0.004 | 0.004 | 0.003 |
| Exph5 | 0.004 | 0.002 | 0.004 |
| Gm11264 | 0.004 | 0.002 | 0.004 |
| Slc2a4 | 0.004 | 0.121 | 0.024 |
| Lrrc10b | 0.004 | 0.002 | 0.011 |
| Gm43951 | 0.004 | 0.004 | 0.003 |
| Gm12689 | 0.004 | 0.010 | 0.011 |
| Il18 | 0.004 | 0.006 | 0.009 |
| Iqub | 0.004 | 0.007 | 0.002 |
| Gm17690 | 0.004 | 0.004 | 0.007 |
| Gm31763 | 0.004 | 0.004 | 0.004 |
| Osgin1 | 0.004 | 0.002 | 0.003 |

| | | | |
|---|---|---|---|
| Phf11c | 0.004 | 0.002 | 0.005 |
| 1700105P06R | 0.004 | 0.005 | 0.003 |
| Sema4a | 0.004 | 0.007 | 0.011 |
| Gm12925 | 0.004 | 0.004 | 0.003 |
| Tbx15 | 0.004 | 0.051 | 0.017 |
| Tvp23a | 0.004 | 0.022 | 0.041 |
| Kif21b | 0.004 | 0.029 | 0.032 |
| Iqck | 0.004 | 0.004 | 0.008 |
| 1700010I14R | 0.004 | 0.002 | 0.001 |
| Lmod3 | 0.004 | 0.007 | 0.001 |
| Gm49249 | 0.004 | 0.001 | 0.001 |
| Faxc | 0.004 | 0.025 | 0.021 |
| Aqp5 | 0.004 | 0.004 | 0.006 |
| A330041J22R | 0.004 | 0.004 | 0.003 |
| Duxbl3 | 0.004 | 0.006 | 0.005 |
| Srsf12 | 0.004 | 0.012 | 0.009 |
| Elavl2 | 0.004 | 0.002 | 0.004 |
| A830082K12R | 0.004 | 0.012 | 0.023 |
| Lamc2 | 0.004 | 0.013 | 0.005 |
| Ngf | 0.004 | 0.003 | 0.078 |
| Gm26648 | 0.004 | 0.004 | 0.013 |
| Catsperg1 | 0.004 | 0.003 | 0.005 |
| Npm2 | 0.004 | 0.004 | 0.003 |
| Trpm5 | 0.004 | 0.005 | 0.056 |
| Hist1h2ab | 0.004 | 0.003 | 0.002 |
| Ttc41 | 0.004 | 0.002 | 0.004 |
| Gm12108 | 0.004 | 0.002 | 0.005 |
| Scd4 | 0.004 | 0.004 | 0.003 |
| Tbx6 | 0.004 | 0.004 | 0.007 |
| Slc23a3 | 0.004 | 0.007 | 0.005 |
| Stac | 0.004 | 0.115 | 0.031 |
| Neil2 | 0.004 | 0.001 | 0.002 |
| Calr3 | 0.004 | 0.003 | 0.005 |
| Dgat2 | 0.004 | 0.022 | 0.024 |
| Xirp1 | 0.004 | 0.087 | 0.007 |
| Shisa3 | 0.004 | 0.015 | 0.133 |
| Catsper3 | 0.004 | 0.001 | 0.001 |
| AC129178.1 | 0.004 | 0.000 | 0.001 |
| Gm14226 | 0.004 | 0.125 | 0.011 |
| Padi3 | 0.004 | 0.000 | 0.001 |
| Nrg4 | 0.004 | 0.001 | 0.001 |
| Zfp977 | 0.004 | 0.004 | 0.003 |
| Gm13054 | 0.004 | 0.002 | 0.002 |
| Stra6 | 0.004 | 0.000 | 0.001 |
| Gm26640 | 0.004 | 0.000 | 0.001 |
| Syngr3 | 0.004 | 0.004 | 0.005 |
| Gm26810 | 0.004 | 0.002 | 0.002 |
| Proscos | 0.004 | 0.003 | 0.003 |

| | | | |
|---|---|---|---|
| Lhx6 | 0.004 | 0.000 | 0.001 |
| Gm15445 | 0.004 | 0.005 | 0.004 |
| Gm525 | 0.004 | 0.000 | 0.000 |
| Arhgef33 | 0.004 | 0.003 | 0.002 |
| Chodl | 0.004 | 0.083 | 0.039 |
| Zfp872 | 0.004 | 0.002 | 0.001 |
| Gm13375 | 0.004 | 0.004 | 0.008 |
| Fgd2 | 0.004 | 0.002 | 0.002 |
| Gm48882 | 0.004 | 0.004 | 0.002 |
| Slc4a11 | 0.004 | 0.005 | 0.005 |
| Enox1 | 0.004 | 0.008 | 0.026 |
| Ccnb1ip1 | 0.004 | 0.001 | 0.001 |
| Gm21814 | 0.004 | 0.013 | 0.004 |
| Gdf5 | 0.004 | 0.010 | 0.006 |
| A430073D23I | 0.004 | 0.008 | 0.009 |
| Gm29609 | 0.004 | 0.005 | 0.003 |
| Gulo | 0.004 | 0.001 | 0.000 |
| Xlr3a | 0.004 | 0.001 | 0.001 |
| Mutyh | 0.004 | 0.006 | 0.012 |
| Hoxb8 | 0.004 | 0.001 | 0.001 |
| Wfikkn1 | 0.004 | 0.007 | 0.007 |
| Slc22a23 | 0.004 | 0.004 | 0.003 |
| Brinp1 | 0.004 | 0.031 | 0.109 |
| Catsper2 | 0.004 | 0.009 | 0.011 |
| Gm26879 | 0.004 | 0.013 | 0.006 |
| Mmp19 | 0.004 | 0.020 | 0.087 |
| Gm37756 | 0.004 | 0.002 | 0.002 |
| Cracr2a | 0.004 | 0.003 | 0.008 |
| Gria1 | 0.004 | 0.067 | 0.045 |
| Rnf182 | 0.004 | 0.029 | 0.037 |
| Rasl10b | 0.004 | 0.065 | 0.012 |
| 8030442B05F | 0.004 | 0.000 | 0.000 |
| 4930432J09R | 0.004 | 0.000 | 0.000 |
| Gm26645 | 0.004 | 0.007 | 0.011 |
| Gm42722 | 0.004 | 0.000 | 0.000 |
| Tfcp2l1 | 0.004 | 0.003 | 0.002 |
| Fam221b | 0.004 | 0.005 | 0.007 |
| Gm36266 | 0.004 | 0.005 | 0.008 |
| Hs3st3a1 | 0.004 | 0.008 | 0.023 |
| 3930402G23I | 0.004 | 0.034 | 0.026 |
| Fgl1 | 0.004 | 0.001 | 0.004 |
| Lgi4 | 0.004 | 0.009 | 0.018 |
| Gm20735 | 0.004 | 0.003 | 0.002 |
| Slc10a1 | 0.004 | 0.016 | 0.008 |
| Arhgap22 | 0.004 | 0.005 | 0.005 |
| Gm12500 | 0.004 | 0.004 | 0.004 |
| Mpzl3 | 0.004 | 0.011 | 0.007 |
| Acss3 | 0.004 | 0.008 | 0.015 |

| | | | |
|---|---|---|---|
| Bcas1 | 0.004 | 0.006 | 0.007 |
| Gm16485 | 0.004 | 0.006 | 0.003 |
| Hsf3 | 0.004 | 0.001 | 0.002 |
| Ercc6l | 0.004 | 0.009 | 0.014 |
| Saxo2 | 0.004 | 0.010 | 0.014 |
| Gm6297 | 0.004 | 0.005 | 0.007 |
| Zfp786 | 0.004 | 0.003 | 0.004 |
| Rhox5 | 0.004 | 0.000 | 0.003 |
| Med9os | 0.004 | 0.006 | 0.006 |
| Aipl1 | 0.004 | 0.001 | 0.001 |
| Kcna5 | 0.004 | 0.103 | 0.020 |
| Thbs4 | 0.004 | 0.012 | 0.024 |
| Tex45 | 0.004 | 0.000 | 0.001 |
| 4930440I19R | 0.004 | 0.000 | 0.000 |
| Adarb2 | 0.004 | 0.001 | 0.001 |
| 9330121J05R | 0.004 | 0.008 | 0.007 |
| Naip5 | 0.004 | 0.000 | 0.001 |
| Shisa8 | 0.004 | 0.001 | 0.001 |
| Gm16062 | 0.004 | 0.012 | 0.006 |
| Gm11767 | 0.004 | 0.001 | 0.002 |
| Catip | 0.004 | 0.009 | 0.016 |
| Ccdc103 | 0.004 | 0.005 | 0.004 |
| Rnft2 | 0.004 | 0.004 | 0.008 |
| Gm17089 | 0.004 | 0.000 | 0.000 |
| Gm15063 | 0.004 | 0.006 | 0.004 |
| Gypa | 0.004 | 0.001 | 0.001 |
| Chrdl1 | 0.004 | 0.048 | 0.053 |
| Gm26734 | 0.004 | 0.004 | 0.005 |
| AC161165.3 | 0.004 | 0.002 | 0.002 |
| Cyfip2 | 0.004 | 0.007 | 0.003 |
| Lipc | 0.004 | 0.002 | 0.000 |
| AC116487.2 | 0.004 | 0.005 | 0.005 |
| Mcf2 | 0.004 | 0.001 | 0.001 |
| Galnt12 | 0.004 | 0.009 | 0.017 |
| Gsto2 | 0.004 | 0.003 | 0.007 |
| Ryr1 | 0.004 | 0.001 | 0.001 |
| 4833423E24F | 0.004 | 0.000 | 0.000 |
| Mfsd7a | 0.004 | 0.011 | 0.022 |
| Me3 | 0.004 | 0.007 | 0.007 |
| Gm10330 | 0.004 | 0.003 | 0.003 |
| Gm17552 | 0.004 | 0.003 | 0.003 |
| Kcnab2 | 0.004 | 0.001 | 0.002 |
| Dok5 | 0.004 | 0.019 | 0.043 |
| Gm26649 | 0.004 | 0.005 | 0.005 |
| Aldh1a2 | 0.004 | 0.005 | 0.044 |
| Pak6 | 0.004 | 0.012 | 0.002 |
| Gm36756 | 0.004 | 0.000 | 0.000 |
| Hgd | 0.004 | 0.001 | 0.002 |

| | | | |
|---|---|---|---|
| Dcdc2c | 0.004 | 0.002 | 0.000 |
| Myo1f | 0.004 | 0.001 | 0.002 |
| Tmem125 | 0.004 | 0.000 | 0.000 |
| Bmpr1b | 0.004 | 0.023 | 0.021 |
| Gm45474 | 0.004 | 0.001 | 0.001 |
| Srrm3 | 0.004 | 0.010 | 0.009 |
| Gm13842 | 0.004 | 0.000 | 0.000 |
| Cldn10 | 0.004 | 0.055 | 0.089 |
| Hcar1 | 0.004 | 0.008 | 0.004 |
| Gm45447 | 0.004 | 0.005 | 0.005 |
| Gm12843 | 0.004 | 0.002 | 0.001 |
| B3gnt7 | 0.004 | 0.001 | 0.002 |
| Soat2 | 0.004 | 0.001 | 0.001 |
| Zkscan2 | 0.004 | 0.002 | 0.002 |
| Gm10131 | 0.004 | 0.007 | 0.006 |
| Phf21b | 0.004 | 0.004 | 0.003 |
| Wfdc6a | 0.004 | 0.004 | 0.001 |
| Hist1h3a | 0.004 | 0.003 | 0.004 |
| Cpa1 | 0.004 | 0.000 | 0.007 |
| Cutal | 0.004 | 0.000 | 0.000 |
| D830024N08I | 0.004 | 0.005 | 0.008 |
| Olfr57 | 0.004 | 0.007 | 0.006 |
| Lpxn | 0.004 | 0.002 | 0.002 |
| Klf1 | 0.004 | 0.002 | 0.003 |
| Rxrg | 0.004 | 0.002 | 0.004 |
| Rbm44 | 0.004 | 0.018 | 0.006 |
| Il23a | 0.004 | 0.012 | 0.019 |
| AC161607.1 | 0.004 | 0.002 | 0.002 |
| Prob1 | 0.004 | 0.011 | 0.006 |
| Gprc5c | 0.004 | 0.047 | 0.039 |
| Cmtm5 | 0.004 | 0.003 | 0.004 |
| Spire2 | 0.004 | 0.000 | 0.001 |
| Krt13 | 0.004 | 0.005 | 0.002 |
| Nlrp10 | 0.004 | 0.006 | 0.003 |
| Igfbpl1 | 0.004 | 0.007 | 0.001 |
| Fgf16 | 0.004 | 0.003 | 0.013 |
| Rgs17 | 0.004 | 0.031 | 0.060 |
| Pilrb2 | 0.004 | 0.000 | 0.000 |
| Vnn1 | 0.004 | 0.042 | 0.015 |
| Gm14154 | 0.004 | 0.000 | 0.001 |
| Gm1123 | 0.004 | 0.000 | 0.000 |
| Fcgr1 | 0.004 | 0.003 | 0.003 |
| Ggn | 0.004 | 0.005 | 0.004 |
| Kcna4 | 0.004 | 0.009 | 0.009 |
| Frem1 | 0.004 | 0.052 | 0.015 |
| 1810019N24I | 0.004 | 0.002 | 0.002 |
| Cyp2u1 | 0.004 | 0.019 | 0.009 |
| Add2 | 0.004 | 0.007 | 0.009 |

| | | | |
|---|---|---|---|
| Gm43137 | 0.004 | 0.003 | 0.002 |
| Fam19a1 | 0.004 | 0.003 | 0.007 |
| Gm26909 | 0.004 | 0.009 | 0.004 |
| Gm28857 | 0.004 | 0.000 | 0.000 |
| Gm26533 | 0.004 | 0.001 | 0.003 |
| Crym | 0.004 | 0.015 | 0.035 |
| Tex52 | 0.004 | 0.004 | 0.004 |
| Tnmd | 0.004 | 0.049 | 0.033 |
| Gm38037 | 0.004 | 0.001 | 0.002 |
| 4933427E11[illegible] | 0.004 | 0.001 | 0.000 |
| Prkg2 | 0.004 | 0.024 | 0.035 |
| B230312C02[illegible] | 0.004 | 0.012 | 0.004 |
| 8430432A02[illegible] | 0.004 | 0.003 | 0.002 |
| Adrb1 | 0.004 | 0.012 | 0.005 |
| Hist1h4a | 0.004 | 0.005 | 0.005 |
| Stk32c | 0.004 | 0.008 | 0.009 |
| Lrrc4c | 0.004 | 0.001 | 0.004 |
| Adam4 | 0.004 | 0.004 | 0.004 |
| Tekt1 | 0.004 | 0.001 | 0.002 |
| Mroh2a | 0.004 | 0.001 | 0.001 |
| Gp1bb | 0.004 | 0.016 | 0.008 |
| Slc7a3 | 0.003 | 0.004 | 0.004 |
| Mymk | 0.003 | 0.000 | 0.004 |
| Fetub | 0.003 | 0.003 | 0.001 |
| Atad3aos | 0.003 | 0.009 | 0.009 |
| 4930432K21[illegible] | 0.003 | 0.003 | 0.005 |
| Lat | 0.003 | 0.003 | 0.002 |
| A330074K22[illegible] | 0.003 | 0.001 | 0.007 |
| Wdr86 | 0.003 | 0.009 | 0.041 |
| Gm26569 | 0.003 | 0.002 | 0.005 |
| Gm20659 | 0.003 | 0.000 | 0.001 |
| Gm15991 | 0.003 | 0.006 | 0.003 |
| Hist1h1a | 0.003 | 0.003 | 0.002 |
| 1700028K03[illegible] | 0.003 | 0.002 | 0.003 |
| Gm44066 | 0.003 | 0.004 | 0.005 |
| 2610206C17[illegible] | 0.003 | 0.000 | 0.000 |
| Fbxo16 | 0.003 | 0.004 | 0.001 |
| Slc8a2 | 0.003 | 0.105 | 0.027 |
| Gm16174 | 0.003 | 0.004 | 0.003 |
| Gm43042 | 0.003 | 0.000 | 0.000 |
| Pfn4 | 0.003 | 0.004 | 0.008 |
| Clec4a2 | 0.003 | 0.004 | 0.003 |
| E130018N17[illegible] | 0.003 | 0.001 | 0.001 |
| Gm4787 | 0.003 | 0.002 | 0.003 |
| Ptprc | 0.003 | 0.003 | 0.004 |
| Gm44731 | 0.003 | 0.003 | 0.000 |
| Gm13346 | 0.003 | 0.000 | 0.000 |
| Ccl17 | 0.003 | 0.005 | 0.001 |

| | | | |
|---|---|---|---|
| Akr1c19 | 0.003 | 0.003 | 0.043 |
| Fsd1 | 0.003 | 0.015 | 0.019 |
| Steap2 | 0.003 | 0.059 | 0.058 |
| Xylb | 0.003 | 0.006 | 0.008 |
| Serpinb5 | 0.003 | 0.001 | 0.000 |
| Gm15594 | 0.003 | 0.002 | 0.002 |
| 4933433G15I | 0.003 | 0.003 | 0.008 |
| Olfr655 | 0.003 | 0.000 | 0.001 |
| 1300014J16R | 0.003 | 0.003 | 0.002 |
| Nsl1 | 0.003 | 0.004 | 0.004 |
| Tmem51os1 | 0.003 | 0.006 | 0.009 |
| Prkar1b | 0.003 | 0.037 | 0.044 |
| Cela1 | 0.003 | 0.004 | 0.008 |
| Map7d2 | 0.003 | 0.006 | 0.009 |
| Slc29a2 | 0.003 | 0.006 | 0.008 |
| Il1a | 0.003 | 0.001 | 0.000 |
| Tas1r3 | 0.003 | 0.002 | 0.001 |
| Plppr3 | 0.003 | 0.010 | 0.019 |
| Flrt1 | 0.003 | 0.010 | 0.010 |
| Ptprcap | 0.003 | 0.002 | 0.003 |
| Rragb | 0.003 | 0.013 | 0.004 |
| Tgtp2 | 0.003 | 0.001 | 0.000 |
| Pif1 | 0.003 | 0.003 | 0.003 |
| Spx | 0.003 | 0.004 | 0.003 |
| 2310040G07I | 0.003 | 0.006 | 0.006 |
| Nfam1 | 0.003 | 0.001 | 0.002 |
| Pln | 0.003 | 0.025 | 0.004 |
| A930012L18F | 0.003 | 0.003 | 0.006 |
| Arhgap19 | 0.003 | 0.006 | 0.005 |
| Styk1 | 0.003 | 0.002 | 0.002 |
| Erich2os | 0.003 | 0.000 | 0.000 |
| Pacsin1 | 0.003 | 0.003 | 0.002 |
| Colec10 | 0.003 | 0.032 | 0.026 |
| Ptafr | 0.003 | 0.002 | 0.002 |
| Gm44148 | 0.003 | 0.003 | 0.005 |
| Gm41077 | 0.003 | 0.002 | 0.001 |
| Tspo2 | 0.003 | 0.000 | 0.000 |
| Slc6a20a | 0.003 | 0.003 | 0.005 |
| Clec4a1 | 0.003 | 0.004 | 0.004 |
| Megf10 | 0.003 | 0.058 | 0.032 |
| 9130213A22F | 0.003 | 0.003 | 0.005 |
| Rap1gap | 0.003 | 0.013 | 0.008 |
| Dyrk3 | 0.003 | 0.002 | 0.002 |
| 8430423G03I | 0.003 | 0.001 | 0.001 |
| Cx3cr1 | 0.003 | 0.005 | 0.006 |
| Ccr10 | 0.003 | 0.001 | 0.002 |
| Zfp109 | 0.003 | 0.001 | 0.001 |
| Naip2 | 0.003 | 0.001 | 0.001 |

| | | | |
|---|---|---|---|
| Efnb3 | 0.003 | 0.010 | 0.022 |
| Gm13199 | 0.003 | 0.005 | 0.004 |
| Gm29684 | 0.003 | 0.001 | 0.000 |
| Gm37634 | 0.003 | 0.004 | 0.004 |
| Thap8 | 0.003 | 0.005 | 0.004 |
| Frmpd2 | 0.003 | 0.000 | 0.001 |
| Tnfaip8l2 | 0.003 | 0.003 | 0.005 |
| Mef2b | 0.003 | 0.011 | 0.007 |
| Hsbp1l1 | 0.003 | 0.004 | 0.002 |
| Sync | 0.003 | 0.041 | 0.015 |
| Dnah11 | 0.003 | 0.000 | 0.000 |
| Gip | 0.003 | 0.000 | 0.000 |
| Fam169a | 0.003 | 0.002 | 0.001 |
| Gm44949 | 0.003 | 0.003 | 0.002 |
| Dpysl4 | 0.003 | 0.003 | 0.011 |
| Parvg | 0.003 | 0.000 | 0.001 |
| Gm45184 | 0.003 | 0.004 | 0.005 |
| Gm48780 | 0.003 | 0.002 | 0.001 |
| Gm14443 | 0.003 | 0.006 | 0.006 |
| Fut4 | 0.003 | 0.004 | 0.006 |
| A730081D07I | 0.003 | 0.001 | 0.002 |
| 4930540M05 | 0.003 | 0.006 | 0.009 |
| Hck | 0.003 | 0.002 | 0.018 |
| Cebpa | 0.003 | 0.016 | 0.069 |
| A330084C13I | 0.003 | 0.005 | 0.007 |
| AI464131 | 0.003 | 0.014 | 0.082 |
| Pcdhb11 | 0.003 | 0.004 | 0.005 |
| Creg2 | 0.003 | 0.001 | 0.003 |
| I830134H01R | 0.003 | 0.001 | 0.001 |
| Ccdc13 | 0.003 | 0.002 | 0.002 |
| Pla1a | 0.003 | 0.001 | 0.002 |
| Gm10640 | 0.003 | 0.000 | 0.000 |
| Gpr20 | 0.003 | 0.024 | 0.012 |
| Gm19276 | 0.003 | 0.001 | 0.004 |
| Far2 | 0.003 | 0.002 | 0.001 |
| Zfp652os | 0.003 | 0.005 | 0.008 |
| Myom2 | 0.003 | 0.003 | 0.001 |
| Gm14455 | 0.003 | 0.001 | 0.000 |
| Gm26801 | 0.003 | 0.000 | 0.000 |
| Xlr4a | 0.003 | 0.001 | 0.003 |
| Rhpn1 | 0.003 | 0.002 | 0.002 |
| Ttll13 | 0.003 | 0.003 | 0.004 |
| Fbxw10 | 0.003 | 0.008 | 0.009 |
| C5ar1 | 0.003 | 0.003 | 0.005 |
| Gm10647 | 0.003 | 0.002 | 0.002 |
| Clcnkb | 0.003 | 0.000 | 0.001 |
| Nutf2-ps1 | 0.003 | 0.002 | 0.001 |
| Dhrs13os | 0.003 | 0.004 | 0.002 |

| | | | |
|---|---|---|---|
| Enpep | 0.003 | 0.019 | 0.010 |
| Steap4 | 0.003 | 0.006 | 0.057 |
| Mfap3l | 0.003 | 0.003 | 0.007 |
| Gm26798 | 0.003 | 0.003 | 0.004 |
| Gm36569 | 0.003 | 0.001 | 0.000 |
| Ppp1r1c | 0.003 | 0.002 | 0.001 |
| AC161165.2 | 0.003 | 0.000 | 0.000 |
| Gm15866 | 0.003 | 0.003 | 0.004 |
| Ankle1 | 0.003 | 0.003 | 0.005 |
| Tmem171 | 0.003 | 0.000 | 0.000 |
| 3110006O06I | 0.003 | 0.004 | 0.004 |
| 4933406P04F | 0.003 | 0.006 | 0.008 |
| D930030I03R | 0.003 | 0.003 | 0.006 |
| Gm48727 | 0.003 | 0.001 | 0.001 |
| Grik2 | 0.003 | 0.040 | 0.022 |
| Ppm1j | 0.003 | 0.010 | 0.003 |
| Il1rn | 0.003 | 0.001 | 0.000 |
| 1600010M07 | 0.003 | 0.000 | 0.001 |
| 1700084C06F | 0.003 | 0.002 | 0.001 |
| Sp5 | 0.003 | 0.000 | 0.000 |
| Zfp473 | 0.003 | 0.004 | 0.003 |
| 6430550D23I | 0.003 | 0.002 | 0.003 |
| Acvr1c | 0.003 | 0.008 | 0.003 |
| Tceal6 | 0.003 | 0.035 | 0.004 |
| Gm21833 | 0.003 | 0.001 | 0.001 |
| Lgr5 | 0.003 | 0.015 | 0.022 |
| Nr6a1os | 0.003 | 0.002 | 0.001 |
| Lat2 | 0.003 | 0.008 | 0.005 |
| Prrg4 | 0.003 | 0.001 | 0.001 |
| Neb | 0.003 | 0.004 | 0.004 |
| Dnah8 | 0.003 | 0.012 | 0.013 |
| Ttyh1 | 0.003 | 0.029 | 0.012 |
| Gpc2 | 0.003 | 0.002 | 0.001 |
| Ackr1 | 0.003 | 0.001 | 0.012 |
| Slc16a6 | 0.003 | 0.014 | 0.055 |
| B430219N15I | 0.003 | 0.003 | 0.002 |
| Gm43660 | 0.003 | 0.006 | 0.005 |
| Syt14 | 0.003 | 0.001 | 0.001 |
| 2010300C02F | 0.003 | 0.023 | 0.006 |
| Pcdhb15 | 0.003 | 0.005 | 0.005 |
| Gm48809 | 0.003 | 0.001 | 0.001 |
| Dnah7b | 0.003 | 0.010 | 0.022 |
| Prr18 | 0.003 | 0.005 | 0.004 |
| Ptger3 | 0.003 | 0.021 | 0.070 |
| Cyp2s1 | 0.003 | 0.004 | 0.010 |
| Gm7967 | 0.003 | 0.000 | 0.000 |
| Ccdc42 | 0.003 | 0.047 | 0.036 |
| Hoxb7 | 0.003 | 0.001 | 0.002 |

| | | | |
|---|---|---|---|
| Zap70 | 0.003 | 0.001 | 0.000 |
| Plekhb1 | 0.003 | 0.005 | 0.006 |
| Gm11963 | 0.003 | 0.002 | 0.002 |
| 4933405D12I | 0.003 | 0.028 | 0.019 |
| Gm15491 | 0.003 | 0.007 | 0.005 |
| Fndc11 | 0.003 | 0.000 | 0.000 |
| S1pr3 | 0.003 | 0.035 | 0.119 |
| Gm35315 | 0.003 | 0.006 | 0.004 |
| Gm10184 | 0.003 | 0.002 | 0.002 |
| Mesp2 | 0.003 | 0.000 | 0.002 |
| Rims2 | 0.003 | 0.044 | 0.010 |
| A930007I19R | 0.003 | 0.007 | 0.010 |
| Fut2 | 0.003 | 0.001 | 0.001 |
| Sema4g | 0.003 | 0.013 | 0.012 |
| Ttr | 0.003 | 0.004 | 0.007 |
| Gm29562 | 0.003 | 0.001 | 0.001 |
| Gm266 | 0.003 | 0.007 | 0.108 |
| Gm4425 | 0.003 | 0.002 | 0.004 |
| Naip6 | 0.003 | 0.000 | 0.000 |
| Dnd1 | 0.003 | 0.005 | 0.004 |
| Syngap1 | 0.003 | 0.026 | 0.018 |
| Chst10 | 0.003 | 0.017 | 0.028 |
| Rsph10b | 0.003 | 0.001 | 0.001 |
| Rgs14 | 0.003 | 0.000 | 0.000 |
| Gm13861 | 0.003 | 0.022 | 0.003 |
| Gm2479 | 0.003 | 0.001 | 0.003 |
| Nek5 | 0.003 | 0.001 | 0.002 |
| Inca1 | 0.003 | 0.007 | 0.009 |
| Gm15934 | 0.003 | 0.002 | 0.000 |
| Nalcn | 0.003 | 0.021 | 0.008 |
| Tnfaip8l3 | 0.003 | 0.001 | 0.002 |
| Tac1 | 0.003 | 0.008 | 0.020 |
| Cyp7b1 | 0.003 | 0.029 | 0.047 |
| Areg | 0.003 | 0.007 | 0.002 |
| Scgb3a1 | 0.003 | 0.001 | 0.003 |
| Vma21-ps | 0.003 | 0.001 | 0.002 |
| Phex | 0.003 | 0.015 | 0.034 |
| Cabcoco1 | 0.003 | 0.021 | 0.016 |
| Syt4 | 0.003 | 0.005 | 0.003 |
| Sec31b | 0.003 | 0.001 | 0.001 |
| Sybu | 0.003 | 0.002 | 0.007 |
| Fam184b | 0.003 | 0.049 | 0.040 |
| Fbxw26 | 0.003 | 0.000 | 0.000 |
| Syt9 | 0.003 | 0.002 | 0.002 |
| Gabrb2 | 0.003 | 0.031 | 0.037 |
| Gm31513 | 0.003 | 0.004 | 0.007 |
| Mos | 0.003 | 0.001 | 0.001 |
| Dnah6 | 0.003 | 0.000 | 0.000 |

| | | | |
|---|---|---|---|
| Erc2 | 0.003 | 0.001 | 0.000 |
| Stum | 0.003 | 0.028 | 0.007 |
| Slc16a14 | 0.003 | 0.000 | 0.002 |
| Foxd2 | 0.003 | 0.003 | 0.006 |
| Gm37986 | 0.003 | 0.005 | 0.004 |
| Ttc30a2 | 0.003 | 0.001 | 0.001 |
| Gm11149 | 0.003 | 0.002 | 0.000 |
| Mkx | 0.003 | 0.003 | 0.004 |
| Grip1 | 0.003 | 0.008 | 0.015 |
| Trpm1 | 0.003 | 0.000 | 0.001 |
| Galnt2 | 0.003 | 0.009 | 0.004 |
| Nepn | 0.003 | 0.000 | 0.000 |
| Csrnp3 | 0.003 | 0.010 | 0.025 |
| 1700016F12F | 0.003 | 0.001 | 0.001 |
| Gm33091 | 0.003 | 0.001 | 0.000 |
| Irf5 | 0.003 | 0.013 | 0.026 |
| Begain | 0.003 | 0.014 | 0.047 |
| 2210008F06F | 0.003 | 0.003 | 0.006 |
| 4933423P22F | 0.003 | 0.001 | 0.000 |
| Gm20703 | 0.003 | 0.000 | 0.000 |
| Zfand4 | 0.003 | 0.006 | 0.009 |
| Unc13a | 0.003 | 0.004 | 0.003 |
| Strc | 0.003 | 0.000 | 0.000 |
| Abcg4 | 0.003 | 0.022 | 0.014 |
| Gal3st1 | 0.003 | 0.004 | 0.004 |
| Gm26522 | 0.003 | 0.009 | 0.003 |
| Prickle4 | 0.003 | 0.001 | 0.001 |
| E130114P18F | 0.003 | 0.006 | 0.011 |
| Cnnm1 | 0.003 | 0.007 | 0.005 |
| Rnf207 | 0.003 | 0.006 | 0.002 |
| Rgs1 | 0.003 | 0.004 | 0.003 |
| Gm27003 | 0.003 | 0.009 | 0.007 |
| Pard6b | 0.003 | 0.003 | 0.004 |
| Gm44623 | 0.003 | 0.002 | 0.002 |
| Gm35558.1 | 0.003 | 0.003 | 0.006 |
| 2900055J20R | 0.003 | 0.029 | 0.026 |
| Cdh12 | 0.003 | 0.013 | 0.033 |
| Ebi3 | 0.003 | 0.001 | 0.001 |
| Amhr2 | 0.003 | 0.002 | 0.003 |
| Glb1l2 | 0.003 | 0.021 | 0.008 |
| 0610025J13R | 0.003 | 0.002 | 0.005 |
| Gm17275 | 0.003 | 0.004 | 0.005 |
| Tmem132c | 0.003 | 0.012 | 0.060 |
| Gm32051 | 0.003 | 0.000 | 0.001 |
| Gm10767 | 0.003 | 0.003 | 0.003 |
| Faah | 0.003 | 0.001 | 0.002 |
| Gm17324 | 0.003 | 0.005 | 0.008 |
| Gm26804 | 0.003 | 0.001 | 0.001 |

| | | | |
|---|---|---|---|
| 1810034E14F | 0.003 | 0.011 | 0.014 |
| Fam46b | 0.003 | 0.027 | 0.009 |
| Ggt7 | 0.003 | 0.021 | 0.017 |
| Myl2 | 0.003 | 0.002 | 0.005 |
| Mfsd2b | 0.003 | 0.004 | 0.002 |
| Myh7 | 0.003 | 0.015 | 0.005 |
| Cfap77 | 0.003 | 0.005 | 0.003 |
| Gm16907 | 0.003 | 0.009 | 0.008 |
| Rnf39 | 0.003 | 0.000 | 0.000 |
| Fanca | 0.003 | 0.002 | 0.004 |
| Baiap2l2 | 0.003 | 0.030 | 0.007 |
| Gm45733 | 0.003 | 0.002 | 0.002 |
| Clic6 | 0.003 | 0.000 | 0.006 |
| Il17re | 0.003 | 0.002 | 0.002 |
| Dlg2 | 0.003 | 0.037 | 0.108 |
| Aqp3 | 0.003 | 0.001 | 0.000 |
| Slc27a6 | 0.003 | 0.008 | 0.020 |
| Card14 | 0.003 | 0.001 | 0.004 |
| Abcc8 | 0.003 | 0.013 | 0.001 |
| Gm12207 | 0.003 | 0.004 | 0.004 |
| Phf2os1 | 0.003 | 0.002 | 0.002 |
| Rasl2-9 | 0.003 | 0.001 | 0.001 |
| Gm12031 | 0.003 | 0.006 | 0.008 |
| Lhfpl4 | 0.003 | 0.005 | 0.004 |
| Gm7854 | 0.003 | 0.000 | 0.001 |
| Ccdc85a | 0.003 | 0.013 | 0.007 |
| Kcnn1 | 0.003 | 0.004 | 0.012 |
| Irx3 | 0.003 | 0.010 | 0.005 |
| Gpnmb | 0.003 | 0.008 | 0.026 |
| Gm16279 | 0.003 | 0.002 | 0.002 |
| Avp | 0.003 | 0.001 | 0.001 |
| A530040E14F | 0.003 | 0.000 | 0.000 |
| D030025P21I | 0.003 | 0.010 | 0.010 |
| Slain1 | 0.003 | 0.002 | 0.001 |
| Depdc1b | 0.003 | 0.024 | 0.002 |
| Gm10457 | 0.003 | 0.006 | 0.014 |
| A930024E05F | 0.003 | 0.002 | 0.002 |
| Recql4 | 0.003 | 0.001 | 0.003 |
| 1700030J22R | 0.003 | 0.002 | 0.003 |
| Rrh | 0.003 | 0.002 | 0.002 |
| Ubxn10 | 0.003 | 0.001 | 0.000 |
| Gbx2 | 0.003 | 0.000 | 0.000 |
| Gm26944 | 0.003 | 0.001 | 0.002 |
| Samd13 | 0.003 | 0.001 | 0.002 |
| Gm26610 | 0.003 | 0.002 | 0.002 |
| Gm2115 | 0.003 | 0.015 | 0.007 |
| Best1 | 0.003 | 0.003 | 0.020 |
| Platr1 | 0.003 | 0.003 | 0.003 |

| | | | |
|---|---|---|---|
| Ces2e | 0.003 | 0.000 | 0.000 |
| Sftpb | 0.003 | 0.000 | 0.000 |
| Ifi27l2b | 0.003 | 0.000 | 0.000 |
| Bhlhe41 | 0.003 | 0.030 | 0.047 |
| Gm13134 | 0.003 | 0.006 | 0.001 |
| Rbakdn | 0.003 | 0.000 | 0.000 |
| Nppb | 0.003 | 0.001 | 0.000 |
| Hephl1 | 0.003 | 0.000 | 0.000 |
| Gm45620 | 0.003 | 0.005 | 0.004 |
| Hapln3 | 0.003 | 0.004 | 0.003 |
| Gm10584 | 0.003 | 0.000 | 0.000 |
| Lrp8 | 0.003 | 0.008 | 0.003 |
| Kcns3 | 0.003 | 0.060 | 0.051 |
| Glis1 | 0.003 | 0.031 | 0.011 |
| Gm45187 | 0.003 | 0.001 | 0.002 |
| Grid2 | 0.003 | 0.008 | 0.007 |
| Il18r1 | 0.003 | 0.002 | 0.002 |
| C130060C02F | 0.003 | 0.001 | 0.001 |
| Ubxn11 | 0.003 | 0.003 | 0.007 |
| Pip5k1b | 0.003 | 0.052 | 0.007 |
| Slc26a10 | 0.003 | 0.005 | 0.007 |
| Icos | 0.003 | 0.001 | 0.004 |
| Gm26779 | 0.003 | 0.001 | 0.000 |
| Fv1 | 0.003 | 0.001 | 0.001 |
| A730071L15F | 0.003 | 0.001 | 0.001 |
| Gm32219 | 0.003 | 0.001 | 0.004 |
| Hp | 0.003 | 0.007 | 0.068 |
| Gm15839 | 0.003 | 0.003 | 0.005 |
| Gm36899 | 0.003 | 0.003 | 0.003 |
| Gpx2 | 0.003 | 0.000 | 0.001 |
| Ano3 | 0.003 | 0.006 | 0.004 |
| Ska3 | 0.003 | 0.006 | 0.007 |
| Slc23a1 | 0.003 | 0.003 | 0.002 |
| Bpifb5 | 0.003 | 0.000 | 0.000 |
| Gm43050 | 0.003 | 0.004 | 0.006 |
| Gm16548 | 0.003 | 0.000 | 0.000 |
| Upk1b | 0.003 | 0.004 | 0.004 |
| Mroh4 | 0.003 | 0.000 | 0.000 |
| Crybb1 | 0.003 | 0.001 | 0.001 |
| Hist1h4c | 0.003 | 0.003 | 0.002 |
| Gm37584 | 0.003 | 0.003 | 0.002 |
| Dzank1 | 0.003 | 0.010 | 0.009 |
| Gm8104 | 0.003 | 0.000 | 0.000 |
| Xk | 0.003 | 0.025 | 0.005 |
| Olfr691 | 0.003 | 0.000 | 0.000 |
| Gm45833 | 0.003 | 0.011 | 0.004 |
| Ccr5 | 0.003 | 0.002 | 0.001 |
| Zfp781 | 0.003 | 0.006 | 0.008 |

| | | | |
|---|---|---|---|
| Rgs11 | 0.003 | 0.015 | 0.006 |
| Runx2os2 | 0.003 | 0.000 | 0.000 |
| Gm42903 | 0.003 | 0.002 | 0.001 |
| Syt8 | 0.003 | 0.002 | 0.002 |
| Miat | 0.003 | 0.000 | 0.002 |
| Gm48610 | 0.003 | 0.002 | 0.002 |
| Frmd8os | 0.003 | 0.010 | 0.007 |
| Ctrl | 0.003 | 0.001 | 0.001 |
| Fanci | 0.003 | 0.002 | 0.003 |
| Gm44672 | 0.003 | 0.002 | 0.005 |
| Aldh1b1 | 0.003 | 0.009 | 0.005 |
| Scn11a | 0.003 | 0.001 | 0.000 |
| Fam178b | 0.003 | 0.009 | 0.009 |
| Tll1 | 0.003 | 0.001 | 0.002 |
| Necab3 | 0.003 | 0.002 | 0.008 |
| Slc16a11 | 0.003 | 0.000 | 0.000 |
| Shox2 | 0.003 | 0.002 | 0.039 |
| Pbp2 | 0.003 | 0.000 | 0.002 |
| Rbm47 | 0.003 | 0.001 | 0.001 |
| AC154509.1 | 0.003 | 0.034 | 0.023 |
| Xlr4b | 0.003 | 0.001 | 0.002 |
| 4930589L23F | 0.003 | 0.003 | 0.003 |
| Gm44700 | 0.003 | 0.004 | 0.005 |
| Gm14762 | 0.003 | 0.001 | 0.001 |
| Cntn1 | 0.003 | 0.001 | 0.002 |
| Nxph3 | 0.003 | 0.003 | 0.002 |
| Fxyd2 | 0.003 | 0.012 | 0.058 |
| Gm31683 | 0.003 | 0.005 | 0.003 |
| Dapl1 | 0.003 | 0.002 | 0.001 |
| Arhgap4 | 0.003 | 0.003 | 0.003 |
| Gm39214 | 0.003 | 0.005 | 0.003 |
| Il20ra | 0.003 | 0.004 | 0.001 |
| BC030499 | 0.003 | 0.008 | 0.017 |
| Mageb16 | 0.003 | 0.000 | 0.000 |
| Thy1 | 0.003 | 0.007 | 0.036 |
| Tmem25 | 0.003 | 0.006 | 0.006 |
| Lrrd1 | 0.003 | 0.004 | 0.005 |
| Sytl2 | 0.003 | 0.022 | 0.024 |
| Gm34425 | 0.003 | 0.005 | 0.002 |
| A030014E15F | 0.003 | 0.000 | 0.001 |
| 9430085M18 | 0.003 | 0.002 | 0.006 |
| Gng7 | 0.003 | 0.004 | 0.010 |
| Gm26594 | 0.003 | 0.001 | 0.000 |
| AC163633.2 | 0.003 | 0.001 | 0.001 |
| Gm47167 | 0.003 | 0.001 | 0.001 |
| Tyr | 0.003 | 0.000 | 0.000 |
| Fam229a | 0.003 | 0.002 | 0.001 |
| Plet1 | 0.003 | 0.001 | 0.000 |

| | | | |
|---|---|---|---|
| Gm11444 | 0.003 | 0.006 | 0.008 |
| Ms4a6c | 0.003 | 0.010 | 0.008 |
| Tex35 | 0.003 | 0.001 | 0.001 |
| Lrmp | 0.003 | 0.001 | 0.001 |
| Kcnc4 | 0.003 | 0.002 | 0.001 |
| Dennd1c | 0.003 | 0.002 | 0.001 |
| Gm26767 | 0.003 | 0.006 | 0.004 |
| Gna15 | 0.003 | 0.001 | 0.001 |
| Stap1 | 0.003 | 0.004 | 0.003 |
| Nckap1l | 0.003 | 0.003 | 0.002 |
| Susd3 | 0.003 | 0.002 | 0.001 |
| Tlr6 | 0.003 | 0.004 | 0.009 |
| Gm26620 | 0.003 | 0.007 | 0.012 |
| Slc17a7 | 0.003 | 0.006 | 0.007 |
| Gm45847 | 0.003 | 0.011 | 0.005 |
| 4930579P08F | 0.003 | 0.008 | 0.001 |
| Gm44622 | 0.003 | 0.000 | 0.000 |
| Npl | 0.003 | 0.004 | 0.012 |
| AC163720.3 | 0.003 | 0.006 | 0.007 |
| 4933408J17R | 0.003 | 0.001 | 0.001 |
| Ntf5 | 0.003 | 0.003 | 0.014 |
| Adam18 | 0.003 | 0.000 | 0.000 |
| Fam131b | 0.003 | 0.008 | 0.024 |
| AI839979 | 0.003 | 0.007 | 0.005 |
| Htr1b | 0.003 | 0.038 | 0.008 |
| Spp1 | 0.003 | 0.005 | 0.004 |
| Iqcg | 0.003 | 0.002 | 0.004 |
| Trpv2 | 0.003 | 0.039 | 0.024 |
| 2210406O10I | 0.003 | 0.003 | 0.005 |
| Ccr1 | 0.003 | 0.002 | 0.002 |
| Zpbp | 0.003 | 0.005 | 0.005 |
| Gad2 | 0.003 | 0.001 | 0.000 |
| Bpifb1 | 0.003 | 0.000 | 0.001 |
| Cxcl14 | 0.003 | 0.039 | 0.013 |
| Gm47761 | 0.003 | 0.002 | 0.003 |
| Tmsb15l | 0.003 | 0.006 | 0.004 |
| Foxa2 | 0.003 | 0.002 | 0.001 |
| 5031434C07F | 0.003 | 0.001 | 0.000 |
| Amph | 0.003 | 0.006 | 0.007 |
| Zfp879 | 0.003 | 0.006 | 0.004 |
| Cdh18 | 0.003 | 0.038 | 0.018 |
| Izumo1 | 0.003 | 0.000 | 0.000 |
| A530016L24F | 0.003 | 0.000 | 0.003 |
| Tuba3b | 0.003 | 0.000 | 0.000 |
| Gpat3 | 0.003 | 0.040 | 0.015 |
| E130006D01F | 0.003 | 0.016 | 0.021 |
| Gm9796 | 0.003 | 0.001 | 0.004 |
| Prss53 | 0.003 | 0.005 | 0.011 |

| | | | |
|---|---|---|---|
| 9530046B11F | 0.003 | 0.004 | 0.004 |
| D630003M21 | 0.003 | 0.003 | 0.040 |
| Gm49201 | 0.003 | 0.002 | 0.002 |
| Pcdhgb2 | 0.003 | 0.004 | 0.004 |
| Gpha2 | 0.003 | 0.001 | 0.000 |
| Cd79b | 0.003 | 0.001 | 0.001 |
| Wdr38 | 0.003 | 0.001 | 0.003 |
| Ttll11 | 0.003 | 0.010 | 0.006 |
| Olfr552 | 0.003 | 0.004 | 0.003 |
| Gm15420 | 0.003 | 0.002 | 0.003 |
| Trim63 | 0.003 | 0.014 | 0.010 |
| Phox2a | 0.003 | 0.009 | 0.002 |
| Gm48435 | 0.003 | 0.000 | 0.000 |
| Gja6 | 0.003 | 0.003 | 0.002 |
| D930048G16I | 0.003 | 0.004 | 0.005 |
| Gm48079 | 0.003 | 0.002 | 0.001 |
| Trap1a | 0.003 | 0.000 | 0.000 |
| Gm3468 | 0.003 | 0.007 | 0.003 |
| Gm10874 | 0.003 | 0.001 | 0.001 |
| Slc18a1 | 0.003 | 0.000 | 0.000 |
| Hist1h2br | 0.003 | 0.001 | 0.001 |
| 4930444M15 | 0.003 | 0.016 | 0.005 |
| Lrrn4cl | 0.003 | 0.014 | 0.099 |
| B230112J18R | 0.003 | 0.010 | 0.003 |
| Gm37748 | 0.003 | 0.002 | 0.003 |
| Gm32849 | 0.003 | 0.001 | 0.003 |
| Sptb | 0.003 | 0.001 | 0.001 |
| Gm44257 | 0.003 | 0.001 | 0.001 |
| Gm42918 | 0.003 | 0.011 | 0.008 |
| Slc13a2 | 0.003 | 0.000 | 0.000 |
| Hgf | 0.002 | 0.005 | 0.026 |
| Gm45345 | 0.002 | 0.001 | 0.001 |
| Scgb3a2 | 0.002 | 0.003 | 0.004 |
| Gm26869 | 0.002 | 0.004 | 0.006 |
| Gm31410 | 0.002 | 0.000 | 0.002 |
| 4930594M22 | 0.002 | 0.001 | 0.001 |
| Gm29361 | 0.002 | 0.003 | 0.002 |
| Tmem132b | 0.002 | 0.001 | 0.001 |
| Gm44759 | 0.002 | 0.004 | 0.001 |
| 2010308F09F | 0.002 | 0.001 | 0.002 |
| Abca4 | 0.002 | 0.000 | 0.002 |
| Gm16208 | 0.002 | 0.003 | 0.002 |
| Bdkrb2 | 0.002 | 0.006 | 0.016 |
| Adra1b | 0.002 | 0.001 | 0.011 |
| Cxcr4 | 0.002 | 0.010 | 0.002 |
| 4930505A04F | 0.002 | 0.002 | 0.002 |
| Gm11205 | 0.002 | 0.002 | 0.003 |
| Gm15859 | 0.002 | 0.001 | 0.001 |

| | | | |
|---|---|---|---|
| Flicr | 0.002 | 0.002 | 0.004 |
| Gm45669 | 0.002 | 0.006 | 0.004 |
| 4732496C06F | 0.002 | 0.003 | 0.003 |
| Ccdc63 | 0.002 | 0.003 | 0.002 |
| Txk | 0.002 | 0.002 | 0.001 |
| Mettl21c | 0.002 | 0.003 | 0.002 |
| F2 | 0.002 | 0.000 | 0.001 |
| Gm16049 | 0.002 | 0.001 | 0.002 |
| Cldn22 | 0.002 | 0.002 | 0.025 |
| Gsc | 0.002 | 0.008 | 0.087 |
| Ltc4s | 0.002 | 0.004 | 0.003 |
| Gm47232 | 0.002 | 0.001 | 0.001 |
| BC067074 | 0.002 | 0.024 | 0.036 |
| AL589692.1 | 0.002 | 0.001 | 0.001 |
| Ccdc142os | 0.002 | 0.004 | 0.004 |
| Egfl6 | 0.002 | 0.001 | 0.005 |
| Gm48754 | 0.002 | 0.000 | 0.000 |
| Itgb1bp2 | 0.002 | 0.002 | 0.002 |
| Plcxd3 | 0.002 | 0.001 | 0.023 |
| Gm16315 | 0.002 | 0.003 | 0.001 |
| Gm47545 | 0.002 | 0.001 | 0.001 |
| Phactr3 | 0.002 | 0.010 | 0.003 |
| Npb | 0.002 | 0.003 | 0.008 |
| Gm16537 | 0.002 | 0.006 | 0.003 |
| Tgfbr3l | 0.002 | 0.008 | 0.008 |
| Ccdc88b | 0.002 | 0.001 | 0.001 |
| Atp8b5 | 0.002 | 0.003 | 0.002 |
| Ipcef1 | 0.002 | 0.002 | 0.000 |
| Iqcd | 0.002 | 0.005 | 0.008 |
| 4933424M12 | 0.002 | 0.000 | 0.000 |
| Fhit | 0.002 | 0.007 | 0.007 |
| Afmid | 0.002 | 0.004 | 0.005 |
| Xntrpc | 0.002 | 0.004 | 0.005 |
| Gm13166 | 0.002 | 0.004 | 0.005 |
| Art5 | 0.002 | 0.002 | 0.002 |
| Gm15524 | 0.002 | 0.000 | 0.000 |
| Rpp25 | 0.002 | 0.002 | 0.015 |
| Gm47428 | 0.002 | 0.002 | 0.002 |
| St8sia4 | 0.002 | 0.001 | 0.001 |
| Slc19a3 | 0.002 | 0.000 | 0.001 |
| Tstd1 | 0.002 | 0.002 | 0.003 |
| L3mbtl4 | 0.002 | 0.000 | 0.001 |
| F2rl3 | 0.002 | 0.001 | 0.001 |
| Dpp10 | 0.002 | 0.013 | 0.002 |
| Tex16 | 0.002 | 0.001 | 0.000 |
| Mageb3 | 0.002 | 0.002 | 0.001 |
| Selenbp2 | 0.002 | 0.000 | 0.000 |
| 2010310C07F | 0.002 | 0.004 | 0.006 |

| | | | |
|---|---|---|---|
| Cfap206 | 0.002 | 0.001 | 0.000 |
| Drc1 | 0.002 | 0.003 | 0.002 |
| Gjc3 | 0.002 | 0.005 | 0.003 |
| Gm12185 | 0.002 | 0.000 | 0.000 |
| S100a14 | 0.002 | 0.001 | 0.001 |
| Mamdc2 | 0.002 | 0.030 | 0.019 |
| Il31ra | 0.002 | 0.010 | 0.027 |
| Tmc3 | 0.002 | 0.000 | 0.001 |
| Crb3 | 0.002 | 0.001 | 0.000 |
| Ache | 0.002 | 0.004 | 0.002 |
| Adamts18 | 0.002 | 0.000 | 0.009 |
| Adam2 | 0.002 | 0.001 | 0.000 |
| Gdnf | 0.002 | 0.015 | 0.005 |
| AC163018.1 | 0.002 | 0.000 | 0.001 |
| C78859 | 0.002 | 0.004 | 0.004 |
| 1700001C19F | 0.002 | 0.000 | 0.000 |
| Mylk2 | 0.002 | 0.005 | 0.003 |
| Hist1h2ag | 0.002 | 0.001 | 0.001 |
| Gm28727 | 0.002 | 0.002 | 0.002 |
| 4932416K20F | 0.002 | 0.000 | 0.001 |
| Zkscan16 | 0.002 | 0.001 | 0.002 |
| Hmcn2 | 0.002 | 0.003 | 0.024 |
| Slc1a2 | 0.002 | 0.008 | 0.006 |
| Axdnd1 | 0.002 | 0.002 | 0.002 |
| Orc1 | 0.002 | 0.003 | 0.005 |
| Amdhd1 | 0.002 | 0.000 | 0.001 |
| 1700124L16F | 0.002 | 0.007 | 0.007 |
| Asb16 | 0.002 | 0.001 | 0.001 |
| Ccna1 | 0.002 | 0.001 | 0.000 |
| Gpr39 | 0.002 | 0.005 | 0.043 |
| Crb2 | 0.002 | 0.001 | 0.003 |
| Rab11fip4 | 0.002 | 0.001 | 0.000 |
| Lrrc52 | 0.002 | 0.000 | 0.000 |
| Kbtbd8 | 0.002 | 0.001 | 0.001 |
| Gm45151 | 0.002 | 0.002 | 0.003 |
| Gm16712 | 0.002 | 0.001 | 0.001 |
| Gm9949 | 0.002 | 0.000 | 0.001 |
| C3ar1 | 0.002 | 0.008 | 0.005 |
| Fbxl16 | 0.002 | 0.009 | 0.008 |
| Asb5 | 0.002 | 0.018 | 0.003 |
| Gm47662 | 0.002 | 0.001 | 0.001 |
| 3110045C21F | 0.002 | 0.003 | 0.005 |
| Gm17334 | 0.002 | 0.012 | 0.012 |
| Gm10366 | 0.002 | 0.002 | 0.002 |
| Ppfia2 | 0.002 | 0.012 | 0.030 |
| Rab15 | 0.002 | 0.011 | 0.039 |
| Rtl8c | 0.002 | 0.002 | 0.003 |
| Hs3st3b1 | 0.002 | 0.003 | 0.007 |

| | | | |
|---|---|---|---|
| Bloc1s6os | 0.002 | 0.004 | 0.004 |
| Gm16401 | 0.002 | 0.000 | 0.000 |
| Gm8126 | 0.002 | 0.000 | 0.000 |
| Fbxo47 | 0.002 | 0.001 | 0.001 |
| Apol10b | 0.002 | 0.000 | 0.000 |
| 4930599N23I | 0.002 | 0.002 | 0.003 |
| Gm10629 | 0.002 | 0.000 | 0.000 |
| Abra | 0.002 | 0.062 | 0.009 |
| Ptges | 0.002 | 0.037 | 0.053 |
| Nkx6-1 | 0.002 | 0.005 | 0.003 |
| Spag6l | 0.002 | 0.000 | 0.000 |
| Dnhd1 | 0.002 | 0.003 | 0.003 |
| Gm16741 | 0.002 | 0.000 | 0.001 |
| Pou3f2 | 0.002 | 0.001 | 0.002 |
| Ncam2 | 0.002 | 0.001 | 0.003 |
| Olfr506 | 0.002 | 0.000 | 0.001 |
| Pla2g2c | 0.002 | 0.000 | 0.000 |
| Bbof1 | 0.002 | 0.005 | 0.005 |
| Ncf4 | 0.002 | 0.003 | 0.003 |
| Slc45a2 | 0.002 | 0.000 | 0.000 |
| Galnt14 | 0.002 | 0.023 | 0.010 |
| Magee2 | 0.002 | 0.002 | 0.007 |
| Gm38405 | 0.002 | 0.000 | 0.001 |
| Chrna5 | 0.002 | 0.001 | 0.001 |
| Adcy1 | 0.002 | 0.010 | 0.009 |
| A730049H05I | 0.002 | 0.021 | 0.008 |
| Slc4a1 | 0.002 | 0.000 | 0.000 |
| Gm26674 | 0.002 | 0.001 | 0.002 |
| Arhgap30 | 0.002 | 0.002 | 0.001 |
| 4930524O07I | 0.002 | 0.004 | 0.002 |
| Mmrn1 | 0.002 | 0.001 | 0.002 |
| AC163354.1 | 0.002 | 0.000 | 0.001 |
| Gm38336 | 0.002 | 0.007 | 0.004 |
| Vwce | 0.002 | 0.002 | 0.002 |
| Gm26912 | 0.002 | 0.004 | 0.003 |
| Sh3rf3 | 0.002 | 0.019 | 0.043 |
| Bdh2 | 0.002 | 0.013 | 0.036 |
| Erbb3 | 0.002 | 0.004 | 0.005 |
| Gm48089 | 0.002 | 0.001 | 0.001 |
| Gpr132 | 0.002 | 0.000 | 0.001 |
| Gpld1 | 0.002 | 0.003 | 0.004 |
| Olfr1385 | 0.002 | 0.002 | 0.001 |
| A030005L19F | 0.002 | 0.004 | 0.005 |
| Hist1h2bb | 0.002 | 0.002 | 0.001 |
| Wdfy4 | 0.002 | 0.001 | 0.004 |
| Ncf2 | 0.002 | 0.006 | 0.004 |
| Srpk3 | 0.002 | 0.053 | 0.012 |
| Nckap5los | 0.002 | 0.005 | 0.005 |

| | | | |
|---|---|---|---|
| Sash3 | 0.002 | 0.001 | 0.001 |
| Barx1 | 0.002 | 0.015 | 0.041 |
| Pgpep1l | 0.002 | 0.003 | 0.001 |
| Lrrc3 | 0.002 | 0.004 | 0.014 |
| Exd1 | 0.002 | 0.003 | 0.003 |
| Tff2 | 0.002 | 0.000 | 0.000 |
| Ucp1 | 0.002 | 0.006 | 0.004 |
| Car13 | 0.002 | 0.005 | 0.017 |
| Gm14291 | 0.002 | 0.002 | 0.001 |
| Gm16120 | 0.002 | 0.000 | 0.000 |
| Klk9 | 0.002 | 0.000 | 0.000 |
| Gm31805 | 0.002 | 0.004 | 0.004 |
| Nlrc5 | 0.002 | 0.002 | 0.001 |
| Gm3055 | 0.002 | 0.005 | 0.006 |
| Cpa3 | 0.002 | 0.006 | 0.006 |
| C130071C03F | 0.002 | 0.001 | 0.001 |
| 5730522E02F | 0.002 | 0.002 | 0.003 |
| Gm16059 | 0.002 | 0.001 | 0.001 |
| Gm6569 | 0.002 | 0.005 | 0.002 |
| Cfap70 | 0.002 | 0.001 | 0.001 |
| Gm26671 | 0.002 | 0.001 | 0.001 |
| Rtp3 | 0.002 | 0.000 | 0.000 |
| Gm16299 | 0.002 | 0.001 | 0.003 |
| Actn2 | 0.002 | 0.006 | 0.007 |
| A330094K24F | 0.002 | 0.001 | 0.001 |
| 4930509G22I | 0.002 | 0.003 | 0.004 |
| Dkk1 | 0.002 | 0.003 | 0.001 |
| Nol3 | 0.002 | 0.009 | 0.015 |
| Tmem156 | 0.002 | 0.000 | 0.001 |
| Tulp2 | 0.002 | 0.002 | 0.004 |
| Mbnl3 | 0.002 | 0.002 | 0.004 |
| Trim67 | 0.002 | 0.021 | 0.016 |
| Spata20 | 0.002 | 0.001 | 0.003 |
| Ndp | 0.002 | 0.007 | 0.002 |
| Sbk3 | 0.002 | 0.001 | 0.001 |
| Fgf14 | 0.002 | 0.000 | 0.001 |
| A630009H07I | 0.002 | 0.003 | 0.001 |
| 4933406C10F | 0.002 | 0.003 | 0.007 |
| Ppp1r3g | 0.002 | 0.003 | 0.003 |
| Kcnn2 | 0.002 | 0.015 | 0.010 |
| Ttc36 | 0.002 | 0.005 | 0.003 |
| Gm13483 | 0.002 | 0.006 | 0.004 |
| 1700001L05F | 0.002 | 0.010 | 0.008 |
| Igdcc3 | 0.002 | 0.003 | 0.005 |
| D030040B21I | 0.002 | 0.007 | 0.009 |
| Lrrc18 | 0.002 | 0.003 | 0.006 |
| Khdc3 | 0.002 | 0.002 | 0.000 |
| Cfhr2 | 0.002 | 0.000 | 0.000 |

| | | | |
|---|---|---|---|
| A530001N23I | 0.002 | 0.001 | 0.001 |
| Cnr1 | 0.002 | 0.004 | 0.007 |
| Adipoq | 0.002 | 0.002 | 0.004 |
| Ttll8 | 0.002 | 0.001 | 0.002 |
| Dock5 | 0.002 | 0.022 | 0.030 |
| Lrp1b | 0.002 | 0.036 | 0.030 |
| Crygb | 0.002 | 0.002 | 0.000 |
| 1190001M18 | 0.002 | 0.001 | 0.001 |
| Pcna-ps2 | 0.002 | 0.002 | 0.001 |
| 1810010D01I | 0.002 | 0.001 | 0.003 |
| Tcap | 0.002 | 0.048 | 0.010 |
| Cd244 | 0.002 | 0.000 | 0.000 |
| Gm48682 | 0.002 | 0.003 | 0.006 |
| Fmo4 | 0.002 | 0.017 | 0.027 |
| Nsun7 | 0.002 | 0.012 | 0.015 |
| Ak7 | 0.002 | 0.000 | 0.001 |
| Gm38246 | 0.002 | 0.001 | 0.000 |
| Pcdhb12 | 0.002 | 0.003 | 0.005 |
| Bmp7 | 0.002 | 0.001 | 0.050 |
| Gm35696 | 0.002 | 0.001 | 0.000 |
| 2810429I04R | 0.002 | 0.005 | 0.003 |
| Nrip2 | 0.002 | 0.006 | 0.008 |
| Chrnb2 | 0.002 | 0.001 | 0.001 |
| Hoxd9 | 0.002 | 0.001 | 0.000 |
| Gm43140 | 0.002 | 0.001 | 0.000 |
| Nrxn1 | 0.002 | 0.000 | 0.002 |
| Gm26538 | 0.002 | 0.002 | 0.000 |
| Pdzd9 | 0.002 | 0.002 | 0.001 |
| Ttc29 | 0.002 | 0.000 | 0.001 |
| Fam163a | 0.002 | 0.000 | 0.000 |
| Morn5 | 0.002 | 0.001 | 0.001 |
| Gm13387 | 0.002 | 0.000 | 0.000 |
| 1700069L16F | 0.002 | 0.000 | 0.000 |
| Kcnip4 | 0.002 | 0.031 | 0.042 |
| Gm34397 | 0.002 | 0.000 | 0.000 |
| Gm15915 | 0.002 | 0.000 | 0.001 |
| H2-Ob | 0.002 | 0.000 | 0.000 |
| Map3k15 | 0.002 | 0.001 | 0.001 |
| 4933422A05F | 0.002 | 0.000 | 0.001 |
| Gm13703 | 0.002 | 0.000 | 0.000 |
| Gad1os | 0.002 | 0.001 | 0.000 |
| Jsrp1 | 0.002 | 0.007 | 0.012 |
| Ryr3 | 0.002 | 0.009 | 0.006 |
| Cep112os1 | 0.002 | 0.002 | 0.001 |
| Gm4841 | 0.002 | 0.001 | 0.002 |
| Plac9a | 0.002 | 0.003 | 0.012 |
| Gm26691 | 0.002 | 0.001 | 0.003 |
| Gm15635 | 0.002 | 0.001 | 0.002 |

| | | | |
|---|---|---|---|
| Tns4 | 0.002 | 0.001 | 0.001 |
| Gm15938 | 0.002 | 0.002 | 0.001 |
| Gm31816 | 0.002 | 0.000 | 0.000 |
| Gm4779 | 0.002 | 0.000 | 0.000 |
| Gm44686 | 0.002 | 0.000 | 0.000 |
| Amy1 | 0.002 | 0.004 | 0.005 |
| Gm38947 | 0.002 | 0.002 | 0.003 |
| Gm35835 | 0.002 | 0.000 | 0.003 |
| Gm9873 | 0.002 | 0.001 | 0.004 |
| Fam241b | 0.002 | 0.002 | 0.002 |
| Cyb561a3 | 0.002 | 0.002 | 0.001 |
| Hcst | 0.002 | 0.001 | 0.001 |
| Slc38a5 | 0.002 | 0.003 | 0.073 |
| Gm16033 | 0.002 | 0.003 | 0.001 |
| Kl | 0.002 | 0.001 | 0.002 |
| Olfr742 | 0.002 | 0.000 | 0.001 |
| E230025N22R | 0.002 | 0.000 | 0.001 |
| Deup1 | 0.002 | 0.000 | 0.000 |
| Gm44837 | 0.002 | 0.008 | 0.005 |
| Tdrd6 | 0.002 | 0.004 | 0.002 |
| Fam228a | 0.002 | 0.003 | 0.005 |
| Gm11520 | 0.002 | 0.001 | 0.002 |
| Gm26816 | 0.002 | 0.000 | 0.001 |
| Gm1667 | 0.002 | 0.000 | 0.000 |
| Lrfn2 | 0.002 | 0.003 | 0.011 |
| Fam180a | 0.002 | 0.068 | 0.041 |
| Erfe | 0.002 | 0.001 | 0.001 |
| 2810032G03R | 0.002 | 0.002 | 0.006 |
| Msr1 | 0.002 | 0.002 | 0.015 |
| Pabpn1l | 0.002 | 0.001 | 0.001 |
| Kcnq5 | 0.002 | 0.025 | 0.005 |
| Cldn1 | 0.002 | 0.002 | 0.007 |
| Gm26541 | 0.002 | 0.010 | 0.015 |
| C430042M11 | 0.002 | 0.001 | 0.001 |
| Gm10489 | 0.002 | 0.000 | 0.000 |
| Camk1g | 0.002 | 0.000 | 0.006 |
| B230334C09R | 0.002 | 0.017 | 0.010 |
| Fads6 | 0.002 | 0.002 | 0.010 |
| Fam209 | 0.002 | 0.007 | 0.004 |
| Slc2a5 | 0.002 | 0.000 | 0.001 |
| Acot10 | 0.002 | 0.002 | 0.002 |
| Hgfac | 0.002 | 0.001 | 0.001 |
| Gm16075 | 0.002 | 0.000 | 0.000 |
| Prok2 | 0.002 | 0.000 | 0.000 |
| Gm17518 | 0.002 | 0.002 | 0.002 |
| Rnase12 | 0.002 | 0.000 | 0.000 |
| Gm28154 | 0.002 | 0.008 | 0.004 |
| Aldh1l1 | 0.002 | 0.005 | 0.007 |

| | | | |
|---|---|---|---|
| Gm26717 | 0.002 | 0.001 | 0.002 |
| Gm16052 | 0.002 | 0.000 | 0.001 |
| Zfp82 | 0.002 | 0.001 | 0.001 |
| Scn7a | 0.002 | 0.005 | 0.084 |
| Wdr49 | 0.002 | 0.000 | 0.000 |
| Gm7628 | 0.002 | 0.020 | 0.009 |
| Trp63 | 0.002 | 0.001 | 0.001 |
| Stac2 | 0.002 | 0.002 | 0.004 |
| Gm26867 | 0.002 | 0.003 | 0.007 |
| Pigz | 0.002 | 0.003 | 0.007 |
| Clec2l | 0.002 | 0.004 | 0.011 |
| Mfsd4b3 | 0.002 | 0.002 | 0.004 |
| Tctex1d4 | 0.002 | 0.001 | 0.002 |
| Gm33195 | 0.002 | 0.002 | 0.004 |
| Oprl1 | 0.002 | 0.003 | 0.002 |
| Stfa3 | 0.002 | 0.000 | 0.000 |
| Gm34093 | 0.002 | 0.000 | 0.000 |
| Dscam | 0.002 | 0.000 | 0.000 |
| Gm15169 | 0.002 | 0.000 | 0.001 |
| Guca1b | 0.002 | 0.004 | 0.002 |
| Tesmin | 0.002 | 0.004 | 0.005 |
| Gm14137 | 0.002 | 0.000 | 0.000 |
| 4930573H18I | 0.002 | 0.000 | 0.000 |
| AI182371 | 0.002 | 0.001 | 0.003 |
| Gm28730 | 0.002 | 0.002 | 0.006 |
| Gm17111 | 0.002 | 0.002 | 0.002 |
| Tbx5 | 0.002 | 0.016 | 0.032 |
| Bdkrb1 | 0.002 | 0.006 | 0.006 |
| Gpr150 | 0.002 | 0.001 | 0.000 |
| Epsti1 | 0.002 | 0.002 | 0.002 |
| Tlx1 | 0.002 | 0.000 | 0.000 |
| Epn3 | 0.002 | 0.002 | 0.001 |
| 2310074N15I | 0.002 | 0.003 | 0.003 |
| Tmsb15b2 | 0.002 | 0.005 | 0.006 |
| Fam196a | 0.002 | 0.002 | 0.001 |
| Aspa | 0.002 | 0.005 | 0.019 |
| Tfap2a | 0.002 | 0.012 | 0.004 |
| Ppef1 | 0.002 | 0.007 | 0.009 |
| Pex5l | 0.002 | 0.009 | 0.021 |
| CT033750.2 | 0.002 | 0.002 | 0.000 |
| Gm12204 | 0.002 | 0.001 | 0.002 |
| Madcam1 | 0.002 | 0.000 | 0.000 |
| Gm44214 | 0.002 | 0.001 | 0.000 |
| Mmp24 | 0.002 | 0.000 | 0.000 |
| Sv2c | 0.002 | 0.005 | 0.013 |
| Dact2 | 0.002 | 0.001 | 0.031 |
| Scn2b | 0.002 | 0.022 | 0.028 |
| Gm26668 | 0.002 | 0.002 | 0.002 |

| | | | |
|---|---|---|---|
| Ttyh2 | 0.002 | 0.022 | 0.093 |
| Muc2 | 0.002 | 0.002 | 0.001 |
| Gm26739 | 0.002 | 0.002 | 0.003 |
| Gm12264 | 0.002 | 0.001 | 0.001 |
| 4933433G19I | 0.002 | 0.001 | 0.001 |
| Slitrk5 | 0.002 | 0.002 | 0.001 |
| Zic2 | 0.002 | 0.000 | 0.000 |
| Gnrh1 | 0.002 | 0.002 | 0.004 |
| Snap25 | 0.002 | 0.001 | 0.001 |
| Fkbp6 | 0.002 | 0.000 | 0.000 |
| Pacrg | 0.002 | 0.005 | 0.005 |
| Gm26892 | 0.002 | 0.001 | 0.001 |
| 2410004P03F | 0.002 | 0.003 | 0.007 |
| Mdga2 | 0.002 | 0.019 | 0.019 |
| Wnk2 | 0.002 | 0.005 | 0.002 |
| Zfp457 | 0.002 | 0.001 | 0.001 |
| 4930592C13F | 0.002 | 0.001 | 0.001 |
| 1700102P08F | 0.002 | 0.004 | 0.004 |
| Cyp46a1 | 0.002 | 0.008 | 0.017 |
| Gm38287 | 0.002 | 0.001 | 0.000 |
| Prdm1 | 0.002 | 0.001 | 0.001 |
| Elavl4 | 0.002 | 0.007 | 0.010 |
| AA543186 | 0.002 | 0.003 | 0.003 |
| 1700067K01F | 0.002 | 0.002 | 0.002 |
| Samsn1 | 0.002 | 0.002 | 0.002 |
| 9430002A10F | 0.002 | 0.004 | 0.004 |
| Gm45231 | 0.002 | 0.001 | 0.001 |
| Polq | 0.002 | 0.003 | 0.003 |
| Insc | 0.002 | 0.037 | 0.022 |
| Gnat1 | 0.002 | 0.002 | 0.001 |
| Mapk4 | 0.002 | 0.002 | 0.010 |
| T2 | 0.002 | 0.001 | 0.001 |
| Gm43604 | 0.002 | 0.001 | 0.000 |
| Ccdc24 | 0.002 | 0.004 | 0.005 |
| Slc1a7 | 0.002 | 0.001 | 0.002 |
| Wdr72 | 0.002 | 0.012 | 0.007 |
| Dyx1c1 | 0.002 | 0.007 | 0.005 |
| 1700112J16R | 0.002 | 0.004 | 0.005 |
| Igsf5 | 0.002 | 0.002 | 0.000 |
| C2cd4b | 0.002 | 0.002 | 0.003 |
| Col26a1 | 0.002 | 0.005 | 0.039 |
| Gm17660 | 0.002 | 0.000 | 0.000 |
| Apela | 0.002 | 0.003 | 0.006 |
| F2rl2 | 0.002 | 0.000 | 0.000 |
| Wnt2b | 0.002 | 0.001 | 0.021 |
| Adra2b | 0.002 | 0.001 | 0.002 |
| Zfp459 | 0.002 | 0.002 | 0.002 |
| Cd33 | 0.002 | 0.002 | 0.001 |

| | | | |
|---|---|---|---|
| Tmem163 | 0.002 | 0.001 | 0.000 |
| Ackr2 | 0.002 | 0.000 | 0.007 |
| D6Ertd527e | 0.002 | 0.005 | 0.006 |
| Gm40437 | 0.002 | 0.001 | 0.001 |
| 4921539H07I | 0.002 | 0.002 | 0.006 |
| Gm13499 | 0.002 | 0.000 | 0.000 |
| Slc25a18 | 0.002 | 0.000 | 0.000 |
| Sorcs1 | 0.002 | 0.024 | 0.022 |
| Fam26f | 0.002 | 0.003 | 0.003 |
| Elfn1 | 0.002 | 0.001 | 0.008 |
| Gm10277 | 0.002 | 0.002 | 0.001 |
| Gm42851 | 0.002 | 0.001 | 0.002 |
| Gm11899 | 0.002 | 0.000 | 0.000 |
| Oxtr | 0.002 | 0.002 | 0.003 |
| Gm11412 | 0.002 | 0.002 | 0.005 |
| Sla | 0.002 | 0.001 | 0.001 |
| Gm32283 | 0.002 | 0.000 | 0.000 |
| Dnah2 | 0.002 | 0.001 | 0.004 |
| Dnah17 | 0.002 | 0.001 | 0.001 |
| Sycp3 | 0.002 | 0.001 | 0.001 |
| Myo1g | 0.002 | 0.002 | 0.001 |
| Cpxm2 | 0.002 | 0.022 | 0.009 |
| Gm13582 | 0.002 | 0.000 | 0.000 |
| Gm614 | 0.002 | 0.000 | 0.000 |
| Ccl6 | 0.002 | 0.005 | 0.004 |
| Rbm20 | 0.002 | 0.015 | 0.016 |
| Noxred1 | 0.002 | 0.001 | 0.002 |
| Mmp3 | 0.002 | 0.002 | 0.014 |
| Xkrx | 0.002 | 0.001 | 0.002 |
| Garem2 | 0.002 | 0.001 | 0.000 |
| Nphp4 | 0.002 | 0.005 | 0.006 |
| Spef2 | 0.002 | 0.000 | 0.001 |
| Syt5 | 0.002 | 0.037 | 0.012 |
| Nrn1l | 0.002 | 0.005 | 0.001 |
| Gm28183 | 0.002 | 0.000 | 0.000 |
| Gm42809 | 0.002 | 0.001 | 0.002 |
| 1700007K13I | 0.002 | 0.001 | 0.001 |
| Itih3 | 0.002 | 0.008 | 0.002 |
| Rtl1 | 0.002 | 0.003 | 0.023 |
| Oas2 | 0.002 | 0.000 | 0.001 |
| Mterf1b | 0.002 | 0.005 | 0.002 |
| Npy | 0.002 | 0.009 | 0.004 |
| Gm20109 | 0.002 | 0.003 | 0.004 |
| Cacna1e | 0.002 | 0.007 | 0.008 |
| Gpr162 | 0.002 | 0.009 | 0.014 |
| Mc4r | 0.002 | 0.000 | 0.001 |
| Gm33940 | 0.002 | 0.001 | 0.000 |
| Gm12963 | 0.002 | 0.003 | 0.005 |

| | | | |
|---|---|---|---|
| Foxj1 | 0.002 | 0.008 | 0.004 |
| Gm15326 | 0.002 | 0.001 | 0.002 |
| Rubie | 0.002 | 0.002 | 0.000 |
| Gm37401 | 0.002 | 0.001 | 0.001 |
| Dner | 0.002 | 0.005 | 0.098 |
| Gm11175 | 0.002 | 0.003 | 0.004 |
| Nfasc | 0.002 | 0.009 | 0.005 |
| Lrrc75aos2 | 0.002 | 0.006 | 0.007 |
| Icam5 | 0.002 | 0.007 | 0.011 |
| Clcn1 | 0.002 | 0.001 | 0.001 |
| Slc11a1 | 0.002 | 0.021 | 0.043 |
| 3110015C05F | 0.002 | 0.000 | 0.002 |
| Hao1 | 0.002 | 0.004 | 0.001 |
| Cldn19 | 0.002 | 0.006 | 0.002 |
| Spag8 | 0.002 | 0.002 | 0.001 |
| Cnksr2 | 0.002 | 0.001 | 0.002 |
| Lyzl4os | 0.002 | 0.000 | 0.000 |
| Gm20753 | 0.002 | 0.000 | 0.000 |
| Wnt6 | 0.002 | 0.006 | 0.009 |
| Disp2 | 0.002 | 0.000 | 0.000 |
| Gm9903 | 0.002 | 0.001 | 0.002 |
| Gm42372 | 0.002 | 0.001 | 0.000 |
| Phyhip | 0.002 | 0.001 | 0.002 |
| Myoc | 0.002 | 0.002 | 0.011 |
| Olfr259 | 0.002 | 0.000 | 0.000 |
| Rab11fip1 | 0.002 | 0.001 | 0.001 |
| Ccdc116 | 0.002 | 0.001 | 0.002 |
| Pou6f2 | 0.002 | 0.000 | 0.000 |
| Dock2 | 0.002 | 0.001 | 0.001 |
| AC113595.1 | 0.002 | 0.002 | 0.002 |
| Sult4a1 | 0.002 | 0.001 | 0.001 |
| A930003A15F | 0.002 | 0.000 | 0.000 |
| B230344G16I | 0.002 | 0.002 | 0.004 |
| Cd200r3 | 0.002 | 0.000 | 0.000 |
| Gm5532 | 0.002 | 0.002 | 0.002 |
| Thrsp | 0.002 | 0.007 | 0.002 |
| Gm37035 | 0.002 | 0.000 | 0.000 |
| Plac1 | 0.002 | 0.004 | 0.010 |
| Ccdc89 | 0.002 | 0.002 | 0.003 |
| Gm29183 | 0.002 | 0.000 | 0.000 |
| Gm28386 | 0.002 | 0.001 | 0.001 |
| Gm47551 | 0.002 | 0.002 | 0.002 |
| Nap1l5 | 0.002 | 0.015 | 0.022 |
| Fgf10 | 0.002 | 0.004 | 0.142 |
| Spag17 | 0.002 | 0.001 | 0.001 |
| Srcin1 | 0.002 | 0.001 | 0.001 |
| Gm29340 | 0.002 | 0.001 | 0.003 |
| Hgs | 0.002 | 0.002 | 0.002 |

| | | | |
|---|---|---|---|
| Pcdh20 | 0.002 | 0.005 | 0.026 |
| AL732506.1 | 0.002 | 0.002 | 0.002 |
| Clstn3 | 0.002 | 0.001 | 0.001 |
| Dleu7 | 0.002 | 0.003 | 0.019 |
| Prtg | 0.002 | 0.009 | 0.008 |
| Stfa1 | 0.002 | 0.002 | 0.001 |
| Foxd1 | 0.002 | 0.008 | 0.014 |
| Gm26852 | 0.002 | 0.002 | 0.001 |
| Morn3 | 0.002 | 0.000 | 0.000 |
| 1110017D15I | 0.002 | 0.003 | 0.002 |
| Ism1 | 0.002 | 0.005 | 0.019 |
| Syn1 | 0.002 | 0.004 | 0.009 |
| Scg3 | 0.002 | 0.002 | 0.019 |
| Gm13256 | 0.002 | 0.000 | 0.000 |
| 1700008O03I | 0.002 | 0.004 | 0.005 |
| Hs6st3 | 0.002 | 0.000 | 0.003 |
| Eef1a2 | 0.002 | 0.002 | 0.002 |
| Ina | 0.002 | 0.001 | 0.000 |
| Wasf3 | 0.002 | 0.016 | 0.022 |
| Gm9725 | 0.002 | 0.001 | 0.001 |
| Sult2b1 | 0.002 | 0.001 | 0.001 |
| Wt1 | 0.002 | 0.005 | 0.057 |
| Slc7a8 | 0.002 | 0.007 | 0.016 |
| Gm14308 | 0.002 | 0.002 | 0.002 |
| Srrm4 | 0.002 | 0.011 | 0.009 |
| Gm20597 | 0.002 | 0.001 | 0.001 |
| C030010L15F | 0.002 | 0.004 | 0.004 |
| 4930550L24F | 0.002 | 0.000 | 0.000 |
| 1700034I23R | 0.002 | 0.001 | 0.000 |
| Gm13287 | 0.002 | 0.000 | 0.000 |
| Pnliprp1 | 0.002 | 0.000 | 0.000 |
| Tomm6os | 0.002 | 0.001 | 0.001 |
| Chst8 | 0.002 | 0.001 | 0.001 |
| Klhl33 | 0.002 | 0.007 | 0.021 |
| Gm15721 | 0.002 | 0.012 | 0.002 |
| Gm26582 | 0.002 | 0.002 | 0.001 |
| Gm12968 | 0.002 | 0.000 | 0.000 |
| Zfyve28 | 0.002 | 0.012 | 0.012 |
| Gm31812 | 0.002 | 0.002 | 0.003 |
| Phyhd1 | 0.002 | 0.006 | 0.004 |
| Gm35208 | 0.002 | 0.000 | 0.000 |
| Crlf1 | 0.002 | 0.016 | 0.018 |
| Dcst1 | 0.002 | 0.001 | 0.000 |
| Uchl4 | 0.002 | 0.003 | 0.001 |
| Nanos2 | 0.002 | 0.000 | 0.000 |
| Prr22 | 0.002 | 0.001 | 0.002 |
| Lrrn3 | 0.002 | 0.041 | 0.054 |
| Gm9484 | 0.002 | 0.002 | 0.005 |

| | | | |
|---|---|---|---|
| Gm31282 | 0.002 | 0.002 | 0.001 |
| Capns2 | 0.002 | 0.001 | 0.000 |
| Clstn2 | 0.002 | 0.004 | 0.007 |
| Alpk2 | 0.002 | 0.002 | 0.003 |
| Olfr212 | 0.002 | 0.002 | 0.001 |
| Igfbp1 | 0.002 | 0.000 | 0.000 |
| 4930444P10F | 0.002 | 0.000 | 0.001 |
| Lrrc24 | 0.002 | 0.001 | 0.002 |
| Rsg1 | 0.002 | 0.001 | 0.001 |
| Apoa1 | 0.002 | 0.004 | 0.004 |
| 5930422O12I | 0.002 | 0.003 | 0.002 |
| Vmn1r81 | 0.002 | 0.001 | 0.001 |
| Scrg1 | 0.002 | 0.001 | 0.001 |
| Gm15492 | 0.002 | 0.004 | 0.002 |
| Il2rb | 0.002 | 0.003 | 0.001 |
| Gm5608 | 0.002 | 0.020 | 0.011 |
| Gm47450 | 0.002 | 0.002 | 0.003 |
| Olfr775 | 0.002 | 0.001 | 0.000 |
| Ptpro | 0.002 | 0.001 | 0.002 |
| Ccl28 | 0.002 | 0.002 | 0.001 |
| Hcrtr1 | 0.002 | 0.000 | 0.000 |
| Tmcc3os | 0.002 | 0.000 | 0.001 |
| Nr5a2 | 0.002 | 0.000 | 0.000 |
| Gm14703 | 0.002 | 0.001 | 0.002 |
| Gm49106 | 0.002 | 0.000 | 0.000 |
| Gm47162 | 0.002 | 0.000 | 0.000 |
| Six1 | 0.002 | 0.010 | 0.078 |
| Gpr84 | 0.002 | 0.002 | 0.001 |
| Slc22a1 | 0.002 | 0.003 | 0.001 |
| Gsta1 | 0.002 | 0.000 | 0.000 |
| 4930579J19R | 0.002 | 0.000 | 0.000 |
| Mir670hg | 0.002 | 0.031 | 0.016 |
| Grtp1 | 0.002 | 0.017 | 0.012 |
| Gm5148 | 0.002 | 0.002 | 0.005 |
| Lrrc39 | 0.002 | 0.010 | 0.002 |
| Olfr56 | 0.002 | 0.001 | 0.001 |
| 9230104M06 | 0.002 | 0.002 | 0.002 |
| Olfr46 | 0.002 | 0.003 | 0.003 |
| 4930413F20F | 0.002 | 0.006 | 0.008 |
| Prima1 | 0.002 | 0.002 | 0.004 |
| Gm43109 | 0.002 | 0.001 | 0.001 |
| Adam32 | 0.002 | 0.003 | 0.003 |
| Sycp2 | 0.002 | 0.007 | 0.001 |
| Gm16364 | 0.002 | 0.003 | 0.002 |
| Gm53 | 0.002 | 0.000 | 0.000 |
| Gm15651 | 0.002 | 0.001 | 0.000 |
| Ldlrad1 | 0.002 | 0.000 | 0.000 |
| Pla2g2e | 0.002 | 0.000 | 0.003 |

| | | | |
|---|---|---|---|
| Cox6a2 | 0.002 | 0.002 | 0.005 |
| Gm15247 | 0.002 | 0.003 | 0.003 |
| Cavin4 | 0.002 | 0.009 | 0.008 |
| Zfp641 | 0.002 | 0.013 | 0.008 |
| Zan | 0.002 | 0.001 | 0.004 |
| Ky | 0.002 | 0.034 | 0.010 |
| Fcho1 | 0.002 | 0.001 | 0.000 |
| Gm38025 | 0.002 | 0.001 | 0.001 |
| Ell3 | 0.002 | 0.000 | 0.000 |
| Pdzd7 | 0.002 | 0.000 | 0.001 |
| Fut7 | 0.002 | 0.000 | 0.000 |
| Gm47725 | 0.002 | 0.001 | 0.000 |
| Sez6l2 | 0.002 | 0.001 | 0.003 |
| Plcxd2 | 0.002 | 0.008 | 0.009 |
| Gm15688 | 0.002 | 0.003 | 0.002 |
| Gm42031 | 0.002 | 0.004 | 0.002 |
| Tm4sf5 | 0.002 | 0.001 | 0.002 |
| Fam83h | 0.002 | 0.002 | 0.002 |
| Frmpd1 | 0.002 | 0.000 | 0.000 |
| Eqtn | 0.002 | 0.002 | 0.001 |
| Cks1brt | 0.002 | 0.002 | 0.002 |
| A630014C17F | 0.002 | 0.001 | 0.001 |
| Sh2d1b1 | 0.002 | 0.001 | 0.000 |
| 4933430I17R | 0.002 | 0.001 | 0.001 |
| 1810041H14I | 0.002 | 0.001 | 0.001 |
| Hepacam | 0.002 | 0.000 | 0.000 |
| Dph1 | 0.002 | 0.001 | 0.001 |
| Gm4221 | 0.002 | 0.002 | 0.004 |
| Gm26808 | 0.002 | 0.003 | 0.002 |
| Cpne1 | 0.002 | 0.002 | 0.001 |
| Gm20449 | 0.002 | 0.001 | 0.001 |
| Cacna1d | 0.002 | 0.021 | 0.025 |
| Fscn2 | 0.002 | 0.007 | 0.005 |
| Capn3 | 0.002 | 0.005 | 0.002 |
| E330013P04F | 0.002 | 0.001 | 0.001 |
| 9330158H04I | 0.002 | 0.009 | 0.011 |
| Clec10a | 0.002 | 0.001 | 0.002 |
| Ace2 | 0.002 | 0.004 | 0.056 |
| 4930513D17I | 0.002 | 0.000 | 0.000 |
| Lmntd1 | 0.002 | 0.015 | 0.031 |
| Gng13 | 0.002 | 0.001 | 0.002 |
| Cartpt | 0.002 | 0.001 | 0.001 |
| Gm26756 | 0.002 | 0.000 | 0.000 |
| 1700020D05I | 0.002 | 0.002 | 0.002 |
| Tmem270 | 0.002 | 0.000 | 0.000 |
| Rgn | 0.002 | 0.002 | 0.000 |
| Plau | 0.002 | 0.005 | 0.002 |
| Gnmt | 0.002 | 0.004 | 0.002 |

| | | | |
|---|---|---|---|
| Gm27042 | 0.002 | 0.002 | 0.002 |
| Gm15691 | 0.002 | 0.001 | 0.001 |
| Ccbe1 | 0.002 | 0.004 | 0.110 |
| Gpr165 | 0.002 | 0.003 | 0.002 |
| Rnase6 | 0.002 | 0.001 | 0.000 |
| Rtn4r | 0.002 | 0.003 | 0.024 |
| Fgd3 | 0.002 | 0.000 | 0.000 |
| Ica1l | 0.002 | 0.003 | 0.003 |
| 5930438M14 | 0.002 | 0.000 | 0.000 |
| Dnajc22 | 0.002 | 0.000 | 0.000 |
| Gm43126 | 0.002 | 0.001 | 0.000 |
| Gramd2 | 0.002 | 0.004 | 0.002 |
| Gm16677 | 0.002 | 0.002 | 0.002 |
| Lingo4 | 0.002 | 0.000 | 0.000 |
| Spns3 | 0.002 | 0.000 | 0.001 |
| Hypm | 0.002 | 0.001 | 0.001 |
| Spata9 | 0.002 | 0.003 | 0.003 |
| Kcng4 | 0.002 | 0.000 | 0.002 |
| Hspa1l | 0.002 | 0.012 | 0.002 |
| Gm48336 | 0.002 | 0.000 | 0.000 |
| Gm36757 | 0.002 | 0.000 | 0.000 |
| Hist1h2bm | 0.002 | 0.001 | 0.000 |
| Dpf3 | 0.002 | 0.003 | 0.015 |
| Runx2os1 | 0.002 | 0.004 | 0.003 |
| Kcnab3os | 0.002 | 0.002 | 0.002 |
| 9330136K24F | 0.002 | 0.002 | 0.002 |
| Tmem211 | 0.002 | 0.000 | 0.000 |
| Cntfr | 0.002 | 0.026 | 0.131 |
| Gm29707 | 0.002 | 0.001 | 0.000 |
| Bst1 | 0.002 | 0.002 | 0.001 |
| Foxd3 | 0.002 | 0.003 | 0.004 |
| Rnasek | 0.002 | 0.005 | 0.002 |
| Mpp4 | 0.002 | 0.002 | 0.003 |
| Dbil5 | 0.002 | 0.001 | 0.003 |
| Mtfr2 | 0.002 | 0.007 | 0.008 |
| Kbtbd6 | 0.002 | 0.002 | 0.002 |
| 9930004E17F | 0.002 | 0.003 | 0.003 |
| Gm43597 | 0.002 | 0.011 | 0.008 |
| Gm35558 | 0.002 | 0.002 | 0.002 |
| 2810414N06I | 0.002 | 0.002 | 0.004 |
| Ciita | 0.002 | 0.003 | 0.001 |
| Gm47903 | 0.002 | 0.000 | 0.001 |
| AC160562.1 | 0.002 | 0.000 | 0.001 |
| Mmp9 | 0.002 | 0.001 | 0.000 |
| Acoxl | 0.002 | 0.000 | 0.001 |
| Slitrk4 | 0.002 | 0.022 | 0.063 |
| 0610038B21F | 0.002 | 0.012 | 0.017 |
| Mtmr7 | 0.002 | 0.031 | 0.008 |

| | | | |
|---|---|---|---|
| Synpr | 0.002 | 0.005 | 0.008 |
| Ttc16 | 0.002 | 0.004 | 0.007 |
| Tdrd1 | 0.002 | 0.000 | 0.001 |
| A830018L16F | 0.002 | 0.023 | 0.018 |
| Gm48742 | 0.002 | 0.002 | 0.002 |
| Cyp4f18 | 0.002 | 0.002 | 0.001 |
| Spata31d1a | 0.002 | 0.001 | 0.002 |
| Scamp4 | 0.002 | 0.001 | 0.001 |
| Gm5544 | 0.002 | 0.002 | 0.002 |
| Gm8206 | 0.002 | 0.000 | 0.000 |
| Fam110c | 0.002 | 0.001 | 0.000 |
| Psmb11 | 0.002 | 0.001 | 0.001 |
| Asgr1 | 0.002 | 0.002 | 0.001 |
| Zfp239 | 0.002 | 0.015 | 0.015 |
| Rimkla | 0.002 | 0.001 | 0.000 |
| Napsa | 0.002 | 0.001 | 0.001 |
| Slc6a15 | 0.002 | 0.007 | 0.004 |
| 5730507C01F | 0.002 | 0.006 | 0.011 |
| Rhcg | 0.002 | 0.005 | 0.000 |
| Fbxw25 | 0.002 | 0.000 | 0.000 |
| Fstl4 | 0.002 | 0.013 | 0.002 |
| Serac1 | 0.002 | 0.005 | 0.007 |
| Il10ra | 0.002 | 0.001 | 0.001 |
| Slc10a4 | 0.002 | 0.000 | 0.000 |
| Gm30934 | 0.002 | 0.002 | 0.002 |
| Muc16 | 0.002 | 0.002 | 0.001 |
| Tnfrsf13c | 0.002 | 0.001 | 0.001 |
| Gpr160 | 0.002 | 0.000 | 0.001 |
| Gm44709 | 0.002 | 0.000 | 0.000 |
| Sh2d4a | 0.002 | 0.004 | 0.003 |
| Gm32913 | 0.002 | 0.000 | 0.001 |
| Gm49173 | 0.002 | 0.000 | 0.000 |
| Gm37065 | 0.002 | 0.001 | 0.001 |
| Zfp354b | 0.002 | 0.008 | 0.009 |
| Mesp1 | 0.002 | 0.000 | 0.000 |
| C1ql3 | 0.002 | 0.030 | 0.024 |
| Scn8a | 0.002 | 0.002 | 0.011 |
| Plxdc1 | 0.002 | 0.007 | 0.004 |
| Gm31577 | 0.002 | 0.000 | 0.000 |
| Gm26590 | 0.002 | 0.002 | 0.002 |
| Syt2 | 0.002 | 0.006 | 0.007 |
| Kif5a | 0.002 | 0.007 | 0.010 |
| Dbndd1 | 0.002 | 0.002 | 0.001 |
| Gm34350 | 0.002 | 0.000 | 0.000 |
| Pcsk1n | 0.002 | 0.004 | 0.006 |
| Serpina9 | 0.002 | 0.001 | 0.001 |
| Unc79 | 0.002 | 0.001 | 0.002 |
| Cstad | 0.002 | 0.021 | 0.016 |

| | | | |
|---|---|---|---|
| Tnfrsf19 | 0.002 | 0.009 | 0.002 |
| Sirpb1c | 0.002 | 0.001 | 0.000 |
| Mcemp1 | 0.002 | 0.000 | 0.000 |
| Lrrc23 | 0.002 | 0.001 | 0.001 |
| Gm29243 | 0.002 | 0.000 | 0.000 |
| Nxph4 | 0.002 | 0.005 | 0.004 |
| Onecut2 | 0.002 | 0.003 | 0.001 |
| Gm40909 | 0.002 | 0.001 | 0.002 |
| Fut1 | 0.002 | 0.000 | 0.000 |
| Impg2 | 0.002 | 0.004 | 0.004 |
| Gm15624 | 0.002 | 0.000 | 0.000 |
| Gm29019 | 0.002 | 0.001 | 0.001 |
| Fat3 | 0.002 | 0.007 | 0.014 |
| Prr36 | 0.002 | 0.002 | 0.004 |
| Ankrd45 | 0.002 | 0.017 | 0.003 |
| Efcab5 | 0.002 | 0.001 | 0.002 |
| Gm136 | 0.002 | 0.000 | 0.000 |
| Gm16183 | 0.002 | 0.001 | 0.004 |
| Tmem215 | 0.002 | 0.116 | 0.035 |
| Reln | 0.002 | 0.003 | 0.003 |
| Gm32401 | 0.002 | 0.000 | 0.000 |
| Gm47128 | 0.002 | 0.001 | 0.001 |
| Apol6 | 0.002 | 0.012 | 0.005 |
| Dhrs7c | 0.002 | 0.031 | 0.008 |
| Ccdc78 | 0.002 | 0.002 | 0.003 |
| Tmem86b | 0.002 | 0.005 | 0.006 |
| Gm35248 | 0.002 | 0.001 | 0.002 |
| AL611987.1 | 0.002 | 0.000 | 0.000 |
| Gm48293 | 0.002 | 0.000 | 0.000 |
| Nrip3 | 0.002 | 0.001 | 0.001 |
| Rem2 | 0.002 | 0.002 | 0.002 |
| Ear2 | 0.002 | 0.000 | 0.001 |
| Kctd4 | 0.002 | 0.000 | 0.000 |
| AU023762 | 0.002 | 0.000 | 0.000 |
| Plet1os | 0.002 | 0.000 | 0.000 |
| Was | 0.002 | 0.002 | 0.002 |
| Gm43689 | 0.002 | 0.002 | 0.004 |
| Pck1 | 0.002 | 0.001 | 0.000 |
| 4921511C10I | 0.002 | 0.000 | 0.000 |
| Gm21738 | 0.002 | 0.000 | 0.000 |
| Fam186b | 0.002 | 0.001 | 0.000 |
| Gm15976 | 0.002 | 0.000 | 0.001 |
| Gm26676 | 0.002 | 0.001 | 0.004 |
| Cyp4x1 | 0.002 | 0.001 | 0.000 |
| Atp10b | 0.002 | 0.006 | 0.014 |
| Gm26877 | 0.002 | 0.002 | 0.003 |
| Aoah | 0.002 | 0.002 | 0.002 |
| 5430402O13I | 0.002 | 0.004 | 0.008 |

| | | | |
|---|---|---|---|
| Nefh | 0.002 | 0.001 | 0.000 |
| Gm36423 | 0.002 | 0.004 | 0.004 |
| Gm44037 | 0.002 | 0.001 | 0.001 |
| Plekhg6 | 0.002 | 0.001 | 0.001 |
| Foxred2 | 0.002 | 0.005 | 0.006 |
| Gm29155 | 0.002 | 0.000 | 0.001 |
| Ifit1bl1 | 0.002 | 0.000 | 0.001 |
| Amh | 0.002 | 0.002 | 0.002 |
| Tmem132e | 0.002 | 0.007 | 0.024 |
| Arhgap9 | 0.002 | 0.001 | 0.001 |
| Gm45441 | 0.002 | 0.001 | 0.005 |
| Rnf222 | 0.002 | 0.000 | 0.000 |
| Gm42846 | 0.002 | 0.001 | 0.000 |
| Apom | 0.002 | 0.001 | 0.000 |
| Nkain3 | 0.002 | 0.005 | 0.086 |
| Lrguk | 0.002 | 0.005 | 0.002 |
| AL589670.3 | 0.002 | 0.001 | 0.002 |
| Cd2 | 0.002 | 0.000 | 0.001 |
| Cysltr1 | 0.002 | 0.003 | 0.003 |
| D830036C21I | 0.002 | 0.014 | 0.017 |
| Ddn | 0.002 | 0.001 | 0.002 |
| C2cd4d | 0.002 | 0.003 | 0.008 |
| Ccdc158 | 0.002 | 0.009 | 0.012 |
| Gm39121 | 0.002 | 0.001 | 0.001 |
| Olfr1238 | 0.002 | 0.001 | 0.001 |
| 4732471J01R | 0.001 | 0.003 | 0.002 |
| Mobp | 0.001 | 0.000 | 0.001 |
| BC018473 | 0.001 | 0.002 | 0.001 |
| 1700065D16I | 0.001 | 0.001 | 0.002 |
| Insl6 | 0.001 | 0.010 | 0.004 |
| A630072L19F | 0.001 | 0.000 | 0.000 |
| Dqx1 | 0.001 | 0.003 | 0.004 |
| Hist1h2bf | 0.001 | 0.001 | 0.001 |
| Matn3 | 0.001 | 0.000 | 0.001 |
| Slc26a8 | 0.001 | 0.004 | 0.004 |
| Asb2 | 0.001 | 0.030 | 0.003 |
| Chst9 | 0.001 | 0.000 | 0.000 |
| Gm28043 | 0.001 | 0.001 | 0.002 |
| Gm13441 | 0.001 | 0.000 | 0.000 |
| Gm10062 | 0.001 | 0.000 | 0.000 |
| Gm26707 | 0.001 | 0.000 | 0.000 |
| A930002C04F | 0.001 | 0.014 | 0.006 |
| AC154683.1 | 0.001 | 0.000 | 0.000 |
| Ccl24 | 0.001 | 0.004 | 0.003 |
| Adcy2 | 0.001 | 0.025 | 0.015 |
| Hr | 0.001 | 0.007 | 0.006 |
| AI463229 | 0.001 | 0.000 | 0.000 |
| 2900045O20I | 0.001 | 0.002 | 0.001 |

| | | | |
|---|---|---|---|
| Gabrb3 | 0.001 | 0.004 | 0.010 |
| Gm26811 | 0.001 | 0.001 | 0.001 |
| Bhlhe22 | 0.001 | 0.000 | 0.002 |
| AC133498.1 | 0.001 | 0.000 | 0.000 |
| Ticam2 | 0.001 | 0.002 | 0.002 |
| Kif16bos | 0.001 | 0.000 | 0.000 |
| Gm46212 | 0.001 | 0.001 | 0.001 |
| Gm30382 | 0.001 | 0.019 | 0.007 |
| B9d1os | 0.001 | 0.001 | 0.001 |
| Tex14 | 0.001 | 0.000 | 0.002 |
| Itga11 | 0.001 | 0.017 | 0.032 |
| Adap2 | 0.001 | 0.001 | 0.001 |
| Gm26821 | 0.001 | 0.000 | 0.000 |
| Prf1 | 0.001 | 0.000 | 0.000 |
| 5930412G12I | 0.001 | 0.000 | 0.000 |
| Gm11940 | 0.001 | 0.001 | 0.003 |
| Jakmip2 | 0.001 | 0.000 | 0.002 |
| Havcr2 | 0.001 | 0.001 | 0.001 |
| Armc2 | 0.001 | 0.009 | 0.019 |
| Zfp985 | 0.001 | 0.000 | 0.000 |
| Gm20756 | 0.001 | 0.000 | 0.000 |
| Itgb7 | 0.001 | 0.002 | 0.001 |
| Plxnb3 | 0.001 | 0.001 | 0.000 |
| Capn12 | 0.001 | 0.006 | 0.008 |
| 4931422A03F | 0.001 | 0.000 | 0.000 |
| Gm44210 | 0.001 | 0.000 | 0.002 |
| B130046B21F | 0.001 | 0.002 | 0.002 |
| Fbxl13 | 0.001 | 0.005 | 0.017 |
| Olfr539 | 0.001 | 0.001 | 0.001 |
| Capn11 | 0.001 | 0.002 | 0.000 |
| Ly6g6c | 0.001 | 0.002 | 0.002 |
| Gm47882 | 0.001 | 0.001 | 0.001 |
| Gm43518 | 0.001 | 0.000 | 0.000 |
| Fhad1 | 0.001 | 0.005 | 0.002 |
| 4933424G06I | 0.001 | 0.003 | 0.001 |
| Cdh9 | 0.001 | 0.005 | 0.004 |
| Gm32926 | 0.001 | 0.000 | 0.000 |
| I830077J02Ri | 0.001 | 0.001 | 0.001 |
| Gm44081 | 0.001 | 0.001 | 0.001 |
| Prss50 | 0.001 | 0.000 | 0.000 |
| Gm17197 | 0.001 | 0.000 | 0.001 |
| 1700020L24F | 0.001 | 0.001 | 0.003 |
| Gm15956 | 0.001 | 0.004 | 0.004 |
| Gm26749 | 0.001 | 0.002 | 0.001 |
| Gm49179 | 0.001 | 0.009 | 0.011 |
| Wdr95 | 0.001 | 0.017 | 0.006 |
| Henmt1 | 0.001 | 0.001 | 0.001 |
| Elmod1 | 0.001 | 0.002 | 0.027 |

| | | | |
|---|---|---|---|
| Gm44751 | 0.001 | 0.004 | 0.001 |
| Tmem178b | 0.001 | 0.001 | 0.007 |
| Gm7879 | 0.001 | 0.001 | 0.002 |
| AC139347.1 | 0.001 | 0.002 | 0.004 |
| Obp2a | 0.001 | 0.000 | 0.000 |
| Tmem117 | 0.001 | 0.008 | 0.003 |
| Cblc | 0.001 | 0.001 | 0.002 |
| Gm19345 | 0.001 | 0.000 | 0.000 |
| Sstr2 | 0.001 | 0.000 | 0.000 |
| Ptgir | 0.001 | 0.011 | 0.029 |
| Gm13431 | 0.001 | 0.001 | 0.001 |
| Syndig1 | 0.001 | 0.003 | 0.003 |
| Gm10791 | 0.001 | 0.000 | 0.001 |
| Klkb1 | 0.001 | 0.001 | 0.000 |
| Sema3e | 0.001 | 0.008 | 0.002 |
| Zmynd10 | 0.001 | 0.001 | 0.000 |
| Fbxw27 | 0.001 | 0.000 | 0.000 |
| Gm128 | 0.001 | 0.004 | 0.003 |
| Asprv1 | 0.001 | 0.003 | 0.004 |
| Gm15632 | 0.001 | 0.001 | 0.000 |
| Fgfr4 | 0.001 | 0.001 | 0.001 |
| Gm41307 | 0.001 | 0.003 | 0.001 |
| Nectin4 | 0.001 | 0.000 | 0.001 |
| Gm28551 | 0.001 | 0.003 | 0.003 |
| Gm38708 | 0.001 | 0.000 | 0.000 |
| Wdr17 | 0.001 | 0.000 | 0.000 |
| Prss27 | 0.001 | 0.006 | 0.005 |
| Gm29113 | 0.001 | 0.001 | 0.001 |
| Gm11379 | 0.001 | 0.000 | 0.000 |
| 2810442N19I | 0.001 | 0.006 | 0.008 |
| Mymx | 0.001 | 0.000 | 0.001 |
| Gm13563 | 0.001 | 0.001 | 0.002 |
| Gm17216 | 0.001 | 0.002 | 0.006 |
| Gm49024 | 0.001 | 0.000 | 0.000 |
| G630030J09F | 0.001 | 0.001 | 0.001 |
| Cend1 | 0.001 | 0.012 | 0.012 |
| Ptpn22 | 0.001 | 0.001 | 0.000 |
| Krt20 | 0.001 | 0.001 | 0.009 |
| Hist1h3g | 0.001 | 0.001 | 0.001 |
| Gm37850 | 0.001 | 0.001 | 0.001 |
| Lrr1 | 0.001 | 0.002 | 0.006 |
| Gm11527 | 0.001 | 0.000 | 0.000 |
| Gm17088 | 0.001 | 0.000 | 0.000 |
| Gm45339 | 0.001 | 0.000 | 0.000 |
| Gm49375 | 0.001 | 0.000 | 0.000 |
| Eddm3b | 0.001 | 0.000 | 0.001 |
| Nkpd1 | 0.001 | 0.000 | 0.001 |
| Rgs13 | 0.001 | 0.000 | 0.002 |

| | | | |
|---|---|---|---|
| Fmr1os | 0.001 | 0.000 | 0.001 |
| Styxl1 | 0.001 | 0.001 | 0.000 |
| Gm38250 | 0.001 | 0.001 | 0.001 |
| Cma2 | 0.001 | 0.000 | 0.000 |
| Cyp11a1 | 0.001 | 0.001 | 0.000 |
| Unc5a | 0.001 | 0.001 | 0.003 |
| Hoxb9 | 0.001 | 0.000 | 0.000 |
| 6430531B16F | 0.001 | 0.002 | 0.002 |
| Bco1 | 0.001 | 0.001 | 0.000 |
| Gm47418 | 0.001 | 0.000 | 0.000 |
| Gm47079 | 0.001 | 0.001 | 0.002 |
| 4933407L21F | 0.001 | 0.001 | 0.002 |
| Ankdd1b | 0.001 | 0.000 | 0.020 |
| Gm49032 | 0.001 | 0.002 | 0.002 |
| Ccl5 | 0.001 | 0.002 | 0.002 |
| Arg2 | 0.001 | 0.001 | 0.001 |
| Cd300lf | 0.001 | 0.000 | 0.001 |
| Gm17597 | 0.001 | 0.000 | 0.001 |
| Gm17058 | 0.001 | 0.003 | 0.003 |
| Nkx2-5 | 0.001 | 0.008 | 0.006 |
| AC091463.1 | 0.001 | 0.000 | 0.000 |
| Gm26551 | 0.001 | 0.001 | 0.002 |
| Klk7 | 0.001 | 0.000 | 0.000 |
| Apod | 0.001 | 0.003 | 0.044 |
| Gm10093 | 0.001 | 0.004 | 0.002 |
| Marco | 0.001 | 0.000 | 0.000 |
| Foxa3 | 0.001 | 0.000 | 0.000 |
| Il5 | 0.001 | 0.000 | 0.000 |
| Abcc2 | 0.001 | 0.001 | 0.000 |
| Hist1h4n | 0.001 | 0.000 | 0.000 |
| Phospho1 | 0.001 | 0.000 | 0.000 |
| Gm46332 | 0.001 | 0.001 | 0.001 |
| Hoxa6 | 0.001 | 0.000 | 0.000 |
| Magel2 | 0.001 | 0.000 | 0.019 |
| Tmem30b | 0.001 | 0.003 | 0.016 |
| Gm26586 | 0.001 | 0.003 | 0.004 |
| Cdsn | 0.001 | 0.004 | 0.002 |
| Gm11615 | 0.001 | 0.000 | 0.000 |
| Gm34788 | 0.001 | 0.003 | 0.004 |
| Ms4a4d | 0.001 | 0.001 | 0.044 |
| Rslcan18 | 0.001 | 0.000 | 0.001 |
| Gm15138 | 0.001 | 0.001 | 0.000 |
| Pcdhgb7 | 0.001 | 0.006 | 0.005 |
| C330021F23F | 0.001 | 0.000 | 0.000 |
| Gm30074 | 0.001 | 0.003 | 0.003 |
| Gm28513 | 0.001 | 0.000 | 0.000 |
| Gm27032 | 0.001 | 0.000 | 0.000 |
| 1-Mar | 0.001 | 0.001 | 0.000 |

| | | | |
|---|---|---|---|
| Tspan10 | 0.001 | 0.000 | 0.000 |
| Tcrg-C1 | 0.001 | 0.000 | 0.000 |
| Ccdc162 | 0.001 | 0.009 | 0.005 |
| C1ql1 | 0.001 | 0.002 | 0.003 |
| Nptx2 | 0.001 | 0.046 | 0.005 |
| Gm44226 | 0.001 | 0.000 | 0.000 |
| 1700095J03R | 0.001 | 0.002 | 0.003 |
| AB124611 | 0.001 | 0.009 | 0.004 |
| Gm11335 | 0.001 | 0.001 | 0.000 |
| Zfp981 | 0.001 | 0.000 | 0.001 |
| Pilra | 0.001 | 0.001 | 0.001 |
| Ak8 | 0.001 | 0.000 | 0.001 |
| Msmp | 0.001 | 0.002 | 0.002 |
| Sla2 | 0.001 | 0.001 | 0.000 |
| Gm44626 | 0.001 | 0.000 | 0.000 |
| Gm44421 | 0.001 | 0.005 | 0.002 |
| Cyp2j8 | 0.001 | 0.000 | 0.000 |
| Mirt1 | 0.001 | 0.002 | 0.001 |
| Nlrp5 | 0.001 | 0.001 | 0.002 |
| B3galt2 | 0.001 | 0.001 | 0.001 |
| Gm34653 | 0.001 | 0.000 | 0.000 |
| Gm13285 | 0.001 | 0.001 | 0.001 |
| Lrch4 | 0.001 | 0.001 | 0.001 |
| Gm26795 | 0.001 | 0.001 | 0.000 |
| Aqp9 | 0.001 | 0.001 | 0.001 |
| Gm47093 | 0.001 | 0.002 | 0.006 |
| Gm5602 | 0.001 | 0.002 | 0.001 |
| Tmc5 | 0.001 | 0.000 | 0.000 |
| Bmp8a | 0.001 | 0.002 | 0.001 |
| Eno4 | 0.001 | 0.000 | 0.000 |
| Gm44275 | 0.001 | 0.000 | 0.000 |
| Bcas3os1 | 0.001 | 0.003 | 0.003 |
| 4930578C19F | 0.001 | 0.000 | 0.000 |
| Gm21994 | 0.001 | 0.003 | 0.007 |
| Ctnna2 | 0.001 | 0.001 | 0.002 |
| Mroh7 | 0.001 | 0.001 | 0.003 |
| Mlxipl | 0.001 | 0.004 | 0.003 |
| Ldhc | 0.001 | 0.001 | 0.001 |
| Them7 | 0.001 | 0.000 | 0.000 |
| Zfp965 | 0.001 | 0.002 | 0.002 |
| Gm10532 | 0.001 | 0.000 | 0.000 |
| Slc26a1 | 0.001 | 0.004 | 0.003 |
| Unc5cl | 0.001 | 0.000 | 0.000 |
| Gm16302 | 0.001 | 0.001 | 0.000 |
| Lgi1 | 0.001 | 0.004 | 0.072 |
| Avil | 0.001 | 0.006 | 0.008 |
| Gm19610 | 0.001 | 0.001 | 0.000 |
| A230065N10I | 0.001 | 0.000 | 0.001 |

| | | | |
|---|---|---|---|
| Gm48488 | 0.001 | 0.001 | 0.002 |
| Lif | 0.001 | 0.011 | 0.009 |
| Mrgprf | 0.001 | 0.014 | 0.065 |
| 1700034H15I | 0.001 | 0.002 | 0.000 |
| 1700092M07 | 0.001 | 0.001 | 0.000 |
| Lrrc2 | 0.001 | 0.001 | 0.000 |
| Ube2u | 0.001 | 0.004 | 0.003 |
| Gm38948 | 0.001 | 0.000 | 0.000 |
| Dydc1 | 0.001 | 0.000 | 0.000 |
| Dock3 | 0.001 | 0.001 | 0.001 |
| Zmynd12 | 0.001 | 0.002 | 0.002 |
| Pabpc5 | 0.001 | 0.007 | 0.009 |
| Gm29542 | 0.001 | 0.000 | 0.000 |
| Angptl7 | 0.001 | 0.039 | 0.121 |
| Snx22 | 0.001 | 0.001 | 0.002 |
| Gm45022 | 0.001 | 0.002 | 0.002 |
| Gm38414 | 0.001 | 0.000 | 0.000 |
| Tusc5 | 0.001 | 0.001 | 0.002 |
| Trim69 | 0.001 | 0.001 | 0.003 |
| Gm29590 | 0.001 | 0.000 | 0.000 |
| 4933430M04 | 0.001 | 0.000 | 0.000 |
| A530020G20I | 0.001 | 0.003 | 0.046 |
| Gm48942 | 0.001 | 0.001 | 0.000 |
| Gjc2 | 0.001 | 0.002 | 0.009 |
| Tmem45a2 | 0.001 | 0.001 | 0.000 |
| Gm17234 | 0.001 | 0.000 | 0.000 |
| Nell2 | 0.001 | 0.007 | 0.011 |
| Ptcra | 0.001 | 0.003 | 0.002 |
| Ankrd31 | 0.001 | 0.000 | 0.000 |
| Vav1 | 0.001 | 0.001 | 0.001 |
| Tmem254c | 0.001 | 0.001 | 0.002 |
| 4930512P04F | 0.001 | 0.000 | 0.000 |
| Xrra1 | 0.001 | 0.002 | 0.002 |
| Olfr433 | 0.001 | 0.000 | 0.000 |
| Tprg | 0.001 | 0.000 | 0.000 |
| Gm48004 | 0.001 | 0.000 | 0.000 |
| Thpo | 0.001 | 0.002 | 0.003 |
| 1700113H08I | 0.001 | 0.000 | 0.000 |
| Mpig6b | 0.001 | 0.000 | 0.001 |
| Gm12996 | 0.001 | 0.002 | 0.002 |
| Ccdc170 | 0.001 | 0.000 | 0.001 |
| Kcnk12 | 0.001 | 0.000 | 0.001 |
| Gm43507 | 0.001 | 0.000 | 0.000 |
| Hrk | 0.001 | 0.000 | 0.003 |
| Gm32834 | 0.001 | 0.004 | 0.001 |
| Tmem253 | 0.001 | 0.005 | 0.008 |
| Lct | 0.001 | 0.000 | 0.000 |
| Gm29430 | 0.001 | 0.003 | 0.002 |

| | | | |
|---|---|---|---|
| Fzd10 | 0.001 | 0.000 | 0.000 |
| Shank1 | 0.001 | 0.022 | 0.030 |
| Gm14827 | 0.001 | 0.001 | 0.001 |
| Col28a1 | 0.001 | 0.008 | 0.009 |
| Gm21284 | 0.001 | 0.001 | 0.001 |
| Kctd8 | 0.001 | 0.012 | 0.010 |
| Nppa | 0.001 | 0.001 | 0.000 |
| Fezf1 | 0.001 | 0.000 | 0.000 |
| Gpr141 | 0.001 | 0.000 | 0.000 |
| Gm33148 | 0.001 | 0.002 | 0.001 |
| Fam129c | 0.001 | 0.007 | 0.007 |
| Zfp972 | 0.001 | 0.000 | 0.000 |
| Kdm4d | 0.001 | 0.002 | 0.002 |
| Nap1l2 | 0.001 | 0.001 | 0.001 |
| Gemin4 | 0.001 | 0.001 | 0.001 |
| 2010013B24I | 0.001 | 0.000 | 0.000 |
| H1foo | 0.001 | 0.000 | 0.000 |
| Arhgap15 | 0.001 | 0.002 | 0.004 |
| Hist1h3b | 0.001 | 0.001 | 0.001 |
| Traf3ip3 | 0.001 | 0.001 | 0.002 |
| Ankrd61 | 0.001 | 0.001 | 0.001 |
| Dsg2 | 0.001 | 0.011 | 0.003 |
| Trim10 | 0.001 | 0.000 | 0.000 |
| Gm31793 | 0.001 | 0.000 | 0.000 |
| Hist2h2bb | 0.001 | 0.003 | 0.002 |
| Wif1 | 0.001 | 0.014 | 0.017 |
| Gpc5 | 0.001 | 0.000 | 0.000 |
| Vat1l | 0.001 | 0.001 | 0.009 |
| Gm20554 | 0.001 | 0.003 | 0.006 |
| Gm44710 | 0.001 | 0.001 | 0.000 |
| Cdrt4 | 0.001 | 0.001 | 0.007 |
| Abcg5 | 0.001 | 0.000 | 0.002 |
| Krt28 | 0.001 | 0.000 | 0.000 |
| Gm26884 | 0.001 | 0.002 | 0.001 |
| Gm26515 | 0.001 | 0.002 | 0.002 |
| 9130204K15I | 0.001 | 0.001 | 0.001 |
| Acat3 | 0.001 | 0.003 | 0.002 |
| Cacna1h | 0.001 | 0.030 | 0.017 |
| Fam166b | 0.001 | 0.001 | 0.001 |
| Gm17484 | 0.001 | 0.004 | 0.005 |
| Fam84a | 0.001 | 0.028 | 0.019 |
| P2rx3 | 0.001 | 0.000 | 0.000 |
| Masp2 | 0.001 | 0.001 | 0.002 |
| Gm26918 | 0.001 | 0.000 | 0.000 |
| Sst | 0.001 | 0.000 | 0.000 |
| Gm960 | 0.001 | 0.001 | 0.001 |
| Gm26825 | 0.001 | 0.009 | 0.007 |
| Car14 | 0.001 | 0.008 | 0.001 |

| | | | |
|---|---|---|---|
| Lax1 | 0.001 | 0.015 | 0.011 |
| Platr27 | 0.001 | 0.000 | 0.000 |
| Gm35082 | 0.001 | 0.003 | 0.003 |
| Foxg1 | 0.001 | 0.013 | 0.029 |
| Gm15731 | 0.001 | 0.001 | 0.000 |
| Macc1 | 0.001 | 0.002 | 0.001 |
| Gm27017 | 0.001 | 0.001 | 0.003 |
| Ggt6 | 0.001 | 0.000 | 0.000 |
| Cpne9 | 0.001 | 0.002 | 0.001 |
| Atp1a3 | 0.001 | 0.001 | 0.001 |
| Olfr77 | 0.001 | 0.003 | 0.002 |
| Gm3512 | 0.001 | 0.002 | 0.002 |
| Tctex1d1 | 0.001 | 0.000 | 0.000 |
| A930009A15F | 0.001 | 0.000 | 0.000 |
| Hist1h3f | 0.001 | 0.001 | 0.002 |
| Lrrc34 | 0.001 | 0.000 | 0.003 |
| Lrrc8e | 0.001 | 0.002 | 0.002 |
| Car2 | 0.001 | 0.001 | 0.001 |
| Gm13710 | 0.001 | 0.001 | 0.001 |
| Rgs9 | 0.001 | 0.002 | 0.003 |
| Gm16008 | 0.001 | 0.002 | 0.003 |
| Lbx2 | 0.001 | 0.002 | 0.001 |
| 1810011H11I | 0.001 | 0.000 | 0.000 |
| Gm15829 | 0.001 | 0.001 | 0.001 |
| AC167229.1 | 0.001 | 0.000 | 0.003 |
| Gm34623 | 0.001 | 0.000 | 0.000 |
| Gm32352 | 0.001 | 0.003 | 0.001 |
| Nog | 0.001 | 0.030 | 0.008 |
| Csf2rb2 | 0.001 | 0.001 | 0.007 |
| Itgax | 0.001 | 0.000 | 0.000 |
| Tsnaxip1 | 0.001 | 0.000 | 0.000 |
| Scn4a | 0.001 | 0.000 | 0.000 |
| Gm2093 | 0.001 | 0.000 | 0.000 |
| Lamp5 | 0.001 | 0.000 | 0.000 |
| Gm20716 | 0.001 | 0.000 | 0.000 |
| Gm46516 | 0.001 | 0.001 | 0.000 |
| Gm11168 | 0.001 | 0.000 | 0.000 |
| AC165271.1 | 0.001 | 0.004 | 0.002 |
| Gm37240 | 0.001 | 0.002 | 0.000 |
| Fabp12 | 0.001 | 0.000 | 0.000 |
| Sel1l3 | 0.001 | 0.015 | 0.005 |
| Gm43791 | 0.001 | 0.000 | 0.000 |
| Mfsd2a | 0.001 | 0.017 | 0.005 |
| Gm16230 | 0.001 | 0.001 | 0.002 |
| Gm11831 | 0.001 | 0.000 | 0.000 |
| Kcnh8 | 0.001 | 0.000 | 0.001 |
| Bin2 | 0.001 | 0.001 | 0.001 |
| Asz1 | 0.001 | 0.000 | 0.000 |

| | | | |
|---|---|---|---|
| Gm39526 | 0.001 | 0.000 | 0.000 |
| Gm11032 | 0.001 | 0.001 | 0.001 |
| Gdap1l1 | 0.001 | 0.001 | 0.002 |
| Fstl5 | 0.001 | 0.000 | 0.000 |
| Car5a | 0.001 | 0.002 | 0.001 |
| Gm34923 | 0.001 | 0.000 | 0.000 |
| Gm10778 | 0.001 | 0.000 | 0.000 |
| 2310043M15 | 0.001 | 0.000 | 0.000 |
| Coro2a | 0.001 | 0.001 | 0.000 |
| Tmem95 | 0.001 | 0.007 | 0.016 |
| Aknad1 | 0.001 | 0.002 | 0.006 |
| Gm48163 | 0.001 | 0.000 | 0.001 |
| Paqr9 | 0.001 | 0.002 | 0.002 |
| Gm35439 | 0.001 | 0.002 | 0.003 |
| Gm13465 | 0.001 | 0.000 | 0.000 |
| Fgf7 | 0.001 | 0.003 | 0.105 |
| Psg17 | 0.001 | 0.000 | 0.000 |
| Gm40761 | 0.001 | 0.000 | 0.000 |
| Ltb4r1 | 0.001 | 0.003 | 0.003 |
| Scin | 0.001 | 0.007 | 0.000 |
| Gm10800 | 0.001 | 0.001 | 0.001 |
| Gm13391 | 0.001 | 0.000 | 0.000 |
| Arl11 | 0.001 | 0.001 | 0.001 |
| Rab33a | 0.001 | 0.006 | 0.009 |
| Cd300a | 0.001 | 0.003 | 0.003 |
| Bend6 | 0.001 | 0.007 | 0.011 |
| Olfr365 | 0.001 | 0.000 | 0.000 |
| Gm14493 | 0.001 | 0.000 | 0.000 |
| Dscaml1 | 0.001 | 0.000 | 0.001 |
| Hist1h2an | 0.001 | 0.001 | 0.001 |
| Htr5b | 0.001 | 0.001 | 0.001 |
| Gm16010 | 0.001 | 0.001 | 0.000 |
| Gm44793 | 0.001 | 0.000 | 0.000 |
| Olfr731 | 0.001 | 0.000 | 0.001 |
| Gm20463 | 0.001 | 0.001 | 0.002 |
| Slc3a1 | 0.001 | 0.001 | 0.001 |
| Chrm1 | 0.001 | 0.001 | 0.001 |
| Lime1 | 0.001 | 0.000 | 0.000 |
| Ybx2 | 0.001 | 0.003 | 0.001 |
| Zfp385c | 0.001 | 0.001 | 0.002 |
| Gm17949 | 0.001 | 0.003 | 0.003 |
| Gm9945 | 0.001 | 0.001 | 0.001 |
| Stxbp5l | 0.001 | 0.002 | 0.002 |
| Rab25 | 0.001 | 0.000 | 0.001 |
| 4930579F01F | 0.001 | 0.000 | 0.000 |
| Cpt1b | 0.001 | 0.001 | 0.001 |
| Gpr65 | 0.001 | 0.002 | 0.001 |
| Mast1 | 0.001 | 0.000 | 0.001 |

| | | | |
|---|---|---|---|
| Rab17 | 0.001 | 0.005 | 0.001 |
| Car1 | 0.001 | 0.000 | 0.000 |
| Gm44296 | 0.001 | 0.001 | 0.000 |
| Klhl40 | 0.001 | 0.007 | 0.007 |
| 1700094M23 | 0.001 | 0.000 | 0.000 |
| Lbhd2 | 0.001 | 0.017 | 0.010 |
| Gm26508 | 0.001 | 0.004 | 0.002 |
| Snph | 0.001 | 0.000 | 0.001 |
| Gm8773 | 0.001 | 0.004 | 0.007 |
| 1700025N21I | 0.001 | 0.000 | 0.000 |
| Hcrtr2 | 0.001 | 0.001 | 0.001 |
| Fgf9 | 0.001 | 0.003 | 0.014 |
| 1700061I17R | 0.001 | 0.000 | 0.000 |
| Tigd4 | 0.001 | 0.004 | 0.001 |
| Hsd17b1 | 0.001 | 0.001 | 0.002 |
| Olfr129 | 0.001 | 0.000 | 0.000 |
| Lrrc9 | 0.001 | 0.000 | 0.000 |
| Gm26721 | 0.001 | 0.005 | 0.004 |
| Dkkl1 | 0.001 | 0.001 | 0.001 |
| Fmo3 | 0.001 | 0.008 | 0.004 |
| Gpr37l1 | 0.001 | 0.001 | 0.001 |
| Gm5128 | 0.001 | 0.000 | 0.000 |
| Cyp2e1 | 0.001 | 0.000 | 0.000 |
| Tmem116 | 0.001 | 0.003 | 0.004 |
| Gm9984 | 0.001 | 0.000 | 0.000 |
| Rnf208 | 0.001 | 0.002 | 0.004 |
| Clec5a | 0.001 | 0.000 | 0.001 |
| Cfap46 | 0.001 | 0.001 | 0.000 |
| Gm26706 | 0.001 | 0.001 | 0.000 |
| Gm27253 | 0.001 | 0.001 | 0.002 |
| Gm1070 | 0.001 | 0.000 | 0.001 |
| Slc9b2 | 0.001 | 0.007 | 0.004 |
| Pilrb1 | 0.001 | 0.000 | 0.000 |
| 4833412C05F | 0.001 | 0.021 | 0.030 |
| Prox2os | 0.001 | 0.004 | 0.004 |
| B430305J03R | 0.001 | 0.000 | 0.001 |
| Ifi205 | 0.001 | 0.001 | 0.002 |
| Gm10785 | 0.001 | 0.004 | 0.005 |
| Gm34574 | 0.001 | 0.000 | 0.000 |
| Aloxe3 | 0.001 | 0.001 | 0.003 |
| Gm13420 | 0.001 | 0.000 | 0.000 |
| P2ry10 | 0.001 | 0.000 | 0.004 |
| Gm44698 | 0.001 | 0.000 | 0.001 |
| Car4 | 0.001 | 0.000 | 0.000 |
| B230311B06F | 0.001 | 0.000 | 0.004 |
| Islr2 | 0.001 | 0.001 | 0.026 |
| 1700061N14I | 0.001 | 0.001 | 0.001 |
| Gcgr | 0.001 | 0.000 | 0.002 |

| | | | |
|---|---|---|---|
| Pkib | 0.001 | 0.009 | 0.002 |
| Brsk2 | 0.001 | 0.001 | 0.001 |
| A230107N01I | 0.001 | 0.001 | 0.000 |
| Snx31 | 0.001 | 0.000 | 0.002 |
| AC166832.3 | 0.001 | 0.000 | 0.000 |
| Slfn4 | 0.001 | 0.000 | 0.000 |
| Gm45670 | 0.001 | 0.037 | 0.005 |
| Gm28051 | 0.001 | 0.000 | 0.000 |
| Gm28424 | 0.001 | 0.002 | 0.003 |
| Ifitm7 | 0.001 | 0.003 | 0.006 |
| Slitrk6 | 0.001 | 0.006 | 0.014 |
| Tmed6 | 0.001 | 0.001 | 0.001 |
| Gm32014 | 0.001 | 0.000 | 0.001 |
| Msx2 | 0.001 | 0.054 | 0.009 |
| Gm47701 | 0.001 | 0.001 | 0.001 |
| Lrrc46 | 0.001 | 0.002 | 0.003 |
| Gm16143 | 0.001 | 0.000 | 0.000 |
| 4921514A10I | 0.001 | 0.002 | 0.001 |
| Kcng3 | 0.001 | 0.000 | 0.000 |
| Hist2h3b | 0.001 | 0.001 | 0.001 |
| Slc27a5 | 0.001 | 0.000 | 0.000 |
| Actbl2 | 0.001 | 0.000 | 0.001 |
| Relt | 0.001 | 0.006 | 0.007 |
| Sptbn2 | 0.001 | 0.001 | 0.001 |
| Hnf1b | 0.001 | 0.000 | 0.000 |
| Myom3 | 0.001 | 0.017 | 0.008 |
| Kcnip1 | 0.001 | 0.001 | 0.005 |
| Caps2 | 0.001 | 0.003 | 0.002 |
| Zar1 | 0.001 | 0.000 | 0.000 |
| Dchs2 | 0.001 | 0.007 | 0.046 |
| Otoa | 0.001 | 0.004 | 0.011 |
| Tmem121b | 0.001 | 0.001 | 0.001 |
| Spib | 0.001 | 0.000 | 0.000 |
| Zfp488 | 0.001 | 0.000 | 0.000 |
| Gm3331 | 0.001 | 0.001 | 0.001 |
| Edaradd | 0.001 | 0.001 | 0.002 |
| Gm26863 | 0.001 | 0.000 | 0.000 |
| AC144408.3 | 0.001 | 0.002 | 0.001 |
| Vwa3b | 0.001 | 0.017 | 0.009 |
| 9230111E07I | 0.001 | 0.002 | 0.001 |
| Pinc | 0.001 | 0.000 | 0.000 |
| Arhgdig | 0.001 | 0.009 | 0.023 |
| Gm14341 | 0.001 | 0.001 | 0.000 |
| Tal2 | 0.001 | 0.002 | 0.001 |
| Gm17203 | 0.001 | 0.003 | 0.003 |
| Elf3 | 0.001 | 0.001 | 0.000 |
| Nr2e1 | 0.001 | 0.000 | 0.000 |
| Gm6685 | 0.001 | 0.000 | 0.000 |

| | | | |
|---|---|---|---|
| Pga5 | 0.001 | 0.000 | 0.000 |
| Olfr1555-ps1 | 0.001 | 0.000 | 0.000 |
| Gm43936 | 0.001 | 0.000 | 0.002 |
| Gm40538 | 0.001 | 0.000 | 0.000 |
| 1110002E22F | 0.001 | 0.003 | 0.008 |
| Cdh23 | 0.001 | 0.003 | 0.002 |
| Trim29 | 0.001 | 0.003 | 0.003 |
| Agbl2 | 0.001 | 0.003 | 0.003 |
| Ptprn | 0.001 | 0.005 | 0.002 |
| Gm32585 | 0.001 | 0.001 | 0.001 |
| Gm17281 | 0.001 | 0.001 | 0.001 |
| Eif2c5 | 0.001 | 0.004 | 0.002 |
| Gm40557 | 0.001 | 0.001 | 0.001 |
| Ifi211 | 0.001 | 0.002 | 0.017 |
| Gm48653 | 0.001 | 0.001 | 0.002 |
| Prlh | 0.001 | 0.000 | 0.000 |
| Gm48708 | 0.001 | 0.001 | 0.002 |
| Adam28 | 0.001 | 0.001 | 0.001 |
| Gm16552 | 0.001 | 0.005 | 0.003 |
| Creb3l3 | 0.001 | 0.002 | 0.048 |
| Rgs20 | 0.001 | 0.000 | 0.001 |
| Gm32569 | 0.001 | 0.000 | 0.002 |
| Gm38102 | 0.001 | 0.001 | 0.000 |
| Vmn1r58 | 0.001 | 0.000 | 0.000 |
| Gm4316 | 0.001 | 0.001 | 0.001 |
| Pla2g1b | 0.001 | 0.000 | 0.000 |
| Gm28035 | 0.001 | 0.001 | 0.001 |
| Gm48250 | 0.001 | 0.000 | 0.000 |
| Il19 | 0.001 | 0.000 | 0.000 |
| Kncn | 0.001 | 0.000 | 0.001 |
| A430057M04 | 0.001 | 0.001 | 0.001 |
| Lilr4b | 0.001 | 0.002 | 0.003 |
| Gm5431 | 0.001 | 0.000 | 0.000 |
| Sez6l | 0.001 | 0.003 | 0.002 |
| Gm47950 | 0.001 | 0.001 | 0.000 |
| Gm44264 | 0.001 | 0.000 | 0.000 |
| Acox2 | 0.001 | 0.000 | 0.000 |
| Gm43653 | 0.001 | 0.002 | 0.002 |
| Gm19385 | 0.001 | 0.000 | 0.000 |
| Trh | 0.001 | 0.007 | 0.005 |
| Prss8 | 0.001 | 0.000 | 0.000 |
| Slc24a1 | 0.001 | 0.001 | 0.001 |
| Gm31600 | 0.001 | 0.000 | 0.000 |
| Gm10389 | 0.001 | 0.000 | 0.000 |
| Oacyl | 0.001 | 0.006 | 0.003 |
| Gm17096 | 0.001 | 0.002 | 0.001 |
| Gm42420 | 0.001 | 0.000 | 0.000 |
| Prss57 | 0.001 | 0.005 | 0.003 |

| | | | |
|---|---|---|---|
| Lair1 | 0.001 | 0.002 | 0.001 |
| Lmx1a | 0.001 | 0.000 | 0.004 |
| Gm38335 | 0.001 | 0.000 | 0.000 |
| Fam159a | 0.001 | 0.001 | 0.002 |
| Slc5a9 | 0.001 | 0.000 | 0.000 |
| Krtdap | 0.001 | 0.002 | 0.050 |
| Avpr1a | 0.001 | 0.001 | 0.012 |
| Ankrd55 | 0.001 | 0.000 | 0.003 |
| Tdrd9 | 0.001 | 0.000 | 0.001 |
| Mcoln2 | 0.001 | 0.001 | 0.001 |
| Myh3 | 0.001 | 0.012 | 0.002 |
| Frs3os | 0.001 | 0.000 | 0.000 |
| Nlrp3 | 0.001 | 0.001 | 0.001 |
| 4930573C15F | 0.001 | 0.001 | 0.000 |
| Gm44067 | 0.001 | 0.000 | 0.000 |
| Agrp | 0.001 | 0.002 | 0.003 |
| Cideb | 0.001 | 0.003 | 0.003 |
| Chia1 | 0.001 | 0.000 | 0.000 |
| Tdrd12 | 0.001 | 0.001 | 0.001 |
| Gm14252 | 0.001 | 0.000 | 0.000 |
| C130026L21F | 0.001 | 0.002 | 0.009 |
| Pla2g4b | 0.001 | 0.000 | 0.000 |
| Hsd17b13 | 0.001 | 0.001 | 0.006 |
| Mybphl | 0.001 | 0.002 | 0.001 |
| Hfe2 | 0.001 | 0.001 | 0.000 |
| Gm13067 | 0.001 | 0.000 | 0.000 |
| A530095I07R | 0.001 | 0.024 | 0.012 |
| Gm3248 | 0.001 | 0.002 | 0.001 |
| Pgc | 0.001 | 0.000 | 0.000 |
| AL592187.3 | 0.001 | 0.000 | 0.000 |
| Gm20548 | 0.001 | 0.002 | 0.000 |
| Lin7a | 0.001 | 0.010 | 0.011 |
| F930015N05F | 0.001 | 0.002 | 0.006 |
| Ly6k | 0.001 | 0.000 | 0.004 |
| 2010110K18F | 0.001 | 0.000 | 0.000 |
| Gm17102 | 0.001 | 0.000 | 0.001 |
| AC166344.1 | 0.001 | 0.000 | 0.000 |
| Efcab8 | 0.001 | 0.001 | 0.001 |
| Clec4n | 0.001 | 0.004 | 0.005 |
| Klhl38 | 0.001 | 0.004 | 0.001 |
| Lrrc7 | 0.001 | 0.018 | 0.010 |
| Olfr986 | 0.001 | 0.001 | 0.001 |
| A630023A22F | 0.001 | 0.012 | 0.002 |
| Gm10306 | 0.001 | 0.001 | 0.001 |
| Gabrg3 | 0.001 | 0.000 | 0.023 |
| Gm13610 | 0.001 | 0.002 | 0.001 |
| Ankrd2 | 0.001 | 0.001 | 0.001 |
| AC140186.1 | 0.001 | 0.000 | 0.000 |

| | | | |
|---|---|---|---|
| Adcy10 | 0.001 | 0.002 | 0.002 |
| Gm43915 | 0.001 | 0.004 | 0.004 |
| Ptchd4 | 0.001 | 0.001 | 0.003 |
| Gm5799 | 0.001 | 0.000 | 0.000 |
| Cyp2d26 | 0.001 | 0.002 | 0.000 |
| Gm15489 | 0.001 | 0.001 | 0.000 |
| 4930477G07I | 0.001 | 0.000 | 0.000 |
| Gm17546 | 0.001 | 0.000 | 0.000 |
| B4galnt2 | 0.001 | 0.000 | 0.000 |
| Mapk10 | 0.001 | 0.003 | 0.004 |
| Agbl1 | 0.001 | 0.000 | 0.000 |
| Krt14 | 0.001 | 0.002 | 0.001 |
| Olfr1167 | 0.001 | 0.000 | 0.000 |
| Nudt8 | 0.001 | 0.001 | 0.001 |
| Calcb | 0.001 | 0.003 | 0.005 |
| Gm2564 | 0.001 | 0.006 | 0.019 |
| Gm42504 | 0.001 | 0.002 | 0.004 |
| Plcxd1 | 0.001 | 0.001 | 0.001 |
| Gm26525 | 0.001 | 0.001 | 0.003 |
| Esrrg | 0.001 | 0.004 | 0.021 |
| Gm30270 | 0.001 | 0.000 | 0.000 |
| Gm26751 | 0.001 | 0.001 | 0.001 |
| Atp1a4 | 0.001 | 0.000 | 0.001 |
| Gm26748 | 0.001 | 0.000 | 0.000 |
| Gm30954 | 0.001 | 0.000 | 0.000 |
| 4933406G16I | 0.001 | 0.001 | 0.002 |
| Gm48770 | 0.001 | 0.000 | 0.000 |
| Armc3 | 0.001 | 0.004 | 0.001 |
| Adig | 0.001 | 0.001 | 0.001 |
| Gm42901 | 0.001 | 0.004 | 0.003 |
| Slc22a13 | 0.001 | 0.001 | 0.000 |
| Cfap53 | 0.001 | 0.004 | 0.007 |
| 5730559C18I | 0.001 | 0.000 | 0.000 |
| 4930429F24I | 0.001 | 0.001 | 0.000 |
| Rgs9bp | 0.001 | 0.002 | 0.001 |
| Catsperd | 0.001 | 0.001 | 0.001 |
| Gm7932 | 0.001 | 0.000 | 0.000 |
| Gm15929 | 0.001 | 0.002 | 0.002 |
| Gm43221 | 0.001 | 0.003 | 0.005 |
| Olfr1322 | 0.001 | 0.000 | 0.001 |
| Gm45774 | 0.001 | 0.002 | 0.004 |
| Trat1 | 0.001 | 0.000 | 0.002 |
| Jph4 | 0.001 | 0.007 | 0.024 |
| Chst4 | 0.001 | 0.001 | 0.000 |
| Mmp27 | 0.001 | 0.000 | 0.001 |
| Gm5464 | 0.001 | 0.001 | 0.003 |
| Mgat5b | 0.001 | 0.001 | 0.000 |
| Gm34868 | 0.001 | 0.001 | 0.001 |

| | | | |
|---|---|---|---|
| 1700109K24F | 0.001 | 0.000 | 0.000 |
| 4930430E12F | 0.001 | 0.000 | 0.000 |
| Pitpnm2os2 | 0.001 | 0.001 | 0.000 |
| Gm4961 | 0.001 | 0.000 | 0.000 |
| Zic5 | 0.001 | 0.000 | 0.000 |
| Klhl10 | 0.001 | 0.002 | 0.003 |
| Spata46 | 0.001 | 0.000 | 0.000 |
| Gm29261 | 0.001 | 0.000 | 0.000 |
| Tmem184a | 0.001 | 0.001 | 0.001 |
| Gm10069 | 0.001 | 0.008 | 0.011 |
| Gm20512 | 0.001 | 0.003 | 0.002 |
| Pnldc1 | 0.001 | 0.005 | 0.004 |
| Kcnt1 | 0.001 | 0.010 | 0.003 |
| Nefl | 0.001 | 0.002 | 0.009 |
| Gal3st3 | 0.001 | 0.028 | 0.004 |
| Cntnap2 | 0.001 | 0.001 | 0.003 |
| Trp53cor1 | 0.001 | 0.003 | 0.004 |
| Gm29292 | 0.001 | 0.002 | 0.002 |
| Tfap2c | 0.001 | 0.000 | 0.000 |
| Gm21541 | 0.001 | 0.000 | 0.000 |
| 1700011L22F | 0.001 | 0.001 | 0.001 |
| Lcp2 | 0.001 | 0.002 | 0.002 |
| Rbmxl2 | 0.001 | 0.000 | 0.000 |
| Tcp10a | 0.001 | 0.000 | 0.000 |
| Vmn1r4 | 0.001 | 0.001 | 0.001 |
| A830052D11I | 0.001 | 0.001 | 0.003 |
| Fdx1l | 0.001 | 0.001 | 0.001 |
| Gm33785 | 0.001 | 0.000 | 0.000 |
| Gm12905 | 0.001 | 0.000 | 0.001 |
| Hnf1a | 0.001 | 0.000 | 0.000 |
| Phf11a | 0.001 | 0.001 | 0.000 |
| Mettl7a3 | 0.001 | 0.000 | 0.000 |
| Gm15217 | 0.001 | 0.000 | 0.000 |
| Gm13383 | 0.001 | 0.000 | 0.000 |
| Fam83f | 0.001 | 0.000 | 0.001 |
| S100a7a | 0.001 | 0.003 | 0.009 |
| Armh1 | 0.001 | 0.002 | 0.012 |
| Gucy1b2 | 0.001 | 0.001 | 0.005 |
| Hoxd4.1 | 0.001 | 0.000 | 0.000 |
| Klhl41 | 0.001 | 0.001 | 0.001 |
| Gm13074 | 0.001 | 0.000 | 0.000 |
| Gm29773 | 0.001 | 0.001 | 0.001 |
| Kif18b | 0.001 | 0.002 | 0.003 |
| Gm43713 | 0.001 | 0.005 | 0.003 |
| Abca6 | 0.001 | 0.004 | 0.062 |
| 1700064M15 | 0.001 | 0.001 | 0.003 |
| Gm42902 | 0.001 | 0.000 | 0.001 |
| Sox5os3 | 0.001 | 0.000 | 0.000 |

| | | | |
|---|---|---|---|
| Gm45768 | 0.001 | 0.000 | 0.002 |
| Nr1i2 | 0.001 | 0.000 | 0.000 |
| 4930500M09 | 0.001 | 0.002 | 0.001 |
| D5Ertd615e | 0.001 | 0.000 | 0.000 |
| Gm26851 | 0.001 | 0.000 | 0.000 |
| Gm9999 | 0.001 | 0.000 | 0.000 |
| Dbpht2 | 0.001 | 0.002 | 0.004 |
| Ifi204 | 0.001 | 0.003 | 0.027 |
| Mpzl2 | 0.001 | 0.001 | 0.001 |
| Gm16933 | 0.001 | 0.002 | 0.002 |
| Ephx4 | 0.001 | 0.004 | 0.013 |
| Gm10719 | 0.001 | 0.000 | 0.000 |
| 1810008I18R | 0.001 | 0.000 | 0.000 |
| Adra1a | 0.001 | 0.000 | 0.001 |
| Gm20732 | 0.001 | 0.000 | 0.000 |
| Gm3667 | 0.001 | 0.001 | 0.000 |
| Prr15l | 0.001 | 0.001 | 0.001 |
| Pcdha4 | 0.001 | 0.000 | 0.000 |
| Gsdma | 0.001 | 0.001 | 0.001 |
| Wbscr25 | 0.001 | 0.002 | 0.004 |
| Gm44175 | 0.001 | 0.006 | 0.004 |
| Cox8b | 0.001 | 0.002 | 0.007 |
| 1700013M08 | 0.001 | 0.000 | 0.000 |
| 9130015G15I | 0.001 | 0.005 | 0.001 |
| Gm49123 | 0.001 | 0.001 | 0.003 |
| Higd1b | 0.001 | 0.000 | 0.001 |
| 2310026L22F | 0.001 | 0.001 | 0.001 |
| Fbxo24 | 0.001 | 0.001 | 0.001 |
| Etv4 | 0.001 | 0.003 | 0.013 |
| Ccl19 | 0.001 | 0.006 | 0.025 |
| Ccdc105 | 0.001 | 0.000 | 0.000 |
| Cldn9 | 0.001 | 0.001 | 0.006 |
| E030044B06F | 0.001 | 0.000 | 0.000 |
| Gm42742 | 0.001 | 0.001 | 0.003 |
| BC030343 | 0.001 | 0.001 | 0.003 |
| Kif27 | 0.001 | 0.002 | 0.003 |
| Gm20139 | 0.001 | 0.000 | 0.000 |
| 1700016K19F | 0.001 | 0.000 | 0.000 |
| Plekhg4 | 0.001 | 0.000 | 0.001 |
| Ephx3 | 0.001 | 0.005 | 0.005 |
| 1700128A07F | 0.001 | 0.000 | 0.000 |
| Rhov | 0.001 | 0.004 | 0.004 |
| Zfp296 | 0.001 | 0.001 | 0.001 |
| Myo15 | 0.001 | 0.001 | 0.001 |
| Gm28041 | 0.001 | 0.000 | 0.001 |
| Lipg | 0.001 | 0.001 | 0.002 |
| Slc35f3 | 0.001 | 0.000 | 0.000 |
| Gm45538 | 0.001 | 0.000 | 0.000 |

| | | | |
|---|---|---|---|
| Gm12111 | 0.001 | 0.002 | 0.006 |
| Fcgr4 | 0.001 | 0.002 | 0.001 |
| Tifab | 0.001 | 0.001 | 0.001 |
| Gm26675 | 0.001 | 0.001 | 0.000 |
| Uchl1os | 0.001 | 0.000 | 0.000 |
| AC153364.1 | 0.001 | 0.000 | 0.000 |
| Inpp5j | 0.001 | 0.007 | 0.001 |
| Rasgrp1 | 0.001 | 0.001 | 0.001 |
| Gm20518 | 0.001 | 0.000 | 0.000 |
| Gm15690 | 0.001 | 0.004 | 0.002 |
| Alkal2 | 0.001 | 0.001 | 0.027 |
| Gm11008 | 0.001 | 0.001 | 0.001 |
| Gm17130 | 0.001 | 0.000 | 0.000 |
| Evi2 | 0.001 | 0.000 | 0.000 |
| Clec4d | 0.001 | 0.001 | 0.000 |
| Clcnka | 0.001 | 0.000 | 0.000 |
| Gm10490 | 0.001 | 0.000 | 0.000 |
| Zfp92 | 0.001 | 0.000 | 0.000 |
| Dusp9 | 0.001 | 0.007 | 0.003 |
| Cdcp1 | 0.001 | 0.000 | 0.000 |
| Gm21269 | 0.001 | 0.002 | 0.002 |
| Gm28940 | 0.001 | 0.000 | 0.000 |
| Myo18b | 0.001 | 0.007 | 0.003 |
| Gast | 0.001 | 0.002 | 0.001 |
| 1810013D15I | 0.001 | 0.000 | 0.000 |
| Rasef | 0.001 | 0.000 | 0.002 |
| AC158554.1 | 0.001 | 0.003 | 0.002 |
| Pramef12os | 0.001 | 0.000 | 0.001 |
| Gm45437 | 0.001 | 0.000 | 0.000 |
| Hist2h2aa2 | 0.001 | 0.000 | 0.000 |
| Dusp13 | 0.001 | 0.000 | 0.001 |
| Atp2a1 | 0.001 | 0.000 | 0.001 |
| Gm35520 | 0.001 | 0.000 | 0.000 |
| Slc22a20 | 0.001 | 0.000 | 0.000 |
| Gm26578 | 0.001 | 0.000 | 0.000 |
| Upk3bl | 0.001 | 0.000 | 0.007 |
| Rnf43 | 0.001 | 0.000 | 0.001 |
| Hist1h2ak | 0.001 | 0.001 | 0.001 |
| Mx1 | 0.001 | 0.001 | 0.001 |
| Slpi | 0.001 | 0.000 | 0.000 |
| Gm34411 | 0.001 | 0.000 | 0.000 |
| Hrh1 | 0.001 | 0.000 | 0.000 |
| Clec12a | 0.001 | 0.001 | 0.000 |
| Stfa2l1 | 0.001 | 0.000 | 0.001 |
| BB123696 | 0.001 | 0.019 | 0.011 |
| Rfx8 | 0.001 | 0.002 | 0.001 |
| Aldob | 0.001 | 0.000 | 0.000 |
| Insl5 | 0.001 | 0.000 | 0.000 |

| | | | |
|---|---|---|---|
| 4933428P19F | 0.001 | 0.002 | 0.002 |
| Gjd4 | 0.001 | 0.000 | 0.001 |
| Zfp558 | 0.001 | 0.003 | 0.004 |
| Gm26841 | 0.001 | 0.001 | 0.002 |
| Prrt2 | 0.001 | 0.002 | 0.002 |
| Gm35584 | 0.001 | 0.000 | 0.000 |
| Platr22 | 0.001 | 0.000 | 0.000 |
| Gm38134 | 0.001 | 0.000 | 0.001 |
| Gm11541 | 0.001 | 0.001 | 0.030 |
| Gm10801 | 0.001 | 0.000 | 0.000 |
| Gm16240 | 0.001 | 0.000 | 0.000 |
| Gm26860 | 0.001 | 0.001 | 0.001 |
| Cyp17a1 | 0.001 | 0.000 | 0.000 |
| Gm15601 | 0.001 | 0.001 | 0.001 |
| C130073E24F | 0.001 | 0.003 | 0.001 |
| Gm16630 | 0.001 | 0.001 | 0.001 |
| Vmn2r97 | 0.001 | 0.001 | 0.000 |
| Gm29325 | 0.001 | 0.000 | 0.003 |
| 9330175M20 | 0.001 | 0.001 | 0.001 |
| Anxa10 | 0.001 | 0.000 | 0.000 |
| Sftpc | 0.001 | 0.001 | 0.000 |
| Krt76 | 0.001 | 0.000 | 0.000 |
| Awat2 | 0.001 | 0.000 | 0.001 |
| Lmo3 | 0.001 | 0.000 | 0.005 |
| Gm26887 | 0.001 | 0.001 | 0.002 |
| Ntn5 | 0.001 | 0.001 | 0.001 |
| Gm15609 | 0.001 | 0.000 | 0.000 |
| Gipr | 0.001 | 0.028 | 0.019 |
| Cfap74 | 0.001 | 0.003 | 0.013 |
| Pla2g4f | 0.001 | 0.000 | 0.000 |
| A330015K06F | 0.001 | 0.001 | 0.002 |
| Krtap28-13 | 0.001 | 0.002 | 0.001 |
| Gpr3 | 0.001 | 0.006 | 0.003 |
| Gm44763 | 0.001 | 0.003 | 0.007 |
| Vmn1r5 | 0.001 | 0.000 | 0.000 |
| Agtr1b | 0.001 | 0.003 | 0.008 |
| Kif17 | 0.001 | 0.003 | 0.009 |
| Plin1 | 0.001 | 0.004 | 0.004 |
| Hist1h3i | 0.001 | 0.002 | 0.001 |
| Tmem265 | 0.001 | 0.000 | 0.000 |
| Gm27241 | 0.001 | 0.001 | 0.000 |
| Gm44000 | 0.001 | 0.000 | 0.001 |
| Gm40614 | 0.001 | 0.000 | 0.000 |
| Vtn | 0.001 | 0.001 | 0.002 |
| Gm3417 | 0.001 | 0.000 | 0.000 |
| Gm8994 | 0.001 | 0.000 | 0.001 |
| Gm11592 | 0.001 | 0.000 | 0.000 |
| 9330111N05I | 0.001 | 0.001 | 0.004 |

| | | | |
|---|---|---|---|
| Adgrg5 | 0.001 | 0.000 | 0.000 |
| Cryba1 | 0.001 | 0.001 | 0.000 |
| Gm20635 | 0.001 | 0.002 | 0.004 |
| Gm26619 | 0.001 | 0.001 | 0.003 |
| Gm15612 | 0.001 | 0.001 | 0.001 |
| Cd19 | 0.001 | 0.000 | 0.000 |
| Grm6 | 0.001 | 0.000 | 0.000 |
| Tmem200b | 0.001 | 0.001 | 0.002 |
| Gm47578 | 0.001 | 0.000 | 0.000 |
| A330102I10R | 0.001 | 0.002 | 0.001 |
| Gm47520 | 0.001 | 0.000 | 0.000 |
| Npffr1 | 0.001 | 0.003 | 0.000 |
| Arpp21 | 0.001 | 0.020 | 0.005 |
| Aqp8 | 0.001 | 0.000 | 0.000 |
| Slc17a8 | 0.001 | 0.001 | 0.001 |
| Hsh2d | 0.001 | 0.001 | 0.001 |
| Zdhhc11 | 0.001 | 0.000 | 0.000 |
| Serpina3c | 0.001 | 0.001 | 0.027 |
| Npas4 | 0.001 | 0.016 | 0.004 |
| Tagln3 | 0.001 | 0.010 | 0.004 |
| Gm37662 | 0.001 | 0.001 | 0.000 |
| Uckl1os | 0.001 | 0.001 | 0.000 |
| Ppbp | 0.001 | 0.000 | 0.000 |
| Gm49373 | 0.001 | 0.001 | 0.001 |
| Ccdc153 | 0.001 | 0.000 | 0.000 |
| Gm36752 | 0.001 | 0.000 | 0.000 |
| Cyp2d11 | 0.001 | 0.000 | 0.000 |
| Serinc4 | 0.001 | 0.002 | 0.002 |
| Cadm2 | 0.001 | 0.002 | 0.002 |
| Cox7b2 | 0.001 | 0.000 | 0.001 |
| 4930579G18I | 0.001 | 0.001 | 0.001 |
| Klhl31 | 0.001 | 0.006 | 0.006 |
| Mlana | 0.001 | 0.001 | 0.002 |
| 1700022I11R | 0.001 | 0.001 | 0.001 |
| Gm12827 | 0.001 | 0.000 | 0.000 |
| Sgpp2 | 0.001 | 0.000 | 0.000 |
| Gpa33 | 0.001 | 0.000 | 0.000 |
| Olfr1195 | 0.001 | 0.000 | 0.000 |
| Clvs1 | 0.001 | 0.001 | 0.000 |
| Gm12536 | 0.001 | 0.000 | 0.000 |
| Dlgap3 | 0.001 | 0.002 | 0.007 |
| Grid2ip | 0.001 | 0.001 | 0.001 |
| Gm45058 | 0.001 | 0.000 | 0.000 |
| B4galnt4 | 0.001 | 0.003 | 0.007 |
| 4933411E08F | 0.001 | 0.000 | 0.002 |
| Grb7 | 0.001 | 0.000 | 0.001 |
| Hoxd3 | 0.001 | 0.016 | 0.010 |
| Pnmal1 | 0.001 | 0.000 | 0.001 |

| | | | |
|---|---|---|---|
| Gm32468 | 0.001 | 0.000 | 0.000 |
| Gm26611 | 0.001 | 0.000 | 0.000 |
| Nrap | 0.001 | 0.002 | 0.002 |
| Fndc9 | 0.001 | 0.000 | 0.000 |
| Hsd11b2 | 0.001 | 0.003 | 0.061 |
| Mapk8ip2 | 0.001 | 0.001 | 0.003 |
| Gm36584 | 0.001 | 0.000 | 0.000 |
| AC061963.1 | 0.001 | 0.001 | 0.002 |
| 6720483E21F | 0.001 | 0.001 | 0.001 |
| Gm16754 | 0.001 | 0.001 | 0.001 |
| Gm37637 | 0.001 | 0.001 | 0.001 |
| Krt15 | 0.001 | 0.016 | 0.005 |
| Gm47112 | 0.001 | 0.001 | 0.002 |
| Col10a1 | 0.001 | 0.013 | 0.001 |
| Gm11454 | 0.001 | 0.000 | 0.000 |
| Mylk4 | 0.001 | 0.052 | 0.005 |
| AC101921.1 | 0.001 | 0.001 | 0.000 |
| Mc2r | 0.001 | 0.000 | 0.004 |
| 1700093K21F | 0.001 | 0.004 | 0.002 |
| Sgca | 0.001 | 0.008 | 0.004 |
| F630042J09R | 0.001 | 0.000 | 0.000 |
| Myoz1 | 0.001 | 0.005 | 0.002 |
| Gm26792 | 0.001 | 0.001 | 0.001 |
| Rho | 0.001 | 0.000 | 0.001 |
| Vwa5b2 | 0.001 | 0.002 | 0.001 |
| Vtcn1 | 0.001 | 0.008 | 0.001 |
| Cngb3 | 0.001 | 0.002 | 0.001 |
| Trim71 | 0.001 | 0.000 | 0.000 |
| Chit1 | 0.001 | 0.000 | 0.001 |
| Gm15774 | 0.001 | 0.001 | 0.000 |
| Palm2 | 0.001 | 0.052 | 0.017 |
| Eppk1 | 0.001 | 0.003 | 0.005 |
| Klhl30 | 0.001 | 0.008 | 0.001 |
| Efcab12 | 0.001 | 0.001 | 0.001 |
| Myoz2 | 0.001 | 0.030 | 0.006 |
| Gm43916 | 0.001 | 0.000 | 0.000 |
| Mag | 0.001 | 0.002 | 0.002 |
| Luzp2 | 0.001 | 0.002 | 0.007 |
| Zc3h12d | 0.001 | 0.001 | 0.001 |
| Dclk3 | 0.001 | 0.006 | 0.001 |
| 1700047I17R | 0.001 | 0.001 | 0.001 |
| Gm14055 | 0.001 | 0.000 | 0.000 |
| Gm12974 | 0.001 | 0.000 | 0.000 |
| Hapln4 | 0.001 | 0.000 | 0.000 |
| Gm9821 | 0.001 | 0.001 | 0.000 |
| Gm17399 | 0.001 | 0.001 | 0.003 |
| Pla2g3 | 0.001 | 0.001 | 0.002 |
| Elovl2 | 0.001 | 0.001 | 0.002 |

| | | | |
|---|---|---|---|
| Gdap1 | 0.001 | 0.003 | 0.003 |
| Gm5082 | 0.001 | 0.000 | 0.000 |
| Plekhd1os | 0.001 | 0.000 | 0.000 |
| Syne4 | 0.001 | 0.003 | 0.005 |
| Gm44899 | 0.001 | 0.001 | 0.001 |
| Gm47003 | 0.001 | 0.001 | 0.001 |
| Gm15943 | 0.001 | 0.002 | 0.001 |
| Gm20163 | 0.001 | 0.002 | 0.006 |
| Skor1 | 0.001 | 0.001 | 0.001 |
| Trdn | 0.001 | 0.001 | 0.001 |
| Rbp3 | 0.001 | 0.000 | 0.000 |
| Tmem198 | 0.001 | 0.014 | 0.013 |
| Gpr83 | 0.001 | 0.001 | 0.003 |
| Sec1 | 0.001 | 0.001 | 0.001 |
| Slc22a2 | 0.001 | 0.000 | 0.001 |
| Gm15543 | 0.001 | 0.001 | 0.001 |
| Kcnh3 | 0.001 | 0.003 | 0.001 |
| Tescl | 0.001 | 0.000 | 0.001 |
| Adcyap1r1 | 0.001 | 0.001 | 0.012 |
| Dnmt3aos | 0.001 | 0.001 | 0.001 |
| Mccc1os | 0.001 | 0.001 | 0.001 |
| Chrna1 | 0.001 | 0.000 | 0.000 |
| Gm10401 | 0.001 | 0.004 | 0.003 |
| Snai3 | 0.001 | 0.000 | 0.000 |
| Gm20652 | 0.001 | 0.000 | 0.001 |
| Gm15582 | 0.001 | 0.000 | 0.000 |
| Gm26574 | 0.001 | 0.000 | 0.000 |
| Gm15232 | 0.001 | 0.000 | 0.000 |
| Gm9754 | 0.001 | 0.001 | 0.001 |
| Stag3 | 0.001 | 0.002 | 0.003 |
| Gm49324 | 0.001 | 0.000 | 0.000 |
| Gm47398 | 0.001 | 0.001 | 0.000 |
| Gm5447 | 0.001 | 0.000 | 0.000 |
| Gm16892 | 0.001 | 0.000 | 0.001 |
| Gm15340 | 0.001 | 0.000 | 0.000 |
| Gm11672 | 0.001 | 0.000 | 0.000 |
| Atxn7l1os1 | 0.001 | 0.000 | 0.000 |
| Hrasls | 0.001 | 0.002 | 0.000 |
| Kcnj13 | 0.001 | 0.003 | 0.002 |
| Gm8066 | 0.001 | 0.001 | 0.001 |
| Gm15816 | 0.001 | 0.001 | 0.000 |
| Gfap | 0.001 | 0.000 | 0.001 |
| Gpr62 | 0.001 | 0.003 | 0.006 |
| Gpr75 | 0.001 | 0.001 | 0.001 |
| 9330161L09F | 0.001 | 0.000 | 0.000 |
| 5033417F24F | 0.001 | 0.004 | 0.003 |
| Plppr1 | 0.001 | 0.008 | 0.012 |
| Slc28a1 | 0.001 | 0.000 | 0.001 |

| | | | |
|---|---|---|---|
| Slc44a3 | 0.001 | 0.003 | 0.019 |
| Gm17300 | 0.001 | 0.002 | 0.001 |
| Tas2r137 | 0.001 | 0.000 | 0.000 |
| D830030K20I | 0.001 | 0.008 | 0.006 |
| Gm26836 | 0.001 | 0.003 | 0.002 |
| Flvcr2 | 0.001 | 0.002 | 0.006 |
| Gm29455 | 0.001 | 0.000 | 0.000 |
| Gabra2 | 0.001 | 0.003 | 0.009 |
| Gm7173 | 0.001 | 0.000 | 0.001 |
| Chil1 | 0.001 | 0.001 | 0.001 |
| 5430427O19I | 0.001 | 0.001 | 0.000 |
| Gm13269 | 0.001 | 0.015 | 0.003 |
| 4931423N10I | 0.001 | 0.001 | 0.001 |
| Klk1 | 0.001 | 0.000 | 0.000 |
| Gm28989 | 0.001 | 0.000 | 0.000 |
| 6430710C18F | 0.001 | 0.000 | 0.001 |
| Gm29417 | 0.001 | 0.002 | 0.003 |
| Gm3488 | 0.001 | 0.000 | 0.000 |
| Fa2h | 0.001 | 0.001 | 0.000 |
| A830009L08F | 0.001 | 0.001 | 0.001 |
| Tmem89 | 0.001 | 0.001 | 0.001 |
| Gm43064 | 0.001 | 0.000 | 0.000 |
| Ccdc40 | 0.001 | 0.002 | 0.004 |
| B230359F08F | 0.001 | 0.000 | 0.000 |
| Rnf151 | 0.001 | 0.001 | 0.001 |
| Zmiz1os1 | 0.001 | 0.003 | 0.003 |
| Irx5 | 0.001 | 0.002 | 0.007 |
| Muc3a | 0.001 | 0.000 | 0.000 |
| Lcn8 | 0.001 | 0.004 | 0.007 |
| Ins2 | 0.001 | 0.001 | 0.002 |
| Bbox1 | 0.001 | 0.002 | 0.004 |
| A730017C20F | 0.001 | 0.006 | 0.017 |
| Nyap1 | 0.001 | 0.033 | 0.023 |
| Gm37168 | 0.001 | 0.001 | 0.000 |
| Gabbr2 | 0.001 | 0.000 | 0.001 |
| Fgfbp1 | 0.001 | 0.000 | 0.001 |
| Gm3739 | 0.001 | 0.005 | 0.002 |
| Ank1 | 0.001 | 0.003 | 0.007 |
| Gm20275 | 0.001 | 0.000 | 0.001 |
| Cfap44 | 0.001 | 0.000 | 0.000 |
| Atp8b4 | 0.001 | 0.000 | 0.002 |
| Gm4793 | 0.001 | 0.003 | 0.003 |
| Fam71f2 | 0.001 | 0.003 | 0.002 |
| Sv2b | 0.001 | 0.008 | 0.005 |
| Adcyap1 | 0.001 | 0.002 | 0.000 |
| Ptpn7 | 0.001 | 0.001 | 0.000 |
| Gm43674 | 0.001 | 0.001 | 0.001 |
| AC125351.1 | 0.001 | 0.000 | 0.000 |

| | | | |
|---|---|---|---|
| Gm20658 | 0.001 | 0.000 | 0.000 |
| Hist1h3c | 0.001 | 0.002 | 0.002 |
| Gm12932 | 0.001 | 0.001 | 0.000 |
| Gm26641 | 0.001 | 0.001 | 0.001 |
| Fam69c | 0.001 | 0.011 | 0.009 |
| Gfra3 | 0.001 | 0.002 | 0.001 |
| Acrv1 | 0.001 | 0.000 | 0.000 |
| Gm10549 | 0.001 | 0.000 | 0.001 |
| Tlr5 | 0.001 | 0.004 | 0.011 |
| 2900011O08I | 0.001 | 0.002 | 0.003 |
| Gm47127 | 0.001 | 0.001 | 0.001 |
| 2310047D07I | 0.001 | 0.002 | 0.003 |
| Gm32250 | 0.001 | 0.001 | 0.001 |
| Fmr1nb | 0.001 | 0.001 | 0.001 |
| Kcnmb2 | 0.001 | 0.000 | 0.000 |
| Gm46204 | 0.001 | 0.000 | 0.000 |
| Gm26850 | 0.001 | 0.001 | 0.002 |
| Pcdhgb1 | 0.001 | 0.003 | 0.002 |
| P2ry12 | 0.001 | 0.001 | 0.001 |
| Folr2 | 0.001 | 0.004 | 0.004 |
| AC133868.2 | 0.001 | 0.002 | 0.001 |
| Gm13133 | 0.001 | 0.000 | 0.000 |
| Anks1b | 0.001 | 0.002 | 0.003 |
| Esm1 | 0.001 | 0.001 | 0.006 |
| Soga3 | 0.001 | 0.000 | 0.000 |
| Gm12088 | 0.001 | 0.000 | 0.000 |
| Smim22 | 0.001 | 0.001 | 0.001 |
| Tspan1 | 0.001 | 0.000 | 0.000 |
| Gm49380 | 0.001 | 0.002 | 0.002 |
| Gm39459 | 0.001 | 0.002 | 0.001 |
| Lonrf2 | 0.001 | 0.012 | 0.028 |
| Gm2163 | 0.001 | 0.002 | 0.004 |
| H60c | 0.001 | 0.000 | 0.000 |
| Gm31036 | 0.001 | 0.000 | 0.000 |
| Lrrc66 | 0.001 | 0.000 | 0.000 |
| Vmn1r201 | 0.001 | 0.000 | 0.000 |
| Kif6 | 0.001 | 0.008 | 0.004 |
| Gm29237 | 0.001 | 0.001 | 0.002 |
| Baiap3 | 0.001 | 0.002 | 0.001 |
| Gm26981 | 0.001 | 0.005 | 0.008 |
| Atp6v1b1 | 0.001 | 0.000 | 0.000 |
| 2010005H15I | 0.001 | 0.001 | 0.000 |
| Gm16093 | 0.001 | 0.001 | 0.001 |
| Ibsp | 0.001 | 0.001 | 0.000 |
| Ido1 | 0.001 | 0.000 | 0.000 |
| C7 | 0.001 | 0.000 | 0.001 |
| Cntn6 | 0.001 | 0.007 | 0.046 |
| 5430427N15I | 0.001 | 0.000 | 0.002 |

| | | | |
|---|---|---|---|
| Gm28048 | 0.001 | 0.001 | 0.001 |
| Hist2h2ab | 0.001 | 0.000 | 0.000 |
| Crhr2 | 0.001 | 0.002 | 0.000 |
| A730043L09F | 0.001 | 0.000 | 0.000 |
| Spem1 | 0.001 | 0.009 | 0.017 |
| Gm35842 | 0.001 | 0.001 | 0.001 |
| Itgae | 0.001 | 0.001 | 0.001 |
| Lrrc74b | 0.001 | 0.000 | 0.001 |
| Gm26840 | 0.001 | 0.002 | 0.003 |
| Gm11802 | 0.001 | 0.000 | 0.000 |
| Cdhr1 | 0.001 | 0.004 | 0.002 |
| A830005F24F | 0.001 | 0.000 | 0.001 |
| Gm48899 | 0.001 | 0.002 | 0.003 |
| Tmc1 | 0.001 | 0.000 | 0.000 |
| Nxph1 | 0.001 | 0.006 | 0.016 |
| Gm31718 | 0.001 | 0.001 | 0.002 |
| Hils1 | 0.001 | 0.000 | 0.002 |
| Slc22a8 | 0.001 | 0.000 | 0.000 |
| Sh2d6 | 0.001 | 0.001 | 0.002 |
| Ifi208 | 0.001 | 0.000 | 0.000 |
| Gm27222 | 0.001 | 0.002 | 0.005 |
| Myl3 | 0.001 | 0.001 | 0.002 |
| Gm48265 | 0.001 | 0.000 | 0.000 |
| Kcnq3 | 0.001 | 0.004 | 0.001 |
| Gm36028 | 0.001 | 0.001 | 0.000 |
| Il1r2 | 0.001 | 0.001 | 0.028 |
| Spint1 | 0.001 | 0.000 | 0.000 |
| Gm30873 | 0.001 | 0.004 | 0.010 |
| Tjp3 | 0.001 | 0.000 | 0.000 |
| Prss34 | 0.001 | 0.000 | 0.000 |
| Lbx1 | 0.001 | 0.000 | 0.000 |
| Gm15990 | 0.001 | 0.000 | 0.000 |
| Tenm2 | 0.001 | 0.001 | 0.003 |
| Psg16 | 0.001 | 0.000 | 0.000 |
| 6030466F02F | 0.001 | 0.000 | 0.000 |
| Large2 | 0.001 | 0.001 | 0.001 |
| Ctcfl | 0.001 | 0.001 | 0.000 |
| Chl1 | 0.001 | 0.002 | 0.014 |
| Gm16105 | 0.001 | 0.001 | 0.000 |
| Chdh | 0.001 | 0.004 | 0.002 |
| Dppa5a | 0.001 | 0.002 | 0.000 |
| Gm46411 | 0.001 | 0.000 | 0.000 |
| Stx19 | 0.001 | 0.002 | 0.002 |
| Gpr152 | 0.001 | 0.001 | 0.001 |
| Gm43154 | 0.001 | 0.000 | 0.000 |
| 1700007G11I | 0.001 | 0.001 | 0.001 |
| Nos1 | 0.001 | 0.001 | 0.001 |
| Ptx4 | 0.001 | 0.003 | 0.011 |

| | | | |
|---|---|---|---|
| 4930519A11F | 0.001 | 0.000 | 0.000 |
| Gm1992 | 0.001 | 0.000 | 0.000 |
| Gm38304 | 0.001 | 0.001 | 0.001 |
| Gm11549 | 0.001 | 0.033 | 0.005 |
| Grm8 | 0.001 | 0.000 | 0.000 |
| Gm20110 | 0.001 | 0.000 | 0.000 |
| Gm16196 | 0.001 | 0.001 | 0.001 |
| Gm17224 | 0.001 | 0.001 | 0.000 |
| Gm12580 | 0.001 | 0.000 | 0.002 |
| Gm47770 | 0.001 | 0.000 | 0.000 |
| Gm47922 | 0.001 | 0.000 | 0.000 |
| 0610012D04F | 0.001 | 0.000 | 0.001 |
| Gm26785 | 0.001 | 0.004 | 0.004 |
| Omp | 0.001 | 0.000 | 0.001 |
| 4930486L24F | 0.001 | 0.000 | 0.000 |
| Gm26742 | 0.001 | 0.000 | 0.000 |
| Sox2 | 0.001 | 0.003 | 0.002 |
| Gm15657 | 0.001 | 0.000 | 0.000 |
| Olfr1393 | 0.001 | 0.001 | 0.002 |
| Cacna2d4 | 0.001 | 0.000 | 0.000 |
| Tex15 | 0.001 | 0.012 | 0.012 |
| Gm9946 | 0.001 | 0.001 | 0.000 |
| Selenov | 0.001 | 0.003 | 0.005 |
| Lncbate1 | 0.001 | 0.002 | 0.002 |
| Ms4a8a | 0.001 | 0.000 | 0.000 |
| Qprt | 0.001 | 0.001 | 0.017 |
| Phldb3 | 0.001 | 0.002 | 0.000 |
| Ascl3 | 0.001 | 0.000 | 0.000 |
| Efhc2 | 0.001 | 0.003 | 0.000 |
| 3110039M20 | 0.001 | 0.007 | 0.011 |
| Gm17276 | 0.001 | 0.000 | 0.000 |
| Tex26 | 0.001 | 0.002 | 0.006 |
| Gm17029 | 0.001 | 0.001 | 0.001 |
| Aire | 0.001 | 0.000 | 0.000 |
| 4930425P05F | 0.001 | 0.001 | 0.003 |
| Gm12236 | 0.001 | 0.000 | 0.000 |
| 6530409C15F | 0.001 | 0.001 | 0.001 |
| Gm33497 | 0.001 | 0.000 | 0.000 |
| Lzts1 | 0.001 | 0.003 | 0.004 |
| Gm8225 | 0.001 | 0.001 | 0.001 |
| Gm12927 | 0.001 | 0.001 | 0.000 |
| D030047H15I | 0.001 | 0.000 | 0.000 |
| Mogat2 | 0.001 | 0.001 | 0.001 |
| Mchr1 | 0.001 | 0.002 | 0.002 |
| Oas3 | 0.001 | 0.001 | 0.001 |
| 9430078K24F | 0.001 | 0.001 | 0.003 |
| 3-Sep | 0.001 | 0.005 | 0.005 |
| Slitrk2 | 0.001 | 0.001 | 0.004 |

| | | | |
|---|---|---|---|
| Casq2 | 0.001 | 0.005 | 0.003 |
| Pm20d1 | 0.001 | 0.000 | 0.001 |
| Odf1 | 0.001 | 0.000 | 0.000 |
| Khdrbs2 | 0.001 | 0.022 | 0.016 |
| Gm15265 | 0.001 | 0.003 | 0.003 |
| Gm37696 | 0.001 | 0.000 | 0.000 |
| 0610005C13F | 0.001 | 0.002 | 0.002 |
| Mettl7b | 0.001 | 0.000 | 0.000 |
| Btn2a2 | 0.001 | 0.000 | 0.001 |
| Gp5 | 0.001 | 0.001 | 0.001 |
| Gm36888 | 0.001 | 0.000 | 0.000 |
| Zbtb32 | 0.001 | 0.000 | 0.001 |
| 4933428G20I | 0.001 | 0.000 | 0.001 |
| Gm15879 | 0.001 | 0.002 | 0.003 |
| Sspo | 0.001 | 0.001 | 0.000 |
| 2600014E21F | 0.001 | 0.001 | 0.010 |
| B430218F22F | 0.001 | 0.001 | 0.001 |
| Map3k19 | 0.001 | 0.000 | 0.000 |
| Aox3 | 0.001 | 0.001 | 0.010 |
| 6530437J22R | 0.001 | 0.006 | 0.006 |
| Gm15965 | 0.001 | 0.002 | 0.000 |
| Lrtm1 | 0.001 | 0.000 | 0.000 |
| Esr2 | 0.001 | 0.001 | 0.000 |
| Coro6 | 0.001 | 0.010 | 0.003 |
| 4930548H24I | 0.001 | 0.002 | 0.002 |
| Gm41231 | 0.001 | 0.000 | 0.001 |
| Grm3 | 0.001 | 0.006 | 0.012 |
| Gm26700 | 0.001 | 0.000 | 0.000 |
| Pirt | 0.001 | 0.001 | 0.003 |
| Gm28192 | 0.001 | 0.003 | 0.005 |
| 4632411P08F | 0.001 | 0.001 | 0.001 |
| Cd48 | 0.001 | 0.001 | 0.001 |
| Tlr7 | 0.001 | 0.000 | 0.001 |
| Clec4a3 | 0.001 | 0.002 | 0.003 |
| Klra2 | 0.001 | 0.001 | 0.000 |
| Ccr2 | 0.001 | 0.001 | 0.001 |
| Gm44053 | 0.001 | 0.003 | 0.001 |
| AC134548.2 | 0.001 | 0.002 | 0.001 |
| Padi1 | 0.001 | 0.001 | 0.001 |
| Klk5 | 0.001 | 0.000 | 0.000 |
| Slc35g1 | 0.001 | 0.001 | 0.004 |
| Gm28913 | 0.001 | 0.000 | 0.002 |
| Zfp536 | 0.001 | 0.002 | 0.004 |
| Dpy19l2 | 0.001 | 0.000 | 0.000 |
| Grin2a | 0.001 | 0.004 | 0.006 |
| 4930445N18I | 0.001 | 0.000 | 0.001 |
| Cyp26c1 | 0.001 | 0.000 | 0.000 |
| Kif24 | 0.001 | 0.001 | 0.001 |

| | | | |
|---|---|---|---|
| Gm17705 | 0.001 | 0.001 | 0.001 |
| Camkv | 0.001 | 0.000 | 0.001 |
| 2810459M11 | 0.001 | 0.000 | 0.001 |
| Olfr1263 | 0.001 | 0.000 | 0.000 |
| Gm14434 | 0.001 | 0.000 | 0.000 |
| Gm12790 | 0.001 | 0.000 | 0.000 |
| Ksr2 | 0.001 | 0.001 | 0.002 |
| Fgf21 | 0.001 | 0.000 | 0.000 |
| D7Ertd443e | 0.001 | 0.000 | 0.001 |
| Cd3g | 0.001 | 0.000 | 0.000 |
| Gm48678 | 0.001 | 0.000 | 0.000 |
| Gm29168 | 0.001 | 0.000 | 0.000 |
| Ifnlr1 | 0.001 | 0.001 | 0.001 |
| Edn2 | 0.001 | 0.008 | 0.002 |
| Tmem182 | 0.001 | 0.001 | 0.000 |
| Opn1sw | 0.001 | 0.007 | 0.005 |
| Kcnb2 | 0.001 | 0.008 | 0.052 |
| Pax6 | 0.001 | 0.003 | 0.003 |
| Gm26799 | 0.001 | 0.001 | 0.001 |
| Nps | 0.001 | 0.001 | 0.001 |
| Gm20234 | 0.001 | 0.000 | 0.000 |
| Gm15978 | 0.001 | 0.000 | 0.000 |
| Rnf17 | 0.001 | 0.002 | 0.004 |
| Adam21 | 0.001 | 0.000 | 0.000 |
| Odf4 | 0.001 | 0.000 | 0.000 |
| 9430018G01I | 0.001 | 0.000 | 0.000 |
| Cd37 | 0.001 | 0.001 | 0.003 |
| Gm2694 | 0.001 | 0.001 | 0.008 |
| Ddx25 | 0.001 | 0.002 | 0.006 |
| Krt78 | 0.001 | 0.000 | 0.001 |
| 1500035N22I | 0.001 | 0.005 | 0.001 |
| Gm17590 | 0.001 | 0.004 | 0.002 |
| Gm13185 | 0.001 | 0.000 | 0.000 |
| Cass4 | 0.001 | 0.000 | 0.000 |
| Gm43118 | 0.001 | 0.001 | 0.000 |
| Vsnl1 | 0.001 | 0.002 | 0.002 |
| AC154507.3 | 0.001 | 0.004 | 0.001 |
| Gm45194 | 0.001 | 0.000 | 0.000 |
| Gm9970 | 0.001 | 0.001 | 0.001 |
| Psors1c2 | 0.001 | 0.003 | 0.002 |
| Gm12295 | 0.001 | 0.003 | 0.000 |
| Nanp | 0.001 | 0.001 | 0.002 |
| Pou4f1 | 0.001 | 0.002 | 0.001 |
| Dlx2 | 0.001 | 0.013 | 0.002 |
| Myo1a | 0.001 | 0.001 | 0.001 |
| Bcl2a1b | 0.001 | 0.005 | 0.005 |
| Pcdhb6 | 0.001 | 0.005 | 0.006 |
| Gm47565 | 0.001 | 0.001 | 0.000 |

| | | | |
|---|---|---|---|
| S100a3 | 0.001 | 0.000 | 0.001 |
| Cpne4 | 0.001 | 0.001 | 0.000 |
| Ggt5 | 0.001 | 0.042 | 0.053 |
| Catspere2 | 0.001 | 0.004 | 0.004 |
| Catsperz | 0.001 | 0.002 | 0.001 |
| Oscar | 0.001 | 0.001 | 0.000 |
| Gm3227 | 0.001 | 0.001 | 0.001 |
| C130026I21R | 0.001 | 0.000 | 0.002 |
| S100a5 | 0.001 | 0.000 | 0.000 |
| Rsf1os2 | 0.001 | 0.000 | 0.000 |
| Fam159b | 0.001 | 0.001 | 0.000 |
| Gm47439 | 0.001 | 0.001 | 0.001 |
| A230083N12I | 0.001 | 0.002 | 0.002 |
| Tepp | 0.001 | 0.001 | 0.002 |
| Gm2670 | 0.001 | 0.000 | 0.000 |
| Olfr1391 | 0.001 | 0.000 | 0.000 |
| Gm47096 | 0.001 | 0.001 | 0.003 |
| R3hdml | 0.001 | 0.000 | 0.000 |
| Zfp978 | 0.001 | 0.000 | 0.000 |
| A930018P22F | 0.001 | 0.001 | 0.000 |
| 4930455H04I | 0.001 | 0.000 | 0.000 |
| Trpm2 | 0.001 | 0.000 | 0.000 |
| Nat1 | 0.001 | 0.000 | 0.000 |
| Gm43625 | 0.001 | 0.006 | 0.006 |
| P2ry13 | 0.001 | 0.000 | 0.001 |
| Pla2g4c | 0.001 | 0.002 | 0.002 |
| Ahrr | 0.001 | 0.002 | 0.003 |
| Gm1141 | 0.001 | 0.001 | 0.001 |
| Ido2 | 0.001 | 0.000 | 0.000 |
| Gm29724 | 0.001 | 0.000 | 0.001 |
| Vpreb2 | 0.001 | 0.000 | 0.000 |
| Fgf23 | 0.001 | 0.001 | 0.008 |
| Gm12278 | 0.001 | 0.001 | 0.001 |
| Ankrd35 | 0.001 | 0.010 | 0.025 |
| Slc25a31 | 0.001 | 0.001 | 0.001 |
| Ccl11 | 0.001 | 0.003 | 0.052 |
| Ifnz | 0.001 | 0.000 | 0.001 |
| 5730460C07F | 0.001 | 0.000 | 0.000 |
| Elfn2 | 0.001 | 0.000 | 0.000 |
| Gm47400 | 0.001 | 0.000 | 0.001 |
| Gm12474 | 0.001 | 0.000 | 0.000 |
| Zfp957 | 0.001 | 0.000 | 0.000 |
| Zswim5 | 0.001 | 0.003 | 0.001 |
| Dlec1 | 0.001 | 0.001 | 0.001 |
| 4930539E08F | 0.001 | 0.000 | 0.002 |
| Gm45606 | 0.001 | 0.000 | 0.000 |
| Rtl9 | 0.001 | 0.000 | 0.001 |
| Gm44696 | 0.001 | 0.001 | 0.001 |

| | | | |
|---|---|---|---|
| AC157566.4 | 0.001 | 0.000 | 0.001 |
| Stpg1 | 0.001 | 0.001 | 0.001 |
| Slc23a4 | 0.001 | 0.003 | 0.003 |
| Brinp3 | 0.001 | 0.004 | 0.023 |
| Abca14 | 0.001 | 0.000 | 0.000 |
| AI463170 | 0.001 | 0.007 | 0.002 |
| Chrna4 | 0.001 | 0.001 | 0.020 |
| Dsc2 | 0.001 | 0.006 | 0.005 |
| Gm13782 | 0.001 | 0.000 | 0.000 |
| Cpa6 | 0.001 | 0.001 | 0.000 |
| AI115009 | 0.001 | 0.000 | 0.002 |
| Gm26762 | 0.001 | 0.001 | 0.000 |
| Gm39090 | 0.001 | 0.002 | 0.004 |
| Apba2 | 0.001 | 0.001 | 0.002 |
| 4933411E06F | 0.001 | 0.004 | 0.004 |
| Egr4 | 0.001 | 0.011 | 0.014 |
| Il7r | 0.001 | 0.000 | 0.001 |
| Rgs22 | 0.001 | 0.000 | 0.001 |
| Jchain | 0.001 | 0.001 | 0.001 |
| Sct | 0.001 | 0.001 | 0.031 |
| Plscr5 | 0.001 | 0.000 | 0.000 |
| Gm11992 | 0.001 | 0.000 | 0.000 |
| Olfr328 | 0.001 | 0.002 | 0.002 |
| Gm36287 | 0.001 | 0.002 | 0.001 |
| Mucl2 | 0.001 | 0.002 | 0.001 |
| Actl9 | 0.001 | 0.002 | 0.001 |
| Ctcflos | 0.001 | 0.002 | 0.001 |
| Impg1 | 0.001 | 0.004 | 0.000 |
| Sema4f | 0.001 | 0.002 | 0.002 |
| Gm5767 | 0.001 | 0.001 | 0.000 |
| Muc3 | 0.001 | 0.000 | 0.000 |
| Ccdc83 | 0.001 | 0.000 | 0.001 |
| Gm43598 | 0.001 | 0.010 | 0.008 |
| Chrne | 0.001 | 0.004 | 0.003 |
| 4930448D08I | 0.001 | 0.000 | 0.000 |
| Tmem74b | 0.001 | 0.001 | 0.004 |
| Epgn | 0.001 | 0.012 | 0.001 |
| C78197 | 0.001 | 0.001 | 0.002 |
| Asb17os | 0.001 | 0.001 | 0.000 |
| Gpr63 | 0.001 | 0.002 | 0.002 |
| Gm5868 | 0.001 | 0.000 | 0.000 |
| Tmem229a | 0.001 | 0.002 | 0.004 |
| E030018B13F | 0.001 | 0.000 | 0.000 |
| Rp1l1 | 0.001 | 0.000 | 0.000 |
| Isl2 | 0.001 | 0.009 | 0.004 |
| Hand1 | 0.001 | 0.004 | 0.003 |
| Gm29591 | 0.001 | 0.001 | 0.000 |
| Rhox7b | 0.001 | 0.000 | 0.000 |

| | | | |
|---|---|---|---|
| Olfr718-ps1 | 0.001 | 0.002 | 0.002 |
| Edar | 0.001 | 0.000 | 0.000 |
| Gm26520 | 0.001 | 0.001 | 0.001 |
| Tmem74bos | 0.001 | 0.001 | 0.000 |
| Gm15945 | 0.001 | 0.000 | 0.000 |
| Pcdha5 | 0.001 | 0.000 | 0.000 |
| Ccdc113 | 0.001 | 0.003 | 0.002 |
| Gm20663 | 0.001 | 0.000 | 0.000 |
| Bcl2l10 | 0.001 | 0.001 | 0.000 |
| Gm45897 | 0.001 | 0.002 | 0.000 |
| Mslnl | 0.001 | 0.005 | 0.002 |
| Gm13523 | 0.001 | 0.004 | 0.001 |
| Gm34921 | 0.001 | 0.000 | 0.000 |
| Gm26593 | 0.001 | 0.000 | 0.000 |
| Phox2b | 0.001 | 0.001 | 0.000 |
| Gm30373 | 0.001 | 0.001 | 0.002 |
| Zbtb8b | 0.001 | 0.001 | 0.001 |
| B3gnt4 | 0.001 | 0.001 | 0.002 |
| Slc27a2 | 0.001 | 0.000 | 0.000 |
| Fgf3 | 0.001 | 0.000 | 0.000 |
| Gm10717 | 0.001 | 0.000 | 0.000 |
| AC112943.2 | 0.001 | 0.000 | 0.000 |
| Retn | 0.001 | 0.000 | 0.000 |
| Tbx4 | 0.001 | 0.000 | 0.000 |
| Gm32884 | 0.001 | 0.000 | 0.000 |
| Trdc | 0.001 | 0.002 | 0.001 |
| Gm40983 | 0.001 | 0.000 | 0.000 |
| Slc26a4 | 0.001 | 0.000 | 0.000 |
| Cd96 | 0.001 | 0.000 | 0.002 |
| Retnlg | 0.001 | 0.001 | 0.000 |
| Kremen2 | 0.001 | 0.000 | 0.001 |
| Gm10309 | 0.001 | 0.001 | 0.000 |
| Btbd11 | 0.001 | 0.015 | 0.014 |
| Adgrv1 | 0.001 | 0.003 | 0.010 |
| Wfdc17 | 0.001 | 0.008 | 0.016 |
| I730028E13Ri | 0.001 | 0.000 | 0.001 |
| Apcdd1 | 0.001 | 0.005 | 0.002 |
| Sall3 | 0.001 | 0.000 | 0.000 |
| Gm43409 | 0.001 | 0.000 | 0.000 |
| Gm39228 | 0.001 | 0.002 | 0.001 |
| Cyct | 0.001 | 0.001 | 0.001 |
| Prr19 | 0.001 | 0.000 | 0.001 |
| Olfr804 | 0.001 | 0.001 | 0.000 |
| AC124739.1 | 0.001 | 0.001 | 0.001 |
| 1700125G22I | 0.001 | 0.000 | 0.000 |
| Gm26625 | 0.001 | 0.003 | 0.004 |
| Nup62cl | 0.001 | 0.001 | 0.001 |
| Gm49092 | 0.001 | 0.000 | 0.000 |

| | | | |
|---|---|---|---|
| Tmem200a | 0.001 | 0.002 | 0.007 |
| Gm14285 | 0.001 | 0.004 | 0.001 |
| Ralyl | 0.001 | 0.001 | 0.003 |
| AI507597 | 0.001 | 0.001 | 0.001 |
| Fbxo2 | 0.001 | 0.033 | 0.009 |
| 4930500F10F | 0.001 | 0.000 | 0.000 |
| Ccdc42os | 0.001 | 0.005 | 0.004 |
| Dhx58os | 0.001 | 0.001 | 0.003 |
| Gm49282 | 0.001 | 0.001 | 0.000 |
| AL583891.1 | 0.001 | 0.003 | 0.018 |
| Megf11 | 0.001 | 0.001 | 0.008 |
| Smyd1 | 0.001 | 0.002 | 0.002 |
| Hoxd4 | 0.001 | 0.000 | 0.000 |
| Gm20605 | 0.001 | 0.000 | 0.001 |
| Chrnb3 | 0.001 | 0.000 | 0.000 |
| AC138306.1 | 0.001 | 0.000 | 0.000 |
| Hpx | 0.001 | 0.003 | 0.002 |
| Nr5a1 | 0.001 | 0.000 | 0.000 |
| 4732490B19F | 0.001 | 0.000 | 0.000 |
| Hist1h2bk | 0.001 | 0.001 | 0.001 |
| Gm13262 | 0.001 | 0.000 | 0.000 |
| Sult1e1 | 0.001 | 0.000 | 0.002 |
| 1700063D05I | 0.001 | 0.000 | 0.000 |
| Gm996 | 0.001 | 0.004 | 0.003 |
| Gm2464 | 0.001 | 0.001 | 0.001 |
| Gm20457 | 0.001 | 0.001 | 0.001 |
| F7 | 0.001 | 0.000 | 0.000 |
| Gm11630 | 0.001 | 0.000 | 0.000 |
| Nxf2 | 0.001 | 0.001 | 0.003 |
| Dio3 | 0.001 | 0.001 | 0.007 |
| Grin2c | 0.001 | 0.000 | 0.000 |
| Gm15747 | 0.000 | 0.002 | 0.002 |
| Sgcz | 0.000 | 0.000 | 0.001 |
| Gm29771 | 0.000 | 0.000 | 0.000 |
| Abcb4 | 0.000 | 0.001 | 0.001 |
| Bco2 | 0.000 | 0.000 | 0.001 |
| Gm11373 | 0.000 | 0.000 | 0.000 |
| Prss56 | 0.000 | 0.000 | 0.001 |
| 5031425F14F | 0.000 | 0.000 | 0.001 |
| Cd79a | 0.000 | 0.002 | 0.001 |
| Kcnj5 | 0.000 | 0.000 | 0.000 |
| Asgr2 | 0.000 | 0.000 | 0.000 |
| Krt16 | 0.000 | 0.001 | 0.000 |
| Gm30198 | 0.000 | 0.001 | 0.001 |
| 4933427D06I | 0.000 | 0.000 | 0.000 |
| Gm44773 | 0.000 | 0.001 | 0.001 |
| Gm45571 | 0.000 | 0.000 | 0.000 |
| Gm48492 | 0.000 | 0.000 | 0.000 |

| | | | |
|---|---|---|---|
| 2610204G07I | 0.000 | 0.008 | 0.002 |
| F530104D19F | 0.000 | 0.000 | 0.000 |
| Gm34866 | 0.000 | 0.000 | 0.001 |
| Sp6 | 0.000 | 0.000 | 0.001 |
| Gm19866 | 0.000 | 0.000 | 0.000 |
| Tex22 | 0.000 | 0.000 | 0.000 |
| Elovl3 | 0.000 | 0.000 | 0.000 |
| Bcan | 0.000 | 0.000 | 0.000 |
| Cxcr2 | 0.000 | 0.002 | 0.002 |
| Crtac1 | 0.000 | 0.001 | 0.002 |
| Cntd1 | 0.000 | 0.002 | 0.003 |
| Kcnj10 | 0.000 | 0.001 | 0.001 |
| Caly | 0.000 | 0.001 | 0.004 |
| Kmo | 0.000 | 0.000 | 0.000 |
| Phf24 | 0.000 | 0.004 | 0.016 |
| 5430435K18F | 0.000 | 0.000 | 0.000 |
| Klf14 | 0.000 | 0.002 | 0.022 |
| Grm1 | 0.000 | 0.003 | 0.002 |
| Nek11 | 0.000 | 0.000 | 0.000 |
| 9630002D21I | 0.000 | 0.001 | 0.008 |
| Spry3 | 0.000 | 0.001 | 0.004 |
| Csf3 | 0.000 | 0.001 | 0.001 |
| Pla2r1 | 0.000 | 0.012 | 0.033 |
| Gm1965 | 0.000 | 0.000 | 0.000 |
| 6430571L13F | 0.000 | 0.001 | 0.002 |
| Gm48584 | 0.000 | 0.000 | 0.000 |
| 1700034P13F | 0.000 | 0.001 | 0.001 |
| Lcn12 | 0.000 | 0.000 | 0.000 |
| Gm42984 | 0.000 | 0.002 | 0.002 |
| Fam19a2 | 0.000 | 0.005 | 0.002 |
| Srp54b | 0.000 | 0.002 | 0.002 |
| Gm5475 | 0.000 | 0.001 | 0.004 |
| Gpr82 | 0.000 | 0.000 | 0.000 |
| E130304I02Ri | 0.000 | 0.000 | 0.000 |
| 5430419D17I | 0.000 | 0.000 | 0.000 |
| 1700104B16F | 0.000 | 0.000 | 0.000 |
| 4930572G02I | 0.000 | 0.001 | 0.001 |
| Lctl | 0.000 | 0.000 | 0.000 |
| Gm48707 | 0.000 | 0.000 | 0.000 |
| A330048O09I | 0.000 | 0.013 | 0.002 |
| St6gal2 | 0.000 | 0.004 | 0.000 |
| Gm14025 | 0.000 | 0.000 | 0.000 |
| Hcar2 | 0.000 | 0.000 | 0.000 |
| Gm47416 | 0.000 | 0.000 | 0.000 |
| Prss41 | 0.000 | 0.000 | 0.000 |
| Cpn1 | 0.000 | 0.000 | 0.000 |
| Tmem236 | 0.000 | 0.001 | 0.001 |
| Pdzk1 | 0.000 | 0.000 | 0.000 |

| | | | |
|---|---|---|---|
| Kcna3 | 0.000 | 0.000 | 0.000 |
| Gm44117 | 0.000 | 0.001 | 0.000 |
| Adamts20 | 0.000 | 0.004 | 0.014 |
| Gm2885 | 0.000 | 0.004 | 0.002 |
| Katnal2 | 0.000 | 0.001 | 0.002 |
| Kcnip2 | 0.000 | 0.002 | 0.001 |
| Gm20631 | 0.000 | 0.000 | 0.000 |
| 3110079O15I | 0.000 | 0.006 | 0.003 |
| Pou3f4 | 0.000 | 0.002 | 0.001 |
| Slc10a5 | 0.000 | 0.000 | 0.000 |
| Sptbn4 | 0.000 | 0.004 | 0.002 |
| Gm14435 | 0.000 | 0.000 | 0.000 |
| Sox3 | 0.000 | 0.001 | 0.001 |
| Gm40117 | 0.000 | 0.000 | 0.001 |
| Gm38944 | 0.000 | 0.001 | 0.000 |
| Ptprt | 0.000 | 0.000 | 0.001 |
| Gm31881 | 0.000 | 0.002 | 0.001 |
| Gm29681 | 0.000 | 0.001 | 0.001 |
| Kcnmb4os1 | 0.000 | 0.000 | 0.000 |
| Ttc23l | 0.000 | 0.001 | 0.002 |
| BC051537 | 0.000 | 0.001 | 0.001 |
| Ihh | 0.000 | 0.005 | 0.001 |
| Olfr351 | 0.000 | 0.000 | 0.000 |
| Gm15350 | 0.000 | 0.000 | 0.000 |
| Greb1 | 0.000 | 0.001 | 0.001 |
| Hhipl2 | 0.000 | 0.001 | 0.000 |
| Lrrtm4 | 0.000 | 0.000 | 0.004 |
| Gm36595 | 0.000 | 0.000 | 0.000 |
| Gm49016 | 0.000 | 0.000 | 0.000 |
| Vmn1r202 | 0.000 | 0.000 | 0.000 |
| AC126055.1 | 0.000 | 0.000 | 0.003 |
| Gm5426 | 0.000 | 0.001 | 0.001 |
| 4930412O13I | 0.000 | 0.000 | 0.000 |
| Gm14230 | 0.000 | 0.000 | 0.001 |
| 4930522O17I | 0.000 | 0.000 | 0.000 |
| Ptprh | 0.000 | 0.000 | 0.000 |
| Gm26710 | 0.000 | 0.004 | 0.003 |
| Gm3235 | 0.000 | 0.000 | 0.000 |
| Gm48940 | 0.000 | 0.000 | 0.000 |
| Gm13283 | 0.000 | 0.001 | 0.001 |
| Gm26911 | 0.000 | 0.001 | 0.000 |
| Tgm1 | 0.000 | 0.002 | 0.002 |
| Capsl | 0.000 | 0.000 | 0.000 |
| Ogdhl | 0.000 | 0.001 | 0.002 |
| Ankmy1 | 0.000 | 0.002 | 0.002 |
| 6330409D20I | 0.000 | 0.001 | 0.000 |
| Plcb2 | 0.000 | 0.001 | 0.001 |
| Zfp345 | 0.000 | 0.000 | 0.001 |

| | | | |
|---|---|---|---|
| Msh4 | 0.000 | 0.001 | 0.001 |
| Gm43303 | 0.000 | 0.000 | 0.000 |
| Gm26725 | 0.000 | 0.000 | 0.000 |
| 2810425M01 | 0.000 | 0.000 | 0.000 |
| Lcn11 | 0.000 | 0.000 | 0.000 |
| Gm26711 | 0.000 | 0.001 | 0.001 |
| Slco1b2 | 0.000 | 0.002 | 0.001 |
| Gipc3 | 0.000 | 0.009 | 0.001 |
| Foxf2 | 0.000 | 0.001 | 0.002 |
| Myrfl | 0.000 | 0.000 | 0.000 |
| Golt1a | 0.000 | 0.000 | 0.000 |
| Gm26906 | 0.000 | 0.003 | 0.007 |
| AC140304.2 | 0.000 | 0.003 | 0.001 |
| Als2cr12 | 0.000 | 0.000 | 0.000 |
| Gm43136 | 0.000 | 0.003 | 0.003 |
| AY358078 | 0.000 | 0.000 | 0.000 |
| Gm12052 | 0.000 | 0.001 | 0.004 |
| Nsg2 | 0.000 | 0.001 | 0.002 |
| Pde6g | 0.000 | 0.001 | 0.001 |
| Gm17115 | 0.000 | 0.001 | 0.001 |
| Il1rapl1 | 0.000 | 0.001 | 0.017 |
| Gm49396 | 0.000 | 0.001 | 0.001 |
| Fitm1 | 0.000 | 0.016 | 0.006 |
| H2-T3 | 0.000 | 0.000 | 0.000 |
| Spesp1 | 0.000 | 0.000 | 0.000 |
| Atoh7 | 0.000 | 0.001 | 0.001 |
| Gm26876 | 0.000 | 0.002 | 0.001 |
| Ptpn20 | 0.000 | 0.000 | 0.001 |
| 1810028F09F | 0.000 | 0.000 | 0.000 |
| C530050E15F | 0.000 | 0.002 | 0.003 |
| Gm29459 | 0.000 | 0.000 | 0.000 |
| Kng2 | 0.000 | 0.000 | 0.001 |
| Musk | 0.000 | 0.001 | 0.003 |
| Gm12763 | 0.000 | 0.000 | 0.000 |
| Gm48202 | 0.000 | 0.000 | 0.000 |
| Rxfp3 | 0.000 | 0.000 | 0.001 |
| Gm37768 | 0.000 | 0.004 | 0.002 |
| Gpr158 | 0.000 | 0.000 | 0.000 |
| Gm19710 | 0.000 | 0.001 | 0.001 |
| Vstm2b | 0.000 | 0.062 | 0.069 |
| Olfr648 | 0.000 | 0.000 | 0.000 |
| AC109617.2 | 0.000 | 0.000 | 0.010 |
| Gm16618 | 0.000 | 0.000 | 0.000 |
| Hmx2 | 0.000 | 0.000 | 0.000 |
| Car6 | 0.000 | 0.000 | 0.000 |
| Slc6a21 | 0.000 | 0.000 | 0.000 |
| Gm27227 | 0.000 | 0.000 | 0.000 |
| Glp2r | 0.000 | 0.000 | 0.000 |

| | | | |
|---|---|---|---|
| Sorcs3 | 0.000 | 0.004 | 0.012 |
| Cel | 0.000 | 0.000 | 0.000 |
| Plppr5 | 0.000 | 0.001 | 0.008 |
| Tbxas1 | 0.000 | 0.013 | 0.003 |
| Rsph6a | 0.000 | 0.000 | 0.001 |
| Rasgrp4 | 0.000 | 0.001 | 0.000 |
| Gm2044 | 0.000 | 0.000 | 0.000 |
| 2310039L15F | 0.000 | 0.001 | 0.000 |
| Ppp1r27 | 0.000 | 0.000 | 0.000 |
| Tigit | 0.000 | 0.000 | 0.000 |
| Tmem30c | 0.000 | 0.002 | 0.003 |
| Chgb | 0.000 | 0.001 | 0.003 |
| Ano4 | 0.000 | 0.000 | 0.004 |
| Cntnap4 | 0.000 | 0.003 | 0.001 |
| Gm42743 | 0.000 | 0.001 | 0.001 |
| Hpdl | 0.000 | 0.002 | 0.001 |
| Clec2h | 0.000 | 0.002 | 0.000 |
| Gm12764 | 0.000 | 0.000 | 0.000 |
| Gm11497 | 0.000 | 0.000 | 0.000 |
| Gm31946 | 0.000 | 0.001 | 0.001 |
| Gm20683 | 0.000 | 0.000 | 0.000 |
| Olfr1178 | 0.000 | 0.000 | 0.000 |
| Cdh15 | 0.000 | 0.001 | 0.002 |
| Gm10646 | 0.000 | 0.000 | 0.001 |
| Mycbpap | 0.000 | 0.001 | 0.002 |
| Zpbp2 | 0.000 | 0.000 | 0.000 |
| Platr17 | 0.000 | 0.000 | 0.001 |
| Olfr1369-ps1 | 0.000 | 0.003 | 0.006 |
| Gm5415 | 0.000 | 0.000 | 0.000 |
| Klre1 | 0.000 | 0.000 | 0.000 |
| Lin7b | 0.000 | 0.014 | 0.009 |
| Elavl3 | 0.000 | 0.001 | 0.002 |
| Gm15860 | 0.000 | 0.009 | 0.007 |
| M1ap | 0.000 | 0.001 | 0.003 |
| Gm44892 | 0.000 | 0.000 | 0.000 |
| Gm39377 | 0.000 | 0.000 | 0.000 |
| Gm10635 | 0.000 | 0.000 | 0.000 |
| Sox15 | 0.000 | 0.000 | 0.000 |
| Apoh | 0.000 | 0.001 | 0.001 |
| Adcy8 | 0.000 | 0.007 | 0.009 |
| Asb11 | 0.000 | 0.002 | 0.001 |
| Ptprq | 0.000 | 0.000 | 0.000 |
| Vmn1r218 | 0.000 | 0.000 | 0.001 |
| Gm17746 | 0.000 | 0.000 | 0.000 |
| Gm16596 | 0.000 | 0.000 | 0.000 |
| Gm16344 | 0.000 | 0.000 | 0.000 |
| Otc | 0.000 | 0.000 | 0.000 |
| Tmprss11d | 0.000 | 0.000 | 0.000 |

| | | | |
|---|---|---|---|
| Folh1 | 0.000 | 0.002 | 0.022 |
| Rdh16 | 0.000 | 0.000 | 0.000 |
| Gm15743 | 0.000 | 0.001 | 0.000 |
| Cabyr | 0.000 | 0.001 | 0.000 |
| 4930431L21F | 0.000 | 0.000 | 0.000 |
| Alb | 0.000 | 0.000 | 0.000 |
| Gpr183 | 0.000 | 0.002 | 0.001 |
| Gm15983 | 0.000 | 0.001 | 0.001 |
| Gm21149 | 0.000 | 0.000 | 0.000 |
| Gm26690 | 0.000 | 0.009 | 0.010 |
| Gm2245 | 0.000 | 0.000 | 0.000 |
| Adam1a | 0.000 | 0.001 | 0.000 |
| D030051J21F | 0.000 | 0.003 | 0.003 |
| Hgs.1 | 0.000 | 0.001 | 0.000 |
| Shc3 | 0.000 | 0.001 | 0.001 |
| Gm5678 | 0.000 | 0.000 | 0.000 |
| Alkal1 | 0.000 | 0.003 | 0.001 |
| Gm43077 | 0.000 | 0.000 | 0.000 |
| Gm45092 | 0.000 | 0.013 | 0.003 |
| 4921522P10F | 0.000 | 0.000 | 0.000 |
| Gm45542 | 0.000 | 0.000 | 0.000 |
| Kcnk5 | 0.000 | 0.002 | 0.007 |
| Hist1h2ai | 0.000 | 0.000 | 0.000 |
| C230034O21I | 0.000 | 0.001 | 0.001 |
| Gm20594 | 0.000 | 0.000 | 0.000 |
| Kcnc3 | 0.000 | 0.002 | 0.001 |
| Cbln3 | 0.000 | 0.002 | 0.004 |
| Gm15907 | 0.000 | 0.000 | 0.000 |
| C230088H06I | 0.000 | 0.000 | 0.000 |
| 4931440J10R | 0.000 | 0.001 | 0.001 |
| Rab39 | 0.000 | 0.000 | 0.000 |
| Slc5a10 | 0.000 | 0.000 | 0.000 |
| Gm49392 | 0.000 | 0.000 | 0.000 |
| Rgs8 | 0.000 | 0.001 | 0.001 |
| 3110099E03F | 0.000 | 0.000 | 0.003 |
| Tgm3 | 0.000 | 0.001 | 0.000 |
| Mir124-2hg | 0.000 | 0.000 | 0.000 |
| Dab1 | 0.000 | 0.001 | 0.002 |
| Gm42596 | 0.000 | 0.000 | 0.000 |
| Ppp1r3a | 0.000 | 0.001 | 0.001 |
| 6430584L05F | 0.000 | 0.000 | 0.000 |
| Gm20686 | 0.000 | 0.003 | 0.001 |
| Gm38150 | 0.000 | 0.001 | 0.001 |
| Gm12022 | 0.000 | 0.003 | 0.066 |
| Kdm6bos | 0.000 | 0.015 | 0.015 |
| Gm48829 | 0.000 | 0.000 | 0.000 |
| Rspo2 | 0.000 | 0.002 | 0.023 |
| Ppp3r2 | 0.000 | 0.000 | 0.001 |

| | | | |
|---|---|---|---|
| Gm12916 | 0.000 | 0.000 | 0.000 |
| Mmd2 | 0.000 | 0.001 | 0.002 |
| Clhc1 | 0.000 | 0.001 | 0.002 |
| Dmrtc1a | 0.000 | 0.001 | 0.003 |
| Gm28087 | 0.000 | 0.001 | 0.000 |
| Gm39460 | 0.000 | 0.000 | 0.000 |
| Dpcr1 | 0.000 | 0.001 | 0.002 |
| Gm14124 | 0.000 | 0.000 | 0.001 |
| Actl7b | 0.000 | 0.000 | 0.002 |
| Cpvl | 0.000 | 0.000 | 0.000 |
| Wdr93 | 0.000 | 0.001 | 0.002 |
| Tph2 | 0.000 | 0.000 | 0.001 |
| Gm30246 | 0.000 | 0.001 | 0.000 |
| 1810064F22F | 0.000 | 0.000 | 0.000 |
| Gm28379 | 0.000 | 0.001 | 0.000 |
| Gm20537 | 0.000 | 0.000 | 0.000 |
| Gm26555 | 0.000 | 0.001 | 0.000 |
| Adgrf2 | 0.000 | 0.000 | 0.000 |
| Gm15337 | 0.000 | 0.001 | 0.002 |
| Pifo | 0.000 | 0.000 | 0.000 |
| Slc6a16 | 0.000 | 0.000 | 0.000 |
| Gm48607 | 0.000 | 0.000 | 0.001 |
| Neu2 | 0.000 | 0.001 | 0.001 |
| Gm26831 | 0.000 | 0.001 | 0.002 |
| Mroh8 | 0.000 | 0.001 | 0.000 |
| Tomt | 0.000 | 0.000 | 0.001 |
| Hcn1 | 0.000 | 0.001 | 0.011 |
| Nphs2 | 0.000 | 0.000 | 0.000 |
| Crp | 0.000 | 0.000 | 0.000 |
| Nyx | 0.000 | 0.001 | 0.001 |
| Gm37855 | 0.000 | 0.001 | 0.001 |
| 4933438K21F | 0.000 | 0.000 | 0.000 |
| Nrsn1 | 0.000 | 0.001 | 0.001 |
| Gm6938 | 0.000 | 0.001 | 0.003 |
| Gm20422 | 0.000 | 0.000 | 0.000 |
| Pebp4 | 0.000 | 0.000 | 0.000 |
| Gm37988 | 0.000 | 0.000 | 0.000 |
| Grin1os | 0.000 | 0.000 | 0.000 |
| Gm33023 | 0.000 | 0.009 | 0.004 |
| Izumo1r | 0.000 | 0.000 | 0.000 |
| Rab44 | 0.000 | 0.000 | 0.000 |
| Gm13571 | 0.000 | 0.001 | 0.000 |
| Gm12473 | 0.000 | 0.001 | 0.000 |
| Plpp4 | 0.000 | 0.008 | 0.003 |
| 9230102K24F | 0.000 | 0.000 | 0.000 |
| Gm26614 | 0.000 | 0.003 | 0.000 |
| Gjd3 | 0.000 | 0.000 | 0.000 |
| Frmpd1os | 0.000 | 0.000 | 0.000 |

| | | | |
|---|---|---|---|
| Trim80 | 0.000 | 0.000 | 0.000 |
| Gm5483 | 0.000 | 0.002 | 0.001 |
| Maats1 | 0.000 | 0.000 | 0.001 |
| Cnih3 | 0.000 | 0.001 | 0.002 |
| Gm35164 | 0.000 | 0.000 | 0.000 |
| Cacng1 | 0.000 | 0.000 | 0.001 |
| Gm35818 | 0.000 | 0.000 | 0.000 |
| Gm15738 | 0.000 | 0.000 | 0.000 |
| Epcam | 0.000 | 0.001 | 0.001 |
| Cd28 | 0.000 | 0.001 | 0.007 |
| Gm39662 | 0.000 | 0.000 | 0.000 |
| 9830004L10F | 0.000 | 0.000 | 0.000 |
| Plppr4 | 0.000 | 0.002 | 0.003 |
| Ermap | 0.000 | 0.001 | 0.002 |
| Gm20629 | 0.000 | 0.003 | 0.000 |
| Prrt3 | 0.000 | 0.002 | 0.001 |
| Ano2 | 0.000 | 0.003 | 0.002 |
| 1700034J05R | 0.000 | 0.000 | 0.000 |
| Timm8a2 | 0.000 | 0.001 | 0.001 |
| Gm49395 | 0.000 | 0.000 | 0.000 |
| Mreg | 0.000 | 0.005 | 0.002 |
| Gm21411 | 0.000 | 0.000 | 0.000 |
| 1300017J02R | 0.000 | 0.001 | 0.001 |
| Hbq1b | 0.000 | 0.000 | 0.000 |
| Tmem102 | 0.000 | 0.001 | 0.001 |
| Rhag | 0.000 | 0.000 | 0.000 |
| Sell | 0.000 | 0.000 | 0.000 |
| Gm21962 | 0.000 | 0.000 | 0.001 |
| Gm43948 | 0.000 | 0.003 | 0.000 |
| Grik4 | 0.000 | 0.003 | 0.006 |
| Gm10425 | 0.000 | 0.000 | 0.001 |
| H2-T24 | 0.000 | 0.000 | 0.001 |
| Gm29040 | 0.000 | 0.000 | 0.000 |
| D330045A20I | 0.000 | 0.005 | 0.006 |
| Gm43134 | 0.000 | 0.000 | 0.000 |
| 6330403N20I | 0.000 | 0.004 | 0.005 |
| 1700006J14R | 0.000 | 0.000 | 0.000 |
| Ighm | 0.000 | 0.002 | 0.002 |
| Sytl3 | 0.000 | 0.000 | 0.001 |
| Gm38097 | 0.000 | 0.000 | 0.001 |
| Gm26883 | 0.000 | 0.006 | 0.010 |
| Gm5129 | 0.000 | 0.000 | 0.000 |
| Tbx3os1 | 0.000 | 0.003 | 0.004 |
| Gm10655 | 0.000 | 0.002 | 0.001 |
| Slc36a3os | 0.000 | 0.003 | 0.001 |
| G930009F23F | 0.000 | 0.000 | 0.000 |
| Glyat | 0.000 | 0.000 | 0.002 |
| Epb42 | 0.000 | 0.000 | 0.001 |

| | | | |
|---|---|---|---|
| Sap25 | 0.000 | 0.000 | 0.000 |
| 1700001P01I | 0.000 | 0.000 | 0.000 |
| Gm47467 | 0.000 | 0.002 | 0.002 |
| Dtnbos | 0.000 | 0.001 | 0.002 |
| AC126280.1 | 0.000 | 0.001 | 0.002 |
| Ccdc160 | 0.000 | 0.004 | 0.001 |
| Lrrc31 | 0.000 | 0.000 | 0.000 |
| Gm45081 | 0.000 | 0.000 | 0.001 |
| Gm10190 | 0.000 | 0.000 | 0.000 |
| Akr1cl | 0.000 | 0.001 | 0.007 |
| Gm2453 | 0.000 | 0.001 | 0.000 |
| Peak1os | 0.000 | 0.000 | 0.000 |
| AC133488.1 | 0.000 | 0.001 | 0.001 |
| Il18rap | 0.000 | 0.000 | 0.001 |
| Ifi206 | 0.000 | 0.000 | 0.000 |
| Card9 | 0.000 | 0.000 | 0.000 |
| Myo3b | 0.000 | 0.000 | 0.000 |
| Slc9a7 | 0.000 | 0.001 | 0.000 |
| F630028O10I | 0.000 | 0.000 | 0.000 |
| Tlr13 | 0.000 | 0.000 | 0.001 |
| Gm9733 | 0.000 | 0.000 | 0.000 |
| Gm5150 | 0.000 | 0.000 | 0.000 |
| Skint3 | 0.000 | 0.000 | 0.000 |
| Csf3r | 0.000 | 0.001 | 0.001 |
| Gm5106 | 0.000 | 0.000 | 0.000 |
| Gm16251 | 0.000 | 0.000 | 0.000 |
| Siglecg | 0.000 | 0.000 | 0.000 |
| Il4i1 | 0.000 | 0.001 | 0.002 |
| Il21r | 0.000 | 0.000 | 0.001 |
| Ifitm6 | 0.000 | 0.000 | 0.001 |
| Gm26740 | 0.000 | 0.000 | 0.001 |
| Gm27021 | 0.000 | 0.000 | 0.000 |
| S1pr5 | 0.000 | 0.000 | 0.000 |
| Bcl2a1d | 0.000 | 0.002 | 0.002 |
| Adora2b | 0.000 | 0.009 | 0.014 |
| Mir142hg | 0.000 | 0.000 | 0.000 |
| Ly86 | 0.000 | 0.004 | 0.005 |
| Mefv | 0.000 | 0.000 | 0.000 |
| Cd86 | 0.000 | 0.002 | 0.001 |
| 4932438H23I | 0.000 | 0.003 | 0.004 |
| Gm15964 | 0.000 | 0.000 | 0.000 |
| Trem1 | 0.000 | 0.000 | 0.000 |
| Treml4 | 0.000 | 0.000 | 0.000 |
| Adgre4 | 0.000 | 0.000 | 0.000 |
| 1700011I03R | 0.000 | 0.002 | 0.000 |
| Pdcd1lg2 | 0.000 | 0.001 | 0.000 |
| Prkag3 | 0.000 | 0.003 | 0.002 |
| Gm14342 | 0.000 | 0.000 | 0.000 |

| | | | |
|---|---|---|---|
| Trim30c | 0.000 | 0.000 | 0.000 |
| Notum | 0.000 | 0.003 | 0.001 |
| Gm33682 | 0.000 | 0.000 | 0.000 |
| Gm648 | 0.000 | 0.001 | 0.000 |
| Tyrp1 | 0.000 | 0.002 | 0.001 |
| 0610040J01R | 0.000 | 0.001 | 0.001 |
| Slc5a5 | 0.000 | 0.001 | 0.002 |
| Spaca3 | 0.000 | 0.000 | 0.000 |
| Tbr1 | 0.000 | 0.001 | 0.002 |
| Gm14235 | 0.000 | 0.000 | 0.000 |
| Gm13288 | 0.000 | 0.000 | 0.000 |
| Nr2e3 | 0.000 | 0.007 | 0.001 |
| Gm47320 | 0.000 | 0.002 | 0.000 |
| 2410004I01R | 0.000 | 0.002 | 0.009 |
| Lypd6b | 0.000 | 0.000 | 0.000 |
| Myb | 0.000 | 0.000 | 0.001 |
| Amn | 0.000 | 0.001 | 0.001 |
| 4930558K02F | 0.000 | 0.001 | 0.001 |
| Gimap3 | 0.000 | 0.000 | 0.000 |
| Ggt1 | 0.000 | 0.002 | 0.001 |
| Slc38a8 | 0.000 | 0.001 | 0.001 |
| Gm31508 | 0.000 | 0.000 | 0.000 |
| Aifm3 | 0.000 | 0.001 | 0.001 |
| 1700052H01I | 0.000 | 0.000 | 0.000 |
| D030045P18I | 0.000 | 0.000 | 0.002 |
| Terb1 | 0.000 | 0.001 | 0.000 |
| Abcd2 | 0.000 | 0.004 | 0.031 |
| AI847159 | 0.000 | 0.000 | 0.000 |
| Grpr | 0.000 | 0.003 | 0.001 |
| Gm16685 | 0.000 | 0.000 | 0.001 |
| Gm16282 | 0.000 | 0.000 | 0.000 |
| Gm48342 | 0.000 | 0.001 | 0.002 |
| 1700064H15I | 0.000 | 0.001 | 0.001 |
| Atp6v0d2 | 0.000 | 0.001 | 0.001 |
| Ccdc81 | 0.000 | 0.001 | 0.002 |
| Gm26705 | 0.000 | 0.003 | 0.001 |
| Gm26765 | 0.000 | 0.001 | 0.000 |
| Gm47231 | 0.000 | 0.001 | 0.001 |
| Gm36278 | 0.000 | 0.000 | 0.000 |
| Gm10775 | 0.000 | 0.001 | 0.001 |
| 4930557J02R | 0.000 | 0.000 | 0.000 |
| Dnah10 | 0.000 | 0.001 | 0.000 |
| Hoxb1 | 0.000 | 0.002 | 0.002 |
| D130043K22I | 0.000 | 0.001 | 0.001 |
| Gm26723 | 0.000 | 0.001 | 0.000 |
| Gm4737 | 0.000 | 0.002 | 0.003 |
| Gm10300 | 0.000 | 0.000 | 0.000 |
| Irs3 | 0.000 | 0.001 | 0.001 |

| | | | |
|---|---|---|---|
| 2310002D06I | 0.000 | 0.001 | 0.001 |
| 3300005D01I | 0.000 | 0.001 | 0.002 |
| Cd274 | 0.000 | 0.002 | 0.001 |
| Gm35986 | 0.000 | 0.000 | 0.000 |
| Clec9a | 0.000 | 0.000 | 0.000 |
| Lin28b | 0.000 | 0.000 | 0.000 |
| Gm36198 | 0.000 | 0.001 | 0.000 |
| Gm13496 | 0.000 | 0.000 | 0.000 |
| Tdo2 | 0.000 | 0.001 | 0.003 |
| Spata21 | 0.000 | 0.000 | 0.000 |
| 1700003E16F | 0.000 | 0.001 | 0.001 |
| Cd69 | 0.000 | 0.001 | 0.000 |
| Gm17535 | 0.000 | 0.000 | 0.000 |
| Gm35371 | 0.000 | 0.001 | 0.001 |
| Krt88 | 0.000 | 0.000 | 0.000 |
| Olfr167 | 0.000 | 0.000 | 0.000 |
| Prox1os | 0.000 | 0.000 | 0.000 |
| Rag1 | 0.000 | 0.000 | 0.000 |
| Gnb3 | 0.000 | 0.004 | 0.004 |
| Mybpc2 | 0.000 | 0.005 | 0.002 |
| Gm21119 | 0.000 | 0.000 | 0.000 |
| Gm35256 | 0.000 | 0.000 | 0.000 |
| Nupl1 | 0.000 | 0.001 | 0.000 |
| Gm27028 | 0.000 | 0.000 | 0.000 |
| Gm35521 | 0.000 | 0.000 | 0.000 |
| Pde4c | 0.000 | 0.002 | 0.005 |
| 5330417C22F | 0.000 | 0.000 | 0.000 |
| Gm27188 | 0.000 | 0.000 | 0.000 |
| 9230116L04F | 0.000 | 0.001 | 0.001 |
| Angpt4 | 0.000 | 0.004 | 0.042 |
| Slc6a13 | 0.000 | 0.005 | 0.015 |
| Onecut3 | 0.000 | 0.000 | 0.000 |
| Slitrk1 | 0.000 | 0.001 | 0.013 |
| Lhx1 | 0.000 | 0.000 | 0.000 |
| Gm20390 | 0.000 | 0.000 | 0.000 |
| Krt2 | 0.000 | 0.000 | 0.000 |
| Gm37323 | 0.000 | 0.000 | 0.000 |
| Tcf24 | 0.000 | 0.001 | 0.002 |
| Gm26781 | 0.000 | 0.000 | 0.000 |
| Gm4788 | 0.000 | 0.000 | 0.000 |
| A430078G23I | 0.000 | 0.000 | 0.000 |
| 1700006E09F | 0.000 | 0.000 | 0.000 |
| Hist1h2bn | 0.000 | 0.002 | 0.000 |
| Gm34667 | 0.000 | 0.000 | 0.000 |
| D330050G23I | 0.000 | 0.001 | 0.001 |
| Gm37876 | 0.000 | 0.001 | 0.001 |
| Ampd1 | 0.000 | 0.003 | 0.005 |
| Gm12953 | 0.000 | 0.001 | 0.000 |

| | | | |
|---|---|---|---|
| 9430041J12R | 0.000 | 0.001 | 0.001 |
| Tspoap1 | 0.000 | 0.001 | 0.000 |
| AC125526.1 | 0.000 | 0.000 | 0.000 |
| Mettl7a2 | 0.000 | 0.000 | 0.001 |
| Noxa1 | 0.000 | 0.000 | 0.000 |
| Slc2a6 | 0.000 | 0.001 | 0.002 |
| Zfp804a | 0.000 | 0.001 | 0.002 |
| A730032A03R | 0.000 | 0.000 | 0.000 |
| Artn | 0.000 | 0.003 | 0.006 |
| Syce1 | 0.000 | 0.000 | 0.000 |
| Adprhl1 | 0.000 | 0.002 | 0.001 |
| Rtbdn | 0.000 | 0.000 | 0.001 |
| Stmn4 | 0.000 | 0.004 | 0.039 |
| Gm28959 | 0.000 | 0.001 | 0.001 |
| Vmo1 | 0.000 | 0.006 | 0.017 |
| Gm17473 | 0.000 | 0.000 | 0.000 |
| Grik1 | 0.000 | 0.000 | 0.009 |
| Gm10118 | 0.000 | 0.002 | 0.000 |
| Gck | 0.000 | 0.001 | 0.002 |
| Samd11 | 0.000 | 0.003 | 0.002 |
| Rasal1 | 0.000 | 0.000 | 0.000 |
| Npvf | 0.000 | 0.001 | 0.000 |
| Klk14 | 0.000 | 0.000 | 0.000 |
| Gm4353 | 0.000 | 0.000 | 0.000 |
| Gm10322 | 0.000 | 0.000 | 0.000 |
| D030068K23R | 0.000 | 0.004 | 0.001 |
| Esyt3 | 0.000 | 0.000 | 0.001 |
| Fbxo48 | 0.000 | 0.000 | 0.000 |
| D630024D03R | 0.000 | 0.000 | 0.000 |
| AI504432 | 0.000 | 0.002 | 0.003 |
| Ccdc180 | 0.000 | 0.000 | 0.000 |
| Gpr37 | 0.000 | 0.001 | 0.000 |
| Gm15401 | 0.000 | 0.000 | 0.001 |
| Etv2 | 0.000 | 0.000 | 0.000 |
| 4933408N05R | 0.000 | 0.000 | 0.001 |
| Slc39a2 | 0.000 | 0.001 | 0.002 |
| Fabp6 | 0.000 | 0.000 | 0.000 |
| Slc26a3 | 0.000 | 0.006 | 0.001 |
| Csta1 | 0.000 | 0.000 | 0.000 |
| Tmem169 | 0.000 | 0.003 | 0.005 |
| Tmprss11b | 0.000 | 0.000 | 0.000 |
| Gm15406 | 0.000 | 0.003 | 0.002 |
| Trim66 | 0.000 | 0.000 | 0.002 |
| Pcdh15 | 0.000 | 0.000 | 0.000 |
| Gm28722 | 0.000 | 0.000 | 0.004 |
| Gm34113 | 0.000 | 0.001 | 0.000 |
| Crhbp | 0.000 | 0.012 | 0.001 |
| Spdef | 0.000 | 0.002 | 0.002 |

| | | | |
|---|---|---|---|
| Pde11a | 0.000 | 0.001 | 0.002 |
| Gm43549 | 0.000 | 0.000 | 0.000 |
| Gm4349 | 0.000 | 0.001 | 0.002 |
| Sytl1 | 0.000 | 0.003 | 0.002 |
| 3100003L05F | 0.000 | 0.012 | 0.013 |
| Btbd16 | 0.000 | 0.000 | 0.000 |
| Slc47a1 | 0.000 | 0.000 | 0.001 |
| Car10 | 0.000 | 0.014 | 0.005 |
| Hist1h4b | 0.000 | 0.000 | 0.000 |
| Fbxo43 | 0.000 | 0.000 | 0.000 |
| CT025584.1 | 0.000 | 0.000 | 0.000 |
| Gm42949 | 0.000 | 0.000 | 0.000 |
| AU023070 | 0.000 | 0.000 | 0.000 |
| 5830468F06F | 0.000 | 0.000 | 0.000 |
| 1700020M21 | 0.000 | 0.000 | 0.000 |
| Tgm4 | 0.000 | 0.001 | 0.001 |
| Gm47695 | 0.000 | 0.000 | 0.000 |
| 4930474N09I | 0.000 | 0.001 | 0.002 |
| Gm26904 | 0.000 | 0.000 | 0.000 |
| Rax | 0.000 | 0.001 | 0.003 |
| Slamf7 | 0.000 | 0.000 | 0.001 |
| Gpat2 | 0.000 | 0.000 | 0.001 |
| Grk1 | 0.000 | 0.000 | 0.000 |
| Tmem266 | 0.000 | 0.000 | 0.001 |
| Gm48071 | 0.000 | 0.000 | 0.000 |
| Gm6970 | 0.000 | 0.000 | 0.000 |
| Gm9047 | 0.000 | 0.000 | 0.000 |
| AI854703 | 0.000 | 0.002 | 0.004 |
| C1s2 | 0.000 | 0.000 | 0.002 |
| Gm49086 | 0.000 | 0.001 | 0.001 |
| Panx3 | 0.000 | 0.000 | 0.000 |
| Gm47237 | 0.000 | 0.011 | 0.005 |
| Gm48116 | 0.000 | 0.000 | 0.000 |
| Papln | 0.000 | 0.000 | 0.004 |
| Syndig1l | 0.000 | 0.000 | 0.000 |
| Gm16381 | 0.000 | 0.000 | 0.000 |
| Gm26766 | 0.000 | 0.001 | 0.001 |
| Dppa2 | 0.000 | 0.002 | 0.002 |
| Gm28285 | 0.000 | 0.001 | 0.002 |
| Baat | 0.000 | 0.001 | 0.000 |
| Gm2366 | 0.000 | 0.001 | 0.001 |
| Cadps | 0.000 | 0.001 | 0.002 |
| Gm26805 | 0.000 | 0.000 | 0.001 |
| Bpifb2 | 0.000 | 0.000 | 0.000 |
| 1700060C20F | 0.000 | 0.001 | 0.001 |
| Arhgap40 | 0.000 | 0.000 | 0.000 |
| Cd209c | 0.000 | 0.001 | 0.000 |
| Gm31518 | 0.000 | 0.000 | 0.000 |

| | | | |
|---|---|---|---|
| AC114585.1 | 0.000 | 0.000 | 0.000 |
| Esrrb | 0.000 | 0.000 | 0.000 |
| Rtl10 | 0.000 | 0.001 | 0.001 |
| Gm15718 | 0.000 | 0.001 | 0.001 |
| Grin3b | 0.000 | 0.003 | 0.002 |
| Gm11681 | 0.000 | 0.000 | 0.000 |
| Fam50b | 0.000 | 0.000 | 0.000 |
| 5330416C01R | 0.000 | 0.000 | 0.001 |
| Gm49169 | 0.000 | 0.000 | 0.000 |
| Pla2g10 | 0.000 | 0.000 | 0.000 |
| Atp13a4 | 0.000 | 0.000 | 0.000 |
| Fbxo15 | 0.000 | 0.000 | 0.000 |
| Atp6v0a4 | 0.000 | 0.001 | 0.011 |
| Gm31479 | 0.000 | 0.000 | 0.000 |
| Gm44812 | 0.000 | 0.003 | 0.006 |
| Gm48654 | 0.000 | 0.000 | 0.000 |
| Gstt4 | 0.000 | 0.000 | 0.000 |
| Nr1h4 | 0.000 | 0.000 | 0.002 |
| Sorbs2os | 0.000 | 0.001 | 0.000 |
| Gm39464 | 0.000 | 0.000 | 0.000 |
| Gm11973 | 0.000 | 0.003 | 0.002 |
| Gm26829 | 0.000 | 0.000 | 0.002 |
| Gm10814 | 0.000 | 0.002 | 0.004 |
| Gm15872 | 0.000 | 0.000 | 0.000 |
| Olfr1220 | 0.000 | 0.000 | 0.000 |
| Gstm6 | 0.000 | 0.002 | 0.004 |
| Gm16001 | 0.000 | 0.001 | 0.001 |
| A830011K09R | 0.000 | 0.000 | 0.002 |
| Gm26714 | 0.000 | 0.000 | 0.000 |
| Gm41030 | 0.000 | 0.000 | 0.000 |
| Tnr | 0.000 | 0.001 | 0.000 |
| Gm15423 | 0.000 | 0.000 | 0.000 |
| Vstm2l | 0.000 | 0.000 | 0.001 |
| 2610028E06R | 0.000 | 0.000 | 0.002 |
| 4930458D05R | 0.000 | 0.026 | 0.012 |
| Arg1 | 0.000 | 0.001 | 0.002 |
| Gm45866 | 0.000 | 0.000 | 0.000 |
| Agt | 0.000 | 0.005 | 0.039 |
| Mhrt | 0.000 | 0.000 | 0.000 |
| D630033O11R | 0.000 | 0.000 | 0.020 |
| Sarm1 | 0.000 | 0.006 | 0.002 |
| Slc16a5 | 0.000 | 0.000 | 0.000 |
| Ube2ql1 | 0.000 | 0.005 | 0.007 |
| Kcnf1 | 0.000 | 0.001 | 0.000 |
| Sim1 | 0.000 | 0.000 | 0.000 |
| Prtn3 | 0.000 | 0.005 | 0.007 |
| Olfr921 | 0.000 | 0.002 | 0.005 |
| Mrap2 | 0.000 | 0.000 | 0.000 |

| | | | |
|---|---|---|---|
| D730003K21I | 0.000 | 0.000 | 0.000 |
| Olfr1383 | 0.000 | 0.000 | 0.000 |
| Gm11738 | 0.000 | 0.001 | 0.001 |
| Pou1f1 | 0.000 | 0.000 | 0.000 |
| Gm973 | 0.000 | 0.004 | 0.002 |
| Gm16701.1 | 0.000 | 0.005 | 0.002 |
| Olfr624 | 0.000 | 0.000 | 0.000 |
| Zg16 | 0.000 | 0.000 | 0.000 |
| Bfsp2 | 0.000 | 0.003 | 0.001 |
| Gm17200 | 0.000 | 0.000 | 0.000 |
| Gm48508 | 0.000 | 0.000 | 0.000 |
| 4930563E18F | 0.000 | 0.000 | 0.000 |
| Slc9a2 | 0.000 | 0.004 | 0.001 |
| Gm13999 | 0.000 | 0.000 | 0.000 |
| Pax5 | 0.000 | 0.000 | 0.000 |
| Trim54 | 0.000 | 0.000 | 0.000 |
| Pkp3 | 0.000 | 0.001 | 0.000 |
| Slc35d3 | 0.000 | 0.002 | 0.003 |
| Dhrs2 | 0.000 | 0.000 | 0.001 |
| Ccr3 | 0.000 | 0.000 | 0.001 |
| 4930565N06I | 0.000 | 0.001 | 0.000 |
| Gm5160 | 0.000 | 0.001 | 0.000 |
| Stkld1 | 0.000 | 0.001 | 0.001 |
| L3mbtl1 | 0.000 | 0.002 | 0.003 |
| Gm26652 | 0.000 | 0.002 | 0.002 |
| Gm12524 | 0.000 | 0.001 | 0.002 |
| Gm33466 | 0.000 | 0.000 | 0.000 |
| 0610043K17F | 0.000 | 0.003 | 0.005 |
| Mpl | 0.000 | 0.000 | 0.000 |
| Gm15396 | 0.000 | 0.000 | 0.000 |
| Il27 | 0.000 | 0.001 | 0.000 |
| 4930463O16I | 0.000 | 0.004 | 0.007 |
| Lrriq1 | 0.000 | 0.000 | 0.000 |
| Fbxw24 | 0.000 | 0.000 | 0.000 |
| 10-Mar | 0.000 | 0.001 | 0.001 |
| BC048507 | 0.000 | 0.000 | 0.001 |
| Slc8a3 | 0.000 | 0.024 | 0.013 |
| AC168977.1 | 0.000 | 0.001 | 0.001 |
| Grem2 | 0.000 | 0.004 | 0.013 |
| Pnma3 | 0.000 | 0.000 | 0.000 |
| Guca2a | 0.000 | 0.001 | 0.001 |
| Olfr558 | 0.000 | 0.001 | 0.000 |
| Cldn23 | 0.000 | 0.001 | 0.000 |
| Slmapos2 | 0.000 | 0.000 | 0.001 |
| AF529169 | 0.000 | 0.022 | 0.015 |
| A530099J19R | 0.000 | 0.000 | 0.000 |
| Psca | 0.000 | 0.000 | 0.000 |
| 4931409K22F | 0.000 | 0.000 | 0.000 |

| | | | |
|---|---|---|---|
| Gm20619 | 0.000 | 0.000 | 0.000 |
| Gm30978 | 0.000 | 0.001 | 0.000 |
| 5330439A09F | 0.000 | 0.000 | 0.000 |
| Tpo | 0.000 | 0.001 | 0.001 |
| Gm26865 | 0.000 | 0.000 | 0.000 |
| Gm28536 | 0.000 | 0.000 | 0.000 |
| Hsd3b2 | 0.000 | 0.001 | 0.001 |
| Cntn4 | 0.000 | 0.001 | 0.015 |
| Gm31108 | 0.000 | 0.000 | 0.000 |
| Gm32687 | 0.000 | 0.002 | 0.002 |
| Olfr49 | 0.000 | 0.000 | 0.000 |
| Klhdc7b | 0.000 | 0.000 | 0.000 |
| Gm17382 | 0.000 | 0.000 | 0.001 |
| Il17f | 0.000 | 0.000 | 0.001 |
| Gm15469 | 0.000 | 0.000 | 0.001 |
| Atp8b3 | 0.000 | 0.001 | 0.002 |
| Lrrc10 | 0.000 | 0.000 | 0.000 |
| Olfr318 | 0.000 | 0.000 | 0.000 |
| Ttc9b | 0.000 | 0.015 | 0.003 |
| Olfr1535 | 0.000 | 0.000 | 0.000 |
| AC140300.1 | 0.000 | 0.001 | 0.000 |
| 4930558J18R | 0.000 | 0.001 | 0.001 |
| Gm45069 | 0.000 | 0.001 | 0.001 |
| Gm47917 | 0.000 | 0.000 | 0.001 |
| 2210011C24F | 0.000 | 0.001 | 0.001 |
| Pld6 | 0.000 | 0.001 | 0.002 |
| Ece2 | 0.000 | 0.009 | 0.007 |
| Prok1 | 0.000 | 0.001 | 0.000 |
| Rims3 | 0.000 | 0.001 | 0.001 |
| Plch2 | 0.000 | 0.001 | 0.002 |
| Adamts16 | 0.000 | 0.023 | 0.081 |
| Gm34471 | 0.000 | 0.003 | 0.000 |
| Slc22a22 | 0.000 | 0.000 | 0.000 |
| AC161199.1 | 0.000 | 0.000 | 0.000 |
| A230028O05I | 0.000 | 0.003 | 0.002 |
| Ankub1 | 0.000 | 0.000 | 0.000 |
| Gm12525 | 0.000 | 0.000 | 0.000 |
| Gm12863 | 0.000 | 0.000 | 0.000 |
| Gm36208 | 0.000 | 0.000 | 0.001 |
| 2810455O05I | 0.000 | 0.005 | 0.003 |
| Gm47814 | 0.000 | 0.000 | 0.000 |
| Gm15221 | 0.000 | 0.000 | 0.000 |
| Gm34829 | 0.000 | 0.007 | 0.001 |
| Gm30709 | 0.000 | 0.000 | 0.000 |
| Gm4211 | 0.000 | 0.001 | 0.001 |
| Gm34388 | 0.000 | 0.000 | 0.000 |
| Pnma1 | 0.000 | 0.007 | 0.007 |
| 9230102O04I | 0.000 | 0.000 | 0.000 |

| | | | |
|---|---|---|---|
| Gm39929 | 0.000 | 0.001 | 0.002 |
| Hcn3 | 0.000 | 0.004 | 0.003 |
| Gm35911 | 0.000 | 0.000 | 0.000 |
| Gm26770 | 0.000 | 0.000 | 0.000 |
| D830039M14 | 0.000 | 0.001 | 0.000 |
| Gm16222 | 0.000 | 0.000 | 0.000 |
| Ubap1l | 0.000 | 0.002 | 0.002 |
| 5730403I07R | 0.000 | 0.000 | 0.000 |
| 1700001L19R | 0.000 | 0.002 | 0.003 |
| Gm32828 | 0.000 | 0.000 | 0.000 |
| Npas3 | 0.000 | 0.020 | 0.028 |
| Pabpc2 | 0.000 | 0.000 | 0.000 |
| Gm42567 | 0.000 | 0.001 | 0.001 |
| Gm11210 | 0.000 | 0.000 | 0.000 |
| Gm3519 | 0.000 | 0.000 | 0.000 |
| Muc6 | 0.000 | 0.001 | 0.002 |
| Gm5463 | 0.000 | 0.000 | 0.001 |
| Gm47270 | 0.000 | 0.001 | 0.001 |
| Gm3764 | 0.000 | 0.002 | 0.000 |
| Gm8013 | 0.000 | 0.000 | 0.001 |
| Gm26750 | 0.000 | 0.000 | 0.000 |
| Gm3460 | 0.000 | 0.000 | 0.000 |
| AC166061.1 | 0.000 | 0.000 | 0.000 |
| Olfr398 | 0.000 | 0.000 | 0.000 |
| Gm47904 | 0.000 | 0.001 | 0.003 |
| Cdkl1 | 0.000 | 0.005 | 0.007 |
| 4930526L06R | 0.000 | 0.000 | 0.000 |
| Syp | 0.000 | 0.003 | 0.009 |
| Gm6525 | 0.000 | 0.001 | 0.002 |
| 9530052C20R | 0.000 | 0.000 | 0.001 |
| Cyp3a13 | 0.000 | 0.000 | 0.001 |
| 1700120K04R | 0.000 | 0.000 | 0.000 |
| Csl | 0.000 | 0.000 | 0.000 |
| Gm47230 | 0.000 | 0.000 | 0.000 |
| Gm36264 | 0.000 | 0.000 | 0.000 |
| Gm48799 | 0.000 | 0.000 | 0.001 |
| Hus1b | 0.000 | 0.000 | 0.001 |
| Lgals2 | 0.000 | 0.000 | 0.001 |
| Kcne1 | 0.000 | 0.000 | 0.000 |
| AC168220.3 | 0.000 | 0.000 | 0.000 |
| Vmn2r90 | 0.000 | 0.000 | 0.000 |
| Chd5 | 0.000 | 0.003 | 0.001 |
| Prkcb | 0.000 | 0.001 | 0.003 |
| Jakmip3 | 0.000 | 0.001 | 0.002 |
| C230099D08R | 0.000 | 0.000 | 0.000 |
| Gm49004 | 0.000 | 0.000 | 0.000 |
| Rubcnl | 0.000 | 0.001 | 0.001 |
| Pou2f3 | 0.000 | 0.002 | 0.002 |

| | | | |
|---|---|---|---|
| Efcab9 | 0.000 | 0.001 | 0.001 |
| Galr2 | 0.000 | 0.004 | 0.002 |
| Gm11209 | 0.000 | 0.000 | 0.000 |
| Zfp853 | 0.000 | 0.008 | 0.014 |
| Gm26827 | 0.000 | 0.001 | 0.001 |
| Fank1 | 0.000 | 0.001 | 0.001 |
| Gm26752 | 0.000 | 0.000 | 0.000 |
| Gm48053 | 0.000 | 0.002 | 0.001 |
| Gm35166 | 0.000 | 0.001 | 0.002 |
| Alox12e | 0.000 | 0.000 | 0.000 |
| Meioc | 0.000 | 0.000 | 0.001 |
| Gm17039 | 0.000 | 0.000 | 0.000 |
| Gm26698 | 0.000 | 0.002 | 0.002 |
| Gm28068 | 0.000 | 0.000 | 0.001 |
| Tcp10b | 0.000 | 0.001 | 0.000 |
| Platr26 | 0.000 | 0.000 | 0.000 |
| Pdyn | 0.000 | 0.000 | 0.000 |
| 6030498E09F | 0.000 | 0.000 | 0.000 |
| Slc24a2 | 0.000 | 0.000 | 0.001 |
| Grhl3 | 0.000 | 0.009 | 0.001 |
| Cryba4 | 0.000 | 0.001 | 0.001 |
| Reep6 | 0.000 | 0.003 | 0.002 |
| 2610528A11F | 0.000 | 0.003 | 0.002 |
| Gm12158 | 0.000 | 0.000 | 0.001 |
| Gm15893 | 0.000 | 0.001 | 0.000 |
| Krt23 | 0.000 | 0.000 | 0.001 |
| 4931403G20I | 0.000 | 0.000 | 0.000 |
| Adgrf1 | 0.000 | 0.000 | 0.000 |
| Gm16146 | 0.000 | 0.000 | 0.000 |
| Sphkap | 0.000 | 0.008 | 0.013 |
| Gm15298 | 0.000 | 0.000 | 0.000 |
| Gm37854 | 0.000 | 0.000 | 0.000 |
| A730090N16I | 0.000 | 0.001 | 0.001 |
| 1700015F17F | 0.000 | 0.001 | 0.000 |
| Gm10010 | 0.000 | 0.001 | 0.001 |
| Tmem145 | 0.000 | 0.001 | 0.002 |
| 4930467K11F | 0.000 | 0.000 | 0.000 |
| Triml2 | 0.000 | 0.001 | 0.001 |
| Olfr904 | 0.000 | 0.000 | 0.000 |
| 4732414G09I | 0.000 | 0.007 | 0.005 |
| Hist1h1t | 0.000 | 0.000 | 0.000 |
| Lamp3 | 0.000 | 0.000 | 0.000 |
| Smim17 | 0.000 | 0.000 | 0.000 |
| Aurkc | 0.000 | 0.001 | 0.001 |
| Zfp773 | 0.000 | 0.004 | 0.001 |
| Ascl1 | 0.000 | 0.001 | 0.000 |
| Olfr784 | 0.000 | 0.000 | 0.000 |
| Trank1 | 0.000 | 0.004 | 0.001 |

| | | | |
|---|---|---|---|
| Gm28401 | 0.000 | 0.000 | 0.000 |
| Mroh5 | 0.000 | 0.002 | 0.002 |
| Zcchc12 | 0.000 | 0.005 | 0.010 |
| Fndc3c1 | 0.000 | 0.001 | 0.001 |
| Gm33370 | 0.000 | 0.000 | 0.001 |
| 1810020O05I | 0.000 | 0.000 | 0.000 |
| Tuba8 | 0.000 | 0.001 | 0.000 |
| Gm44702 | 0.000 | 0.002 | 0.001 |
| Gm15880 | 0.000 | 0.002 | 0.001 |
| Fsd2 | 0.000 | 0.000 | 0.000 |
| Serpinf2 | 0.000 | 0.001 | 0.001 |
| Smco1 | 0.000 | 0.000 | 0.000 |
| Spdya | 0.000 | 0.005 | 0.006 |
| Gm6741 | 0.000 | 0.000 | 0.000 |
| Drp2 | 0.000 | 0.002 | 0.004 |
| Gm20647 | 0.000 | 0.000 | 0.000 |
| Gm3294 | 0.000 | 0.001 | 0.001 |
| Hsd17b14 | 0.000 | 0.008 | 0.006 |
| Gm33586 | 0.000 | 0.000 | 0.000 |
| Lrrc63 | 0.000 | 0.000 | 0.000 |
| Slc36a2 | 0.000 | 0.002 | 0.004 |
| Sap30bpos | 0.000 | 0.000 | 0.000 |
| Hist1h2af | 0.000 | 0.001 | 0.001 |
| Clca3b | 0.000 | 0.003 | 0.001 |
| 2610528J11R | 0.000 | 0.000 | 0.000 |
| Igsf11 | 0.000 | 0.001 | 0.002 |
| Gm4577 | 0.000 | 0.001 | 0.001 |
| Txndc2 | 0.000 | 0.001 | 0.001 |
| Cdkl4 | 0.000 | 0.000 | 0.001 |
| Gm37381 | 0.000 | 0.000 | 0.000 |
| Gm16041 | 0.000 | 0.000 | 0.000 |
| Oprk1 | 0.000 | 0.000 | 0.000 |
| Npbwr1 | 0.000 | 0.000 | 0.000 |
| St18 | 0.000 | 0.000 | 0.000 |
| Sntg1 | 0.000 | 0.003 | 0.015 |
| 3110035E14F | 0.000 | 0.000 | 0.000 |
| Ppp1r42 | 0.000 | 0.000 | 0.000 |
| Gm17644 | 0.000 | 0.000 | 0.000 |
| Gm29663 | 0.000 | 0.000 | 0.000 |
| Prdm14 | 0.000 | 0.000 | 0.000 |
| Gm29570 | 0.000 | 0.000 | 0.001 |
| Gm9947 | 0.000 | 0.000 | 0.001 |
| Msc | 0.000 | 0.000 | 0.004 |
| Trpa1 | 0.000 | 0.000 | 0.000 |
| Gm7568 | 0.000 | 0.000 | 0.000 |
| Gm28376 | 0.000 | 0.000 | 0.000 |
| Gm28783 | 0.000 | 0.000 | 0.000 |
| Gm28153 | 0.000 | 0.000 | 0.000 |

| | | | |
|---|---|---|---|
| Gm28340 | 0.000 | 0.000 | 0.000 |
| Pkhd1 | 0.000 | 0.000 | 0.000 |
| Gm28653 | 0.000 | 0.000 | 0.000 |
| Il17a | 0.000 | 0.000 | 0.000 |
| Gm28836 | 0.000 | 0.000 | 0.001 |
| Khdc1a | 0.000 | 0.000 | 0.000 |
| Khdc1c | 0.000 | 0.000 | 0.000 |
| Khdc1b | 0.000 | 0.000 | 0.000 |
| Gm26580 | 0.000 | 0.001 | 0.000 |
| 4933415F23F | 0.000 | 0.000 | 0.000 |
| Col19a1 | 0.000 | 0.014 | 0.001 |
| Adgrb3 | 0.000 | 0.005 | 0.011 |
| Gm9898 | 0.000 | 0.000 | 0.000 |
| Gm37724 | 0.000 | 0.000 | 0.000 |
| Gm28306 | 0.000 | 0.000 | 0.000 |
| 4930568A12F | 0.000 | 0.000 | 0.000 |
| Prss39 | 0.000 | 0.000 | 0.000 |
| Cfc1 | 0.000 | 0.000 | 0.000 |
| Amer3 | 0.000 | 0.000 | 0.000 |
| Gm33222 | 0.000 | 0.000 | 0.000 |
| Gm37335 | 0.000 | 0.000 | 0.000 |
| Gm33280 | 0.000 | 0.000 | 0.001 |
| Fer1l5 | 0.000 | 0.000 | 0.000 |
| Gm38033 | 0.000 | 0.002 | 0.002 |
| Ankrd23 | 0.000 | 0.004 | 0.003 |
| Gm42417 | 0.000 | 0.001 | 0.002 |
| Gm33533 | 0.000 | 0.000 | 0.000 |
| Gm37020 | 0.000 | 0.000 | 0.000 |
| Cnga3 | 0.000 | 0.000 | 0.000 |
| 4930556I23R | 0.000 | 0.000 | 0.000 |
| Gm15457 | 0.000 | 0.000 | 0.000 |
| Gm5099 | 0.000 | 0.000 | 0.000 |
| Gm16150 | 0.000 | 0.000 | 0.000 |
| Gm16151 | 0.000 | 0.000 | 0.000 |
| D930019O06I | 0.000 | 0.000 | 0.000 |
| Gm3646 | 0.000 | 0.005 | 0.000 |
| 1700066B17F | 0.000 | 0.002 | 0.001 |
| Gm16894 | 0.000 | 0.001 | 0.002 |
| Il1rl1 | 0.000 | 0.001 | 0.000 |
| Slc9a4 | 0.000 | 0.001 | 0.001 |
| Gm37915 | 0.000 | 0.000 | 0.000 |
| Gm29260 | 0.000 | 0.000 | 0.000 |
| Pantr1 | 0.000 | 0.000 | 0.001 |
| Pou3f3 | 0.000 | 0.000 | 0.001 |
| Gm29156 | 0.000 | 0.000 | 0.001 |
| Mettl21e | 0.000 | 0.000 | 0.001 |
| 4930521E06F | 0.000 | 0.000 | 0.000 |
| Dnah7c | 0.000 | 0.000 | 0.000 |

| | | | |
|---|---|---|---|
| Gm28826 | 0.000 | 0.000 | 0.000 |
| Gm28322 | 0.000 | 0.000 | 0.000 |
| 1700072G22I | 0.000 | 0.000 | 0.000 |
| Gm17767 | 0.000 | 0.000 | 0.000 |
| Gm28055 | 0.000 | 0.000 | 0.000 |
| Gm553 | 0.000 | 0.001 | 0.000 |
| Mstn | 0.000 | 0.000 | 0.000 |
| 1700019A02F | 0.000 | 0.000 | 0.000 |
| Gm28777 | 0.000 | 0.000 | 0.000 |
| Dnah7a | 0.000 | 0.001 | 0.002 |
| Ccdc150 | 0.000 | 0.000 | 0.000 |
| 4930444A19F | 0.000 | 0.000 | 0.000 |
| Boll | 0.000 | 0.000 | 0.000 |
| 9130227L01F | 0.000 | 0.000 | 0.000 |
| Gm28240 | 0.000 | 0.000 | 0.000 |
| Aox4 | 0.000 | 0.000 | 0.001 |
| Gm15759 | 0.000 | 0.000 | 0.000 |
| Aox2 | 0.000 | 0.000 | 0.002 |
| C2cd6b | 0.000 | 0.000 | 0.000 |
| Gm29018 | 0.000 | 0.000 | 0.000 |
| C2cd6 | 0.000 | 0.000 | 0.000 |
| Cdk15 | 0.000 | 0.002 | 0.002 |
| Gm29016 | 0.000 | 0.000 | 0.000 |
| Gm26813 | 0.000 | 0.001 | 0.000 |
| Gm28411 | 0.000 | 0.000 | 0.000 |
| Gm11579 | 0.000 | 0.000 | 0.001 |
| 2310016D23I | 0.000 | 0.000 | 0.000 |
| Ctla4 | 0.000 | 0.000 | 0.000 |
| Gm11587 | 0.000 | 0.000 | 0.000 |
| 9530026F06F | 0.000 | 0.000 | 0.001 |
| Gm11588 | 0.000 | 0.000 | 0.000 |
| Pard3bos1 | 0.000 | 0.001 | 0.000 |
| Pard3bos2 | 0.000 | 0.000 | 0.000 |
| Pard3bos3 | 0.000 | 0.000 | 0.000 |
| Gm29084 | 0.000 | 0.000 | 0.000 |
| Gm29083 | 0.000 | 0.000 | 0.000 |
| Gm4208 | 0.000 | 0.000 | 0.000 |
| Gpr1 | 0.000 | 0.000 | 0.001 |
| Gm39653 | 0.000 | 0.000 | 0.000 |
| Mdh1b | 0.000 | 0.000 | 0.000 |
| 4933402D24I | 0.000 | 0.000 | 0.000 |
| Gm13749 | 0.000 | 0.001 | 0.004 |
| Gm28981 | 0.000 | 0.000 | 0.000 |
| Gm28982 | 0.000 | 0.000 | 0.000 |
| Gm29152 | 0.000 | 0.000 | 0.000 |
| Cryge | 0.000 | 0.000 | 0.001 |
| Crygd | 0.000 | 0.000 | 0.000 |
| Crygc | 0.000 | 0.000 | 0.000 |

| | | | |
|---|---|---|---|
| Cryga | 0.000 | 0.000 | 0.000 |
| D630023F18F | 0.000 | 0.001 | 0.000 |
| Pth2r | 0.000 | 0.000 | 0.000 |
| Crygf | 0.000 | 0.000 | 0.000 |
| Gm10558 | 0.000 | 0.000 | 0.000 |
| Unc80 | 0.000 | 0.000 | 0.000 |
| Gm15669 | 0.000 | 0.000 | 0.000 |
| Gm28497 | 0.000 | 0.000 | 0.000 |
| Gm29114 | 0.000 | 0.000 | 0.000 |
| Spag16 | 0.000 | 0.000 | 0.000 |
| Abca12 | 0.000 | 0.000 | 0.000 |
| Gm8883 | 0.000 | 0.000 | 0.000 |
| Gm28818 | 0.000 | 0.002 | 0.001 |
| 4933417E11F | 0.000 | 0.000 | 0.000 |
| Gm4319 | 0.000 | 0.000 | 0.000 |
| 4-Mar | 0.000 | 0.000 | 0.000 |
| Ankar | 0.000 | 0.000 | 0.000 |
| Gm29185 | 0.000 | 0.000 | 0.000 |
| C530043A13F | 0.000 | 0.000 | 0.000 |
| 6030407O03I | 0.000 | 0.001 | 0.001 |
| Gpbar1 | 0.000 | 0.005 | 0.003 |
| Plcd4 | 0.000 | 0.002 | 0.001 |
| Gm29539 | 0.000 | 0.000 | 0.000 |
| Wnt10a | 0.000 | 0.000 | 0.000 |
| Cdk5r2 | 0.000 | 0.000 | 0.001 |
| Cfap65 | 0.000 | 0.000 | 0.000 |
| A630095N17I | 0.000 | 0.000 | 0.000 |
| Gm28294 | 0.000 | 0.000 | 0.000 |
| Resp18 | 0.000 | 0.000 | 0.001 |
| Asic4 | 0.000 | 0.002 | 0.000 |
| Gm816 | 0.000 | 0.000 | 0.000 |
| Pax3 | 0.000 | 0.001 | 0.001 |
| BC035947 | 0.000 | 0.000 | 0.000 |
| Mogat1 | 0.000 | 0.000 | 0.001 |
| 5730419F03F | 0.000 | 0.000 | 0.000 |
| Gm29536 | 0.000 | 0.000 | 0.000 |
| Gm29187 | 0.000 | 0.000 | 0.000 |
| Scg2 | 0.000 | 0.002 | 0.007 |
| 2310015K22F | 0.000 | 0.000 | 0.000 |
| 1700016L21F | 0.000 | 0.000 | 0.000 |
| Nyap2 | 0.000 | 0.000 | 0.000 |
| Gm9747 | 0.000 | 0.000 | 0.000 |
| Gm6217 | 0.000 | 0.000 | 0.000 |
| Daw1 | 0.000 | 0.000 | 0.000 |
| 4933436I20R | 0.000 | 0.000 | 0.000 |
| A530032D15I | 0.000 | 0.000 | 0.000 |
| Gm16028 | 0.000 | 0.000 | 0.000 |
| Gm16025 | 0.000 | 0.000 | 0.000 |

| | | | |
|---|---|---|---|
| G530012D18I | 0.000 | 0.000 | 0.000 |
| Gm16094 | 0.000 | 0.000 | 0.000 |
| 9930111H07I | 0.000 | 0.000 | 0.000 |
| Gm28884 | 0.000 | 0.000 | 0.000 |
| Gpr55 | 0.000 | 0.000 | 0.000 |
| Spata3 | 0.000 | 0.000 | 0.000 |
| Htr2b | 0.000 | 0.000 | 0.000 |
| Gm21972 | 0.000 | 0.001 | 0.000 |
| Gm28626 | 0.000 | 0.000 | 0.000 |
| Nmur1 | 0.000 | 0.000 | 0.000 |
| Tex44 | 0.000 | 0.000 | 0.000 |
| Gm28375 | 0.000 | 0.000 | 0.000 |
| Gm29371 | 0.000 | 0.000 | 0.000 |
| Alppl2 | 0.000 | 0.000 | 0.000 |
| Gm29374 | 0.000 | 0.000 | 0.000 |
| Ecel1 | 0.000 | 0.001 | 0.002 |
| Chrng | 0.000 | 0.000 | 0.001 |
| Gm19582 | 0.000 | 0.000 | 0.000 |
| Ugt1a7c | 0.000 | 0.000 | 0.001 |
| Ugt1a6b | 0.000 | 0.000 | 0.000 |
| Ugt1a6a | 0.000 | 0.000 | 0.002 |
| Ugt1a5 | 0.000 | 0.000 | 0.000 |
| Ugt1a2 | 0.000 | 0.000 | 0.000 |
| Spp2 | 0.000 | 0.000 | 0.000 |
| Gm29538 | 0.000 | 0.000 | 0.000 |
| Glrp1 | 0.000 | 0.000 | 0.001 |
| Gm19589 | 0.000 | 0.000 | 0.000 |
| Gm29336 | 0.000 | 0.000 | 0.000 |
| Platr5 | 0.000 | 0.000 | 0.000 |
| Asb18 | 0.000 | 0.000 | 0.000 |
| Iqca | 0.000 | 0.000 | 0.000 |
| 4930434B07I | 0.000 | 0.000 | 0.000 |
| Gm28199 | 0.000 | 0.000 | 0.000 |
| Gm26683 | 0.000 | 0.000 | 0.000 |
| Gm17090 | 0.000 | 0.000 | 0.000 |
| Espnl | 0.000 | 0.000 | 0.000 |
| Gm29100 | 0.000 | 0.002 | 0.009 |
| Olfr12 | 0.000 | 0.000 | 0.000 |
| Otos | 0.000 | 0.000 | 0.000 |
| Gm29480 | 0.000 | 0.000 | 0.000 |
| Gm28086 | 0.000 | 0.000 | 0.000 |
| Agxt | 0.000 | 0.000 | 0.000 |
| 2310007B03I | 0.000 | 0.000 | 0.000 |
| Crocc2 | 0.000 | 0.000 | 0.002 |
| Ano7 | 0.000 | 0.000 | 0.001 |
| Gm10550 | 0.000 | 0.000 | 0.000 |
| Gal3st2 | 0.000 | 0.000 | 0.000 |
| Gal3st2b | 0.000 | 0.000 | 0.000 |

| | | | |
|---|---|---|---|
| Gal3st2c | 0.000 | 0.000 | 0.000 |
| Pdcd1 | 0.000 | 0.000 | 0.000 |
| Slco4c1 | 0.000 | 0.000 | 0.000 |
| Panct2 | 0.000 | 0.000 | 0.000 |
| Gm28901 | 0.000 | 0.000 | 0.000 |
| Slco6d1 | 0.000 | 0.000 | 0.000 |
| Cntnap5b | 0.000 | 0.000 | 0.000 |
| Cdh20 | 0.000 | 0.000 | 0.001 |
| A530053M12 | 0.000 | 0.000 | 0.000 |
| D630008O14I | 0.000 | 0.000 | 0.000 |
| Serpinb12 | 0.000 | 0.000 | 0.000 |
| Serpinb11 | 0.000 | 0.000 | 0.000 |
| Serpinb7 | 0.000 | 0.000 | 0.001 |
| Serpinb2 | 0.000 | 0.000 | 0.000 |
| Gm15389 | 0.000 | 0.000 | 0.000 |
| D830032E09F | 0.000 | 0.000 | 0.000 |
| Cdh7 | 0.000 | 0.000 | 0.000 |
| Cdh19 | 0.000 | 0.002 | 0.001 |
| Gm29088 | 0.000 | 0.000 | 0.001 |
| 9330185C12F | 0.000 | 0.000 | 0.000 |
| Cntnap5a | 0.000 | 0.000 | 0.001 |
| Gm28867 | 0.000 | 0.001 | 0.001 |
| Gm27184 | 0.000 | 0.001 | 0.000 |
| 3830432H09I | 0.000 | 0.000 | 0.000 |
| Cfap221 | 0.000 | 0.000 | 0.000 |
| Sctr | 0.000 | 0.000 | 0.000 |
| C1ql2 | 0.000 | 0.000 | 0.002 |
| En1 | 0.000 | 0.000 | 0.000 |
| 2610027F03F | 0.000 | 0.000 | 0.000 |
| Gm29348 | 0.000 | 0.000 | 0.000 |
| Celrr | 0.000 | 0.000 | 0.000 |
| 2210011K15F | 0.000 | 0.000 | 0.000 |
| Acmsd | 0.000 | 0.000 | 0.000 |
| Gm16081 | 0.000 | 0.000 | 0.000 |
| Cd55b | 0.000 | 0.000 | 0.000 |
| Gm15674 | 0.000 | 0.000 | 0.000 |
| Zp3r | 0.000 | 0.000 | 0.000 |
| C4bp | 0.000 | 0.002 | 0.001 |
| Gm29427 | 0.000 | 0.001 | 0.001 |
| Gm28856 | 0.000 | 0.000 | 0.000 |
| Gm15848 | 0.000 | 0.000 | 0.000 |
| Pigr | 0.000 | 0.000 | 0.000 |
| Fcmr | 0.000 | 0.000 | 0.000 |
| Il10 | 0.000 | 0.000 | 0.000 |
| Avpr1b | 0.000 | 0.000 | 0.000 |
| Slc26a9 | 0.000 | 0.000 | 0.000 |
| Gm29629 | 0.000 | 0.000 | 0.000 |
| 1700037F24F | 0.000 | 0.000 | 0.000 |

| | | | |
|---|---|---|---|
| F730311O21F | 0.000 | 0.000 | 0.000 |
| Cntn2 | 0.000 | 0.001 | 0.000 |
| Gm28609 | 0.000 | 0.000 | 0.000 |
| Gm19461 | 0.000 | 0.000 | 0.000 |
| Zc3h11a.1 | 0.000 | 0.000 | 0.000 |
| Gm15851 | 0.000 | 0.000 | 0.000 |
| Gm1627 | 0.000 | 0.000 | 0.000 |
| Mybph | 0.000 | 0.000 | 0.000 |
| Adora1 | 0.000 | 0.004 | 0.001 |
| Myog | 0.000 | 0.000 | 0.000 |
| Gm26783 | 0.000 | 0.000 | 0.000 |
| Gm10535 | 0.000 | 0.000 | 0.000 |
| Pkp1 | 0.000 | 0.000 | 0.000 |
| Igfn1 | 0.000 | 0.000 | 0.000 |
| Cacna1s | 0.000 | 0.000 | 0.000 |
| Gm37759 | 0.000 | 0.000 | 0.000 |
| Mroh3 | 0.000 | 0.000 | 0.000 |
| A430034D21I | 0.000 | 0.000 | 0.000 |
| Gm37552 | 0.000 | 0.000 | 0.000 |
| Gm37799 | 0.000 | 0.000 | 0.000 |
| Gm33994 | 0.000 | 0.000 | 0.000 |
| Platr23 | 0.000 | 0.001 | 0.001 |
| Gm26936 | 0.000 | 0.000 | 0.000 |
| Gm28556 | 0.000 | 0.000 | 0.000 |
| Atp6v1g3 | 0.000 | 0.000 | 0.000 |
| Gm28501 | 0.000 | 0.000 | 0.000 |
| Lhx9 | 0.000 | 0.001 | 0.000 |
| Crb1 | 0.000 | 0.000 | 0.000 |
| F13b | 0.000 | 0.000 | 0.000 |
| 4930590L20F | 0.000 | 0.000 | 0.000 |
| Rgs18 | 0.000 | 0.001 | 0.000 |
| Gm15584 | 0.000 | 0.000 | 0.000 |
| Ptgs2os2 | 0.000 | 0.000 | 0.003 |
| Pdc | 0.000 | 0.000 | 0.000 |
| 2310030A07F | 0.000 | 0.000 | 0.000 |
| Prg4 | 0.000 | 0.001 | 0.008 |
| Gm47985 | 0.000 | 0.000 | 0.000 |
| Gm8947 | 0.000 | 0.000 | 0.000 |
| Apobec4 | 0.000 | 0.000 | 0.000 |
| Shcbp1l | 0.000 | 0.001 | 0.001 |
| C230024C17F | 0.000 | 0.000 | 0.001 |
| Rgsl1 | 0.000 | 0.001 | 0.001 |
| Teddm1b | 0.000 | 0.000 | 0.000 |
| Gm29290 | 0.000 | 0.000 | 0.000 |
| Gm29291 | 0.000 | 0.000 | 0.000 |
| A830008E24F | 0.000 | 0.000 | 0.001 |
| Gm29282 | 0.000 | 0.000 | 0.000 |
| Lhx4 | 0.000 | 0.000 | 0.001 |

| | | | |
|---|---|---|---|
| Tdrd5 | 0.000 | 0.000 | 0.000 |
| 4930518J20R | 0.000 | 0.000 | 0.000 |
| 4930439D14I | 0.000 | 0.000 | 0.000 |
| Gm15486 | 0.000 | 0.000 | 0.000 |
| Brinp2 | 0.000 | 0.000 | 0.000 |
| Gm37679 | 0.000 | 0.000 | 0.001 |
| Pappa2 | 0.000 | 0.014 | 0.017 |
| 4930562F17F | 0.000 | 0.000 | 0.000 |
| Gm37294 | 0.000 | 0.000 | 0.000 |
| 4930562F07F | 0.000 | 0.003 | 0.001 |
| Gm36975 | 0.000 | 0.000 | 0.001 |
| Gm37083 | 0.000 | 0.000 | 0.000 |
| Gm37362 | 0.000 | 0.000 | 0.000 |
| Tex50 | 0.000 | 0.000 | 0.000 |
| Tnfsf4 | 0.000 | 0.005 | 0.001 |
| Tnfsf18 | 0.000 | 0.000 | 0.000 |
| Fasl | 0.000 | 0.000 | 0.000 |
| Myocos | 0.000 | 0.000 | 0.001 |
| Fmo6 | 0.000 | 0.000 | 0.003 |
| Mroh9 | 0.000 | 0.000 | 0.000 |
| Gm29500 | 0.000 | 0.000 | 0.000 |
| Mettl11b | 0.000 | 0.000 | 0.000 |
| Gm16587 | 0.000 | 0.000 | 0.000 |
| Gm26685 | 0.000 | 0.001 | 0.000 |
| Gm37754 | 0.000 | 0.000 | 0.000 |
| Gm37184 | 0.000 | 0.000 | 0.001 |
| Gm36945 | 0.000 | 0.000 | 0.000 |
| Xcl1 | 0.000 | 0.000 | 0.000 |
| 4930568G15I | 0.000 | 0.000 | 0.000 |
| Tbx19 | 0.000 | 0.000 | 0.000 |
| Gm38325 | 0.000 | 0.000 | 0.000 |
| Gm37073 | 0.000 | 0.000 | 0.000 |
| Gm26665 | 0.000 | 0.000 | 0.000 |
| Mael | 0.000 | 0.000 | 0.000 |
| Gm16701 | 0.000 | 0.000 | 0.000 |
| Gm37982 | 0.000 | 0.000 | 0.000 |
| Gm20711 | 0.000 | 0.001 | 0.001 |
| Ccdc190 | 0.000 | 0.007 | 0.005 |
| Gm37502 | 0.000 | 0.001 | 0.001 |
| Fcrlb | 0.000 | 0.000 | 0.000 |
| Fcrla | 0.000 | 0.000 | 0.000 |
| Gm10522 | 0.000 | 0.000 | 0.000 |
| Nr1i3 | 0.000 | 0.000 | 0.000 |
| Ly9 | 0.000 | 0.000 | 0.000 |
| Gm37787 | 0.000 | 0.000 | 0.000 |
| Gm10521 | 0.000 | 0.000 | 0.000 |
| Slamf6 | 0.000 | 0.000 | 0.000 |
| Nhlh1 | 0.000 | 0.000 | 0.000 |

| | | | |
|---|---|---|---|
| Gm37950 | 0.000 | 0.000 | 0.000 |
| Kcnj9 | 0.000 | 0.000 | 0.000 |
| Gm36937 | 0.000 | 0.000 | 0.000 |
| Slamf8 | 0.000 | 0.001 | 0.000 |
| Fcrl6 | 0.000 | 0.000 | 0.000 |
| Fcer1a | 0.000 | 0.000 | 0.000 |
| Ifi214 | 0.000 | 0.000 | 0.000 |
| Ifi213 | 0.000 | 0.000 | 0.000 |
| Ifi209 | 0.000 | 0.001 | 0.001 |
| Olfr432 | 0.000 | 0.000 | 0.000 |
| Olfr414 | 0.000 | 0.000 | 0.000 |
| Wdr64 | 0.000 | 0.000 | 0.000 |
| Gm38100 | 0.000 | 0.000 | 0.000 |
| Gm38101 | 0.000 | 0.000 | 0.000 |
| Gm37306 | 0.000 | 0.000 | 0.000 |
| 2310043L19F | 0.000 | 0.000 | 0.000 |
| Gm16564 | 0.000 | 0.000 | 0.000 |
| Kif28 | 0.000 | 0.001 | 0.001 |
| Gm10518 | 0.000 | 0.000 | 0.000 |
| Gm37336 | 0.000 | 0.001 | 0.001 |
| Gm31728 | 0.000 | 0.000 | 0.001 |
| Mixl1 | 0.000 | 0.000 | 0.000 |
| Gm38293 | 0.000 | 0.000 | 0.000 |
| Lefty2 | 0.000 | 0.000 | 0.001 |
| 2210411M09 | 0.000 | 0.000 | 0.000 |
| Ccdc121 | 0.000 | 0.001 | 0.002 |
| Gm5533 | 0.000 | 0.001 | 0.001 |
| 1700047M11 | 0.000 | 0.001 | 0.004 |
| Gm37885 | 0.000 | 0.002 | 0.001 |
| Capn8 | 0.000 | 0.000 | 0.000 |
| Gm38079 | 0.000 | 0.001 | 0.001 |
| Gm33973 | 0.000 | 0.000 | 0.000 |
| Gm37223 | 0.000 | 0.000 | 0.001 |
| 1700112H15I | 0.000 | 0.000 | 0.000 |
| Gm37800 | 0.000 | 0.000 | 0.000 |
| 1-Mar | 0.000 | 0.001 | 0.002 |
| Gm38251 | 0.000 | 0.000 | 0.000 |
| Gm34732 | 0.000 | 0.000 | 0.000 |
| 9630028B13F | 0.000 | 0.000 | 0.000 |
| Slc30a10 | 0.000 | 0.000 | 0.000 |
| A730004F24F | 0.000 | 0.000 | 0.000 |
| D1Pas1 | 0.000 | 0.000 | 0.000 |
| Spata17 | 0.000 | 0.000 | 0.001 |
| 1700007P06F | 0.000 | 0.000 | 0.001 |
| Gm38155 | 0.000 | 0.000 | 0.000 |
| Gm37896 | 0.000 | 0.000 | 0.001 |
| Gm36388 | 0.000 | 0.000 | 0.000 |
| Ush2a | 0.000 | 0.000 | 0.000 |

| | | | |
|---|---|---|---|
| A430027H14I | 0.000 | 0.000 | 0.000 |
| Gm28172 | 0.000 | 0.000 | 0.000 |
| Gm31373 | 0.000 | 0.000 | 0.000 |
| A230020J21R | 0.000 | 0.000 | 0.000 |
| Spata45 | 0.000 | 0.000 | 0.000 |
| Gm38161 | 0.000 | 0.000 | 0.000 |
| Gm32200 | 0.000 | 0.000 | 0.000 |
| Rd3 | 0.000 | 0.000 | 0.000 |
| 1700065J18R | 0.000 | 0.000 | 0.000 |
| Gm31406 | 0.000 | 0.000 | 0.000 |
| 4930570N18I | 0.000 | 0.000 | 0.000 |
| Gm37650 | 0.000 | 0.000 | 0.000 |
| Gm37027 | 0.000 | 0.000 | 0.000 |
| Acbd7 | 0.000 | 0.001 | 0.001 |
| Olah | 0.000 | 0.000 | 0.000 |
| Meig1 | 0.000 | 0.001 | 0.000 |
| Gm13184 | 0.000 | 0.000 | 0.000 |
| Gm13180 | 0.000 | 0.000 | 0.000 |
| 1700080N15I | 0.000 | 0.000 | 0.000 |
| Gm13179 | 0.000 | 0.001 | 0.003 |
| Ucma | 0.000 | 0.000 | 0.000 |
| 4930551O13I | 0.000 | 0.000 | 0.000 |
| Gm26776 | 0.000 | 0.000 | 0.000 |
| Gm13267 | 0.000 | 0.000 | 0.000 |
| Gm13389 | 0.000 | 0.000 | 0.000 |
| Gm26565 | 0.000 | 0.001 | 0.003 |
| Gm28641 | 0.000 | 0.000 | 0.000 |
| Itih2 | 0.000 | 0.003 | 0.006 |
| Gm13293 | 0.000 | 0.000 | 0.000 |
| Gm10851 | 0.000 | 0.000 | 0.000 |
| Il2ra | 0.000 | 0.000 | 0.000 |
| Slc39a12 | 0.000 | 0.000 | 0.000 |
| Gm10848 | 0.000 | 0.019 | 0.006 |
| Malrd1 | 0.000 | 0.000 | 0.000 |
| H2al2a | 0.000 | 0.000 | 0.000 |
| Spag6 | 0.000 | 0.001 | 0.000 |
| Carlr | 0.000 | 0.000 | 0.000 |
| 4930447M23 | 0.000 | 0.000 | 0.000 |
| Ptf1a | 0.000 | 0.000 | 0.000 |
| Gm17171 | 0.000 | 0.000 | 0.000 |
| Gm13362 | 0.000 | 0.000 | 0.000 |
| Gm13376 | 0.000 | 0.000 | 0.001 |
| Nxph2 | 0.000 | 0.000 | 0.000 |
| Tbpl2 | 0.000 | 0.000 | 0.000 |
| Il1f9 | 0.000 | 0.000 | 0.000 |
| Il1f10 | 0.000 | 0.000 | 0.000 |
| Gm13411 | 0.000 | 0.000 | 0.000 |
| Psd4 | 0.000 | 0.001 | 0.003 |

| | | | |
|---|---|---|---|
| Cacna1b | 0.000 | 0.001 | 0.001 |
| Entpd8 | 0.000 | 0.000 | 0.000 |
| Stpg3 | 0.000 | 0.000 | 0.001 |
| Fam166a | 0.000 | 0.001 | 0.000 |
| Slc34a3 | 0.000 | 0.000 | 0.000 |
| Cysrt1 | 0.000 | 0.000 | 0.000 |
| AL732309.1 | 0.000 | 0.000 | 0.000 |
| Grin1 | 0.000 | 0.000 | 0.002 |
| Gm26702 | 0.000 | 0.000 | 0.000 |
| Fcnaos | 0.000 | 0.000 | 0.000 |
| Ccdc183 | 0.000 | 0.000 | 0.000 |
| Lcn5 | 0.000 | 0.000 | 0.000 |
| Lcn10 | 0.000 | 0.000 | 0.000 |
| Lcn9 | 0.000 | 0.000 | 0.000 |
| Sohlh1 | 0.000 | 0.000 | 0.003 |
| Gm13554 | 0.000 | 0.000 | 0.000 |
| 1810012K08F | 0.000 | 0.000 | 0.000 |
| Lhx3 | 0.000 | 0.000 | 0.000 |
| Ccdc187 | 0.000 | 0.002 | 0.003 |
| Snhg7os | 0.000 | 0.000 | 0.001 |
| Lcn4 | 0.000 | 0.000 | 0.000 |
| Adamts13 | 0.000 | 0.000 | 0.000 |
| Gm13398 | 0.000 | 0.000 | 0.000 |
| Fam163b | 0.000 | 0.001 | 0.001 |
| Dbhos | 0.000 | 0.000 | 0.000 |
| Dbh | 0.000 | 0.002 | 0.001 |
| Sardhos | 0.000 | 0.000 | 0.000 |
| Fcnb | 0.000 | 0.000 | 0.000 |
| F730016J06R | 0.000 | 0.000 | 0.000 |
| Gm13373 | 0.000 | 0.000 | 0.000 |
| Gbgt1 | 0.000 | 0.000 | 0.002 |
| Gm10134 | 0.000 | 0.000 | 0.000 |
| Gm13381 | 0.000 | 0.002 | 0.001 |
| Gfi1b | 0.000 | 0.000 | 0.000 |
| Barhl1 | 0.000 | 0.001 | 0.000 |
| Ntng2 | 0.000 | 0.000 | 0.000 |
| Gm13547 | 0.000 | 0.000 | 0.000 |
| Gm16323 | 0.000 | 0.000 | 0.000 |
| Gm14487 | 0.000 | 0.001 | 0.000 |
| 9330198N18I | 0.000 | 0.002 | 0.002 |
| Gm14486 | 0.000 | 0.001 | 0.001 |
| 4930527E20F | 0.000 | 0.000 | 0.000 |
| Gm13425 | 0.000 | 0.000 | 0.000 |
| Prdm12 | 0.000 | 0.000 | 0.000 |
| Gm13427 | 0.000 | 0.000 | 0.000 |
| Qrfp | 0.000 | 0.000 | 0.000 |
| Fibcd1 | 0.000 | 0.000 | 0.000 |
| Gm13609 | 0.000 | 0.000 | 0.000 |

| | | | |
|---|---|---|---|
| 9430097D07I | 0.000 | 0.000 | 0.000 |
| Pip5kl1 | 0.000 | 0.001 | 0.001 |
| Cfap157 | 0.000 | 0.000 | 0.000 |
| Gm13528 | 0.000 | 0.000 | 0.000 |
| Lmx1b | 0.000 | 0.000 | 0.000 |
| C130021I20R | 0.000 | 0.000 | 0.000 |
| 9430024E24F | 0.000 | 0.000 | 0.000 |
| Nron | 0.000 | 0.000 | 0.000 |
| Gm13403 | 0.000 | 0.000 | 0.000 |
| Gm13404 | 0.000 | 0.000 | 0.000 |
| C230014O12I | 0.000 | 0.000 | 0.000 |
| Hc | 0.000 | 0.000 | 0.000 |
| Gm13605 | 0.000 | 0.000 | 0.000 |
| 4930402F06F | 0.000 | 0.000 | 0.000 |
| 9030204H09I | 0.000 | 0.000 | 0.000 |
| Olfr338 | 0.000 | 0.000 | 0.000 |
| Olfr344 | 0.000 | 0.000 | 0.000 |
| Olfr354 | 0.000 | 0.000 | 0.000 |
| Olfr362 | 0.000 | 0.000 | 0.000 |
| Olfr364-ps1 | 0.000 | 0.000 | 0.000 |
| Olfr366 | 0.000 | 0.000 | 0.000 |
| Gm13556 | 0.000 | 0.000 | 0.000 |
| Lhx2 | 0.000 | 0.000 | 0.000 |
| Gm13589 | 0.000 | 0.000 | 0.000 |
| Nr5a1os | 0.000 | 0.000 | 0.000 |
| Kynu | 0.000 | 0.000 | 0.000 |
| Arhgap15os | 0.000 | 0.000 | 0.000 |
| Gm13479 | 0.000 | 0.000 | 0.000 |
| 1700019E08F | 0.000 | 0.000 | 0.000 |
| Gm13481 | 0.000 | 0.000 | 0.000 |
| Gm48908 | 0.000 | 0.001 | 0.000 |
| Gm13497 | 0.000 | 0.000 | 0.000 |
| Gm13490 | 0.000 | 0.000 | 0.000 |
| Gm13522 | 0.000 | 0.000 | 0.000 |
| Gm26603 | 0.000 | 0.000 | 0.000 |
| Galnt13 | 0.000 | 0.001 | 0.001 |
| Gm26794 | 0.000 | 0.000 | 0.000 |
| Kcnj3 | 0.000 | 0.000 | 0.000 |
| Gm33594 | 0.000 | 0.000 | 0.000 |
| Ermn | 0.000 | 0.000 | 0.000 |
| Gm13546 | 0.000 | 0.000 | 0.000 |
| Gm17409 | 0.000 | 0.000 | 0.000 |
| Gm13572 | 0.000 | 0.000 | 0.000 |
| Gm13583 | 0.000 | 0.000 | 0.000 |
| Itgb6 | 0.000 | 0.000 | 0.000 |
| Slc4a10 | 0.000 | 0.003 | 0.003 |
| Gm13561 | 0.000 | 0.000 | 0.000 |
| Gm13565 | 0.000 | 0.000 | 0.000 |

| | | | |
|---|---|---|---|
| Gm13564 | 0.000 | 0.000 | 0.000 |
| Gcg | 0.000 | 0.001 | 0.001 |
| Kcnh7 | 0.000 | 0.001 | 0.000 |
| Gm13575 | 0.000 | 0.000 | 0.000 |
| Slc38a11 | 0.000 | 0.012 | 0.001 |
| Scn3a | 0.000 | 0.004 | 0.006 |
| Gm13595 | 0.000 | 0.000 | 0.000 |
| Gm13619 | 0.000 | 0.000 | 0.000 |
| Gm13617 | 0.000 | 0.000 | 0.000 |
| Scn1a | 0.000 | 0.001 | 0.000 |
| Gm13629 | 0.000 | 0.000 | 0.000 |
| Scn9a | 0.000 | 0.000 | 0.001 |
| Gm1322 | 0.000 | 0.000 | 0.000 |
| Xirp2 | 0.000 | 0.000 | 0.000 |
| Gm26743 | 0.000 | 0.000 | 0.000 |
| Gm13596 | 0.000 | 0.000 | 0.001 |
| Abcb11 | 0.000 | 0.000 | 0.000 |
| Gm13613 | 0.000 | 0.000 | 0.000 |
| Dhrs9 | 0.000 | 0.002 | 0.003 |
| Lrp2 | 0.000 | 0.001 | 0.001 |
| Gad1 | 0.000 | 0.001 | 0.000 |
| Gm13632 | 0.000 | 0.001 | 0.000 |
| Dlx1as | 0.000 | 0.000 | 0.000 |
| Dlx1 | 0.000 | 0.006 | 0.002 |
| Gm13647 | 0.000 | 0.000 | 0.000 |
| Gm17250 | 0.000 | 0.000 | 0.000 |
| Rapgef4os3 | 0.000 | 0.000 | 0.000 |
| 8430437L04F | 0.000 | 0.000 | 0.000 |
| Sp9 | 0.000 | 0.000 | 0.000 |
| Gm13709 | 0.000 | 0.000 | 0.000 |
| Gm13707 | 0.000 | 0.000 | 0.000 |
| Gm10822 | 0.000 | 0.000 | 0.000 |
| A630050E04F | 0.000 | 0.000 | 0.000 |
| 4930445N08I | 0.000 | 0.000 | 0.000 |
| Hoxd13 | 0.000 | 0.000 | 0.000 |
| Hoxd10 | 0.000 | 0.000 | 0.000 |
| Gm28793 | 0.000 | 0.000 | 0.000 |
| Gm28230 | 0.000 | 0.000 | 0.000 |
| Gm14424 | 0.000 | 0.001 | 0.001 |
| Gm14426 | 0.000 | 0.000 | 0.000 |
| Hoxd1 | 0.000 | 0.000 | 0.000 |
| Gm13652 | 0.000 | 0.000 | 0.000 |
| A330043C09F | 0.000 | 0.000 | 0.000 |
| Gm13656 | 0.000 | 0.000 | 0.000 |
| Pjvk | 0.000 | 0.001 | 0.002 |
| Gm14461 | 0.000 | 0.000 | 0.000 |
| Neurod1 | 0.000 | 0.000 | 0.000 |
| Prdx6b | 0.000 | 0.000 | 0.000 |

| | | | |
|---|---|---|---|
| Fsip2 | 0.000 | 0.000 | 0.000 |
| Btbd18 | 0.000 | 0.000 | 0.000 |
| Rtn4rl2 | 0.000 | 0.001 | 0.001 |
| Prg2 | 0.000 | 0.000 | 0.000 |
| Prg3 | 0.000 | 0.000 | 0.000 |
| Gm13716 | 0.000 | 0.000 | 0.000 |
| Olfr1014 | 0.000 | 0.000 | 0.000 |
| Olfr1023 | 0.000 | 0.000 | 0.000 |
| Olfr1028 | 0.000 | 0.000 | 0.000 |
| Olfr1029 | 0.000 | 0.000 | 0.000 |
| Olfr1030 | 0.000 | 0.000 | 0.000 |
| Olfr1036 | 0.000 | 0.000 | 0.000 |
| Olfr52 | 0.000 | 0.000 | 0.000 |
| Olfr1087 | 0.000 | 0.000 | 0.000 |
| Olfr1090 | 0.000 | 0.000 | 0.000 |
| Olfr1094 | 0.000 | 0.000 | 0.000 |
| Olfr1116 | 0.000 | 0.000 | 0.000 |
| Olfr1122 | 0.000 | 0.000 | 0.001 |
| Olfr1124 | 0.000 | 0.001 | 0.000 |
| Pramel7 | 0.000 | 0.000 | 0.000 |
| Olfr1129 | 0.000 | 0.000 | 0.000 |
| Olfr1137 | 0.000 | 0.000 | 0.000 |
| Olfr152 | 0.000 | 0.000 | 0.000 |
| Olfr1152 | 0.000 | 0.000 | 0.000 |
| Olfr1156 | 0.000 | 0.000 | 0.000 |
| Olfr1160 | 0.000 | 0.001 | 0.000 |
| Olfr73 | 0.000 | 0.000 | 0.000 |
| Olfr1173 | 0.000 | 0.000 | 0.000 |
| Olfr1175-ps | 0.000 | 0.000 | 0.000 |
| Olfr1186 | 0.000 | 0.000 | 0.000 |
| Olfr1191-ps1 | 0.000 | 0.000 | 0.000 |
| Olfr1192-ps1 | 0.000 | 0.000 | 0.000 |
| Olfr1202 | 0.000 | 0.000 | 0.000 |
| Olfr1205 | 0.000 | 0.000 | 0.000 |
| Olfr1209 | 0.000 | 0.000 | 0.000 |
| Olfr1215 | 0.000 | 0.000 | 0.000 |
| Olfr1217 | 0.000 | 0.000 | 0.000 |
| Olfr1221 | 0.000 | 0.000 | 0.000 |
| Olfr1226 | 0.000 | 0.000 | 0.000 |
| Olfr1247 | 0.000 | 0.000 | 0.000 |
| Olfr1256 | 0.000 | 0.000 | 0.000 |
| Olfr48 | 0.000 | 0.000 | 0.000 |
| Olfr1258 | 0.000 | 0.000 | 0.000 |
| Olfr1260 | 0.000 | 0.001 | 0.004 |
| Olfr1261 | 0.000 | 0.000 | 0.000 |
| Olfr1262 | 0.000 | 0.000 | 0.000 |
| Olfr1269 | 0.000 | 0.000 | 0.000 |
| Olfr1270 | 0.000 | 0.000 | 0.000 |

| | | | |
|---|---|---|---|
| Olfr1271 | 0.000 | 0.000 | 0.000 |
| Gm13769 | 0.000 | 0.000 | 0.000 |
| Olfr1274-ps | 0.000 | 0.000 | 0.000 |
| Chrm4 | 0.000 | 0.000 | 0.000 |
| 2900072N19I | 0.000 | 0.001 | 0.001 |
| Gm13814 | 0.000 | 0.000 | 0.000 |
| Gm13816 | 0.000 | 0.000 | 0.000 |
| Syt13 | 0.000 | 0.001 | 0.002 |
| Gm10803 | 0.000 | 0.000 | 0.000 |
| Alx4 | 0.000 | 0.000 | 0.000 |
| Accsl | 0.000 | 0.000 | 0.000 |
| Gm13920 | 0.000 | 0.000 | 0.000 |
| Gm13874 | 0.000 | 0.000 | 0.000 |
| Gm13869 | 0.000 | 0.000 | 0.000 |
| A930006I01R | 0.000 | 0.000 | 0.000 |
| BC016548 | 0.000 | 0.000 | 0.000 |
| Elf5 | 0.000 | 0.000 | 0.000 |
| Gm13881 | 0.000 | 0.000 | 0.000 |
| Gm10912 | 0.000 | 0.000 | 0.000 |
| D430041D05I | 0.000 | 0.000 | 0.000 |
| Pin1rt1 | 0.000 | 0.000 | 0.000 |
| Wt1os | 0.000 | 0.000 | 0.006 |
| Pax6os1 | 0.000 | 0.000 | 0.000 |
| Paupar | 0.000 | 0.000 | 0.000 |
| Gm14014 | 0.000 | 0.000 | 0.000 |
| Gm13936 | 0.000 | 0.000 | 0.000 |
| Slc5a12 | 0.000 | 0.000 | 0.000 |
| Muc15 | 0.000 | 0.000 | 0.000 |
| Olfr1277 | 0.000 | 0.000 | 0.000 |
| Olfr1279 | 0.000 | 0.000 | 0.000 |
| Olfr1281 | 0.000 | 0.000 | 0.000 |
| Olfr1284 | 0.000 | 0.000 | 0.000 |
| Olfr1291-ps1 | 0.000 | 0.000 | 0.000 |
| Olfr1305 | 0.000 | 0.000 | 0.000 |
| Olfr1307 | 0.000 | 0.001 | 0.000 |
| Olfr1309 | 0.000 | 0.000 | 0.000 |
| Olfr1313 | 0.000 | 0.000 | 0.000 |
| Olfr1314 | 0.000 | 0.000 | 0.000 |
| Olfr1315-ps1 | 0.000 | 0.000 | 0.000 |
| Olfr1316 | 0.000 | 0.000 | 0.000 |
| Olfr1318 | 0.000 | 0.000 | 0.000 |
| Nutm1 | 0.000 | 0.000 | 0.000 |
| Gm21985 | 0.000 | 0.000 | 0.000 |
| Chrm5 | 0.000 | 0.000 | 0.000 |
| Gm13963 | 0.000 | 0.000 | 0.000 |
| Gm13964 | 0.000 | 0.000 | 0.000 |
| Gjd2 | 0.000 | 0.000 | 0.000 |
| C130080G10I | 0.000 | 0.000 | 0.000 |

| | | | |
|---|---|---|---|
| Gm13974 | 0.000 | 0.000 | 0.000 |
| Gm28494 | 0.000 | 0.000 | 0.000 |
| Gm13977 | 0.000 | 0.000 | 0.000 |
| Gm13985 | 0.000 | 0.000 | 0.000 |
| 4930412B13F | 0.000 | 0.000 | 0.000 |
| Gm29233 | 0.000 | 0.000 | 0.000 |
| Fsip1 | 0.000 | 0.000 | 0.001 |
| Gpr176 | 0.000 | 0.003 | 0.004 |
| Ankrd63 | 0.000 | 0.000 | 0.000 |
| Phgr1 | 0.000 | 0.000 | 0.000 |
| Gm14091 | 0.000 | 0.000 | 0.000 |
| Ppp1r14d | 0.000 | 0.000 | 0.000 |
| Ltk | 0.000 | 0.001 | 0.000 |
| Pla2g4e | 0.000 | 0.007 | 0.002 |
| Pla2g4d | 0.000 | 0.000 | 0.000 |
| Gm26899 | 0.000 | 0.001 | 0.001 |
| Tgm7 | 0.000 | 0.000 | 0.000 |
| Gm14978 | 0.000 | 0.000 | 0.000 |
| Ckmt1 | 0.000 | 0.001 | 0.001 |
| Gm14019 | 0.000 | 0.001 | 0.001 |
| Patl2 | 0.000 | 0.000 | 0.000 |
| Duoxa2 | 0.000 | 0.000 | 0.000 |
| Duoxa1 | 0.000 | 0.000 | 0.003 |
| Duox1 | 0.000 | 0.000 | 0.002 |
| Slc28a2 | 0.000 | 0.000 | 0.000 |
| Gm14085 | 0.000 | 0.000 | 0.000 |
| Ctxn2 | 0.000 | 0.000 | 0.000 |
| Slc12a1 | 0.000 | 0.000 | 0.000 |
| Gm14002 | 0.000 | 0.001 | 0.005 |
| A530010F05F | 0.000 | 0.000 | 0.000 |
| Gm14003 | 0.000 | 0.000 | 0.000 |
| Fam227b | 0.000 | 0.000 | 0.000 |
| Gm17555 | 0.000 | 0.000 | 0.000 |
| Astl | 0.000 | 0.000 | 0.000 |
| Prom2 | 0.000 | 0.000 | 0.000 |
| Gm14009 | 0.000 | 0.000 | 0.000 |
| Gm14010 | 0.000 | 0.000 | 0.000 |
| Gm29010 | 0.000 | 0.000 | 0.000 |
| Spdye4c | 0.000 | 0.000 | 0.000 |
| Gm14011 | 0.000 | 0.000 | 0.000 |
| 4933427J07R | 0.000 | 0.000 | 0.000 |
| Gm14029 | 0.000 | 0.000 | 0.000 |
| A730036I17R | 0.000 | 0.000 | 0.000 |
| Il1bos | 0.000 | 0.000 | 0.000 |
| Gm28196 | 0.000 | 0.001 | 0.001 |
| Tgm6 | 0.000 | 0.000 | 0.000 |
| Tmc2 | 0.000 | 0.000 | 0.000 |
| 1700020A23F | 0.000 | 0.000 | 0.000 |

| | | | |
|---|---|---|---|
| 4930473A02F | 0.000 | 0.000 | 0.000 |
| Siglec1 | 0.000 | 0.002 | 0.001 |
| Adra1d | 0.000 | 0.002 | 0.000 |
| Gm14280 | 0.000 | 0.000 | 0.000 |
| 5330413P13F | 0.000 | 0.000 | 0.000 |
| 1700026D11I | 0.000 | 0.001 | 0.001 |
| AU019990 | 0.000 | 0.000 | 0.000 |
| Gm14097 | 0.000 | 0.000 | 0.000 |
| Gm14104 | 0.000 | 0.000 | 0.000 |
| Fermt1 | 0.000 | 0.000 | 0.000 |
| 4930545L23F | 0.000 | 0.000 | 0.001 |
| 9630028H03I | 0.000 | 0.001 | 0.000 |
| Platr3 | 0.000 | 0.000 | 0.000 |
| Gm14210 | 0.000 | 0.000 | 0.000 |
| Gm14064 | 0.000 | 0.000 | 0.000 |
| Gm14061 | 0.000 | 0.000 | 0.000 |
| Macrod2os2 | 0.000 | 0.000 | 0.000 |
| Macrod2os1 | 0.000 | 0.000 | 0.000 |
| Otor | 0.000 | 0.007 | 0.004 |
| Banf2 | 0.000 | 0.000 | 0.000 |
| Banf2os | 0.000 | 0.000 | 0.000 |
| Gm5535 | 0.000 | 0.000 | 0.001 |
| 4930444E06F | 0.000 | 0.000 | 0.000 |
| Ovol2 | 0.000 | 0.000 | 0.000 |
| Gm14092 | 0.000 | 0.000 | 0.000 |
| Cfap61 | 0.000 | 0.000 | 0.000 |
| Insm1 | 0.000 | 0.000 | 0.000 |
| A930019D19I | 0.000 | 0.000 | 0.000 |
| Nkx2-4 | 0.000 | 0.000 | 0.000 |
| Nkx2-2 | 0.000 | 0.000 | 0.000 |
| AI646519 | 0.000 | 0.003 | 0.003 |
| Pax1 | 0.000 | 0.005 | 0.009 |
| A530006G24I | 0.000 | 0.000 | 0.000 |
| 9030622O22I | 0.000 | 0.000 | 0.000 |
| Sstr4 | 0.000 | 0.000 | 0.001 |
| Gm14120 | 0.000 | 0.000 | 0.000 |
| Cst11 | 0.000 | 0.000 | 0.000 |
| Cst12 | 0.000 | 0.000 | 0.000 |
| Gm14133 | 0.000 | 0.000 | 0.000 |
| Gm14139 | 0.000 | 0.000 | 0.000 |
| Gm10130 | 0.000 | 0.000 | 0.000 |
| Cst7 | 0.000 | 0.000 | 0.000 |
| Vsx1 | 0.000 | 0.000 | 0.000 |
| Platr15 | 0.000 | 0.000 | 0.000 |
| Gm26681 | 0.000 | 0.000 | 0.000 |
| Gm14149 | 0.000 | 0.000 | 0.000 |
| Rad21l | 0.000 | 0.000 | 0.000 |
| Rspo4 | 0.000 | 0.000 | 0.001 |

| | | | |
|---|---|---|---|
| Rspo4os | 0.000 | 0.000 | 0.000 |
| AA387200 | 0.000 | 0.000 | 0.000 |
| Nrsn2 | 0.000 | 0.000 | 0.001 |
| 6820408C15I | 0.000 | 0.001 | 0.001 |
| Defb20 | 0.000 | 0.000 | 0.000 |
| Defb22 | 0.000 | 0.000 | 0.000 |
| Gm17416 | 0.000 | 0.000 | 0.000 |
| Defb19 | 0.000 | 0.000 | 0.000 |
| Defb45 | 0.000 | 0.000 | 0.000 |
| Defb36 | 0.000 | 0.000 | 0.000 |
| 1700030C14I | 0.000 | 0.000 | 0.000 |
| Dusp15 | 0.000 | 0.003 | 0.004 |
| Ttll9 | 0.000 | 0.000 | 0.000 |
| Xkr7 | 0.000 | 0.000 | 0.000 |
| 4930404H24I | 0.000 | 0.000 | 0.000 |
| Dnmt3bos | 0.000 | 0.000 | 0.000 |
| Bpifb6 | 0.000 | 0.000 | 0.000 |
| Bpifb4 | 0.000 | 0.000 | 0.000 |
| Bpifa2 | 0.000 | 0.000 | 0.000 |
| Actl10 | 0.000 | 0.000 | 0.000 |
| 4930519P11I | 0.000 | 0.000 | 0.000 |
| Gm14198 | 0.000 | 0.000 | 0.000 |
| Gm14214 | 0.000 | 0.000 | 0.000 |
| a | 0.000 | 0.000 | 0.001 |
| Gm45609 | 0.000 | 0.000 | 0.000 |
| 2310005A03I | 0.000 | 0.000 | 0.000 |
| Gssos2 | 0.000 | 0.000 | 0.000 |
| Gssos1 | 0.000 | 0.000 | 0.000 |
| Myh7b | 0.000 | 0.001 | 0.001 |
| Gm16098 | 0.000 | 0.001 | 0.001 |
| Fam83c | 0.000 | 0.000 | 0.000 |
| Gm15557 | 0.000 | 0.000 | 0.000 |
| Fer1l4 | 0.000 | 0.000 | 0.000 |
| Gm28036 | 0.000 | 0.001 | 0.000 |
| Gm14224 | 0.000 | 0.000 | 0.000 |
| Tldc2 | 0.000 | 0.000 | 0.000 |
| Ghrh | 0.000 | 0.000 | 0.000 |
| 2010009K17I | 0.000 | 0.000 | 0.000 |
| Bpi | 0.000 | 0.000 | 0.000 |
| Gm23925 | 0.000 | 0.000 | 0.000 |
| Snhg11 | 0.000 | 0.001 | 0.002 |
| Slc32a1 | 0.000 | 0.000 | 0.000 |
| Gm826 | 0.000 | 0.001 | 0.001 |
| Gm27206 | 0.000 | 0.001 | 0.000 |
| Emilin3 | 0.000 | 0.001 | 0.004 |
| Gm14228 | 0.000 | 0.000 | 0.000 |
| Gm14243 | 0.000 | 0.000 | 0.000 |
| Sgk2 | 0.000 | 0.000 | 0.000 |

| | | | |
|---|---|---|---|
| 2310001K24F | 0.000 | 0.000 | 0.001 |
| Gm16316 | 0.000 | 0.000 | 0.001 |
| Kcnk15 | 0.000 | 0.000 | 0.000 |
| Rims4 | 0.000 | 0.000 | 0.000 |
| Gm11455 | 0.000 | 0.000 | 0.000 |
| Kcns1 | 0.000 | 0.000 | 0.000 |
| Wfdc5 | 0.000 | 0.000 | 0.000 |
| Wfdc12 | 0.000 | 0.000 | 0.000 |
| Wfdc15a | 0.000 | 0.000 | 0.000 |
| Wfdc15b | 0.000 | 0.000 | 0.000 |
| Rbpjl | 0.000 | 0.006 | 0.001 |
| Gm14317 | 0.000 | 0.000 | 0.000 |
| Wfdc8 | 0.000 | 0.000 | 0.000 |
| Wfdc6b | 0.000 | 0.000 | 0.000 |
| Wfdc16 | 0.000 | 0.000 | 0.000 |
| Wfdc9 | 0.000 | 0.000 | 0.000 |
| Wfdc13 | 0.000 | 0.000 | 0.001 |
| Spint4 | 0.000 | 0.000 | 0.000 |
| Gm11457 | 0.000 | 0.000 | 0.001 |
| Spata25 | 0.000 | 0.000 | 0.000 |
| Cdh22 | 0.000 | 0.000 | 0.000 |
| Zfp663 | 0.000 | 0.000 | 0.000 |
| Ocstamp | 0.000 | 0.000 | 0.000 |
| Slc13a3 | 0.000 | 0.001 | 0.000 |
| Gm28163 | 0.000 | 0.000 | 0.000 |
| Platr29 | 0.000 | 0.000 | 0.000 |
| Gm11464 | 0.000 | 0.000 | 0.000 |
| Gm11468 | 0.000 | 0.000 | 0.000 |
| Gm14268 | 0.000 | 0.000 | 0.000 |
| Gm14320 | 0.000 | 0.001 | 0.001 |
| Gm14321 | 0.000 | 0.000 | 0.000 |
| Gm14319 | 0.000 | 0.001 | 0.001 |
| Kcng1 | 0.000 | 0.000 | 0.000 |
| Sall4 | 0.000 | 0.000 | 0.000 |
| Gm14258 | 0.000 | 0.000 | 0.000 |
| 9430093N23I | 0.000 | 0.000 | 0.000 |
| Gm14249 | 0.000 | 0.000 | 0.000 |
| A630075F10F | 0.000 | 0.000 | 0.000 |
| Gm17619 | 0.000 | 0.000 | 0.001 |
| Bcas1os2 | 0.000 | 0.000 | 0.000 |
| Cyp24a1 | 0.000 | 0.000 | 0.000 |
| Gm16796 | 0.000 | 0.000 | 0.000 |
| Gm14264 | 0.000 | 0.000 | 0.000 |
| Cbln4 | 0.000 | 0.000 | 0.000 |
| Mc3r | 0.000 | 0.000 | 0.000 |
| Gm14275 | 0.000 | 0.000 | 0.000 |
| Gm14273 | 0.000 | 0.000 | 0.000 |
| Gcnt7 | 0.000 | 0.000 | 0.000 |

| | | | |
|---|---|---|---|
| Spo11 | 0.000 | 0.000 | 0.000 |
| Ankrd60 | 0.000 | 0.000 | 0.000 |
| 1700010B08F | 0.000 | 0.000 | 0.000 |
| Gm10714 | 0.000 | 0.000 | 0.000 |
| Zfp831 | 0.000 | 0.000 | 0.000 |
| Edn3 | 0.000 | 0.001 | 0.008 |
| Gm14444 | 0.000 | 0.000 | 0.000 |
| Gm14393 | 0.000 | 0.002 | 0.001 |
| Gm14440 | 0.000 | 0.000 | 0.000 |
| Gm11007 | 0.000 | 0.000 | 0.000 |
| Gm14288 | 0.000 | 0.000 | 0.000 |
| Gm2026 | 0.000 | 0.001 | 0.002 |
| Gm2004 | 0.000 | 0.000 | 0.000 |
| 2210418O10I | 0.000 | 0.000 | 0.000 |
| Gm14408 | 0.000 | 0.000 | 0.000 |
| Gm14401 | 0.000 | 0.001 | 0.000 |
| Gm14409 | 0.000 | 0.001 | 0.001 |
| Gm14412 | 0.000 | 0.001 | 0.001 |
| Gm14406 | 0.000 | 0.000 | 0.000 |
| Hrh3 | 0.000 | 0.000 | 0.000 |
| Gata5os | 0.000 | 0.000 | 0.000 |
| Slco4a1 | 0.000 | 0.000 | 0.000 |
| Ntsr1 | 0.000 | 0.000 | 0.000 |
| Bhlhe23 | 0.000 | 0.000 | 0.000 |
| Gm14340 | 0.000 | 0.000 | 0.000 |
| Birc7 | 0.000 | 0.000 | 0.001 |
| Kcnq2 | 0.000 | 0.000 | 0.000 |
| Abhd16b | 0.000 | 0.000 | 0.000 |
| Gm29797 | 0.000 | 0.000 | 0.000 |
| Myt1 | 0.000 | 0.000 | 0.000 |
| AU022751 | 0.000 | 0.000 | 0.000 |
| Ccnb3 | 0.000 | 0.000 | 0.000 |
| Foxp3 | 0.000 | 0.000 | 0.000 |
| Cacna1f | 0.000 | 0.000 | 0.000 |
| Eras | 0.000 | 0.001 | 0.000 |
| Gata1 | 0.000 | 0.000 | 0.000 |
| Glod5 | 0.000 | 0.000 | 0.000 |
| Gm14820 | 0.000 | 0.000 | 0.000 |
| Ssxb9 | 0.000 | 0.000 | 0.000 |
| Gm5751 | 0.000 | 0.000 | 0.000 |
| Gm14862 | 0.000 | 0.001 | 0.000 |
| H2al1m | 0.000 | 0.001 | 0.000 |
| H2al1k | 0.000 | 0.000 | 0.000 |
| H2al3 | 0.000 | 0.000 | 0.000 |
| Gm14635 | 0.000 | 0.000 | 0.000 |
| Gpr34 | 0.000 | 0.001 | 0.001 |
| Drr1 | 0.000 | 0.000 | 0.000 |
| Spaca5 | 0.000 | 0.000 | 0.000 |

| | | | |
|---|---|---|---|
| Slc6a14 | 0.000 | 0.000 | 0.000 |
| Gm28268 | 0.000 | 0.000 | 0.000 |
| Gm4907 | 0.000 | 0.000 | 0.000 |
| Gm6268 | 0.000 | 0.000 | 0.000 |
| Gm14569 | 0.000 | 0.000 | 0.000 |
| Gm14549 | 0.000 | 0.000 | 0.000 |
| Gm15008 | 0.000 | 0.000 | 0.000 |
| Sowahd | 0.000 | 0.000 | 0.000 |
| Akap14 | 0.000 | 0.000 | 0.000 |
| Rhox2d | 0.000 | 0.000 | 0.000 |
| Rhox2g | 0.000 | 0.000 | 0.000 |
| Rhox8 | 0.000 | 0.001 | 0.000 |
| Rhox10 | 0.000 | 0.000 | 0.000 |
| Atp1b4 | 0.000 | 0.000 | 0.000 |
| Gm7598 | 0.000 | 0.003 | 0.002 |
| Gm14565 | 0.000 | 0.000 | 0.000 |
| Sh2d1a | 0.000 | 0.000 | 0.000 |
| Tenm1 | 0.000 | 0.000 | 0.005 |
| Dcaf12l2 | 0.000 | 0.002 | 0.006 |
| Prr32 | 0.000 | 0.000 | 0.000 |
| Xpnpep2 | 0.000 | 0.001 | 0.020 |
| 9530027J09R | 0.000 | 0.000 | 0.001 |
| Gpr119 | 0.000 | 0.000 | 0.000 |
| Gm595 | 0.000 | 0.000 | 0.000 |
| Arhgap36 | 0.000 | 0.002 | 0.006 |
| Igsf1 | 0.000 | 0.001 | 0.007 |
| Olfr1324 | 0.000 | 0.000 | 0.000 |
| Usp26 | 0.000 | 0.000 | 0.000 |
| 1700080O16I | 0.000 | 0.000 | 0.000 |
| Gm14582 | 0.000 | 0.000 | 0.000 |
| A630012P03F | 0.000 | 0.000 | 0.000 |
| Fam122c | 0.000 | 0.000 | 0.000 |
| Etd | 0.000 | 0.000 | 0.000 |
| Gm14597 | 0.000 | 0.000 | 0.000 |
| 4930502E18F | 0.000 | 0.000 | 0.000 |
| 1700013H16I | 0.000 | 0.000 | 0.000 |
| Zfp36l3 | 0.000 | 0.000 | 0.000 |
| Gm2155 | 0.000 | 0.000 | 0.000 |
| Gm2174 | 0.000 | 0.000 | 0.000 |
| Adgrg4 | 0.000 | 0.000 | 0.000 |
| Gm14718 | 0.000 | 0.000 | 0.000 |
| Cd40lg | 0.000 | 0.000 | 0.000 |
| Zic3 | 0.000 | 0.000 | 0.000 |
| F9 | 0.000 | 0.000 | 0.000 |
| Gm14662 | 0.000 | 0.000 | 0.000 |
| Gm14664 | 0.000 | 0.000 | 0.000 |
| C230004F18F | 0.000 | 0.015 | 0.017 |
| Cdr1 | 0.000 | 0.000 | 0.000 |

| | | | |
|---|---|---|---|
| Ldoc1 | 0.000 | 0.000 | 0.000 |
| 4933402E13F | 0.000 | 0.000 | 0.000 |
| Tslrn1 | 0.000 | 0.000 | 0.000 |
| 3830417A13F | 0.000 | 0.000 | 0.000 |
| 4930447F04F | 0.000 | 0.000 | 0.000 |
| 4933436I01R | 0.000 | 0.000 | 0.000 |
| Gm14705 | 0.000 | 0.001 | 0.001 |
| 1700111N16I | 0.000 | 0.000 | 0.000 |
| Gm16189 | 0.000 | 0.000 | 0.000 |
| Gpr50 | 0.000 | 0.000 | 0.001 |
| Fate1 | 0.000 | 0.000 | 0.001 |
| Cnga2 | 0.000 | 0.001 | 0.001 |
| Magea10 | 0.000 | 0.000 | 0.000 |
| Gabrq | 0.000 | 0.008 | 0.004 |
| Xlr5b | 0.000 | 0.000 | 0.000 |
| Xlr4c | 0.000 | 0.000 | 0.000 |
| Xlr3c | 0.000 | 0.000 | 0.000 |
| Xlr5c | 0.000 | 0.000 | 0.000 |
| Gm18336 | 0.000 | 0.000 | 0.000 |
| Atp2b3 | 0.000 | 0.000 | 0.000 |
| Avpr2 | 0.000 | 0.000 | 0.000 |
| Tktl1 | 0.000 | 0.001 | 0.000 |
| Gm6880 | 0.000 | 0.000 | 0.000 |
| Gm5640 | 0.000 | 0.000 | 0.000 |
| Gm5936 | 0.000 | 0.000 | 0.000 |
| Smim9 | 0.000 | 0.000 | 0.000 |
| Gm15384 | 0.000 | 0.000 | 0.000 |
| Cldn34b3 | 0.000 | 0.000 | 0.000 |
| Gm14742 | 0.000 | 0.000 | 0.000 |
| Tsga8 | 0.000 | 0.000 | 0.000 |
| Fthl17a | 0.000 | 0.000 | 0.000 |
| Gm14764 | 0.000 | 0.000 | 0.000 |
| Nr0b1 | 0.000 | 0.000 | 0.000 |
| Gm27000 | 0.000 | 0.000 | 0.001 |
| Mageb18 | 0.000 | 0.000 | 0.001 |
| Gm5941 | 0.000 | 0.000 | 0.000 |
| 1700003E24F | 0.000 | 0.000 | 0.000 |
| Arx | 0.000 | 0.000 | 0.000 |
| AU015836 | 0.000 | 0.000 | 0.000 |
| Gm14798 | 0.000 | 0.000 | 0.000 |
| Fam90a1b | 0.000 | 0.000 | 0.000 |
| Asb12 | 0.000 | 0.001 | 0.000 |
| Vsig4 | 0.000 | 0.000 | 0.000 |
| Gm14812 | 0.000 | 0.000 | 0.005 |
| Gm14809 | 0.000 | 0.000 | 0.000 |
| Gm14808 | 0.000 | 0.000 | 0.000 |
| Tmem28 | 0.000 | 0.000 | 0.001 |
| Dgat2l6 | 0.000 | 0.000 | 0.000 |

| | | | |
|---|---|---|---|
| Arr3 | 0.000 | 0.000 | 0.000 |
| Gm14902 | 0.000 | 0.005 | 0.000 |
| Tex11 | 0.000 | 0.000 | 0.000 |
| Gjb1 | 0.000 | 0.000 | 0.000 |
| Cxcr3 | 0.000 | 0.000 | 0.000 |
| 8030474K03F | 0.000 | 0.000 | 0.001 |
| Dmrtc1b | 0.000 | 0.000 | 0.000 |
| Gm26952 | 0.000 | 0.000 | 0.000 |
| Zcchc13 | 0.000 | 0.000 | 0.000 |
| 5330434G04I | 0.000 | 0.000 | 0.000 |
| Cypt2 | 0.000 | 0.000 | 0.000 |
| Gm5127 | 0.000 | 0.001 | 0.000 |
| Gpr174 | 0.000 | 0.000 | 0.000 |
| Fam46d | 0.000 | 0.000 | 0.000 |
| Gm379 | 0.000 | 0.000 | 0.000 |
| 4933403O08I | 0.000 | 0.000 | 0.000 |
| 2010106E10F | 0.000 | 0.000 | 0.000 |
| Pof1b | 0.000 | 0.000 | 0.000 |
| Dach2 | 0.000 | 0.001 | 0.002 |
| Gm14929 | 0.000 | 0.000 | 0.000 |
| Gm14951 | 0.000 | 0.000 | 0.000 |
| Gm14950 | 0.000 | 0.000 | 0.000 |
| Cldn34c3 | 0.000 | 0.000 | 0.000 |
| 4932411N23I | 0.000 | 0.000 | 0.000 |
| Nox1 | 0.000 | 0.001 | 0.002 |
| Taf7l | 0.000 | 0.000 | 0.000 |
| Btk | 0.000 | 0.001 | 0.001 |
| Nxf7 | 0.000 | 0.000 | 0.000 |
| Tcp11x2 | 0.000 | 0.000 | 0.000 |
| Tmsb15a | 0.000 | 0.000 | 0.000 |
| Nxf3 | 0.000 | 0.000 | 0.000 |
| Tceal7 | 0.000 | 0.000 | 0.000 |
| Rab9b | 0.000 | 0.001 | 0.001 |
| Esx1 | 0.000 | 0.000 | 0.000 |
| Il1rapl2 | 0.000 | 0.001 | 0.003 |
| 4930513O06I | 0.000 | 0.000 | 0.000 |
| 4933428M09 | 0.000 | 0.000 | 0.000 |
| Ripply1 | 0.000 | 0.000 | 0.000 |
| Pih1h3b | 0.000 | 0.000 | 0.000 |
| Gm15046 | 0.000 | 0.000 | 0.000 |
| Frmpd3 | 0.000 | 0.006 | 0.008 |
| Tex13b | 0.000 | 0.000 | 0.000 |
| Vsig1 | 0.000 | 0.001 | 0.000 |
| Irs4 | 0.000 | 0.009 | 0.008 |
| Gm15295 | 0.000 | 0.000 | 0.000 |
| Gm15294 | 0.000 | 0.000 | 0.000 |
| Gucy2f | 0.000 | 0.000 | 0.000 |
| Trpc5os | 0.000 | 0.000 | 0.000 |

| | | | |
|---|---|---|---|
| Lhfpl1 | 0.000 | 0.000 | 0.002 |
| Htr2c | 0.000 | 0.000 | 0.001 |
| Gm15104 | 0.000 | 0.000 | 0.000 |
| Gpr143 | 0.000 | 0.000 | 0.000 |
| Foxr2 | 0.000 | 0.000 | 0.000 |
| Magea5 | 0.000 | 0.000 | 0.000 |
| Cldn34b2 | 0.000 | 0.000 | 0.000 |
| Smpx | 0.000 | 0.003 | 0.003 |
| Klhl34 | 0.000 | 0.000 | 0.000 |
| Gm15241 | 0.000 | 0.000 | 0.000 |
| Rs1 | 0.000 | 0.000 | 0.000 |
| Scml1 | 0.000 | 0.000 | 0.000 |
| S100g | 0.000 | 0.000 | 0.001 |
| Rnf138rt1 | 0.000 | 0.001 | 0.000 |
| Tmem27 | 0.000 | 0.000 | 0.000 |
| Asb9 | 0.000 | 0.000 | 0.000 |
| Glra2 | 0.000 | 0.000 | 0.000 |
| Gm1720 | 0.000 | 0.000 | 0.000 |
| Gm8817 | 0.000 | 0.000 | 0.000 |
| Gm15228 | 0.000 | 0.000 | 0.000 |
| Tlr8 | 0.000 | 0.000 | 0.000 |
| Gm15239 | 0.000 | 0.000 | 0.000 |
| Frmpd4 | 0.000 | 0.001 | 0.002 |
| Amelx | 0.000 | 0.000 | 0.000 |
| Gm15245 | 0.000 | 0.000 | 0.000 |
| 4933400A11F | 0.000 | 0.001 | 0.001 |
| Gm15726 | 0.000 | 0.001 | 0.003 |
| Asmt | 0.000 | 0.000 | 0.000 |
| Hnf4g | 0.000 | 0.000 | 0.000 |
| 1110015O18I | 0.000 | 0.000 | 0.000 |
| Gm17308 | 0.000 | 0.000 | 0.001 |
| 2700069I18R | 0.000 | 0.000 | 0.000 |
| Gm37350 | 0.000 | 0.000 | 0.000 |
| C230057A21F | 0.000 | 0.001 | 0.001 |
| C030034L19F | 0.000 | 0.002 | 0.001 |
| Gm16337 | 0.000 | 0.000 | 0.000 |
| Pmp2 | 0.000 | 0.000 | 0.000 |
| Fabp9 | 0.000 | 0.000 | 0.000 |
| Gm38303 | 0.000 | 0.000 | 0.000 |
| Gm2464.1 | 0.000 | 0.000 | 0.000 |
| Sirpb1a | 0.000 | 0.000 | 0.000 |
| Sirpb1b | 0.000 | 0.000 | 0.000 |
| Gm30340 | 0.000 | 0.000 | 0.000 |
| Dnajc5b | 0.000 | 0.000 | 0.000 |
| Trim55 | 0.000 | 0.000 | 0.000 |
| Cpb1 | 0.000 | 0.010 | 0.019 |
| Gm31693 | 0.000 | 0.000 | 0.000 |
| A830092H15I | 0.000 | 0.002 | 0.001 |

| | | | |
|---|---|---|---|
| Ghsr | 0.000 | 0.000 | 0.000 |
| Gm42196 | 0.000 | 0.000 | 0.000 |
| 1700112D23I | 0.000 | 0.001 | 0.001 |
| Gm32950 | 0.000 | 0.000 | 0.000 |
| Gm33051 | 0.000 | 0.000 | 0.001 |
| Gm37867 | 0.000 | 0.000 | 0.000 |
| Egfem1 | 0.000 | 0.000 | 0.000 |
| Gm33206 | 0.000 | 0.000 | 0.000 |
| Actrt3 | 0.000 | 0.000 | 0.000 |
| Lrriq4 | 0.000 | 0.000 | 0.000 |
| Gm15462 | 0.000 | 0.001 | 0.000 |
| Gm15496 | 0.000 | 0.000 | 0.000 |
| Kcnmb3 | 0.000 | 0.000 | 0.001 |
| Gm37592 | 0.000 | 0.000 | 0.000 |
| Gm37640 | 0.000 | 0.000 | 0.000 |
| Gm37459 | 0.000 | 0.000 | 0.000 |
| 4930419G24I | 0.000 | 0.000 | 0.000 |
| Gm43666 | 0.000 | 0.000 | 0.000 |
| Gm20515 | 0.000 | 0.000 | 0.000 |
| Gm42695 | 0.000 | 0.000 | 0.000 |
| Gm21388 | 0.000 | 0.000 | 0.000 |
| 1700017M07 | 0.000 | 0.000 | 0.000 |
| Ccdc144b | 0.000 | 0.000 | 0.000 |
| Qrfpr | 0.000 | 0.000 | 0.001 |
| Trpc3 | 0.000 | 0.003 | 0.002 |
| Adad1 | 0.000 | 0.000 | 0.000 |
| Gm12534 | 0.000 | 0.000 | 0.000 |
| Fgf2os | 0.000 | 0.002 | 0.001 |
| Gm42921 | 0.000 | 0.000 | 0.000 |
| Gm20755 | 0.000 | 0.000 | 0.000 |
| Gm36823 | 0.000 | 0.000 | 0.000 |
| Gm43539 | 0.000 | 0.000 | 0.001 |
| 1700017G19I | 0.000 | 0.000 | 0.000 |
| Gm16508 | 0.000 | 0.000 | 0.000 |
| 2400006E01F | 0.000 | 0.000 | 0.000 |
| Gm31266 | 0.000 | 0.000 | 0.000 |
| Gm5103 | 0.000 | 0.001 | 0.000 |
| Mgarp | 0.000 | 0.001 | 0.001 |
| Gm37229 | 0.000 | 0.000 | 0.000 |
| Gm20089 | 0.000 | 0.001 | 0.000 |
| Gm37170 | 0.000 | 0.000 | 0.002 |
| Gm20402 | 0.000 | 0.000 | 0.000 |
| Gm30173 | 0.000 | 0.000 | 0.000 |
| Gm2447 | 0.000 | 0.000 | 0.000 |
| Gm30292 | 0.000 | 0.000 | 0.000 |
| Gm20750 | 0.000 | 0.000 | 0.000 |
| Gm43803 | 0.000 | 0.002 | 0.001 |
| Gm16206 | 0.000 | 0.000 | 0.000 |

| | | | |
|---|---|---|---|
| Stoml3 | 0.000 | 0.002 | 0.001 |
| C820005J03R | 0.000 | 0.000 | 0.000 |
| Trpc4 | 0.000 | 0.000 | 0.001 |
| Gm26678 | 0.000 | 0.000 | 0.000 |
| Gm21958 | 0.000 | 0.000 | 0.000 |
| Sertm1 | 0.000 | 0.000 | 0.000 |
| Sohlh2 | 0.000 | 0.000 | 0.000 |
| Gm42609 | 0.000 | 0.000 | 0.000 |
| Mab21l1 | 0.000 | 0.001 | 0.001 |
| Gm43376 | 0.000 | 0.000 | 0.000 |
| Gm40055 | 0.000 | 0.000 | 0.000 |
| Erich6 | 0.000 | 0.001 | 0.000 |
| 4930593A02F | 0.000 | 0.000 | 0.000 |
| Clrn1 | 0.000 | 0.000 | 0.000 |
| Gm43570 | 0.000 | 0.001 | 0.001 |
| Gpr171 | 0.000 | 0.001 | 0.000 |
| Gpr87 | 0.000 | 0.000 | 0.000 |
| Sucnr1 | 0.000 | 0.000 | 0.000 |
| Gm38326 | 0.000 | 0.000 | 0.000 |
| Gpr149 | 0.000 | 0.001 | 0.000 |
| Plch1 | 0.000 | 0.001 | 0.000 |
| Vmn2r1 | 0.000 | 0.001 | 0.000 |
| Vmn2r5 | 0.000 | 0.000 | 0.002 |
| Vmn2r6 | 0.000 | 0.000 | 0.000 |
| Gm37359 | 0.000 | 0.000 | 0.000 |
| Il12a | 0.000 | 0.001 | 0.001 |
| 4930535E02F | 0.000 | 0.000 | 0.000 |
| Gm1647 | 0.000 | 0.000 | 0.000 |
| Arl14 | 0.000 | 0.000 | 0.000 |
| Sptssb | 0.000 | 0.003 | 0.002 |
| Otol1 | 0.000 | 0.000 | 0.000 |
| Slitrk3 | 0.000 | 0.002 | 0.000 |
| Gm20754 | 0.000 | 0.004 | 0.000 |
| Gm38353 | 0.000 | 0.000 | 0.000 |
| 4930509J09R | 0.000 | 0.000 | 0.000 |
| Platr10 | 0.000 | 0.000 | 0.000 |
| Gm29133 | 0.000 | 0.000 | 0.000 |
| Gm37256 | 0.000 | 0.000 | 0.000 |
| Gm17359 | 0.000 | 0.000 | 0.000 |
| Rxfp1 | 0.000 | 0.000 | 0.000 |
| A330069K06F | 0.000 | 0.001 | 0.000 |
| Gria2 | 0.000 | 0.000 | 0.002 |
| Gm16000 | 0.000 | 0.000 | 0.000 |
| A830029E22F | 0.000 | 0.000 | 0.000 |
| Asic5 | 0.000 | 0.000 | 0.001 |
| Npy2r | 0.000 | 0.000 | 0.001 |
| Rbm46 | 0.000 | 0.005 | 0.027 |
| Rbm46os | 0.000 | 0.000 | 0.000 |

| | | | |
|---|---|---|---|
| Lrat | 0.000 | 0.000 | 0.001 |
| Gm30097 | 0.000 | 0.000 | 0.000 |
| Fgg | 0.000 | 0.001 | 0.000 |
| Fga | 0.000 | 0.000 | 0.000 |
| Fgb | 0.000 | 0.000 | 0.000 |
| Gm10710 | 0.000 | 0.001 | 0.010 |
| Gm38096 | 0.000 | 0.000 | 0.006 |
| 1700028M03 | 0.000 | 0.000 | 0.000 |
| 1700036G14I | 0.000 | 0.002 | 0.000 |
| Gm42812 | 0.000 | 0.000 | 0.000 |
| Prss48 | 0.000 | 0.000 | 0.000 |
| Gm37933 | 0.000 | 0.001 | 0.001 |
| Cd5l | 0.000 | 0.000 | 0.000 |
| Fcrl1 | 0.000 | 0.000 | 0.000 |
| Fcrl5 | 0.000 | 0.000 | 0.000 |
| Ntrk1 | 0.000 | 0.000 | 0.008 |
| Insrr | 0.000 | 0.000 | 0.001 |
| Sh2d2a | 0.000 | 0.000 | 0.000 |
| Hapln2 | 0.000 | 0.000 | 0.001 |
| AI849053 | 0.000 | 0.003 | 0.002 |
| Rhbg | 0.000 | 0.000 | 0.000 |
| Gm38392 | 0.000 | 0.000 | 0.000 |
| Bglap3 | 0.000 | 0.000 | 0.000 |
| Bglap2 | 0.000 | 0.000 | 0.001 |
| Bglap | 0.000 | 0.001 | 0.001 |
| Gm43738 | 0.000 | 0.000 | 0.000 |
| Pklr | 0.000 | 0.000 | 0.000 |
| Gm16069 | 0.000 | 0.000 | 0.000 |
| 4731419I09R | 0.000 | 0.001 | 0.001 |
| Dcst2 | 0.000 | 0.001 | 0.001 |
| Lenep | 0.000 | 0.000 | 0.000 |
| S100a2 | 0.000 | 0.000 | 0.000 |
| Gm43735 | 0.000 | 0.000 | 0.000 |
| Sprr1a | 0.000 | 0.000 | 0.001 |
| Ivl | 0.000 | 0.000 | 0.000 |
| Lce1d | 0.000 | 0.000 | 0.000 |
| Crct1 | 0.000 | 0.000 | 0.000 |
| Gm37245 | 0.000 | 0.000 | 0.000 |
| Them5 | 0.000 | 0.000 | 0.000 |
| Gm38411 | 0.000 | 0.000 | 0.000 |
| 1700040D17I | 0.000 | 0.000 | 0.000 |
| Oaz3 | 0.000 | 0.002 | 0.002 |
| Celf3 | 0.000 | 0.001 | 0.000 |
| Gm15234 | 0.000 | 0.000 | 0.001 |
| Tmod4 | 0.000 | 0.002 | 0.003 |
| Bnipl | 0.000 | 0.001 | 0.001 |
| Hormad1 | 0.000 | 0.000 | 0.000 |
| Gm15444 | 0.000 | 0.000 | 0.000 |

| | | | |
|---|---|---|---|
| Gm20628 | 0.000 | 0.001 | 0.001 |
| 4930477E14F | 0.000 | 0.000 | 0.000 |
| Gm10685 | 0.000 | 0.000 | 0.000 |
| Gm16253 | 0.000 | 0.000 | 0.002 |
| Ankrd34a | 0.000 | 0.016 | 0.008 |
| Cd160 | 0.000 | 0.000 | 0.000 |
| Gm31305 | 0.000 | 0.004 | 0.002 |
| Gm15999 | 0.000 | 0.000 | 0.000 |
| Reg4 | 0.000 | 0.000 | 0.000 |
| Gm4450 | 0.000 | 0.000 | 0.000 |
| Hsd3b5 | 0.000 | 0.000 | 0.000 |
| Hsd3b3 | 0.000 | 0.000 | 0.000 |
| Hsd3b6 | 0.000 | 0.000 | 0.001 |
| Hsd3b1 | 0.000 | 0.000 | 0.001 |
| Hao2 | 0.000 | 0.000 | 0.000 |
| 5730437C11F | 0.000 | 0.000 | 0.000 |
| Gm43464 | 0.000 | 0.000 | 0.000 |
| Cd101 | 0.000 | 0.001 | 0.001 |
| Gm42538 | 0.000 | 0.000 | 0.000 |
| Mab21l3 | 0.000 | 0.000 | 0.001 |
| Gm16160 | 0.000 | 0.000 | 0.000 |
| A230001M10 | 0.000 | 0.000 | 0.002 |
| Gm43242 | 0.000 | 0.000 | 0.000 |
| Gm42678 | 0.000 | 0.000 | 0.000 |
| Tshb | 0.000 | 0.001 | 0.000 |
| Gm42681 | 0.000 | 0.001 | 0.000 |
| Sycp1 | 0.000 | 0.000 | 0.001 |
| Nr1h5 | 0.000 | 0.000 | 0.000 |
| Syt6 | 0.000 | 0.000 | 0.000 |
| Atg4a-ps | 0.000 | 0.000 | 0.000 |
| Gm15886 | 0.000 | 0.000 | 0.000 |
| Bcl2l15 | 0.000 | 0.000 | 0.001 |
| Gm5546 | 0.000 | 0.000 | 0.000 |
| Gm29561 | 0.000 | 0.000 | 0.000 |
| Fam19a3 | 0.000 | 0.000 | 0.001 |
| Kcnd3os | 0.000 | 0.001 | 0.001 |
| Gm5547 | 0.000 | 0.000 | 0.001 |
| Adora3 | 0.000 | 0.000 | 0.000 |
| Tmigd3 | 0.000 | 0.000 | 0.000 |
| Chil3 | 0.000 | 0.000 | 0.000 |
| Chil6 | 0.000 | 0.000 | 0.000 |
| Cym | 0.000 | 0.001 | 0.000 |
| A630076J17R | 0.000 | 0.000 | 0.000 |
| Alx3 | 0.000 | 0.000 | 0.001 |
| Gm36211 | 0.000 | 0.000 | 0.000 |
| Eps8l3 | 0.000 | 0.000 | 0.000 |
| Gstm3 | 0.000 | 0.000 | 0.000 |
| Gm12498 | 0.000 | 0.000 | 0.000 |

| | | | |
|---|---|---|---|
| Gpr61 | 0.000 | 0.000 | 0.000 |
| Sypl2 | 0.000 | 0.001 | 0.000 |
| 1700010K24F | 0.000 | 0.000 | 0.000 |
| 1700013F07F | 0.000 | 0.000 | 0.000 |
| Fndc7 | 0.000 | 0.002 | 0.001 |
| Gm9857 | 0.000 | 0.000 | 0.001 |
| Gpr88 | 0.000 | 0.002 | 0.005 |
| Gm29151 | 0.000 | 0.000 | 0.000 |
| Gm26530 | 0.000 | 0.001 | 0.000 |
| Gm42457 | 0.000 | 0.000 | 0.000 |
| Gm9916 | 0.000 | 0.000 | 0.000 |
| 6530403H02I | 0.000 | 0.000 | 0.001 |
| A730020M07 | 0.000 | 0.000 | 0.004 |
| Gm43607 | 0.000 | 0.000 | 0.000 |
| Gm4610 | 0.000 | 0.000 | 0.000 |
| Gm42836 | 0.000 | 0.000 | 0.000 |
| Gm43569 | 0.000 | 0.000 | 0.000 |
| Fabp2 | 0.000 | 0.000 | 0.000 |
| Gm35065 | 0.000 | 0.003 | 0.004 |
| 1700003H04I | 0.000 | 0.000 | 0.000 |
| Gm42825 | 0.000 | 0.000 | 0.000 |
| Ndst4 | 0.000 | 0.000 | 0.000 |
| Gm15551 | 0.000 | 0.003 | 0.000 |
| Gm16958 | 0.000 | 0.000 | 0.009 |
| Neurog2 | 0.000 | 0.001 | 0.000 |
| Gm35585 | 0.000 | 0.000 | 0.000 |
| 9830132P13F | 0.000 | 0.000 | 0.000 |
| 6330410L21F | 0.000 | 0.000 | 0.000 |
| Lrit3 | 0.000 | 0.000 | 0.000 |
| Etnppl | 0.000 | 0.000 | 0.000 |
| Gm43522 | 0.000 | 0.000 | 0.000 |
| Gm36520 | 0.000 | 0.000 | 0.000 |
| 4930534D22I | 0.000 | 0.000 | 0.000 |
| Gm29865 | 0.000 | 0.000 | 0.000 |
| Gm42880 | 0.000 | 0.000 | 0.000 |
| Gimd1 | 0.000 | 0.000 | 0.000 |
| Gm29811 | 0.000 | 0.000 | 0.000 |
| Gm26820 | 0.000 | 0.000 | 0.000 |
| Gm31243 | 0.000 | 0.000 | 0.000 |
| Bank1 | 0.000 | 0.001 | 0.000 |
| Gm40155 | 0.000 | 0.001 | 0.001 |
| Gm5105 | 0.000 | 0.000 | 0.001 |
| 0610031O16I | 0.000 | 0.000 | 0.001 |
| Gm16559 | 0.000 | 0.000 | 0.000 |
| Stpg2 | 0.000 | 0.000 | 0.000 |
| Gbp2b | 0.000 | 0.000 | 0.000 |
| Gm15540 | 0.000 | 0.000 | 0.000 |
| Gm43614 | 0.000 | 0.000 | 0.000 |

| | | | |
|---|---|---|---|
| A830019L24F | 0.000 | 0.000 | 0.000 |
| Gm43616 | 0.000 | 0.000 | 0.000 |
| Gm33651 | 0.000 | 0.000 | 0.001 |
| Gm6260 | 0.000 | 0.000 | 0.000 |
| Gm42705 | 0.000 | 0.000 | 0.000 |
| Gm42706 | 0.000 | 0.000 | 0.000 |
| Gm42708 | 0.000 | 0.000 | 0.000 |
| Gm43447 | 0.000 | 0.000 | 0.000 |
| Gm43240 | 0.000 | 0.001 | 0.000 |
| Gm16233 | 0.000 | 0.000 | 0.000 |
| Clca1 | 0.000 | 0.000 | 0.000 |
| Clca2 | 0.000 | 0.004 | 0.003 |
| 9530034A14F | 0.000 | 0.001 | 0.001 |
| Gm35409 | 0.000 | 0.000 | 0.000 |
| Wdr63 | 0.000 | 0.001 | 0.004 |
| Mcoln3 | 0.000 | 0.000 | 0.001 |
| Gm10636 | 0.000 | 0.000 | 0.000 |
| Uox | 0.000 | 0.000 | 0.000 |
| Dnase2b | 0.000 | 0.000 | 0.000 |
| 4930503B20F | 0.000 | 0.000 | 0.000 |
| Gm10287 | 0.000 | 0.001 | 0.000 |
| Ifi44l | 0.000 | 0.000 | 0.000 |
| Asb17 | 0.000 | 0.000 | 0.000 |
| Gm43400 | 0.000 | 0.000 | 0.000 |
| Slc44a5 | 0.000 | 0.007 | 0.003 |
| Erich3 | 0.000 | 0.000 | 0.000 |
| Gm20389 | 0.000 | 0.000 | 0.000 |
| Tnni3k | 0.000 | 0.001 | 0.000 |
| Vmn1r3 | 0.000 | 0.000 | 0.000 |
| Gm26857 | 0.000 | 0.000 | 0.000 |
| Gm11782 | 0.000 | 0.000 | 0.000 |
| 4930423M02 | 0.000 | 0.000 | 0.000 |
| 8430436N08I | 0.000 | 0.000 | 0.005 |
| Gm26548 | 0.000 | 0.000 | 0.001 |
| Gm11816 | 0.000 | 0.000 | 0.001 |
| 1700123O12I | 0.000 | 0.000 | 0.000 |
| Gm11815 | 0.000 | 0.000 | 0.000 |
| Gm26663 | 0.000 | 0.000 | 0.000 |
| Esrp1 | 0.000 | 0.000 | 0.000 |
| Cdh17 | 0.000 | 0.003 | 0.001 |
| Gm10604 | 0.000 | 0.000 | 0.000 |
| Gm26895 | 0.000 | 0.000 | 0.000 |
| Gm11842 | 0.000 | 0.000 | 0.000 |
| Gm11823 | 0.000 | 0.000 | 0.000 |
| Lrrc69 | 0.000 | 0.000 | 0.000 |
| Necab1 | 0.000 | 0.001 | 0.006 |
| Calb1 | 0.000 | 0.001 | 0.001 |
| Gm11851 | 0.000 | 0.000 | 0.000 |

| | | | |
|---|---|---|---|
| Cnbd1 | 0.000 | 0.000 | 0.000 |
| Gm12353 | 0.000 | 0.000 | 0.000 |
| Slc7a13 | 0.000 | 0.000 | 0.000 |
| Gm11872 | 0.000 | 0.000 | 0.000 |
| Prdm13 | 0.000 | 0.000 | 0.000 |
| Gm30731 | 0.000 | 0.000 | 0.000 |
| Fhl5 | 0.000 | 0.000 | 0.001 |
| Fut9 | 0.000 | 0.000 | 0.004 |
| Gm11915 | 0.000 | 0.000 | 0.000 |
| Bach2os | 0.000 | 0.000 | 0.000 |
| Gja10 | 0.000 | 0.000 | 0.000 |
| Gabrr2 | 0.000 | 0.001 | 0.001 |
| Gabrr1 | 0.000 | 0.000 | 0.000 |
| Cga | 0.000 | 0.000 | 0.000 |
| Gm12364 | 0.000 | 0.000 | 0.000 |
| Lingo2 | 0.000 | 0.000 | 0.002 |
| Gm12371 | 0.000 | 0.000 | 0.000 |
| Gm12381 | 0.000 | 0.000 | 0.000 |
| Gm17094 | 0.000 | 0.000 | 0.000 |
| 1700009N14I | 0.000 | 0.000 | 0.000 |
| 4933428C19I | 0.000 | 0.000 | 0.000 |
| Spink4 | 0.000 | 0.000 | 0.000 |
| Aqp7 | 0.000 | 0.002 | 0.002 |
| Fam219aos | 0.000 | 0.000 | 0.000 |
| Gm12406 | 0.000 | 0.000 | 0.000 |
| Gm12405 | 0.000 | 0.000 | 0.000 |
| Arid3c | 0.000 | 0.000 | 0.000 |
| Gm13307 | 0.000 | 0.005 | 0.006 |
| Gm3892 | 0.000 | 0.000 | 0.000 |
| Gm13306 | 0.000 | 0.002 | 0.002 |
| Gm10593 | 0.000 | 0.000 | 0.000 |
| Ccl27b | 0.000 | 0.000 | 0.000 |
| 4930578G10I | 0.000 | 0.001 | 0.001 |
| Gm12395 | 0.000 | 0.001 | 0.001 |
| Gm12394 | 0.000 | 0.000 | 0.001 |
| Fam205c | 0.000 | 0.000 | 0.000 |
| Gm26643 | 0.000 | 0.000 | 0.001 |
| Sit1 | 0.000 | 0.002 | 0.001 |
| Gm12472 | 0.000 | 0.000 | 0.000 |
| Olfr71 | 0.000 | 0.000 | 0.000 |
| Olfr156 | 0.000 | 0.000 | 0.000 |
| Olfr157 | 0.000 | 0.000 | 0.000 |
| Ccin | 0.000 | 0.000 | 0.000 |
| Gm12678 | 0.000 | 0.000 | 0.000 |
| Gm12493 | 0.000 | 0.000 | 0.000 |
| Gm12679 | 0.000 | 0.000 | 0.000 |
| Gm12408 | 0.000 | 0.000 | 0.000 |
| Gm12410 | 0.000 | 0.000 | 0.000 |

| | | | |
|---|---|---|---|
| Stra6l | 0.000 | 0.000 | 0.000 |
| Gm12446 | 0.000 | 0.000 | 0.000 |
| Foxe1 | 0.000 | 0.000 | 0.001 |
| Hemgn | 0.000 | 0.000 | 0.000 |
| Gm568 | 0.000 | 0.001 | 0.001 |
| Acnat1 | 0.000 | 0.000 | 0.000 |
| Nipsnap3a | 0.000 | 0.000 | 0.000 |
| Gm12480 | 0.000 | 0.000 | 0.000 |
| Gm12511 | 0.000 | 0.000 | 0.001 |
| 2310081O03I | 0.000 | 0.000 | 0.000 |
| Gm26657 | 0.000 | 0.000 | 0.000 |
| Frrs1l | 0.000 | 0.000 | 0.001 |
| Gm12530 | 0.000 | 0.000 | 0.001 |
| Pakap | 0.000 | 0.004 | 0.001 |
| Txndc8 | 0.000 | 0.000 | 0.000 |
| Gm26889 | 0.000 | 0.000 | 0.000 |
| Mup5 | 0.000 | 0.000 | 0.000 |
| Rnf183 | 0.000 | 0.000 | 0.000 |
| Bspry | 0.000 | 0.000 | 0.001 |
| Ambp | 0.000 | 0.000 | 0.000 |
| Kif12 | 0.000 | 0.000 | 0.000 |
| Orm1 | 0.000 | 0.002 | 0.001 |
| Orm3 | 0.000 | 0.004 | 0.001 |
| Orm2 | 0.000 | 0.002 | 0.000 |
| Gm11214 | 0.000 | 0.000 | 0.000 |
| Tnfsf8 | 0.000 | 0.003 | 0.005 |
| Gm11217 | 0.000 | 0.000 | 0.000 |
| Gm11232 | 0.000 | 0.000 | 0.000 |
| Aldoart1 | 0.000 | 0.000 | 0.000 |
| Gm42303 | 0.000 | 0.000 | 0.000 |
| Gm11261 | 0.000 | 0.000 | 0.000 |
| Gm11269 | 0.000 | 0.000 | 0.000 |
| Cer1 | 0.000 | 0.000 | 0.001 |
| Gm12414 | 0.000 | 0.000 | 0.000 |
| Saxo1 | 0.000 | 0.000 | 0.000 |
| Ifnb1 | 0.000 | 0.000 | 0.000 |
| Ifnab | 0.000 | 0.000 | 0.000 |
| Gm13274 | 0.000 | 0.000 | 0.000 |
| Gm16686 | 0.000 | 0.000 | 0.000 |
| Gm13272 | 0.000 | 0.001 | 0.001 |
| Gm13276 | 0.000 | 0.000 | 0.001 |
| 4930553M12 | 0.000 | 0.000 | 0.000 |
| Wincr1 | 0.000 | 0.001 | 0.001 |
| Cdkn2a | 0.000 | 0.001 | 0.003 |
| Dmrta1 | 0.000 | 0.000 | 0.002 |
| Gm12629 | 0.000 | 0.000 | 0.000 |
| Gm12666 | 0.000 | 0.000 | 0.000 |
| Gm12648 | 0.000 | 0.000 | 0.000 |

| | | | |
|---|---|---|---|
| Lrrc19 | 0.000 | 0.000 | 0.001 |
| Gm12694 | 0.000 | 0.000 | 0.000 |
| 4930551L18R | 0.000 | 0.000 | 0.000 |
| Gm12708 | 0.000 | 0.000 | 0.000 |
| Gm29064 | 0.000 | 0.000 | 0.000 |
| Cyp2j13 | 0.000 | 0.000 | 0.000 |
| Gm12695 | 0.000 | 0.000 | 0.000 |
| L1td1 | 0.000 | 0.001 | 0.001 |
| Kank4os | 0.000 | 0.000 | 0.000 |
| I0C0044D17R | 0.000 | 0.000 | 0.000 |
| Angptl3 | 0.000 | 0.001 | 0.001 |
| Gm12705 | 0.000 | 0.000 | 0.000 |
| Gm12681 | 0.000 | 0.000 | 0.000 |
| Gm12688 | 0.000 | 0.001 | 0.001 |
| Ube2uos | 0.000 | 0.000 | 0.000 |
| Gm12798 | 0.000 | 0.000 | 0.000 |
| Dnajc6 | 0.000 | 0.000 | 0.000 |
| Gm12709 | 0.000 | 0.000 | 0.000 |
| BB031773 | 0.000 | 0.000 | 0.000 |
| Gm12718 | 0.000 | 0.000 | 0.000 |
| C8b | 0.000 | 0.000 | 0.000 |
| Fyb2 | 0.000 | 0.000 | 0.001 |
| Gm12728 | 0.000 | 0.001 | 0.001 |
| Pcsk9 | 0.000 | 0.001 | 0.007 |
| Bsnd | 0.000 | 0.000 | 0.000 |
| Tmem61 | 0.000 | 0.000 | 0.000 |
| 2700068H02I | 0.000 | 0.000 | 0.000 |
| Gm12744 | 0.000 | 0.000 | 0.000 |
| Ttc22 | 0.000 | 0.009 | 0.002 |
| Fam151a | 0.000 | 0.000 | 0.001 |
| Gm12746 | 0.000 | 0.000 | 0.000 |
| Gm12786 | 0.000 | 0.000 | 0.000 |
| Dio1 | 0.000 | 0.000 | 0.000 |
| Lrp8os2 | 0.000 | 0.000 | 0.000 |
| 4930407G08I | 0.000 | 0.000 | 0.000 |
| Zyg11a | 0.000 | 0.000 | 0.000 |
| Calr4 | 0.000 | 0.001 | 0.002 |
| Ttc39aos1 | 0.000 | 0.000 | 0.000 |
| 4930522H14I | 0.000 | 0.000 | 0.000 |
| Agbl4 | 0.000 | 0.001 | 0.000 |
| Skint8 | 0.000 | 0.000 | 0.000 |
| Gm12829 | 0.000 | 0.000 | 0.000 |
| 9130410C08R | 0.000 | 0.000 | 0.000 |
| Gm12830 | 0.000 | 0.000 | 0.000 |
| Cyp4a31 | 0.000 | 0.000 | 0.000 |
| Tex38 | 0.000 | 0.000 | 0.000 |
| Gm12847 | 0.000 | 0.000 | 0.000 |
| 6430628N08I | 0.000 | 0.000 | 0.000 |

| | | | |
|---|---|---|---|
| Dmbx1 | 0.000 | 0.000 | 0.000 |
| Gm12951 | 0.000 | 0.000 | 0.000 |
| 1700021J08R | 0.000 | 0.000 | 0.000 |
| Gm12841 | 0.000 | 0.001 | 0.000 |
| Cfap57 | 0.000 | 0.001 | 0.002 |
| Olfr1341 | 0.000 | 0.000 | 0.000 |
| Olfr1330 | 0.000 | 0.000 | 0.000 |
| Gm12868 | 0.000 | 0.000 | 0.000 |
| Tmem269 | 0.000 | 0.000 | 0.000 |
| Foxo6os | 0.000 | 0.001 | 0.001 |
| Slfnl1 | 0.000 | 0.000 | 0.000 |
| Gm12860 | 0.000 | 0.000 | 0.000 |
| Gm45579 | 0.000 | 0.000 | 0.000 |
| Tmco2 | 0.000 | 0.000 | 0.000 |
| Bmp8b | 0.000 | 0.000 | 0.000 |
| Oxct2b | 0.000 | 0.000 | 0.000 |
| Hpcal4 | 0.000 | 0.003 | 0.001 |
| Nt5c1a | 0.000 | 0.000 | 0.000 |
| Oxct2a | 0.000 | 0.000 | 0.000 |
| 1700121C08F | 0.000 | 0.000 | 0.000 |
| 4933427I04R | 0.000 | 0.000 | 0.000 |
| 4930535I16R | 0.000 | 0.000 | 0.000 |
| 1700057H15I | 0.000 | 0.000 | 0.000 |
| 4933407E24F | 0.000 | 0.000 | 0.000 |
| 3100002H09I | 0.000 | 0.000 | 0.000 |
| Maneal | 0.000 | 0.003 | 0.004 |
| Epha10 | 0.000 | 0.000 | 0.000 |
| Dnali1 | 0.000 | 0.000 | 0.000 |
| Grik3 | 0.000 | 0.000 | 0.000 |
| Gm12946 | 0.000 | 0.000 | 0.000 |
| Tfap2e | 0.000 | 0.001 | 0.000 |
| Gjb3 | 0.000 | 0.001 | 0.002 |
| Gjb4 | 0.000 | 0.000 | 0.001 |
| 4930471C06F | 0.000 | 0.000 | 0.000 |
| A630031M04 | 0.000 | 0.000 | 0.000 |
| Csmd2 | 0.000 | 0.000 | 0.001 |
| Tlr12 | 0.000 | 0.000 | 0.000 |
| 1700086P04F | 0.000 | 0.003 | 0.002 |
| Tmem54 | 0.000 | 0.000 | 0.000 |
| Hpca | 0.000 | 0.001 | 0.006 |
| Tssk3 | 0.000 | 0.000 | 0.000 |
| Gm12979 | 0.000 | 0.000 | 0.000 |
| E330017L17R | 0.000 | 0.000 | 0.001 |
| Spocd1 | 0.000 | 0.000 | 0.000 |
| 1700003M07 | 0.000 | 0.001 | 0.001 |
| Gm853 | 0.000 | 0.000 | 0.000 |
| Gm10570 | 0.000 | 0.001 | 0.000 |
| Nkain1 | 0.000 | 0.001 | 0.005 |

| | | | |
|---|---|---|---|
| Gm26516 | 0.000 | 0.000 | 0.000 |
| Gm12972 | 0.000 | 0.000 | 0.000 |
| Gm26716 | 0.000 | 0.000 | 0.000 |
| Matn1 | 0.000 | 0.000 | 0.000 |
| Gm831 | 0.000 | 0.000 | 0.000 |
| Gm12962 | 0.000 | 0.000 | 0.000 |
| Gm13063 | 0.000 | 0.000 | 0.000 |
| Themis2 | 0.000 | 0.001 | 0.001 |
| Cd164l2 | 0.000 | 0.005 | 0.002 |
| Gm13257 | 0.000 | 0.000 | 0.000 |
| Kdf1 | 0.000 | 0.000 | 0.000 |
| Nr0b2 | 0.000 | 0.000 | 0.000 |
| Lin28a | 0.000 | 0.000 | 0.000 |
| Crybg2 | 0.000 | 0.000 | 0.000 |
| Catsper4 | 0.000 | 0.000 | 0.000 |
| E130218I03Ri | 0.000 | 0.000 | 0.000 |
| Gm30191 | 0.000 | 0.000 | 0.000 |
| Slc30a2 | 0.000 | 0.000 | 0.000 |
| Extl1 | 0.000 | 0.001 | 0.000 |
| 1700021N21I | 0.000 | 0.000 | 0.000 |
| Gm16225 | 0.000 | 0.000 | 0.000 |
| Runx3 | 0.000 | 0.001 | 0.001 |
| Cnr2 | 0.000 | 0.000 | 0.000 |
| Gm13008 | 0.000 | 0.000 | 0.000 |
| Gm27022 | 0.000 | 0.001 | 0.001 |
| Htr1d | 0.000 | 0.003 | 0.019 |
| Tex46 | 0.000 | 0.000 | 0.000 |
| Lactbl1 | 0.000 | 0.000 | 0.000 |
| Epha8 | 0.000 | 0.000 | 0.000 |
| Cela3b | 0.000 | 0.000 | 0.000 |
| 1700013G24I | 0.000 | 0.001 | 0.000 |
| Ldlrad2 | 0.000 | 0.000 | 0.000 |
| Rap1gapos | 0.000 | 0.000 | 0.000 |
| Gm13012 | 0.000 | 0.000 | 0.000 |
| 1700095J12R | 0.000 | 0.000 | 0.000 |
| Fam43b | 0.000 | 0.001 | 0.000 |
| AB041806 | 0.000 | 0.000 | 0.000 |
| Vwa5b1 | 0.000 | 0.000 | 0.000 |
| Pla2g2d | 0.000 | 0.000 | 0.000 |
| Htr6 | 0.000 | 0.001 | 0.001 |
| Gm21969 | 0.000 | 0.000 | 0.000 |
| Tas1r2 | 0.000 | 0.000 | 0.000 |
| Klhdc7a | 0.000 | 0.000 | 0.000 |
| Igsf21 | 0.000 | 0.000 | 0.000 |
| Gm13016 | 0.000 | 0.000 | 0.000 |
| Gm13025 | 0.000 | 0.000 | 0.000 |
| Padi2 | 0.000 | 0.001 | 0.000 |
| Gm13031 | 0.000 | 0.001 | 0.002 |

| | | | |
|---|---|---|---|
| Gm13055 | 0.000 | 0.000 | 0.000 |
| Srarp | 0.000 | 0.000 | 0.000 |
| Agmat | 0.000 | 0.000 | 0.000 |
| Cela2a | 0.000 | 0.000 | 0.000 |
| Fhad1os2 | 0.000 | 0.000 | 0.000 |
| Gm13052 | 0.000 | 0.000 | 0.002 |
| Lrrc38 | 0.000 | 0.000 | 0.000 |
| Gm13102 | 0.000 | 0.000 | 0.000 |
| Gm13178 | 0.000 | 0.000 | 0.000 |
| Tnfrsf8 | 0.000 | 0.000 | 0.000 |
| Zfp990 | 0.000 | 0.000 | 0.000 |
| Zfp986 | 0.000 | 0.000 | 0.000 |
| Zfp987 | 0.000 | 0.000 | 0.000 |
| Gm26573 | 0.000 | 0.000 | 0.000 |
| Gm26880 | 0.000 | 0.000 | 0.000 |
| Zfp989 | 0.000 | 0.000 | 0.000 |
| Zfp988 | 0.000 | 0.000 | 0.000 |
| Gm16211 | 0.000 | 0.000 | 0.000 |
| Gm13201 | 0.000 | 0.000 | 0.000 |
| Draxin | 0.000 | 0.000 | 0.001 |
| Disp3 | 0.000 | 0.000 | 0.000 |
| Gm13206 | 0.000 | 0.000 | 0.000 |
| Gm13209 | 0.000 | 0.000 | 0.000 |
| Gm13203 | 0.000 | 0.001 | 0.001 |
| Cort | 0.000 | 0.000 | 0.001 |
| Ube4bos3 | 0.000 | 0.000 | 0.000 |
| Ube4bos2 | 0.000 | 0.000 | 0.000 |
| Ube4bos1 | 0.000 | 0.001 | 0.001 |
| Gm13066 | 0.000 | 0.000 | 0.000 |
| Gm13073 | 0.000 | 0.000 | 0.000 |
| Gm13070 | 0.000 | 0.000 | 0.000 |
| Slc2a7 | 0.000 | 0.000 | 0.000 |
| Slc45a1 | 0.000 | 0.000 | 0.001 |
| 1700045H11I | 0.000 | 0.000 | 0.000 |
| Gm13090 | 0.000 | 0.000 | 0.000 |
| 4930589P08F | 0.000 | 0.000 | 0.000 |
| Tas1r1 | 0.000 | 0.000 | 0.000 |
| Hes2 | 0.000 | 0.000 | 0.000 |
| Hes3 | 0.000 | 0.000 | 0.000 |
| Ajap1 | 0.000 | 0.002 | 0.002 |
| BC039966 | 0.000 | 0.000 | 0.000 |
| Ccdc27 | 0.000 | 0.000 | 0.001 |
| Trp73 | 0.000 | 0.000 | 0.000 |
| Gm27202 | 0.000 | 0.000 | 0.000 |
| Mmel1 | 0.000 | 0.000 | 0.000 |
| Hes5 | 0.000 | 0.001 | 0.001 |
| Gm27200 | 0.000 | 0.000 | 0.000 |
| Gabrd | 0.000 | 0.000 | 0.001 |

| | | | |
|---|---|---|---|
| Tmem52 | 0.000 | 0.000 | 0.004 |
| Gm10563 | 0.000 | 0.000 | 0.000 |
| Tmem88b | 0.000 | 0.001 | 0.001 |
| Ankrd65 | 0.000 | 0.000 | 0.000 |
| Mxra8os | 0.000 | 0.000 | 0.000 |
| Ttll10 | 0.000 | 0.000 | 0.000 |
| Rnf223 | 0.000 | 0.001 | 0.000 |
| Perm1 | 0.000 | 0.004 | 0.001 |
| Gm47252 | 0.000 | 0.000 | 0.000 |
| Gm36548 | 0.000 | 0.000 | 0.000 |
| Gm42435 | 0.000 | 0.000 | 0.000 |
| Gm43825 | 0.000 | 0.001 | 0.001 |
| Tmbim7 | 0.000 | 0.000 | 0.000 |
| Gm42744 | 0.000 | 0.000 | 0.000 |
| Steap1 | 0.000 | 0.006 | 0.006 |
| Zfp804b | 0.000 | 0.000 | 0.000 |
| Gm30835 | 0.000 | 0.000 | 0.000 |
| Gm15611 | 0.000 | 0.000 | 0.000 |
| Gm40264 | 0.000 | 0.000 | 0.000 |
| Gm15734 | 0.000 | 0.000 | 0.000 |
| Gm10482 | 0.000 | 0.000 | 0.000 |
| Gm9758 | 0.000 | 0.000 | 0.000 |
| Speer4e | 0.000 | 0.000 | 0.000 |
| Gm10354 | 0.000 | 0.000 | 0.000 |
| Gm17019 | 0.000 | 0.000 | 0.000 |
| Gm43391 | 0.000 | 0.000 | 0.000 |
| Gm21190 | 0.000 | 0.000 | 0.000 |
| Gm21083 | 0.000 | 0.000 | 0.000 |
| 4930572O03I | 0.000 | 0.000 | 0.000 |
| Speer4c | 0.000 | 0.000 | 0.000 |
| Gm3510 | 0.000 | 0.000 | 0.000 |
| Gm43000 | 0.000 | 0.000 | 0.000 |
| Gm28710 | 0.000 | 0.000 | 0.000 |
| Speer4f2 | 0.000 | 0.000 | 0.000 |
| Gnat3 | 0.000 | 0.000 | 0.000 |
| Gm31752 | 0.000 | 0.000 | 0.000 |
| Slc26a5 | 0.000 | 0.000 | 0.000 |
| Gm10475 | 0.000 | 0.000 | 0.000 |
| Gm43657 | 0.000 | 0.000 | 0.000 |
| A930003O13I | 0.000 | 0.000 | 0.000 |
| Lhfpl3 | 0.000 | 0.002 | 0.004 |
| 4930580E04F | 0.000 | 0.000 | 0.000 |
| Asic3 | 0.000 | 0.001 | 0.002 |
| Gbx1 | 0.000 | 0.000 | 0.000 |
| Asb10 | 0.000 | 0.002 | 0.003 |
| Prkag2os2 | 0.000 | 0.000 | 0.000 |
| Galntl5 | 0.000 | 0.000 | 0.000 |
| E130116L18R | 0.000 | 0.000 | 0.000 |

| | | | |
|---|---|---|---|
| Cct8l1 | 0.000 | 0.000 | 0.000 |
| Gm43972 | 0.000 | 0.000 | 0.000 |
| Gm21680 | 0.000 | 0.000 | 0.000 |
| Speer4a | 0.000 | 0.000 | 0.000 |
| Gm10471 | 0.000 | 0.000 | 0.000 |
| 5031410I06R | 0.000 | 0.000 | 0.000 |
| Gm7361 | 0.000 | 0.000 | 0.000 |
| Dpp6 | 0.000 | 0.000 | 0.002 |
| Gm26608 | 0.000 | 0.000 | 0.000 |
| En2 | 0.000 | 0.000 | 0.000 |
| Shh | 0.000 | 0.000 | 0.000 |
| 9530036O11I | 0.000 | 0.000 | 0.000 |
| Gm26894 | 0.000 | 0.000 | 0.000 |
| Gm42450 | 0.000 | 0.000 | 0.000 |
| A230098N10I | 0.000 | 0.000 | 0.001 |
| 3110082J24R | 0.000 | 0.000 | 0.000 |
| Adgrf3 | 0.000 | 0.004 | 0.003 |
| Otof | 0.000 | 0.001 | 0.001 |
| 1700001C02F | 0.000 | 0.000 | 0.000 |
| Cib4 | 0.000 | 0.001 | 0.000 |
| Gm42765 | 0.000 | 0.000 | 0.000 |
| Gm15461 | 0.000 | 0.000 | 0.000 |
| Slc30a3 | 0.000 | 0.000 | 0.001 |
| Ucn | 0.000 | 0.000 | 0.000 |
| Gckr | 0.000 | 0.000 | 0.000 |
| 4930566F21F | 0.000 | 0.000 | 0.000 |
| Gm7596 | 0.000 | 0.000 | 0.000 |
| Slc5a1 | 0.000 | 0.001 | 0.000 |
| Gm20465 | 0.000 | 0.000 | 0.000 |
| Gm42849 | 0.000 | 0.001 | 0.000 |
| Gm42848 | 0.000 | 0.000 | 0.000 |
| Cfap99 | 0.000 | 0.000 | 0.000 |
| Gm15522 | 0.000 | 0.000 | 0.000 |
| Gm42560 | 0.000 | 0.000 | 0.000 |
| Hmx1 | 0.000 | 0.000 | 0.000 |
| 4931431C16F | 0.000 | 0.000 | 0.000 |
| Psapl1 | 0.000 | 0.000 | 0.000 |
| Gm43701 | 0.000 | 0.000 | 0.000 |
| Ppp2r2c | 0.000 | 0.000 | 0.001 |
| Gm20052 | 0.000 | 0.000 | 0.000 |
| Otop1 | 0.000 | 0.000 | 0.000 |
| 4930487D11I | 0.000 | 0.000 | 0.000 |
| Clnk | 0.000 | 0.000 | 0.000 |
| Gm40293 | 0.000 | 0.000 | 0.000 |
| Gm42524 | 0.000 | 0.000 | 0.000 |
| Gm35191 | 0.000 | 0.000 | 0.000 |
| Nkx3-2 | 0.000 | 0.000 | 0.000 |
| Gm43699 | 0.000 | 0.000 | 0.000 |

| | | | |
|---|---|---|---|
| Gm43021 | 0.000 | 0.000 | 0.000 |
| Gm42555 | 0.000 | 0.000 | 0.000 |
| Gm42982 | 0.000 | 0.000 | 0.000 |
| Gm36017 | 0.000 | 0.000 | 0.000 |
| 4930431F12F | 0.000 | 0.000 | 0.000 |
| 9630001P10F | 0.000 | 0.004 | 0.004 |
| 4930518C09F | 0.000 | 0.000 | 0.000 |
| 4930448I18R | 0.000 | 0.000 | 0.000 |
| Gm43177 | 0.000 | 0.000 | 0.000 |
| 8030423F21F | 0.000 | 0.001 | 0.011 |
| Slc34a2 | 0.000 | 0.000 | 0.000 |
| Gm42745 | 0.000 | 0.000 | 0.000 |
| Gm10441 | 0.000 | 0.000 | 0.000 |
| Gm40304 | 0.000 | 0.000 | 0.000 |
| Cckar | 0.000 | 0.000 | 0.000 |
| Gm10440 | 0.000 | 0.000 | 0.001 |
| G6pd2 | 0.000 | 0.000 | 0.000 |
| Nwd2 | 0.000 | 0.005 | 0.005 |
| Gm15819 | 0.000 | 0.000 | 0.000 |
| 5830416I19R | 0.000 | 0.000 | 0.000 |
| Gm31363 | 0.000 | 0.000 | 0.000 |
| 1700027F09F | 0.000 | 0.000 | 0.000 |
| Gm26761 | 0.000 | 0.000 | 0.000 |
| Tlr1 | 0.000 | 0.000 | 0.001 |
| Klb | 0.000 | 0.000 | 0.000 |
| Chrna9 | 0.000 | 0.000 | 0.000 |
| 9130230L23F | 0.000 | 0.000 | 0.000 |
| Gm43698 | 0.000 | 0.000 | 0.000 |
| Gm15478 | 0.000 | 0.000 | 0.000 |
| Gm15477 | 0.000 | 0.000 | 0.000 |
| Yipf7 | 0.000 | 0.001 | 0.000 |
| Gabrg1 | 0.000 | 0.000 | 0.000 |
| Gabra4 | 0.000 | 0.021 | 0.009 |
| Cnga1 | 0.000 | 0.000 | 0.000 |
| 4933408A14F | 0.000 | 0.000 | 0.000 |
| Gm10135 | 0.000 | 0.000 | 0.000 |
| 1700071G01I | 0.000 | 0.000 | 0.000 |
| 1700025M24 | 0.000 | 0.000 | 0.000 |
| Cwh43 | 0.000 | 0.000 | 0.000 |
| Spata18 | 0.000 | 0.003 | 0.001 |
| Gm15984 | 0.000 | 0.000 | 0.000 |
| Gm42575 | 0.000 | 0.000 | 0.001 |
| Gm6116 | 0.000 | 0.001 | 0.001 |
| Gm42802 | 0.000 | 0.001 | 0.001 |
| Gm42800 | 0.000 | 0.000 | 0.000 |
| Gm28865 | 0.000 | 0.000 | 0.000 |
| Pdcl2 | 0.000 | 0.000 | 0.000 |
| Nmu | 0.000 | 0.001 | 0.000 |

| | | | |
|---|---|---|---|
| Arl9 | 0.000 | 0.001 | 0.001 |
| Thegl | 0.000 | 0.000 | 0.000 |
| Hopxos | 0.000 | 0.000 | 0.001 |
| Gm34648 | 0.000 | 0.000 | 0.000 |
| Gm16054 | 0.000 | 0.000 | 0.000 |
| Tecrl | 0.000 | 0.003 | 0.001 |
| Gm43532 | 0.000 | 0.000 | 0.000 |
| Gm43567 | 0.000 | 0.000 | 0.000 |
| Gnrhr | 0.000 | 0.001 | 0.010 |
| Tmprss11a | 0.000 | 0.000 | 0.000 |
| Tmprss11f | 0.000 | 0.000 | 0.000 |
| Gm43056 | 0.000 | 0.000 | 0.000 |
| Tmprss11e | 0.000 | 0.000 | 0.000 |
| Gm16579 | 0.000 | 0.000 | 0.000 |
| Gm43057 | 0.000 | 0.000 | 0.000 |
| Ugt2b1 | 0.000 | 0.000 | 0.000 |
| Ugt2b36 | 0.000 | 0.000 | 0.001 |
| Sult1d1 | 0.000 | 0.000 | 0.000 |
| Csn1s1 | 0.000 | 0.000 | 0.000 |
| Csn3 | 0.000 | 0.000 | 0.000 |
| 2310003L06F | 0.000 | 0.000 | 0.000 |
| Cabs1 | 0.000 | 0.000 | 0.000 |
| Smr3a | 0.000 | 0.000 | 0.001 |
| Amtn | 0.000 | 0.000 | 0.000 |
| Ambn | 0.000 | 0.000 | 0.000 |
| Gm42912 | 0.000 | 0.000 | 0.000 |
| Gc | 0.000 | 0.000 | 0.000 |
| Npffr2 | 0.000 | 0.000 | 0.000 |
| Afp | 0.000 | 0.000 | 0.000 |
| Afm | 0.000 | 0.000 | 0.000 |
| 5830473C10F | 0.000 | 0.000 | 0.000 |
| Gm19619 | 0.000 | 0.000 | 0.000 |
| Gm43688 | 0.000 | 0.000 | 0.000 |
| Ppef2 | 0.000 | 0.001 | 0.002 |
| Gm43599 | 0.000 | 0.000 | 0.000 |
| Cxcl9 | 0.000 | 0.001 | 0.006 |
| Fam47e | 0.000 | 0.005 | 0.002 |
| 4932430I15R | 0.000 | 0.001 | 0.000 |
| Sowahb | 0.000 | 0.000 | 0.000 |
| AA792892 | 0.000 | 0.000 | 0.000 |
| Gm3183 | 0.000 | 0.000 | 0.000 |
| Gm33050 | 0.000 | 0.000 | 0.000 |
| Gm33609 | 0.000 | 0.000 | 0.000 |
| Gm2861 | 0.000 | 0.000 | 0.000 |
| Naa11 | 0.000 | 0.000 | 0.000 |
| Prdm8 | 0.000 | 0.006 | 0.011 |
| A730035I17R | 0.000 | 0.000 | 0.000 |
| Fgf5 | 0.000 | 0.001 | 0.005 |

| | | | |
|---|---|---|---|
| 1700010H22I | 0.000 | 0.000 | 0.000 |
| Tmem150cos | 0.000 | 0.000 | 0.000 |
| Gm15777 | 0.000 | 0.000 | 0.000 |
| 4930429D17I | 0.000 | 0.000 | 0.000 |
| Gm42620 | 0.000 | 0.000 | 0.000 |
| 1700016H13I | 0.000 | 0.000 | 0.000 |
| Gm26703 | 0.000 | 0.001 | 0.000 |
| Dspp | 0.000 | 0.001 | 0.000 |
| Dmp1 | 0.000 | 0.012 | 0.001 |
| Gm26519 | 0.000 | 0.000 | 0.000 |
| Lrrc8dos | 0.000 | 0.000 | 0.000 |
| Barhl2 | 0.000 | 0.000 | 0.000 |
| Gm26872 | 0.000 | 0.000 | 0.000 |
| Hfm1 | 0.000 | 0.000 | 0.001 |
| Gm33474 | 0.000 | 0.001 | 0.003 |
| Lpcat2b | 0.000 | 0.000 | 0.000 |
| Gm42669 | 0.000 | 0.001 | 0.001 |
| A930041C12I | 0.000 | 0.000 | 0.000 |
| Gm26692 | 0.000 | 0.000 | 0.000 |
| 4930428O21I | 0.000 | 0.000 | 0.000 |
| Gfi1 | 0.000 | 0.001 | 0.000 |
| Gm42518 | 0.000 | 0.000 | 0.000 |
| Pde6b | 0.000 | 0.000 | 0.000 |
| Cplx1 | 0.000 | 0.000 | 0.000 |
| Vmn2r10 | 0.000 | 0.000 | 0.000 |
| Vmn2r12 | 0.000 | 0.000 | 0.000 |
| Gm26718 | 0.000 | 0.001 | 0.001 |
| Gm15788 | 0.000 | 0.000 | 0.000 |
| P2rx2 | 0.000 | 0.000 | 0.001 |
| Lrcol1 | 0.000 | 0.000 | 0.000 |
| A630023P12I | 0.000 | 0.000 | 0.000 |
| Galnt9 | 0.000 | 0.000 | 0.000 |
| Gm42489 | 0.000 | 0.000 | 0.000 |
| Gm36535 | 0.000 | 0.000 | 0.000 |
| Gm42488 | 0.000 | 0.000 | 0.001 |
| 1700028D13I | 0.000 | 0.000 | 0.000 |
| Gm6583 | 0.000 | 0.000 | 0.000 |
| Gm6588 | 0.000 | 0.000 | 0.000 |
| Gm27004 | 0.000 | 0.000 | 0.000 |
| Crybb2 | 0.000 | 0.000 | 0.000 |
| Gm42161 | 0.000 | 0.000 | 0.000 |
| Gm15736 | 0.000 | 0.000 | 0.000 |
| Gm17122 | 0.000 | 0.000 | 0.000 |
| Gm42797 | 0.000 | 0.000 | 0.000 |
| Dao | 0.000 | 0.001 | 0.001 |
| Svop | 0.000 | 0.000 | 0.000 |
| Foxn4 | 0.000 | 0.000 | 0.000 |
| 4930519G04I | 0.000 | 0.000 | 0.000 |

| | | | |
|---|---|---|---|
| Gm9936 | 0.000 | 0.000 | 0.001 |
| Gm13822 | 0.000 | 0.000 | 0.000 |
| Hnf1aos1 | 0.000 | 0.000 | 0.000 |
| 4930401G09I | 0.000 | 0.000 | 0.000 |
| Gm13832 | 0.000 | 0.000 | 0.000 |
| Tmem233 | 0.000 | 0.000 | 0.000 |
| C330018A13F | 0.000 | 0.000 | 0.001 |
| Gm13837 | 0.000 | 0.000 | 0.000 |
| Gm13838 | 0.000 | 0.001 | 0.001 |
| 4930562A09F | 0.000 | 0.001 | 0.000 |
| Srrm4os | 0.000 | 0.003 | 0.001 |
| Gm43122 | 0.000 | 0.000 | 0.000 |
| BC051077 | 0.000 | 0.002 | 0.002 |
| Gm7538 | 0.000 | 0.000 | 0.000 |
| Gm16063 | 0.000 | 0.000 | 0.000 |
| Gm16064 | 0.000 | 0.000 | 0.000 |
| 1700021F13F | 0.000 | 0.001 | 0.001 |
| Gm43269 | 0.000 | 0.000 | 0.000 |
| Gm38554 | 0.000 | 0.000 | 0.000 |
| Gm10390 | 0.000 | 0.000 | 0.000 |
| Gm42654 | 0.000 | 0.000 | 0.000 |
| Lhx5 | 0.000 | 0.000 | 0.000 |
| Gm43579 | 0.000 | 0.000 | 0.000 |
| Gm43069 | 0.000 | 0.000 | 0.000 |
| Oas1e | 0.000 | 0.000 | 0.000 |
| Oas1g | 0.000 | 0.000 | 0.001 |
| Rph3a | 0.000 | 0.001 | 0.000 |
| Gm42917 | 0.000 | 0.000 | 0.000 |
| Adam1b | 0.000 | 0.001 | 0.001 |
| 1700008B11F | 0.000 | 0.000 | 0.000 |
| Gm3970 | 0.000 | 0.000 | 0.000 |
| Gm15637 | 0.000 | 0.000 | 0.000 |
| Gm26745 | 0.000 | 0.000 | 0.000 |
| Hpd | 0.000 | 0.000 | 0.000 |
| Il31 | 0.000 | 0.000 | 0.000 |
| Lrrc43 | 0.000 | 0.000 | 0.000 |
| Gm49027 | 0.000 | 0.000 | 0.000 |
| Gm34086 | 0.000 | 0.000 | 0.000 |
| Pitpnm2os1 | 0.000 | 0.000 | 0.000 |
| 4930404A12F | 0.000 | 0.000 | 0.000 |
| Gm42838 | 0.000 | 0.000 | 0.000 |
| Gm42839 | 0.000 | 0.000 | 0.000 |
| 1700048F04F | 0.000 | 0.000 | 0.000 |
| Gm42980 | 0.000 | 0.000 | 0.000 |
| Glt1d1 | 0.000 | 0.000 | 0.000 |
| Tmem132d | 0.000 | 0.000 | 0.000 |
| Gm42817 | 0.000 | 0.000 | 0.000 |
| Piwil1 | 0.000 | 0.000 | 0.000 |

| | | | |
|---|---|---|---|
| Rimbp2 | 0.000 | 0.000 | 0.000 |
| Gm42875 | 0.000 | 0.000 | 0.000 |
| Caln1 | 0.000 | 0.005 | 0.003 |
| 2810432F15F | 0.000 | 0.000 | 0.000 |
| 4930563F08F | 0.000 | 0.000 | 0.000 |
| Gm42439 | 0.000 | 0.000 | 0.000 |
| Gm28616 | 0.000 | 0.000 | 0.000 |
| Gm42626 | 0.000 | 0.000 | 0.000 |
| Gm15627 | 0.000 | 0.002 | 0.001 |
| Gm42882 | 0.000 | 0.000 | 0.000 |
| Gm42884 | 0.000 | 0.000 | 0.000 |
| Syna | 0.000 | 0.000 | 0.000 |
| Cldn13 | 0.000 | 0.000 | 0.000 |
| Cldn4 | 0.000 | 0.001 | 0.000 |
| Gm43091 | 0.000 | 0.000 | 0.000 |
| Zp3 | 0.000 | 0.001 | 0.004 |
| Gm16599 | 0.000 | 0.000 | 0.000 |
| Myl10 | 0.000 | 0.000 | 0.000 |
| Vgf | 0.000 | 0.001 | 0.001 |
| Gm42456 | 0.000 | 0.000 | 0.000 |
| Gm17112 | 0.000 | 0.000 | 0.000 |
| Actl6b | 0.000 | 0.000 | 0.000 |
| Tfr2 | 0.000 | 0.007 | 0.011 |
| Gm15498 | 0.000 | 0.000 | 0.001 |
| Azgp1 | 0.000 | 0.000 | 0.000 |
| Nxpe5 | 0.000 | 0.000 | 0.001 |
| Gal3st4 | 0.000 | 0.002 | 0.005 |
| Gm43720 | 0.000 | 0.000 | 0.000 |
| Pvrig | 0.000 | 0.000 | 0.000 |
| Gm5294 | 0.000 | 0.000 | 0.000 |
| Uncx | 0.000 | 0.000 | 0.000 |
| Gm16121 | 0.000 | 0.000 | 0.000 |
| Gm4869 | 0.000 | 0.000 | 0.000 |
| Grifin | 0.000 | 0.000 | 0.001 |
| Card11 | 0.000 | 0.000 | 0.000 |
| Gm16036 | 0.000 | 0.000 | 0.000 |
| Spdye4b | 0.000 | 0.000 | 0.000 |
| Spdye4a | 0.000 | 0.000 | 0.000 |
| Gm17135 | 0.000 | 0.000 | 0.000 |
| Gm42421 | 0.000 | 0.001 | 0.001 |
| Bhlha15 | 0.000 | 0.001 | 0.000 |
| Tmem130 | 0.000 | 0.000 | 0.000 |
| Kpna7 | 0.000 | 0.000 | 0.000 |
| Cyp3a57 | 0.000 | 0.000 | 0.000 |
| 1700001J03R | 0.000 | 0.001 | 0.001 |
| 1700041I07R | 0.000 | 0.000 | 0.000 |
| Gsx1 | 0.000 | 0.000 | 0.000 |
| Flt3 | 0.000 | 0.000 | 0.000 |

| | | | |
|---|---|---|---|
| Gm26597 | 0.000 | 0.000 | 0.000 |
| Gm43156 | 0.000 | 0.000 | 0.000 |
| Gm36186 | 0.000 | 0.000 | 0.000 |
| 2210417A02F | 0.000 | 0.000 | 0.000 |
| Gm15410 | 0.000 | 0.002 | 0.000 |
| Gm15409 | 0.000 | 0.000 | 0.000 |
| 5730422E09F | 0.000 | 0.001 | 0.001 |
| Gm42788 | 0.000 | 0.004 | 0.005 |
| Gm43332 | 0.000 | 0.000 | 0.000 |
| Gm29264 | 0.000 | 0.000 | 0.000 |
| Gm19719 | 0.000 | 0.001 | 0.001 |
| BC028471 | 0.000 | 0.000 | 0.005 |
| Gm43151 | 0.000 | 0.000 | 0.000 |
| Gm43150 | 0.000 | 0.000 | 0.000 |
| Gm20005 | 0.000 | 0.000 | 0.000 |
| Rxfp2 | 0.000 | 0.000 | 0.000 |
| Gm42556 | 0.000 | 0.000 | 0.000 |
| Gm5 | 0.000 | 0.000 | 0.000 |
| Gm42529 | 0.000 | 0.001 | 0.000 |
| Gm43690 | 0.000 | 0.000 | 0.000 |
| Gm36447 | 0.000 | 0.000 | 0.000 |
| Gm42906 | 0.000 | 0.000 | 0.000 |
| Vmn2r18 | 0.000 | 0.000 | 0.000 |
| Gm43129 | 0.000 | 0.000 | 0.000 |
| Gm8579 | 0.000 | 0.001 | 0.000 |
| Hepacam2 | 0.000 | 0.000 | 0.000 |
| Calcr | 0.000 | 0.000 | 0.000 |
| Gngt1 | 0.000 | 0.000 | 0.000 |
| 4930528H21I | 0.000 | 0.000 | 0.000 |
| 1700019G24I | 0.000 | 0.000 | 0.000 |
| Gm20618 | 0.000 | 0.001 | 0.000 |
| Gm26774 | 0.000 | 0.000 | 0.000 |
| Gm19272 | 0.000 | 0.000 | 0.000 |
| Gm16055 | 0.000 | 0.000 | 0.000 |
| Gm44129 | 0.000 | 0.000 | 0.000 |
| Vwde | 0.000 | 0.000 | 0.000 |
| Tfec | 0.000 | 0.000 | 0.001 |
| Gm15473 | 0.000 | 0.001 | 0.000 |
| Gm15581 | 0.000 | 0.000 | 0.000 |
| Ankrd7 | 0.000 | 0.000 | 0.000 |
| Kcnd2 | 0.000 | 0.004 | 0.002 |
| Gm3289 | 0.000 | 0.000 | 0.000 |
| Gm26719 | 0.000 | 0.002 | 0.003 |
| Aass | 0.000 | 0.008 | 0.008 |
| Rnf148 | 0.000 | 0.000 | 0.000 |
| Slc13a1 | 0.000 | 0.000 | 0.000 |
| Asb15 | 0.000 | 0.000 | 0.000 |
| Lmod2 | 0.000 | 0.001 | 0.001 |

| | | | |
|---|---|---|---|
| Fscn3 | 0.000 | 0.001 | 0.001 |
| Lep | 0.000 | 0.000 | 0.000 |
| Fam71f1 | 0.000 | 0.000 | 0.000 |
| Gm26627 | 0.000 | 0.000 | 0.002 |
| 1700095J07R | 0.000 | 0.000 | 0.000 |
| Ssmem1 | 0.000 | 0.000 | 0.000 |
| Cpa4 | 0.000 | 0.000 | 0.002 |
| Gm13781 | 0.000 | 0.000 | 0.000 |
| Cpa5 | 0.000 | 0.000 | 0.000 |
| 4930412F09R | 0.000 | 0.000 | 0.000 |
| Tsga13 | 0.000 | 0.001 | 0.005 |
| Gm27019 | 0.000 | 0.000 | 0.000 |
| Gm6117 | 0.000 | 0.000 | 0.000 |
| Gm13848 | 0.000 | 0.001 | 0.002 |
| Plxna4os2 | 0.000 | 0.000 | 0.000 |
| BC049739 | 0.000 | 0.000 | 0.000 |
| Gm14540 | 0.000 | 0.000 | 0.000 |
| Gm13853 | 0.000 | 0.000 | 0.000 |
| Gm13856 | 0.000 | 0.000 | 0.000 |
| Gm14546 | 0.000 | 0.000 | 0.000 |
| Npn2 | 0.000 | 0.000 | 0.000 |
| Gm14547 | 0.000 | 0.000 | 0.000 |
| Stra8 | 0.000 | 0.000 | 0.000 |
| Slc13a4 | 0.000 | 0.001 | 0.001 |
| Svopl | 0.000 | 0.001 | 0.010 |
| Tmem213 | 0.000 | 0.000 | 0.000 |
| 1700025N23I | 0.000 | 0.000 | 0.000 |
| Gm16272 | 0.000 | 0.000 | 0.000 |
| Gm26807 | 0.000 | 0.000 | 0.000 |
| Wee2 | 0.000 | 0.000 | 0.000 |
| Olfr461 | 0.000 | 0.000 | 0.000 |
| Mgam | 0.000 | 0.000 | 0.001 |
| Gm4744 | 0.000 | 0.000 | 0.000 |
| 2210010C04I | 0.000 | 0.000 | 0.000 |
| 5830405F06R | 0.000 | 0.000 | 0.000 |
| Trbv16 | 0.000 | 0.000 | 0.000 |
| Trbv29 | 0.000 | 0.000 | 0.000 |
| Prss2 | 0.000 | 0.000 | 0.000 |
| Trbc1 | 0.000 | 0.000 | 0.000 |
| Trpv6 | 0.000 | 0.000 | 0.000 |
| Trpv5 | 0.000 | 0.000 | 0.000 |
| Kel | 0.000 | 0.000 | 0.000 |
| Pip | 0.000 | 0.000 | 0.000 |
| Sva | 0.000 | 0.000 | 0.000 |
| Tmem139 | 0.000 | 0.000 | 0.000 |
| Gm44284 | 0.000 | 0.000 | 0.000 |
| Tas2r143 | 0.000 | 0.000 | 0.001 |
| Tas2r135 | 0.000 | 0.000 | 0.000 |

| | | | |
|---|---|---|---|
| Tas2r126 | 0.000 | 0.000 | 0.000 |
| Olfr458 | 0.000 | 0.000 | 0.000 |
| Olfr457 | 0.000 | 0.000 | 0.000 |
| Olfr456 | 0.000 | 0.000 | 0.000 |
| Tcaf3 | 0.000 | 0.000 | 0.000 |
| Olfr453 | 0.000 | 0.000 | 0.000 |
| Olfr452 | 0.000 | 0.000 | 0.000 |
| Nobox | 0.000 | 0.000 | 0.000 |
| 1700026J14R | 0.000 | 0.000 | 0.000 |
| Gm5111 | 0.000 | 0.000 | 0.001 |
| Gimap7 | 0.000 | 0.000 | 0.000 |
| Aoc1 | 0.000 | 0.000 | 0.000 |
| 1600015I10R | 0.000 | 0.000 | 0.000 |
| Stk31 | 0.000 | 0.001 | 0.000 |
| Gm3455 | 0.000 | 0.000 | 0.000 |
| G930045G22 | 0.000 | 0.000 | 0.000 |
| Gm32479 | 0.000 | 0.000 | 0.000 |
| Gm6559 | 0.000 | 0.000 | 0.000 |
| 0610033M10 | 0.000 | 0.000 | 0.000 |
| Gm38811 | 0.000 | 0.000 | 0.000 |
| Gm44077 | 0.000 | 0.000 | 0.000 |
| Gm28308 | 0.000 | 0.000 | 0.000 |
| Hoxa13 | 0.000 | 0.000 | 0.000 |
| Evx1os | 0.000 | 0.000 | 0.000 |
| Evx1 | 0.000 | 0.000 | 0.000 |
| Gm44028 | 0.000 | 0.000 | 0.000 |
| Gm16499 | 0.000 | 0.000 | 0.001 |
| Gm28402 | 0.000 | 0.000 | 0.000 |
| Inmt | 0.000 | 0.000 | 0.002 |
| Ccdc129 | 0.000 | 0.000 | 0.000 |
| Ppp1r17 | 0.000 | 0.000 | 0.000 |
| Vmn1r6 | 0.000 | 0.000 | 0.000 |
| Vmn1r8 | 0.000 | 0.000 | 0.000 |
| Vmn1r9 | 0.000 | 0.000 | 0.000 |
| Vmn1r12 | 0.000 | 0.001 | 0.000 |
| Vmn1r13 | 0.000 | 0.000 | 0.000 |
| Vmn1r15 | 0.000 | 0.000 | 0.000 |
| Vmn1r16 | 0.000 | 0.000 | 0.000 |
| Vmn1r18 | 0.000 | 0.000 | 0.000 |
| Vmn1r20 | 0.000 | 0.000 | 0.000 |
| 4930533I22R | 0.000 | 0.000 | 0.000 |
| Vmn1r24 | 0.000 | 0.000 | 0.000 |
| Vmn1r28 | 0.000 | 0.000 | 0.000 |
| Gm45060 | 0.000 | 0.000 | 0.001 |
| BB365896 | 0.000 | 0.000 | 0.000 |
| Atoh1 | 0.000 | 0.000 | 0.000 |
| C130060K24 | 0.000 | 0.000 | 0.000 |
| Tnip3 | 0.000 | 0.001 | 0.001 |

| | | | |
|---|---|---|---|
| Gm16838 | 0.000 | 0.003 | 0.002 |
| 4930544G11I | 0.000 | 0.000 | 0.000 |
| 2610300M13 | 0.000 | 0.000 | 0.002 |
| Vmn1r32 | 0.000 | 0.000 | 0.000 |
| Vmn1r38 | 0.000 | 0.000 | 0.000 |
| Vmn1r39 | 0.000 | 0.000 | 0.000 |
| Il12rb2 | 0.000 | 0.001 | 0.000 |
| Il23r | 0.000 | 0.000 | 0.000 |
| Tacstd2 | 0.000 | 0.000 | 0.000 |
| Igkv12-44 | 0.000 | 0.000 | 0.000 |
| Igkv1-35 | 0.000 | 0.000 | 0.000 |
| Igkc | 0.000 | 0.001 | 0.000 |
| Gm44427 | 0.000 | 0.000 | 0.000 |
| Fabp1 | 0.000 | 0.000 | 0.000 |
| Cd8b1 | 0.000 | 0.000 | 0.001 |
| Gm44110 | 0.000 | 0.000 | 0.000 |
| Cd8a | 0.000 | 0.000 | 0.000 |
| 1700040L08F | 0.000 | 0.000 | 0.000 |
| 4933431G14I | 0.000 | 0.000 | 0.000 |
| Gm20560 | 0.000 | 0.000 | 0.000 |
| Gm44166 | 0.000 | 0.000 | 0.000 |
| Reg3g | 0.000 | 0.001 | 0.001 |
| Gm44271 | 0.000 | 0.000 | 0.000 |
| 1700009C05F | 0.000 | 0.000 | 0.000 |
| Gm20383 | 0.000 | 0.003 | 0.001 |
| Tacr1 | 0.000 | 0.009 | 0.002 |
| Gm17034 | 0.000 | 0.001 | 0.001 |
| Gm38843 | 0.000 | 0.000 | 0.001 |
| Gm32591 | 0.000 | 0.000 | 0.000 |
| 2310069B03F | 0.000 | 0.000 | 0.000 |
| Gm45145 | 0.000 | 0.000 | 0.000 |
| Tlx2 | 0.000 | 0.000 | 0.000 |
| Ccdc142 | 0.000 | 0.000 | 0.000 |
| Gm42688 | 0.000 | 0.000 | 0.000 |
| Gm44287 | 0.000 | 0.000 | 0.000 |
| Slc4a5 | 0.000 | 0.001 | 0.001 |
| Clec4f | 0.000 | 0.000 | 0.000 |
| Cd207 | 0.000 | 0.000 | 0.000 |
| Vax2 | 0.000 | 0.000 | 0.000 |
| Ankrd53 | 0.000 | 0.000 | 0.000 |
| Gm33024 | 0.000 | 0.000 | 0.000 |
| Gm15475 | 0.000 | 0.000 | 0.000 |
| Gm44012 | 0.000 | 0.000 | 0.000 |
| Gm5878 | 0.000 | 0.000 | 0.000 |
| Emx1 | 0.000 | 0.000 | 0.000 |
| Fbxo41 | 0.000 | 0.003 | 0.003 |
| Nat8f3 | 0.000 | 0.000 | 0.001 |
| Nat8f6 | 0.000 | 0.000 | 0.000 |

| | | | |
|---|---|---|---|
| Nat8f5 | 0.000 | 0.000 | 0.000 |
| Nat8 | 0.000 | 0.000 | 0.000 |
| Nat8f2 | 0.000 | 0.000 | 0.000 |
| Gm19265 | 0.000 | 0.001 | 0.001 |
| 4930553P18F | 0.000 | 0.000 | 0.001 |
| Gm44089 | 0.000 | 0.000 | 0.000 |
| Gm44941 | 0.000 | 0.000 | 0.000 |
| Gm44386 | 0.000 | 0.000 | 0.000 |
| Gm44036 | 0.000 | 0.000 | 0.000 |
| Gm44418 | 0.000 | 0.001 | 0.000 |
| Gkn1 | 0.000 | 0.001 | 0.000 |
| Gkn2 | 0.000 | 0.000 | 0.000 |
| Gm34312 | 0.000 | 0.000 | 0.001 |
| Gm43904 | 0.000 | 0.000 | 0.000 |
| Gm45140 | 0.000 | 0.000 | 0.000 |
| Gm26636 | 0.000 | 0.000 | 0.000 |
| 1700031F10F | 0.000 | 0.000 | 0.000 |
| Gm44170 | 0.000 | 0.001 | 0.000 |
| Kbtbd12 | 0.000 | 0.000 | 0.000 |
| Gm26588 | 0.000 | 0.000 | 0.000 |
| Gm839 | 0.000 | 0.000 | 0.000 |
| 4930512J16R | 0.000 | 0.000 | 0.000 |
| Vmn1r41 | 0.000 | 0.000 | 0.000 |
| Vmn1r43 | 0.000 | 0.000 | 0.000 |
| Vmn1r46 | 0.000 | 0.000 | 0.000 |
| Vmn1r47 | 0.000 | 0.000 | 0.000 |
| Vmn1r49 | 0.000 | 0.000 | 0.000 |
| Vmn1r53 | 0.000 | 0.000 | 0.000 |
| Vmn1r54 | 0.000 | 0.000 | 0.000 |
| Chst13 | 0.000 | 0.000 | 0.000 |
| Uroc1 | 0.000 | 0.000 | 0.000 |
| Cfap100 | 0.000 | 0.001 | 0.000 |
| Gm45901 | 0.000 | 0.000 | 0.000 |
| Gm15756 | 0.000 | 0.000 | 0.000 |
| Gm44123 | 0.000 | 0.000 | 0.001 |
| 4930471M09 | 0.000 | 0.000 | 0.000 |
| Gm45218 | 0.000 | 0.000 | 0.000 |
| Gm15737 | 0.000 | 0.001 | 0.000 |
| 4930511E03F | 0.000 | 0.001 | 0.000 |
| 1700123L14F | 0.000 | 0.000 | 0.000 |
| Fam19a4 | 0.000 | 0.000 | 0.000 |
| Gm32592 | 0.000 | 0.001 | 0.001 |
| Gm765 | 0.000 | 0.000 | 0.000 |
| Gm44169 | 0.000 | 0.000 | 0.000 |
| Gm9871 | 0.000 | 0.000 | 0.000 |
| Cntn3 | 0.000 | 0.017 | 0.002 |
| Gm19757 | 0.000 | 0.000 | 0.000 |
| Gm15631 | 0.000 | 0.000 | 0.000 |

| | | | |
|---|---|---|---|
| Il5ra | 0.000 | 0.000 | 0.000 |
| 4933431M02 | 0.000 | 0.000 | 0.000 |
| Gm44040 | 0.000 | 0.000 | 0.000 |
| 0610040F04F | 0.000 | 0.001 | 0.001 |
| Gm20387 | 0.000 | 0.000 | 0.000 |
| Grm7 | 0.000 | 0.001 | 0.004 |
| Cidec | 0.000 | 0.001 | 0.001 |
| Fancd2os | 0.000 | 0.000 | 0.000 |
| Gm44199 | 0.000 | 0.000 | 0.000 |
| 1700015O11I | 0.000 | 0.000 | 0.000 |
| Ghrl | 0.000 | 0.001 | 0.001 |
| Gm15083 | 0.000 | 0.000 | 0.000 |
| Slc6a11 | 0.000 | 0.000 | 0.000 |
| Slc6a1 | 0.000 | 0.000 | 0.000 |
| Gm44079 | 0.000 | 0.000 | 0.000 |
| Mkrn2os | 0.000 | 0.000 | 0.001 |
| Gm44206 | 0.000 | 0.000 | 0.000 |
| Gm20404 | 0.000 | 0.000 | 0.000 |
| Alox5 | 0.000 | 0.000 | 0.001 |
| Gm5580 | 0.000 | 0.000 | 0.000 |
| Tmem72 | 0.000 | 0.001 | 0.000 |
| Gm4640 | 0.000 | 0.001 | 0.000 |
| Gm45083 | 0.000 | 0.000 | 0.000 |
| 1700030F04F | 0.000 | 0.000 | 0.000 |
| Gm7292 | 0.000 | 0.000 | 0.000 |
| Gm44317 | 0.000 | 0.001 | 0.000 |
| Ninj2 | 0.000 | 0.000 | 0.000 |
| B4galnt3 | 0.000 | 0.005 | 0.001 |
| D330020A13I | 0.000 | 0.003 | 0.001 |
| Gm20692 | 0.000 | 0.000 | 0.000 |
| Gm15856 | 0.000 | 0.000 | 0.000 |
| Slc6a12 | 0.000 | 0.002 | 0.003 |
| Iqsec3 | 0.000 | 0.000 | 0.000 |
| Klrg1 | 0.000 | 0.000 | 0.000 |
| 1700063H04I | 0.000 | 0.000 | 0.000 |
| Aicda | 0.000 | 0.000 | 0.000 |
| Gdf3 | 0.000 | 0.000 | 0.000 |
| Dppa3 | 0.000 | 0.000 | 0.000 |
| Nanog | 0.000 | 0.000 | 0.000 |
| Clec4a4 | 0.000 | 0.000 | 0.000 |
| Clec4b1 | 0.000 | 0.000 | 0.000 |
| Gm43971 | 0.000 | 0.000 | 0.000 |
| Cd163 | 0.000 | 0.003 | 0.003 |
| 1700013D24I | 0.000 | 0.000 | 0.000 |
| 1700027F06F | 0.000 | 0.000 | 0.000 |
| Gm44096 | 0.000 | 0.000 | 0.000 |
| Gm16567 | 0.000 | 0.000 | 0.000 |
| C1rb | 0.000 | 0.000 | 0.001 |

| | | | |
|---|---|---|---|
| Gm15884 | 0.000 | 0.000 | 0.000 |
| Gm20531 | 0.000 | 0.000 | 0.000 |
| Cd4 | 0.000 | 0.000 | 0.001 |
| A230083G16I | 0.000 | 0.001 | 0.001 |
| 4930557K07F | 0.000 | 0.000 | 0.000 |
| Tuba3a | 0.000 | 0.000 | 0.000 |
| Gm45869 | 0.000 | 0.000 | 0.000 |
| Kcna1 | 0.000 | 0.002 | 0.004 |
| Gm26673 | 0.000 | 0.000 | 0.000 |
| Akap3 | 0.000 | 0.000 | 0.000 |
| Dyrk4 | 0.000 | 0.000 | 0.000 |
| Platr31 | 0.000 | 0.000 | 0.000 |
| Prmt8 | 0.000 | 0.000 | 0.001 |
| Gm38901 | 0.000 | 0.000 | 0.002 |
| 9330102E08F | 0.000 | 0.001 | 0.001 |
| Pzp | 0.000 | 0.000 | 0.000 |
| A2ml1 | 0.000 | 0.000 | 0.001 |
| Gm44009 | 0.000 | 0.000 | 0.000 |
| Klrb1a | 0.000 | 0.000 | 0.000 |
| Klrb1c | 0.000 | 0.000 | 0.000 |
| Klrb1b | 0.000 | 0.000 | 0.000 |
| Gm26656 | 0.000 | 0.001 | 0.001 |
| BC035044 | 0.000 | 0.000 | 0.000 |
| Clec2i | 0.000 | 0.000 | 0.000 |
| Clec2g | 0.000 | 0.000 | 0.000 |
| Gm15987 | 0.000 | 0.000 | 0.000 |
| Klrb1f | 0.000 | 0.000 | 0.000 |
| Clec2e | 0.000 | 0.000 | 0.001 |
| Gm47861 | 0.000 | 0.000 | 0.000 |
| Tmem52b | 0.000 | 0.000 | 0.000 |
| Klrd1 | 0.000 | 0.000 | 0.000 |
| Klrk1 | 0.000 | 0.000 | 0.000 |
| Klrc2 | 0.000 | 0.000 | 0.000 |
| Klrc1 | 0.000 | 0.000 | 0.000 |
| Klri2 | 0.000 | 0.000 | 0.000 |
| Klra17 | 0.000 | 0.000 | 0.000 |
| Klra5 | 0.000 | 0.000 | 0.000 |
| Klra1 | 0.000 | 0.000 | 0.000 |
| Tas2r136 | 0.000 | 0.000 | 0.000 |
| 4933406J09R | 0.000 | 0.000 | 0.000 |
| Bcl2l14 | 0.000 | 0.000 | 0.000 |
| Gm36328 | 0.000 | 0.000 | 0.000 |
| Gm44238 | 0.000 | 0.001 | 0.000 |
| Gprc5d | 0.000 | 0.001 | 0.000 |
| Gm19434 | 0.000 | 0.000 | 0.000 |
| Gm36640 | 0.000 | 0.000 | 0.000 |
| Grin2b | 0.000 | 0.000 | 0.000 |
| Gm14329 | 0.000 | 0.000 | 0.000 |

| | | | |
|---|---|---|---|
| Gm26653 | 0.000 | 0.000 | 0.000 |
| E330021D16F | 0.000 | 0.000 | 0.000 |
| Gucy2c | 0.000 | 0.000 | 0.000 |
| Smco3 | 0.000 | 0.001 | 0.004 |
| Slc15a5 | 0.000 | 0.000 | 0.001 |
| 4922502N22I | 0.000 | 0.000 | 0.000 |
| Rergl | 0.000 | 0.001 | 0.000 |
| Pik3c2g | 0.000 | 0.000 | 0.002 |
| Plcz1 | 0.000 | 0.000 | 0.000 |
| Slco1a6 | 0.000 | 0.000 | 0.000 |
| Gm20400 | 0.000 | 0.004 | 0.002 |
| Slco1a5 | 0.000 | 0.000 | 0.000 |
| Gys2 | 0.000 | 0.000 | 0.000 |
| 5330439B14F | 0.000 | 0.000 | 0.000 |
| Sult6b2 | 0.000 | 0.000 | 0.000 |
| St8sia1 | 0.000 | 0.000 | 0.000 |
| 1700126G02I | 0.000 | 0.000 | 0.000 |
| D6Ertd474e | 0.000 | 0.000 | 0.000 |
| Gm15687 | 0.000 | 0.000 | 0.000 |
| Gm15499 | 0.000 | 0.000 | 0.000 |
| Gm15704 | 0.000 | 0.002 | 0.003 |
| Sspnos | 0.000 | 0.000 | 0.001 |
| Gm32914 | 0.000 | 0.000 | 0.000 |
| 4930479D17I | 0.000 | 0.000 | 0.000 |
| Smco2 | 0.000 | 0.001 | 0.000 |
| Mansc4 | 0.000 | 0.001 | 0.001 |
| 1700049E15F | 0.000 | 0.006 | 0.007 |
| Far2os1 | 0.000 | 0.000 | 0.000 |
| Gm6313 | 0.000 | 0.000 | 0.000 |
| 4930528G23I | 0.000 | 0.000 | 0.000 |
| Gm43913 | 0.000 | 0.000 | 0.000 |
| 1700003I16R | 0.000 | 0.000 | 0.000 |
| Nlrp12 | 0.000 | 0.000 | 0.000 |
| Cacng6 | 0.000 | 0.000 | 0.000 |
| Tarm1 | 0.000 | 0.000 | 0.001 |
| Gm44920 | 0.000 | 0.000 | 0.000 |
| Pira2 | 0.000 | 0.000 | 0.000 |
| Gm14548 | 0.000 | 0.000 | 0.000 |
| Lilra6 | 0.000 | 0.000 | 0.000 |
| Lilra5 | 0.000 | 0.000 | 0.000 |
| Ncr1 | 0.000 | 0.000 | 0.000 |
| Gp6 | 0.000 | 0.000 | 0.000 |
| Gm15494 | 0.000 | 0.001 | 0.000 |
| Tmem150b | 0.000 | 0.001 | 0.000 |
| Tmem190 | 0.000 | 0.000 | 0.000 |
| Shisa7 | 0.000 | 0.000 | 0.000 |
| Gm36210 | 0.000 | 0.000 | 0.002 |
| Gm44983 | 0.000 | 0.000 | 0.000 |

| | | | |
|---|---|---|---|
| Sbk2 | 0.000 | 0.001 | 0.001 |
| Rfpl4 | 0.000 | 0.000 | 0.000 |
| Gm5065 | 0.000 | 0.000 | 0.000 |
| Vmn2r28 | 0.000 | 0.000 | 0.000 |
| Vmn1r65 | 0.000 | 0.000 | 0.000 |
| Gm20715 | 0.000 | 0.000 | 0.000 |
| Olfr1336 | 0.000 | 0.000 | 0.000 |
| Olfr1346 | 0.000 | 0.000 | 0.000 |
| Vmn2r29 | 0.000 | 0.001 | 0.000 |
| Gm45631 | 0.000 | 0.000 | 0.000 |
| Vmn2r30 | 0.000 | 0.000 | 0.000 |
| Vmn1r66 | 0.000 | 0.000 | 0.000 |
| Vmn1r72 | 0.000 | 0.000 | 0.000 |
| Vmn1r73 | 0.000 | 0.000 | 0.000 |
| Vmn1r74 | 0.000 | 0.000 | 0.000 |
| Vmn1r78 | 0.000 | 0.000 | 0.000 |
| Vmn1r79 | 0.000 | 0.000 | 0.000 |
| Vmn1r83 | 0.000 | 0.000 | 0.000 |
| Vmn1r84 | 0.000 | 0.000 | 0.000 |
| Gm31152 | 0.000 | 0.001 | 0.000 |
| Gm29638 | 0.000 | 0.000 | 0.000 |
| Vmn1r86 | 0.000 | 0.000 | 0.000 |
| Vmn1r87 | 0.000 | 0.000 | 0.000 |
| Cabp5 | 0.000 | 0.000 | 0.000 |
| Sult2a8 | 0.000 | 0.000 | 0.000 |
| Sult2a7 | 0.000 | 0.000 | 0.000 |
| Obox6 | 0.000 | 0.000 | 0.000 |
| Crx | 0.000 | 0.000 | 0.000 |
| Crxos | 0.000 | 0.000 | 0.000 |
| Zfp541 | 0.000 | 0.000 | 0.000 |
| Gm45510 | 0.000 | 0.000 | 0.000 |
| Npas1 | 0.000 | 0.000 | 0.000 |
| Gm29443 | 0.000 | 0.000 | 0.000 |
| Gm29442 | 0.000 | 0.000 | 0.000 |
| Ceacam15 | 0.000 | 0.001 | 0.001 |
| Ceacam9 | 0.000 | 0.000 | 0.000 |
| Gm32846 | 0.000 | 0.000 | 0.000 |
| Psg27 | 0.000 | 0.000 | 0.000 |
| Psg23 | 0.000 | 0.000 | 0.000 |
| Psg22 | 0.000 | 0.000 | 0.000 |
| Gm4969 | 0.000 | 0.000 | 0.000 |
| 1700058P15R | 0.000 | 0.004 | 0.002 |
| A930016O22I | 0.000 | 0.000 | 0.000 |
| Apoc2 | 0.000 | 0.000 | 0.000 |
| Apoc4 | 0.000 | 0.001 | 0.001 |
| Gm26600 | 0.000 | 0.001 | 0.000 |
| Gm16175 | 0.000 | 0.000 | 0.000 |
| Gm44658 | 0.000 | 0.000 | 0.000 |

| | | | |
|---|---|---|---|
| Ceacam16 | 0.000 | 0.000 | 0.000 |
| Ceacam19 | 0.000 | 0.000 | 0.000 |
| Nlrp9b | 0.000 | 0.000 | 0.000 |
| Vmn1r180 | 0.000 | 0.000 | 0.000 |
| Vmn1r181 | 0.000 | 0.000 | 0.000 |
| Lypd5 | 0.000 | 0.000 | 0.000 |
| Irgc1 | 0.000 | 0.001 | 0.000 |
| Pinlyp | 0.000 | 0.000 | 0.000 |
| Lypd3 | 0.000 | 0.000 | 0.000 |
| Gm4598 | 0.000 | 0.000 | 0.000 |
| Tex101 | 0.000 | 0.000 | 0.000 |
| BC049730 | 0.000 | 0.000 | 0.000 |
| Cd177 | 0.000 | 0.000 | 0.000 |
| D930028M14 | 0.000 | 0.000 | 0.000 |
| 4933430L12F | 0.000 | 0.000 | 0.000 |
| Cnfn | 0.000 | 0.002 | 0.001 |
| Cxcl17 | 0.000 | 0.001 | 0.000 |
| Gm38569 | 0.000 | 0.000 | 0.000 |
| Erich4 | 0.000 | 0.001 | 0.001 |
| Gm30146 | 0.000 | 0.000 | 0.000 |
| Cyp2b10 | 0.000 | 0.004 | 0.001 |
| BC026762 | 0.000 | 0.000 | 0.000 |
| Cyp2b13 | 0.000 | 0.000 | 0.000 |
| Nlrp4a | 0.000 | 0.000 | 0.000 |
| Vmn1r185 | 0.000 | 0.000 | 0.000 |
| Cyp2b23 | 0.000 | 0.000 | 0.000 |
| Cyp2a5 | 0.000 | 0.000 | 0.000 |
| Cyp2f2 | 0.000 | 0.000 | 0.000 |
| Gm15883 | 0.000 | 0.000 | 0.000 |
| Cyp2t4 | 0.000 | 0.000 | 0.000 |
| Gm21983 | 0.000 | 0.000 | 0.000 |
| Gm15567 | 0.000 | 0.001 | 0.000 |
| Gm26891 | 0.000 | 0.000 | 0.000 |
| Gm44744 | 0.000 | 0.000 | 0.000 |
| Fcgbp | 0.000 | 0.000 | 0.000 |
| 9530053A07F | 0.000 | 0.000 | 0.000 |
| Dll3 | 0.000 | 0.001 | 0.001 |
| Sycn | 0.000 | 0.000 | 0.000 |
| Nccrp1 | 0.000 | 0.000 | 0.001 |
| Gm44618 | 0.000 | 0.000 | 0.000 |
| Ccer2 | 0.000 | 0.000 | 0.000 |
| Gm10648 | 0.000 | 0.000 | 0.001 |
| Gm26604 | 0.000 | 0.000 | 0.001 |
| Gm44632 | 0.000 | 0.000 | 0.000 |
| A330087D11I | 0.000 | 0.000 | 0.000 |
| Gm26554 | 0.000 | 0.000 | 0.000 |
| Ovol3 | 0.000 | 0.000 | 0.000 |
| Kirrel2 | 0.000 | 0.000 | 0.000 |

| | | | |
|---|---|---|---|
| Nphs1 | 0.000 | 0.001 | 0.001 |
| Prodh2 | 0.000 | 0.000 | 0.000 |
| Igflr1 | 0.000 | 0.000 | 0.000 |
| 2200002J24R | 0.000 | 0.000 | 0.000 |
| Atp4a | 0.000 | 0.001 | 0.000 |
| Gapdhs | 0.000 | 0.000 | 0.000 |
| Ffar2 | 0.000 | 0.000 | 0.000 |
| Ffar3 | 0.000 | 0.000 | 0.000 |
| Ffar1 | 0.000 | 0.001 | 0.000 |
| Cd22 | 0.000 | 0.000 | 0.000 |
| D7Ertd128e | 0.000 | 0.000 | 0.000 |
| Hamp | 0.000 | 0.001 | 0.000 |
| Gm17077 | 0.000 | 0.000 | 0.000 |
| Fxyd3 | 0.000 | 0.003 | 0.001 |
| Scgb1b30 | 0.000 | 0.000 | 0.000 |
| Gm12762 | 0.000 | 0.000 | 0.001 |
| Gm6096 | 0.000 | 0.000 | 0.000 |
| Gm12766 | 0.000 | 0.000 | 0.000 |
| Gm12784 | 0.000 | 0.001 | 0.000 |
| Gm12781 | 0.000 | 0.001 | 0.001 |
| Gm35611 | 0.000 | 0.000 | 0.000 |
| Slc7a9 | 0.000 | 0.000 | 0.000 |
| Gm26790 | 0.000 | 0.001 | 0.001 |
| Gm28075 | 0.000 | 0.000 | 0.000 |
| D530033B14I | 0.000 | 0.000 | 0.000 |
| Gm44767 | 0.000 | 0.000 | 0.000 |
| 4930558N11I | 0.000 | 0.000 | 0.000 |
| Gm44992 | 0.000 | 0.000 | 0.000 |
| A230077H06I | 0.000 | 0.001 | 0.001 |
| Gm28307 | 0.000 | 0.000 | 0.000 |
| Vmn2r57 | 0.000 | 0.000 | 0.000 |
| Gm6871 | 0.000 | 0.000 | 0.000 |
| Gm21028 | 0.000 | 0.000 | 0.000 |
| Gm2381 | 0.000 | 0.000 | 0.000 |
| Siglecf | 0.000 | 0.000 | 0.000 |
| Nkg7 | 0.000 | 0.000 | 0.000 |
| Gm45233 | 0.000 | 0.000 | 0.000 |
| 4931406B18I | 0.000 | 0.000 | 0.000 |
| Gm2511 | 0.000 | 0.000 | 0.000 |
| Zfp819 | 0.000 | 0.000 | 0.000 |
| Ceacam18 | 0.000 | 0.000 | 0.000 |
| Siglece | 0.000 | 0.001 | 0.000 |
| Klk13 | 0.000 | 0.000 | 0.000 |
| Klk11 | 0.000 | 0.000 | 0.000 |
| Klk6 | 0.000 | 0.000 | 0.000 |
| Klk1b3 | 0.000 | 0.000 | 0.000 |
| Acp4 | 0.000 | 0.000 | 0.000 |
| Gm15517 | 0.000 | 0.001 | 0.001 |

| | | | |
|---|---|---|---|
| Gm44780 | 0.000 | 0.000 | 0.000 |
| Aspdh | 0.000 | 0.001 | 0.001 |
| Gm44832 | 0.000 | 0.000 | 0.000 |
| Myh14 | 0.000 | 0.000 | 0.000 |
| Tsks | 0.000 | 0.001 | 0.001 |
| Gm45552 | 0.000 | 0.000 | 0.000 |
| Gm45713 | 0.000 | 0.000 | 0.000 |
| Gfy | 0.000 | 0.000 | 0.000 |
| Pth2 | 0.000 | 0.000 | 0.000 |
| Ccdc155 | 0.000 | 0.000 | 0.000 |
| Mtag2 | 0.000 | 0.001 | 0.000 |
| Gm45619 | 0.000 | 0.000 | 0.000 |
| Kcna7 | 0.000 | 0.000 | 0.000 |
| Lhb | 0.000 | 0.000 | 0.000 |
| Gm45564 | 0.000 | 0.000 | 0.000 |
| Syngr4 | 0.000 | 0.000 | 0.002 |
| Gm45442 | 0.000 | 0.001 | 0.000 |
| Abcc6 | 0.000 | 0.001 | 0.002 |
| Kcnj11 | 0.000 | 0.004 | 0.001 |
| Gm45486 | 0.000 | 0.001 | 0.000 |
| Ush1c | 0.000 | 0.000 | 0.000 |
| 1700025L06F | 0.000 | 0.000 | 0.000 |
| Kcnc1 | 0.000 | 0.001 | 0.000 |
| Gm15700 | 0.000 | 0.000 | 0.000 |
| Tph1 | 0.000 | 0.000 | 0.000 |
| Saa1 | 0.000 | 0.000 | 0.001 |
| Saa2 | 0.000 | 0.000 | 0.001 |
| C86187 | 0.000 | 0.001 | 0.001 |
| Ptpn5 | 0.000 | 0.000 | 0.000 |
| Mrgpra2a | 0.000 | 0.000 | 0.000 |
| Mrgpra2b | 0.000 | 0.000 | 0.000 |
| Mrgprb1 | 0.000 | 0.000 | 0.000 |
| Mrgprx2 | 0.000 | 0.000 | 0.000 |
| Mrgprb2 | 0.000 | 0.000 | 0.000 |
| Gm26856 | 0.000 | 0.000 | 0.000 |
| Nell1os | 0.000 | 0.001 | 0.000 |
| 4933405O20I | 0.000 | 0.001 | 0.000 |
| Ano5 | 0.000 | 0.000 | 0.000 |
| Slc17a6 | 0.000 | 0.000 | 0.000 |
| Siglech | 0.000 | 0.000 | 0.000 |
| Oca2 | 0.000 | 0.000 | 0.000 |
| Gm9962 | 0.000 | 0.000 | 0.000 |
| Gabra5 | 0.000 | 0.000 | 0.001 |
| Gm44644 | 0.000 | 0.000 | 0.000 |
| B230209E15F | 0.000 | 0.000 | 0.000 |
| Gm32061 | 0.000 | 0.000 | 0.000 |
| Chrna7 | 0.000 | 0.002 | 0.001 |
| 4930554H23I | 0.000 | 0.000 | 0.000 |

| | | | |
|---|---|---|---|
| Gm44718 | 0.000 | 0.000 | 0.000 |
| Gm45052 | 0.000 | 0.000 | 0.000 |
| Gm32633 | 0.000 | 0.000 | 0.000 |
| Fam189a1 | 0.000 | 0.000 | 0.000 |
| Cers3 | 0.000 | 0.001 | 0.000 |
| Gm33926 | 0.000 | 0.000 | 0.000 |
| Gm39027 | 0.000 | 0.000 | 0.000 |
| Gm16157 | 0.000 | 0.000 | 0.000 |
| Fam169b | 0.000 | 0.009 | 0.003 |
| Gm44889 | 0.000 | 0.000 | 0.000 |
| 4933436H12I | 0.000 | 0.000 | 0.000 |
| Gm34549 | 0.000 | 0.000 | 0.000 |
| Gm44886 | 0.000 | 0.000 | 0.000 |
| Gm44691 | 0.000 | 0.000 | 0.000 |
| Gm34838 | 0.000 | 0.000 | 0.000 |
| Gm38584 | 0.000 | 0.000 | 0.000 |
| Gm29683 | 0.000 | 0.001 | 0.001 |
| Gm44946 | 0.000 | 0.000 | 0.001 |
| Gm44532 | 0.000 | 0.000 | 0.001 |
| Gm42397 | 0.000 | 0.000 | 0.006 |
| Gm36011 | 0.000 | 0.000 | 0.000 |
| Gm39033 | 0.000 | 0.000 | 0.000 |
| 6030442E23F | 0.000 | 0.000 | 0.001 |
| Gm29328 | 0.000 | 0.000 | 0.000 |
| 1700011C11F | 0.000 | 0.000 | 0.000 |
| Mctp2 | 0.000 | 0.001 | 0.004 |
| Gm30075 | 0.000 | 0.001 | 0.002 |
| A730056A06F | 0.000 | 0.003 | 0.007 |
| Gm44737 | 0.000 | 0.000 | 0.000 |
| Gm44736 | 0.000 | 0.000 | 0.000 |
| Gm44739 | 0.000 | 0.001 | 0.000 |
| Gm10619 | 0.000 | 0.000 | 0.000 |
| Gm45004 | 0.000 | 0.000 | 0.000 |
| Gm7580 | 0.000 | 0.000 | 0.000 |
| Gm30459 | 0.000 | 0.000 | 0.000 |
| 0610006L08F | 0.000 | 0.000 | 0.000 |
| 4930533N22I | 0.000 | 0.000 | 0.000 |
| 4921513I08R | 0.000 | 0.000 | 0.000 |
| 5330411O13I | 0.000 | 0.000 | 0.000 |
| 1700011D18I | 0.000 | 0.001 | 0.000 |
| Gm26633 | 0.000 | 0.000 | 0.000 |
| Rlbp1 | 0.000 | 0.000 | 0.000 |
| Mir9-3hg | 0.000 | 0.000 | 0.000 |
| Gm35040 | 0.000 | 0.000 | 0.000 |
| Gm45169 | 0.000 | 0.000 | 0.000 |
| 9330171B17F | 0.000 | 0.000 | 0.000 |
| Gm44706 | 0.000 | 0.000 | 0.000 |
| Gm26646 | 0.000 | 0.000 | 0.000 |

| | | | |
|---|---|---|---|
| Gm21057 | 0.000 | 0.000 | 0.000 |
| Gm44851 | 0.000 | 0.000 | 0.000 |
| Gm44649 | 0.000 | 0.000 | 0.000 |
| Ap3b2 | 0.000 | 0.001 | 0.003 |
| Gm45016 | 0.000 | 0.000 | 0.000 |
| Gm10160 | 0.000 | 0.000 | 0.000 |
| Gm45014 | 0.000 | 0.000 | 0.000 |
| Bnc1 | 0.000 | 0.000 | 0.001 |
| Gm20744 | 0.000 | 0.000 | 0.000 |
| 4933406J10R | 0.000 | 0.000 | 0.000 |
| A530021J07R | 0.000 | 0.000 | 0.000 |
| Gm44724 | 0.000 | 0.000 | 0.000 |
| Gm44991 | 0.000 | 0.000 | 0.000 |
| Gm10610 | 0.000 | 0.000 | 0.000 |
| Cfap161 | 0.000 | 0.000 | 0.000 |
| Gm39043 | 0.000 | 0.000 | 0.000 |
| Gm44827 | 0.000 | 0.000 | 0.000 |
| Olfr291 | 0.000 | 0.000 | 0.000 |
| Olfr307 | 0.000 | 0.000 | 0.000 |
| Grm5 | 0.000 | 0.000 | 0.000 |
| Gm44995 | 0.000 | 0.000 | 0.000 |
| AI314278 | 0.000 | 0.000 | 0.001 |
| Prss23os | 0.000 | 0.000 | 0.000 |
| Gm45159 | 0.000 | 0.001 | 0.001 |
| 4930567K12F | 0.000 | 0.000 | 0.000 |
| Gm32647 | 0.000 | 0.000 | 0.001 |
| Gm15416 | 0.000 | 0.000 | 0.000 |
| Rsf1os1 | 0.000 | 0.000 | 0.000 |
| Gdpd4 | 0.000 | 0.000 | 0.000 |
| B3gnt6 | 0.000 | 0.000 | 0.000 |
| Gucy2d | 0.000 | 0.000 | 0.000 |
| Gm19656 | 0.000 | 0.000 | 0.000 |
| Gm45188 | 0.000 | 0.000 | 0.001 |
| Gm34280 | 0.000 | 0.000 | 0.000 |
| Olfr520 | 0.000 | 0.000 | 0.000 |
| F730035P03F | 0.000 | 0.000 | 0.000 |
| Gm34821 | 0.000 | 0.000 | 0.000 |
| Gm10603 | 0.000 | 0.001 | 0.000 |
| Ucp3 | 0.000 | 0.000 | 0.000 |
| Dnajb13 | 0.000 | 0.000 | 0.000 |
| Gm39059 | 0.000 | 0.000 | 0.000 |
| Gm20476 | 0.000 | 0.000 | 0.000 |
| Pde2a | 0.000 | 0.000 | 0.000 |
| Gm35363 | 0.000 | 0.000 | 0.000 |
| Folr1 | 0.000 | 0.003 | 0.005 |
| Art1 | 0.000 | 0.002 | 0.002 |
| Chrna10 | 0.000 | 0.000 | 0.000 |
| Olfr543 | 0.000 | 0.000 | 0.002 |

| | | | |
|---|---|---|---|
| Olfr544 | 0.000 | 0.000 | 0.000 |
| Olfr545 | 0.000 | 0.001 | 0.001 |
| Olfr553 | 0.000 | 0.000 | 0.000 |
| Olfr78 | 0.000 | 0.001 | 0.001 |
| Olfr571 | 0.000 | 0.000 | 0.000 |
| Olfr592 | 0.000 | 0.000 | 0.000 |
| Usp17ld | 0.000 | 0.000 | 0.000 |
| Olfr596 | 0.000 | 0.000 | 0.001 |
| Olfr600 | 0.000 | 0.000 | 0.000 |
| Olfr607 | 0.000 | 0.000 | 0.000 |
| Olfr608 | 0.000 | 0.000 | 0.000 |
| Olfr613 | 0.000 | 0.000 | 0.000 |
| Olfr620 | 0.000 | 0.000 | 0.000 |
| Olfr243 | 0.000 | 0.000 | 0.000 |
| Hbb-bh1 | 0.000 | 0.000 | 0.000 |
| Olfr633 | 0.000 | 0.000 | 0.000 |
| Olfr635 | 0.000 | 0.001 | 0.001 |
| Olfr639 | 0.000 | 0.001 | 0.001 |
| Olfr640 | 0.000 | 0.000 | 0.000 |
| Olfr642 | 0.000 | 0.000 | 0.000 |
| 4930516K23F | 0.000 | 0.000 | 0.000 |
| Olfr646 | 0.000 | 0.000 | 0.000 |
| Gm47248 | 0.000 | 0.000 | 0.000 |
| Trim34b | 0.000 | 0.000 | 0.000 |
| Trim30b | 0.000 | 0.000 | 0.000 |
| Olfr652 | 0.000 | 0.000 | 0.000 |
| Olfr658 | 0.000 | 0.000 | 0.000 |
| Usp17la | 0.000 | 0.000 | 0.000 |
| Olfr667 | 0.000 | 0.000 | 0.000 |
| Olfr671 | 0.000 | 0.000 | 0.000 |
| Olfr675 | 0.000 | 0.000 | 0.000 |
| Olfr678 | 0.000 | 0.000 | 0.000 |
| Olfr679 | 0.000 | 0.000 | 0.000 |
| Gm5901 | 0.000 | 0.000 | 0.000 |
| Cnga4 | 0.000 | 0.000 | 0.000 |
| Cckbr | 0.000 | 0.000 | 0.000 |
| Olfr695 | 0.000 | 0.000 | 0.000 |
| Olfr697 | 0.000 | 0.000 | 0.000 |
| 4930458B22F | 0.000 | 0.000 | 0.000 |
| Olfr701 | 0.000 | 0.000 | 0.000 |
| Olfr709-ps1 | 0.000 | 0.000 | 0.000 |
| Olfr710 | 0.000 | 0.000 | 0.000 |
| Olfr715b | 0.000 | 0.000 | 0.000 |
| Olfr716 | 0.000 | 0.000 | 0.000 |
| Platr28 | 0.000 | 0.000 | 0.000 |
| Cyb5r2 | 0.000 | 0.000 | 0.001 |
| Gm45040 | 0.000 | 0.000 | 0.000 |
| Olfr472 | 0.000 | 0.000 | 0.000 |

| | | | |
|---|---|---|---|
| Olfr481 | 0.000 | 0.000 | 0.000 |
| Olfr510 | 0.000 | 0.000 | 0.000 |
| Gm26599 | 0.000 | 0.001 | 0.000 |
| Stk33 | 0.000 | 0.006 | 0.007 |
| BC051019 | 0.000 | 0.000 | 0.000 |
| Gm45515 | 0.000 | 0.000 | 0.000 |
| 5430402P08F | 0.000 | 0.000 | 0.000 |
| Far1os | 0.000 | 0.000 | 0.000 |
| Gm45025 | 0.000 | 0.000 | 0.000 |
| Gm45155 | 0.000 | 0.000 | 0.000 |
| Gpr139 | 0.000 | 0.000 | 0.002 |
| Acsm2 | 0.000 | 0.000 | 0.000 |
| Acsm1 | 0.000 | 0.000 | 0.000 |
| Gm19950 | 0.000 | 0.000 | 0.000 |
| Gm45792 | 0.000 | 0.000 | 0.000 |
| Acsm4 | 0.000 | 0.000 | 0.000 |
| Acsm3 | 0.000 | 0.000 | 0.000 |
| Dnah3 | 0.000 | 0.000 | 0.000 |
| Zp2 | 0.000 | 0.000 | 0.000 |
| 4930505K13F | 0.000 | 0.000 | 0.000 |
| Anks4b | 0.000 | 0.000 | 0.000 |
| Abca16 | 0.000 | 0.000 | 0.000 |
| Hs3st2 | 0.000 | 0.000 | 0.003 |
| 4933432K03F | 0.000 | 0.000 | 0.000 |
| 1700069B07F | 0.000 | 0.001 | 0.000 |
| Scnn1b | 0.000 | 0.000 | 0.001 |
| Gm44987 | 0.000 | 0.000 | 0.000 |
| Ern2 | 0.000 | 0.000 | 0.000 |
| Cacng3 | 0.000 | 0.000 | 0.000 |
| Gm45846 | 0.000 | 0.001 | 0.001 |
| Slc5a11 | 0.000 | 0.000 | 0.000 |
| Gm27040 | 0.000 | 0.000 | 0.000 |
| Hs3st4 | 0.000 | 0.000 | 0.001 |
| 4933440M02 | 0.000 | 0.000 | 0.000 |
| Gm30717 | 0.000 | 0.000 | 0.001 |
| Gsg1l | 0.000 | 0.000 | 0.000 |
| 4930448A20F | 0.000 | 0.000 | 0.000 |
| Aldoa.1 | 0.000 | 0.000 | 0.000 |
| Gm44854 | 0.000 | 0.000 | 0.000 |
| Doc2a | 0.000 | 0.000 | 0.000 |
| Asphd1 | 0.000 | 0.000 | 0.000 |
| Pagr1b | 0.000 | 0.000 | 0.000 |
| Gm31749 | 0.000 | 0.000 | 0.000 |
| Gm31897 | 0.000 | 0.000 | 0.000 |
| Gm28198 | 0.000 | 0.000 | 0.000 |
| Ctf2 | 0.000 | 0.001 | 0.001 |
| Gm15533 | 0.000 | 0.002 | 0.002 |
| Trim72 | 0.000 | 0.001 | 0.001 |

| | | | |
|---|---|---|---|
| Itgam | 0.000 | 0.001 | 0.002 |
| Gm49368 | 0.000 | 0.000 | 0.000 |
| Itgad | 0.000 | 0.000 | 0.000 |
| Gm44850 | 0.000 | 0.000 | 0.000 |
| Slc5a2 | 0.000 | 0.000 | 0.001 |
| Gm40457 | 0.000 | 0.000 | 0.000 |
| Gm45121 | 0.000 | 0.000 | 0.000 |
| Gm33027 | 0.000 | 0.000 | 0.000 |
| Gm39094 | 0.000 | 0.000 | 0.001 |
| Gm4265 | 0.000 | 0.000 | 0.000 |
| Gm44647 | 0.000 | 0.000 | 0.001 |
| Etos1 | 0.000 | 0.000 | 0.000 |
| Dmbt1 | 0.000 | 0.000 | 0.000 |
| Cuzd1 | 0.000 | 0.001 | 0.001 |
| Fam24b | 0.000 | 0.001 | 0.001 |
| Fam24a | 0.000 | 0.000 | 0.000 |
| Hmx3 | 0.000 | 0.000 | 0.000 |
| Gm15677 | 0.000 | 0.000 | 0.000 |
| Gm34908 | 0.000 | 0.000 | 0.000 |
| 4930483O08I | 0.000 | 0.000 | 0.000 |
| Gm28747 | 0.000 | 0.000 | 0.000 |
| Mmp21 | 0.000 | 0.000 | 0.000 |
| Gm4593 | 0.000 | 0.000 | 0.000 |
| 1700120G07I | 0.000 | 0.000 | 0.000 |
| Gm36356 | 0.000 | 0.000 | 0.000 |
| 5830432E09F | 0.000 | 0.000 | 0.000 |
| 6330420H09I | 0.000 | 0.000 | 0.000 |
| Gm36737 | 0.000 | 0.000 | 0.000 |
| C030029H02I | 0.000 | 0.000 | 0.000 |
| C230079O03I | 0.000 | 0.000 | 0.000 |
| Gm6249 | 0.000 | 0.000 | 0.001 |
| Gm45278 | 0.000 | 0.000 | 0.000 |
| Gm10578 | 0.000 | 0.001 | 0.007 |
| Gm45397 | 0.000 | 0.000 | 0.002 |
| Gm45682 | 0.000 | 0.000 | 0.000 |
| Tcerg1l | 0.000 | 0.000 | 0.000 |
| Gm45507 | 0.000 | 0.000 | 0.000 |
| Gm32486 | 0.000 | 0.000 | 0.000 |
| Nkx6-2 | 0.000 | 0.001 | 0.001 |
| Adgra1 | 0.000 | 0.000 | 0.001 |
| Kndc1 | 0.000 | 0.006 | 0.001 |
| Utf1 | 0.000 | 0.000 | 0.000 |
| Prap1 | 0.000 | 0.000 | 0.000 |
| Sprn | 0.000 | 0.002 | 0.005 |
| Olfr523 | 0.000 | 0.000 | 0.000 |
| Cd163l1 | 0.000 | 0.000 | 0.000 |
| Olfr531 | 0.000 | 0.000 | 0.000 |
| Olfr535 | 0.000 | 0.000 | 0.000 |

| | | | |
|---|---|---|---|
| Olfr45 | 0.000 | 0.000 | 0.000 |
| Urah | 0.000 | 0.001 | 0.001 |
| C330022C24R | 0.000 | 0.000 | 0.000 |
| Scgb1c1 | 0.000 | 0.000 | 0.000 |
| Odf3 | 0.000 | 0.000 | 0.000 |
| Nlrp6 | 0.000 | 0.000 | 0.000 |
| Gm45717 | 0.000 | 0.000 | 0.000 |
| Ano9 | 0.000 | 0.000 | 0.000 |
| Lmntd2 | 0.000 | 0.000 | 0.000 |
| Cdhr5 | 0.000 | 0.000 | 0.000 |
| Drd4 | 0.000 | 0.000 | 0.001 |
| Gm16982 | 0.000 | 0.001 | 0.001 |
| Muc5ac | 0.000 | 0.000 | 0.000 |
| Muc5b | 0.000 | 0.000 | 0.000 |
| Krtap5-5 | 0.000 | 0.000 | 0.000 |
| Gm45889 | 0.000 | 0.000 | 0.001 |
| Prr33 | 0.000 | 0.001 | 0.001 |
| Gm49394 | 0.000 | 0.000 | 0.000 |
| Gm6471 | 0.000 | 0.000 | 0.000 |
| Gm39117 | 0.000 | 0.000 | 0.000 |
| Th | 0.000 | 0.000 | 0.000 |
| Ascl2 | 0.000 | 0.002 | 0.004 |
| Tspan32os | 0.000 | 0.000 | 0.000 |
| 4933417O13I | 0.000 | 0.000 | 0.004 |
| Gm44732 | 0.000 | 0.000 | 0.001 |
| Gm15579 | 0.000 | 0.001 | 0.001 |
| Mrgprg | 0.000 | 0.000 | 0.001 |
| Shank2 | 0.000 | 0.001 | 0.007 |
| Gm44930 | 0.000 | 0.000 | 0.000 |
| Gm44929 | 0.000 | 0.000 | 0.000 |
| Gm26793 | 0.000 | 0.000 | 0.000 |
| Fgf15 | 0.000 | 0.000 | 0.000 |
| Gm39119 | 0.000 | 0.000 | 0.000 |
| Mrgprd | 0.000 | 0.000 | 0.000 |
| Iyd | 0.000 | 0.008 | 0.009 |
| Gm30570 | 0.000 | 0.000 | 0.000 |
| Gm10097 | 0.000 | 0.002 | 0.000 |
| Oprm1 | 0.000 | 0.000 | 0.000 |
| Gm48614 | 0.000 | 0.000 | 0.000 |
| Gm30906 | 0.000 | 0.000 | 0.000 |
| Gm48728 | 0.000 | 0.000 | 0.000 |
| Gm48748 | 0.000 | 0.000 | 0.000 |
| Gm48759 | 0.000 | 0.000 | 0.000 |
| Gm46210 | 0.000 | 0.000 | 0.000 |
| Gm48278 | 0.000 | 0.000 | 0.000 |
| Gm48279 | 0.000 | 0.000 | 0.000 |
| Adgb | 0.000 | 0.000 | 0.000 |
| Gm16577 | 0.000 | 0.000 | 0.000 |

| | | | |
|---|---|---|---|
| Gm48697 | 0.000 | 0.000 | 0.000 |
| Gm48721 | 0.000 | 0.000 | 0.000 |
| Gm48724 | 0.000 | 0.000 | 0.000 |
| Zc2hc1b | 0.000 | 0.000 | 0.000 |
| Gm32172 | 0.000 | 0.000 | 0.000 |
| Gm32105 | 0.000 | 0.000 | 0.000 |
| Gm26835 | 0.000 | 0.001 | 0.001 |
| Gm28289 | 0.000 | 0.000 | 0.000 |
| Gm48843 | 0.000 | 0.000 | 0.000 |
| Gm16234 | 0.000 | 0.000 | 0.000 |
| Nmbr | 0.000 | 0.000 | 0.000 |
| Gje1 | 0.000 | 0.000 | 0.000 |
| Gm47710 | 0.000 | 0.000 | 0.000 |
| Gm47769 | 0.000 | 0.000 | 0.000 |
| Ect2l | 0.000 | 0.005 | 0.004 |
| Gm48483 | 0.000 | 0.000 | 0.000 |
| Il22ra2 | 0.000 | 0.000 | 0.000 |
| AC153891.1 | 0.000 | 0.000 | 0.000 |
| 4930405J17R | 0.000 | 0.000 | 0.000 |
| Gm48249 | 0.000 | 0.000 | 0.000 |
| Gm20149 | 0.000 | 0.000 | 0.000 |
| Gm48189 | 0.000 | 0.000 | 0.000 |
| Gm26577 | 0.000 | 0.000 | 0.000 |
| Gm33619 | 0.000 | 0.000 | 0.000 |
| 4930455C13F | 0.000 | 0.000 | 0.000 |
| Aldh8a1 | 0.000 | 0.002 | 0.001 |
| 1700021A07F | 0.000 | 0.000 | 0.000 |
| Gm5420 | 0.000 | 0.000 | 0.000 |
| 4930444G20I | 0.000 | 0.000 | 0.000 |
| Gm10824 | 0.000 | 0.002 | 0.009 |
| Gm47843 | 0.000 | 0.001 | 0.000 |
| Vnn3 | 0.000 | 0.000 | 0.000 |
| Gm15271 | 0.000 | 0.001 | 0.000 |
| Gm47715 | 0.000 | 0.000 | 0.000 |
| Gm40617 | 0.000 | 0.002 | 0.002 |
| Smlr1 | 0.000 | 0.000 | 0.000 |
| Gm29571 | 0.000 | 0.000 | 0.000 |
| Gm36908 | 0.000 | 0.000 | 0.000 |
| 4930401C15F | 0.000 | 0.000 | 0.000 |
| Gm47939 | 0.000 | 0.000 | 0.000 |
| Gm9767 | 0.000 | 0.000 | 0.000 |
| Samd3 | 0.000 | 0.000 | 0.000 |
| Gm48084 | 0.000 | 0.000 | 0.000 |
| 4930579H20I | 0.000 | 0.000 | 0.000 |
| Themis | 0.000 | 0.000 | 0.001 |
| Gm49353 | 0.000 | 0.000 | 0.000 |
| 9330159F19F | 0.000 | 0.001 | 0.003 |
| Gm40621 | 0.000 | 0.000 | 0.000 |

| | | | |
|---|---|---|---|
| Gm48159 | 0.000 | 0.002 | 0.000 |
| Gm48215 | 0.000 | 0.000 | 0.000 |
| Gm48094 | 0.000 | 0.000 | 0.000 |
| Nkain2 | 0.000 | 0.002 | 0.001 |
| Clvs2 | 0.000 | 0.001 | 0.000 |
| Rsph4a | 0.000 | 0.001 | 0.000 |
| Trappc3l | 0.000 | 0.000 | 0.000 |
| Fam26d | 0.000 | 0.000 | 0.000 |
| Gm26564 | 0.000 | 0.000 | 0.000 |
| Hs3st5 | 0.000 | 0.003 | 0.010 |
| Gm31378 | 0.000 | 0.000 | 0.003 |
| Gm48197 | 0.000 | 0.000 | 0.000 |
| Rfpl4b | 0.000 | 0.000 | 0.000 |
| Wisp3 | 0.000 | 0.000 | 0.001 |
| D030034A15I | 0.000 | 0.000 | 0.000 |
| Gm31562 | 0.000 | 0.000 | 0.000 |
| Gm26824 | 0.000 | 0.000 | 0.000 |
| Mfsd4b2 | 0.000 | 0.000 | 0.000 |
| Mfsd4b5 | 0.000 | 0.001 | 0.000 |
| Mfsd4b1 | 0.000 | 0.009 | 0.001 |
| Gm47598 | 0.000 | 0.000 | 0.000 |
| A930033M14 | 0.000 | 0.000 | 0.000 |
| Cdk19os | 0.000 | 0.001 | 0.001 |
| Ddo | 0.000 | 0.005 | 0.002 |
| AC166165.1 | 0.000 | 0.000 | 0.000 |
| Ak9 | 0.000 | 0.000 | 0.000 |
| Gm17196 | 0.000 | 0.002 | 0.002 |
| BC048559 | 0.000 | 0.000 | 0.000 |
| Gm15200 | 0.000 | 0.000 | 0.000 |
| Gm15199 | 0.000 | 0.000 | 0.000 |
| Gm48671 | 0.000 | 0.000 | 0.000 |
| Gm47815 | 0.000 | 0.000 | 0.000 |
| Gm29246 | 0.000 | 0.000 | 0.000 |
| Gm47889 | 0.000 | 0.001 | 0.000 |
| F930017D23I | 0.000 | 0.000 | 0.000 |
| Gm47927 | 0.000 | 0.000 | 0.000 |
| Gm40634 | 0.000 | 0.000 | 0.000 |
| 4933404K13I | 0.000 | 0.000 | 0.000 |
| Gm35028 | 0.000 | 0.000 | 0.000 |
| Gm48065 | 0.000 | 0.000 | 0.000 |
| Gm35154 | 0.000 | 0.000 | 0.000 |
| Gm48066 | 0.000 | 0.000 | 0.000 |
| Gm47389 | 0.000 | 0.000 | 0.000 |
| Gm47392 | 0.000 | 0.000 | 0.000 |
| Gm48275 | 0.000 | 0.000 | 0.000 |
| Gm49339 | 0.000 | 0.000 | 0.000 |
| Gm40645 | 0.000 | 0.000 | 0.000 |
| Lilrb4a | 0.000 | 0.001 | 0.001 |

| | | | |
|---|---|---|---|
| Gm46224 | 0.000 | 0.000 | 0.000 |
| Fam162b | 0.000 | 0.000 | 0.000 |
| Vgll2 | 0.000 | 0.000 | 0.000 |
| Gm40649 | 0.000 | 0.000 | 0.000 |
| Gm36173 | 0.000 | 0.000 | 0.000 |
| Gm36229 | 0.000 | 0.000 | 0.001 |
| AC155941.1 | 0.000 | 0.000 | 0.000 |
| 4930452L12F | 0.000 | 0.000 | 0.000 |
| Gm36827 | 0.000 | 0.000 | 0.000 |
| Duxf3 | 0.000 | 0.000 | 0.000 |
| Pla2g12b | 0.000 | 0.000 | 0.000 |
| Gm17059 | 0.000 | 0.000 | 0.000 |
| Pldi | 0.000 | 0.000 | 0.000 |
| Gm26947 | 0.000 | 0.000 | 0.000 |
| Tbata | 0.000 | 0.000 | 0.005 |
| Nodal | 0.000 | 0.000 | 0.000 |
| Gm47595 | 0.000 | 0.000 | 0.000 |
| Tacr2 | 0.000 | 0.000 | 0.000 |
| Hkdc1 | 0.000 | 0.001 | 0.002 |
| 4930507D05I | 0.000 | 0.002 | 0.003 |
| Gm16135 | 0.000 | 0.000 | 0.000 |
| Mypn | 0.000 | 0.000 | 0.000 |
| Gm26789 | 0.000 | 0.004 | 0.002 |
| Gm47896 | 0.000 | 0.000 | 0.000 |
| Gm47897 | 0.000 | 0.000 | 0.000 |
| Gm47900 | 0.000 | 0.000 | 0.000 |
| 1110002J07R | 0.000 | 0.001 | 0.001 |
| Gm26576 | 0.000 | 0.000 | 0.000 |
| AC155712.1 | 0.000 | 0.000 | 0.000 |
| Gm32023 | 0.000 | 0.000 | 0.000 |
| Gm32255 | 0.000 | 0.000 | 0.000 |
| 4930563J15R | 0.000 | 0.000 | 0.000 |
| Gm32364 | 0.000 | 0.000 | 0.000 |
| AC127568.1 | 0.000 | 0.000 | 0.000 |
| Rtkn2 | 0.000 | 0.002 | 0.004 |
| Gm16212 | 0.000 | 0.000 | 0.000 |
| Gm47078 | 0.000 | 0.000 | 0.000 |
| Gm33035 | 0.000 | 0.000 | 0.000 |
| Gm40685 | 0.000 | 0.002 | 0.001 |
| Phyhipl | 0.000 | 0.003 | 0.004 |
| Gm33979 | 0.000 | 0.000 | 0.000 |
| Gm48022 | 0.000 | 0.000 | 0.000 |
| Gm48023 | 0.000 | 0.000 | 0.000 |
| Gm48024 | 0.000 | 0.000 | 0.002 |
| 4930533K18F | 0.000 | 0.000 | 0.003 |
| Gm15647 | 0.000 | 0.000 | 0.000 |
| C730027H18I | 0.000 | 0.000 | 0.000 |
| AC079680.2 | 0.000 | 0.000 | 0.000 |

| | | | |
|---|---|---|---|
| AC079680.1 | 0.000 | 0.000 | 0.000 |
| 4930551I15R | 0.000 | 0.000 | 0.000 |
| A330049N07I | 0.000 | 0.000 | 0.000 |
| Rsph14 | 0.000 | 0.000 | 0.000 |
| Gm867 | 0.000 | 0.000 | 0.000 |
| Vpreb3 | 0.000 | 0.000 | 0.000 |
| Slc5a4b | 0.000 | 0.000 | 0.000 |
| 1700094J05R | 0.000 | 0.000 | 0.000 |
| 2610028H24I | 0.000 | 0.001 | 0.000 |
| Spatc1l | 0.000 | 0.000 | 0.000 |
| Ftcd | 0.000 | 0.000 | 0.000 |
| Gm35608 | 0.000 | 0.000 | 0.000 |
| Gm48276 | 0.000 | 0.000 | 0.000 |
| Gm3137 | 0.000 | 0.000 | 0.000 |
| Gm49325 | 0.000 | 0.000 | 0.000 |
| Tspear | 0.000 | 0.000 | 0.000 |
| 1700009J07R | 0.000 | 0.000 | 0.000 |
| Dnmt3l | 0.000 | 0.000 | 0.002 |
| 4921516A02F | 0.000 | 0.001 | 0.001 |
| Olfr1358 | 0.000 | 0.000 | 0.000 |
| Olfr1357 | 0.000 | 0.000 | 0.000 |
| Olfr1356 | 0.000 | 0.000 | 0.001 |
| Olfr8 | 0.000 | 0.000 | 0.000 |
| Gm47343 | 0.000 | 0.000 | 0.000 |
| Olfr1351 | 0.000 | 0.000 | 0.000 |
| Gm47015 | 0.000 | 0.000 | 0.000 |
| C2cd4c | 0.000 | 0.001 | 0.001 |
| Odf3l2 | 0.000 | 0.000 | 0.000 |
| Fgf22 | 0.000 | 0.002 | 0.003 |
| Gm48495 | 0.000 | 0.000 | 0.000 |
| Misp | 0.000 | 0.002 | 0.002 |
| Gm17134 | 0.000 | 0.000 | 0.000 |
| Elane | 0.000 | 0.000 | 0.000 |
| Cfd | 0.000 | 0.000 | 0.000 |
| Gm26602 | 0.000 | 0.000 | 0.000 |
| Plk5 | 0.000 | 0.000 | 0.000 |
| Gm15124 | 0.000 | 0.000 | 0.000 |
| Lingo3 | 0.000 | 0.000 | 0.000 |
| Tmprss9 | 0.000 | 0.000 | 0.000 |
| Diras1 | 0.000 | 0.000 | 0.000 |
| Gm16099 | 0.000 | 0.000 | 0.000 |
| 4930442H23I | 0.000 | 0.000 | 0.000 |
| Nmrk2 | 0.000 | 0.002 | 0.002 |
| Atcay | 0.000 | 0.004 | 0.001 |
| Gm15917 | 0.000 | 0.000 | 0.000 |
| Gm4767 | 0.000 | 0.000 | 0.000 |
| AC158605.1 | 0.000 | 0.001 | 0.001 |
| Gm47561 | 0.000 | 0.000 | 0.000 |

| | | | |
|---|---|---|---|
| 1700028I16R | 0.000 | 0.000 | 0.000 |
| Gm40723 | 0.000 | 0.000 | 0.000 |
| Gm34184 | 0.000 | 0.000 | 0.000 |
| E230014E18R | 0.000 | 0.000 | 0.001 |
| Rfx4 | 0.000 | 0.000 | 0.000 |
| Ascl4 | 0.000 | 0.000 | 0.000 |
| Bpifc | 0.000 | 0.000 | 0.000 |
| A230060F14R | 0.000 | 0.000 | 0.000 |
| Gm48485 | 0.000 | 0.000 | 0.000 |
| Gm16268 | 0.000 | 0.000 | 0.000 |
| Stab2 | 0.000 | 0.001 | 0.001 |
| Pah | 0.000 | 0.000 | 0.000 |
| Igf1os | 0.000 | 0.000 | 0.001 |
| Gm48306 | 0.000 | 0.000 | 0.001 |
| Pmch | 0.000 | 0.000 | 0.000 |
| Gm26546 | 0.000 | 0.000 | 0.000 |
| Gm16235 | 0.000 | 0.000 | 0.000 |
| Mybpc1 | 0.000 | 0.000 | 0.000 |
| Spic | 0.000 | 0.000 | 0.001 |
| Slc5a8 | 0.000 | 0.000 | 0.001 |
| 4930534H03I | 0.000 | 0.000 | 0.000 |
| Gm47082 | 0.000 | 0.000 | 0.001 |
| Gm31592 | 0.000 | 0.000 | 0.000 |
| 4930401A07F | 0.000 | 0.000 | 0.000 |
| Hal | 0.000 | 0.001 | 0.000 |
| Ccdc38 | 0.000 | 0.001 | 0.000 |
| Gm15963 | 0.000 | 0.000 | 0.000 |
| Usp44 | 0.000 | 0.000 | 0.000 |
| Gm4792 | 0.000 | 0.000 | 0.000 |
| Gm48657 | 0.000 | 0.000 | 0.001 |
| Gm48689 | 0.000 | 0.000 | 0.000 |
| Gm47599 | 0.000 | 0.000 | 0.000 |
| Gm33336 | 0.000 | 0.000 | 0.000 |
| Anapc15-ps | 0.000 | 0.000 | 0.000 |
| 1110019B22F | 0.000 | 0.000 | 0.000 |
| Gm47708 | 0.000 | 0.001 | 0.000 |
| 4732465J04R | 0.000 | 0.000 | 0.000 |
| Gm47718 | 0.000 | 0.000 | 0.000 |
| Gm33543 | 0.000 | 0.000 | 0.000 |
| 4930459C07F | 0.000 | 0.000 | 0.000 |
| Gm48427 | 0.000 | 0.000 | 0.000 |
| Gm33782 | 0.000 | 0.000 | 0.000 |
| Gm48505 | 0.000 | 0.000 | 0.000 |
| Gm48521 | 0.000 | 0.000 | 0.000 |
| 4930556N09I | 0.000 | 0.000 | 0.000 |
| 4930473O22I | 0.000 | 0.000 | 0.000 |
| Kera | 0.000 | 0.000 | 0.004 |
| Gm10754 | 0.000 | 0.000 | 0.000 |

| | | | |
|---|---|---|---|
| Ccer1 | 0.000 | 0.000 | 0.000 |
| 4930525C09F | 0.000 | 0.000 | 0.000 |
| C030005K15F | 0.000 | 0.000 | 0.000 |
| Gm8633 | 0.000 | 0.000 | 0.000 |
| Gm34297 | 0.000 | 0.000 | 0.000 |
| Gm5427 | 0.000 | 0.000 | 0.000 |
| Phxr2 | 0.000 | 0.000 | 0.001 |
| Gm16239 | 0.000 | 0.000 | 0.000 |
| Gm34777 | 0.000 | 0.000 | 0.000 |
| B530045E10F | 0.000 | 0.000 | 0.000 |
| Gm34983 | 0.000 | 0.000 | 0.000 |
| Gm35206 | 0.000 | 0.000 | 0.000 |
| Mgat4c | 0.000 | 0.000 | 0.001 |
| Alx1 | 0.000 | 0.003 | 0.016 |
| Gm17028 | 0.000 | 0.000 | 0.000 |
| Gm47221 | 0.000 | 0.000 | 0.000 |
| C230021G24I | 0.000 | 0.001 | 0.001 |
| 4930532I03R | 0.000 | 0.000 | 0.000 |
| Gm29674 | 0.000 | 0.000 | 0.001 |
| Myf5 | 0.000 | 0.000 | 0.000 |
| Myf6 | 0.000 | 0.000 | 0.000 |
| Gm29685 | 0.000 | 0.000 | 0.000 |
| Gm5136 | 0.000 | 0.007 | 0.004 |
| 1700020G17I | 0.000 | 0.000 | 0.000 |
| Gm28592 | 0.000 | 0.001 | 0.001 |
| Gm30262 | 0.000 | 0.000 | 0.000 |
| Gm30624 | 0.000 | 0.000 | 0.000 |
| Gm47865 | 0.000 | 0.000 | 0.000 |
| Trhde | 0.000 | 0.001 | 0.001 |
| Gm31182 | 0.000 | 0.000 | 0.000 |
| Gm26579 | 0.000 | 0.000 | 0.000 |
| 1700030O20I | 0.000 | 0.000 | 0.000 |
| D630029K05I | 0.000 | 0.001 | 0.000 |
| Best3 | 0.000 | 0.000 | 0.000 |
| Gm10747 | 0.000 | 0.000 | 0.000 |
| 9530003J23R | 0.000 | 0.000 | 0.000 |
| Lyz1 | 0.000 | 0.000 | 0.000 |
| Gm48903 | 0.000 | 0.000 | 0.000 |
| Gm32552 | 0.000 | 0.000 | 0.000 |
| Gm40773 | 0.000 | 0.000 | 0.000 |
| 5330439M10 | 0.000 | 0.001 | 0.000 |
| Gm47958 | 0.000 | 0.000 | 0.000 |
| Il22 | 0.000 | 0.000 | 0.000 |
| Gm32717 | 0.000 | 0.000 | 0.000 |
| Gm32802 | 0.000 | 0.000 | 0.000 |
| Gm9049 | 0.000 | 0.000 | 0.000 |
| Ifng | 0.000 | 0.000 | 0.000 |
| Gm47425 | 0.000 | 0.000 | 0.000 |

| | | | |
|---|---|---|---|
| Gm4065 | 0.000 | 0.000 | 0.000 |
| Gm38403 | 0.000 | 0.000 | 0.002 |
| Gm34045 | 0.000 | 0.000 | 0.000 |
| 4930477N07I | 0.000 | 0.000 | 0.000 |
| Grip1os3 | 0.000 | 0.000 | 0.000 |
| Gm47027 | 0.000 | 0.000 | 0.000 |
| Grip1os1 | 0.000 | 0.000 | 0.000 |
| 9230105E05F | 0.000 | 0.000 | 0.000 |
| 4930471E19F | 0.000 | 0.000 | 0.000 |
| 4921513I03R | 0.000 | 0.001 | 0.000 |
| Gm40787 | 0.000 | 0.000 | 0.000 |
| D930020B18I | 0.000 | 0.000 | 0.000 |
| Gm4489 | 0.000 | 0.000 | 0.000 |
| Gm36041 | 0.000 | 0.000 | 0.001 |
| A130077B15F | 0.000 | 0.001 | 0.001 |
| 4930484H19I | 0.000 | 0.000 | 0.000 |
| Gm49335 | 0.000 | 0.001 | 0.001 |
| Gm47972 | 0.000 | 0.001 | 0.003 |
| Cyp27b1 | 0.000 | 0.001 | 0.002 |
| Agap2 | 0.000 | 0.001 | 0.001 |
| Gm47200 | 0.000 | 0.000 | 0.000 |
| Inhbe | 0.000 | 0.000 | 0.000 |
| Inhbc | 0.000 | 0.000 | 0.000 |
| Gm16217 | 0.000 | 0.000 | 0.000 |
| Gm16229 | 0.000 | 0.001 | 0.000 |
| Rdh1 | 0.000 | 0.000 | 0.001 |
| Rdh9 | 0.000 | 0.000 | 0.000 |
| Rdh16f2 | 0.000 | 0.000 | 0.000 |
| Mip | 0.000 | 0.000 | 0.000 |
| Apon | 0.000 | 0.000 | 0.000 |
| Apof | 0.000 | 0.000 | 0.000 |
| Gm17201 | 0.000 | 0.001 | 0.000 |
| Slc39a5 | 0.000 | 0.000 | 0.000 |
| Gm29585 | 0.000 | 0.000 | 0.000 |
| Gm47540 | 0.000 | 0.000 | 0.000 |
| AC117232.1 | 0.000 | 0.000 | 0.000 |
| Olfr767 | 0.000 | 0.000 | 0.000 |
| Olfr788 | 0.000 | 0.000 | 0.000 |
| Olfr802 | 0.000 | 0.000 | 0.000 |
| Olfr808 | 0.000 | 0.000 | 0.000 |
| Gm10310 | 0.000 | 0.000 | 0.000 |
| Olfr822 | 0.000 | 0.000 | 0.000 |
| Olfr825 | 0.000 | 0.000 | 0.000 |
| Tespa1 | 0.000 | 0.000 | 0.000 |
| Vmn2r85 | 0.000 | 0.000 | 0.000 |
| Vmn2r86 | 0.000 | 0.000 | 0.000 |
| Vmn2r87 | 0.000 | 0.000 | 0.000 |
| Pcp2 | 0.000 | 0.000 | 0.000 |

| | | | |
|---|---|---|---|
| Fcer2a | 0.000 | 0.000 | 0.000 |
| Cd209a | 0.000 | 0.000 | 0.000 |
| Cd209f | 0.000 | 0.000 | 0.000 |
| Cd209g | 0.000 | 0.000 | 0.000 |
| 4932443L11F | 0.000 | 0.000 | 0.000 |
| Gm6410 | 0.000 | 0.001 | 0.001 |
| Slc10a2 | 0.000 | 0.000 | 0.000 |
| D530014G21I | 0.000 | 0.000 | 0.000 |
| 4930453L07F | 0.000 | 0.000 | 0.000 |
| Gm44516 | 0.000 | 0.000 | 0.000 |
| Gm10067 | 0.000 | 0.000 | 0.000 |
| Gm39129 | 0.000 | 0.000 | 0.000 |
| Gm32540 | 0.000 | 0.000 | 0.000 |
| B930025P03I | 0.000 | 0.000 | 0.000 |
| Gm2814 | 0.000 | 0.000 | 0.002 |
| Gm44955 | 0.000 | 0.000 | 0.000 |
| Gm44956 | 0.000 | 0.001 | 0.001 |
| B830042I05R | 0.000 | 0.000 | 0.000 |
| Gm44717 | 0.000 | 0.000 | 0.000 |
| Gm15418 | 0.000 | 0.000 | 0.000 |
| 1700016D06I | 0.000 | 0.000 | 0.000 |
| Tex29 | 0.000 | 0.001 | 0.003 |
| Gm39132 | 0.000 | 0.000 | 0.000 |
| B020031H02I | 0.000 | 0.000 | 0.000 |
| Gm29175 | 0.000 | 0.000 | 0.000 |
| Gm15347 | 0.000 | 0.000 | 0.000 |
| Gm15351 | 0.000 | 0.000 | 0.000 |
| Proz | 0.000 | 0.000 | 0.000 |
| Gm17023 | 0.000 | 0.000 | 0.000 |
| Atp4b | 0.000 | 0.000 | 0.000 |
| 1700029H14I | 0.000 | 0.001 | 0.000 |
| 4932443I19R | 0.000 | 0.000 | 0.000 |
| AF366264 | 0.000 | 0.000 | 0.000 |
| Gm5907 | 0.000 | 0.000 | 0.000 |
| Dlgap2 | 0.000 | 0.001 | 0.000 |
| Gm16350 | 0.000 | 0.000 | 0.000 |
| Csmd1 | 0.000 | 0.000 | 0.000 |
| Gm16346 | 0.000 | 0.000 | 0.000 |
| Gm16725 | 0.000 | 0.001 | 0.000 |
| Gm35934 | 0.000 | 0.000 | 0.000 |
| 4930467E23F | 0.000 | 0.000 | 0.000 |
| Gm15319 | 0.000 | 0.000 | 0.000 |
| Gm14851 | 0.000 | 0.000 | 0.000 |
| AY761184 | 0.000 | 0.000 | 0.000 |
| Defb1 | 0.000 | 0.001 | 0.000 |
| Defb50 | 0.000 | 0.000 | 0.000 |
| Defb2 | 0.000 | 0.001 | 0.000 |
| Defb9 | 0.000 | 0.000 | 0.000 |

| | | | |
|---|---|---|---|
| Defb11 | 0.000 | 0.000 | 0.000 |
| Defb15 | 0.000 | 0.000 | 0.000 |
| Atp7b | 0.000 | 0.000 | 0.000 |
| Tpte | 0.000 | 0.000 | 0.000 |
| Dkk4 | 0.000 | 0.000 | 0.000 |
| Gm15346 | 0.000 | 0.000 | 0.000 |
| Nkx6-3 | 0.000 | 0.000 | 0.000 |
| Zmat4 | 0.000 | 0.001 | 0.002 |
| Gm45163 | 0.000 | 0.001 | 0.000 |
| Gm30692 | 0.000 | 0.000 | 0.000 |
| Adam3 | 0.000 | 0.000 | 0.000 |
| 5430421F17F | 0.000 | 0.000 | 0.000 |
| Gm45237 | 0.000 | 0.001 | 0.000 |
| Gm31045 | 0.000 | 0.000 | 0.000 |
| Gm16159 | 0.000 | 0.000 | 0.000 |
| Kcnu1 | 0.000 | 0.000 | 0.001 |
| Gm45411 | 0.000 | 0.000 | 0.000 |
| Gm31784 | 0.000 | 0.000 | 0.000 |
| Gm45307 | 0.000 | 0.000 | 0.000 |
| Gm31898 | 0.000 | 0.000 | 0.000 |
| Gm45580 | 0.000 | 0.000 | 0.000 |
| Gm39149 | 0.000 | 0.000 | 0.000 |
| Gm32050 | 0.000 | 0.000 | 0.000 |
| Gm32098 | 0.000 | 0.000 | 0.000 |
| 2310008N11I | 0.000 | 0.000 | 0.000 |
| Gm45572 | 0.000 | 0.002 | 0.001 |
| Gm45371 | 0.000 | 0.000 | 0.001 |
| Gm45370 | 0.000 | 0.000 | 0.000 |
| Gm32389 | 0.000 | 0.001 | 0.000 |
| Gm45267 | 0.000 | 0.000 | 0.000 |
| Got1l1 | 0.000 | 0.000 | 0.000 |
| Adrb3 | 0.000 | 0.004 | 0.004 |
| Gm45470 | 0.000 | 0.001 | 0.001 |
| Tex24 | 0.000 | 0.000 | 0.000 |
| Gm45861 | 0.000 | 0.000 | 0.000 |
| Gm1698 | 0.000 | 0.000 | 0.000 |
| 7420700N18I | 0.000 | 0.000 | 0.000 |
| Gm45587 | 0.000 | 0.000 | 0.000 |
| Smim18 | 0.000 | 0.000 | 0.000 |
| Gm26768 | 0.000 | 0.001 | 0.001 |
| Mboat4 | 0.000 | 0.001 | 0.000 |
| Gm39158 | 0.000 | 0.000 | 0.000 |
| B930018H19I | 0.000 | 0.000 | 0.000 |
| Gm45627 | 0.000 | 0.000 | 0.000 |
| Gm34474 | 0.000 | 0.000 | 0.000 |
| Gm34597 | 0.000 | 0.000 | 0.000 |
| Gm19410 | 0.000 | 0.000 | 0.000 |
| 5430403N17I | 0.000 | 0.000 | 0.001 |

| | | | |
|---|---|---|---|
| 4933430A20R | 0.000 | 0.000 | 0.000 |
| G630064G18I | 0.000 | 0.000 | 0.001 |
| Adam24 | 0.000 | 0.000 | 0.000 |
| Adam39 | 0.000 | 0.000 | 0.000 |
| Gm26584 | 0.000 | 0.001 | 0.000 |
| AY512931 | 0.000 | 0.000 | 0.000 |
| Mtnr1a | 0.000 | 0.000 | 0.000 |
| Gm6329 | 0.000 | 0.000 | 0.000 |
| Lrp2bp | 0.000 | 0.001 | 0.001 |
| Helt | 0.000 | 0.000 | 0.000 |
| Gm39168 | 0.000 | 0.000 | 0.000 |
| Gm30931 | 0.000 | 0.000 | 0.000 |
| Gm45607 | 0.000 | 0.000 | 0.000 |
| Gm45479 | 0.000 | 0.000 | 0.000 |
| Gm31172 | 0.000 | 0.000 | 0.001 |
| 1700011L03R | 0.000 | 0.000 | 0.000 |
| E330018M18I | 0.000 | 0.000 | 0.000 |
| Gm31786 | 0.000 | 0.000 | 0.000 |
| Gm45864 | 0.000 | 0.001 | 0.001 |
| Cldn24 | 0.000 | 0.000 | 0.000 |
| 4930555F03R | 0.000 | 0.000 | 0.000 |
| Gm45322 | 0.000 | 0.000 | 0.000 |
| Gm45388 | 0.000 | 0.000 | 0.001 |
| Spata4 | 0.000 | 0.000 | 0.000 |
| Glra3 | 0.000 | 0.000 | 0.000 |
| Gm34099 | 0.000 | 0.001 | 0.001 |
| Gm45359 | 0.000 | 0.000 | 0.000 |
| Gm45356 | 0.000 | 0.000 | 0.000 |
| Galntl6 | 0.000 | 0.000 | 0.001 |
| Gm26905 | 0.000 | 0.000 | 0.000 |
| BC030500 | 0.000 | 0.000 | 0.000 |
| Aadat | 0.000 | 0.000 | 0.001 |
| Gm45650 | 0.000 | 0.000 | 0.000 |
| 1700001D01I | 0.000 | 0.000 | 0.000 |
| 4930512H18I | 0.000 | 0.000 | 0.000 |
| Gm45518 | 0.000 | 0.001 | 0.001 |
| Trim61 | 0.000 | 0.000 | 0.000 |
| BC030870 | 0.000 | 0.000 | 0.000 |
| Gm15356 | 0.000 | 0.000 | 0.000 |
| Gm32568 | 0.000 | 0.000 | 0.000 |
| Npy5r | 0.000 | 0.000 | 0.000 |
| Gm10660 | 0.000 | 0.000 | 0.000 |
| Nat3 | 0.000 | 0.000 | 0.000 |
| Gm26659 | 0.000 | 0.000 | 0.000 |
| 1700125H03I | 0.000 | 0.000 | 0.000 |
| Gm33103 | 0.000 | 0.000 | 0.000 |
| Gm15716 | 0.000 | 0.000 | 0.000 |
| Gm9495 | 0.000 | 0.000 | 0.000 |

| | | | |
|---|---|---|---|
| Gm43263 | 0.000 | 0.001 | 0.000 |
| Ncan | 0.000 | 0.000 | 0.000 |
| Cers1 | 0.000 | 0.001 | 0.002 |
| Tmem59l | 0.000 | 0.000 | 0.000 |
| Il12rb1 | 0.000 | 0.000 | 0.000 |
| Nxnl1 | 0.000 | 0.000 | 0.001 |
| Insl3 | 0.000 | 0.000 | 0.000 |
| Gm35572 | 0.000 | 0.000 | 0.000 |
| Olfr373 | 0.000 | 0.001 | 0.000 |
| Olfr374 | 0.000 | 0.000 | 0.000 |
| Cib3 | 0.000 | 0.002 | 0.000 |
| Isx | 0.000 | 0.000 | 0.000 |
| Gm45822 | 0.000 | 0.000 | 0.000 |
| Gm45899 | 0.000 | 0.000 | 0.000 |
| Gm10358 | 0.000 | 0.000 | 0.000 |
| Iqcm | 0.000 | 0.000 | 0.000 |
| Gm45406 | 0.000 | 0.001 | 0.001 |
| Gm29895 | 0.000 | 0.000 | 0.000 |
| Pou4f2 | 0.000 | 0.000 | 0.000 |
| Gm45326 | 0.000 | 0.000 | 0.000 |
| Gm30329 | 0.000 | 0.000 | 0.000 |
| Gm45601 | 0.000 | 0.000 | 0.001 |
| Gm45602 | 0.000 | 0.000 | 0.000 |
| Gm45435 | 0.000 | 0.000 | 0.000 |
| Gm45431 | 0.000 | 0.000 | 0.000 |
| Frem3 | 0.000 | 0.000 | 0.000 |
| Gm27048 | 0.000 | 0.000 | 0.002 |
| Gm38590 | 0.000 | 0.000 | 0.000 |
| Gm31545 | 0.000 | 0.000 | 0.000 |
| Gm45449 | 0.000 | 0.001 | 0.000 |
| Clgn | 0.000 | 0.000 | 0.000 |
| Gm10644 | 0.000 | 0.001 | 0.001 |
| Rln3 | 0.000 | 0.000 | 0.000 |
| Podnl1 | 0.000 | 0.000 | 0.001 |
| Best2 | 0.000 | 0.000 | 0.001 |
| Gm5741 | 0.000 | 0.000 | 0.000 |
| Gm45425 | 0.000 | 0.000 | 0.000 |
| 2010110E17F | 0.000 | 0.000 | 0.000 |
| Gm5118 | 0.000 | 0.000 | 0.000 |
| Gm10637 | 0.000 | 0.000 | 0.000 |
| Cbln1 | 0.000 | 0.001 | 0.026 |
| Gm45639 | 0.000 | 0.000 | 0.000 |
| Gm35850 | 0.000 | 0.000 | 0.000 |
| Gm35963 | 0.000 | 0.000 | 0.000 |
| Gm19935 | 0.000 | 0.000 | 0.001 |
| Gm45641 | 0.000 | 0.000 | 0.000 |
| Gm36325 | 0.000 | 0.000 | 0.000 |
| Gm36380 | 0.000 | 0.000 | 0.000 |

| | | | |
|---|---|---|---|
| Irx3os | 0.000 | 0.001 | 0.000 |
| Gm28515 | 0.000 | 0.000 | 0.000 |
| Gm36670 | 0.000 | 0.000 | 0.000 |
| Gm21817 | 0.000 | 0.000 | 0.000 |
| Gm45472 | 0.000 | 0.000 | 0.000 |
| Slc6a2 | 0.000 | 0.001 | 0.002 |
| Ces1a | 0.000 | 0.000 | 0.000 |
| Ces1d | 0.000 | 0.000 | 0.000 |
| Ces1f | 0.000 | 0.000 | 0.000 |
| Ces1g | 0.000 | 0.000 | 0.000 |
| Ces5a | 0.000 | 0.000 | 0.000 |
| Gm45708 | 0.000 | 0.000 | 0.000 |
| Gm30606 | 0.000 | 0.000 | 0.000 |
| 4930488L21R | 0.000 | 0.000 | 0.000 |
| Slc12a3 | 0.000 | 0.000 | 0.001 |
| Gm15889 | 0.000 | 0.000 | 0.000 |
| Gm15890 | 0.000 | 0.000 | 0.000 |
| 9330175E14R | 0.000 | 0.002 | 0.001 |
| Ccl22 | 0.000 | 0.000 | 0.000 |
| Drc7 | 0.000 | 0.000 | 0.000 |
| 4933406B17R | 0.000 | 0.000 | 0.000 |
| Gm31659 | 0.000 | 0.000 | 0.000 |
| Gm32005 | 0.000 | 0.000 | 0.000 |
| Gm15679 | 0.000 | 0.000 | 0.000 |
| Cdh8 | 0.000 | 0.005 | 0.004 |
| A330008L17R | 0.000 | 0.001 | 0.000 |
| Gm45391 | 0.000 | 0.000 | 0.000 |
| Gm32531 | 0.000 | 0.000 | 0.000 |
| 1600027J07R | 0.000 | 0.000 | 0.000 |
| Gm29682 | 0.000 | 0.000 | 0.000 |
| Gm10631 | 0.000 | 0.000 | 0.000 |
| Gm45892 | 0.000 | 0.000 | 0.000 |
| Cmtm2a | 0.000 | 0.000 | 0.000 |
| Cmtm1 | 0.000 | 0.000 | 0.000 |
| Cmtm2b | 0.000 | 0.000 | 0.000 |
| Cdh16 | 0.000 | 0.002 | 0.000 |
| 1700082M22 | 0.000 | 0.000 | 0.000 |
| Ces2a | 0.000 | 0.000 | 0.000 |
| Ces2f | 0.000 | 0.000 | 0.000 |
| Ces3b | 0.000 | 0.000 | 0.000 |
| Gm3830 | 0.000 | 0.003 | 0.019 |
| Hsf4 | 0.000 | 0.001 | 0.001 |
| Kctd19 | 0.000 | 0.000 | 0.001 |
| Carmil2 | 0.000 | 0.001 | 0.002 |
| 4933405L10R | 0.000 | 0.000 | 0.000 |
| Gm16156 | 0.000 | 0.001 | 0.001 |
| Gm45752 | 0.000 | 0.000 | 0.000 |
| Dpep3 | 0.000 | 0.000 | 0.000 |

| | | | |
|---|---|---|---|
| Dpep2nb | 0.000 | 0.000 | 0.000 |
| Gm26786 | 0.000 | 0.001 | 0.001 |
| 1110028F18F | 0.000 | 0.001 | 0.001 |
| Cdh1 | 0.000 | 0.001 | 0.000 |
| 5033426E14F | 0.000 | 0.000 | 0.000 |
| Gm39244 | 0.000 | 0.000 | 0.000 |
| Pmfbp1 | 0.000 | 0.001 | 0.001 |
| Pkd1l3 | 0.000 | 0.000 | 0.000 |
| Gm17344 | 0.000 | 0.001 | 0.001 |
| Marveld3 | 0.000 | 0.002 | 0.001 |
| Tat | 0.000 | 0.000 | 0.000 |
| 8030455M16 | 0.000 | 0.000 | 0.000 |
| Il34 | 0.000 | 0.002 | 0.006 |
| Gm15894 | 0.000 | 0.000 | 0.000 |
| Gm15895 | 0.000 | 0.000 | 0.000 |
| Clec18a | 0.000 | 0.000 | 0.000 |
| Ctrb1 | 0.000 | 0.000 | 0.000 |
| Chst5 | 0.000 | 0.000 | 0.000 |
| Gm45904 | 0.000 | 0.000 | 0.000 |
| Gm10280 | 0.000 | 0.000 | 0.000 |
| Syce1l | 0.000 | 0.000 | 0.000 |
| Gm30052 | 0.000 | 0.000 | 0.000 |
| Clec3a | 0.000 | 0.000 | 0.000 |
| Gm16116 | 0.000 | 0.001 | 0.000 |
| Gm16117 | 0.000 | 0.000 | 0.000 |
| Gm16118 | 0.000 | 0.002 | 0.007 |
| 4930488N15I | 0.000 | 0.000 | 0.000 |
| Gm15655 | 0.000 | 0.003 | 0.004 |
| Dynlrb2 | 0.000 | 0.001 | 0.001 |
| Pkd1l2 | 0.000 | 0.000 | 0.000 |
| Gm15395 | 0.000 | 0.000 | 0.000 |
| Hsd17b2 | 0.000 | 0.000 | 0.000 |
| Gm31774 | 0.000 | 0.000 | 0.000 |
| Necab2 | 0.000 | 0.000 | 0.001 |
| Dnaaf1 | 0.000 | 0.000 | 0.000 |
| Adad2 | 0.000 | 0.000 | 0.000 |
| Atp2c2 | 0.000 | 0.000 | 0.000 |
| Gm15684 | 0.000 | 0.000 | 0.000 |
| Fam92b | 0.000 | 0.000 | 0.000 |
| Gm26815 | 0.000 | 0.000 | 0.000 |
| Gm26747 | 0.000 | 0.000 | 0.000 |
| Gm26784 | 0.000 | 0.000 | 0.000 |
| 1700018B08F | 0.000 | 0.001 | 0.000 |
| Jph3 | 0.000 | 0.000 | 0.000 |
| Gm20406 | 0.000 | 0.000 | 0.000 |
| Gm26812 | 0.000 | 0.001 | 0.002 |
| Trhr2 | 0.000 | 0.000 | 0.000 |
| Gm20681 | 0.000 | 0.000 | 0.000 |

| | | | |
|---|---|---|---|
| Il17c | 0.000 | 0.000 | 0.000 |
| Gm10612 | 0.000 | 0.000 | 0.000 |
| A530010L16F | 0.000 | 0.000 | 0.000 |
| Gm45894 | 0.000 | 0.001 | 0.001 |
| Gm45842 | 0.000 | 0.000 | 0.000 |
| 4933417D19I | 0.000 | 0.001 | 0.001 |
| 4732419C18F | 0.000 | 0.001 | 0.003 |
| 1810008B01F | 0.000 | 0.001 | 0.000 |
| Gm15775 | 0.000 | 0.000 | 0.000 |
| Pgbd5 | 0.000 | 0.002 | 0.003 |
| Capn9 | 0.000 | 0.002 | 0.003 |
| Gm45709 | 0.000 | 0.000 | 0.000 |
| Gm16237 | 0.000 | 0.000 | 0.000 |
| Pcnx2 | 0.000 | 0.000 | 0.000 |
| A730098A19F | 0.000 | 0.000 | 0.000 |
| Map3k21 | 0.000 | 0.001 | 0.001 |
| Gm45805 | 0.000 | 0.001 | 0.000 |
| A630001O12I | 0.000 | 0.000 | 0.000 |
| Ccdc7a | 0.000 | 0.000 | 0.000 |
| Gm2897 | 0.000 | 0.000 | 0.000 |
| Gm5795 | 0.000 | 0.000 | 0.000 |
| 1700110I01R | 0.000 | 0.000 | 0.000 |
| Gm2956 | 0.000 | 0.000 | 0.000 |
| Gm3002 | 0.000 | 0.000 | 0.000 |
| Gm5796 | 0.000 | 0.000 | 0.000 |
| Gm8108 | 0.000 | 0.000 | 0.000 |
| Gm2974 | 0.000 | 0.000 | 0.000 |
| 2610042L04F | 0.000 | 0.000 | 0.000 |
| Gm3164 | 0.000 | 0.002 | 0.000 |
| Gm3252 | 0.000 | 0.000 | 0.000 |
| Gm3264 | 0.000 | 0.000 | 0.000 |
| Gm3278 | 0.000 | 0.000 | 0.000 |
| Gm8281 | 0.000 | 0.000 | 0.000 |
| Gm8237 | 0.000 | 0.000 | 0.000 |
| Gm3453 | 0.000 | 0.000 | 0.000 |
| Gm3411 | 0.000 | 0.001 | 0.000 |
| Gm3591 | 0.000 | 0.000 | 0.000 |
| Gm8050 | 0.000 | 0.000 | 0.000 |
| Gm10406 | 0.000 | 0.000 | 0.001 |
| Gm3696 | 0.000 | 0.002 | 0.002 |
| Gm10339 | 0.000 | 0.000 | 0.000 |
| 4933406F09F | 0.000 | 0.000 | 0.000 |
| Gm3755 | 0.000 | 0.000 | 0.000 |
| Gm3752 | 0.000 | 0.000 | 0.000 |
| Gm3558 | 0.000 | 0.000 | 0.001 |
| Gm45521 | 0.000 | 0.000 | 0.000 |
| Dnase1l3 | 0.000 | 0.000 | 0.000 |
| Oit1 | 0.000 | 0.000 | 0.000 |

| | | | |
|---|---|---|---|
| 4930455B14F | 0.000 | 0.000 | 0.000 |
| Gm48439 | 0.000 | 0.000 | 0.000 |
| Fhitos | 0.000 | 0.000 | 0.000 |
| Gm3848 | 0.000 | 0.000 | 0.000 |
| Gm49355 | 0.000 | 0.000 | 0.000 |
| Gm48239 | 0.000 | 0.000 | 0.000 |
| Sntn | 0.000 | 0.000 | 0.000 |
| Gm281 | 0.000 | 0.000 | 0.000 |
| Olfr720 | 0.000 | 0.000 | 0.000 |
| Gm48857 | 0.000 | 0.000 | 0.000 |
| Olfr31 | 0.000 | 0.000 | 0.000 |
| Olfr721-ps1 | 0.000 | 0.000 | 0.000 |
| Nek10 | 0.000 | 0.000 | 0.000 |
| AC117608.1 | 0.000 | 0.000 | 0.000 |
| Gm31804 | 0.000 | 0.000 | 0.000 |
| 1700062C10F | 0.000 | 0.000 | 0.000 |
| Gm47794 | 0.000 | 0.000 | 0.000 |
| Gm48162 | 0.000 | 0.000 | 0.000 |
| Gm48321 | 0.000 | 0.001 | 0.000 |
| Gm2244 | 0.000 | 0.000 | 0.000 |
| Gm5458 | 0.000 | 0.000 | 0.000 |
| D14Ertd670e | 0.000 | 0.000 | 0.000 |
| Gm48283 | 0.000 | 0.000 | 0.000 |
| Gm48395 | 0.000 | 0.000 | 0.000 |
| Gm30054 | 0.000 | 0.001 | 0.001 |
| 7330404K18F | 0.000 | 0.000 | 0.000 |
| Gm15935 | 0.000 | 0.000 | 0.000 |
| 4931407E12F | 0.000 | 0.000 | 0.000 |
| Gm7480 | 0.000 | 0.000 | 0.000 |
| Gm10248 | 0.000 | 0.000 | 0.000 |
| CT025624.1 | 0.000 | 0.000 | 0.000 |
| E330034G19F | 0.000 | 0.000 | 0.000 |
| Gm38408 | 0.000 | 0.000 | 0.000 |
| Gm17747 | 0.000 | 0.000 | 0.000 |
| Ppifos | 0.000 | 0.001 | 0.001 |
| Anxa11os | 0.000 | 0.000 | 0.000 |
| Duxbl1 | 0.000 | 0.002 | 0.003 |
| Cphx2 | 0.000 | 0.000 | 0.000 |
| Cphx3 | 0.000 | 0.000 | 0.000 |
| Duxbl2 | 0.000 | 0.000 | 0.000 |
| Gm34059 | 0.000 | 0.000 | 0.000 |
| 4933413J09R | 0.000 | 0.002 | 0.001 |
| Dnah12 | 0.000 | 0.002 | 0.002 |
| Asb14 | 0.000 | 0.000 | 0.001 |
| Hesx1 | 0.000 | 0.000 | 0.001 |
| Gm45645 | 0.000 | 0.000 | 0.000 |
| Gm41118 | 0.000 | 0.000 | 0.000 |
| Gm47414 | 0.000 | 0.000 | 0.000 |

| | | | |
|---|---|---|---|
| Il17rb | 0.000 | 0.002 | 0.003 |
| AC154646.1 | 0.000 | 0.002 | 0.001 |
| 1700087M22 | 0.000 | 0.000 | 0.000 |
| Gm49217 | 0.000 | 0.000 | 0.000 |
| Colq | 0.000 | 0.000 | 0.001 |
| Gm49387 | 0.000 | 0.000 | 0.000 |
| D830044D21I | 0.000 | 0.000 | 0.000 |
| Msmb | 0.000 | 0.001 | 0.001 |
| Gm17210 | 0.000 | 0.000 | 0.000 |
| Chat | 0.000 | 0.000 | 0.000 |
| Slc18a3 | 0.000 | 0.000 | 0.000 |
| Gm2990 | 0.000 | 0.000 | 0.000 |
| Prrxl1 | 0.000 | 0.000 | 0.001 |
| 3425401B19I | 0.000 | 0.003 | 0.005 |
| Gm28651 | 0.000 | 0.000 | 0.000 |
| Fam170b | 0.000 | 0.000 | 0.001 |
| Gm15512 | 0.000 | 0.001 | 0.000 |
| Gm49108 | 0.000 | 0.000 | 0.000 |
| Gm30083 | 0.000 | 0.000 | 0.000 |
| Gm30556 | 0.000 | 0.001 | 0.000 |
| Npy4r | 0.000 | 0.002 | 0.000 |
| Fam25c | 0.000 | 0.000 | 0.000 |
| Gm17110 | 0.000 | 0.000 | 0.000 |
| 9230112D13I | 0.000 | 0.000 | 0.000 |
| Opn4 | 0.000 | 0.000 | 0.000 |
| Gm49012 | 0.000 | 0.000 | 0.000 |
| Gm49025 | 0.000 | 0.000 | 0.000 |
| A930038B10I | 0.000 | 0.001 | 0.001 |
| Lrit2 | 0.000 | 0.000 | 0.000 |
| Nrg3 | 0.000 | 0.000 | 0.001 |
| Dydc2 | 0.000 | 0.000 | 0.000 |
| Mat1a | 0.000 | 0.000 | 0.000 |
| Sftpa1 | 0.000 | 0.000 | 0.000 |
| Mbl1 | 0.000 | 0.000 | 0.000 |
| Sftpd | 0.000 | 0.000 | 0.000 |
| Gm2832 | 0.000 | 0.000 | 0.000 |
| Gm6401 | 0.000 | 0.000 | 0.000 |
| Gm8180 | 0.000 | 0.000 | 0.000 |
| Ear1 | 0.000 | 0.000 | 0.000 |
| Ang5 | 0.000 | 0.000 | 0.000 |
| Gm49087 | 0.000 | 0.000 | 0.000 |
| Gm49101 | 0.000 | 0.000 | 0.000 |
| 1700047E10I | 0.000 | 0.000 | 0.000 |
| Gm8212 | 0.000 | 0.000 | 0.000 |
| Gm8220 | 0.000 | 0.000 | 0.000 |
| Gm8229 | 0.000 | 0.000 | 0.000 |
| Ptgdr | 0.000 | 0.000 | 0.000 |
| Gm17173 | 0.000 | 0.000 | 0.000 |

| | | | |
|---|---|---|---|
| Ptger2 | 0.000 | 0.002 | 0.001 |
| Gm34250 | 0.000 | 0.000 | 0.000 |
| Gm41144 | 0.000 | 0.000 | 0.000 |
| Gm49124 | 0.000 | 0.000 | 0.000 |
| A530076I17R | 0.000 | 0.000 | 0.000 |
| 4933425B07F | 0.000 | 0.001 | 0.001 |
| D330046F09F | 0.000 | 0.002 | 0.001 |
| Gm48924 | 0.000 | 0.000 | 0.000 |
| Gm49150 | 0.000 | 0.000 | 0.000 |
| Gm49189 | 0.000 | 0.000 | 0.000 |
| Gm48975 | 0.000 | 0.000 | 0.000 |
| Gm49302 | 0.000 | 0.000 | 0.000 |
| Gm35360 | 0.000 | 0.000 | 0.000 |
| 4930447J18R | 0.000 | 0.000 | 0.000 |
| Gm49303 | 0.000 | 0.000 | 0.000 |
| Gm49304 | 0.000 | 0.000 | 0.000 |
| Gm49120 | 0.000 | 0.000 | 0.000 |
| Gm49151 | 0.000 | 0.000 | 0.000 |
| Otx2 | 0.000 | 0.000 | 0.000 |
| Otx2os1 | 0.000 | 0.000 | 0.000 |
| Gm48935 | 0.000 | 0.001 | 0.000 |
| 1700011H14I | 0.000 | 0.000 | 0.000 |
| Slc35f4 | 0.000 | 0.000 | 0.000 |
| Gm36146 | 0.000 | 0.000 | 0.000 |
| Olfr727 | 0.000 | 0.000 | 0.000 |
| Olfr729 | 0.000 | 0.000 | 0.000 |
| Olfr733 | 0.000 | 0.000 | 0.000 |
| Olfr734 | 0.000 | 0.000 | 0.000 |
| Olfr735 | 0.000 | 0.000 | 0.000 |
| Tlr11 | 0.000 | 0.000 | 0.000 |
| Olfr736 | 0.000 | 0.000 | 0.000 |
| Olfr739 | 0.000 | 0.000 | 0.000 |
| Olfr740 | 0.000 | 0.000 | 0.000 |
| Olfr741 | 0.000 | 0.000 | 0.000 |
| Olfr750 | 0.000 | 0.000 | 0.000 |
| Gm5622 | 0.000 | 0.000 | 0.000 |
| Gm5800 | 0.000 | 0.000 | 0.000 |
| Ang4 | 0.000 | 0.000 | 0.000 |
| Ear6 | 0.000 | 0.000 | 0.000 |
| Rnase13 | 0.000 | 0.000 | 0.000 |
| Gm49076 | 0.000 | 0.000 | 0.000 |
| Gm16617 | 0.000 | 0.000 | 0.000 |
| Gm49256 | 0.000 | 0.000 | 0.001 |
| Olfr221 | 0.000 | 0.000 | 0.000 |
| Olfr1507 | 0.000 | 0.000 | 0.000 |
| Gm30214 | 0.000 | 0.000 | 0.000 |
| Trav14d-3-dv8 | 0.000 | 0.000 | 0.000 |
| Trav4-4-dv10 | 0.000 | 0.000 | 0.000 |

| | | | |
|---|---|---|---|
| Trav6-7-dv9 | 0.000 | 0.000 | 0.000 |
| Trav17 | 0.000 | 0.000 | 0.000 |
| Trav19 | 0.000 | 0.000 | 0.000 |
| Trdv1 | 0.000 | 0.001 | 0.000 |
| Gm43647 | 0.000 | 0.000 | 0.000 |
| Trav22 | 0.000 | 0.002 | 0.000 |
| Trdv2-2 | 0.000 | 0.000 | 0.000 |
| Gm30275 | 0.000 | 0.000 | 0.002 |
| Trdv4 | 0.000 | 0.001 | 0.001 |
| Gm43434 | 0.000 | 0.001 | 0.007 |
| 5330426L24F | 0.000 | 0.000 | 0.000 |
| Gm29776 | 0.000 | 0.000 | 0.000 |
| Cebpe | 0.000 | 0.000 | 0.000 |
| Rnf212b | 0.000 | 0.000 | 0.000 |
| Gm20521 | 0.000 | 0.000 | 0.000 |
| Il25 | 0.000 | 0.000 | 0.001 |
| Gm49130 | 0.000 | 0.000 | 0.000 |
| Gm31251 | 0.000 | 0.000 | 0.000 |
| Zfhx2os | 0.000 | 0.000 | 0.000 |
| Gm20687 | 0.000 | 0.000 | 0.000 |
| Thtpa | 0.000 | 0.000 | 0.001 |
| Cpne6 | 0.000 | 0.009 | 0.002 |
| Nrl | 0.000 | 0.000 | 0.000 |
| Gm10876 | 0.000 | 0.000 | 0.000 |
| A730061H03I | 0.000 | 0.000 | 0.000 |
| Gm15932 | 0.000 | 0.000 | 0.000 |
| Gm49378 | 0.000 | 0.000 | 0.000 |
| Gm49067 | 0.000 | 0.000 | 0.000 |
| Ltb4r2 | 0.000 | 0.003 | 0.002 |
| Mcpt1 | 0.000 | 0.000 | 0.000 |
| Mcpt2 | 0.000 | 0.000 | 0.000 |
| Mcpt8 | 0.000 | 0.000 | 0.000 |
| Ctsg | 0.000 | 0.000 | 0.000 |
| Gzme | 0.000 | 0.000 | 0.002 |
| Gzmc | 0.000 | 0.000 | 0.000 |
| Gzmb | 0.000 | 0.000 | 0.000 |
| Atp12a | 0.000 | 0.000 | 0.000 |
| Gm10873 | 0.000 | 0.000 | 0.000 |
| B020004C17F | 0.000 | 0.000 | 0.000 |
| Gja3 | 0.000 | 0.012 | 0.001 |
| Gjb2 | 0.000 | 0.001 | 0.000 |
| Gjb6 | 0.000 | 0.000 | 0.000 |
| Ift88os | 0.000 | 0.000 | 0.000 |
| Gm49361 | 0.000 | 0.000 | 0.000 |
| Gm33321 | 0.000 | 0.000 | 0.001 |
| Gm33472 | 0.000 | 0.000 | 0.000 |
| Gm48453 | 0.000 | 0.000 | 0.000 |
| Gm5142 | 0.000 | 0.000 | 0.000 |

| | | | |
|---|---|---|---|
| Gm6904 | 0.000 | 0.000 | 0.000 |
| Gm29266 | 0.000 | 0.000 | 0.000 |
| Mipepos | 0.000 | 0.000 | 0.000 |
| 1700109G14I | 0.000 | 0.000 | 0.000 |
| 4930556J02R | 0.000 | 0.000 | 0.000 |
| Gm26916 | 0.000 | 0.001 | 0.001 |
| Gm35419 | 0.000 | 0.000 | 0.000 |
| Gm4131 | 0.000 | 0.000 | 0.000 |
| Serpine3 | 0.000 | 0.000 | 0.000 |
| Defb48 | 0.000 | 0.000 | 0.000 |
| Defb47 | 0.000 | 0.000 | 0.000 |
| Defb43 | 0.000 | 0.000 | 0.000 |
| Defb30 | 0.000 | 0.000 | 0.000 |
| Gm49390 | 0.000 | 0.000 | 0.000 |
| Blk | 0.000 | 0.000 | 0.000 |
| Tdh | 0.000 | 0.000 | 0.000 |
| Gm17232 | 0.000 | 0.000 | 0.000 |
| Gm15918 | 0.000 | 0.000 | 0.000 |
| Prss55 | 0.000 | 0.000 | 0.000 |
| 4930471C04I | 0.000 | 0.000 | 0.000 |
| Gm17116 | 0.000 | 0.000 | 0.000 |
| 9630015K15I | 0.000 | 0.002 | 0.001 |
| Mir124a-1hg | 0.000 | 0.000 | 0.000 |
| Gm48433 | 0.000 | 0.000 | 0.000 |
| Gm48449 | 0.000 | 0.000 | 0.000 |
| Nuggc | 0.000 | 0.001 | 0.002 |
| Gm10032 | 0.000 | 0.000 | 0.000 |
| Pnma2 | 0.000 | 0.001 | 0.006 |
| Gm20675 | 0.000 | 0.000 | 0.001 |
| Gm6878 | 0.000 | 0.000 | 0.003 |
| 4930438E09I | 0.000 | 0.000 | 0.000 |
| Gm10860 | 0.000 | 0.000 | 0.000 |
| Gm38409 | 0.000 | 0.000 | 0.000 |
| Gm31107 | 0.000 | 0.000 | 0.000 |
| Nefm | 0.000 | 0.003 | 0.008 |
| Adamdec1 | 0.000 | 0.000 | 0.000 |
| Gm41192 | 0.000 | 0.000 | 0.000 |
| Nkx2-6 | 0.000 | 0.000 | 0.000 |
| Nkx3-1 | 0.000 | 0.004 | 0.000 |
| Synb | 0.000 | 0.000 | 0.000 |
| Entpd4 | 0.000 | 0.000 | 0.000 |
| Entpd4b | 0.000 | 0.000 | 0.000 |
| Gm27177 | 0.000 | 0.000 | 0.000 |
| Gm33524 | 0.000 | 0.000 | 0.000 |
| Fgf17 | 0.000 | 0.001 | 0.001 |
| 2410012E07I | 0.000 | 0.000 | 0.000 |
| Gm49289 | 0.000 | 0.000 | 0.000 |
| Gm9195 | 0.000 | 0.000 | 0.000 |

| | | | |
|---|---|---|---|
| Gm16549 | 0.000 | 0.000 | 0.000 |
| Cysltr2 | 0.000 | 0.000 | 0.000 |
| Gm17233 | 0.000 | 0.000 | 0.000 |
| Gm21750 | 0.000 | 0.000 | 0.000 |
| Gm49165 | 0.000 | 0.000 | 0.000 |
| Gm49171 | 0.000 | 0.000 | 0.000 |
| Gm10847 | 0.000 | 0.000 | 0.000 |
| Cpb2 | 0.000 | 0.000 | 0.000 |
| Siah3 | 0.000 | 0.000 | 0.001 |
| 2900040C04F | 0.000 | 0.000 | 0.000 |
| E130202H07F | 0.000 | 0.002 | 0.002 |
| 4930431P22F | 0.000 | 0.002 | 0.001 |
| Gm48969 | 0.000 | 0.001 | 0.001 |
| Gm6994 | 0.000 | 0.000 | 0.000 |
| Gm1587 | 0.000 | 0.000 | 0.000 |
| Tnfsf11 | 0.000 | 0.001 | 0.000 |
| Gm49015 | 0.000 | 0.000 | 0.000 |
| Gm4632 | 0.000 | 0.000 | 0.000 |
| Cnmd | 0.000 | 0.000 | 0.021 |
| 4930452G13I | 0.000 | 0.001 | 0.001 |
| Pcdh8 | 0.000 | 0.000 | 0.001 |
| Gm10845 | 0.000 | 0.000 | 0.000 |
| AC121960.2 | 0.000 | 0.000 | 0.000 |
| Olfm4 | 0.000 | 0.000 | 0.000 |
| Gm49048 | 0.000 | 0.000 | 0.000 |
| 9630013A20F | 0.000 | 0.000 | 0.000 |
| Gm32729 | 0.000 | 0.000 | 0.000 |
| Klhl1 | 0.000 | 0.000 | 0.000 |
| 4930517O19I | 0.000 | 0.000 | 0.000 |
| 6330576A10F | 0.000 | 0.000 | 0.000 |
| Gm9922 | 0.000 | 0.001 | 0.000 |
| AC102815.1 | 0.000 | 0.000 | 0.000 |
| Acod1 | 0.000 | 0.000 | 0.000 |
| Gm34907 | 0.000 | 0.000 | 0.000 |
| Scel | 0.000 | 0.000 | 0.000 |
| D130079A08I | 0.000 | 0.000 | 0.000 |
| D130009I18R | 0.000 | 0.000 | 0.000 |
| 5430440P10F | 0.000 | 0.000 | 0.000 |
| Gm35909 | 0.000 | 0.000 | 0.000 |
| Gm48970 | 0.000 | 0.000 | 0.000 |
| Trim52 | 0.000 | 0.000 | 0.000 |
| Gm36107 | 0.000 | 0.000 | 0.001 |
| Gm48992 | 0.000 | 0.000 | 0.000 |
| 1700100I10R | 0.000 | 0.000 | 0.000 |
| 4930435M08 | 0.000 | 0.000 | 0.000 |
| Gm26773 | 0.000 | 0.000 | 0.000 |
| 4933431J24R | 0.000 | 0.000 | 0.000 |
| Sox21 | 0.000 | 0.000 | 0.000 |

| | | | |
|---|---|---|---|
| Gm4675 | 0.000 | 0.000 | 0.000 |
| Gm9376 | 0.000 | 0.000 | 0.000 |
| Oxgr1 | 0.000 | 0.000 | 0.000 |
| Gm48997 | 0.000 | 0.000 | 0.000 |
| Gm17613 | 0.000 | 0.000 | 0.000 |
| Slc15a1 | 0.000 | 0.000 | 0.000 |
| Gm49035 | 0.000 | 0.000 | 0.000 |
| Gpr18 | 0.000 | 0.000 | 0.000 |
| 1700108J01R | 0.000 | 0.000 | 0.000 |
| Gm5089 | 0.000 | 0.000 | 0.000 |
| Gm10837 | 0.000 | 0.000 | 0.000 |
| Gm28932 | 0.000 | 0.000 | 0.000 |
| 1700024B18F | 0.000 | 0.000 | 0.000 |
| Gm10721 | 0.000 | 0.000 | 0.000 |
| Gm10718 | 0.000 | 0.000 | 0.000 |
| Mmp13 | 0.000 | 0.000 | 0.000 |
| Mmp12 | 0.000 | 0.001 | 0.001 |
| Mmp10 | 0.000 | 0.000 | 0.000 |
| Mmp8 | 0.000 | 0.000 | 0.000 |
| Gm16833 | 0.000 | 0.000 | 0.000 |
| Cntn5 | 0.000 | 0.000 | 0.000 |
| 4930568E12F | 0.000 | 0.000 | 0.000 |
| Gm47557 | 0.000 | 0.000 | 0.000 |
| 1700019J19R | 0.000 | 0.001 | 0.000 |
| Gm34069 | 0.000 | 0.001 | 0.000 |
| Gm47326 | 0.000 | 0.000 | 0.000 |
| Gm7607 | 0.000 | 0.000 | 0.000 |
| Piwil4 | 0.000 | 0.001 | 0.003 |
| Gm10706 | 0.000 | 0.000 | 0.000 |
| 1700012B09F | 0.000 | 0.000 | 0.000 |
| Gm16568 | 0.000 | 0.000 | 0.000 |
| Gm34973 | 0.000 | 0.000 | 0.000 |
| Mbd3l2 | 0.000 | 0.000 | 0.000 |
| Mbd3l1 | 0.000 | 0.000 | 0.000 |
| Olfr24 | 0.000 | 0.000 | 0.000 |
| Olfr839-ps1 | 0.000 | 0.000 | 0.000 |
| Olfr857 | 0.000 | 0.000 | 0.000 |
| Olfr859 | 0.000 | 0.000 | 0.000 |
| Olfr872 | 0.000 | 0.000 | 0.000 |
| Olfr39 | 0.000 | 0.000 | 0.000 |
| Olfr873 | 0.000 | 0.001 | 0.001 |
| Olfr18 | 0.000 | 0.000 | 0.001 |
| Gm26592 | 0.000 | 0.001 | 0.002 |
| Zglp1 | 0.000 | 0.002 | 0.002 |
| Gm38431 | 0.000 | 0.001 | 0.000 |
| Ap1m2 | 0.000 | 0.000 | 0.000 |
| Gm39302 | 0.000 | 0.001 | 0.001 |
| Gm16853 | 0.000 | 0.003 | 0.003 |

| | | | |
|---|---|---|---|
| Angptl8 | 0.000 | 0.000 | 0.001 |
| Ccdc151 | 0.000 | 0.001 | 0.001 |
| Npsr1 | 0.000 | 0.000 | 0.000 |
| Gm29824 | 0.000 | 0.000 | 0.000 |
| E130101E03R | 0.000 | 0.000 | 0.000 |
| Gm29642 | 0.000 | 0.001 | 0.000 |
| Gm48362 | 0.000 | 0.000 | 0.000 |
| Gm39312 | 0.000 | 0.000 | 0.000 |
| Gm48393 | 0.000 | 0.000 | 0.000 |
| B3gat1 | 0.000 | 0.001 | 0.000 |
| Glb1l3 | 0.000 | 0.000 | 0.000 |
| Gm1110 | 0.000 | 0.000 | 0.000 |
| Gm48803 | 0.000 | 0.000 | 0.000 |
| Spata19 | 0.000 | 0.000 | 0.000 |
| Gm15520 | 0.000 | 0.000 | 0.001 |
| Gm31013 | 0.000 | 0.000 | 0.000 |
| Tmem45b | 0.000 | 0.000 | 0.000 |
| Gm39317 | 0.000 | 0.000 | 0.000 |
| Gm27201 | 0.000 | 0.000 | 0.000 |
| Gm27240 | 0.000 | 0.000 | 0.000 |
| Gm27166 | 0.000 | 0.000 | 0.000 |
| Gm47775 | 0.000 | 0.000 | 0.000 |
| Gm47785 | 0.000 | 0.000 | 0.000 |
| Gm39318 | 0.000 | 0.000 | 0.000 |
| Gm33838 | 0.000 | 0.003 | 0.003 |
| Gm1113 | 0.000 | 0.000 | 0.000 |
| Pate4 | 0.000 | 0.000 | 0.000 |
| Pate1 | 0.000 | 0.000 | 0.000 |
| D730048I06R | 0.000 | 0.000 | 0.000 |
| 9230113P08F | 0.000 | 0.000 | 0.000 |
| Gm26787 | 0.000 | 0.000 | 0.000 |
| Gm3896 | 0.000 | 0.000 | 0.000 |
| Gm48081 | 0.000 | 0.000 | 0.000 |
| Gm47633 | 0.000 | 0.000 | 0.000 |
| Gm34885 | 0.000 | 0.001 | 0.001 |
| Robo3 | 0.000 | 0.000 | 0.000 |
| Olfr160 | 0.000 | 0.000 | 0.000 |
| Olfr877 | 0.000 | 0.000 | 0.000 |
| Olfr889 | 0.000 | 0.002 | 0.003 |
| Olfr893 | 0.000 | 0.000 | 0.000 |
| Olfr905 | 0.000 | 0.001 | 0.001 |
| Olfr906 | 0.000 | 0.000 | 0.000 |
| Olfr912 | 0.000 | 0.000 | 0.000 |
| Olfr914 | 0.000 | 0.000 | 0.000 |
| Olfr924 | 0.000 | 0.000 | 0.000 |
| Olfr948 | 0.000 | 0.000 | 0.000 |
| Olfr959 | 0.000 | 0.000 | 0.002 |
| Olfr961 | 0.000 | 0.000 | 0.000 |

| | | | |
|---|---|---|---|
| Olfr967 | 0.000 | 0.000 | 0.000 |
| Olfr969 | 0.000 | 0.000 | 0.000 |
| Olfr974 | 0.000 | 0.000 | 0.000 |
| Olfr975 | 0.000 | 0.000 | 0.000 |
| Olfr976 | 0.000 | 0.000 | 0.000 |
| Olfr981 | 0.000 | 0.000 | 0.000 |
| Olfr984 | 0.000 | 0.000 | 0.000 |
| 1700110K17F | 0.000 | 0.000 | 0.000 |
| Gm48284 | 0.000 | 0.000 | 0.000 |
| Bsx | 0.000 | 0.000 | 0.000 |
| Jhy | 0.000 | 0.000 | 0.000 |
| Crtam | 0.000 | 0.000 | 0.000 |
| Gm48740 | 0.000 | 0.001 | 0.001 |
| Gm48784 | 0.000 | 0.000 | 0.000 |
| Gm40513 | 0.000 | 0.000 | 0.000 |
| Gm16214 | 0.000 | 0.000 | 0.000 |
| Gm39321 | 0.000 | 0.000 | 0.000 |
| 4930546K05F | 0.000 | 0.000 | 0.000 |
| Tecta | 0.000 | 0.000 | 0.000 |
| Gm47629 | 0.000 | 0.000 | 0.000 |
| Gm29961 | 0.000 | 0.000 | 0.000 |
| Gm3898 | 0.000 | 0.000 | 0.000 |
| Gm10688 | 0.000 | 0.000 | 0.000 |
| Gm39323 | 0.000 | 0.000 | 0.000 |
| Gm39325 | 0.000 | 0.000 | 0.000 |
| Mfrp | 0.000 | 0.000 | 0.001 |
| Pdzd3 | 0.000 | 0.005 | 0.001 |
| Gm48853 | 0.000 | 0.000 | 0.000 |
| Foxr1 | 0.000 | 0.000 | 0.000 |
| Cxcr5 | 0.000 | 0.000 | 0.000 |
| BC049987 | 0.000 | 0.000 | 0.001 |
| Treh | 0.000 | 0.000 | 0.000 |
| Gm48702 | 0.000 | 0.000 | 0.000 |
| Cd3d | 0.000 | 0.000 | 0.000 |
| Cd3e | 0.000 | 0.000 | 0.000 |
| Jaml | 0.000 | 0.003 | 0.002 |
| Gm10684 | 0.000 | 0.000 | 0.000 |
| Scn4b | 0.000 | 0.000 | 0.000 |
| Tmprss4 | 0.000 | 0.000 | 0.000 |
| BC049352 | 0.000 | 0.000 | 0.000 |
| Apoa4 | 0.000 | 0.000 | 0.000 |
| Apoa5 | 0.000 | 0.000 | 0.000 |
| 4931429L15F | 0.000 | 0.000 | 0.000 |
| Gm39329 | 0.000 | 0.000 | 0.000 |
| Gm31374 | 0.000 | 0.000 | 0.000 |
| Gm31614 | 0.000 | 0.000 | 0.000 |
| 2900052N01I | 0.000 | 0.000 | 0.000 |
| Gm47153 | 0.000 | 0.000 | 0.000 |

| | | | |
|---|---|---|---|
| Gm40518 | 0.000 | 0.000 | 0.000 |
| Htr3a | 0.000 | 0.000 | 0.000 |
| Htr3b | 0.000 | 0.000 | 0.000 |
| Tmprss5 | 0.000 | 0.001 | 0.002 |
| Drd2 | 0.000 | 0.000 | 0.000 |
| Ankk1 | 0.000 | 0.000 | 0.001 |
| Gm47543 | 0.000 | 0.000 | 0.000 |
| 2310003N18I | 0.000 | 0.000 | 0.000 |
| Gm47077 | 0.000 | 0.000 | 0.000 |
| Tex12 | 0.000 | 0.000 | 0.000 |
| Btg4 | 0.000 | 0.000 | 0.000 |
| 4833427G06I | 0.000 | 0.000 | 0.000 |
| Gm32742 | 0.000 | 0.000 | 0.000 |
| Pou2af1 | 0.000 | 0.000 | 0.000 |
| Arhgap20os | 0.000 | 0.000 | 0.000 |
| Gm6980 | 0.000 | 0.000 | 0.000 |
| Gm47957 | 0.000 | 0.000 | 0.000 |
| AI593442 | 0.000 | 0.000 | 0.000 |
| 4930510E17F | 0.000 | 0.000 | 0.000 |
| Slc35f2 | 0.000 | 0.001 | 0.000 |
| Gldn | 0.000 | 0.001 | 0.002 |
| Sh2d7 | 0.000 | 0.000 | 0.000 |
| Acsbg1 | 0.000 | 0.001 | 0.000 |
| Chrna3 | 0.000 | 0.000 | 0.000 |
| Chrnb4 | 0.000 | 0.001 | 0.000 |
| Gm17226 | 0.000 | 0.000 | 0.000 |
| Odf3l1 | 0.000 | 0.002 | 0.002 |
| Trcg1 | 0.000 | 0.000 | 0.000 |
| Cplx3 | 0.000 | 0.000 | 0.000 |
| Lman1l | 0.000 | 0.000 | 0.000 |
| Cyp1a1 | 0.000 | 0.000 | 0.000 |
| Gm17231 | 0.000 | 0.000 | 0.000 |
| Gm17322 | 0.000 | 0.000 | 0.000 |
| Ccdc33 | 0.000 | 0.000 | 0.000 |
| Gm16130 | 0.000 | 0.000 | 0.000 |
| 4930461G14I | 0.000 | 0.000 | 0.000 |
| Gm47271 | 0.000 | 0.000 | 0.000 |
| Gm33180 | 0.000 | 0.000 | 0.000 |
| Celf6 | 0.000 | 0.000 | 0.001 |
| Gm15507 | 0.000 | 0.002 | 0.000 |
| 2010001M07 | 0.000 | 0.000 | 0.000 |
| 9230112J17R | 0.000 | 0.001 | 0.000 |
| Gm49334 | 0.000 | 0.001 | 0.000 |
| 1700036A12F | 0.000 | 0.001 | 0.000 |
| Gm9869 | 0.000 | 0.001 | 0.001 |
| Gm47923 | 0.000 | 0.000 | 0.000 |
| Gm5122 | 0.000 | 0.000 | 0.000 |
| Gm34004 | 0.000 | 0.000 | 0.000 |

| | | | |
|---|---|---|---|
| B930092H01I | 0.000 | 0.001 | 0.000 |
| Gm47262 | 0.000 | 0.000 | 0.000 |
| Gm35288 | 0.000 | 0.000 | 0.000 |
| Gm48193 | 0.000 | 0.000 | 0.000 |
| Iqch | 0.000 | 0.001 | 0.001 |
| Gm47350 | 0.000 | 0.000 | 0.001 |
| Cilp | 0.000 | 0.002 | 0.047 |
| Gm39363 | 0.000 | 0.001 | 0.000 |
| Kbtbd13 | 0.000 | 0.003 | 0.002 |
| Ankdd1a | 0.000 | 0.000 | 0.000 |
| Gm15563 | 0.000 | 0.000 | 0.000 |
| BC050972 | 0.000 | 0.000 | 0.000 |
| Gm17098 | 0.000 | 0.000 | 0.000 |
| Gm47047 | 0.000 | 0.000 | 0.000 |
| M5C1000I18F | 0.000 | 0.000 | 0.000 |
| C2cd4a | 0.000 | 0.000 | 0.000 |
| Gm20745 | 0.000 | 0.000 | 0.000 |
| Gm15511 | 0.000 | 0.000 | 0.000 |
| Gm16144 | 0.000 | 0.000 | 0.000 |
| Gcnt3 | 0.000 | 0.000 | 0.000 |
| Gm26849 | 0.000 | 0.000 | 0.000 |
| Gm32511 | 0.000 | 0.000 | 0.000 |
| Gm27159 | 0.000 | 0.000 | 0.000 |
| BC065403 | 0.000 | 0.002 | 0.009 |
| 4930509E16F | 0.000 | 0.000 | 0.000 |
| Gm27232 | 0.000 | 0.000 | 0.000 |
| Gm27204 | 0.000 | 0.000 | 0.000 |
| Onecut1 | 0.000 | 0.000 | 0.000 |
| Myo5c | 0.000 | 0.001 | 0.001 |
| Gm48677 | 0.000 | 0.000 | 0.000 |
| Bmp5 | 0.000 | 0.000 | 0.000 |
| Gm26903 | 0.000 | 0.000 | 0.000 |
| Gfral | 0.000 | 0.000 | 0.000 |
| Gm27234 | 0.000 | 0.000 | 0.000 |
| Fam83b | 0.000 | 0.000 | 0.000 |
| Gm34158 | 0.000 | 0.000 | 0.000 |
| Tinag | 0.000 | 0.000 | 0.000 |
| Mlip | 0.000 | 0.002 | 0.001 |
| Gm34654 | 0.000 | 0.000 | 0.000 |
| Gm47818 | 0.000 | 0.000 | 0.000 |
| Gm39375 | 0.000 | 0.000 | 0.000 |
| Gcm1 | 0.000 | 0.000 | 0.000 |
| Gm3776 | 0.000 | 0.000 | 0.000 |
| Gm10639 | 0.000 | 0.000 | 0.000 |
| Gsta2 | 0.000 | 0.000 | 0.000 |
| 4930542C12F | 0.000 | 0.000 | 0.000 |
| Ddx43 | 0.000 | 0.001 | 0.000 |
| Gm47430 | 0.000 | 0.000 | 0.000 |

| | | | |
|---|---|---|---|
| 4921509A06F | 0.000 | 0.000 | 0.000 |
| Gm35024 | 0.000 | 0.000 | 0.002 |
| 4930562D21F | 0.000 | 0.001 | 0.001 |
| Gm27224 | 0.000 | 0.000 | 0.000 |
| D430036J16F | 0.000 | 0.008 | 0.003 |
| Mei4 | 0.000 | 0.002 | 0.001 |
| Gm39383 | 0.000 | 0.000 | 0.000 |
| Gm38398 | 0.000 | 0.000 | 0.000 |
| Gm46123 | 0.000 | 0.000 | 0.000 |
| Gm2109 | 0.000 | 0.000 | 0.000 |
| Gm48830 | 0.000 | 0.000 | 0.000 |
| Gm48831 | 0.000 | 0.000 | 0.000 |
| Gm48832 | 0.000 | 0.000 | 0.000 |
| Gm48833 | 0.000 | 0.000 | 0.000 |
| Gm48834 | 0.000 | 0.000 | 0.000 |
| Snap91 | 0.000 | 0.004 | 0.009 |
| Ripply2 | 0.000 | 0.000 | 0.000 |
| Gm28231 | 0.000 | 0.000 | 0.000 |
| Trim43a | 0.000 | 0.000 | 0.000 |
| Gm10634 | 0.000 | 0.000 | 0.000 |
| Trim43c | 0.000 | 0.000 | 0.000 |
| Bcl2a1a | 0.000 | 0.001 | 0.002 |
| Trim43b | 0.000 | 0.000 | 0.000 |
| 4930579C12F | 0.000 | 0.000 | 0.000 |
| Gm29094 | 0.000 | 0.000 | 0.000 |
| Gm47403 | 0.000 | 0.001 | 0.001 |
| Gm47409 | 0.000 | 0.000 | 0.000 |
| Rasgrf1 | 0.000 | 0.000 | 0.001 |
| Gm30849 | 0.000 | 0.000 | 0.000 |
| Gm28703 | 0.000 | 0.000 | 0.000 |
| Zic1 | 0.000 | 0.000 | 0.001 |
| Zic4 | 0.000 | 0.000 | 0.000 |
| 1700057G04F | 0.000 | 0.000 | 0.000 |
| Pls1 | 0.000 | 0.001 | 0.001 |
| C78334 | 0.000 | 0.000 | 0.000 |
| Gm28085 | 0.000 | 0.000 | 0.000 |
| Gm28729 | 0.000 | 0.000 | 0.000 |
| A930006L05F | 0.000 | 0.000 | 0.000 |
| Gm16185 | 0.000 | 0.002 | 0.001 |
| Trim42 | 0.000 | 0.000 | 0.000 |
| 4921534H16F | 0.000 | 0.000 | 0.000 |
| Foxl2 | 0.000 | 0.000 | 0.001 |
| Nme9 | 0.000 | 0.000 | 0.000 |
| A4gnt | 0.000 | 0.000 | 0.000 |
| Cldn18 | 0.000 | 0.000 | 0.000 |
| Gm28166 | 0.000 | 0.000 | 0.000 |
| Gm28979 | 0.000 | 0.000 | 0.000 |
| 9630041A04F | 0.000 | 0.000 | 0.000 |

| | | | |
|---|---|---|---|
| Gm26846 | 0.000 | 0.000 | 0.000 |
| 4930533D04I | 0.000 | 0.000 | 0.000 |
| Gm5627 | 0.000 | 0.000 | 0.000 |
| 5830418P13I | 0.000 | 0.000 | 0.000 |
| 4932413F04I | 0.000 | 0.000 | 0.000 |
| Gm29154 | 0.000 | 0.001 | 0.000 |
| Gm33054 | 0.000 | 0.000 | 0.000 |
| Gm47106 | 0.000 | 0.000 | 0.000 |
| Gm19667 | 0.000 | 0.000 | 0.000 |
| Col6a5 | 0.000 | 0.000 | 0.004 |
| Col6a4 | 0.000 | 0.000 | 0.003 |
| D030055H07I | 0.000 | 0.000 | 0.000 |
| Tlr9 | 0.000 | 0.000 | 0.000 |
| Gm29123 | 0.000 | 0.000 | 0.000 |
| Iqcf1 | 0.000 | 0.000 | 0.000 |
| Iqcf4 | 0.000 | 0.000 | 0.000 |
| Grm2 | 0.000 | 0.000 | 0.000 |
| 1700039M15 | 0.000 | 0.000 | 0.000 |
| Gm17041 | 0.000 | 0.000 | 0.000 |
| 4930429P21I | 0.000 | 0.000 | 0.000 |
| Gm34106 | 0.000 | 0.000 | 0.000 |
| Lsmem2 | 0.000 | 0.000 | 0.000 |
| Actl11 | 0.000 | 0.000 | 0.000 |
| 4921523L03I | 0.000 | 0.001 | 0.001 |
| Gm20662 | 0.000 | 0.000 | 0.000 |
| 4930535L15I | 0.000 | 0.000 | 0.000 |
| Mst1 | 0.000 | 0.000 | 0.000 |
| Bsn | 0.000 | 0.000 | 0.002 |
| Ccdc36 | 0.000 | 0.003 | 0.004 |
| Celsr3 | 0.000 | 0.000 | 0.001 |
| Ucn2 | 0.000 | 0.001 | 0.001 |
| Gm38004 | 0.000 | 0.000 | 0.000 |
| Fbxw20 | 0.000 | 0.000 | 0.000 |
| Fbxw22 | 0.000 | 0.000 | 0.000 |
| Gm42756 | 0.000 | 0.000 | 0.000 |
| Spink8 | 0.000 | 0.000 | 0.000 |
| Camp | 0.000 | 0.000 | 0.001 |
| Gm43732 | 0.000 | 0.000 | 0.000 |
| Ngp | 0.000 | 0.000 | 0.000 |
| Gm42470 | 0.000 | 0.000 | 0.000 |
| Gm35715 | 0.000 | 0.000 | 0.000 |
| Prss42 | 0.000 | 0.000 | 0.000 |
| Prss44 | 0.000 | 0.000 | 0.000 |
| Prss43 | 0.000 | 0.000 | 0.001 |
| Prss45 | 0.000 | 0.000 | 0.001 |
| Prss46 | 0.000 | 0.000 | 0.000 |
| Tdgf1 | 0.000 | 0.000 | 0.000 |
| Gm42523 | 0.000 | 0.000 | 0.000 |

| | | | |
|---|---|---|---|
| 2310075C17F | 0.000 | 0.000 | 0.000 |
| 2900079G21I | 0.000 | 0.001 | 0.001 |
| Gm36251 | 0.000 | 0.000 | 0.000 |
| Gm4657 | 0.000 | 0.002 | 0.001 |
| Gm36367 | 0.000 | 0.000 | 0.000 |
| AC164881.1 | 0.000 | 0.000 | 0.000 |
| Gm47875 | 0.000 | 0.000 | 0.000 |
| Gm36745 | 0.000 | 0.001 | 0.001 |
| Gm29825 | 0.000 | 0.001 | 0.000 |
| Bcl2a1c | 0.000 | 0.000 | 0.000 |
| Ccr4 | 0.000 | 0.000 | 0.000 |
| C130032M10 | 0.000 | 0.000 | 0.000 |
| Gm26962 | 0.000 | 0.000 | 0.000 |
| Gm48007 | 0.000 | 0.000 | 0.000 |
| Gadl1 | 0.000 | 0.000 | 0.001 |
| Gm4668 | 0.000 | 0.000 | 0.000 |
| Gm45881 | 0.000 | 0.000 | 0.000 |
| Eomes | 0.000 | 0.000 | 0.000 |
| Gm33460 | 0.000 | 0.000 | 0.000 |
| Gm16295 | 0.000 | 0.000 | 0.000 |
| Gm47289 | 0.000 | 0.000 | 0.000 |
| 4930516B21F | 0.000 | 0.000 | 0.000 |
| Scn5a | 0.000 | 0.000 | 0.000 |
| Scn10a | 0.000 | 0.000 | 0.000 |
| Gm47043 | 0.000 | 0.000 | 0.000 |
| AC117245.1 | 0.000 | 0.000 | 0.000 |
| Gm33858 | 0.000 | 0.000 | 0.000 |
| Ccr8 | 0.000 | 0.000 | 0.000 |
| Gm5922 | 0.000 | 0.000 | 0.000 |
| Gm38642 | 0.000 | 0.000 | 0.000 |
| Entpd3 | 0.000 | 0.000 | 0.000 |
| Gm47095 | 0.000 | 0.000 | 0.000 |
| Gm38661 | 0.000 | 0.000 | 0.000 |
| Hhatl | 0.000 | 0.004 | 0.008 |
| Gm17163 | 0.000 | 0.001 | 0.000 |
| Gm47113 | 0.000 | 0.001 | 0.001 |
| Cyp8b1 | 0.000 | 0.000 | 0.000 |
| Gm47117 | 0.000 | 0.000 | 0.001 |
| A730085K08F | 0.000 | 0.000 | 0.000 |
| Gm47134 | 0.000 | 0.000 | 0.000 |
| Gm47136 | 0.000 | 0.000 | 0.000 |
| Topaz1 | 0.000 | 0.000 | 0.000 |
| Gm17120 | 0.000 | 0.001 | 0.000 |
| Ccr9 | 0.000 | 0.001 | 0.001 |
| Cxcr6 | 0.000 | 0.000 | 0.000 |
| Xcr1 | 0.000 | 0.001 | 0.001 |
| Gm47181 | 0.000 | 0.000 | 0.000 |
| Gm12735 | 0.000 | 0.000 | 0.000 |

| | | | |
|---|---|---|---|
| 4921536K21I | 0.000 | 0.000 | 0.001 |
| 4930556J24R | 0.000 | 0.000 | 0.000 |
| Sec14l4 | 0.000 | 0.000 | 0.000 |
| Sec14l3 | 0.000 | 0.000 | 0.000 |
| Osm | 0.000 | 0.000 | 0.000 |
| Hormad2 | 0.000 | 0.000 | 0.000 |
| Gm11961 | 0.000 | 0.000 | 0.000 |
| Cabp7 | 0.000 | 0.000 | 0.000 |
| Ankrd36 | 0.000 | 0.002 | 0.000 |
| Gm11967 | 0.000 | 0.001 | 0.000 |
| Npc1l1 | 0.000 | 0.000 | 0.000 |
| Gm11985 | 0.000 | 0.000 | 0.000 |
| Pkd1l1 | 0.000 | 0.000 | 0.000 |
| Gm11998 | 0.000 | 0.000 | 0.000 |
| 4930512M02 | 0.000 | 0.000 | 0.000 |
| 1700042O10I | 0.000 | 0.000 | 0.001 |
| Gm12002 | 0.000 | 0.003 | 0.001 |
| 4930554G24I | 0.000 | 0.000 | 0.000 |
| Vstm2a | 0.000 | 0.001 | 0.005 |
| Egfros | 0.000 | 0.006 | 0.012 |
| Gm12018 | 0.000 | 0.000 | 0.000 |
| Gm12023 | 0.000 | 0.000 | 0.000 |
| Gm16140 | 0.000 | 0.004 | 0.007 |
| Gm12027 | 0.000 | 0.002 | 0.000 |
| Gm12037 | 0.000 | 0.000 | 0.000 |
| Otx1 | 0.000 | 0.000 | 0.002 |
| Gm12050 | 0.000 | 0.000 | 0.000 |
| Gm12051 | 0.000 | 0.000 | 0.000 |
| Gm12056 | 0.000 | 0.000 | 0.000 |
| 1700061J23R | 0.000 | 0.000 | 0.000 |
| Gm12063 | 0.000 | 0.000 | 0.000 |
| Gm12064 | 0.000 | 0.000 | 0.000 |
| 4930538E20I | 0.000 | 0.000 | 0.000 |
| Gm12065 | 0.000 | 0.000 | 0.000 |
| Gm6899 | 0.000 | 0.000 | 0.000 |
| Gm12092 | 0.000 | 0.000 | 0.000 |
| Gm12107 | 0.000 | 0.000 | 0.000 |
| 4930524B15I | 0.000 | 0.000 | 0.000 |
| Il9r | 0.000 | 0.000 | 0.000 |
| Hba-x | 0.000 | 0.000 | 0.001 |
| Hbq1a | 0.000 | 0.000 | 0.000 |
| Tlx3 | 0.000 | 0.000 | 0.001 |
| Gm12116 | 0.000 | 0.001 | 0.000 |
| Gm12117 | 0.000 | 0.000 | 0.000 |
| Gabrp | 0.000 | 0.000 | 0.000 |
| 4930469K13I | 0.000 | 0.001 | 0.001 |
| 4930403D09I | 0.000 | 0.000 | 0.000 |
| Fam196b | 0.000 | 0.002 | 0.001 |

| | | | |
|---|---|---|---|
| Fbll1 | 0.000 | 0.000 | 0.000 |
| Gm12128 | 0.000 | 0.000 | 0.000 |
| Gabrg2 | 0.000 | 0.000 | 0.000 |
| Gm9972 | 0.000 | 0.000 | 0.000 |
| Gm12146 | 0.000 | 0.000 | 0.000 |
| Gm12144 | 0.000 | 0.001 | 0.002 |
| Gm12148 | 0.000 | 0.000 | 0.000 |
| Gm12150 | 0.000 | 0.000 | 0.000 |
| Gm12153 | 0.000 | 0.000 | 0.000 |
| Il12b | 0.000 | 0.000 | 0.000 |
| 4930597A21F | 0.000 | 0.000 | 0.001 |
| Gm12159 | 0.000 | 0.000 | 0.000 |
| Sox30 | 0.000 | 0.004 | 0.003 |
| Gm16034 | 0.000 | 0.001 | 0.000 |
| Itk | 0.000 | 0.000 | 0.000 |
| Gm12167 | 0.000 | 0.000 | 0.000 |
| Fam71b | 0.000 | 0.000 | 0.000 |
| 2310031A07F | 0.000 | 0.000 | 0.000 |
| BC053393 | 0.000 | 0.000 | 0.000 |
| Gm4926 | 0.000 | 0.000 | 0.000 |
| Havcr1 | 0.000 | 0.000 | 0.000 |
| Timd4 | 0.000 | 0.000 | 0.000 |
| Gm16170 | 0.000 | 0.001 | 0.000 |
| Tgtp1 | 0.000 | 0.000 | 0.000 |
| Olfr1396 | 0.000 | 0.000 | 0.000 |
| Btnl9 | 0.000 | 0.000 | 0.001 |
| Olfr1392 | 0.000 | 0.001 | 0.001 |
| Olfr10 | 0.000 | 0.000 | 0.000 |
| Olfr1390 | 0.000 | 0.000 | 0.000 |
| Olfr1389 | 0.000 | 0.000 | 0.000 |
| Olfr1388 | 0.000 | 0.000 | 0.000 |
| Olfr1382 | 0.000 | 0.000 | 0.000 |
| Olfr1381 | 0.000 | 0.000 | 0.000 |
| Rasgef1c | 0.000 | 0.001 | 0.002 |
| Gm12195 | 0.000 | 0.000 | 0.000 |
| Gm26542 | 0.000 | 0.001 | 0.003 |
| Cby3 | 0.000 | 0.000 | 0.000 |
| Gm12199 | 0.000 | 0.000 | 0.000 |
| Gm12200 | 0.000 | 0.000 | 0.000 |
| 4933414I15R | 0.000 | 0.000 | 0.000 |
| Olfr51 | 0.000 | 0.000 | 0.000 |
| Olfr54 | 0.000 | 0.000 | 0.000 |
| 9230009I02R | 0.000 | 0.000 | 0.000 |
| 9430098F02F | 0.000 | 0.001 | 0.001 |
| Olfr1373 | 0.000 | 0.000 | 0.000 |
| Leap2 | 0.000 | 0.000 | 0.000 |
| Gm10447 | 0.000 | 0.000 | 0.000 |
| Gm12212 | 0.000 | 0.000 | 0.000 |

| | | | |
|---|---|---|---|
| Gm9837 | 0.000 | 0.000 | 0.000 |
| Sowaha | 0.000 | 0.000 | 0.000 |
| Il4 | 0.000 | 0.000 | 0.000 |
| Il13 | 0.000 | 0.000 | 0.000 |
| Csf2 | 0.000 | 0.000 | 0.000 |
| Gm12223 | 0.000 | 0.000 | 0.000 |
| Acsl6 | 0.000 | 0.000 | 0.001 |
| Gm12224 | 0.000 | 0.000 | 0.001 |
| Meikin | 0.000 | 0.000 | 0.000 |
| Gm12227 | 0.000 | 0.000 | 0.000 |
| Lyrm7os | 0.000 | 0.000 | 0.000 |
| Slc36a3 | 0.000 | 0.000 | 0.000 |
| Slc36a1os | 0.000 | 0.000 | 0.000 |
| Fat2 | 0.000 | 0.000 | 0.000 |
| Nmur2 | 0.000 | 0.000 | 0.000 |
| Gm12243 | 0.000 | 0.000 | 0.000 |
| 4933426K07R | 0.000 | 0.000 | 0.000 |
| Gm12248 | 0.000 | 0.000 | 0.000 |
| Gm12246 | 0.000 | 0.000 | 0.000 |
| 4930438A08R | 0.000 | 0.000 | 0.000 |
| Lypd8 | 0.000 | 0.000 | 0.000 |
| 1810065E05R | 0.000 | 0.000 | 0.000 |
| Olfr30 | 0.000 | 0.000 | 0.000 |
| Olfr330 | 0.000 | 0.000 | 0.001 |
| Olfr329-ps | 0.000 | 0.000 | 0.001 |
| Olfr224 | 0.000 | 0.001 | 0.001 |
| Olfr325 | 0.000 | 0.000 | 0.000 |
| 2210407C18R | 0.000 | 0.000 | 0.000 |
| Olfr314 | 0.000 | 0.000 | 0.000 |
| Fam183b | 0.000 | 0.000 | 0.000 |
| Olfr313 | 0.000 | 0.000 | 0.000 |
| Gm12256 | 0.000 | 0.000 | 0.000 |
| Trim17 | 0.000 | 0.001 | 0.000 |
| Gm10435 | 0.000 | 0.000 | 0.000 |
| Prss38 | 0.000 | 0.001 | 0.001 |
| Gm15755 | 0.000 | 0.000 | 0.000 |
| Olfr222 | 0.000 | 0.000 | 0.000 |
| Gm12714 | 0.000 | 0.000 | 0.000 |
| 1810063I02R | 0.000 | 0.000 | 0.000 |
| 4930412M03 | 0.000 | 0.000 | 0.000 |
| Gm26837 | 0.000 | 0.001 | 0.000 |
| Kcnj12 | 0.000 | 0.000 | 0.000 |
| Slc47a2 | 0.000 | 0.000 | 0.000 |
| Gm26964 | 0.000 | 0.005 | 0.003 |
| Gm12273 | 0.000 | 0.000 | 0.000 |
| Gm12949 | 0.000 | 0.000 | 0.000 |
| Gm12279 | 0.000 | 0.000 | 0.001 |
| Gm12280 | 0.000 | 0.001 | 0.003 |

| | | | |
|---|---|---|---|
| Lrrc75aos1 | 0.000 | 0.000 | 0.000 |
| Gm27194 | 0.000 | 0.000 | 0.000 |
| Cdrt4os1 | 0.000 | 0.000 | 0.000 |
| Tekt3 | 0.000 | 0.002 | 0.005 |
| B430319H21I | 0.000 | 0.001 | 0.001 |
| B130011K05F | 0.000 | 0.001 | 0.002 |
| Gm12289 | 0.000 | 0.000 | 0.000 |
| Gm12290 | 0.000 | 0.000 | 0.000 |
| Dnah9 | 0.000 | 0.001 | 0.003 |
| Shisa6 | 0.000 | 0.000 | 0.001 |
| Myh2 | 0.000 | 0.000 | 0.001 |
| 9430073C21I | 0.000 | 0.000 | 0.000 |
| Myh1 | 0.000 | 0.000 | 0.001 |
| Myhas | 0.000 | 0.000 | 0.000 |
| Myh8 | 0.000 | 0.000 | 0.000 |
| Myh13 | 0.000 | 0.000 | 0.000 |
| Rcvrn | 0.000 | 0.000 | 0.001 |
| Cfap52 | 0.000 | 0.001 | 0.000 |
| Pik3r5 | 0.000 | 0.002 | 0.003 |
| Mfsd6l | 0.000 | 0.000 | 0.000 |
| Alox8 | 0.000 | 0.000 | 0.000 |
| Gucy2e | 0.000 | 0.000 | 0.000 |
| Shbg | 0.000 | 0.000 | 0.000 |
| Tnfsf13 | 0.000 | 0.000 | 0.000 |
| Gm39566 | 0.000 | 0.001 | 0.002 |
| Spem2 | 0.000 | 0.000 | 0.001 |
| Mgl2 | 0.000 | 0.001 | 0.001 |
| Alox15 | 0.000 | 0.000 | 0.000 |
| Gltpd2 | 0.000 | 0.000 | 0.001 |
| Gm40193 | 0.000 | 0.000 | 0.000 |
| Gm12316 | 0.000 | 0.000 | 0.000 |
| 4930544D05I | 0.000 | 0.000 | 0.000 |
| Gm12315 | 0.000 | 0.000 | 0.000 |
| Gm12319 | 0.000 | 0.001 | 0.000 |
| Gm12320 | 0.000 | 0.000 | 0.000 |
| Scimp | 0.000 | 0.002 | 0.001 |
| Nlrp1a | 0.000 | 0.000 | 0.000 |
| Nlrp1b | 0.000 | 0.000 | 0.000 |
| Slc13a5 | 0.000 | 0.000 | 0.000 |
| Trpv1 | 0.000 | 0.000 | 0.000 |
| Trpv3 | 0.000 | 0.000 | 0.001 |
| Olfr381 | 0.000 | 0.000 | 0.000 |
| Olfr384 | 0.000 | 0.000 | 0.000 |
| Olfr392 | 0.000 | 0.000 | 0.000 |
| Olfr393 | 0.000 | 0.000 | 0.000 |
| Olfr394 | 0.000 | 0.000 | 0.000 |
| Olfr395 | 0.000 | 0.000 | 0.000 |
| Olfr403 | 0.000 | 0.000 | 0.000 |

| | | | |
|---|---|---|---|
| Olfr410 | 0.000 | 0.000 | 0.000 |
| Olfr411 | 0.000 | 0.000 | 0.000 |
| Ccdc92b | 0.000 | 0.002 | 0.001 |
| Gm28180 | 0.000 | 0.000 | 0.000 |
| Gm12333 | 0.000 | 0.000 | 0.000 |
| Gm12336 | 0.000 | 0.000 | 0.000 |
| Gm26780 | 0.000 | 0.000 | 0.000 |
| Bhlha9 | 0.000 | 0.000 | 0.000 |
| Tmigd1 | 0.000 | 0.001 | 0.004 |
| Slc6a4 | 0.000 | 0.003 | 0.002 |
| Sez6 | 0.000 | 0.001 | 0.001 |
| Foxn1 | 0.000 | 0.000 | 0.000 |
| Nos2 | 0.000 | 0.000 | 0.000 |
| Gm11201 | 0.000 | 0.000 | 0.000 |
| Gm44787 | 0.000 | 0.000 | 0.000 |
| Gm9964 | 0.000 | 0.000 | 0.000 |
| AU040972 | 0.000 | 0.000 | 0.000 |
| Omg | 0.000 | 0.001 | 0.001 |
| Evi2b | 0.000 | 0.000 | 0.000 |
| Rab11fip4os1 | 0.000 | 0.000 | 0.000 |
| Adap2os | 0.000 | 0.000 | 0.000 |
| Asic2 | 0.000 | 0.001 | 0.002 |
| 5530401A14I | 0.000 | 0.000 | 0.000 |
| Gm31522 | 0.000 | 0.000 | 0.000 |
| Ccl1 | 0.000 | 0.000 | 0.000 |
| Gm38577 | 0.000 | 0.000 | 0.000 |
| Fndc8 | 0.000 | 0.000 | 0.000 |
| Gm48462 | 0.000 | 0.000 | 0.000 |
| Slfn1 | 0.000 | 0.000 | 0.000 |
| Slfn14 | 0.000 | 0.000 | 0.000 |
| Gas2l2 | 0.000 | 0.000 | 0.000 |
| Heatr9 | 0.000 | 0.000 | 0.000 |
| Wfdc21 | 0.000 | 0.000 | 0.001 |
| Gm11437 | 0.000 | 0.000 | 0.000 |
| Lhx1os | 0.000 | 0.000 | 0.000 |
| Gm11431 | 0.000 | 0.000 | 0.000 |
| Bcas3os2 | 0.000 | 0.001 | 0.001 |
| Gm11479 | 0.000 | 0.000 | 0.000 |
| Mpo | 0.000 | 0.000 | 0.002 |
| Lpo | 0.000 | 0.000 | 0.000 |
| Epx | 0.000 | 0.000 | 0.000 |
| Gm11508 | 0.000 | 0.001 | 0.000 |
| Gm38534 | 0.000 | 0.000 | 0.000 |
| Ccdc182 | 0.000 | 0.000 | 0.000 |
| Gm11493 | 0.000 | 0.000 | 0.000 |
| 4930556N13I | 0.000 | 0.000 | 0.000 |
| Scpep1os | 0.000 | 0.000 | 0.000 |
| Elobl | 0.000 | 0.007 | 0.003 |

| | | | |
|---|---|---|---|
| Gm2018 | 0.000 | 0.000 | 0.000 |
| Ankfn1 | 0.000 | 0.000 | 0.000 |
| Gm11494 | 0.000 | 0.000 | 0.000 |
| 4930405D11I | 0.000 | 0.000 | 0.000 |
| Cdc34b | 0.000 | 0.000 | 0.001 |
| Tmem92 | 0.000 | 0.000 | 0.000 |
| Gm11513 | 0.000 | 0.000 | 0.000 |
| Dlx3 | 0.000 | 0.000 | 0.000 |
| Dlx4 | 0.000 | 0.000 | 0.000 |
| Dlx4os | 0.000 | 0.000 | 0.000 |
| Gm11515 | 0.000 | 0.000 | 0.000 |
| Tac4 | 0.000 | 0.000 | 0.000 |
| Gm11521 | 0.000 | 0.000 | 0.000 |
| Gm11534 | 0.000 | 0.000 | 0.000 |
| Gm29202 | 0.000 | 0.000 | 0.000 |
| Calcoco2 | 0.000 | 0.000 | 0.000 |
| Ttll6 | 0.000 | 0.000 | 0.000 |
| Gm11536 | 0.000 | 0.002 | 0.001 |
| Skap1 | 0.000 | 0.002 | 0.003 |
| 2010300F17F | 0.000 | 0.001 | 0.001 |
| Gm11537 | 0.000 | 0.000 | 0.000 |
| Gm11525 | 0.000 | 0.002 | 0.000 |
| Tbx21 | 0.000 | 0.001 | 0.000 |
| Gm11583 | 0.000 | 0.000 | 0.000 |
| Gpr179 | 0.000 | 0.002 | 0.002 |
| Gm11614 | 0.000 | 0.000 | 0.000 |
| Ppp1r1b | 0.000 | 0.000 | 0.001 |
| Pnmt | 0.000 | 0.003 | 0.000 |
| Gm12352 | 0.000 | 0.000 | 0.001 |
| Ikzf3 | 0.000 | 0.000 | 0.000 |
| Ccr7 | 0.000 | 0.000 | 0.000 |
| Krt222 | 0.000 | 0.000 | 0.002 |
| Krt24 | 0.000 | 0.000 | 0.000 |
| Krt12 | 0.000 | 0.000 | 0.000 |
| A830036E02F | 0.000 | 0.000 | 0.000 |
| Krtap4-9 | 0.000 | 0.000 | 0.000 |
| Krtap17-1 | 0.000 | 0.000 | 0.000 |
| Krt33a | 0.000 | 0.000 | 0.001 |
| Krt33b | 0.000 | 0.000 | 0.000 |
| Krt31 | 0.000 | 0.000 | 0.000 |
| Krt32 | 0.000 | 0.000 | 0.000 |
| Krt35 | 0.000 | 0.000 | 0.000 |
| Krt36 | 0.000 | 0.000 | 0.006 |
| Krt9 | 0.000 | 0.000 | 0.000 |
| Krt42 | 0.000 | 0.000 | 0.000 |
| Gm12349 | 0.000 | 0.000 | 0.000 |
| Ttc25 | 0.000 | 0.002 | 0.006 |
| Hspb9 | 0.000 | 0.000 | 0.000 |

| | | | |
|---|---|---|---|
| Kcnh4 | 0.000 | 0.000 | 0.000 |
| Hcrt | 0.000 | 0.000 | 0.000 |
| Gm28156 | 0.000 | 0.000 | 0.000 |
| G6pc | 0.000 | 0.002 | 0.001 |
| Gm11626 | 0.000 | 0.000 | 0.000 |
| Gm11634 | 0.000 | 0.000 | 0.000 |
| Gm11551 | 0.000 | 0.000 | 0.001 |
| 4930417O22I | 0.000 | 0.004 | 0.000 |
| Cd300lg | 0.000 | 0.001 | 0.002 |
| Nags | 0.000 | 0.000 | 0.000 |
| Gm11636 | 0.000 | 0.003 | 0.009 |
| Fam187a | 0.000 | 0.000 | 0.000 |
| 1700023F06F | 0.000 | 0.001 | 0.002 |
| Spata32 | 0.000 | 0.000 | 0.000 |
| Arhgap27os3 | 0.000 | 0.000 | 0.000 |
| Gm39397 | 0.000 | 0.000 | 0.000 |
| Gm884 | 0.000 | 0.000 | 0.000 |
| Wnt3 | 0.000 | 0.000 | 0.000 |
| Gm11665 | 0.000 | 0.000 | 0.000 |
| Gm11639 | 0.000 | 0.001 | 0.001 |
| Efcab3 | 0.000 | 0.000 | 0.000 |
| Gm10842 | 0.000 | 0.000 | 0.000 |
| Gm11651 | 0.000 | 0.000 | 0.000 |
| Ace3 | 0.000 | 0.000 | 0.000 |
| Kcnh6 | 0.000 | 0.000 | 0.000 |
| Gm11646 | 0.000 | 0.000 | 0.000 |
| Gm10840 | 0.000 | 0.000 | 0.000 |
| Tcam1 | 0.000 | 0.000 | 0.003 |
| Gh | 0.000 | 0.000 | 0.000 |
| Prr29 | 0.000 | 0.000 | 0.000 |
| Gm11706 | 0.000 | 0.000 | 0.000 |
| Gm11707 | 0.000 | 0.000 | 0.000 |
| Gm11715 | 0.000 | 0.000 | 0.000 |
| Cacng5 | 0.000 | 0.000 | 0.000 |
| Gm11655 | 0.000 | 0.000 | 0.000 |
| 9930022D16I | 0.000 | 0.000 | 0.000 |
| Gm11685 | 0.000 | 0.000 | 0.000 |
| 1700012B07F | 0.000 | 0.000 | 0.000 |
| Gm11697 | 0.000 | 0.000 | 0.001 |
| Gm11682 | 0.000 | 0.000 | 0.000 |
| Kcnj16 | 0.000 | 0.000 | 0.000 |
| Gm11680 | 0.000 | 0.000 | 0.000 |
| Cpsf4l | 0.000 | 0.000 | 0.000 |
| 1700025D23I | 0.000 | 0.000 | 0.000 |
| 1700001J04R | 0.000 | 0.000 | 0.000 |
| 4932435O22I | 0.000 | 0.000 | 0.000 |
| Dnaic2 | 0.000 | 0.004 | 0.019 |
| Kif19a | 0.000 | 0.000 | 0.001 |

| | | | |
|---|---|---|---|
| Btbd17 | 0.000 | 0.000 | 0.001 |
| Cd300lb | 0.000 | 0.000 | 0.000 |
| Cd300c | 0.000 | 0.000 | 0.000 |
| Cd300ld | 0.000 | 0.000 | 0.001 |
| Cd300ld2 | 0.000 | 0.000 | 0.000 |
| Cd300ld4 | 0.000 | 0.000 | 0.000 |
| Cd300ld3 | 0.000 | 0.000 | 0.000 |
| Cd300e | 0.000 | 0.000 | 0.000 |
| Otop2 | 0.000 | 0.000 | 0.001 |
| Ush1g | 0.000 | 0.000 | 0.000 |
| Otop3 | 0.000 | 0.000 | 0.000 |
| Ict1os | 0.000 | 0.000 | 0.001 |
| Gm26613 | 0.000 | 0.000 | 0.000 |
| Myo15b | 0.000 | 0.000 | 0.001 |
| Gm26730 | 0.000 | 0.000 | 0.000 |
| Qrich2 | 0.000 | 0.001 | 0.001 |
| Gm11739 | 0.000 | 0.000 | 0.000 |
| Aanat | 0.000 | 0.000 | 0.000 |
| St6galnac2.1 | 0.000 | 0.000 | 0.000 |
| St6galnac1 | 0.000 | 0.000 | 0.000 |
| Gm16045 | 0.000 | 0.002 | 0.002 |
| Gm11729 | 0.000 | 0.000 | 0.000 |
| Gm34418 | 0.000 | 0.000 | 0.000 |
| Gm11733 | 0.000 | 0.000 | 0.000 |
| Gm11734 | 0.000 | 0.000 | 0.000 |
| Gm11724 | 0.000 | 0.000 | 0.000 |
| 6030468B19F | 0.000 | 0.000 | 0.000 |
| Tmem235 | 0.000 | 0.000 | 0.000 |
| Gm11725 | 0.000 | 0.002 | 0.002 |
| Gm11747 | 0.000 | 0.001 | 0.001 |
| Rbfox3 | 0.000 | 0.001 | 0.001 |
| Gm26888 | 0.000 | 0.000 | 0.000 |
| Rptoros | 0.000 | 0.000 | 0.000 |
| Gm11766 | 0.000 | 0.000 | 0.000 |
| Gm16755 | 0.000 | 0.000 | 0.000 |
| Gm11788 | 0.000 | 0.000 | 0.000 |
| Gm11789 | 0.000 | 0.000 | 0.000 |
| Myadml2 | 0.000 | 0.001 | 0.000 |
| Myadml2os | 0.000 | 0.000 | 0.000 |
| Notumos | 0.000 | 0.000 | 0.000 |
| Cd7 | 0.000 | 0.000 | 0.000 |
| Sectm1b | 0.000 | 0.000 | 0.000 |
| Sectm1a | 0.000 | 0.000 | 0.000 |
| Tex19.1 | 0.000 | 0.000 | 0.000 |
| Uts2r | 0.000 | 0.000 | 0.001 |
| Zfp750 | 0.000 | 0.000 | 0.000 |
| Gm12589 | 0.000 | 0.001 | 0.000 |
| Ptchd3 | 0.000 | 0.000 | 0.000 |

| | | | |
|---|---|---|---|
| Gm48530 | 0.000 | 0.000 | 0.000 |
| Calml3 | 0.000 | 0.001 | 0.000 |
| Calm4 | 0.000 | 0.000 | 0.000 |
| Ucn3 | 0.000 | 0.000 | 0.000 |
| Akr1c18 | 0.000 | 0.001 | 0.027 |
| Gm47925 | 0.000 | 0.000 | 0.000 |
| Akr1c12 | 0.000 | 0.000 | 0.002 |
| Akr1c20 | 0.000 | 0.000 | 0.000 |
| Akr1c21 | 0.000 | 0.000 | 0.002 |
| Gm40658 | 0.000 | 0.000 | 0.000 |
| Gm5444 | 0.000 | 0.000 | 0.000 |
| AC151530.1 | 0.000 | 0.000 | 0.000 |
| 1700016G22I | 0.000 | 0.001 | 0.001 |
| Gm47507 | 0.000 | 0.000 | 0.000 |
| Gm47549 | 0.000 | 0.000 | 0.000 |
| Idi2 | 0.000 | 0.000 | 0.001 |
| Gm48825 | 0.000 | 0.000 | 0.000 |
| Gm26861 | 0.000 | 0.000 | 0.000 |
| Gm47408 | 0.000 | 0.000 | 0.000 |
| Gm47486 | 0.000 | 0.000 | 0.000 |
| Gm30239 | 0.000 | 0.000 | 0.000 |
| Prl2c3 | 0.000 | 0.000 | 0.000 |
| Gm48544 | 0.000 | 0.000 | 0.000 |
| Gng4 | 0.000 | 0.002 | 0.008 |
| Gm30893 | 0.000 | 0.001 | 0.001 |
| 4933412O06I | 0.000 | 0.000 | 0.000 |
| A530046M15 | 0.000 | 0.000 | 0.000 |
| Gm48503 | 0.000 | 0.002 | 0.002 |
| Tcrg-V6 | 0.000 | 0.000 | 0.000 |
| Tcrg-V5 | 0.000 | 0.000 | 0.000 |
| Tcrg-C2 | 0.000 | 0.000 | 0.000 |
| Trgv2 | 0.000 | 0.000 | 0.000 |
| Tcrg-C4 | 0.000 | 0.000 | 0.000 |
| 9330199G10I | 0.000 | 0.000 | 0.000 |
| Nme8 | 0.000 | 0.001 | 0.000 |
| Olfr1370 | 0.000 | 0.000 | 0.000 |
| Gm47806 | 0.000 | 0.000 | 0.000 |
| Gpx5 | 0.000 | 0.000 | 0.000 |
| Olfr1366 | 0.000 | 0.000 | 0.000 |
| Hist1h2bl | 0.000 | 0.000 | 0.000 |
| Hist1h3h | 0.000 | 0.000 | 0.000 |
| Hist1h4k | 0.000 | 0.000 | 0.000 |
| Gm11274 | 0.000 | 0.001 | 0.000 |
| Hist1h2bp | 0.000 | 0.001 | 0.000 |
| Hist1h2ao | 0.000 | 0.000 | 0.000 |
| Hist1h4m | 0.000 | 0.000 | 0.000 |
| Pom121l2 | 0.000 | 0.000 | 0.000 |
| Prss16 | 0.000 | 0.000 | 0.001 |

| | | | |
|---|---|---|---|
| Hist1h2ah | 0.000 | 0.000 | 0.000 |
| Vmn1r188 | 0.000 | 0.000 | 0.000 |
| Vmn1r193 | 0.000 | 0.000 | 0.000 |
| Vmn1r197 | 0.000 | 0.000 | 0.000 |
| Vmn1r200 | 0.000 | 0.000 | 0.000 |
| Vmn1r203 | 0.000 | 0.000 | 0.000 |
| Vmn1r206 | 0.000 | 0.000 | 0.000 |
| Vmn1r210 | 0.000 | 0.000 | 0.000 |
| Vmn1r212 | 0.000 | 0.000 | 0.000 |
| Vmn1r216 | 0.000 | 0.000 | 0.000 |
| Vmn1r220 | 0.000 | 0.000 | 0.000 |
| 4930586N03I | 0.000 | 0.000 | 0.000 |
| Btn1a1 | 0.000 | 0.000 | 0.001 |
| Hist1h4f | 0.000 | 0.001 | 0.001 |
| Hist1h2ad | 0.000 | 0.000 | 0.000 |
| 4930558J22R | 0.000 | 0.000 | 0.000 |
| Trim38 | 0.000 | 0.000 | 0.000 |
| Slc17a2 | 0.000 | 0.000 | 0.000 |
| Slc17a3 | 0.000 | 0.000 | 0.000 |
| Slc17a4 | 0.000 | 0.000 | 0.000 |
| Hist1h2aa | 0.000 | 0.000 | 0.000 |
| Gm11342 | 0.000 | 0.000 | 0.001 |
| 1700016G14I | 0.000 | 0.000 | 0.000 |
| 4932702P03I | 0.000 | 0.000 | 0.000 |
| Gm11351 | 0.000 | 0.000 | 0.000 |
| Prl | 0.000 | 0.000 | 0.000 |
| Gm34639 | 0.000 | 0.000 | 0.000 |
| Gm11365 | 0.000 | 0.000 | 0.000 |
| Irf4 | 0.000 | 0.000 | 0.001 |
| Gm29675 | 0.000 | 0.001 | 0.002 |
| A530084C06I | 0.000 | 0.000 | 0.000 |
| Gm11377 | 0.000 | 0.000 | 0.000 |
| 1700018A04I | 0.000 | 0.000 | 0.000 |
| D930007J09I | 0.000 | 0.000 | 0.000 |
| Serpinb1c | 0.000 | 0.000 | 0.000 |
| Serpinb1b | 0.000 | 0.000 | 0.001 |
| Gm6093 | 0.000 | 0.000 | 0.000 |
| Serpinb6e | 0.000 | 0.000 | 0.000 |
| Serpinb6c | 0.000 | 0.001 | 0.004 |
| Gm47076 | 0.000 | 0.000 | 0.000 |
| Gm36500 | 0.000 | 0.000 | 0.000 |
| Gm15908 | 0.000 | 0.000 | 0.000 |
| Gm47151 | 0.000 | 0.000 | 0.000 |
| Gm47157 | 0.000 | 0.000 | 0.000 |
| 4933417A18I | 0.000 | 0.000 | 0.000 |
| Gm16984 | 0.000 | 0.000 | 0.001 |
| Gm40910 | 0.000 | 0.000 | 0.000 |
| Gm48629 | 0.000 | 0.000 | 0.000 |

| | | | |
|---|---|---|---|
| 1700019C18F | 0.000 | 0.000 | 0.000 |
| Gm48703 | 0.000 | 0.000 | 0.000 |
| Gm48746 | 0.000 | 0.000 | 0.000 |
| Gm30127 | 0.000 | 0.000 | 0.000 |
| Gm48765 | 0.000 | 0.000 | 0.000 |
| Gm30177 | 0.000 | 0.000 | 0.000 |
| Gm49144 | 0.000 | 0.002 | 0.001 |
| Gm47675 | 0.000 | 0.000 | 0.000 |
| Gm30489 | 0.000 | 0.000 | 0.000 |
| Gm30600 | 0.000 | 0.000 | 0.000 |
| Gm47709 | 0.000 | 0.000 | 0.000 |
| Gm47711 | 0.000 | 0.000 | 0.000 |
| Gm47732 | 0.000 | 0.000 | 0.000 |
| Gm30918 | 0.000 | 0.000 | 0.000 |
| Gm47754 | 0.000 | 0.000 | 0.000 |
| Gm47990 | 0.000 | 0.000 | 0.000 |
| Gm47509 | 0.000 | 0.000 | 0.000 |
| Gm47348 | 0.000 | 0.000 | 0.000 |
| Gm3509 | 0.000 | 0.000 | 0.000 |
| Gm47061 | 0.000 | 0.002 | 0.000 |
| Gm26688 | 0.000 | 0.001 | 0.001 |
| Mak | 0.000 | 0.000 | 0.000 |
| Gcm2 | 0.000 | 0.000 | 0.000 |
| Gm48510 | 0.000 | 0.000 | 0.000 |
| Gm32184 | 0.000 | 0.000 | 0.000 |
| Adtrp | 0.000 | 0.002 | 0.001 |
| 1700061E18F | 0.000 | 0.000 | 0.000 |
| Gm47118 | 0.000 | 0.000 | 0.000 |
| Gm15810 | 0.000 | 0.000 | 0.000 |
| Gm32939 | 0.000 | 0.000 | 0.000 |
| Gm47728 | 0.000 | 0.000 | 0.000 |
| Gm20751 | 0.000 | 0.000 | 0.001 |
| Gm33115 | 0.000 | 0.000 | 0.000 |
| Gm33489 | 0.000 | 0.000 | 0.000 |
| Gm29676 | 0.000 | 0.000 | 0.000 |
| Gm33958 | 0.000 | 0.000 | 0.000 |
| Gm9817 | 0.000 | 0.002 | 0.000 |
| Gm34466 | 0.000 | 0.000 | 0.000 |
| Gm47460 | 0.000 | 0.000 | 0.000 |
| 4930453C13F | 0.000 | 0.000 | 0.000 |
| Gm10113 | 0.000 | 0.000 | 0.000 |
| Gm47523 | 0.000 | 0.000 | 0.000 |
| 4930471G24I | 0.000 | 0.000 | 0.000 |
| G630093K05I | 0.000 | 0.000 | 0.000 |
| 4931429P17F | 0.000 | 0.000 | 0.000 |
| Gm36101 | 0.000 | 0.000 | 0.000 |
| A330033J07R | 0.000 | 0.000 | 0.000 |
| Gm36346 | 0.000 | 0.000 | 0.000 |

| | | | |
|---|---|---|---|
| Gm49291 | 0.000 | 0.001 | 0.000 |
| Gm48648 | 0.000 | 0.000 | 0.000 |
| Gm26651 | 0.000 | 0.000 | 0.000 |
| Gm48190 | 0.000 | 0.000 | 0.000 |
| Gm33424 | 0.000 | 0.000 | 0.000 |
| Gm2762 | 0.000 | 0.000 | 0.000 |
| Gm34278 | 0.000 | 0.000 | 0.000 |
| Gm48550 | 0.000 | 0.000 | 0.000 |
| Drd1 | 0.000 | 0.012 | 0.004 |
| Hrh2 | 0.000 | 0.000 | 0.000 |
| Gm16248 | 0.000 | 0.000 | 0.000 |
| Cdhr2 | 0.000 | 0.000 | 0.000 |
| Gprin1 | 0.000 | 0.000 | 0.002 |
| Sncb | 0.000 | 0.000 | 0.000 |
| 5330429C05F | 0.000 | 0.000 | 0.000 |
| Hk3 | 0.000 | 0.000 | 0.001 |
| Slc34a1 | 0.000 | 0.000 | 0.000 |
| Pfn3 | 0.000 | 0.000 | 0.000 |
| F12 | 0.000 | 0.000 | 0.000 |
| Gm46416 | 0.000 | 0.000 | 0.000 |
| Pitx1 | 0.000 | 0.000 | 0.001 |
| Gm47071 | 0.000 | 0.000 | 0.001 |
| Neurog1 | 0.000 | 0.000 | 0.000 |
| Slc25a48 | 0.000 | 0.000 | 0.000 |
| Lect2 | 0.000 | 0.000 | 0.000 |
| Gm45623 | 0.000 | 0.000 | 0.000 |
| Trpc7 | 0.000 | 0.000 | 0.000 |
| Spock1 | 0.000 | 0.000 | 0.000 |
| Gm47918 | 0.000 | 0.000 | 0.000 |
| Slc28a3 | 0.000 | 0.001 | 0.001 |
| 4930455J16R | 0.000 | 0.000 | 0.000 |
| Gm47947 | 0.000 | 0.000 | 0.000 |
| Gm47423 | 0.000 | 0.003 | 0.004 |
| Gm34354 | 0.000 | 0.000 | 0.002 |
| Gm47359 | 0.000 | 0.000 | 0.001 |
| Gm34672 | 0.000 | 0.001 | 0.002 |
| Gm34721 | 0.000 | 0.000 | 0.000 |
| 4930455M05 | 0.000 | 0.000 | 0.000 |
| Spata31d1d | 0.000 | 0.000 | 0.000 |
| Gm46419 | 0.000 | 0.000 | 0.000 |
| 4930528D03I | 0.000 | 0.000 | 0.000 |
| 1700014D04I | 0.000 | 0.000 | 0.000 |
| A530065N20I | 0.000 | 0.001 | 0.002 |
| Gm48397 | 0.000 | 0.000 | 0.000 |
| Gm5084 | 0.000 | 0.001 | 0.006 |
| Gm35333 | 0.000 | 0.000 | 0.001 |
| Gm48500 | 0.000 | 0.000 | 0.000 |
| BC051665 | 0.000 | 0.000 | 0.000 |

| | | | |
|---|---|---|---|
| Ctsj | 0.000 | 0.000 | 0.000 |
| Gm49357 | 0.000 | 0.000 | 0.000 |
| Gm48228 | 0.000 | 0.000 | 0.000 |
| Gm40977 | 0.000 | 0.000 | 0.000 |
| Gm19792 | 0.000 | 0.002 | 0.001 |
| Gm48795 | 0.000 | 0.000 | 0.000 |
| Fbp2 | 0.000 | 0.000 | 0.001 |
| Fbp1 | 0.000 | 0.000 | 0.000 |
| Hsd17b3 | 0.000 | 0.000 | 0.001 |
| 1700015C15F | 0.000 | 0.000 | 0.000 |
| Cntnap3 | 0.000 | 0.001 | 0.000 |
| Prss47 | 0.000 | 0.000 | 0.001 |
| Olfr466 | 0.000 | 0.000 | 0.000 |
| Nlrp4f | 0.000 | 0.000 | 0.000 |
| Gm47251 | 0.000 | 0.000 | 0.000 |
| Zfp708 | 0.000 | 0.000 | 0.000 |
| Gm49064 | 0.000 | 0.001 | 0.002 |
| Gm48824 | 0.000 | 0.000 | 0.000 |
| Gm28044 | 0.000 | 0.000 | 0.000 |
| Gm48900 | 0.000 | 0.000 | 0.000 |
| Gm26965 | 0.000 | 0.000 | 0.000 |
| Gm8016 | 0.000 | 0.000 | 0.000 |
| Gm35161 | 0.000 | 0.000 | 0.000 |
| Gm48819 | 0.000 | 0.000 | 0.000 |
| Gm26819 | 0.000 | 0.000 | 0.000 |
| D030007L05F | 0.000 | 0.000 | 0.000 |
| Gm36377 | 0.000 | 0.000 | 0.000 |
| Gm36426 | 0.000 | 0.000 | 0.000 |
| Gm36607 | 0.000 | 0.000 | 0.000 |
| Gm36529 | 0.000 | 0.001 | 0.006 |
| Irx1 | 0.000 | 0.004 | 0.007 |
| Irx4 | 0.000 | 0.000 | 0.000 |
| Eprn | 0.000 | 0.000 | 0.000 |
| Slc6a3 | 0.000 | 0.000 | 0.000 |
| Slc6a19 | 0.000 | 0.000 | 0.000 |
| Slc6a19os | 0.000 | 0.000 | 0.000 |
| Slc9a3 | 0.000 | 0.000 | 0.000 |
| Lrrc14b | 0.000 | 0.004 | 0.001 |
| CT009718.2 | 0.000 | 0.000 | 0.000 |
| CT009718.1 | 0.000 | 0.000 | 0.000 |
| Pcsk1 | 0.000 | 0.000 | 0.009 |
| 1700037F03F | 0.000 | 0.000 | 0.000 |
| Fam81b | 0.000 | 0.000 | 0.000 |
| Gm31219 | 0.000 | 0.000 | 0.000 |
| Gm38604 | 0.000 | 0.001 | 0.001 |
| Gm32067 | 0.000 | 0.000 | 0.000 |
| 5430425K12F | 0.000 | 0.000 | 0.000 |
| Gm48155 | 0.000 | 0.000 | 0.000 |

| | | | |
|---|---|---|---|
| 2310067P03F | 0.000 | 0.000 | 0.000 |
| Gm26803 | 0.000 | 0.000 | 0.000 |
| 2810049E08F | 0.000 | 0.000 | 0.000 |
| Gm17750 | 0.000 | 0.000 | 0.000 |
| Gm29680 | 0.000 | 0.000 | 0.000 |
| 4833422C13F | 0.000 | 0.003 | 0.003 |
| Acot12 | 0.000 | 0.000 | 0.000 |
| Ckmt2 | 0.000 | 0.000 | 0.000 |
| Gm20379 | 0.000 | 0.001 | 0.000 |
| C030017D09I | 0.000 | 0.000 | 0.000 |
| Ankrd34b | 0.000 | 0.000 | 0.000 |
| Gm4814 | 0.000 | 0.000 | 0.000 |
| A630019I02R | 0.000 | 0.001 | 0.000 |
| Bhmt | 0.000 | 0.000 | 0.000 |
| Bhmt2 | 0.000 | 0.000 | 0.001 |
| Dmgdh | 0.000 | 0.000 | 0.000 |
| AW495222 | 0.000 | 0.001 | 0.001 |
| Gm26527 | 0.000 | 0.000 | 0.000 |
| Gm32089 | 0.000 | 0.000 | 0.001 |
| Gm48102 | 0.000 | 0.000 | 0.000 |
| Gm48287 | 0.000 | 0.000 | 0.000 |
| Gm32305 | 0.000 | 0.000 | 0.000 |
| Pde8b | 0.000 | 0.002 | 0.002 |
| S100z | 0.000 | 0.001 | 0.000 |
| Gm48730 | 0.000 | 0.000 | 0.000 |
| Gm17190 | 0.000 | 0.000 | 0.000 |
| Gm29543 | 0.000 | 0.000 | 0.000 |
| Gm48133 | 0.000 | 0.000 | 0.000 |
| Gm48597 | 0.000 | 0.001 | 0.000 |
| C430039J16R | 0.000 | 0.000 | 0.000 |
| Gm33447 | 0.000 | 0.000 | 0.000 |
| Lncenc1 | 0.000 | 0.000 | 0.001 |
| Gm29501 | 0.000 | 0.000 | 0.000 |
| Gm35279 | 0.000 | 0.000 | 0.000 |
| Gm807 | 0.000 | 0.000 | 0.000 |
| BC001981 | 0.000 | 0.001 | 0.000 |
| Naip1 | 0.000 | 0.000 | 0.000 |
| Gm48596 | 0.000 | 0.000 | 0.000 |
| Gm29502 | 0.000 | 0.000 | 0.003 |
| 4932411K12F | 0.000 | 0.000 | 0.000 |
| Gm47533 | 0.000 | 0.000 | 0.000 |
| Gm29927 | 0.000 | 0.000 | 0.000 |
| Cd180 | 0.000 | 0.001 | 0.001 |
| 1700099I09R | 0.000 | 0.001 | 0.000 |
| Gm30551 | 0.000 | 0.000 | 0.000 |
| Gm47849 | 0.000 | 0.001 | 0.000 |
| Gm47851 | 0.000 | 0.000 | 0.000 |
| Rnf180 | 0.000 | 0.008 | 0.007 |

| | | | |
|---|---|---|---|
| Htr1a | 0.000 | 0.000 | 0.000 |
| Gm30411 | 0.000 | 0.000 | 0.000 |
| Gm28988 | 0.000 | 0.000 | 0.000 |
| Gm48684 | 0.000 | 0.001 | 0.002 |
| Gm31452 | 0.000 | 0.000 | 0.000 |
| AI197445 | 0.000 | 0.000 | 0.000 |
| Gm32090 | 0.000 | 0.000 | 0.000 |
| 1700006H21I | 0.000 | 0.000 | 0.000 |
| B230220B15F | 0.000 | 0.000 | 0.000 |
| Rab3c | 0.000 | 0.001 | 0.001 |
| Gapt | 0.000 | 0.000 | 0.000 |
| Gm33172 | 0.000 | 0.000 | 0.000 |
| Gm15288 | 0.000 | 0.000 | 0.000 |
| Gm15287 | 0.000 | 0.000 | 0.000 |
| Gm15324 | 0.000 | 0.000 | 0.000 |
| Gm48802 | 0.000 | 0.000 | 0.000 |
| Gm48837 | 0.000 | 0.000 | 0.000 |
| 2810403G07I | 0.000 | 0.000 | 0.001 |
| Gm48876 | 0.000 | 0.000 | 0.000 |
| Gm48879 | 0.000 | 0.000 | 0.000 |
| Ddx4 | 0.000 | 0.001 | 0.002 |
| Mcidas | 0.000 | 0.000 | 0.000 |
| Gzma | 0.000 | 0.000 | 0.000 |
| Gzmk | 0.000 | 0.000 | 0.000 |
| 4921509O07I | 0.000 | 0.000 | 0.000 |
| Hspb3 | 0.000 | 0.000 | 0.001 |
| Gm34586 | 0.000 | 0.000 | 0.000 |
| Gm47040 | 0.000 | 0.000 | 0.001 |
| 4930544M13 | 0.000 | 0.000 | 0.000 |
| 4930435F18F | 0.000 | 0.000 | 0.000 |
| Gm17509 | 0.000 | 0.000 | 0.000 |
| 4933413L06F | 0.000 | 0.000 | 0.000 |
| Gm10732 | 0.000 | 0.000 | 0.000 |
| Gm16263 | 0.000 | 0.000 | 0.005 |
| Gm47336 | 0.000 | 0.000 | 0.000 |
| 1700074H08I | 0.000 | 0.000 | 0.000 |
| Gm21761 | 0.000 | 0.000 | 0.000 |
| Gm36161 | 0.000 | 0.000 | 0.000 |
| Gm21188 | 0.000 | 0.000 | 0.000 |
| BC147527 | 0.000 | 0.000 | 0.000 |
| Gm48512 | 0.000 | 0.000 | 0.000 |
| Gm48001 | 0.000 | 0.000 | 0.000 |
| 4921501I09R | 0.000 | 0.000 | 0.000 |
| Gm31938 | 0.000 | 0.000 | 0.000 |
| Gm3625 | 0.000 | 0.000 | 0.001 |
| AC159282.1 | 0.000 | 0.000 | 0.001 |
| Gm48633 | 0.000 | 0.000 | 0.000 |
| 5033421B08F | 0.000 | 0.000 | 0.000 |

| | | | |
|---|---|---|---|
| Gm47391 | 0.000 | 0.001 | 0.001 |
| Gm46323 | 0.000 | 0.001 | 0.003 |
| Gm48764 | 0.000 | 0.000 | 0.000 |
| Gm48790 | 0.000 | 0.000 | 0.000 |
| Nt5c1b | 0.000 | 0.000 | 0.000 |
| Gm38407 | 0.000 | 0.000 | 0.002 |
| 9530020I12R | 0.000 | 0.001 | 0.000 |
| Gm47833 | 0.000 | 0.000 | 0.000 |
| Gm34237 | 0.000 | 0.000 | 0.000 |
| Platr19 | 0.000 | 0.000 | 0.000 |
| Gm48311 | 0.000 | 0.000 | 0.000 |
| Gm48558 | 0.000 | 0.001 | 0.000 |
| Gm36235 | 0.000 | 0.000 | 0.000 |
| Gm36372 | 0.000 | 0.001 | 0.000 |
| Gm48606 | 0.000 | 0.000 | 0.000 |
| Gm48605 | 0.000 | 0.000 | 0.000 |
| Ntsr2 | 0.000 | 0.000 | 0.000 |
| Gm48140 | 0.000 | 0.000 | 0.000 |
| Gm36495 | 0.000 | 0.001 | 0.001 |
| Atp6v1c2 | 0.000 | 0.000 | 0.000 |
| Gm48313 | 0.000 | 0.000 | 0.000 |
| Gm10479 | 0.000 | 0.000 | 0.000 |
| Gm49371 | 0.000 | 0.000 | 0.000 |
| Gm47997 | 0.000 | 0.000 | 0.000 |
| Grhl1 | 0.000 | 0.003 | 0.001 |
| Gm47713 | 0.000 | 0.000 | 0.000 |
| 4930480M12 | 0.000 | 0.000 | 0.000 |
| Gm29968 | 0.000 | 0.000 | 0.000 |
| 4930549C15F | 0.000 | 0.000 | 0.000 |
| Gm9866 | 0.000 | 0.000 | 0.000 |
| Gm45941 | 0.000 | 0.000 | 0.000 |
| Gm47872 | 0.000 | 0.000 | 0.000 |
| Allc | 0.000 | 0.000 | 0.000 |
| 4833405L11F | 0.000 | 0.000 | 0.000 |
| Gm31333 | 0.000 | 0.000 | 0.000 |
| Myt1l | 0.000 | 0.001 | 0.001 |
| Gpr22 | 0.000 | 0.001 | 0.003 |
| Gm33111 | 0.000 | 0.000 | 0.000 |
| 5430401H09I | 0.000 | 0.000 | 0.000 |
| Gm47948 | 0.000 | 0.000 | 0.000 |
| Cdhr3 | 0.000 | 0.000 | 0.000 |
| Gm16267 | 0.000 | 0.000 | 0.000 |
| Efcab10 | 0.000 | 0.000 | 0.000 |
| Gm48237 | 0.000 | 0.000 | 0.000 |
| Gm34047 | 0.000 | 0.000 | 0.000 |
| 9130015A21F | 0.000 | 0.000 | 0.000 |
| Lrrc72 | 0.000 | 0.000 | 0.000 |
| Gm34215 | 0.000 | 0.000 | 0.000 |

| | | | |
|---|---|---|---|
| Gm48613 | 0.000 | 0.000 | 0.000 |
| Gm29007 | 0.000 | 0.000 | 0.000 |
| Gm48715 | 0.000 | 0.007 | 0.004 |
| 4930428E07F | 0.000 | 0.000 | 0.000 |
| Gm34611 | 0.000 | 0.000 | 0.000 |
| Arl4aos | 0.000 | 0.001 | 0.000 |
| Gm47376 | 0.000 | 0.000 | 0.000 |
| Gm47373 | 0.000 | 0.000 | 0.000 |
| Gm46348 | 0.000 | 0.000 | 0.000 |
| AC113204.1 | 0.000 | 0.000 | 0.000 |
| Gm46349 | 0.000 | 0.000 | 0.000 |
| Gm33680 | 0.000 | 0.000 | 0.001 |
| Gm48522 | 0.000 | 0.000 | 0.000 |
| B230217J21R | 0.000 | 0.000 | 0.000 |
| Gm34304 | 0.000 | 0.000 | 0.000 |
| Gm43517 | 0.000 | 0.000 | 0.000 |
| Gm48779 | 0.000 | 0.000 | 0.000 |
| Gm26517 | 0.000 | 0.000 | 0.000 |
| Gm35135 | 0.000 | 0.000 | 0.000 |
| Gm35188 | 0.000 | 0.000 | 0.000 |
| 1700031P21F | 0.000 | 0.000 | 0.000 |
| 1700104L18F | 0.000 | 0.000 | 0.000 |
| Gm46328 | 0.000 | 0.000 | 0.000 |
| Gm10465 | 0.000 | 0.000 | 0.000 |
| Gm40884 | 0.000 | 0.000 | 0.000 |
| Aldoart2 | 0.000 | 0.000 | 0.000 |
| Insm2 | 0.000 | 0.000 | 0.000 |
| Nkx2-1 | 0.000 | 0.000 | 0.000 |
| Nkx2-9 | 0.000 | 0.000 | 0.000 |
| Gm16246 | 0.000 | 0.000 | 0.000 |
| Foxa1 | 0.000 | 0.000 | 0.000 |
| Ttc6 | 0.000 | 0.000 | 0.001 |
| Lrfn5 | 0.000 | 0.000 | 0.006 |
| Gm48422 | 0.000 | 0.000 | 0.000 |
| 4930471E15F | 0.000 | 0.000 | 0.000 |
| Gm47515 | 0.000 | 0.000 | 0.000 |
| Gm47518 | 0.000 | 0.000 | 0.000 |
| Gm31447 | 0.000 | 0.000 | 0.000 |
| Gm49383 | 0.000 | 0.000 | 0.000 |
| Gm9887 | 0.000 | 0.000 | 0.000 |
| Abhd12b | 0.000 | 0.000 | 0.000 |
| F730035M05I | 0.000 | 0.000 | 0.000 |
| Gm24474 | 0.000 | 0.001 | 0.000 |
| Gm32369 | 0.000 | 0.001 | 0.001 |
| Gm46355 | 0.000 | 0.000 | 0.001 |
| Tomm20l | 0.000 | 0.000 | 0.000 |
| Gm33016 | 0.000 | 0.000 | 0.000 |
| Gm40438 | 0.000 | 0.000 | 0.000 |

| | | | |
|---|---|---|---|
| 1700083H02I | 0.000 | 0.000 | 0.000 |
| 4930404H11I | 0.000 | 0.001 | 0.002 |
| Gm4756 | 0.000 | 0.000 | 0.000 |
| Six6 | 0.000 | 0.000 | 0.000 |
| D830013O20I | 0.000 | 0.000 | 0.000 |
| Gm48656 | 0.000 | 0.000 | 0.000 |
| Gm34016 | 0.000 | 0.000 | 0.000 |
| 2210039B01F | 0.000 | 0.001 | 0.010 |
| Gm8075 | 0.000 | 0.000 | 0.000 |
| Syt16 | 0.000 | 0.002 | 0.002 |
| Kcnh5 | 0.000 | 0.000 | 0.001 |
| Gphb5 | 0.000 | 0.000 | 0.000 |
| Tex21 | 0.000 | 0.000 | 0.000 |
| Gm35189 | 0.000 | 0.000 | 0.000 |
| Fam71d | 0.000 | 0.000 | 0.000 |
| Gm47766 | 0.000 | 0.002 | 0.001 |
| 4933406B15F | 0.000 | 0.000 | 0.000 |
| Gm47879 | 0.000 | 0.000 | 0.000 |
| Ccdc177 | 0.000 | 0.000 | 0.000 |
| Gm26796 | 0.000 | 0.000 | 0.000 |
| Gm3693 | 0.000 | 0.001 | 0.000 |
| Gm28370 | 0.000 | 0.000 | 0.000 |
| Gm16572 | 0.000 | 0.000 | 0.000 |
| Map3k9 | 0.000 | 0.000 | 0.000 |
| Gm48242 | 0.000 | 0.000 | 0.000 |
| Gm26623 | 0.000 | 0.000 | 0.000 |
| Heatr4 | 0.000 | 0.000 | 0.000 |
| Acot4 | 0.000 | 0.000 | 0.002 |
| 4732463B04F | 0.000 | 0.000 | 0.001 |
| Acot3 | 0.000 | 0.000 | 0.000 |
| Acot5 | 0.000 | 0.000 | 0.001 |
| Gm48709 | 0.000 | 0.000 | 0.000 |
| Gm17139 | 0.000 | 0.001 | 0.000 |
| Gm47819 | 0.000 | 0.000 | 0.000 |
| Gm40477 | 0.000 | 0.002 | 0.000 |
| Gm32296 | 0.000 | 0.000 | 0.000 |
| Gm26531 | 0.000 | 0.000 | 0.000 |
| Gm805 | 0.000 | 0.000 | 0.001 |
| 4732487G21I | 0.000 | 0.000 | 0.000 |
| Zdhhc22 | 0.000 | 0.000 | 0.000 |
| Tmem63c | 0.000 | 0.000 | 0.000 |
| Gm29362 | 0.000 | 0.000 | 0.000 |
| Gm21319 | 0.000 | 0.000 | 0.000 |
| Gm2001 | 0.000 | 0.000 | 0.000 |
| 4930544I03R | 0.000 | 0.000 | 0.000 |
| 5430427M07 | 0.000 | 0.000 | 0.000 |
| Tshr | 0.000 | 0.001 | 0.004 |
| Gm47804 | 0.000 | 0.000 | 0.000 |

| | | | |
|---|---|---|---|
| Gm40552 | 0.000 | 0.000 | 0.000 |
| Gm35326 | 0.000 | 0.001 | 0.005 |
| 3300002A11F | 0.000 | 0.000 | 0.000 |
| Gm47177 | 0.000 | 0.000 | 0.000 |
| Gm19951 | 0.000 | 0.000 | 0.000 |
| Kcnk13 | 0.000 | 0.000 | 0.001 |
| Gm48383 | 0.000 | 0.000 | 0.000 |
| Gpr68 | 0.000 | 0.000 | 0.000 |
| Gm47639 | 0.000 | 0.000 | 0.000 |
| Catsperb | 0.000 | 0.000 | 0.000 |
| Gm16339 | 0.000 | 0.009 | 0.001 |
| Chga | 0.000 | 0.001 | 0.000 |
| Gm20069 | 0.000 | 0.000 | 0.000 |
| Moap1 | 0.000 | 0.000 | 0.000 |
| Cox8c | 0.000 | 0.000 | 0.000 |
| Fam181a | 0.000 | 0.000 | 0.000 |
| Gm29508 | 0.000 | 0.000 | 0.000 |
| Gm15523 | 0.000 | 0.000 | 0.000 |
| Gm47267 | 0.000 | 0.000 | 0.000 |
| Serpina10 | 0.000 | 0.000 | 0.000 |
| Serpina6 | 0.000 | 0.000 | 0.000 |
| Serpina1c | 0.000 | 0.000 | 0.000 |
| Serpina11 | 0.000 | 0.000 | 0.000 |
| Serpina12 | 0.000 | 0.000 | 0.000 |
| Serpina3b | 0.000 | 0.000 | 0.000 |
| AC122556.1 | 0.000 | 0.000 | 0.001 |
| 4930408O17I | 0.000 | 0.000 | 0.000 |
| Tcl1b2 | 0.000 | 0.000 | 0.000 |
| Tunar | 0.000 | 0.000 | 0.000 |
| Gm2721 | 0.000 | 0.000 | 0.000 |
| Gm47796 | 0.000 | 0.001 | 0.001 |
| Gm48807 | 0.000 | 0.001 | 0.000 |
| Gm19554 | 0.000 | 0.000 | 0.000 |
| Gm47646 | 0.000 | 0.000 | 0.000 |
| Gm16087 | 0.000 | 0.000 | 0.000 |
| Gm2800 | 0.000 | 0.000 | 0.000 |
| Gm3234 | 0.000 | 0.000 | 0.000 |
| Bcl11b | 0.000 | 0.000 | 0.001 |
| Gm33467 | 0.000 | 0.000 | 0.000 |
| Gm34081 | 0.000 | 0.000 | 0.000 |
| 4930511J24R | 0.000 | 0.000 | 0.000 |
| 1700001K19F | 0.000 | 0.000 | 0.000 |
| 6030440G07I | 0.000 | 0.000 | 0.000 |
| 4930595D18I | 0.000 | 0.000 | 0.000 |
| Gm40578 | 0.000 | 0.000 | 0.000 |
| Gm36635 | 0.000 | 0.000 | 0.000 |
| Gm15996 | 0.000 | 0.000 | 0.000 |
| Rd3l | 0.000 | 0.000 | 0.000 |

| | | | |
|---|---|---|---|
| E330035G20I | 0.000 | 0.000 | 0.000 |
| Tmem179 | 0.000 | 0.000 | 0.000 |
| Igha | 0.000 | 0.000 | 0.000 |
| Ighg2c | 0.000 | 0.000 | 0.000 |
| Ighg2b | 0.000 | 0.000 | 0.000 |
| Ighg1 | 0.000 | 0.000 | 0.000 |
| Ighd | 0.000 | 0.000 | 0.000 |
| Gm30948 | 0.000 | 0.000 | 0.000 |
| Gm11027 | 0.000 | 0.001 | 0.001 |
| Ptprn2 | 0.000 | 0.000 | 0.000 |
| Tmem196 | 0.000 | 0.006 | 0.007 |
| Gm49231 | 0.000 | 0.000 | 0.000 |
| C6 | 0.000 | 0.000 | 0.000 |
| C9 | 0.000 | 0.000 | 0.000 |
| Gm16029 | 0.000 | 0.000 | 0.000 |
| Ranbp3l | 0.000 | 0.001 | 0.003 |
| Prlr | 0.000 | 0.003 | 0.055 |
| Gm49107 | 0.000 | 0.000 | 0.000 |
| 1700047G03I | 0.000 | 0.000 | 0.000 |
| Gm34759 | 0.000 | 0.000 | 0.002 |
| Gm49240 | 0.000 | 0.000 | 0.000 |
| 1810049J17R | 0.000 | 0.000 | 0.000 |
| C030047K22I | 0.000 | 0.000 | 0.000 |
| Gm49162 | 0.000 | 0.000 | 0.000 |
| AC124561.1 | 0.000 | 0.000 | 0.000 |
| Gm49191 | 0.000 | 0.000 | 0.000 |
| Gm35996 | 0.000 | 0.000 | 0.000 |
| 9230109A22I | 0.000 | 0.000 | 0.000 |
| 4930445E18I | 0.000 | 0.000 | 0.001 |
| Gm5468 | 0.000 | 0.000 | 0.000 |
| 11-Mar | 0.000 | 0.001 | 0.002 |
| Gm49267 | 0.000 | 0.000 | 0.000 |
| Gm31458 | 0.000 | 0.001 | 0.001 |
| Gm49233 | 0.000 | 0.001 | 0.000 |
| Dnah5 | 0.000 | 0.015 | 0.002 |
| 9630009A06I | 0.000 | 0.000 | 0.000 |
| 4930430F21I | 0.000 | 0.000 | 0.000 |
| 4930465K09I | 0.000 | 0.000 | 0.000 |
| Gm41289 | 0.000 | 0.000 | 0.000 |
| 4930592A05I | 0.000 | 0.000 | 0.000 |
| Gm41290 | 0.000 | 0.000 | 0.000 |
| 9430069I07R | 0.000 | 0.000 | 0.000 |
| Erich5 | 0.000 | 0.000 | 0.001 |
| Kcns2 | 0.000 | 0.000 | 0.001 |
| BC048602 | 0.000 | 0.000 | 0.001 |
| Gm41293 | 0.000 | 0.000 | 0.000 |
| Gm34150 | 0.000 | 0.000 | 0.000 |
| Gm49224 | 0.000 | 0.000 | 0.000 |

| | | | |
|---|---|---|---|
| Gm46515 | 0.000 | 0.000 | 0.000 |
| Gm10385 | 0.000 | 0.000 | 0.000 |
| Gm10384 | 0.000 | 0.000 | 0.000 |
| Gm34590 | 0.000 | 0.000 | 0.001 |
| Gm16136 | 0.000 | 0.000 | 0.000 |
| Grhl2 | 0.000 | 0.001 | 0.001 |
| Gm28221 | 0.000 | 0.001 | 0.000 |
| Gm15941 | 0.000 | 0.000 | 0.001 |
| Gm35019 | 0.000 | 0.000 | 0.000 |
| 2310043O21I | 0.000 | 0.000 | 0.000 |
| Gm45924 | 0.000 | 0.000 | 0.001 |
| Dcstamp | 0.000 | 0.001 | 0.000 |
| Dpys | 0.000 | 0.000 | 0.001 |
| Gm16291 | 0.000 | 0.000 | 0.000 |
| AU022793 | 0.000 | 0.000 | 0.000 |
| Gm26854 | 0.000 | 0.000 | 0.000 |
| AC158972.2 | 0.000 | 0.000 | 0.000 |
| Gm49271 | 0.000 | 0.000 | 0.000 |
| Gm10373 | 0.000 | 0.000 | 0.000 |
| Tmem74 | 0.000 | 0.000 | 0.000 |
| Trhr | 0.000 | 0.000 | 0.000 |
| Kcnv1 | 0.000 | 0.000 | 0.000 |
| Csmd3 | 0.000 | 0.000 | 0.006 |
| Gm41318 | 0.000 | 0.000 | 0.000 |
| Gm48913 | 0.000 | 0.000 | 0.000 |
| Gm19303 | 0.000 | 0.000 | 0.000 |
| 1700015H07I | 0.000 | 0.000 | 0.000 |
| Gm7489 | 0.000 | 0.000 | 0.000 |
| Mal2 | 0.000 | 0.001 | 0.000 |
| Gm49212 | 0.000 | 0.000 | 0.001 |
| Gm16006 | 0.000 | 0.000 | 0.002 |
| Fam83a | 0.000 | 0.000 | 0.001 |
| Gm2675 | 0.000 | 0.000 | 0.000 |
| Fer1l6 | 0.000 | 0.001 | 0.001 |
| Gm49356 | 0.000 | 0.000 | 0.000 |
| 4930544F09F | 0.000 | 0.000 | 0.000 |
| AC161172.1 | 0.000 | 0.001 | 0.000 |
| Gm19510 | 0.000 | 0.001 | 0.000 |
| Gm2682 | 0.000 | 0.000 | 0.000 |
| AC140304.1 | 0.000 | 0.000 | 0.000 |
| Gm30159 | 0.000 | 0.000 | 0.000 |
| Gm41335 | 0.000 | 0.000 | 0.000 |
| Gsdmc4 | 0.000 | 0.000 | 0.000 |
| Gm20717 | 0.000 | 0.000 | 0.000 |
| Gm30563 | 0.000 | 0.000 | 0.000 |
| Gm27153 | 0.000 | 0.000 | 0.000 |
| Gm27242 | 0.000 | 0.000 | 0.000 |
| Lrrc6 | 0.000 | 0.000 | 0.000 |

| | | | |
|---|---|---|---|
| Tg | 0.000 | 0.012 | 0.008 |
| Gm17140 | 0.000 | 0.000 | 0.000 |
| Gm17035 | 0.000 | 0.000 | 0.000 |
| AC087116.1 | 0.000 | 0.000 | 0.000 |
| Gm7125 | 0.000 | 0.000 | 0.000 |
| Col22a1 | 0.000 | 0.003 | 0.002 |
| Gm16308 | 0.000 | 0.000 | 0.000 |
| Kcnk9 | 0.000 | 0.000 | 0.000 |
| AC118008.1 | 0.000 | 0.000 | 0.000 |
| Adgrb1 | 0.000 | 0.011 | 0.003 |
| Slurp2 | 0.000 | 0.000 | 0.001 |
| Ly6d | 0.000 | 0.001 | 0.000 |
| 2010109I03R | 0.000 | 0.000 | 0.000 |
| AC139671.1 | 0.000 | 0.000 | 0.000 |
| Ly6i | 0.000 | 0.000 | 0.000 |
| Ly6c2 | 0.000 | 0.000 | 0.000 |
| Ly6g | 0.000 | 0.000 | 0.000 |
| Ly6f | 0.000 | 0.000 | 0.000 |
| AC119264.1 | 0.000 | 0.000 | 0.000 |
| Mroh6 | 0.000 | 0.000 | 0.000 |
| K230010J24R | 0.000 | 0.000 | 0.000 |
| Gm48952 | 0.000 | 0.000 | 0.000 |
| BC024139 | 0.000 | 0.000 | 0.000 |
| AC110211.1 | 0.000 | 0.000 | 0.000 |
| Scrt1 | 0.000 | 0.000 | 0.001 |
| Mb | 0.000 | 0.002 | 0.001 |
| Apol7a | 0.000 | 0.001 | 0.004 |
| Apol10a | 0.000 | 0.000 | 0.000 |
| Apol11a | 0.000 | 0.000 | 0.000 |
| Apol11b | 0.000 | 0.000 | 0.000 |
| Apol7e | 0.000 | 0.000 | 0.000 |
| Gm17638 | 0.000 | 0.000 | 0.001 |
| Cacng2 | 0.000 | 0.000 | 0.000 |
| A730060N03I | 0.000 | 0.000 | 0.000 |
| Pvalb | 0.000 | 0.000 | 0.001 |
| AL590144.3 | 0.000 | 0.001 | 0.001 |
| Sstr3 | 0.000 | 0.000 | 0.000 |
| Gm26634 | 0.000 | 0.000 | 0.000 |
| AL592169.1 | 0.000 | 0.000 | 0.000 |
| 1700027A07F | 0.000 | 0.000 | 0.000 |
| AL589670.5 | 0.000 | 0.000 | 0.000 |
| AL589670.1 | 0.000 | 0.002 | 0.002 |
| Gcat.1 | 0.000 | 0.000 | 0.000 |
| Galr3 | 0.000 | 0.000 | 0.000 |
| AL589670.4 | 0.000 | 0.000 | 0.000 |
| Gm20420 | 0.000 | 0.000 | 0.000 |
| Kcnj4 | 0.000 | 0.002 | 0.002 |
| Gm16575 | 0.000 | 0.000 | 0.000 |

| | | | |
|---|---|---|---|
| Npcd | 0.000 | 0.000 | 0.000 |
| AC113595.2 | 0.000 | 0.000 | 0.000 |
| Cacna1i | 0.000 | 0.000 | 0.000 |
| AC123059.1 | 0.000 | 0.000 | 0.000 |
| Grap2 | 0.000 | 0.000 | 0.001 |
| AC147041.1 | 0.000 | 0.000 | 0.000 |
| AC141880.1 | 0.000 | 0.001 | 0.001 |
| AC102334.1 | 0.000 | 0.001 | 0.001 |
| Dnajb7 | 0.000 | 0.000 | 0.000 |
| Gm17025 | 0.000 | 0.000 | 0.002 |
| AC102103.1 | 0.000 | 0.001 | 0.001 |
| 1700029P11F | 0.000 | 0.000 | 0.000 |
| 4930407I10R | 0.000 | 0.000 | 0.000 |
| Mei1 | 0.000 | 0.000 | 0.000 |
| AC104325.1 | 0.000 | 0.000 | 0.000 |
| 1500009C09F | 0.000 | 0.000 | 0.000 |
| Wbp2nl | 0.000 | 0.000 | 0.000 |
| Cyp2d10 | 0.000 | 0.000 | 0.000 |
| Cyp2d9 | 0.000 | 0.001 | 0.000 |
| AW121686 | 0.000 | 0.000 | 0.000 |
| AL591952.1 | 0.000 | 0.000 | 0.000 |
| Mpped1 | 0.000 | 0.000 | 0.000 |
| Efcab6 | 0.000 | 0.000 | 0.001 |
| AL626769.1 | 0.000 | 0.000 | 0.000 |
| Pnpla3 | 0.000 | 0.000 | 0.000 |
| 1810041L15F | 0.000 | 0.003 | 0.007 |
| AL611986.1 | 0.000 | 0.000 | 0.000 |
| Upk3a | 0.000 | 0.000 | 0.000 |
| Smc1b | 0.000 | 0.000 | 0.002 |
| Ribc2 | 0.000 | 0.000 | 0.000 |
| Wnt7b | 0.000 | 0.000 | 0.000 |
| AU022754 | 0.000 | 0.000 | 0.000 |
| Pkdrej | 0.000 | 0.001 | 0.002 |
| Gm15569 | 0.000 | 0.000 | 0.000 |
| AC124455.1 | 0.000 | 0.000 | 0.000 |
| Mlc1 | 0.000 | 0.000 | 0.000 |
| Mov10l1 | 0.000 | 0.000 | 0.000 |
| Panx2 | 0.000 | 0.001 | 0.000 |
| Adm2 | 0.000 | 0.000 | 0.000 |
| Miox | 0.000 | 0.000 | 0.000 |
| Tymp | 0.000 | 0.000 | 0.000 |
| AC137513.1 | 0.000 | 0.000 | 0.001 |
| Smgc | 0.000 | 0.000 | 0.000 |
| AC109617.1 | 0.000 | 0.000 | 0.000 |
| AC121517.2 | 0.000 | 0.000 | 0.000 |
| AC102910.1 | 0.000 | 0.000 | 0.000 |
| Dbx2 | 0.000 | 0.000 | 0.000 |
| Gm6961 | 0.000 | 0.000 | 0.000 |

| | | | |
|---|---|---|---|
| D030018L15F | 0.000 | 0.000 | 0.000 |
| Col2a1 | 0.000 | 0.003 | 0.002 |
| Gm26513 | 0.000 | 0.001 | 0.001 |
| Ccdc184 | 0.000 | 0.001 | 0.001 |
| Olfr287 | 0.000 | 0.000 | 0.000 |
| Olfr286 | 0.000 | 0.000 | 0.000 |
| Olfr285 | 0.000 | 0.000 | 0.000 |
| OR5BS1P | 0.000 | 0.000 | 0.000 |
| Lalba | 0.000 | 0.000 | 0.000 |
| Olfr279 | 0.000 | 0.001 | 0.001 |
| 4930415O20I | 0.000 | 0.001 | 0.000 |
| Gm29331 | 0.000 | 0.000 | 0.000 |
| Wnt1 | 0.000 | 0.000 | 0.004 |
| C1ql4 | 0.000 | 0.000 | 0.000 |
| Faim2 | 0.000 | 0.003 | 0.007 |
| Aqp2 | 0.000 | 0.000 | 0.001 |
| Gm17349 | 0.000 | 0.000 | 0.000 |
| Gm17057 | 0.000 | 0.001 | 0.006 |
| Fam186a | 0.000 | 0.000 | 0.000 |
| Gm21917 | 0.000 | 0.001 | 0.008 |
| Tmprss12 | 0.000 | 0.000 | 0.000 |
| Methig1 | 0.000 | 0.000 | 0.000 |
| Gm27209 | 0.000 | 0.001 | 0.000 |
| Gm47841 | 0.000 | 0.000 | 0.000 |
| Ankrd33 | 0.000 | 0.000 | 0.000 |
| Gm16031 | 0.000 | 0.000 | 0.000 |
| AC161812.1 | 0.000 | 0.000 | 0.000 |
| AC157583.2 | 0.000 | 0.000 | 0.000 |
| Krt87 | 0.000 | 0.000 | 0.000 |
| Krt83 | 0.000 | 0.000 | 0.000 |
| Krt90 | 0.000 | 0.000 | 0.000 |
| Krt75 | 0.000 | 0.000 | 0.000 |
| Krt5 | 0.000 | 0.000 | 0.000 |
| Krt71 | 0.000 | 0.000 | 0.000 |
| Krt77 | 0.000 | 0.000 | 0.000 |
| Krt4 | 0.000 | 0.000 | 0.000 |
| AC139844.1 | 0.000 | 0.000 | 0.000 |
| Gm9918 | 0.000 | 0.001 | 0.002 |
| Sp7 | 0.000 | 0.002 | 0.004 |
| Gm10337 | 0.000 | 0.000 | 0.000 |
| Gm28047 | 0.000 | 0.000 | 0.000 |
| AC124532.2 | 0.000 | 0.000 | 0.000 |
| Hoxc5 | 0.000 | 0.000 | 0.002 |
| Hoxc6 | 0.000 | 0.000 | 0.001 |
| Hoxc4 | 0.000 | 0.004 | 0.012 |
| Gm28876 | 0.000 | 0.000 | 0.000 |
| Gm10830 | 0.000 | 0.000 | 0.000 |
| D930007P13I | 0.000 | 0.000 | 0.000 |

| | | | |
|---|---|---|---|
| AC162693.1 | 0.000 | 0.000 | 0.000 |
| AC191865.2 | 0.000 | 0.000 | 0.000 |
| Olfr15 | 0.000 | 0.000 | 0.000 |
| Gm20695 | 0.000 | 0.000 | 0.000 |
| Dnase1 | 0.000 | 0.000 | 0.000 |
| Gm15835 | 0.000 | 0.000 | 0.000 |
| 4930562C15I | 0.000 | 0.001 | 0.001 |
| 12-Sep | 0.000 | 0.000 | 0.000 |
| Gm42477 | 0.000 | 0.001 | 0.001 |
| Sec14l5 | 0.000 | 0.000 | 0.000 |
| AC124490.1 | 0.000 | 0.000 | 0.000 |
| Tmem114 | 0.000 | 0.000 | 0.000 |
| AC156026.1 | 0.000 | 0.000 | 0.000 |
| AC136987.1 | 0.000 | 0.000 | 0.000 |
| AC154256.1 | 0.000 | 0.000 | 0.000 |
| Atf7ip2 | 0.000 | 0.001 | 0.001 |
| AC154311.1 | 0.000 | 0.000 | 0.000 |
| Tekt5 | 0.000 | 0.000 | 0.001 |
| Gm15558 | 0.000 | 0.000 | 0.000 |
| Tnp2 | 0.000 | 0.000 | 0.000 |
| Prm3 | 0.000 | 0.001 | 0.000 |
| Prm1 | 0.000 | 0.000 | 0.000 |
| Gm26822 | 0.000 | 0.001 | 0.000 |
| Gm21859 | 0.000 | 0.000 | 0.000 |
| Tnfrsf17 | 0.000 | 0.000 | 0.000 |
| Gm4279 | 0.000 | 0.000 | 0.000 |
| Gm9961 | 0.000 | 0.000 | 0.000 |
| AC117197.3 | 0.000 | 0.000 | 0.000 |
| AC117197.1 | 0.000 | 0.000 | 0.000 |
| 2310015D24I | 0.000 | 0.008 | 0.003 |
| Pla2g10os | 0.000 | 0.000 | 0.000 |
| AC154607.1 | 0.000 | 0.000 | 0.000 |
| Gm15868 | 0.000 | 0.000 | 0.000 |
| A630010A05I | 0.000 | 0.000 | 0.000 |
| AC154606.1 | 0.000 | 0.001 | 0.001 |
| Efcab1 | 0.000 | 0.003 | 0.004 |
| Gm21897 | 0.000 | 0.000 | 0.000 |
| Igll1 | 0.000 | 0.000 | 0.000 |
| Rimbp3 | 0.000 | 0.000 | 0.000 |
| Gm26635 | 0.000 | 0.000 | 0.000 |
| Serpind1 | 0.000 | 0.000 | 0.000 |
| AC078895.1 | 0.000 | 0.000 | 0.001 |
| Car15 | 0.000 | 0.001 | 0.001 |
| Tssk1 | 0.000 | 0.001 | 0.001 |
| Tssk2 | 0.000 | 0.000 | 0.000 |
| Gsc2 | 0.000 | 0.000 | 0.000 |
| 4933432I09R | 0.000 | 0.002 | 0.001 |
| AC118542.1 | 0.000 | 0.000 | 0.000 |

| | | | |
|---|---|---|---|
| Iglc1 | 0.000 | 0.000 | 0.000 |
| Iglc3 | 0.000 | 0.000 | 0.000 |
| Iglc2 | 0.000 | 0.000 | 0.001 |
| Map6d1 | 0.000 | 0.001 | 0.001 |
| Cyp2ab1 | 0.000 | 0.000 | 0.000 |
| AC087898.2 | 0.000 | 0.001 | 0.002 |
| AC123977.1 | 0.000 | 0.000 | 0.000 |
| Gm16863 | 0.000 | 0.000 | 0.001 |
| Gm26744 | 0.000 | 0.000 | 0.000 |
| 2610020F03F | 0.000 | 0.000 | 0.000 |
| Liph | 0.000 | 0.001 | 0.001 |
| 9230117E06F | 0.000 | 0.000 | 0.000 |
| Crygs | 0.000 | 0.004 | 0.001 |
| CT027991.1 | 0.000 | 0.000 | 0.000 |
| Kng1 | 0.000 | 0.000 | 0.000 |
| Rtp1 | 0.000 | 0.000 | 0.000 |
| Rtp2 | 0.000 | 0.000 | 0.000 |
| AC158397.1 | 0.000 | 0.000 | 0.000 |
| AC169509.3 | 0.000 | 0.000 | 0.000 |
| Gm4524 | 0.000 | 0.000 | 0.000 |
| Gmnc | 0.000 | 0.000 | 0.001 |
| Ostn | 0.000 | 0.001 | 0.001 |
| Uts2b | 0.000 | 0.000 | 0.000 |
| AC154438.1 | 0.000 | 0.001 | 0.000 |
| Gm26834 | 0.000 | 0.000 | 0.000 |
| 4632428C04F | 0.000 | 0.000 | 0.000 |
| Gm26562 | 0.000 | 0.001 | 0.002 |
| Cpn2 | 0.000 | 0.000 | 0.000 |
| AC130815.3 | 0.000 | 0.001 | 0.001 |
| AC125371.4 | 0.000 | 0.000 | 0.000 |
| AC125371.2 | 0.000 | 0.000 | 0.000 |
| AC161376.1 | 0.000 | 0.000 | 0.000 |
| AC161376.2 | 0.000 | 0.001 | 0.002 |
| Meltf | 0.000 | 0.000 | 0.000 |
| Bex6 | 0.000 | 0.000 | 0.000 |
| Slc51a | 0.000 | 0.000 | 0.000 |
| Zdhhc19 | 0.000 | 0.000 | 0.000 |
| Tnk2os | 0.000 | 0.000 | 0.000 |
| Muc4 | 0.000 | 0.001 | 0.000 |
| Muc20 | 0.000 | 0.000 | 0.000 |
| Smbd1 | 0.000 | 0.000 | 0.000 |
| Ropn1 | 0.000 | 0.000 | 0.000 |
| 1700119H24I | 0.000 | 0.001 | 0.000 |
| AC110166.1 | 0.000 | 0.004 | 0.002 |
| AC110166.2 | 0.000 | 0.002 | 0.001 |
| Gm15564 | 0.000 | 0.000 | 0.000 |
| Gm5416 | 0.000 | 0.000 | 0.001 |
| BC117090 | 0.000 | 0.000 | 0.000 |

| | | | |
|---|---|---|---|
| BC100530 | 0.000 | 0.002 | 0.001 |
| Gm15845 | 0.000 | 0.000 | 0.000 |
| Stfa2 | 0.000 | 0.001 | 0.000 |
| Casr | 0.000 | 0.000 | 0.000 |
| AC117662.4 | 0.000 | 0.000 | 0.000 |
| Ildr1 | 0.000 | 0.001 | 0.000 |
| Gm21987 | 0.000 | 0.002 | 0.001 |
| Gm9968 | 0.000 | 0.001 | 0.003 |
| Gm15713 | 0.000 | 0.000 | 0.000 |
| Drd3 | 0.000 | 0.000 | 0.000 |
| Sidt1 | 0.000 | 0.000 | 0.000 |
| Gm26732 | 0.000 | 0.000 | 0.000 |
| Cd200r4 | 0.000 | 0.000 | 0.000 |
| Cd200r2 | 0.000 | 0.000 | 0.000 |
| Btla | 0.000 | 0.001 | 0.000 |
| Gm609 | 0.000 | 0.000 | 0.000 |
| Gm17783 | 0.000 | 0.000 | 0.000 |
| Slc9c1 | 0.000 | 0.000 | 0.000 |
| Gcsam | 0.000 | 0.000 | 0.000 |
| BC016579 | 0.000 | 0.003 | 0.001 |
| Tmprss7 | 0.000 | 0.001 | 0.000 |
| Gm15640 | 0.000 | 0.000 | 0.001 |
| Gm15638 | 0.000 | 0.000 | 0.001 |
| Dppa4 | 0.000 | 0.000 | 0.000 |
| Retnla | 0.000 | 0.001 | 0.000 |
| Gm15518 | 0.000 | 0.000 | 0.000 |
| G730013B05I | 0.000 | 0.000 | 0.000 |
| Ccdc54 | 0.000 | 0.000 | 0.000 |
| Zpld1 | 0.000 | 0.000 | 0.000 |
| Adgrg7 | 0.000 | 0.000 | 0.000 |
| CT025521.3 | 0.000 | 0.000 | 0.000 |
| CT025521.2 | 0.000 | 0.002 | 0.001 |
| 4930461C15F | 0.000 | 0.000 | 0.000 |
| E330017A01F | 0.000 | 0.000 | 0.000 |
| AC159200.1 | 0.000 | 0.000 | 0.000 |
| Olfr172 | 0.000 | 0.000 | 0.000 |
| Olfr173 | 0.000 | 0.000 | 0.000 |
| Olfr183 | 0.000 | 0.001 | 0.000 |
| Olfr186 | 0.000 | 0.000 | 0.000 |
| Olfr190 | 0.000 | 0.000 | 0.000 |
| Olfr195 | 0.000 | 0.000 | 0.000 |
| Olfr204 | 0.000 | 0.000 | 0.000 |
| Epha6 | 0.000 | 0.000 | 0.000 |
| AC163285.1 | 0.000 | 0.000 | 0.000 |
| CT025155.1 | 0.000 | 0.000 | 0.000 |
| AC154568.1 | 0.000 | 0.000 | 0.000 |
| Csnka2ip | 0.000 | 0.000 | 0.000 |
| Htr1f | 0.000 | 0.000 | 0.000 |

| | | | |
|---|---|---|---|
| AC118240.2 | 0.000 | 0.000 | 0.000 |
| 1700010K23F | 0.000 | 0.000 | 0.001 |
| D16Ertd519e | 0.000 | 0.000 | 0.000 |
| AC154782.1 | 0.000 | 0.000 | 0.001 |
| AC145744.1 | 0.000 | 0.000 | 0.000 |
| Gm45030 | 0.000 | 0.000 | 0.001 |
| Gm30790 | 0.000 | 0.000 | 0.001 |
| 1700041M19 | 0.000 | 0.000 | 0.000 |
| Gm17333 | 0.000 | 0.000 | 0.000 |
| AC098883.2 | 0.000 | 0.000 | 0.000 |
| AC122375.2 | 0.000 | 0.000 | 0.000 |
| Gm49227 | 0.000 | 0.000 | 0.000 |
| Gm49226 | 0.000 | 0.000 | 0.000 |
| AC126936.1 | 0.000 | 0.000 | 0.000 |
| AC163352.1 | 0.000 | 0.000 | 0.000 |
| AC135964.2 | 0.000 | 0.002 | 0.002 |
| AC135964.1 | 0.000 | 0.001 | 0.002 |
| 2810407A14F | 0.000 | 0.001 | 0.000 |
| AC110241.2 | 0.000 | 0.000 | 0.000 |
| Gm10229 | 0.000 | 0.000 | 0.000 |
| 1110025L11F | 0.000 | 0.000 | 0.000 |
| Krtap8-1 | 0.000 | 0.000 | 0.000 |
| Krtap7-1 | 0.000 | 0.000 | 0.000 |
| Krtap11-1 | 0.000 | 0.000 | 0.000 |
| AC131339.4 | 0.000 | 0.000 | 0.000 |
| AC133505.1 | 0.000 | 0.000 | 0.000 |
| AC160759.1 | 0.000 | 0.000 | 0.000 |
| AC134560.1 | 0.000 | 0.000 | 0.000 |
| 4931406G06I | 0.000 | 0.000 | 0.000 |
| AC034116.3 | 0.000 | 0.000 | 0.000 |
| Olig1 | 0.000 | 0.000 | 0.000 |
| Gm15966 | 0.000 | 0.000 | 0.000 |
| Gm21970 | 0.000 | 0.000 | 0.000 |
| AC150035.3 | 0.000 | 0.000 | 0.000 |
| D430001F17F | 0.000 | 0.001 | 0.000 |
| AC144408.2 | 0.000 | 0.000 | 0.000 |
| AC144408.1 | 0.000 | 0.000 | 0.000 |
| Kcne2 | 0.000 | 0.001 | 0.001 |
| 4930563D23I | 0.000 | 0.000 | 0.000 |
| AC162305.1 | 0.000 | 0.000 | 0.000 |
| AC174448.1 | 0.000 | 0.000 | 0.000 |
| Gm26626 | 0.000 | 0.000 | 0.000 |
| 1700029J03R | 0.000 | 0.000 | 0.001 |
| Cldn14 | 0.000 | 0.000 | 0.000 |
| Sim2 | 0.000 | 0.000 | 0.008 |
| CT030190.1 | 0.000 | 0.000 | 0.000 |
| AC154546.2 | 0.000 | 0.001 | 0.001 |
| AC154691.1 | 0.000 | 0.000 | 0.000 |

| | | | |
|---|---|---|---|
| 2810404F17F | 0.000 | 0.000 | 0.000 |
| B3galt5 | 0.000 | 0.001 | 0.000 |
| Fam3b | 0.000 | 0.000 | 0.000 |
| Tmprss2 | 0.000 | 0.000 | 0.000 |
| Ripk4 | 0.000 | 0.001 | 0.000 |
| AC165953.3 | 0.000 | 0.001 | 0.000 |
| Cldn20 | 0.000 | 0.001 | 0.002 |
| Nox3 | 0.000 | 0.000 | 0.000 |
| AC140300.3 | 0.000 | 0.000 | 0.000 |
| Gm29050 | 0.000 | 0.000 | 0.000 |
| Ldhal6b | 0.000 | 0.001 | 0.001 |
| Gm26848 | 0.000 | 0.000 | 0.000 |
| AC168090.1 | 0.000 | 0.000 | 0.000 |
| Gm1604a | 0.000 | 0.000 | 0.000 |
| Ttll2 | 0.000 | 0.000 | 0.000 |
| Gm9992 | 0.000 | 0.000 | 0.001 |
| AC119998.5 | 0.000 | 0.000 | 0.000 |
| Tagap | 0.000 | 0.006 | 0.001 |
| AC164314.2 | 0.000 | 0.000 | 0.000 |
| Ccr6 | 0.000 | 0.000 | 0.000 |
| Gm16702 | 0.000 | 0.000 | 0.000 |
| T | 0.000 | 0.000 | 0.000 |
| Gm17087 | 0.000 | 0.000 | 0.000 |
| 6530411M01 | 0.000 | 0.000 | 0.000 |
| Gm17728 | 0.000 | 0.000 | 0.000 |
| Pabpc6 | 0.000 | 0.000 | 0.000 |
| 1700110C19F | 0.000 | 0.000 | 0.000 |
| A230009B12F | 0.000 | 0.000 | 0.000 |
| Gm16168 | 0.000 | 0.000 | 0.000 |
| Plg | 0.000 | 0.000 | 0.000 |
| Slc22a3 | 0.000 | 0.000 | 0.000 |
| Mas1 | 0.000 | 0.000 | 0.000 |
| Mrgprh | 0.000 | 0.000 | 0.000 |
| Gpr31b | 0.000 | 0.000 | 0.000 |
| Unc93a | 0.000 | 0.000 | 0.000 |
| Gm16050 | 0.000 | 0.000 | 0.000 |
| Gm7168 | 0.000 | 0.000 | 0.000 |
| CT010437.2 | 0.000 | 0.000 | 0.000 |
| Tcte3 | 0.000 | 0.001 | 0.000 |
| AC154507.2 | 0.000 | 0.000 | 0.000 |
| Gm5091 | 0.000 | 0.000 | 0.000 |
| AC154378.1 | 0.000 | 0.001 | 0.000 |
| AC154200.1 | 0.000 | 0.000 | 0.000 |
| Gm26873 | 0.000 | 0.000 | 0.000 |
| Fpr1 | 0.000 | 0.000 | 0.000 |
| Fpr2 | 0.000 | 0.001 | 0.002 |
| Vmn2r124 | 0.000 | 0.001 | 0.001 |
| Vmn2r92 | 0.000 | 0.000 | 0.000 |

| | | | |
|---|---|---|---|
| Vmn2r93 | 0.000 | 0.000 | 0.000 |
| Vmn2r95 | 0.000 | 0.000 | 0.000 |
| Vmn2r100 | 0.000 | 0.000 | 0.000 |
| Gm5145 | 0.000 | 0.000 | 0.000 |
| AC109204.1 | 0.000 | 0.000 | 0.000 |
| CT010433.1 | 0.000 | 0.000 | 0.000 |
| Gm26753 | 0.000 | 0.000 | 0.000 |
| Gm10509 | 0.000 | 0.000 | 0.000 |
| Gm10226 | 0.000 | 0.000 | 0.000 |
| Gm5493 | 0.000 | 0.000 | 0.001 |
| Mmp25 | 0.000 | 0.000 | 0.000 |
| Bicdl2 | 0.000 | 0.001 | 0.001 |
| Cldn6 | 0.000 | 0.000 | 0.000 |
| 1520401A03F | 0.000 | 0.001 | 0.000 |
| Gm49163 | 0.000 | 0.000 | 0.000 |
| Gm16275 | 0.000 | 0.000 | 0.000 |
| Prss32 | 0.000 | 0.000 | 0.000 |
| Gm15947 | 0.000 | 0.000 | 0.000 |
| Prss21 | 0.000 | 0.000 | 0.000 |
| Sbp | 0.000 | 0.000 | 0.000 |
| Sbpl | 0.000 | 0.000 | 0.000 |
| Prss30 | 0.000 | 0.000 | 0.001 |
| Prss22 | 0.000 | 0.000 | 0.000 |
| Abca17 | 0.000 | 0.000 | 0.000 |
| Bricd5 | 0.000 | 0.000 | 0.000 |
| Hs3st6 | 0.000 | 0.000 | 0.000 |
| Meiob | 0.000 | 0.000 | 0.000 |
| Igfals | 0.000 | 0.000 | 0.001 |
| Ccdc154 | 0.000 | 0.000 | 0.000 |
| Prss28 | 0.000 | 0.000 | 0.000 |
| Tpsab1 | 0.000 | 0.001 | 0.001 |
| Tpsg1 | 0.000 | 0.000 | 0.000 |
| Tekt4 | 0.000 | 0.000 | 0.000 |
| Sstr5 | 0.000 | 0.000 | 0.000 |
| Gm26694 | 0.000 | 0.000 | 0.000 |
| Rhbdl1 | 0.000 | 0.000 | 0.000 |
| A930017K11F | 0.000 | 0.000 | 0.000 |
| D630044L22F | 0.000 | 0.000 | 0.000 |
| 1700022N22I | 0.000 | 0.001 | 0.000 |
| Tmem8 | 0.000 | 0.002 | 0.002 |
| Pdia2 | 0.000 | 0.000 | 0.000 |
| 1700049J03R | 0.000 | 0.000 | 0.000 |
| Gm17218 | 0.000 | 0.000 | 0.000 |
| Gm20468 | 0.000 | 0.000 | 0.000 |
| Zbtb9 | 0.000 | 0.000 | 0.000 |
| AC125141.1 | 0.000 | 0.001 | 0.001 |
| AC125141.3 | 0.000 | 0.000 | 0.000 |
| Grm4 | 0.000 | 0.000 | 0.000 |

| | | | |
|---|---|---|---|
| AC127341.5 | 0.000 | 0.000 | 0.000 |
| Gm15458 | 0.000 | 0.000 | 0.000 |
| Gm15598 | 0.000 | 0.000 | 0.001 |
| Armc12 | 0.000 | 0.000 | 0.000 |
| Clps | 0.000 | 0.000 | 0.000 |
| Lhfpl5 | 0.000 | 0.000 | 0.000 |
| Pnpla1os | 0.000 | 0.000 | 0.000 |
| Pnpla1 | 0.000 | 0.000 | 0.001 |
| Gm16191 | 0.000 | 0.000 | 0.001 |
| Gm16194 | 0.000 | 0.000 | 0.000 |
| Gm26885 | 0.000 | 0.001 | 0.003 |
| Tmem217 | 0.000 | 0.000 | 0.000 |
| Tbc1d22bos | 0.000 | 0.000 | 0.000 |
| Glp1r | 0.000 | 0.000 | 0.000 |
| Umodl1 | 0.000 | 0.000 | 0.000 |
| Tmprss3 | 0.000 | 0.000 | 0.000 |
| Ubash3a | 0.000 | 0.000 | 0.000 |
| 4833413E03F | 0.000 | 0.000 | 0.000 |
| Cryaa | 0.000 | 0.000 | 0.000 |
| Gm26858 | 0.000 | 0.000 | 0.000 |
| Rasal3 | 0.000 | 0.000 | 0.000 |
| Pglyrp2 | 0.000 | 0.000 | 0.000 |
| Cyp4f15 | 0.000 | 0.000 | 0.000 |
| Gm26693 | 0.000 | 0.000 | 0.000 |
| Cyp4f14 | 0.000 | 0.000 | 0.000 |
| Morc2b | 0.000 | 0.001 | 0.002 |
| Olfr239 | 0.000 | 0.000 | 0.000 |
| Olfr1564 | 0.000 | 0.001 | 0.001 |
| Olfr63 | 0.000 | 0.000 | 0.000 |
| Pram1 | 0.000 | 0.002 | 0.002 |
| 4931413I07R | 0.000 | 0.000 | 0.000 |
| H2-Oa | 0.000 | 0.000 | 0.000 |
| H2-DMb2 | 0.000 | 0.000 | 0.000 |
| Gm20496 | 0.000 | 0.000 | 0.000 |
| Gm20506 | 0.000 | 0.000 | 0.000 |
| H2-Eb2 | 0.000 | 0.000 | 0.000 |
| Btnl2 | 0.000 | 0.000 | 0.000 |
| BC051142 | 0.000 | 0.001 | 0.005 |
| Btnl6 | 0.000 | 0.000 | 0.000 |
| Gm20461 | 0.000 | 0.000 | 0.000 |
| Cyp21a1 | 0.000 | 0.000 | 0.002 |
| Slc44a4 | 0.000 | 0.000 | 0.000 |
| Ly6g6f | 0.000 | 0.000 | 0.001 |
| Ly6g5c | 0.000 | 0.000 | 0.000 |
| Lta | 0.000 | 0.001 | 0.000 |
| Gm16181 | 0.000 | 0.000 | 0.000 |
| H2-Q1 | 0.000 | 0.000 | 0.000 |
| H2-Q2 | 0.000 | 0.000 | 0.001 |

| | | | |
|---|---|---|---|
| Gm19553 | 0.000 | 0.000 | 0.000 |
| Sfta2 | 0.000 | 0.000 | 0.001 |
| Gm9573 | 0.000 | 0.000 | 0.000 |
| Gm20443 | 0.000 | 0.000 | 0.000 |
| Gm20442 | 0.000 | 0.001 | 0.011 |
| 4833427F10F | 0.000 | 0.000 | 0.000 |
| Gm20508 | 0.000 | 0.001 | 0.001 |
| Gm6034 | 0.000 | 0.000 | 0.000 |
| Gm11127 | 0.000 | 0.000 | 0.000 |
| Gm19684 | 0.000 | 0.000 | 0.000 |
| Gm8909 | 0.000 | 0.000 | 0.000 |
| Gm20478 | 0.000 | 0.000 | 0.000 |
| Gm20546 | 0.000 | 0.000 | 0.000 |
| H2-M9 | 0.000 | 0.000 | 0.000 |
| H2-M10.5 | 0.000 | 0.000 | 0.000 |
| Trim15 | 0.000 | 0.000 | 0.000 |
| Trim40 | 0.000 | 0.000 | 0.000 |
| Trim31 | 0.000 | 0.000 | 0.000 |
| 1700031A10F | 0.000 | 0.000 | 0.000 |
| 2410137M14 | 0.000 | 0.000 | 0.001 |
| Mog | 0.000 | 0.000 | 0.000 |
| Olfr90 | 0.000 | 0.001 | 0.002 |
| Olfr93 | 0.000 | 0.000 | 0.000 |
| Ubd | 0.000 | 0.000 | 0.000 |
| Olfr95 | 0.000 | 0.000 | 0.000 |
| Olfr96 | 0.000 | 0.000 | 0.000 |
| Olfr107 | 0.000 | 0.001 | 0.001 |
| H2-M2 | 0.000 | 0.000 | 0.000 |
| Olfr110 | 0.000 | 0.000 | 0.000 |
| Olfr111 | 0.000 | 0.000 | 0.000 |
| Olfr113 | 0.000 | 0.000 | 0.000 |
| Olfr125 | 0.000 | 0.000 | 0.000 |
| Olfr127 | 0.000 | 0.000 | 0.000 |
| Olfr130 | 0.000 | 0.000 | 0.000 |
| Olfr131 | 0.000 | 0.000 | 0.000 |
| Esp18 | 0.000 | 0.000 | 0.000 |
| Esp15 | 0.000 | 0.000 | 0.000 |
| Gm44501 | 0.000 | 0.000 | 0.000 |
| Crisp2 | 0.000 | 0.000 | 0.000 |
| Opn5 | 0.000 | 0.000 | 0.000 |
| Adgrf4 | 0.000 | 0.001 | 0.000 |
| Ankrd66 | 0.000 | 0.000 | 0.000 |
| 1700071M16 | 0.000 | 0.000 | 0.000 |
| Spats1 | 0.000 | 0.000 | 0.000 |
| Tcte1 | 0.000 | 0.000 | 0.000 |
| Tmem151b | 0.000 | 0.003 | 0.002 |
| Gm17080 | 0.000 | 0.000 | 0.000 |
| Gm16172 | 0.000 | 0.000 | 0.000 |

| | | | |
|---|---|---|---|
| 1600014C23F | 0.000 | 0.000 | 0.000 |
| F630040K05F | 0.000 | 0.001 | 0.001 |
| Dlk2 | 0.000 | 0.002 | 0.002 |
| Crip3 | 0.000 | 0.002 | 0.002 |
| Slc22a7 | 0.000 | 0.000 | 0.000 |
| Ttbk1 | 0.000 | 0.001 | 0.001 |
| Prph2 | 0.000 | 0.000 | 0.000 |
| Gm21981 | 0.000 | 0.000 | 0.000 |
| Gm15556 | 0.000 | 0.000 | 0.001 |
| 1700122O11I | 0.000 | 0.000 | 0.000 |
| 9830107B12F | 0.000 | 0.000 | 0.000 |
| A530064D06I | 0.000 | 0.000 | 0.000 |
| Trem3 | 0.000 | 0.000 | 0.000 |
| Treml2 | 0.000 | 0.000 | 0.000 |
| B430306N03I | 0.000 | 0.000 | 0.000 |
| Treml1 | 0.000 | 0.000 | 0.001 |
| Gm45330 | 0.000 | 0.000 | 0.000 |
| Gm16555 | 0.000 | 0.000 | 0.000 |
| Dazl | 0.000 | 0.000 | 0.001 |
| Gm19585 | 0.000 | 0.000 | 0.000 |
| Gm27217 | 0.000 | 0.000 | 0.000 |
| Pp2d1 | 0.000 | 0.000 | 0.001 |
| 4932415M13 | 0.000 | 0.000 | 0.000 |
| Sult1c2 | 0.000 | 0.000 | 0.000 |
| Gm26547 | 0.000 | 0.001 | 0.000 |
| Plin5 | 0.000 | 0.001 | 0.000 |
| Acsbg2 | 0.000 | 0.001 | 0.000 |
| Acer1 | 0.000 | 0.000 | 0.000 |
| Pspn | 0.000 | 0.001 | 0.001 |
| Slc25a41 | 0.000 | 0.000 | 0.000 |
| Gm17168 | 0.000 | 0.000 | 0.000 |
| Cd70 | 0.000 | 0.000 | 0.000 |
| Tnfsf14 | 0.000 | 0.000 | 0.000 |
| Cntnap5c | 0.000 | 0.000 | 0.000 |
| Pdzph1 | 0.000 | 0.000 | 0.000 |
| A930002H24I | 0.000 | 0.000 | 0.000 |
| AU016765 | 0.000 | 0.000 | 0.000 |
| Gm17133 | 0.000 | 0.000 | 0.000 |
| Tmem232 | 0.000 | 0.000 | 0.000 |
| Themis3 | 0.000 | 0.001 | 0.000 |
| 1700016K05F | 0.000 | 0.000 | 0.000 |
| Gm47471 | 0.000 | 0.000 | 0.000 |
| Tmem200c | 0.000 | 0.001 | 0.004 |
| 2410021H03I | 0.000 | 0.000 | 0.001 |
| Akain1 | 0.000 | 0.000 | 0.000 |
| 5031415H12I | 0.000 | 0.000 | 0.000 |
| Gm26561 | 0.000 | 0.000 | 0.000 |
| Gm16519 | 0.000 | 0.000 | 0.000 |

| | | | |
|---|---|---|---|
| Togaram2 | 0.000 | 0.001 | 0.001 |
| BC027072 | 0.000 | 0.000 | 0.000 |
| Alk | 0.000 | 0.002 | 0.001 |
| Nlrc4 | 0.000 | 0.001 | 0.001 |
| Sult6b1 | 0.000 | 0.000 | 0.000 |
| Ttc39d | 0.000 | 0.000 | 0.000 |
| C230072F16F | 0.000 | 0.000 | 0.001 |
| Gm29052 | 0.000 | 0.000 | 0.000 |
| Haao | 0.000 | 0.000 | 0.000 |
| Abcg8 | 0.000 | 0.000 | 0.000 |
| 2010106C02F | 0.000 | 0.000 | 0.000 |
| Atp6v1e2 | 0.000 | 0.001 | 0.001 |
| Stpg4 | 0.000 | 0.001 | 0.000 |
| Gm26612 | 0.000 | 0.000 | 0.000 |
| Gtf2a1l | 0.000 | 0.001 | 0.001 |
| Lhcgr | 0.000 | 0.000 | 0.000 |
| Zfy1 | 0.000 | 0.000 | 0.001 |
| Gm28587 | 0.000 | 0.000 | 0.000 |
| Gm21778 | 0.000 | 0.000 | 0.000 |
| Sly | 0.000 | 0.000 | 0.000 |
| Gm21860 | 0.000 | 0.000 | 0.000 |
| G430049J08F | 0.000 | 0.000 | 0.000 |
| Lyzl1 | 0.000 | 0.000 | 0.000 |
| Gm10556 | 0.000 | 0.000 | 0.001 |
| Gm17036 | 0.000 | 0.000 | 0.000 |
| 4921524L21F | 0.000 | 0.000 | 0.000 |
| Armc4 | 0.000 | 0.000 | 0.000 |
| Gm17430 | 0.000 | 0.001 | 0.001 |
| Mir133a-1hg | 0.000 | 0.000 | 0.000 |
| Gm29200 | 0.000 | 0.000 | 0.000 |
| Hrh4 | 0.000 | 0.000 | 0.000 |
| Aqp4 | 0.000 | 0.000 | 0.000 |
| Gm15485 | 0.000 | 0.000 | 0.000 |
| Gm15328 | 0.000 | 0.001 | 0.000 |
| Dsc3 | 0.000 | 0.010 | 0.008 |
| Dsg1c | 0.000 | 0.000 | 0.003 |
| Dsg1a | 0.000 | 0.000 | 0.000 |
| Dsg3 | 0.000 | 0.000 | 0.000 |
| Mep1b | 0.000 | 0.000 | 0.000 |
| Klhl14 | 0.000 | 0.000 | 0.000 |
| 4930426D05I | 0.000 | 0.000 | 0.000 |
| Nol4 | 0.000 | 0.000 | 0.001 |
| 0710001A04F | 0.000 | 0.000 | 0.000 |
| Rit2 | 0.000 | 0.000 | 0.000 |
| Gm26823 | 0.000 | 0.001 | 0.001 |
| Gpr17 | 0.000 | 0.000 | 0.001 |
| Myo7b | 0.000 | 0.000 | 0.000 |
| Proc | 0.000 | 0.000 | 0.000 |

| | | | |
|---|---|---|---|
| Gm35060 | 0.000 | 0.000 | 0.000 |
| 4930455D15I | 0.000 | 0.000 | 0.000 |
| Pkd2l2 | 0.000 | 0.001 | 0.002 |
| Wnt8a | 0.000 | 0.000 | 0.000 |
| Mzb1 | 0.000 | 0.000 | 0.000 |
| Psd2 | 0.000 | 0.000 | 0.001 |
| Slc4a9 | 0.000 | 0.000 | 0.000 |
| Eif4ebp3 | 0.000 | 0.000 | 0.000 |
| Gm37751 | 0.000 | 0.000 | 0.000 |
| Pcdha2 | 0.000 | 0.000 | 0.000 |
| Pcdha3 | 0.000 | 0.000 | 0.000 |
| Pcdha6 | 0.000 | 0.000 | 0.000 |
| Pcdha8 | 0.000 | 0.000 | 0.000 |
| Pcdha9 | 0.000 | 0.000 | 0.000 |
| Pcdha11 | 0.000 | 0.000 | 0.000 |
| Pcdha11.1 | 0.000 | 0.000 | 0.001 |
| Pcdha12 | 0.000 | 0.000 | 0.000 |
| Gm37446 | 0.000 | 0.000 | 0.000 |
| 4930517L18F | 0.000 | 0.000 | 0.000 |
| Slc25a2 | 0.000 | 0.001 | 0.001 |
| Gm37118 | 0.000 | 0.000 | 0.000 |
| Gm37165 | 0.000 | 0.000 | 0.000 |
| Gm15334 | 0.000 | 0.000 | 0.000 |
| Gm15336 | 0.000 | 0.000 | 0.000 |
| Kctd16 | 0.000 | 0.001 | 0.001 |
| Grxcr2 | 0.000 | 0.000 | 0.000 |
| Sh3rf2 | 0.000 | 0.028 | 0.003 |
| Gm5689 | 0.000 | 0.004 | 0.001 |
| Plac8l1 | 0.000 | 0.000 | 0.000 |
| Pou4f3 | 0.000 | 0.000 | 0.000 |
| Gpr151 | 0.000 | 0.000 | 0.000 |
| Spink1 | 0.000 | 0.000 | 0.000 |
| Spink5 | 0.000 | 0.001 | 0.000 |
| Npy6r | 0.000 | 0.000 | 0.000 |
| Myot | 0.000 | 0.000 | 0.000 |
| A330093E20F | 0.000 | 0.000 | 0.000 |
| Trim36 | 0.000 | 0.004 | 0.001 |
| Lvrn | 0.000 | 0.000 | 0.010 |
| Arl14epl | 0.000 | 0.000 | 0.001 |
| Zfp474 | 0.000 | 0.000 | 0.000 |
| 1700034E13F | 0.000 | 0.001 | 0.002 |
| Redrum | 0.000 | 0.000 | 0.000 |
| Tex43 | 0.000 | 0.000 | 0.000 |
| Gm15345 | 0.000 | 0.000 | 0.000 |
| Ctxn3 | 0.000 | 0.004 | 0.000 |
| Gm26507 | 0.000 | 0.001 | 0.001 |
| Adamts19 | 0.000 | 0.001 | 0.031 |
| Chsy3 | 0.000 | 0.006 | 0.004 |

| | | | |
|---|---|---|---|
| Myoz3 | 0.000 | 0.000 | 0.000 |
| Slc6a7 | 0.000 | 0.008 | 0.008 |
| Cdx1 | 0.000 | 0.000 | 0.000 |
| Pde6a | 0.000 | 0.002 | 0.001 |
| Arhgef37 | 0.000 | 0.000 | 0.000 |
| Gm38165 | 0.000 | 0.000 | 0.000 |
| Htr4 | 0.000 | 0.000 | 0.000 |
| Spink13 | 0.000 | 0.000 | 0.000 |
| St8sia3os | 0.000 | 0.000 | 0.000 |
| St8sia3 | 0.000 | 0.000 | 0.000 |
| Cplx4 | 0.000 | 0.000 | 0.000 |
| Mc5r | 0.000 | 0.001 | 0.000 |
| Dcc | 0.000 | 0.006 | 0.002 |
| Mro | 0.000 | 0.000 | 0.000 |
| D730045A05I | 0.000 | 0.000 | 0.000 |
| Gm9925 | 0.000 | 0.000 | 0.000 |
| St8sia5 | 0.000 | 0.000 | 0.000 |
| Loxhd1 | 0.000 | 0.000 | 0.000 |
| Pstpip2 | 0.000 | 0.000 | 0.001 |
| Slc14a1 | 0.000 | 0.001 | 0.000 |
| Slc14a2 | 0.000 | 0.000 | 0.001 |
| Gm27239 | 0.000 | 0.000 | 0.000 |
| Galr1 | 0.000 | 0.000 | 0.000 |
| 2210420H20I | 0.000 | 0.000 | 0.000 |
| Cndp1 | 0.000 | 0.000 | 0.000 |
| Neto1 | 0.000 | 0.001 | 0.006 |
| Cbln2 | 0.000 | 0.001 | 0.008 |
| Gm5096 | 0.000 | 0.000 | 0.000 |
| Cd226 | 0.000 | 0.000 | 0.000 |
| Dok6 | 0.000 | 0.001 | 0.005 |
| 1700030N03I | 0.000 | 0.000 | 0.001 |
| Gal | 0.000 | 0.003 | 0.003 |
| Gm48683 | 0.000 | 0.000 | 0.000 |
| Aldh3b2 | 0.000 | 0.000 | 0.000 |
| Tbx10 | 0.000 | 0.000 | 0.000 |
| Gstp3 | 0.000 | 0.000 | 0.000 |
| Cabp2 | 0.000 | 0.000 | 0.000 |
| Cabp4 | 0.000 | 0.000 | 0.000 |
| 2010003K11F | 0.000 | 0.000 | 0.000 |
| Gm21844 | 0.000 | 0.000 | 0.000 |
| Ccdc87 | 0.000 | 0.000 | 0.000 |
| Peli3 | 0.000 | 0.006 | 0.003 |
| Catsper1 | 0.000 | 0.000 | 0.000 |
| Tsga10ip | 0.000 | 0.000 | 0.000 |
| Ovol1 | 0.000 | 0.000 | 0.000 |
| Tmem262 | 0.000 | 0.000 | 0.000 |
| Naaladl1 | 0.000 | 0.000 | 0.000 |
| Majin | 0.000 | 0.000 | 0.000 |

| | | | |
|---|---|---|---|
| Slc22a12 | 0.000 | 0.000 | 0.000 |
| Kcnk4 | 0.000 | 0.000 | 0.000 |
| Lgals12 | 0.000 | 0.000 | 0.000 |
| Hrasls5 | 0.000 | 0.001 | 0.000 |
| Slc22a29 | 0.000 | 0.000 | 0.000 |
| Slc22a30 | 0.000 | 0.000 | 0.000 |
| Slc22a6 | 0.000 | 0.000 | 0.000 |
| Gm45736 | 0.000 | 0.000 | 0.000 |
| Scgb1a1 | 0.000 | 0.000 | 0.003 |
| Gm26859 | 0.000 | 0.000 | 0.000 |
| Ppp1r32 | 0.000 | 0.000 | 0.000 |
| 2210404E10F | 0.000 | 0.000 | 0.000 |
| Cd6 | 0.000 | 0.000 | 0.000 |
| Zp1 | 0.000 | 0.000 | 0.000 |
| Ptgdr2 | 0.000 | 0.000 | 0.000 |
| Ms4a10 | 0.000 | 0.000 | 0.001 |
| Ms4a15 | 0.000 | 0.000 | 0.000 |
| 1700017D01I | 0.000 | 0.000 | 0.000 |
| Ms4a1 | 0.000 | 0.000 | 0.000 |
| Ms4a14 | 0.000 | 0.000 | 0.000 |
| Ms4a4a | 0.000 | 0.000 | 0.000 |
| Ms4a4c | 0.000 | 0.000 | 0.000 |
| Ms4a4b | 0.000 | 0.000 | 0.000 |
| Gm37387 | 0.000 | 0.000 | 0.000 |
| Gm8369 | 0.000 | 0.000 | 0.000 |
| Ms4a2 | 0.000 | 0.000 | 0.000 |
| Ms4a3 | 0.000 | 0.000 | 0.000 |
| Olfr1419 | 0.000 | 0.000 | 0.000 |
| Olfr1423 | 0.000 | 0.000 | 0.000 |
| Olfr1427 | 0.000 | 0.000 | 0.000 |
| Olfr1440 | 0.000 | 0.000 | 0.000 |
| Gm47242 | 0.000 | 0.000 | 0.000 |
| Gm4952 | 0.000 | 0.000 | 0.001 |
| Keg1 | 0.000 | 0.000 | 0.000 |
| Gm15962 | 0.000 | 0.000 | 0.000 |
| Cntf | 0.000 | 0.000 | 0.000 |
| Olfr1444 | 0.000 | 0.000 | 0.000 |
| Olfr1445 | 0.000 | 0.000 | 0.000 |
| Olfr1448 | 0.000 | 0.000 | 0.000 |
| Olfr1459 | 0.000 | 0.000 | 0.000 |
| Olfr1469 | 0.000 | 0.000 | 0.000 |
| Olfr1484 | 0.000 | 0.000 | 0.000 |
| Olfr1489 | 0.000 | 0.000 | 0.000 |
| Olfr1505 | 0.000 | 0.000 | 0.000 |
| 1500015L24F | 0.000 | 0.000 | 0.000 |
| C730002L08F | 0.000 | 0.000 | 0.000 |
| Gm27151 | 0.000 | 0.000 | 0.000 |
| C330002G04I | 0.000 | 0.000 | 0.000 |

| | | | |
|---|---|---|---|
| 1700028P14F | 0.000 | 0.000 | 0.000 |
| Foxd4 | 0.000 | 0.000 | 0.000 |
| Dmrt3 | 0.000 | 0.000 | 0.000 |
| 2610016A17F | 0.000 | 0.000 | 0.000 |
| Dmrt2 | 0.000 | 0.000 | 0.000 |
| Kcnv2 | 0.000 | 0.001 | 0.004 |
| D930032P07F | 0.000 | 0.000 | 0.000 |
| Slc1a1 | 0.000 | 0.001 | 0.001 |
| Rln1 | 0.000 | 0.001 | 0.001 |
| Mbl2 | 0.000 | 0.000 | 0.001 |
| A1cf | 0.000 | 0.000 | 0.000 |
| Lipo4 | 0.000 | 0.000 | 0.000 |
| Lipo2 | 0.000 | 0.001 | 0.000 |
| Ifit1bl2 | 0.000 | 0.000 | 0.001 |
| Slc16a12 | 0.000 | 0.000 | 0.001 |
| Gm5248 | 0.000 | 0.000 | 0.000 |
| A830019P07F | 0.000 | 0.001 | 0.000 |
| Htr7 | 0.000 | 0.000 | 0.000 |
| Gm47735 | 0.000 | 0.000 | 0.000 |
| A330032B11F | 0.000 | 0.000 | 0.000 |
| Ffar4 | 0.000 | 0.000 | 0.000 |
| Pde6c | 0.000 | 0.000 | 0.000 |
| Gm42723 | 0.000 | 0.000 | 0.000 |
| Gm28991 | 0.000 | 0.000 | 0.000 |
| Cyp2c55 | 0.000 | 0.000 | 0.000 |
| Cyp2c65 | 0.000 | 0.000 | 0.000 |
| Cyp2c70 | 0.000 | 0.000 | 0.000 |
| Cc2d2b | 0.000 | 0.001 | 0.000 |
| Blnk | 0.000 | 0.001 | 0.001 |
| Dntt | 0.000 | 0.000 | 0.000 |
| Tll2 | 0.000 | 0.000 | 0.000 |
| Pik3ap1 | 0.000 | 0.000 | 0.000 |
| Slit1 | 0.000 | 0.000 | 0.000 |
| 4933411K16F | 0.000 | 0.000 | 0.000 |
| Sfrp5 | 0.000 | 0.002 | 0.002 |
| Golga7b | 0.000 | 0.000 | 0.000 |
| Gm16541 | 0.000 | 0.000 | 0.000 |
| Hpse2 | 0.000 | 0.000 | 0.000 |
| Gm47936 | 0.000 | 0.000 | 0.000 |
| Gm47938 | 0.000 | 0.000 | 0.000 |
| Cyp2c23 | 0.000 | 0.000 | 0.002 |
| Scd3 | 0.000 | 0.000 | 0.000 |
| Wnt8b | 0.000 | 0.000 | 0.001 |
| Gm20538 | 0.000 | 0.000 | 0.000 |
| Gm26644 | 0.000 | 0.000 | 0.000 |
| Pax2 | 0.000 | 0.001 | 0.000 |
| Gm20395 | 0.000 | 0.000 | 0.000 |
| Gm29595 | 0.000 | 0.000 | 0.000 |

| | | | |
|---|---|---|---|
| Gm28578 | 0.000 | 0.000 | 0.000 |
| Fgf8 | 0.000 | 0.001 | 0.000 |
| 4930505N22I | 0.000 | 0.000 | 0.001 |
| Pitx3 | 0.000 | 0.000 | 0.000 |
| Cfap58 | 0.000 | 0.000 | 0.000 |
| Gm45352 | 0.000 | 0.000 | 0.000 |
| Gm26629 | 0.000 | 0.000 | 0.000 |
| 1700054A03I | 0.000 | 0.000 | 0.000 |
| Gm10197 | 0.000 | 0.000 | 0.000 |
| 4833407H14I | 0.000 | 0.000 | 0.000 |
| Tectb | 0.000 | 0.000 | 0.000 |
| Gucy2g | 0.000 | 0.002 | 0.000 |
| 4930552P12I | 0.000 | 0.002 | 0.001 |
| Habp2 | 0.000 | 0.000 | 0.001 |
| Plekhs1 | 0.000 | 0.001 | 0.000 |
| Vwa2 | 0.000 | 0.000 | 0.000 |
| Pnlip | 0.000 | 0.000 | 0.001 |
| Pnliprp2 | 0.000 | 0.000 | 0.000 |
| Vax1 | 0.000 | 0.000 | 0.000 |
| Emx2os | 0.000 | 0.000 | 0.000 |
| Emx2 | 0.000 | 0.000 | 0.000 |
| Gm7102 | 0.000 | 0.000 | 0.000 |
| AC133103.1 | 0.000 | 0.000 | 0.000 |
| Ccl21c | 0.000 | 0.000 | 0.000 |
| CR974586.3 | 0.000 | 0.000 | 0.000 |
| CR974586.5 | 0.000 | 0.007 | 0.012 |
| AC132444.1 | 0.000 | 0.000 | 0.000 |
| Csprs | 0.000 | 0.000 | 0.000 |
| AC132444.6 | 0.000 | 0.001 | 0.000 |
| Gm10931 | 0.000 | 0.001 | 0.002 |
| CT868723.1 | 0.000 | 0.000 | 0.000 |
| AC125149.3 | 0.000 | 0.000 | 0.000 |
| AC125149.2 | 0.000 | 0.000 | 0.000 |
| AC125149.4 | 0.000 | 0.000 | 0.000 |
| AC168977.2 | 0.000 | 0.000 | 0.000 |
